# *In Vivo* Large Scale Mapping Of Protein Turnover In The Human Cerebrospinal Fluid

**DOI:** 10.1101/710418

**Authors:** Sylvain Lehmann, Christophe Hirtz, Jérôme Vialaret, Maxence Ory, Guillaume Gras Combes, Marine Le Corre, Stéphanie Badiou, Jean-Paul Cristol, Olivier Hanon, Emmanuel Cornillot, Luc Bauchet, Audrey Gabelle, Jacques Colinge

**Affiliations:** CHU de Montpellier, Montpellier, France; IRMB, INSERM U1183, Laboratoire de Biochimie Protéomique Clinique, Montpellier, France; Université de Montpellier, Montpellier, France; Institut de Recherche en Cancérologie de Montpellier, INSERM U1194, Montpellier, France; CHU de Montpellier, Hôpital Gui de Chauliac, Service de Neurochirurgie, Montpellier, France; INSERM U1051, Montpellier, France; Département de Biochimie et Hormonologie, CHU de Montpellier, Montpellier, France; PhyMedExp, Université de Montpellier, INSERM, CNRS; AP-HP, Hôpital Broca, Service de Gériatrie, Paris, France; Université Paris Descartes, Sorbonne Paris Cité, Paris, France; Centre Mémoire de Ressources et de Recherche Languedoc-Roussillon, Montpellier, France; CHU de Montpellier, Hôpital Gui de Chauliac, Montpellier, France; Institut régional du Cancer de Montpellier, Montpellier, France

**Keywords:** clinical proteomics, dynamic proteome, cerebrospinal fluid, bioinformatics

## Abstract

The extraction of accurate physiological parameters from clinical samples provides a unique perspective to understand disease etiology and evolution, including under therapy. We introduce a new proteomics framework to map patient proteome dynamics *in vivo*, either proteome wide or in large targeted panels. We applied it to ventricular cerebrospinal fluid (CSF) and could determine the turnover parameters of almost 200 proteins, whereas a handful were known previously. We covered a large number of neuron biology- and immune system-related proteins including many biomarkers and drug targets. This first large data set unraveled a significant relationship between turnover and protein origin that relates to our ability to investigate the central nervous system physiology precisely in future studies. Our data constitute a reference in CSF biology as well as a repertoire of peptides for the community to design new proteome dynamics analyses. The disclosed methods apply to other fluids or tissues provided sequential sample collection can be performed.

## INTRODUCTION

Clinical proteomics mostly relies on the absolute quantification of targeted proteins or on global proteome quantification(Doherty and Whitfield, 2011; Meyer and Schilling, 2017). Although highly successful, this type of analysis does not reveal the synthesis and clearance rates behind the observed abundance. Detailed and tissue-specific knowledge of individual protein turnover constitutes a complementary perspective that provides a unique insight in protein regulation. For instance, protein turnover abnormalities were related to disease pathophysiology in certain cases: amyloid-β (Aβ) and Tau in Alzheimer disease(Mawuenyega et al., 2010; Sato et al., 2018) (AD), the superoxide dismutase [Cu-Zn] (SOD1) in amyotrophic lateral sclerosis(Crisp et al., 2015), or the retinol-binding protein 4 (RBP4) in type 1 diabetes(Jourdan et al., 2009). Other authors employed large-scale turnover mapping to identify the regulated processes involved in zebrafish heart morphogenesis(Konzer et al., 2013) or tissue remodeling during early-stage human heart failure(Lam et al., 2014).

Turnover data are commonly obtained by mass spectrometry (MS) and hence isotopic tracers are employed to label the newly synthesized proteins. The ratio of labeled *versus* unlabeled protein peptide abundances (**Fig. 1a**) is called the relative isotope abundance (RIA). Usually, a time course is realized to acquire RIA curves from which synthesis and/or degradation rates can be learnt through mathematical modeling. The tracer is typically introduced in animal or patient diet, or in cell culture media. Commonly used tracers(Claydon and Beynon, 2012) are ^13^C_6_-labeled amino acids, e.g. leucine (Leu) or phenylalanine, [^2^H_2_]O, or ^15^N. Different tracing protocols exist and a common choice consists in delivering the tracer continuously over a rather long period of time (weeks). Labeling thus reaches saturation in most proteins but the longest-lived ones(Hammond et al., 2016; Price et al., 2010). Other protocols, which could be regarded as pulse-chase experiments, provide the tracer over a limited period and keep collecting samples afterwards(Bateman et al., 2006). In every case, performing *in vivo* experiments in tissues causes the measured turnover to be the net result of multiple phenomena and not just a physical property of the proteins. The observed turnover results from local synthesis and degradation (molecular biology) as well as passive and active transport across tissues (physiology). Moreover, for a given protein, its turnover may vary in different tissues. Depending on the tracing protocol, different abilities to separate local *versus* remote contributions might be achieved.

**Figure 1.**
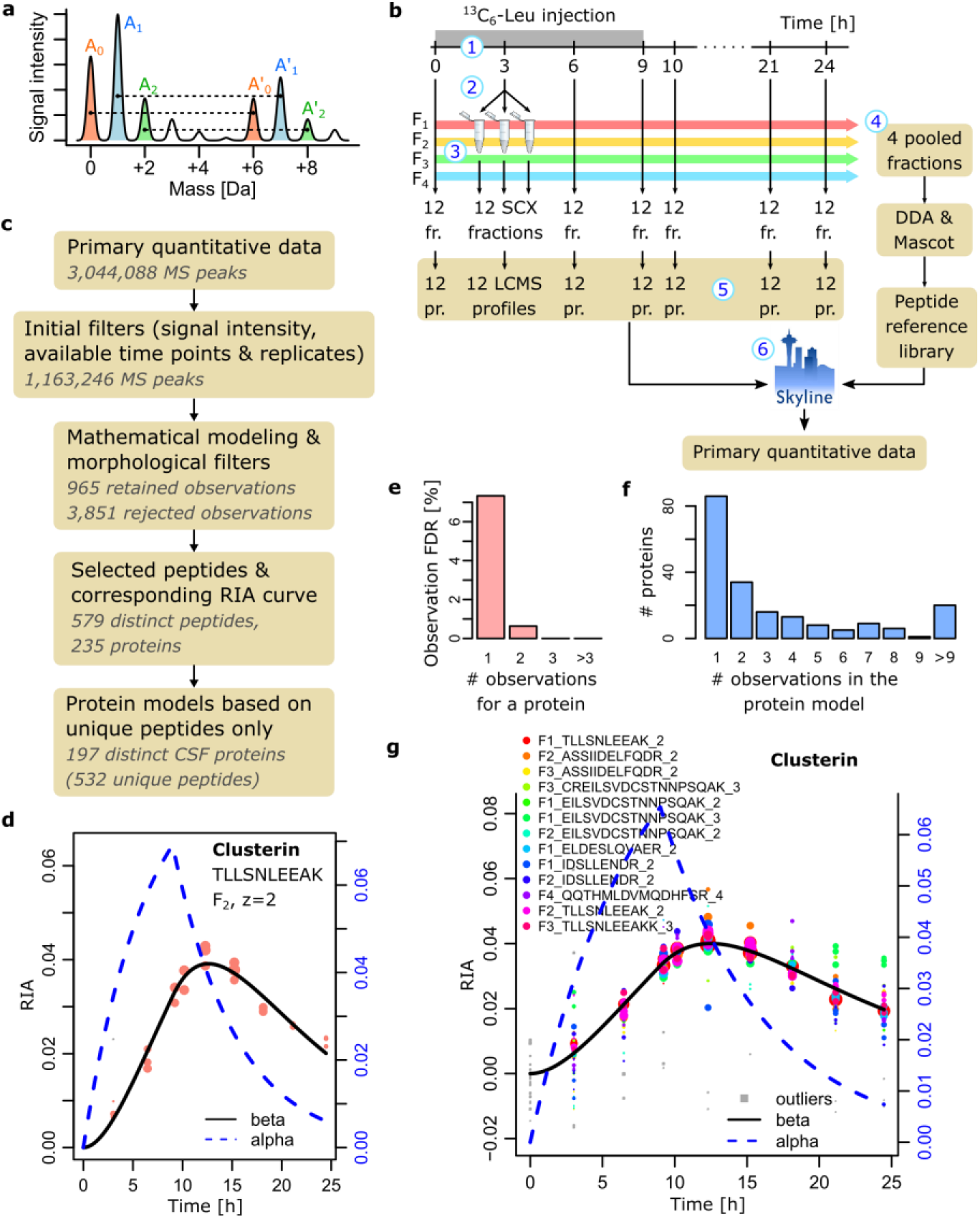
SILK principle. (**a**) Principle of RIA calculation. (**b**) Proteomics workflow: 1, tracer injection for 9 hours; 2, split of each sample in 3 aliquot to increase MS coverage; 3, peptide SCX separation; 5, LC-MS profiles of each fraction of each aliquot at every time point; 6, extraction of MS peaks after the peptide library. The (**c**) Bioinformatics workflow. (**d**) Typical example of a RIA curve featuring gradual incorporation before clearance for a clusterin peptide. The size of the dots represents the labelled peptide MS signal intensity (arbitrary scale). (**e**) False discovery rate estimation of the selected observations (one peptide in a specific fraction and at a specific charge state). (**f**) Number of proteins whose mathematical model was built on 1, 2, etc. observations. (**g**) Example of a protein mathematical model for clustering (observations are denoted fraction_peptide_charge).

Limited *in vivo* data are available in human. Initial efforts aimed at characterizing the total protein dynamics upon uptake of isotopically labeled aminoacids(Waterlow, 1995). More recent work taking advantage of progresses in proteomics determined protein turnover at the single protein level(Bateman et al., 2006; Crisp et al., 2015; Jourdan et al., 2009; Wildsmith et al., 2012), in small-to medium-size panels(Jaleel et al., 2006; Price et al., 2012), or proteome wide(Lam et al., 2014) with a strong bias towards plasma. The protocol called stable isotope labeling kinetics(Bateman et al., 2006) (SILK) was introduced to follow Aβ in the cerebrospinal fluid (CSF). A ^13^C_6_-Leu tracer was intravenously injected during 9 hours while CSF was sequentially collected over 24 hours or more. It is a thus pulse-chase protocol. SILK allowed its authors to determine the turnover of AD molecular actors such as Aβ peptides(Bateman et al., 2006), tau proteins(Sato et al., 2018), and ApoE(Wildsmith et al., 2012). One major finding was AD association with a reduced CSF clearance of Aβ. This observation drew attention to the glymphatic system impacting CSF circulation(Boespflug and Iliff, 2018) and was essential to model the kinetics of amyloid accumulation over the evolution of the disease(Jack and Holtzman, 2013). SILK was also used to confirm the on-target effect of an anti-AD drug(Bateman et al., 2009).

In this work, we introduce *whole proteome* SILK (wpSILK), a novel proteomics framework to perform SILK-like experiments on a large scale. In a first instance, we eliminated any affinity purification step and integrated peptide liquid chromatography (LC) to deal with sample complexity and to perform analyses proteome wide by high-resolution MS (HRMS) profiling. In a second instance, we followed targeted peptides by multiple reaction monitoring(Meyer and Schilling, 2017) (MRM). A sophisticated bioinformatics pipeline including mathematical modeling was developed to process the complex wpSILK data. We demonstrated the potential of wpSILK on ventricular CSF, a fluid of paramount clinical relevance, obtaining the turnover parameters of ^~^200 proteins. Our data constitute the largest reference of this kind. Furthermore, they provide a vast repertoire of easily MS-detectable peptides amenable to turnover experiments that cover a broad range of central nervous system (CNS) functions, known biomarkers, and drug targets. Finally, we tried to dissect the relationship between observed turnover and local *versus* remote protein origin. Our results have the potential to speed up pharmacokinetic/pharmacodynamic (PK/PD) modelling of therapeutic targets related to neurological diseases.

## RESULTS

### Sample analysis and data generation

Ventricular CSF and blood were collected from three post subarachnoid hemorrhage patients over a 24-36 hours period with intravenous injection of ^13^C_6_-Leu according to SILK published procedure(Bateman et al., 2006). The experiment started 8 to 19 days after initial, medical ventricular drainage and normalization of CSF clinical chemistry analysis (normal CSF protein content lies in the 0.2-0.4 g/L range(Roche et al., 2008)). Patient data are reported in **Table S1** and patient 1, who was chosen for HRMS profiling, had the largest post drainage delay. The other two patient samples were used for the MRM variant of wpSILK to confirm 26 selected proteins. CSF and plasma free ^13^C_6_-Leu concentrations were estimated by UPLC-MS/MS at different time points (**Table S2**) and matched previously reported values(Bateman et al., 2006).

Patient 1 CSF samples were submitted to wpSILK HRMS workflow (**Fig. 1b**). Briefly, each sample was split in 3 aliquots and digested (LysC and trypsin). Strong cation exchange (SCX) LC was used to obtain 6 fractions for each. Fractions 5 and 6 were regrouped and eventually discarded due to very limited peptide content (**Fig. S1**). The other 4 fractions were submitted to LC-MS profiling by first pooling the 3 identical SCX fractions of each aliquot and injecting this pool 3 times for better proteome coverage. A few injections were repeated due to poor LC (**Fig. S2**). In parallel, we generated a peptide reference library by pooling identical SCX fractions over all the time points (**Fig 1b**), which were submitted 6 times each to LC-MS-MS/MS. Mascot peptide identifications (<1% FDR) were imported in Skyline software(Pino et al., 2017). In total, we found 4,558 peptides present in 1,001 proteins and 3,196 peptides from 860 proteins were Leu-containing. Skyline was applied extract primary quantitative data from LC-MS, i.e. peptide identity, SCX fraction number, retention time, charge state, nominal and ^13^C_6_-Leu-shifted masses, and peak areas resulting in a large tabular export of 3,044,088 rows (**Table S3**). Time dependent incorporation of ^13^C_6_-Leu in a peptide could be followed by computing RIAs at each time point.

### Peptide selection and protein model construction

The limited duration and amounts of ^13^C_6_-Leu injection due to obvious patient safety considerations precluded full labeling of proteins. Consequently, heavy (^13^C_6_-Leu labeled) peptide signals were 10 to 100 times weaker than their light (unlabeled) counterparts. Moreover, untargeted HRMS profiling and subsequent peptide identification against a reference peptide library may yield wrong assignments. Data processing hence had to ensure that true RIA curves were extracted from Skyline export. Our algorithm follows the general line of bottom-up quantitative proteomics with peptide-level filters and protein-level modeling based on unique – non-shared – peptides only (**Fig. 1c**). Considering occasional low heavy peptide signals and Skyline peak misassignments, we found that RIA were more accurately computed using the most intense isotopic peak only, e.g. 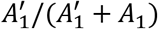 referring to **Fig. 1a**. Initial filters were applied to eliminate incomplete or too weak signals. RIA curves from the same peptide in distinct fractions or at different charge states were treated separately, as *independent observations*. While some peptides gave rise to convincing RIA curves, e.g. **Fig. 1d**, many adopted dubious shapes (**Fig. S3**). We thus reasoned that a mathematical model of ^13^C_6_-Leu incorporation might help eliminating ill-shaped curves and noisy data. Moreover, such a model is necessary to extract turnover parameters.

The so-called 2-compartment model, where tracers from a fee pool integrate a protein-bound pool, has been shown to fit turnover dynamics data accurately(Guan et al., 2012; Rahman et al., 2016). Considering the specifics of SILK, i.e. tracer injection over a limited period of time and free tracer concentration in the range 10-15% as opposed to saturation, we generalized this model. We also wrote the model in a novel fashion, better separating the role of the parameters. Mathematical details, differences with respect to previous literature, and the overall data processing algorithm are provided in Supplemental Information (SI). We only introduce the general principle here. A function *f*(*t*) representing ^13^C_6_-Leu injection is introduced, its value is 0 (no injection) or 1 (injection), i.e. *f*(*t*) = 0 if *t* ≤ 0 or t > 9, and *f*(*t*) = 1 otherwise. Denoting *A* = *A_L_* + *A_H_* the sum of light (unlabeled) and heavy (labeled) free Leu abundance, we can define the free ratio 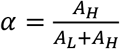. Similarly, the protein-bound total Leu abundance is *P* = *P_L_* + *P_H_*, with bound ratio 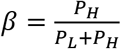. Assuming a rate *λ* of free ^13^C_6_-Leu availability and protein appearance/clearance rate *k_c_*, we obtain the dynamical system:

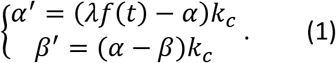

The absence of ^13^C_6_-Leu at *t* = 0 imposes *α*(0) = 0 = *β*(0). The first equation features entrance of new tracers in the free pool and exit into the bound pool, while the second one features corresponding entrance in the bound pool and protein degradation. Note that according to the introduction, we talk about appearance and clearance rates instead of the sole synthesis and degradation rates. Also, the model is written assuming a steady-state, which imposes that appearance and clearance occur at the same rate *k_c_*. Model parameters were fit by minimizing squared-differences between experimental RIA values and *β*(*t*), an example is featured in **Fig. 1d**.

The application of the model to Skyline output allowed us to impose obvious morphological filters, e.g. *β*(*t*) and *α*(*t*) both increasing initially, maximum of *β*(*t*) not reached before 9h, and sufficient correlation between *β*(*t*) and experimental RIAs. To estimate the FDR associated with peptide observations we processed peptides devoid of Leu identically. We estimated a global FDR of 7.3%, which dropped to 0.6%, respectively 0%, when 2, respectively 3 or more, observations were available for a single protein (**Fig. 1e**). At this FDR we selected 965 observations, covering 579 distinct peptides and 235 proteins. Out of these 579 peptides, 532 were unique, i.e. not shared by multiple proteins, and served as the basis of protein turnover calculations for 196 distinct proteins. Peptides from precursors of complement C4 isotypes A and B were found with a large number of selected observations (23), none of which were unique and neither C4A nor C4B were counted. We added C4A to our list as a representative of both isotypes, which raised the total of distinct proteins to 197. Among these 197 proteins, 86 were detected with a single observation at 7.3% FDR, an acceptable risk given the unique nature of the data. The remaining 111 proteins were associated with high-confidence turnover information (**Fig. 1f**).

When more than one observation were available, we integrated them. Although isoforms or chains might display different turnovers(Doherty et al., 2012; Wildsmith et al., 2012), most of the observed variability was expected to be experimental. We implemented a robust algorithm, where all the observations of unique peptides were first combined and fit with our model (Eq. (1)). Then, outlier observations were eventually discarded and the integrated model recomputed. A typical example is featured in **Fig. 1g**. Finally, to obtain estimates of model parameter standard deviations, including 1-observation proteins, we employed a bootstrap procedure(Davison and Hinkley, 1997). Details of the overall procedure, parameter tables as well as plots for all the selected proteins are provided in SI.

The found protein CSF dynamics were very diverse (**Fig. 2**). The distribution of half-lives consisted of a bulk of rather short-lived proteins followed by other proteins harboring slower clearance rates (**Fig. 2a**). Many proteins displayed an increase of their labeled proportion over the whole experiment duration, e.g. KLK6 or ALB (**Fig. 2e-f**) despite a tracer injection stop at 9h.

**Figure 2.**
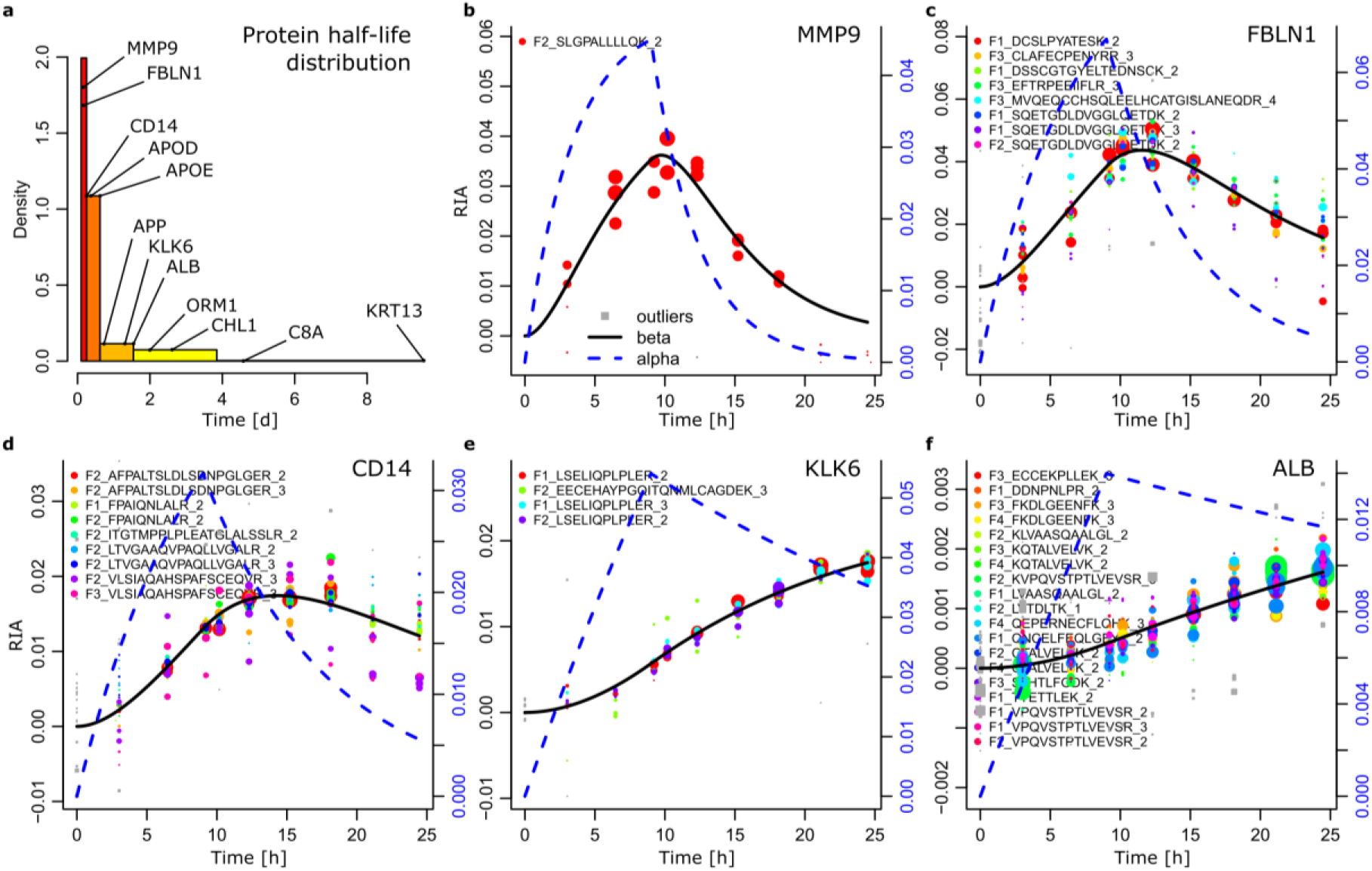
Overview of the observed dynamics. (**a**) Histogram of observed half-lives with selected examples indicated. (**b-f**) Representative examples of distinct dynamics: Matrix metalloproteinase-9 (MMP9); Fibulin-1 (FBLN1), Monocyte differentiation antigen CD14; Kallikrein-6 (KLK6); and Serum albumin (ALB).

### Validation of selected proteins in additional patients

To confirm wpSILK HRMS results we exploited the MRM variant of our framework on 26 chosen proteins. MRM was executed on the CSF samples directly, without prior SCX peptide separation (SI). The mathematical model was applied successfully to MRM data (**Fig. 3a-b**) and the various peptide selection filters applied identically. Starting with patient 1, we found MRM RIA curves highly similar to HRMS data (**Fig. 3c-e**) for all the proteins but two, C9 and GC (**Fig. 3f-g**). The amplitude of the curves were not preserved, reflecting differences in the MS technology, but the shape of the curve governed by the clearance rate *k_c_* was well-preserved. The new parametrization of the 2-compartment model we introduced in Eq. (1) facilitated the estimation of close *k_c_* from similar curves but different amplitudes, yielding reproducible estimations of the turnover parameter *k_c_* (**Fig. 3h**). The parameter *λ*, more related to the amplitude was nonetheless correlated (**Fig. 3i**). Protein dynamics from patients 2 and 4 (**Fig. 3c-g**), were also close and we could estimate inter-individual turnover parameter *k_c_* variability in the range 10-20% for most proteins (**Fig. 3j**), *λ* being more variable (**Fig. 3k**) and reflecting potential variation in ^13^C_6_-Leu patient labelling efficiencies. The 26 protein model parameters are reported in **Table S4**.

**Figure 3.**
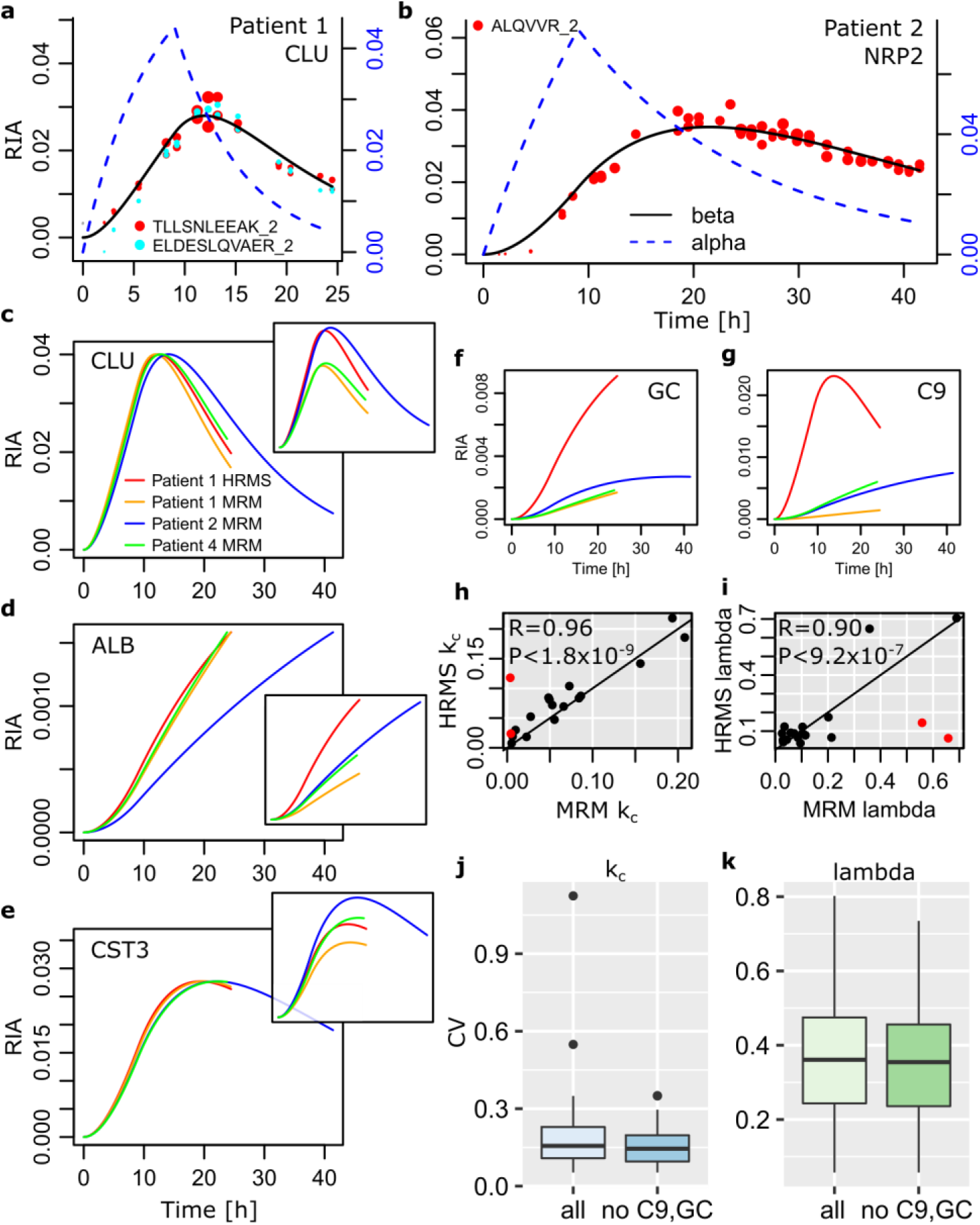
Validation of selected proteins by triple-quad MRM. (**a**) Two clusterin peptides targeted in MRM. (**b**) Neuropilin 2 MRM data in patient 2 who was followed over 41.5 hours. (**c**) Clusterin dynamics in 3 patients, including HRMS and MRM data in patient 1. Original, non-aligned RIA reported in the insert. (d) Serum albumin. (e) Cystatin-C. (f,g) Only 2 cases of strong disagreement between HRMS and TQ data (vitamin D-binding protein, GC, and complement component C9). (h,i) Reproducibility of patient 1 turnover parameters between HRMS and MRM data. Pearson correlation, GC and C9 data excluded (red dots). Pearson’s correlation coefficient and P-value by transforming to a Student’s *t* distribution. (j,k) Relative variability of turnover parameters between the 3 patient MRM data (left, all proteins), and omitting GC and C9 (right).

### Dynamic proteome composition

Among the 197 proteins for which we obtained turnover parameters, 185 were known CSF proteins and 190 known plasma protein. We used as reference plasma proteome the union of the plasma proteome database(Nanjappa et al., 2014) (PPD) and proteins annotated as plasma in UniProtKB/Swissprot. Similarly, we defined a reference CSF proteome by taking the union of CSF proteins reported in two comprehensive lists(Barkovits et al., 2018; Fernandez-Irigoyen et al., 2015). Potentially novel CSF proteins were (HUGO symbols for short) CEP290, FBXW10, KSR2, LOX, SH2D3A, SIK2, SPEN, TMEM212 and immunoglobulins (IGHV3-43, IGHV3-74, IGKV3D-15, IGKV4-1). FBXW10, SH2D3A, and TMEM212 were also absent from our reference plasma proteome. By compiling annotations from Reactome, KEGG, and GOBP, we classified our 197 proteins in general categories and more detailed functions (**Fig. 4a-b**). We covered mainly neuron biology-associated proteins (56), the immune system (44), the extracellular matrix (ECM) (37), intracellular proteins (34) likely reflecting cell leakage, and hemostasis (26).

**Figure 4.**
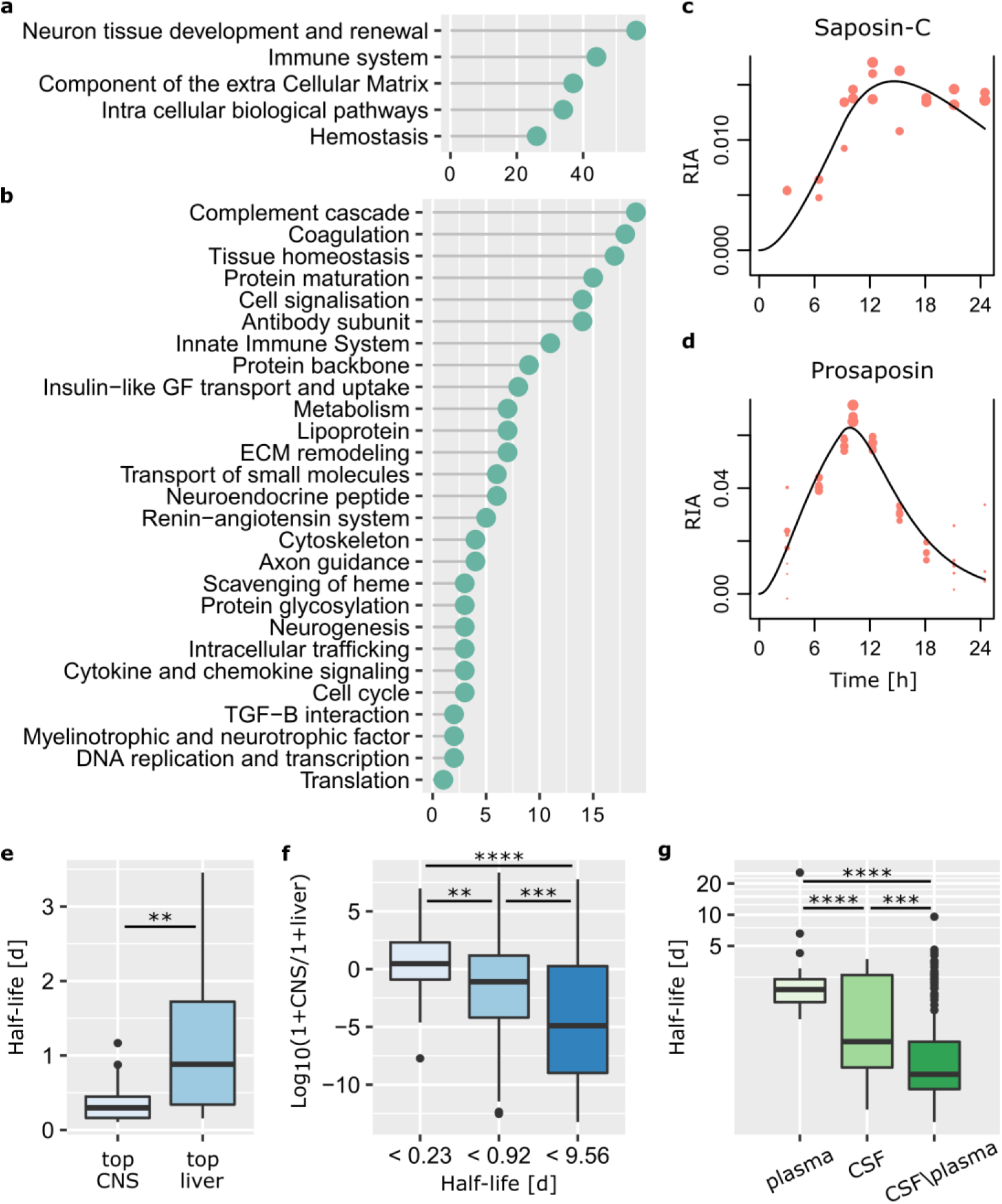
Dynamic proteome composition and relationship with synthesis location. (**a,b**) General and more detailed categories of proteins for which turnover was determined. (**c,d**) Mature chain Saposin-C versus propeptide Prosaposin dynamics. (**e**) Top 22 proteins expressed in the CNS and the liver (GTEx database(GTEx Consortium, 2013)) display very different stability (**P<0.01, Kolmogorov-Smirnov (KS) test). (**f**) Significant trend over the entire data set relating the CNS/liver TPM ratios(GTEx Consortium, 2013) to the first (<0.23), second and third (<0.92), and fourth (<9.56) quartiles of half-lives. (***P<0.005, ****P<10^−5^, KS test). (**g**) Price *et al*., human plasma(Price et al., 2012) halflives (left) *versus* the same protein in our CSF data (middle) and our CSF protein not present in the plasma data (right, KS test).

Neural tissue development and renewal proteins were comprised of different subcategories, e.g. apolipoproteins involved in lipid delivery, growth factors such as IGF-2 and proteins related to IGF transport and uptake, secretogranins, or proteins responsible for small molecule transport such as albumin, transferrin, transthyretin, vitamin D binding protein, and alpha-1-acid glycoprotein 1 involved in drug uptake for instance. Multiple neuropeptides were also profiled, among which the proprotein convertase subtilisin/kexin type 1 inhibitor (PCSK1N), proenkephalin, prosaposin (PSAP), out at first homolog, neural EGFL like 2, chromogranins A and B, etc. Neuronal tissue homeostasis was represented by the amyloid beta precursor protein (APP), cystatin C, follistatin like 1, and calsyntenin 1 among others. Intracellular and extracellular matrix proteins (ECMs, fibulins, VTN) were likely waste or debris of dead or damaged cells in majority. The dynamics of their clearance informs us on the ability of the CSF - and the CNS - to normally eliminate them, deviation from these rates might reflect pathological states.

Interestingly, we could identify distinct dynamics of the precursor and processed peptides of the myelinotrophic and neurotrophic factor PSAP involved in lysosomal degradation of sphingolipids (**Fig. 4cd**) and the prohormone convertase 1 inhibitor PCSK1N (**Fig. S4**). In some cases, peptides from the same protein display very different dynamics as exemplified by SPP1 (**SupFig. 5**). Its peptides segregate according to primary sequence positions coherently, which might suggest the existence of undescribed post-translational processing of this proteins that plays a major role in tumor progression and inflammation(Castello et al., 2012). We also obtained different turnover parameters for the components of the complement system, which might reflect overall control through the shortest-lived components (C1QB, C1QC, C1R, C1S, and C9). Similarly, at the level of a protein complex, FGA has a much higher clearance rate than FGG and might control the fibrinogen assembly in a rate-limiting fashion (SI).

### A repertoire of CSF proteotypic turnover biomarkers

Our data provide a large repertoire of CSF peptides that yielded high-quality MS signals compatible with dynamics profiling. More than 100 of the corresponding proteins harbor mutations or display altered expression in one or several diseases (**Table S5**). Among them, neurological diseases including Alzheimer (APOE, APP), spinocerebellar ataxia (FAT2), leukoencephalopathy (CSF1R, HTRA1, RNASET2, PSAP) and various disorders (AHSG, ATP6AP2, CEP152, CHI3L1, CTSD, DST, RELN, RPS23) are well represented. We also found proteins responsible for amyloidosis (FGA, GSN, LYZ, APOA2, APP, B2M, CST3), a situation where the investigation of the protein fate, i.e. the balance between production, aggregation, and clearance, is essential to understand the underneath pathophysiology(Bateman et al., 2006). We noticed 16 proteins (ADAM9, CEP290, C3, C4a, C9, CFB, CFH, CFI, CST3, DCN, EFEMP1, HTRA1, IGFBP7, LTBP2, RBP4, TGFBI), which are linked to eye diseases, and in particular macular degeneration. We also surveyed the clinical trial database (clinicaltrials.gov) and found 23 proteins that are the direct targets of tested therapies (**Table S6**), 4 of them for diseases affecting the CNS and targets for future AD therapy (APP, CD14, MMP9, SERPING1).

Thanks to omics approaches, networks of proteins considered as potential biomarkers for specific pathologies were assembled. We mapped our protein list using the “Diseases and Functions Annotation” Ingenuity^®^ tool and found a large number of additional Alzheimer disease proteins: A2M, AGT, ALB, APOA1, APOA2, APOA4, APOD, C1R, C4A/C4B, C7, C9, CD14, CFB, CFD, CHI3L1, CHL1, CLEC3B, CLU, CST3, CTSD, CXCL12, GC, GM2A, HP, HPX, HRG, HTRA1, IGF2, IGFBP2, IGFBP6, IGHA2, IGHG1, IGHM, IGKC, KNG1, MMP9, PENK, PLG, PLTP, PROS1, PSAP, PTGDS, RELN, RNASET2, SERPINA1, SERPINA3, SERPINC1, SPARC, TIMP1, TTR. Many of them are in fact believed to play a role in the production and regulation of Aβ peptides. For example, MMP-9 (**Fig. 2b**), involved in neuronal plasticity, acts as α-secretase therefore reducing Aβ production(Talamagas et al., 2007). ApoE, ALB (**Fig. 2f**), A2M are described as Aβ carrier. CD14 protein (**Fig. 2d**), which is a receptor for TREM2, a recently identified genetic risk factor of AD is a marker of microglial cells that mediate the phagocytosis of the amyloid component(Bate et al., 2004) and contribute to neuroinflammation.

### A general relationship between turnover and protein origin

The CSF is relatively poor in cells and most proteins are imported from the blood or the CNS with a double origin for many of them. We continued our analysis by investigating the relationship between the tissue of origin of a protein and its turnover. Due to protein transport in the body, we reasoned that mRNA expression was a better locator of synthesis compared to tissue protein abundance. The Genotype-Tissue Expression (GTEx) project(GTEx Consortium, 2013) served as reference. A majority of our 197 proteins were of CNS or liver origin. We grouped GTEx tissues to obtain CNS, liver, and other tissues median transcripts per million (TPM) for each gene (SI). Selecting the top 25 CNS and liver transcripts, removing common selections, left us with 22 CNS and 22 liver genes. The corresponding CNS proteins displayed significantly shorter half-lives (**Fig. 4e**) suggesting a potential general trend. We hence grouped the whole set of 197 proteins after their half-life in 3 bins: quartiles 1, 4, and 2&3. Comparing the ratio of CNS over liver transcript abundance in these 3 groups revealed significant association with half-life (**Fig. 4f**). To complete this analysis we reprocessed published plasma data(Price et al., 2012) with our mathematical model and algorithm (SI). For the 41 proteins that were shared by our CSF data and at least 2 out of 3 patients of the plasma study, we observed shorter half-life in the CSF (**Fig. 4g**). This significant trend was even stronger for CSF proteins absent from the plasma study, in agreement with **Fig. 4f**.

We believe that the relationship between half-life and protein origin results from the combined effect of CSF renewal on the one hand, and plasma and other tissues acting as a reservoir on the other hand. The CSF is mostly produced by the choroid plexus and reabsorbed by arachnoid granulations, its renewal occurs 3-4 times/day. Proteins entering the CSF, from the CNS or the plasma, experience this flow which adds to protein degradation. Remarkably, median half-life of our 197 proteins is 7.4 hours, a value compatible with CSF renewal rate. Nonetheless, a dissymmetry is created with proteins not produced in the CNS due to the aforementioned reservoir effect, which might even be amplified by frequent higher protein plasma concentrations compared to CSF values. The reservoir effect hides or changes CNS clearance rates of proteins that are not present in the CNS predominantly or not blocked at the CSF barrier resulting in apparent continuous supply of labelled proteins. One could hypothesize that stability could be different for proteins produced in the liver due to the necessity to travel a long way. This nonetheless contradicts murine data, where brain proteins were found more stable than liver proteins(Price et al., 2010), an observation we could confirm reanalyzing these data and considering proteins detected in brain, blood and liver to limit the contribution of intracellular proteins (**Fig. S6**).

## DISCUSSION

To be able to link molecular biology and physiology in a global picture is a very desirable goal. This is the foundation of many past and current efforts and tremendous progresses have been achieved. Nevertheless, to fully embrace the complexity of the relationship between locally occurring molecular interactions and the interplay of several tissues remains an immense challenge. In this report we contributed what we believe could be an important tool to address that task. We propose a methodological framework (wpSILK) to *in vivo* interrogate clinical samples for protein turnover in a highly multiplexed, targeted fashion employing MRM or proteome wide through HRMS. It required the development of innovative bioinformatics and mathematical modeling, which were shown to be applicable to already published data sets in addition to ours. Recent MS instrument developments such as applications ion mobility or further improvements might ultimately alleviate the need for chromatography or allow to analyze all the time points at once exploiting isobaric tagging. In such a case, wpSILK implementation would be similar to the MRM variant we presented and mathematical modeling would remain identical.

We aimed at characterizing human ventricular CSF *in vivo* with wpSILK and we could obtain CSF turnover parameters for 197 proteins, which is a significant increase over previously existing data covering a handful of proteins(Bateman et al., 2006; Crisp et al., 2015; Sato et al., 2018; Wildsmith et al., 2012) (Aβ, APOE, SOD1, tau) to the best of our knowledge. We further showed result reproducibility across a panel of 26 proteins in the same patient and two additional individuals with an orthogonal MS technology (MRM). This panel provided a first estimate of inter-individual turnover variability (10-20%) for late-sixty females. Among previous data, only APOE was also present in our study, we found a turnover parameter *k_c_* in the range 3-5.5%/day for APOE, whereas previous values in younger (22-49 years old) males were in the 1.5-2%/day range. Overall, the inter-individual variability we observed in turnover is compatible with what was reported for APOE, SOD1, and tau (17-40%).

The opportunity to analyze a first large CSF turnover data set unraveled significant correlation between CSF clearance rates and the origin of proteins present in this fluid, i.e. CNS *versus* peripheral. We could support this discovery integrating published human(Guan et al., 2012; Wang et al., 2014) and murine(Price et al., 2010) data. CNS synthesized proteins tended to be cleared faster, which we believe is related to CSF renewal and peripheral tissues (blood, liver, etc.) acting as a reservoir. CSF renewal impacts the clearance of every protein present in the CSF, whereas protein entering the CSF through its barrier with blood experience a concomitant *reservoir effect*. The latter mechanism provides labeled proteins to CSF for a longer time due to peripheral tissues larger volumes, frequent higher concentrations, and delays metabolizing free ^13^C_6_-Leu and exporting the labeled proteins towards the CSF. That is, the obtained turnover data accurately reflect CSF physiology but access to CNS turnover parameters might be limited for proteins that are not CNS-specific, especially if they have a longer halflife. This observation has consequences on the choice of the tracing protocol suggesting that continuous tracing protocols might tend to bias *in vivo* CSF-inferred CNS dynamics with average body dynamics. This is especially true for biomarkers with multiple indications in the CNS and at other organs, e.g. transthyretin (TTR) for AD(Velayudhan et al., 2012) and malnutrition or protein metabolic impairment(Ingenbleek and Bernstein, 2015). Pulse-chase tracing should be preferred to more accurately approach CNS physiology.

Our complete data set constitutes a broad and diverse repertoire of proteotypic peptides amenable to turnover analyses from which the community could compose panels of MRM peptides to conduct research on CNS pathologies or biology. In some cases (PSAP and PCSK1N), we were able to distinguish between propeptides and active chain dynamics to follow enzymatic processing and maturation. This knowledge represents a veritable asset for the community, notably when the metabolism of a specific protein will be at the center of the investigation. This is the case in therapeutic research to evaluate the on-target effect of a pharmaceutical agent as exemplified on a secretase inhibitor acting on Aβ production(Bateman et al., 2009).

## ONLINE METHODS

### Human samples

Samples were generated following the clinical protocol “In Vivo Alzheimer Proteomics (PROMARA)” (ClinicalTrials Identifier: NCT02263235), which was authorized by the French ethical committee CPP Sud-Méditerranée IV (#2011-003926-28) and by the ANSM agency (#121457A-11). Patients hospitalized in neurosurgery unit having in place a temporary ventricular derivation of the CSF were enrolled. The labelling protocol was taken from SILK original publication(Bateman et al., 2006). Briefly, ^13^C_6_-Leu prepared in accordance with the European Pharmacopeia [19] was intravenously administered. After a 10min initial bolus at 2 mg/kg, a 8h50 infusion at 2 mg kg/h was performed. Three to 4 mL of ventricular CSF samples were collected at times: 0, 3.02, 6.47, 9.22, 10.17, 12.3, 15.22, 18.13, 21.13, and 24.47 hours. At times 0 and 9, 3 mL of plasma EDTA were also collected. CSF and blood were aliquoted in polypropylene tubes of 1.5 mL and stored at −80°C until further analysis.

### Proteomics and bioinformatics

Proteomics main steps were briefly mentioned in the Results Section and depicted in **Fig. 1b**. The main steps of the bioinformatics pipeline as well as the mathematical modeling approach are also covered in the Results Section and in **Fig. 1**. Full details for proteomics and bioinformatics are provided as SI.

## Supporting information

MRM parameters

HRMS protein table

TQ protein table

GTEx expression

HRMS protein models

TQ protein models

original TQ protein models

aligned TQ protein models

## Acknowledgements

This work was supported by the National French Alzheimer effort (“Plan Alzheimer”, ProMara PHRC 2010). JC was supported by a Fondation ARC grant PJA 20141201975 for the computer infrastructure. We thank Randall J. Bateman (St Louis, MO, USA) for his help realizing this project.

