## supplemental information for "*In Vivo* Large Scale Mapping Of Protein Turnover In The Human Cerebrospinal Fluid"

Short title: protein turnover in the human cerebrospinal fluid

Sylvain Lehmann[1,2,*], Christophe Hirtz[1,2], Jérôme Vialaret[1], Maxence Ory[3], Guillaume Gras Combes[2,4], Marine Le Corre[2,4], Stéphanie Badiou[2,5], Jean-Paul Cristol[2,5], Olivier Hanon[6], Emmanuel Cornillot[2,3], Luc Bauchet[2,4], Audrey Gabelle[2,7], Jacques Colinge[2,3,8*]

(1) CHU de Montpellier, Montpellier, France ; IRMB, INSERM U1183, Laboratoire de Biochimie Protéomique Clinique, Montpellier, France.

(2) Université de Montpellier, Montpellier, France.

(3) Institut de Recherche en Cancérologie de Montpellier, INSERM U1194, Montpellier, France.

(4) CHU de Montpellier, Hôpital Gui de Chauliac, Service de Neurochirurgie, Montpellier, France ; INSERM U1051, Montpellier, France.

(5) Département de Biochimie et Hormonologie, CHU de Montpellier, Montpellier, France ; PhyMedExp, Université de Montpellier, INSERM, CNRS.

(6) AP-HP, Hôpital Broca, Service de Gériatrie, Paris, France ; Université Paris Descartes, Sorbonne Paris Cité, Paris, France.

(7) Centre Mémoire de Ressources et de Recherche Languedoc-Roussillon, Montpellier, France ; CHU de Montpellier, Hôpital Gui de Chauliac, Montpellier, France.

(8) Institut régional du Cancer de Montpellier, Montpellier, France.

**Contents:**

A. Supplementary Tables
B. Supplementary Figures
C. Supplementary Proteomics Materials and Methods
D. Supplementary Bioinformatics Materials and Methods
E. Supplementary Data (in separated PDF files):
    1. MRM parameters
    2. HRMS Protein table
    3. TQ Protein table
    4. HRMS Protein model plots
    5. TQ Protein model plots
    6. Aligned TQ protein models
    7. Original TQ protein models
    8. mRNA expression in GTEx

### A. Supplementary Tables

**Table S1.** Demographic information.

| Patient | Sex | Age | Diagnosis | Initial CSF protein concentration | Time after set up of ventricular drainage | CSF protein concentration |
|---|---|---|---|---|---|---|
| P1 | F | 67 | Subarachnoid hemorrhage, Fisher scale 4, left carotid aneurysm | 1.39 | 19 days | 0.34 g/L |
| P2 | F | 65 | Subarachnoid hemorrhage, Fisher scale 4, anterior communicating artery aneurysm | 1.97 | 15 days | 0.15 g/L |
| P4 | F | 66 | Subarachnoid hemorrhage, Fisher scale 3, cerebral anterior artery aneurysm | 2.55 | 8 days | 0.50 g/L |

**Table S2.** Free $^{13}C_6$-Leu concentration in ventricular CSF and plasma.

| Patient | Fluid | Time [h] | $^{13}C_6$-Leu concentration |
|---|---|---|---|
| P1 | CSF | 0.00 | 0.000 |
| | CSF | 3.02 | 0.072 |
| | CSF | 9.22 | 0.080 |
| | CSF | 12.30 | 0.035 |
| | CSF | 15.22 | 0.016 |
| | CSF | 18.13 | 0.012 |
| | CSF | 21.13 | 0.010 |
| | CSF | 24.47 | 0.008 |
| | Plasma | 9.22 | 0.156 |
| | Plasma | 24.47 | 0.006 |
| P2 | CSF | 8.50 | 0.091 |
| P4 | CSF | 8.67 | 0.099 |

**Table S3.** Skyline MS peak area export. Skyline export size, i.e. the number of peak areas, is reduced by a factor 2.6 after filtering but peptide and protein diversity is not affected. This means that redundant MS peaks – not the most intense isotopic peaks – were removed only.

| SCX fraction | Skyline export size | Proteins | Peptides | Skyline filtered export size | Proteins | Peptides |
|---|---|---|---|---|---|---|
| F1 | 1,071,020 | 787 | 1,471 | 404,208 | 787 | 1,471 |
| F2 | 1,101,124 | 646 | 1,451 | 426,704 | 646 | 1,451 |
| F3 | 522,632 | 373 | 792 | 208,018 | 373 | 792 |
| F4 | 349,312 | 195 | 409 | 124,316 | 195 | 409 |
| Global* | ∑=3,044,088 | \|U\|=1,032 | \|U\|=2,785 | ∑=1,163,246 | \|U\|=1,032 | \|U\|=2,785 |

*Global numbers can be a sum (denoted by ∑=) or the size of the union of all the proteins or peptides in all the fractions (denoted by |U|=, a protein can have peptides in multiple fractions and a peptide can be detected in several fractions).

**Table S4.** Turnover parameters for the 26 MRM targeted proteins. Empty cells reflect rejected observations that did not pass our quality filters (same criteria as for HRMS data).

| HUGO symbol | $k_c$ ± 2sd [1/h] | | | |
|---|---|---|---|---|
| | Patient 1 | | Patient 2 | Patient 4 |
| | HRMS | MRM | MRM | MRM |
| A1BG | 0.0077 ± 0.0011 | 0.0052 ± 0.0083 | | 0.0069 ± 0.0096 |
| A2M | 0.0835 ± 0.0141 | 0.0840 ± 0.0276 | 0.0529 ± 0.0126 | 0.0822 ± 0.0374 |
| ALB | 0.0192 ± 0.0028 | 0.0064 ± 0.0025 | 0.0077 ± 0.0015 | 0.0078 ± 0.0032 |
| AMBP | 0.0809 ± 0.0149 | 0.0495 ± 0.0340 | | |
| APOA1 | 0.0299 ± 0.0120 | 0.0098 ± 0.0039 | 0.0051 ± 0.0018 | 0.0107 ± 0.0044 |
| APOA2 | 0.0175 ± 0.0080 | | | 0.0600 ± 0.0724 |
| APOA4 | 0.0184 ± 0.0041 | 0.0226 ± 0.0554 | | |
| APOD | 0.0873 ± 0.0059 | 0.0862 ± 0.0233 | 0.0676 ± 0.0080 | 0.0893 ± 0.0311 |
| APOE | 0.0473 ± 0.0053 | 0.0554 ± 0.0208 | 0.0299 ± 0.0055 | 0.0440 ± 0.0264 |
| B2M | 0.0705 ± 0.0137 | | 0.0646 ± 0.0077 | 0.0700 ± 0.0611 |
| C3 | 0.0856 ± 0.0064 | | | 0.1140 ± 0.0420 |
| C4B | 0.0710 ± 0.0049 | 0.0622 ± 0.0227 | 0.0559 ± 0.0073 | 0.0502 ± 0.0290 |
| C9 | 0.1179 ± 0.0250 | 0.0036 ± 0.0016 | 0.0126 ± 0.0063 | 0.0085 ± 0.0036 |
| CFB | 0.0843 ± 0.0090 | 0.0484 ± 0.0413 | | 0.0803 ± 0.0392 |
| CLU | 0.1418 ± 0.0067 | 0.1559 ± 0.0353 | 0.1131 ± 0.0125 | 0.1342 ± 0.0341 |
| CNDP1 | 0.0131 ± 0.0020 | | 0.0237 ± 0.0100 | |
| CST3 | 0.0695 ± 0.0124 | 0.0661 ± 0.0232 | 0.0585 ± 0.0072 | 0.0574 ± 0.0276 |
| F12 | | 0.0076 ± 0.0030 | 0.0078 ± 0.0017 | 0.0071 ± 0.0048 |
| GC | 0.0238 ± 0.0128 | 0.0043 ± 0.0025 | 0.0294 ± 0.0149 | 0.0047 ± 0.0064 |
| GSN | 0.0719 ± 0.0093 | 0.0528 ± 0.0236 | 0.0548 ± 0.0091 | 0.0745 ± 0.0304 |
| NRP2 | | 0.0651 ± 0.0206 | 0.0604 ± 0.0064 | 0.0537 ± 0.0270 |
| RBP4 | 0.0524 ± 0.0193 | 0.0275 ± 0.0226 | 0.0397 ± 0.0091 | 0.0326 ± 0.0631 |
| SERPINA3 | 0.1038 ± 0.0060 | 0.0728 ± 0.0242 | 0.0560 ± 0.0078 | 0.0748 ± 0.0290 |
| SERPINC1 | 0.0078 ± 0.0015 | | 0.0923 ± 0.0190 | |
| SERPINF1 | 0.2182 ± 0.0089 | 0.1934 ± 0.0563 | 0.1695 ± 0.0211 | 0.1648 ± 0.0416 |
| TTR | 0.1856 ± 0.0619 | 0.2079 ± 0.0656 | 0.1637 ± 0.0193 | 0.1826 ± 0.0436 |

**Table S5.** List of 104 proteins directly involved in one or several diseases (see columns) as reported neXtProt medical tab (neXtProt 2018-09-03).

| gene name(s) | protein name | NX acc. code | Disease | Disease | Disease |
|---|---|---|---|---|---|
| **ADAM9** | Disintegrin and metalloprotein ase domain-containing protein 9 | NX_Q13443 | Cone-rod dystrophy 9 (CORD9) | | |
| **AGT** | Angiotensinoge n | NX_P01019 | Essential hypertension (EHT) | Renal tubular dysgenesis (RTD) | |
| **AHSG** | Alpha-2-HS-glycoprotein | NX_P02765 | Alopecia-mental retardation syndrome 1 (APMR1) | | |
| **ALB** | Serum albumin | NX_P02768 | Hyperthyroxinemia, familial dysalbuminemic (FDAH) | Analbuminemia (ANALBA) | |
| **APOA1** | Apolipoprotein A-I | NX_P02647 | High density lipoprotein deficiency 2 (HDLD2) | High density lipoprotein deficiency 1 (HDLD1) | Amyloidosis 8 (AMYL8) |
| **APOA2** | Apolipoprotein A-II | NX_P02652 | Familial renal amyloidosis due to Apolipoprotein AII variant 238269 | | |
| **APOB** | Apolipoprotein B-100 | NX_P04114 | Hypobetalipoproteinemia, familial, 1 (FHBL1) | Familial ligand-defective apolipoprotein B-100 (FDB) | |
| **APOE** | Apolipoprotein E | NX_P02649 | Alzheimer disease 2 (AD2) | Hyperlipoproteinemia 3 (HLPP3 | Sea-blue histiocyte disease (SBHD) |
| **APP** | Amyloid-beta A4 protein | NX_P05067 | Alzheimer disease 1 (AD1) | Cerebral amyloid angiopathy, APP-related (CAA-APP) [MIM:605714 | |
| **ATP6AP1** | V-type proton ATPase subunit S1 | NX_Q15904 | Immunodeficiency 47 (IMD47) | | |
| **ATP6AP2** | Renin receptor | NX_O75787 | Parkinsonism with spasticity, X-linked (XPDS) | Mental retardation, X-linked, with epilepsy (MRXE) | |
| **B2M** | Beta-2-microglobulin | NX_P61769 | Amyloidosis 8 (AMYL8) | Immunodeficiency 43 (IMD43) | |
| **B4GAT1** | Beta-1,4-glucuronyltrans ferase 1 | NX_O43505 | Muscular dystrophy-dystroglycanopathy congenital with brain and eye anomalies A13 (MDDGA13) | Walker-Warburg syndrome 899 | |
| **C1QA** | Complement C1q subcomponent subunit A | NX_P02745 | Complement component C1q deficiency (C1QD) | | |
| **C1QB** | Complement C1q subcomponent subunit B | NX_P02746 | Complement component C1q deficiency (C1QD) | | |
| **C1QC** | Complement C1q subcomponent subunit C | NX_P02747 | Complement component C1q deficiency (C1QD) | | |
| **C1R** | Complement C1r subcomponent | NX_P00736 | Ehlers-Danlos syndrome, periodontal type, 1 (EDSPD1) | | |
| **C1S** | Complement C1s subcomponent | NX_P09871 | Complement component C1s deficiency (C1SD) | Ehlers-Danlos syndrome, periodontal type, 2 (EDSPD2) | |
| **C3** | Complement C3 | NX_P01024 | Macular degeneration, age-related, 9 (ARMD9) | Complement component 3 deficiency (C3D) | Hemolytic uremic syndrome atypical 5 (AHUS5) |

| | | | | | |
|---|---|---|---|---|---|
| **C4A** | Complement C4-A | NX_P0C0L4 | Complement component 4A deficiency (C4AD) | Systemic lupus erythematosus (SLE) | |
| **C5** | Complement C5 | NX_P01031 | Complement component 5 deficiency (C5D) | | |
| **C6** | Complement component C6 | NX_P13671 | Complement component 6 deficiency (C6D) | | |
| **C7** | Complement component C7 | NX_P10643 | Complement component 7 deficiency (C7D) | | |
| **C8A** | Complement component C8 alpha chain | NX_P07357 | Complement component 8 deficiency, 1 (C8D1) | | |
| **C9** | Complement component C9 | NX_P02748 | Macular degeneration, age-related, 15 (ARMD15) | Complement component 9 deficiency (C9D) | |
| **CA2** | Carbonic anhydrase 2 | NX_P00918 | Osteopetrosis, autosomal recessive 3 (OPTB3) | | |
| **CEP152** | Centrosomal protein of 152 kDa | NX_O94986 | Microcephaly 9, primary, autosomal recessive (MCPH9) | Seckel syndrome 5 (SCKL5) | |
| **CEP290** | Centrosomal protein of 290 kDa | NX_O15078 | Joubert syndrome 5 (JBTS5) | Senior-Loken syndrome 6 (SLSN6) | Leber congenital amaurosis 10 (LCA10) |
| **CFB** | Complement factor B | NX_P00751 | Hemolytic uremic syndrome atypical 4 (AHUS4) | Complement factor B deficiency (CFBD) | Age-related macular degeneration 279 |
| **CFD** | Complement factor D | NX_P00746 | Complement factor D deficiency (CFDD) | | |
| **CFH** | Complement factor H | NX_P08603 | Basal laminar drusen (BLD) | Complement factor H deficiency (CFHD) | Macular degeneration, age-related, 4 (ARMD4) |
| **CFI** | Complement factor I | NX_P05156 | Macular degeneration, age-related, 13 (ARMD13 | Complement factor I deficiency (CFI deficiency) | Hemolytic uremic syndrome atypical 3 (AHUS3) |
| **CHI3L1** | Chitinase-3-like protein 1 | NX_P36222 | Schizophrenia (SCZD) | Asthma-related traits 7 (ASRT7) | |
| **CNTN1** | Contactin-1 | NX_Q12860 | Myopathy, congenital, Compton-North (MYPCN) | | |
| **COL1A1** | Collagen alpha-1(I) chain | NX_P02452 | Caffey disease (CAFFD) | Ehlers-Danlos syndrome, classic type (EDS) and 7A | Osteogenesis imperfecta |
| **COL1A2** | Collagen alpha-2(I) chain | NX_P08123 | Ehlers-Danlos syndrome, arthrochalasia type, 2 (EDSARTH2) | Osteogenesis imperfecta | |
| **COL3A1** | Collagen alpha-1(III) chain | NX_P02461 | Ehlers-Danlos syndrome 3 (EDS3) 4 | Familial aneurysm | |
| **COL6A1** | Collagen alpha-1(VI) chain | NX_P12109 | Bethlem myopathy 1 (BTHLM1) | Ullrich congenital muscular dystrophy 1 (UCMD1 | |
| **COL6A3** | Collagen alpha-3(VI) chain | NX_P12111 | Dystonia 27 (DYT27) | Bethlem myopathy 1 (BTHLM1) | Ullrich congenital muscular dystrophy 1 (UCMD1) |
| **CP** | Ceruloplasmin | NX_P00450 | Aceruloplasminemia (ACERULOP) | | |
| **CSF1R** | Macrophage colony-stimulating factor 1 receptor | NX_P07333 | Leukoencephalopathy, diffuse hereditary, with spheroids (HDLS) | Cancers | Inflammatory diseases |
| **CST3** | Cystatin-C | NX_P01034 | Amyloidosis KW-1008 | Age-related macular degeneration KW-0913 | |
| **CTSD** | Cathepsin D | NX_P07339 | Ceroid lipofuscinosis, neuronal, 10 (CLN10) | | |

| | | | | | |
|---|---|---|---|---|---|
| **DAG1** | Dystroglycan | NX_Q14118 | Muscular dystrophy-dystroglycanopathy limb-girdle C9 (MDDGC9) | Muscular dystrophy-dystroglycanopathy congenital with brain and eye anomalies A9 (MDDGA9) | |
| **DCN** | Decorin | NX_P07585 | Corneal dystrophy, congenital stromal (CSCD) | | |
| **DST** | Dystonin | NX_Q03001 | Neuropathy, hereditary sensory and autonomic, 6 (HSAN6) | Epidermolysis bullosa simplex, autosomal recessive 2 (EBSB2) | |
| **ECM1** | Extracellular matrix protein 1 | NX_Q16610 | Lipoid proteinosis (LiP) | | |
| **EFEMP1** | EGF-containing fibulin-like extracellular matrix protein 1 | NX_Q12805 | Doyne honeycomb retinal dystrophy (DHRD) | | |
| **ENPP2** | Ectonucleotide pyrophosphatase/phosphodiesterase family member 2 | NX_Q13822 | Obesity KW-0550 | | |
| **F2** | Prothrombin | NX_P00734 | Factor II deficiency (FA2D) | Ischemic stroke (ISCHSTR) | |
| **F5** | Coagulation factor V | NX_P12259 | Factor V deficiency (FA5D) | Budd-Chiari syndrome (BDCHS) | Ischemic stroke (ISCHSTR) |
| **FAT2** | Protocadherin Fat 2 | NX_Q9NYQ8 | Spinocerebellar ataxia 45 (SCA45) | | |
| **FBLN1** | Fibulin-1 | NX_P23142 | FBLN1-related developmental delay-central nervous system anomaly-syndactyly syndrome 404451 | | |
| **FBLN1** | Fibulin-1 | NX_P23142 | FBLN1-related developmental delay-central nervous system anomaly-syndactyly syndrome 404451 | | |
| **FGA** | Fibrinogen alpha chain | NX_P02671 | Congenital afibrinogenemia (CAFBN) | Amyloidosis 8 (AMYL8) | Dysfibrinogenemia, congenital (DYSFIBRIN) |
| **FGG** | Fibrinogen gamma chain | NX_P02679 | Congenital afibrinogenemia (CAFBN) | Dysfibrinogenemia, congenital (DYSFIBRIN) | |
| **FN1** | Fibronectin | NX_P02751 | Glomerulopathy with fibronectin deposits 2 (GFND2) | Spondylometaphyseal dysplasia, corner fracture type (SMDCF) | |
| **FUCA1** | Tissue alpha-L-fucosidase | NX_P04066 | Fucosidosis (FUCA1D) | | |
| **GM2A** | Ganglioside GM2 activator | NX_P17900 | GM2-gangliosidosis AB (GM2GAB) | | |
| **GSN** | Gelsolin | NX_P06396 | Amyloidosis 5 (AMYL5) | | |
| **HBD** | Hemoglobin subunit delta | NX_P02042 | Delta-beta-thalassemia 231237 | | |
| **HP** | Haptoglobin | NX_P00738 | Anhaptoglobinemia (AHP) | | |
| **HRG** | Histidine-rich glycoprotein | NX_P04196 | Thrombophilia due to histidine-rich glycoprotein deficiency (THPH11) | | |
| **HTRA1** | Serine protease HTRA1 | NX_Q92743 | Cerebral arteriopathy, autosomal recessive, with subcortical infarcts and leukoencephalopathy (CARASIL) | Macular degeneration, age-related, 7 (ARMD7) | |
| **IGF2** | Insulin-like growth factor II | NX_P01344 | Silver-Russell syndrome (SRS) | Growth restriction, severe, with distinctive facies (GRDF) | |
| **IGFALS** | Insulin-like growth factor-binding protein | NX_P35858 | Acid-labile subunit deficiency (ACLSD) | | |

| | | | | | |
|---|---|---|---|---|---|
| | complex acid labile subunit | | | | |
| **IGFBP7** | Insulin-like growth factor-binding protein 7 | NX_Q16270 | Retinal arterial macroaneurysm with supravalvular pulmonic stenosis (RAMSVPS) | | |
| **IGHA1** | Immunoglobulin heavy constant alpha 1 | NX_P01876 | Multiple myeloma (MM) | | |
| **IGHG1** | Immunoglobulin heavy constant gamma 1 | NX_P01857 | Multiple myeloma (MM) | | |
| **IGHM** | Immunoglobulin heavy constant mu | NX_P01871 | Agammaglobulinemia 1, autosomal recessive (AGM1) | | |
| **IGKC** | Immunoglobulin kappa constant | NX_P01834 | Immunoglobulin kappa light chain deficiency (IGKCD) | | |
| **KNG1** | Kininogen-1 | NX_P01042 | High molecular weight kininogen deficiency (HMWK deficiency) | | |
| **KRT1** | Keratin, type II cytoskeletal 1 | NX_P04264 | Epidermolytic hyperkeratosis (EHK) | Ichthyosis hystrix, Curth-Macklin type (IHCM) | Keratoderma, palmoplantar, non-epidermolytic (NEPPK) |
| **KRT10** | Keratin, type I cytoskeletal 10 | NX_P13645 | Epidermolytic hyperkeratosis (EHK) | Ichthyosis annular epidermolytic (AEI) | Erythroderma, ichthyosiform, congenital reticular (CRIE) |
| **KRT13** | Keratin, type I cytoskeletal 13 | NX_P13646 | White sponge nevus 2 (WSN2) | | |
| **LMAN1** | Protein ERGIC-53 | NX_P49257 | Factor V and factor VIII combined deficiency 1 (F5F8D1) | | |
| **LOX** | Protein-lysine 6-oxidase | NX_P28300 | Aortic aneurysm, familial thoracic 10 (AAT10) | | |
| **LTBP2** | Latent-transforming growth factor beta-binding protein 2 | NX_Q14767 | Glaucoma 3, primary congenital, D (GLC3D) | Microspherophakia and/or megalocornea, with ectopia lentis and with or without secondary glaucoma (MSPKA) | Weill-Marchesani syndrome 3 (WMS3) |
| **LYZ** | Lysozyme C | NX_P61626 | Amyloidosis 8 (AMYL8) | | |
| **MEGF8** | Multiple epidermal growth factor-like domains protein 8 | NX_Q7Z7M0 | Carpenter syndrome 2 (CRPT2) | | |
| **MINPP1** | Multiple inositol polyphosphate phosphatase 1 | NX_Q9UNW1 | Thyroid cancer, non-medullary, 2 (NMTC2) | | |
| **MMP9** | Matrix metalloproteinase-9 | NX_P14780 | Intervertebral disc disease (IDD) | Metaphyseal anadysplasia 2 (MANDP2) | |
| **PLG** | Plasminogen | NX_P00747 | Plasminogen deficiency (PLGD) | | |
| **PROS1** | Vitamin K-dependent protein S | NX_P07225 | Thrombophilia due to protein S deficiency, autosomal dominant (THPH5) | Thrombophilia due to protein S deficiency, autosomal recessive (THPH6) | |
| **PSAP** | PSAP | NX_P07602 | Combined saposin deficiency (CSAPD) | Leukodystrophy metachromatic due to saposin-B deficiency (MLD-SAPB) [MIM:249900 | Gaucher disease, atypical, due to saposin C |

| | | | | | |
|---|---|---|---|---|---|
| | | | | | deficiency (AGD) |
| **PSAP** | Prosaposin | NX_P07602 | Combined saposin deficiency (CSAPD) | Leukodystrophy metachromatic due to saposin-B deficiency (MLD-SAPB) [MIM:249900 | |
| **PSAP** | Saposin-C | NX_P07602 | Combined saposin deficiency (CSAPD) | Gaucher disease, atypical, due to saposin C deficiency (AGD) | |
| **RBP4** | Retinol-binding protein 4 | NX_P02753 | Retinal dystrophy, iris coloboma, and comedogenic acne syndrome (RDCCAS) | Microphthalmia, isolated, with coloboma, 10 (MCOPCB10) | |
| **RELN** | Reelin | NX_P78509 | Lissencephaly 2 (LIS2) | Epilepsy, familial temporal lobe, 7 (ETL7) | |
| **RNASET2** | Ribonuclease T2 | NX_O00584 | Leukoencephalopathy, cystic, without megalencephaly (LCWM) | | |
| **RPS23** | 40S ribosomal protein S23 | NX_P62266 | Brachycephaly, trichomegaly, and developmental delay (BTDD) | | |
| **SERPINA1** | Alpha-1-antitrypsin | NX_P01009 | Alpha-1-antitrypsin deficiency (A1ATD) | | |
| **SERPINA3** | Alpha-1-antichymotrypsin | NX_P01011 | Alpha-1-antichymotrypsin deficiency 93594 | | |
| **SERPINA7** | Thyroxine-binding globulin | NX_P05543 | Genetic variants in SERPINA7 influence the serum levels of thyroxine-binding globulin and define the thyroxine-binding globulin quantitaive trait locus (TBGQTL) | | |
| **SERPINC1** | Antithrombin-III | NX_P01008 | Antithrombin III deficiency (AT3D) | | |
| **SERPIND1** | Heparin cofactor 2 | NX_P05546 | Thrombophilia due to heparin cofactor 2 deficiency (THPH10) | | |
| **SERPINF1** | Pigment epithelium-derived factor | NX_P36955 | Osteogenesis imperfecta 6 (OI6) | | |
| **SERPINF2** | Alpha-2-antiplasmin | NX_P08697 | Alpha-2-plasmin inhibitor deficiency (APLID) | | |
| **SERPING1** | Plasma protease C1 inhibitor | NX_P05155 | Hereditary angioedema (HAE) | | |
| **SPARC** | SPARC | NX_P09486 | Osteogenesis imperfecta 17 (OI17) | | |
| **SPP1** | Osteopontin | NX_P10451 | Systemic lupus erythematosus 536 | | |
| **TF** | Serotransferrin | NX_P02787 | Atransferrinemia (ATRAF) | | |
| **TGFBI** | Transforming growth factor-beta-induced protein ig-h3 | NX_Q15582 | Corneal dystrophy, epithelial basement membrane (EBMD) | | |
| **TTR** | Transthyretin | NX_P02766 | Amyloidosis, transthyretin-related (AMYL-TTR) [MIM:105210]: | Hyperthyroxinemia, dystransthyretinemic (DTTRH) | Carpal tunnel syndrome 1 (CTS1) |

**Table S6.** List of 23 proteins which are direct targets of drug(s) (see column) used to treat various pathologies (see column) in past or ongoing therapeutic trials as referenced in the clinical trial database (https://clinicaltrials.gov/, see last column) as of December 2018.

| gene name(s) | protein name | Drug(s) | Pathology | Clinical trial |
|---|---|---|---|---|
| **APOB** | Apolipoprotein B-100 | mipomersen (antisens antiApoB) | Diabetes | NCT00770146 |
| **APP** | Amyloid-beta A4 protein | Therapeutic antibodies, secretase inhibitors… | Alzheimer | NCT00762411 |
| **C1QA** | Complement C1q subcomponent subunit A | ANX007 (C1q inhibitor monoclonal antibody) | Glaucoma | NCT03488550 |
| **CD14** | Monocyte differentiation antigen CD14 | IC14 (monoclonal antibody against human CD14) | Amyloid Lateral Sclerosis | NCT03474263 |
| **CHI3L1** | Chitinase-3-like protein 1 | T-ChOS chitooligosaccharides | Cancer | NCT03320525 |
| **CLU** | Clusterin | Custirsen (OGX-011, antisens against clusterin) | Cancer | NCT00471432 |
| **CSF1R** | Macrophage colony-stimulating factor 1 receptor | DCC-3014, selective inhibitor of CSF1R | Leukemia, cancer | NCT03069469 |
| **CTSD** | Cathepsin D | Cathepsin-S Inhibitor | Rheumatoid Arthritis | NCT00425321 |
| **CXCL12** | Stromal cell-derived factor 1 | NOX-A12 (CXCL12 inhibitor) | Cancer | NCT01486797 |
| **DKK3** | Dickkopf-related protein 3 | ad-REIC/DKK3 vaccine | Prostate Cancer | NCT01197209 |
| **ENPP2** | Ectonucleotide pyrophosphatase/phosphodiesterase family member 2 | Autotaxin (ENPP2) inhibitor | Idiopathic Pulmonary Fibrosis (IPF) | NCT02738801 |
| **FN1** | Fibronectin | ocriplasmin (protease against fibronectin and laminin) | Ophtalmology | NCT02322229 |
| **FOLR2** | Folate receptor beta | OTL38 (Folate receptor inhibithor) | Cancer | NCT02317705 |
| **GM2A** | Ganglioside GM2 activator | Miglustat (Glucocerebrosidase inhibitor) | Gangliosidosis | NCT00418847 |
| **GSN** | Gelsolin | rhu-pGelsolin | Critical care | NCT00671307 |
| **IGF2** | Insulin-like growth factor II | xentuzumab ( humanized anti-IGF monoclonal antibody) | Breast cancer | NCT02123823 |
| **IGFBP2** | Insulin-like growth factor-binding protein 2 | IGFBP-2 Vaccine | Cancer | NCT03029611 |
| **KLK6** | Kallikrein-6 | MDCO-2010, kallikrein inhibitor | Haematology | NCT01535222 |
| **MMP9** | Matrix metalloproteinase-9 | MMP9 inhibitor (AZD1236..) | Cancer, neurodegenerative diseases | NCT01007929 |
| **SERPINF1** | Pigment epithelium-derived factor | AdGVPEDF.11D (adenovirus vector PEDF) | Macular Degeneration | NCT00109499 |
| **SERPINF2** | Alpha-2-antiplasmin | Anti-alpha2-antiplasmin (α2AP) Monoclonal Antibody TS23 | Cadiovascular disease | NCT03001544 |
| **SERPING1** | Plasma protease C1 inhibitor | CINRYZE® (C1 esterase inhibitor) | Neuromyelitis Optica | NCT01759602 |
| **TTR** | Transthyretin | Fx-1006A ( transthyretin chaperone) | Amyloidosis | NCT00630864 |

### B. Supplementary Figures

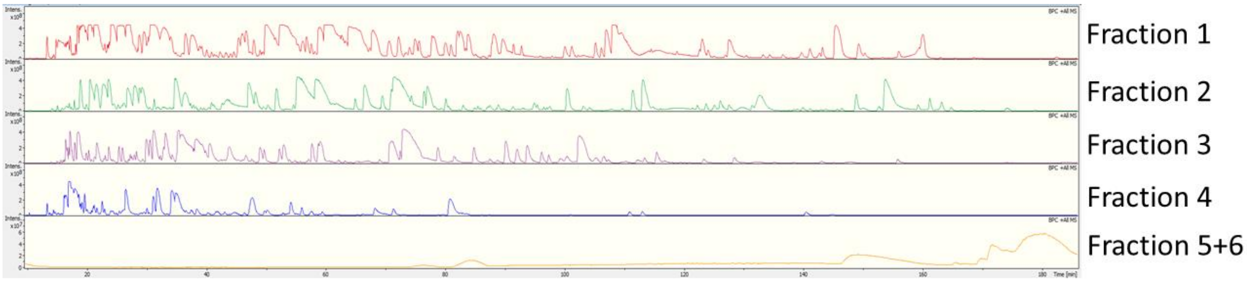

**Figure S1.** Typical base peak chromatograms.

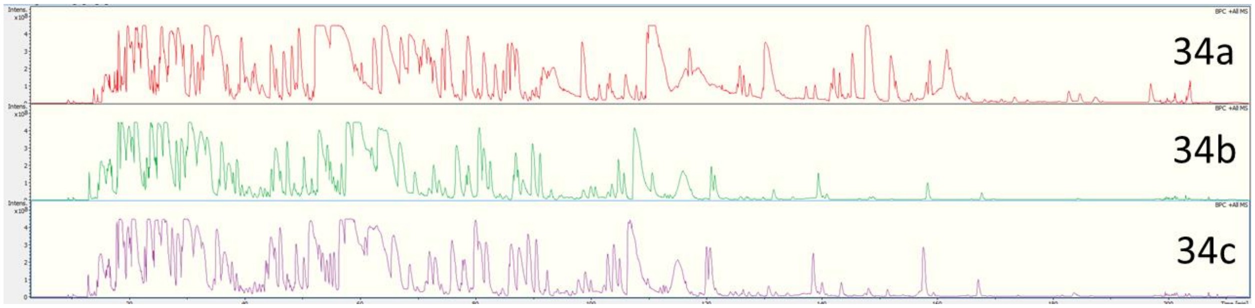

**Figure S2.** Example of a fraction with poor and excessively shifted chromatography in replicate a (top).

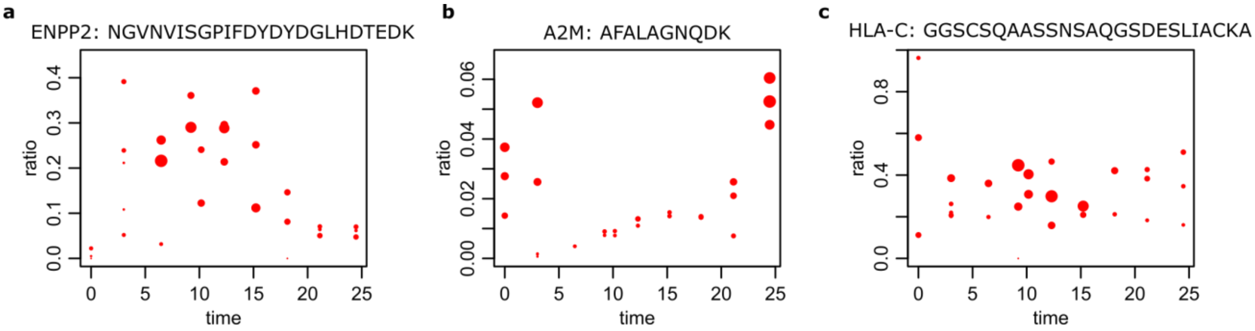

**Figure S3.** Examples to filter out. (**a**) Ectonucleotide pyrophosphatase/phosphodiesterase family member 2 peptide featuring a trend in which one could recognize growth followed by decay but data suffer from excessive dispersion. (**b**-**c**) Two examples of *random* point configurations for Alpha-2-macroglobulin (A2M) and HLA class I histocompatibility antigen, Cw-3 alpha chain (HLA-C), peptides in which no pattern should be sought.

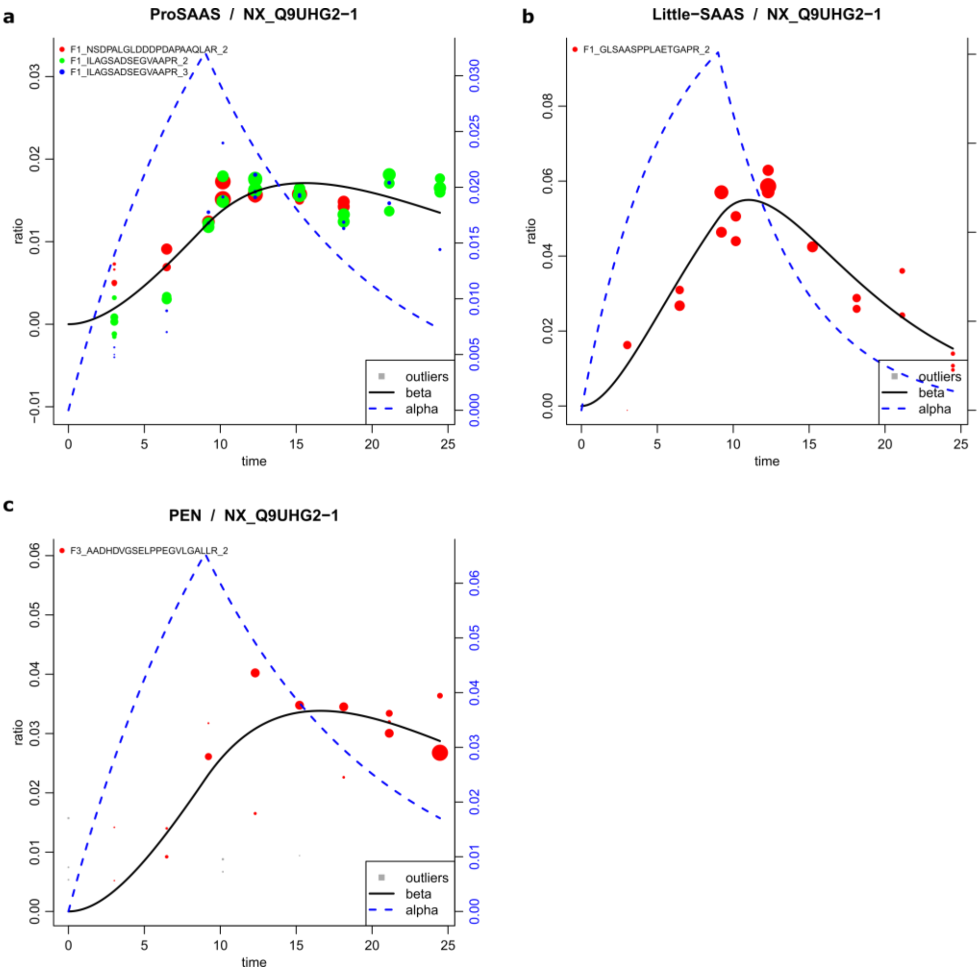

**Figure S4.** Propeptide ProSAAS **(a)**, Little-SAAS **(b)**, and PEN **(c)**.

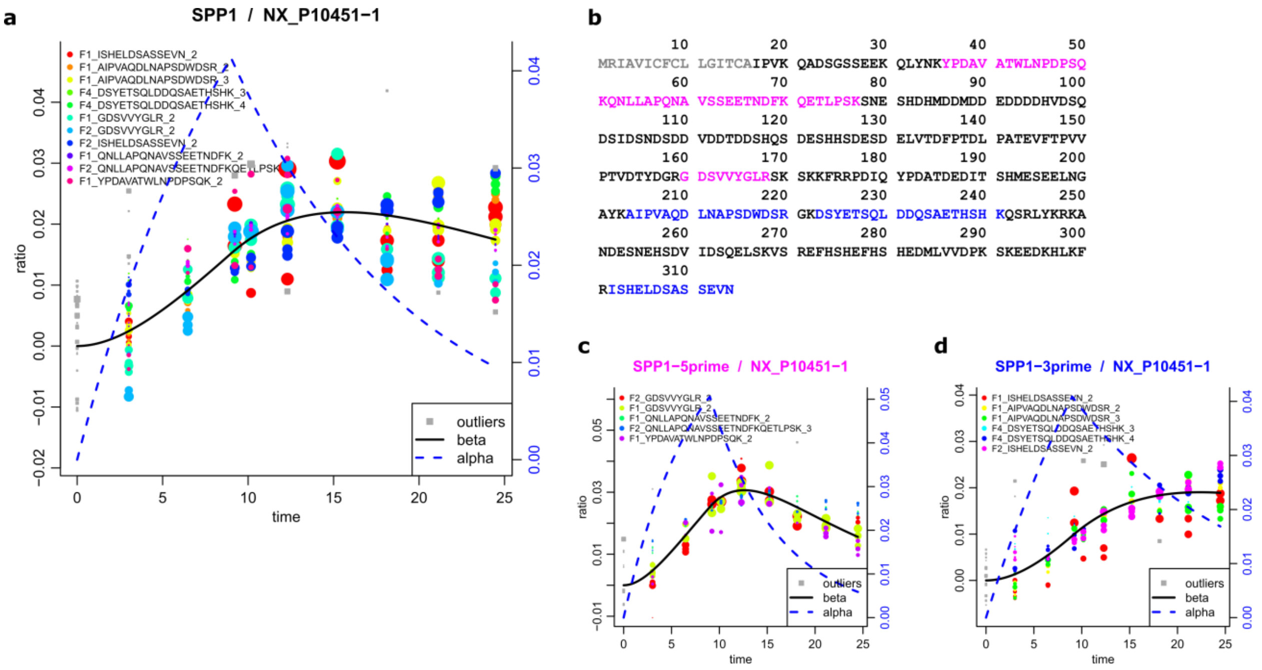

**Figure S5**. Osteopontin (SPP1) peptides **(a)** segregate over the primary sequence according to their dynamics suggesting a potential enzymatic cleavage between the blue and fuchsia regions indicated in panel **(b)**. Signal peptide in gray. **(c,d)** Individual observations split after their N-term ($k_c = 0.139$ 1/h) and C-term ($k_c = 0.057$ 1/h) locations.

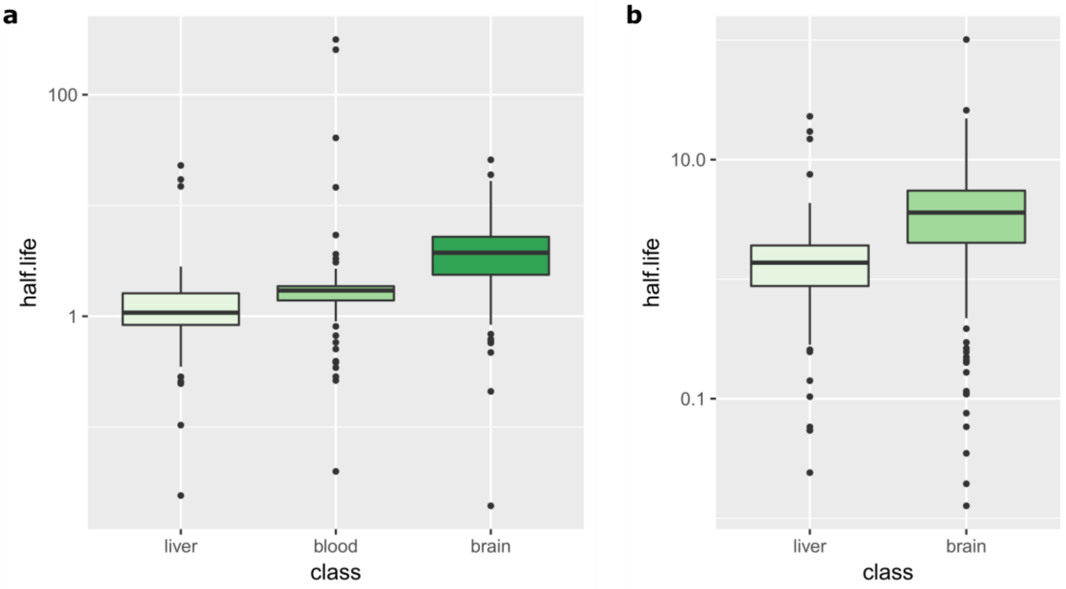

**Figure S6.** Comparison of protein half-life in mouse tissues. **(a)** We selected 124 proteins detected in all 3 tissues. **(b)** We selected 454 proteins detected in brain and liver. Within both boxplots, Kolmogorov-Smirnov test P-values comparing any 2 half-life sets was below $10^{-12}$. The ranking brain > blood > liver was already observed over all the proteins in each tissue by the author of the mouse study (Price *et al.*, *PNAS*, 2010).

### C. Supplementary Proteomics Materials and Methods

**Free $^{13}C_6$-Leu concentration in patient blood and CSF samples**

The ratio $^{13}C_6$-Leu/Leu was determined by ultra-performance liquid chromatography tandem mass spectrometry (UPLC-MS/MS) using a Waters® Xevo-TQD (Waters Corp.,Milford, MA, USA). Ion-pairing chromatography was performed on a Acquity UPLC Waters® BEH C18 1.7µm column at a flow rate of 0.65 ml/min with mobile phases containing Tridecafluoroheptanoic acid (Sigma Saint-Louis, Missouri, USA), adapted from a published method(Waterval et al., 2009). Monitored transitions in MRM and ESI-positive mode were 131.8 >86 for Leu and 137.8 >90.9 for $^{13}C_6$-Leu.

**CSF sample initial preparation**

All the sample preparation steps were performed in triplicate on automation system (BRAVO, Agilent). Three times 20uL of CSF were denaturated and reduced (DTT) (30uL of 8M urea / 20mM DTT / 100mM Tris pH8.5), alkylated (6uL of 400mM IAA / 1M Tris pH11), digested overnight (37°C) with LysC/trypsin mix (diluted with 200uL 20mM Tris pH8.5 / 2mM DTT; and addition of 9uL 0.05ug/uL LysC/trypsin), and acidified by formic acid addition. Desalting was performed by C18 Tips (Agilent Technologies, Santa Clara, CA, USA) primed with 50uL of 70% ACN/ 0.1% TFA, equilibrated with 50uL of 0.1% TFA, loaded with sample, washed two times with 50uL of 0.1% TFA and eluted with 50uL of 70% ACN/ 0.1% TFA.

**Tryptic peptide fractionation**

After peptide clean-up step, samples were dried on speedvac and resuspended with 100uL of the equilibration buffer (10mM H3PO4, 25% ACN, pH 2.6). SCX cartridges (Agilent Technologies, Santa Clara, CA, USA) were primed with 100uL of 350mM KCl/10mM H3PO4/25% ACN/pH 2.6; equilibrated with 50uL of 10mM H3PO4/25% ACN/pH 2.6; loaded with samples, washed with 25uL of 10mM H3PO4/25% ACN/pH 2.6; and sequentially eluted by salt steps gradient in common buffer (10mM H3PO4/25% ACN/pH 2.6) as follows: 115uL of elution buffer 1 (30mM KCl); 115uL of elution buffer 2 (50mM KCl) ; 135uL of elution buffer 3 (85mM KCl) ; 115uL of elution buffer 4 (115mM KCl) ; 90uL of elution buffer 5 (150mM KCl) ; and 40uL of elution buffer 6 (500mM KCl at pH 9.5). First 45uL of the fraction 1 were removed because they did not contain peptides. Fraction 6 was acidified by 2uL of FA and pooled with fraction 5. The 5 resulting fractions were dried and cleaned-up on C18 tips as above.

**Mass Spectrometry Analysis**

Fractionated and cleaned samples were resuspended in 10uL of A phase (A = 0.1% formic acid, 2% acetonitrile in water) and injected on nanoRSLC Ultimate 3000 (Thermo).

NanoFlow LC was coupled to QTOF MS instrument (Impact II, Bruker Daltonics) through captive spray ion source operating with nanobooster. Samples were desalted and pre-concentrated on-line on a PepMap u-precolumn (300 um x 5 mm, C18 PepMap 100, 5 um, 100 Angstrom). The capillary pump worked at 20uL/min with phase constituted by 0.05% TFA, 2% acetonitrile in water for 3 minutes.

Peptides were then transferred to an analytical column (75 um x 500 mm; Acclaim Pepmap RSLC, C18, 2um, 100 Angstrom) to perform separation. A gradient consisting of 5-26% B for 192 min and 90% B for 10 min (A = 0.1% formic acid, 2% acetonitrile in water; B = 0.1% formic acid in acetonitrile) at 400 nL/min, 50°C, was used to elute peptides from the reverse-phase column. Separated peptides were ionized on Captive Spray (1200V, dry gas: 3 l/min at 150°C) with nanobooster (0.2 Bar of Nitrogen boiling in acetonitrile/0.1% FA) and analyzed on the QTOF instrument.

In order to generate a reference peptide library, and considering the limited amount of material available, we pooled 2uL of each fraction at each time point to obtained 5 pooled samples (fraction 1 pool, fraction 2 pool, etc.). Data dependent acquisition was subsequently performed to identify peptides and to generate the peptide library. For all full-scan measurements, a lock-mass (m/z 1222, Hexakis(1H, 1H, 4H-hexafluorobutyloxy) phosphazine, chip cube high mass reference, Agilent) was used as internal calibrant. Instant Expertise software selected as many as possible most intense ions per cycle of 3 seconds. They were fragmented in the collision cell and analyzed in TOF. MS2 accumulation was dependent to the MS1 level (MS1 5000cts = spectra rate of 2Hz, to MS1 50000cts = spectra rate of 50Hz). Peptide fragmentation was conducted with nitrogen gas on the most abundant and at least doubly-charged to 5 times charged ions detected in the initial MS Scan. Active exclusion was performed after 1 spectrum during 2 minutes unless the precursor ion exhibited an intensity higher than times the one of the previous scan. Collision energy for the CID was adapted to the charge and the ion mass.

**Peptide identification**

All the MS/MS spectra were searched against NeXtProt (v 2018-01-17) database by using the Mascot v 2.6.0 algorithm (Matrix Science). Search parameters were the followings, enzyme: trypsin; variable modifications: oxidation (M) and deamidated (N,Q); fixed modifications: carbamidomethyl (C); missed cleavages: maximum of 2; instrument type CID: ESI-QUAD-TOF; peptide tolerance: 10.0 ppm; MS/MS tolerance: 0.05 Da; peptide charge: 1+, 2+ and 3+; mass: monoisotopic; C13: 1; minimum peptide length: 5; peptide decoy: ON; FDR [%]: 1; percolator: on; ions score cut-off: 12; ions score threshold for significant peptide IDs: 12.

**Peptide library generation**

Identified MS/MS spectra by Mascot allowed us to build the peptide library, which was comprised of the peptide sequences, the MS/MS spectra, and the retention times. Skyline 4.1 (MacCoss Lab.) was used to generate it with a cut-off PeptideProphet score of 0.95.

**Profile MS analysis**

$^{13}C_6$-leucine incorporation was followed by LC-MS profiling. A mass range of 300 to 1400 m/z was scanned at 0.5 Hz. The other parameters were the same as in the DDA mode.

Skyline 4.1 was used to process raw MS data. Identified leucine-containing peptides were recovered from the peptide library along with their m/z and retention times to perform profile analysis. Transition settings were adjusted to extract monoisotopic masses for precursor ions with a positive charge of 1-4 in the mass range of 300-1400 m/z.

MS1 filtering was operating on TOF data, resolving power at 60000, isotope analysis based count and on 4 peaks. Retention time tolerance was set to 10 minutes. Peptides contained in the library were directly imported as compounds. The isotope label was specified as “Heavy” corresponding to $^{13}C_6$ Leucine, and the “internal standard type” was set up to “Light”. Automatic peak detection with default parameters and area extraction were performed by the software. A report in csv format was exported, which included the protein name, replicate name, peptide, precursor, product m/z, area under the peak.

**Targeted, triple-quad MS analysis to validate chosen peptide dynamics**

*Sample preparation*

20uL of CSF were prepared in duplicates as described above but without tryptic peptide fractionation step. After clean-up, peptides were resuspended in 20uL of A phase and injected on LC-QqQ (1290 LC – 6490 QqQ, Agilent Technologies).

*MRM method*

LC separation was carried out with a 1290 LC system (Agilent technologies). Peptides were resolved on reversed-phase separative column (RRHD Eclipse Plus C18, 2.1x150mm, 1.8um) maintained at 60°C and at 400uL/min. A 30 min multi-step gradient was used, starting with 2.7% of solvent B (10% water, 90% acetonitrile and 0.1% formic acid), increasing to 9.9% of solvent B in 2 minutes, increasing at 17.1% of solvent B at 15 minutes, 26.1% of solvent B at 22 minutes, 40.5% of solvent B at 25 minutes. The column was then flush by 81% of solvent B at 27 minutes during two minutes and then re-conditionned with 2.7% of solvent B.

Peptide analyses were conducted on a QqQ MS system (6490, Agilent technologies), equipped with a Agilent Jet-Stream ESI interface operated in the positive ion mode. The ESI source was set as follows: capillary tension 3500 V, nozzle voltage 300 V, nebulizer 30 psi, gas flow 15l/min, gas temperature 150°C, sheath gas flow 11l/min, sheath gas temperature 250°C.

The MS instrument worked in dynamic MRM with a retention time window of 4.5 minutes and a maximum cycle time fixed at 700ms. One peptide per protein and 3 transitions by peptide were required. By considering the light and the heavy species, they were hence 6 transitions per proteins.

Precursor ions were transferred inside the first quadrupole with ion funnel RF, high pressure was set to 150 V and low pressure to 60V. Ion transfer was performed with a fragmentor voltage of 380V and with a cell accelerator voltage of 5V. Quadrupoles operated in unit/unit resolution. Collision energies (CE) were optimized transitions dependent. All the monitored transitions, with optimized parameters, are provided as **Supplementary Data**. We implemented MRM analysis using a larger panel of proteins that was originally designed for another project and which was comprised of 54 proteins. Manual inspection of the data showed that only the 26 reported in this study gave an exploitable signal. Skyline was directed to extract data for the 26 detected proteins only.

*Data treatment*

Data processing was realized with Skyline 4.1 Software, parameters were adapted to low resolution data operating in MRM mode. All the replicates were loaded on the software. Automatic method for peak detection (default scoring) was used and validated by manual inspection. Data were exported in the same way as for the high resolution MS data above to a csv file.

### D. Supplementary Bioinformatics Materials and Methods

**Skyline export initial processing and filtering**

A Perl script was written to separate Leu-containing peptides from non-Leu-containing peptides and obtain two separate files. Leu-peptides were used for turnover calculations while non-Leu-peptides allowed us to estimate the false discovery rate of our analysis pipeline (see article text). Beyond this initial script, all the data analysis and modeling was implemented in R.

Initial filtering first regrouped the MS peaks from the same spectra, i.e. the light and heavy signal of a same peptide in a same LC-MS spectrum, as Skyline exports individual peaks on separate lines. Then, the most intense isotope was identified according to the majority rule: most frequently the most intense isotope over times 9.22 to 24.47 in the light species. The maximum intensity of the selected isotope was subsequently required to be higher than $10^5$ (light species again). This threshold was empirically determined to ensure reasonably strong heavy peptide signals, i.e. above noise. Finally, we imposed to have at least 7 out of 10 time points with a minimum of one pair of light/heavy signals available. In most cases, a lot more replicate pairs were available due to the 3 technical LC-MS replicates and potential multiple spectra for a single peptide. Here, by peptide, we mean a peptide in a specific fraction and at a specific charge state, what we called an observation in the article.

**A 2-component mathematical model**

There is an abundant literature on label incorporation dynamics in different settings (*in vitro*, cell lines, animal models) and 2-compartment models are well-established for this purpose. As explained in the article, non-local synthesis is addressed by acknowledging that the appearance and clearance of labelled, newly synthesized proteins and hence their peptides is the net effect of local synthesis and degradation as well as non-local synthesis coupled with transport in both directions through biological compartments, e.g. CSF/blood or CSF/CNS. That is, the mathematical model will follow the logic of local synthesis and degradation but the respective rates must be understood as the net contribution of local and non-local events. Most protocols provide the tracer over the whole experiment(Price et al., 2012, 2010). Following the general line presented by Rahman, *et al*.(Rahman et al., 2016), we adapted this framework to the specifics of SILK: injection of the tracer ($^{13}C_6$-Leu) during the first 9 hours of the experiment that lasted 24.47 hours in total. Limited time injection is introduced in the model through a function $f(t)$ representing $^{13}C_6$-Leu availability over time and taking values 0 or 1. Considering the free

$^{13}C_6$-Leu CSF and plasma concentration curves reported in the original SILK publication(Bateman et al., 2006) and our own data (**SupTable 2**), a simple choice for $f(t)$ is $f(t) = 0$ if $t \leq 0$ or $t > 9$, and $f(t) = 1$ otherwise. For a given protein R, the first stage of the model is an ordinary differential equation (ODE) representing the integration of Leu from the free pool in R synthesis: $A' = k_0 - k_a A$, with $A$ the amount of Leu in the free pool (Leu available protein synthesis but not used yet), $k_0$ the rate of Leu entering the free pool, and $k_a$ the *appearance* rate of R in the CSF (synthesis rate if synthesis would be purely local). Denoting $A = A_L + A_H$ the sum of light and heavy species, we have $A'_H = \lambda f(t) k_0 - k_a A_H$, with $\lambda$ the rate of $^{13}C_6$-Leu availability ($^{13}C_6$-Leu and Leu availability can differ according to where R is synthesized (organ, cell, etc.)). The second stage of the model captures the integration of amino acids in proteins: $P' = k_a A - k_c P$, with $P$ the abundance of R and $k_c$ its CSF *clearance rate*. $P$ is the sum of light and heavy proteins ($P = P_L + P_H$) and we naturally have the relationship $P'_H = k_a A_H - k_c P_H$. Since we were only interested in heavy/light ratios and LC-MS only provides relative quantitation through peak areas, we introduced $\alpha = \frac{A_H}{A_L + A_H}$, the proportion of integrated $^{13}C_6$-Leu, and $\beta = \frac{P_H}{P_L + P_H}$, the proportion of $^{13}C_6$-Leu labelled copies of R in the CSF. Assuming that the system is in a steady state implies $P' = 0 = A'$ and $k_a = k_c$, otherwise proteins would accumulate or disappear. Following the usual rules of derivation, we obtain:

$$\begin{cases} \alpha' = (\lambda f(t) - \alpha) k_c \\ \beta' = (\alpha - \beta) k_c \end{cases} \qquad (1)$$

and $k_0 = k_c A$, which does not need to be computed since we are only interested in the ratio $\alpha$ and not $A$ itself. The absence of $^{13}C_6$-Leu at $t = 0$ imposes the initial conditions $\alpha(0) = 0 = \beta(0)$. By Carathéodory theory and its extensions, Eq. (1) is well-posed despite the discontinuity in $f(t)$. Given the $\frac{P_H}{P_L + P_H}$ ratios (RIAs) obtained experimentally at different times for R, we used a numerical algorithm (quasi-Newton implemented in R optim, method="BFGS") to fit $\lambda, k_a, k_c$ by integrating (1) numerically and minimizing a weighted sum of squared errors. The heavy, $^{13}C_6$-Leu-labelled peptide signal intensity is lower that the light peptide intensity, typically by one order of magnitude (see below). We thus reasoned that among the many replicate experimental RIA values available, the most intense heavy peptide signals provided more reliable data. We empirically found that weights proportional to the square root of the heavy peptide intensity provided satisfying results overall.

For the simple step function $f(t)$ above it is possible to find piecewise exact solutions to Eq. (1) by the method of variation of constants applied before and after 9h separately. Note that to ensure continuity in $\alpha(t)$ and differentiability in $\beta(t)$ would require adjusting the initial conditions of the second part of the solution at 9h. Nonetheless, we preferred employing a numerical integrator to solve Eq. (1) to have the freedom to explore with different $f(t)$. The numerical integrator is Ernst Hairer's radau5 method(Hairer and Wanner, 1996) since $\alpha(t)$ can be stiff for certain proteins, notably when processing bad cases to be rejected. An "event" was set at 9h to deal with the discontinuity properly and maintain the order of the numerical integrator

Price, *et al*. published a study of mouse protein turnover in brain, plasma and liver in which they employed a simpler 1-compartment model(Price et al., 2010). In a paper by Guan, *et al*., the same team revisited its data applying a 2-compartment model(Guan et al., 2012) for better fitting. In this publication, they derived the 2-compartment model equations following a slightly different route than Rahman, *et al*. reasoning on compartment volumes instead of rate of availability, which resulted in equations identical to the non stochastic model in Rahman, *et al*. (Eq. (2) in Rahman, *et al*.; Eqs (16) and (17) in Guan, *et al*.). Our model, by imposing $k_a = k_c$ and modeling different availability of the tracer through $\lambda$, avoids mixing availability with an appearance rate that would be given the freedom to be different from the clearance rate for no obvious reason. According to the nature of Price *et al*. data, no pulse-chase but continuous labeling, it is expected that our model would yield values of $\lambda$ close to 1 for most proteins (food $^{15}N$ labelled >99.5% and long duration of the experiment). Indeed, processing their data with our model we found $\lambda$ close to 1 for a majority of proteins (**Fig. S7**). The analytical solution of $\beta$ in this case is

$$\beta(t) = \lambda\left(1 - k_c t e^{-k_c t} - e^{-k_c t}\right),$$

which can be found by Laplace's variation of constants method.

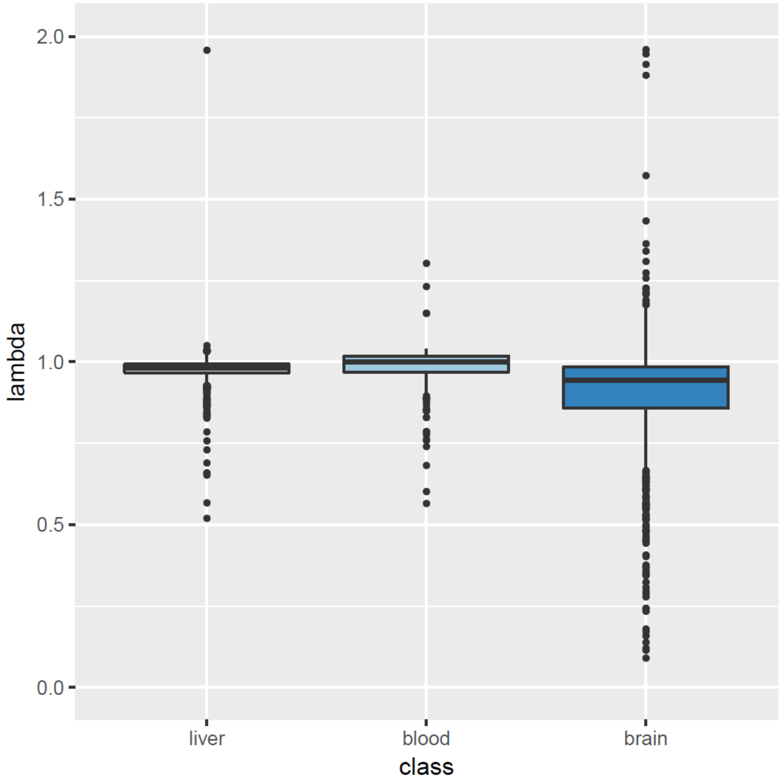

**Figure S7.** Distribution of $\lambda$ for Price *et al*. data sets.

Guan *et al*. introduced a 3-compartment model to account for recycled tracers consecutive to protein degradation, which was applied to their liver data. Rahman *et al*. introduced a Gaussian process to increase the accuracy of their model. In our hands, the simple dynamical system we proposed did not require any extension to accurately fit data, in particular liver data. For instance, the transitional endoplasmic reticulum ATPase (Vcp) example discussed in Guan *et al*. was fit without any difficulty (**Fig. S8**).

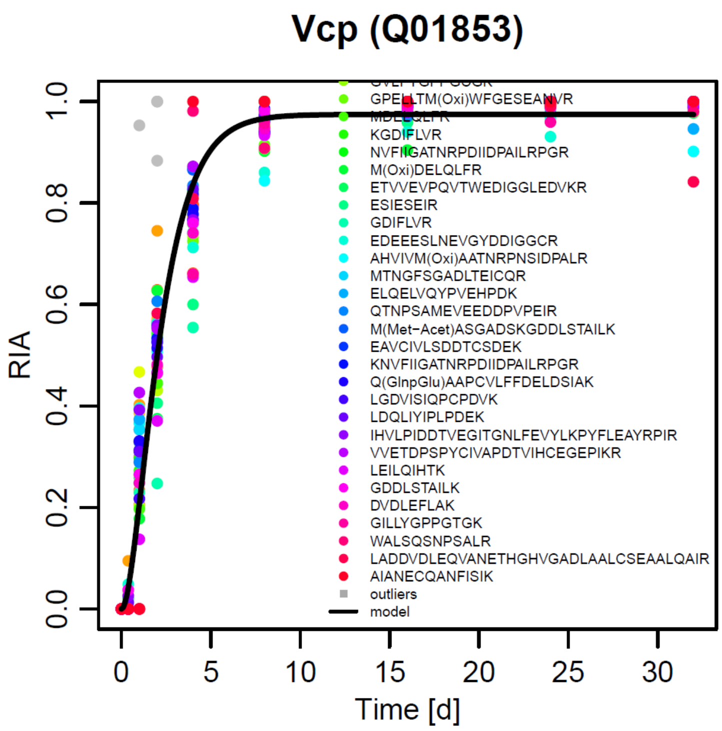

**Figure S8.** Transitional endoplasmic reticulum ATPase (Vcp) liver dynamics in Price *et al*. data.

**The impact of noise in the data**

Comparing multiple observations available for a single protein, we observe vertical shifts in a number of cases (**Fig. S9**). Given the limited amount of $^{13}C_6$-Leu material, labelled peptide MS signals are typically 30 to 100 times smaller thus making the calculation of RIA sensitive to noise occasionally. We believe that this is responsible for the observed vertical shift as indicated by the following estimation. Assuming both multiplicative ($r$), technical, and additive ($n$), chemical and technical, noise, heavy to total MS signal proportion, which is $\frac{H}{L+H}$, becomes $\frac{(1+r)H+n}{(1+r)(L+H)+2n} = \frac{(1+r)H}{(1+r)(L+H)+2n} + \frac{n}{(1+r)(L+H)+2n} = \frac{H}{(L+H)+2n} + \frac{n}{(1+r)(L+H)+2n}$. The first term is close to $\frac{H}{L+H}$, since $n$ is expected to be very small compared to $L$, which we imposed (above) to be rather intense. The second term, instead, cannot be neglected as $H$ can be closer to the noise level. That is, this simple analysis suggest that indeed $\frac{H}{L+H} \cong \frac{H}{L+H} + \varepsilon$, $\varepsilon$ representing a visible vertical shift of the RIA.

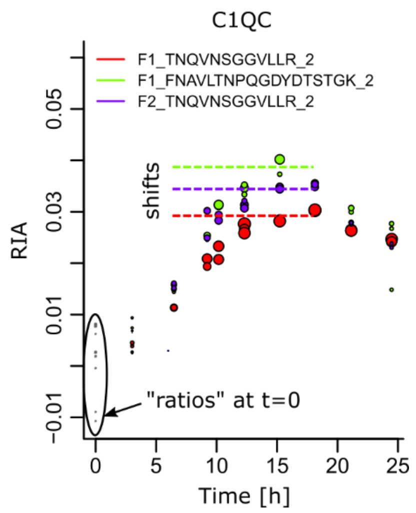

**Figure S9.** Examples of vertical shifts in RIAs. Note that the phenomenon can impact the same peptide in different fractions. Peptides are denoted by the SCX fraction in which they were identified, their sequence, and their charge state. Ratios at t=0 should all be 0.

Practically, the natural solution of "normalizing" RIAs by imposing that the RIA at $t = 0$ is equal to 0 does not work. First of all, for some peptides or proteins the earliest available RIAs start later and, e.g., to impose an RIA at 6 hours to be 0 would be certainly questionable. Moreover, outlier RIAs are frequent and in some cases occur with rather strong signals at $t = 0$, which would ruin the normalization. This later phenomenon is due to co-eluting material obviously.

In theory, it should be possible to estimate noise parameters and adjust data accordingly but in our hand this was not possible. We explain this difficulty by fact that LC-MS data were acquired in a large number (120) of samples, at different times, in different replicates and SCX fractions and despite LC reproducibility the environment, i.e. co-eluting molecules, of a peptide is different each time.

Given the rather simple nature of the problem, occasional small vertical shifts, and the excellent agreement of the 2-component model with experimental data, we reasoned that the mathematical model should be augmented with a shift parameter, the shift magnitude being constrained by the phenomenological 2-compartment model. RIA at time $t = 0$ were not considered in computing the error between experimental RIAs and the model for the reasons we just discussed, they were regarded as *de facto* outliers. In practice, the shift values remained very small and they are reported for each peptide or protein dynamical model in our **Supplementary Data**.

### Final peptide validation

Taking advantage of the mathematical model above, we fit its parameters to each peptide, i.e. observation, which passed the initial filtering described above. We imposed the following criteria to finally validate a peptide:

a) $\alpha(t)$ and $\beta(t)$ increasing at $t = 3.02$ ;

b) $\alpha(t)$ decreasing at $t = 10.17$ ;
c) $\beta(t)$ maximum after 9 hours ;
d) No over-dispersion of the data: taking $\delta = 0.35 \max_t \beta(t)$, RIAs at distance larger than $\delta$ from the curve $\beta(t)$ were considered outliers and peptides with more than 25% outlier RIAs considered over-dispersed ;
e) $\lambda < 1$ ;
f) $\max_t \beta(t) < 0.2$ ;
g) Linear correlation between experimental data (RIAs) and $\beta(t)$ > 0.75.

These parameters are either obvious (a,b,c,e,f) or they have been adjusted empirically, in particular to control the FDR.

**Robust protein model construction**

Proteins detected with a single observation were treated as above.

The first step in assembling multiple observations to construct a protein turnover mathematical model was to "align" available observations. As a matter of fact, we know that each observation might include a small arbitrary vertical displacement. Following the same logic as for other steps of our algorithm, we considered that the higher the heavy peptide signal the more reliable the observation. We thus first identified the most intense observation after its heavy MS intensity. One observation intensity was determined by taking the third quartile of the heavy MS signal between times 9h and 19h considering all available replicates. Using the most intense observation as an anchor, we simply applied vertical shifts to all the other observations by minimizing weighted squared median distances between the anchor and the observation to align, the weight being proportional to the square root of the median of the heavy MS intensities of the observation to align.

Second, a first mathematical model was constructed as above with all the – pooled – observations RIAs.

Third, we determined outliers as above but with a slightly larger $\delta = 0.5 \max_t \beta(t)$ to acknowledge additional variability due to multiple observations. Observations comprised of >25% outliers were discarded from the protein model. They might be nonetheless correct as isoform, chain or active peptide with a different kinetics, meaning that they were maintained as correct as independent observation.

Fourth, the mathematical model was recomputed with all the kept observations.

**Bootstrap**

In order to obtain an estimate of the variability in our observations, we estimated the standard deviation of all the model parameters. We assumed a normal distribution of the differences between the theoretical curve $\beta(t)$ and experimental RIAs. For each protein, we estimated the parameters of this normal distributions and generated 1,000 synthetic data sets. We fit the parameters of 1,000 mathematical models on these data and thus obtained 1,000 estimates of each parameter whose square root of mean squared differences with the original estimates was computed. This is a classical bootstrap procedure(Davison and Hinkley, 1997). We also computed the median absolute deviation (MAD) that is reported in the supplementary tables.

**Fitting MRM data**

These data could be modeled by Eq. (1) unchanged and we applied the same procedure regarding the morphological filters as for HRMS data. Since, by their nature, MRM data contain little noise, we did not use the vertical shift explained above.

**Use of GTEx data**

In order to determine gene tissue specific expression, we made use of the GTEx data(GTEx Consortium, 2013). Namely, we used tissue gene median expression data in TPM (2026-01-15_v7). Tissues were grouped as follows. Liver was only represented by GTEx liver, CNS was represented by GTEx:

- Brain – Amygdala
- Brain - Anterior cingulate cortex (BA24)
- Brain - Caudate (basal ganglia)
- Brain - Cerebellar Hemisphere
- Brain – Cerebellum
- Brain – Cortex
- Brain - Frontal Cortex (BA9)
- Brain – Hippocampus
- Brain - Hypothalamus
- Brain - Nucleus accumbens (basal ganglia)
- Brain - Putamen (basal ganglia)
- Brain - Spinal cord (cervical c-1)
- Brain - Substantia nigra
- Pituitary

Others was comprised of all the remaining GTEx tissues.

**Fitting human plasma data**

The turnover data released by Price et al. for plasma proteins(Price et al., 2012) were reanalyzed with our mathematical model and robust protein model algorithm to extract the clearance rate $k_c$ in a similar fashion (see our discussion above regarding mouse data). In order to obtain a representative rate for each detected protein, we used the 3 patients (4, 7, and 8) that were analyzed on Price et al. 7-day continuous labeling protocol (no pulse-chase like in SILK). For each protein detected in at least 2 patients we calculated the median clearance rate. Clearance rates were in 1/day units for the plasma data set, whereas our CSF rates were in 1/hour units, which required to divide the plasma rates by 24 to obtain values on the same scale. An example of plasma protein model fitting is featured in **Fig. S10**.

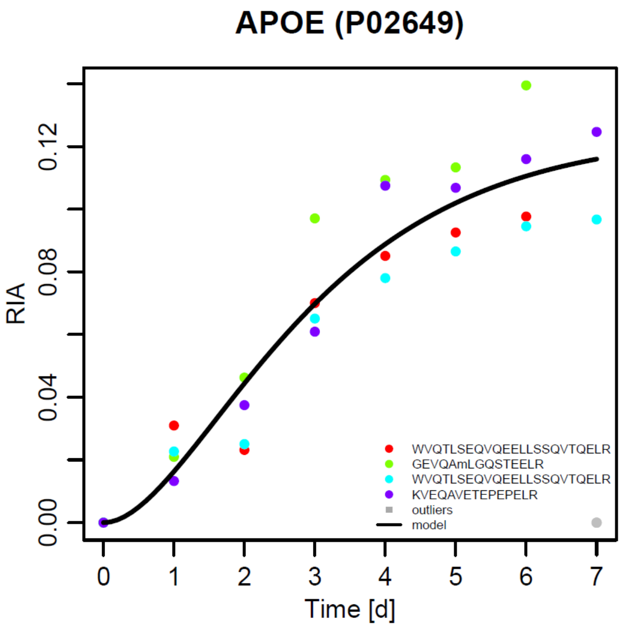

**Figure S10**. Plasma APOE dynamics in Price et al. patient 4 data(Price et al., 2012).

### Supplemental Information References

Bateman RJ, Munsell LY, Morris JC, Swarm R, Yarasheski KE, Holtzman DM. 2006. Human amyloid-beta synthesis and clearance rates as measured in cerebrospinal fluid in vivo. *Nat Med* **12**:856–861. doi:10.1038/nm1438

Davison AC, Hinkley DV. 1997. Bootstrap Methods and their Applications, Cambridge Series in Statistical and Probabilistic Mathematics. Cambridge University Press.

GTEx Consortium. 2013. The Genotype-Tissue Expression (GTEx) project. *Nat Genet* **45**:580–585. doi:10.1038/ng.2653

Guan S, Price JC, Ghaemmaghami S, Prusiner SB, Burlingame AL. 2012. Compartment modeling for mammalian protein turnover studies by stable isotope metabolic labeling. *Anal Chem* **84**:4014–4021. doi:10.1021/ac203330z

Hairer E, Wanner G. 1996. Solving Ordinary Differential Equations II: Stiff and Differential-Algebraic Problems, 2nd ed, Springer Series in Computational Mathematics, Springer Ser.Comp.Mathem. Hairer,E.:Solving Ordinary Diff. Berlin Heidelberg: Springer-Verlag.

Price JC, Guan S, Burlingame A, Prusiner SB, Ghaemmaghami S. 2010. Analysis of proteome dynamics in the mouse brain. *Proc Natl Acad Sci USA* **107**:14508–14513. doi:10.1073/pnas.1006551107

Price JC, Holmes WE, Li KW, Floreani NA, Neese RA, Turner SM, Hellerstein MK. 2012. Measurement of human plasma proteome dynamics with (2)H(2)O and liquid chromatography tandem mass spectrometry. *Anal Biochem* **420**:73–83. doi:10.1016/j.ab.2011.09.007

Rahman M, Previs SF, Kasumov T, Sadygov RG. 2016. Gaussian Process Modeling of Protein Turnover. *J Proteome Res* **15**:2115–2122. doi:10.1021/acs.jproteome.5b00990

Waterval WAH, Scheijen JLJM, Ortmans-Ploemen MMJC, Habets-van der Poel CD, Bierau J. 2009. Quantitative UPLC-MS/MS analysis of underivatised amino acids in body fluids is a reliable tool for the diagnosis and follow-up of patients with inborn errors of metabolism. *Clin Chim Acta* **407**:36–42. doi:10.1016/j.cca.2009.06.023
