## Supplementary material for "*In Vivo* Large Scale Mapping Of Protein Turnover In The Human Cerebrospinal Fluid": MRM parameters

| Column | Description |
|---|---|
| Q1 | precursor m/z setting for quadrupole 1 |
| Q3 | fragment ion m/z setting for quadrupole 3 |
| dwelltime | measurement time spent on this transition (ms) |
| Tr_recalibrated | retention time for scheduled SRM measurement (minutes) |
| transition_group_id | a unique ID for the transitions of one precursor, any format |
| CE | the collision energy used for this precursor |
| protein_name | an identifier for the targeted protein, for example a Uniprot identifier or gene name |
| relative_intensity | the expected relative intensity of this fragment ion compared to the other fragment ions in this transition group |
| transition_name | a name for the transition, any format |
| decoy | true for decoy transitions, false for target transitions (if decoys were measured) |
| sequence | the peptide sequence, including any modifications |
| isotype | a string for the labeling used (heavy or light) |
| prec_z | the precursor charge state |
| frg_type | the fragment ion type (use 'p' for precursor) |
| frg_nr | the fragment number (Excel MID() function useful to split out) (use peptide length for precursor) |
| frg_z | the fragment charge state |
| frg_loss | the fragment neutral loss (expressed as a negative number) |
| DP | the declustering potential |
