## Supplementary material for "*In Vivo* Large Scale Mapping Of Protein Turnover In The Human Cerebrospinal Fluid": HRMS protein table

| # units | | | | 1/h | 1/h | | | | | h | h | | | d | d | 1/h | | | h | | d | TPM | TPM | TPM |
|---|---|---|---|---|---|---|---|---|---|---|---|---|---|---|---|---|---|---|---|---|---|---|---|---|
| **protein** | **gene** | **obs.in.model** | **pept.in.model** | **k.c** | **k.c.MAD** | **lambda** | **lambda.MAD** | **shift** | **shift.MAD** | **apex** | **apex.MAD** | **r.max** | **r.max.MAD** | **half.life** | **half.life.MAD** | **k.c.sd** | **lambda.sd** | **shift.sd** | **apex.sd** | **r.max.sd** | **half.life.sd** | **GTEx.brain** | **GTEx.liver** | **GTEx.rest** |
| NX P04217-1 | A1BG | 4 | 4 | 0.0077 | 0.0003 | 0.7048 | 0.0244 | 0.0011 | 0.0011 | 24.47 | 0.00 | 0.0065 | 0.0004 | 3.73 | 0.16 | 0.0006 | 0.0751 | 0.0012 | 0.0000 | 0.0005 | 0.2861 | 5.19 | 450.20 | 3.86 |
| NX P01023-1 | A2M | 25 | 15 | 0.0835 | 0.0016 | 0.0526 | 0.0012 | 0.0026 | 0.0027 | 17.00 | 0.20 | 0.0142 | 0.0003 | 0.35 | 0.01 | 0.0071 | 0.0196 | 0.0028 | 0.6230 | 0.0007 | 0.2213 | 74.55 | 356.10 | 463.52 |
| NX Q562R1-1 | ACTBL2 | 1 | 1 | 0.0066 | 0.0003 | 0.2164 | 0.0119 | -0.0003 | 0.0003 | 24.47 | 0.00 | 0.0015 | 0.0001 | 4.36 | 0.23 | 0.0006 | 0.0220 | 0.0003 | 0.0000 | 0.0002 | 0.4292 | 0.00 | 0.00 | 0.02 |
| NX Q9UKQ2-1 | ADAM28 | 1 | 1 | 0.0655 | 0.0131 | 0.2293 | 0.0654 | 0.0227 | 0.0272 | 20.20 | 3.00 | 0.0491 | 0.0126 | 0.44 | 0.09 | 0.0305 | 0.1774 | 0.0328 | 3.1150 | 0.0191 | 0.9257 | 2.33 | 0.22 | 3.38 |
| NX Q13443-1 | ADAM9 | 1 | 1 | 0.2055 | 0.0087 | 0.0753 | 0.0059 | -0.0118 | 0.0112 | 10.60 | 0.20 | 0.0449 | 0.0025 | 0.14 | 0.01 | 0.0143 | 0.0088 | 0.0117 | 0.2639 | 0.0038 | 0.0095 | 12.25 | 3.59 | 35.67 |
| NX O75173-1 | ADAMTS4 | 1 | 1 | 0.0662 | 0.0096 | 0.0931 | 0.0133 | -0.0001 | 0.0017 | 20.00 | 2.00 | 0.0201 | 0.0017 | 0.44 | 0.06 | 0.0142 | 0.0391 | 0.0026 | 2.6367 | 0.0028 | 0.1462 | 5.76 | 1.50 | 12.70 |
| NX P01019-1 | AGT | 7 | 3 | 0.0532 | 0.0018 | 0.1268 | 0.0047 | 0.0031 | 0.0035 | 23.60 | 0.60 | 0.0221 | 0.0003 | 0.54 | 0.02 | 0.0027 | 0.0071 | 0.0035 | 0.7484 | 0.0005 | 0.0279 | 172.20 | 1055.00 | 19.92 |
| NX P02765-1 | AHSG | 2 | 2 | 0.0084 | 0.0006 | 0.6923 | 0.0896 | -0.0015 | 0.0011 | 24.47 | 0.00 | 0.0073 | 0.0007 | 3.46 | 0.29 | 0.0009 | 0.1221 | 0.0012 | 0.0000 | 0.0010 | 0.4575 | 0.11 | 1377.00 | 0.16 |
| NX P02768-1 | ALB | 18 | 14 | 0.0192 | 0.0005 | 0.0353 | 0.0015 | 0.0000 | 0.0000 | 24.47 | 0.00 | 0.0016 | 0.0000 | 1.51 | 0.04 | 0.0014 | 0.0061 | 0.0000 | 0.0000 | 0.0001 | 0.1362 | 1.66 | 25520.00 | 11.91 |
| NX P02760-1 | AMBP | 4 | 3 | 0.0809 | 0.0048 | 0.0778 | 0.0054 | 0.0045 | 0.0046 | 17.40 | 0.73 | 0.0204 | 0.0012 | 0.36 | 0.02 | 0.0074 | 0.0085 | 0.0049 | 1.1854 | 0.0018 | 0.0358 | 0.18 | 3299.00 | 1.97 |
| NX Q06481-1 | APLP2 | 1 | 1 | 0.0330 | 0.0190 | 0.7025 | 0.5947 | 0.0364 | 0.0655 | 24.47 | 0.00 | 0.0705 | 0.0561 | 0.88 | 0.43 | 0.1160 | 1.0615 | 0.1525 | 5.5154 | 0.1861 | 359.6621 | 201.86 | 119.10 | 244.65 |
| NX P02647-1 | APOA1 | 11 | 8 | 0.0299 | 0.0029 | 0.0659 | 0.0082 | 0.0009 | 0.0010 | 24.47 | 0.00 | 0.0058 | 0.0002 | 0.97 | 0.09 | 0.0060 | 0.0246 | 0.0011 | 0.0085 | 0.0003 | 0.2225 | 0.82 | 5751.00 | 17.86 |
| NX P02652-1 | APOA2 | 4 | 3 | 0.0175 | 0.0009 | 0.2219 | 0.0147 | 0.0026 | 0.0028 | 24.47 | 0.00 | 0.0086 | 0.0008 | 1.65 | 0.09 | 0.0040 | 0.0517 | 0.0029 | 0.0814 | 0.0011 | 0.2320 | 0.24 | 4629.00 | 0.30 |
| NX P06727-1 | APOA4 | 8 | 6 | 0.0184 | 0.0008 | 0.6478 | 0.0349 | 0.0000 | 0.0009 | 24.47 | 0.00 | 0.0271 | 0.0011 | 1.57 | 0.07 | 0.0021 | 0.1216 | 0.0014 | 0.0000 | 0.0016 | 0.1737 | 0.01 | 60.27 | 20.99 |
| NX P04114-1 | APOB | 3 | 3 | 0.1092 | 0.0068 | 0.1571 | 0.0138 | 0.0328 | 0.0322 | 14.40 | 0.60 | 0.0546 | 0.0048 | 0.26 | 0.02 | 0.0094 | 0.0214 | 0.0327 | 0.7630 | 0.0069 | 0.0238 | 0.05 | 348.40 | 2.75 |
| NX P05090-1 | APOD | 10 | 4 | 0.0873 | 0.0019 | 0.0882 | 0.0027 | 0.0023 | 0.0019 | 16.60 | 0.20 | 0.0248 | 0.0007 | 0.33 | 0.01 | 0.0029 | 0.0038 | 0.0020 | 0.3787 | 0.0010 | 0.0113 | 299.57 | 2.28 | 926.27 |
| NX P02649-1 | APOE | 21 | 11 | 0.0473 | 0.0017 | 0.1238 | 0.0055 | -0.0018 | 0.0024 | 24.47 | 0.00 | 0.0192 | 0.0003 | 0.61 | 0.02 | 0.0027 | 0.0081 | 0.0024 | 0.1757 | 0.0004 | 0.0350 | 921.00 | 3259.00 | 261.74 |
| NX P02749-1 | APOH | 6 | 2 | 0.0199 | 0.0014 | 0.3472 | 0.0361 | 0.0023 | 0.0026 | 24.47 | 0.00 | 0.0166 | 0.0005 | 1.45 | 0.10 | 0.0033 | 0.0954 | 0.0027 | 0.0000 | 0.0008 | 0.2311 | 0.06 | 3046.00 | 0.46 |
| NX P05067-1 | APP | 1 | 1 | 0.0350 | 0.0134 | 0.3187 | 0.1382 | -0.0092 | 0.0124 | 24.47 | 0.00 | 0.0346 | 0.0043 | 0.83 | 0.29 | 0.0173 | 0.4617 | 0.0134 | 1.6920 | 0.0060 | 0.6113 | 335.21 | 67.94 | 243.34 |
| NX Q13508-1 | ART3 | 1 | 1 | 0.0118 | 0.0005 | 0.6360 | 0.0209 | 0.0008 | 0.0010 | 24.47 | 0.00 | 0.0126 | 0.0007 | 2.44 | 0.10 | 0.0008 | 0.0550 | 0.0011 | 0.0000 | 0.0010 | 0.1731 | 4.91 | 0.07 | 7.82 |
| NX Q15904-1 | ATP6AP1 | 1 | 1 | 0.0733 | 0.0029 | 0.1830 | 0.0086 | 0.0135 | 0.0140 | 18.60 | 0.60 | 0.0436 | 0.0026 | 0.39 | 0.02 | 0.0157 | 0.0381 | 0.0158 | 1.2305 | 0.0056 | 0.3653 | 77.09 | 19.96 | 62.28 |
| NX O75787-1 | ATP6AP2 | 1 | 1 | 0.0921 | 0.0048 | 0.0199 | 0.0021 | 0.0016 | 0.0017 | 16.00 | 0.60 | 0.0059 | 0.0007 | 0.31 | 0.02 | 0.0079 | 0.0032 | 0.0019 | 0.9521 | 0.0010 | 0.0308 | 89.79 | 33.37 | 99.30 |
| NX P61769-1 | B2M | 2 | 2 | 0.0705 | 0.0045 | 0.1065 | 0.0063 | 0.0003 | 0.0009 | 19.20 | 0.80 | 0.0244 | 0.0012 | 0.41 | 0.03 | 0.0068 | 0.0103 | 0.0014 | 1.4642 | 0.0017 | 0.0441 | 267.19 | 579.10 | 1202.34 |
| NX O43505-1 | B4GAT1 | 1 | 1 | 0.0173 | 0.0020 | 0.6275 | 0.1015 | 0.0056 | 0.0066 | 24.47 | 0.00 | 0.0237 | 0.0017 | 1.67 | 0.21 | 0.0025 | 0.1380 | 0.0069 | 0.0000 | 0.0025 | 0.2663 | NA | NA | NA |
| NX P02745-1 | C1QA | 1 | 1 | 0.0178 | 0.0034 | 0.6716 | 0.1934 | 0.0076 | 0.0059 | 24.47 | 0.00 | 0.0267 | 0.0021 | 1.62 | 0.35 | 0.0086 | 0.3341 | 0.0062 | 0.2192 | 0.0032 | 0.4994 | 23.38 | 19.33 | 117.37 |
| NX P02746-1 | C1QB | 1 | 1 | 0.0962 | 0.0015 | 0.1066 | 0.0026 | 0.0032 | 0.0030 | 15.60 | 0.20 | 0.0329 | 0.0007 | 0.30 | 0.00 | 0.0022 | 0.0035 | 0.0032 | 0.2419 | 0.0011 | 0.0069 | 30.60 | 18.81 | 130.00 |
| NX P02747-1 | C1QC | 4 | 3 | 0.0928 | 0.0022 | 0.1008 | 0.0030 | 0.0025 | 0.0022 | 15.80 | 0.20 | 0.0301 | 0.0008 | 0.31 | 0.01 | 0.0032 | 0.0044 | 0.0024 | 0.3726 | 0.0013 | 0.0108 | 27.10 | 15.51 | 111.00 |
| NX P00736-1 | C1R | 3 | 1 | 0.1023 | 0.0039 | 0.0923 | 0.0049 | -0.0038 | 0.0048 | 15.00 | 0.40 | 0.0302 | 0.0015 | 0.28 | 0.01 | 0.0061 | 0.0072 | 0.0050 | 0.5650 | 0.0022 | 0.0174 | 12.49 | 922.60 | 362.86 |
| NX P09871-1 | C1S | 3 | 2 | 0.1007 | 0.0039 | 0.1098 | 0.0066 | 0.0039 | 0.0031 | 15.20 | 0.40 | 0.0354 | 0.0019 | 0.29 | 0.01 | 0.0059 | 0.0098 | 0.0038 | 0.5575 | 0.0029 | 0.0170 | 13.07 | 664.70 | 315.78 |
| NX P01024-1 | C3 | 25 | 18 | 0.0856 | 0.0022 | 0.0417 | 0.0014 | 0.0015 | 0.0014 | 16.80 | 0.20 | 0.0115 | 0.0003 | 0.34 | 0.01 | 0.0032 | 0.0020 | 0.0015 | 0.4245 | 0.0005 | 0.0128 | 23.44 | 1660.00 | 198.44 |
| NX P0C0L4-1 | C4A | 23 | 16 | 0.0710 | 0.0017 | 0.1012 | 0.0025 | 0.0041 | 0.0039 | 19.00 | 0.40 | 0.0234 | 0.0005 | 0.41 | 0.01 | 0.0024 | 0.0037 | 0.0040 | 0.4828 | 0.0007 | 0.0141 | 2.11 | 49.85 | 6.11 |
| NX Q5BLP8-1 | C4orf48 | 1 | 1 | 0.1514 | 0.0278 | 0.1180 | 0.0305 | 0.0213 | 0.0199 | 12.00 | 1.00 | 0.0549 | 0.0139 | 0.19 | 0.03 | 0.0494 | 0.1748 | 0.0259 | 4.0192 | 0.0206 | 0.2797 | 40.77 | 1.13 | 8.12 |
| NX P01031-1 | C5 | 2 | 2 | 0.0414 | 0.0106 | 0.0388 | 0.0102 | 0.0000 | 0.0006 | 24.47 | 0.00 | 0.0052 | 0.0006 | 0.70 | 0.16 | 0.0147 | 0.0338 | 0.0009 | 1.7471 | 0.0009 | 0.3848 | 2.52 | 126.40 | 4.72 |
| NX P13671-1 | C6 | 2 | 2 | 0.0130 | 0.0015 | 0.5865 | 0.0422 | 0.0003 | 0.0019 | 24.47 | 0.00 | 0.0137 | 0.0026 | 2.22 | 0.27 | 0.0028 | 0.0765 | 0.0030 | 0.0000 | 0.0039 | 1.1614 | 0.12 | 137.30 | 4.22 |
| NX P10643-1 | C7 | 6 | 3 | 0.0977 | 0.0036 | 0.0851 | 0.0045 | 0.0046 | 0.0059 | 15.40 | 0.40 | 0.0267 | 0.0012 | 0.30 | 0.01 | 0.0056 | 0.0067 | 0.0061 | 0.5720 | 0.0019 | 0.0174 | 3.49 | 49.61 | 226.55 |
| NX P07357-1 | C8A | 1 | 1 | 0.0063 | 0.0003 | 0.6797 | 0.0050 | 0.0002 | 0.0003 | 24.47 | 0.00 | 0.0042 | 0.0003 | 4.60 | 0.20 | 0.0004 | 0.0453 | 0.0004 | 0.0000 | 0.0004 | 0.3628 | 0.00 | 284.10 | 0.01 |
| NX P02748-1 | C9 | 3 | 2 | 0.1179 | 0.0083 | 0.0620 | 0.0077 | -0.0012 | 0.0021 | 13.80 | 0.60 | 0.0231 | 0.0024 | 0.25 | 0.02 | 0.0125 | 0.0115 | 0.0032 | 0.8557 | 0.0037 | 0.0270 | 0.02 | 412.50 | 0.10 |
| NX P00918-1 | CA2 | 1 | 1 | 0.0316 | 0.0221 | 0.7891 | 0.6047 | 0.0356 | 0.1141 | 24.47 | 0.00 | 0.0747 | 0.1192 | 0.92 | 0.44 | 0.0858 | 1.4523 | 0.1506 | 6.5604 | 0.2044 | 2130.5167 | 58.38 | 64.15 | 33.80 |
| NX P08571-1 | CD14 | 9 | 5 | 0.1101 | 0.0025 | 0.0495 | 0.0018 | 0.0019 | 0.0021 | 14.40 | 0.20 | 0.0173 | 0.0006 | 0.26 | 0.01 | 0.0038 | 0.0027 | 0.0022 | 0.3031 | 0.0009 | 0.0090 | 16.52 | 204.40 | 56.13 |
| NX O94986-1 | CEP152 | 1 | 1 | 0.0179 | 0.0018 | 0.7282 | 0.0963 | -0.0005 | 0.0045 | 24.47 | 0.00 | 0.0292 | 0.0049 | 1.61 | 0.17 | 0.0125 | 0.2917 | 0.0090 | 0.8938 | 0.0083 | 0.5370 | 0.83 | 0.56 | 3.22 |
| NX O15078-1 | CEP290 | 1 | 1 | 0.1519 | 0.0051 | 0.0445 | 0.0025 | -0.0074 | 0.0075 | 12.00 | 0.20 | 0.0207 | 0.0010 | 0.19 | 0.01 | 0.0077 | 0.0038 | 0.0076 | 0.3000 | 0.0015 | 0.0096 | 5.34 | 1.57 | 6.57 |
| NX P00751-1 | CFB | 7 | 7 | 0.0843 | 0.0030 | 0.0539 | 0.0026 | -0.0015 | 0.0020 | 17.00 | 0.40 | 0.0147 | 0.0006 | 0.34 | 0.01 | 0.0045 | 0.0036 | 0.0021 | 0.6182 | 0.0009 | 0.0186 | 4.77 | 1146.00 | 25.15 |
| NX P00746-1 | CFD | 1 | 1 | 0.1194 | 0.0074 | 0.0718 | 0.0078 | -0.0020 | 0.0023 | 13.60 | 0.40 | 0.0271 | 0.0024 | 0.24 | 0.01 | 0.0113 | 0.0111 | 0.0034 | 0.7406 | 0.0037 | 0.0233 | 3.68 | 3.67 | 450.51 |
| NX P08603-1 | CFH | 8 | 7 | 0.1205 | 0.0022 | 0.0568 | 0.0017 | 0.0018 | 0.0022 | 13.60 | 0.20 | 0.0216 | 0.0006 | 0.24 | 0.00 | 0.0033 | 0.0026 | 0.0023 | 0.2153 | 0.0009 | 0.0066 | 6.79 | 335.60 | 76.31 |
| NX P05156-1 | CFI | 3 | 2 | 0.1108 | 0.0047 | 0.0379 | 0.0026 | -0.0012 | 0.0017 | 14.20 | 0.40 | 0.0134 | 0.0008 | 0.26 | 0.01 | 0.0072 | 0.0039 | 0.0019 | 0.5595 | 0.0012 | 0.0173 | 3.71 | 259.30 | 25.26 |
| NX P10645-1 | CHGA | 5 | 4 | 0.0109 | 0.0012 | 0.7006 | 0.1354 | 0.0022 | 0.0023 | 24.47 | 0.00 | 0.0119 | 0.0006 | 2.66 | 0.32 | 0.0014 | 0.1971 | 0.0024 | 0.0000 | 0.0008 | 0.4299 | 121.63 | 0.14 | 15.46 |
| NX P05060-1 | CHGB | 6 | 6 | 0.0248 | 0.0012 | 0.2659 | 0.0155 | 0.0043 | 0.0045 | 24.47 | 0.00 | 0.0177 | 0.0005 | 1.17 | 0.06 | 0.0044 | 0.0772 | 0.0046 | 0.0000 | 0.0008 | 0.2070 | 242.76 | 0.12 | 4.61 |
| NX P36222-1 | CHI3L1 | 5 | 3 | 0.1137 | 0.0035 | 0.0772 | 0.0041 | -0.0001 | 0.0012 | 14.00 | 0.20 | 0.0278 | 0.0014 | 0.25 | 0.01 | 0.0055 | 0.0061 | 0.0018 | 0.4032 | 0.0020 | 0.0124 | 36.49 | 115.90 | 23.34 |
| NX O00533-1 | CHL1 | 1 | 1 | 0.0120 | 0.0008 | 0.2073 | 0.0106 | 0.0008 | 0.0008 | 24.47 | 0.00 | 0.0042 | 0.0004 | 2.41 | 0.16 | 0.0013 | 0.0199 | 0.0009 | 0.0000 | 0.0006 | 0.3054 | 14.91 | 0.15 | 5.83 |
| NX P05452-1 | CLEC3B | 7 | 4 | 0.1053 | 0.0033 | 0.0617 | 0.0033 | 0.0021 | 0.0027 | 14.60 | 0.20 | 0.0207 | 0.0011 | 0.27 | 0.01 | 0.0049 | 0.0050 | 0.0030 | 0.4311 | 0.0016 | 0.0129 | 19.80 | 9.98 | 79.99 |
| NX O94985-1 | CLSTN1 | 2 | 2 | 0.0989 | 0.0073 | 0.0758 | 0.0072 | -0.0032 | 0.0044 | 15.22 | 0.62 | 0.0240 | 0.0020 | 0.29 | 0.02 | 0.0107 | 0.0103 | 0.0050 | 1.1436 | 0.0030 | 0.0347 | 172.98 | 5.72 | 81.58 |
| NX P10909-1 | CLU | 13 | 8 | 0.1418 | 0.0023 | 0.0910 | 0.0027 | -0.0013 | 0.0017 | 12.40 | 0.20 | 0.0400 | 0.0009 | 0.20 | 0.00 | 0.0033 | 0.0039 | 0.0020 | 0.1770 | 0.0014 | 0.0048 | 951.05 | 732.90 | 409.08 |
| NX Q96KN2-1 | CNDP1 | 4 | 2 | 0.0131 | 0.0007 | 0.8290 | 0.0783 | 0.0025 | 0.0026 | 24.47 | 0.00 | 0.0195 | 0.0008 | 2.21 | 0.13 | 0.0010 | 0.1071 | 0.0028 | 0.0000 | 0.0012 | 0.1894 | 29.06 | 10.17 | 0.12 |
| NX Q12860-1 | CNTN1 | 1 | 1 | 0.1674 | 0.0099 | 0.0276 | 0.0031 | -0.0038 | 0.0043 | 11.60 | 0.20 | 0.0140 | 0.0013 | 0.17 | 0.01 | 0.0149 | 0.0046 | 0.0045 | 0.4494 | 0.0020 | 0.0155 | 47.41 | 0.11 | 5.94 |
| NX P39060-1 | COL18A1 | 1 | 1 | 0.0714 | 0.0092 | 0.0197 | 0.0038 | 0.0015 | 0.0017 | 19.00 | 1.60 | 0.0046 | 0.0009 | 0.40 | 0.05 | 0.0171 | 0.0124 | 0.0020 | 2.7885 | 0.0014 | 0.2924 | 24.33 | 206.40 | 142.71 |
| NX P02452-1 | COL1A1 | 5 | 5 | 0.1423 | 0.0047 | 0.0880 | 0.0052 | -0.0021 | 0.0016 | 12.40 | 0.20 | 0.0388 | 0.0020 | 0.20 | 0.01 | 0.0068 | 0.0077 | 0.0024 | 0.3006 | 0.0029 | 0.0097 | 12.03 | 14.63 | 356.17 |
| NX P08123-1 | COL1A2 | 1 | 1 | 0.1313 | 0.0037 | 0.0833 | 0.0040 | 0.0008 | 0.0012 | 13.00 | 0.20 | 0.0342 | 0.0014 | 0.22 | 0.01 | 0.0055 | 0.0059 | 0.0018 | 0.2884 | 0.0022 | 0.0092 | 8.24 | 11.95 | 441.99 |
| NX P02461-1 | COL3A1 | 1 | 1 | 0.1109 | 0.0053 | 0.0910 | 0.0066 | 0.0027 | 0.0026 | 14.20 | 0.40 | 0.0321 | 0.0022 | 0.26 | 0.01 | 0.0080 | 0.0101 | 0.0035 | 0.6243 | 0.0033 | 0.0193 | 2.57 | 26.92 | 322.30 |
| NX P12109-1 | COL6A1 | 2 | 2 | 0.1016 | 0.0061 | 0.0755 | 0.0073 | 0.0050 | 0.0046 | 15.00 | 0.60 | 0.0245 | 0.0024 | 0.28 | 0.02 | 0.0105 | 0.0148 | 0.0054 | 0.9823 | 0.0035 | 0.1268 | 101.90 | 59.32 | 682.52 |
| NX P12111-1 | COL6A3 | 2 | 2 | 0.1205 | 0.0048 | 0.0511 | 0.0034 | -0.0013 | 0.0017 | 13.60 | 0.40 | 0.0194 | 0.0011 | 0.24 | 0.01 | 0.0072 | 0.0047 | 0.0021 | 0.4578 | 0.0016 | 0.0145 | 0.61 | 3.46 | 108.98 |
| NX P16870-1 | CPE | 1 | 1 | 0.2501 | 0.0117 | 0.1016 | 0.0073 | -0.0168 | 0.0185 | 10.00 | 0.17 | 0.0697 | 0.0040 | 0.12 | 0.01 | 0.0168 | 0.0104 | 0.0190 | 0.1972 | 0.0060 | 0.0077 | 608.59 | 5.64 | 122.82 |
| NX P22792-1 | CPN2 | 1 | 1 | 0.0071 | 0.0005 | 0.5658 | 0.0183 | 0.0007 | 0.0012 | 24.47 | 0.00 | 0.0044 | 0.0004 | 4.07 | 0.34 | 0.0006 | 0.0421 | 0.0012 | 0.0000 | 0.0006 | 0.4218 | 0.00 | 129.30 | 0.32 |
| NX Q9Y646-1 | CPQ | 2 | 1 | 0.1248 | 0.0052 | 0.1041 | 0.0126 | 0.0125 | 0.0117 | 13.40 | 0.40 | 0.0409 | 0.0046 | 0.23 | 0.01 | 0.0084 | 0.0197 | 0.0130 | 0.4815 | 0.0071 | 0.0153 | 12.01 | 11.51 | 38.06 |
| NX Q9H3G5-1 | CPVL | 1 | 1 | 0.1983 | 0.0118 | 0.1538 | 0.0205 | 0.0133 | 0.0110 | 10.80 | 0.20 | 0.0893 | 0.0092 | 0.15 | 0.01 | 0.0192 | 0.0293 | 0.0151 | 0.3607 | 0.0135 | 0.0135 | 5.91 | 0.84 | 14.68 |
| NX P00450-1 | CP | 7 | 4 | 0.0921 | 0.0034 | 0.0242 | 0.0011 | -0.0015 | 0.0015 | 16.00 | 0.40 | 0.0072 | 0.0003 | 0.31 | 0.01 | 0.0051 | 0.0016 | 0.0015 | 0.5739 | 0.0004 | 0.0175 | 4.20 | 315.80 | 9.15 |
| NX P07333-1 | CSF1R | 1 | 1 | 0.0542 | 0.0023 | 0.0957 | 0.0046 | 0.0001 | 0.0003 | 23.40 | 0.80 | 0.0170 | 0.0004 | 0.53 | 0.02 | 0.0033 | 0.0067 | 0.0005 | 0.9634 | 0.0005 | 0.0328 | 12.66 | 2.73 | 22.90 |
| NX P01034-1 | CST3 | 5 | 2 | 0.0695 | 0.0041 | 0.1224 | 0.0079 | -0.0027 | 0.0039 | 19.40 | 0.80 | 0.0277 | 0.0016 | 0.42 | 0.02 | 0.0062 | 0.0121 | 0.0044 | 1.3114 | 0.0023 | 0.0397 | 630.95 | 159.60 | 380.53 |
| NX P07339-1 | CTSD | 3 | 2 | 0.1899 | 0.0029 | 0.1460 | 0.0037 | 0.0058 | 0.0044 | 11.00 | 0.00 | 0.0820 | 0.0017 | 0.15 | 0.00 | 0.0043 | 0.0055 | 0.0048 | 0.1098 | 0.0026 | 0.0034 | 243.58 | 552.10 | 655.34 |
| NX P48061-1 | CXCL12 | 1 | 1 | 0.1390 | 0.0029 | 0.0976 | 0.0039 | 0.0042 | 0.0041 | 12.60 | 0.20 | 0.0422 | 0.0014 | 0.21 | 0.00 | 0.0044 | 0.0059 | 0.0044 | 0.2070 | 0.0022 | 0.0066 | 4.13 | 34.35 | 58.57 |
| NX Q14118-1 | DAG1 | 2 | 2 | 0.0575 | 0.0080 | 0.1629 | 0.0243 | 0.0019 | 0.0032 | 22.20 | 2.27 | 0.0307 | 0.0028 | 0.50 | 0.07 | 0.0128 | 0.0689 | 0.0045 | 2.2960 | 0.0042 | 0.1718 | 18.60 | 19.70 | 43.21 |
| NX P07585-1 | DCN | 1 | 1 | 0.0484 | 0.0165 | 0.5061 | 0.0911 | 0.0233 | 0.0255 | 24.47 | 0.00 | 0.0804 | 0.0123 | 0.60 | 0.17 | 0.0197 | 0.5193 | 0.0288 | 2.9969 | 0.0209 | 0.4687 | 14.59 | 31.90 | 374.41 |
| NX Q9UBP4-1 | DKK3 | 7 | 4 | 0.0489 | 0.0040 | 0.1505 | 0.0135 | 0.0013 | 0.0011 | 24.47 | 0.00 | 0.0241 | 0.0008 | 0.59 | 0.05 | 0.0059 | 0.0253 | 0.0014 | 0.9210 | 0.0011 | 0.0802 | 216.03 | 2.93 | 107.61 |
| NX Q03001-14 | DST | 1 | 1 | 0.0502 | 0.0066 | 0.1134 | 0.0155 | 0.0003 | 0.0011 | 24.47 | 0.00 | 0.0187 | 0.0012 | 0.58 | 0.07 | 0.0099 | 0.0410 | 0.0017 | 1.8017 | 0.0018 | 0.1570 | 28.08 | 9.47 | 32.01 |
| NX Q16610-1 | ECM1 | 3 | 2 | 0.0214 | 0.0038 | 0.2068 | 0.0626 | -0.0032 | 0.0049 | 24.47 | 0.00 | 0.0111 | 0.0012 | 1.35 | 0.28 | 0.0121 | 0.1222 | 0.0052 | 0.5655 | 0.0018 | 0.5398 | 8.11 | 16.81 | 57.68 |
| NX O94769-1 | ECM2 | 2 | 1 | 0.1480 | 0.0035 | 0.1462 | 0.0066 | 0.0124 | 0.0125 | 12.20 | 0.20 | 0.0667 | 0.0025 | 0.20 | 0.00 | 0.0053 | 0.0096 | 0.0128 | 0.2187 | 0.0038 | 0.0070 | 2.14 | 9.27 | 18.37 |
| NX Q008S8-1 | ECT2L | 1 | 1 | 0.0247 | 0.0027 | 0.0731 | 0.0100 | 0.0008 | 0.0008 | 24.47 | 0.00 | 0.0048 | 0.0005 | 1.17 | 0.13 | 0.0091 | 0.0351 | 0.0009 | 0.3200 | 0.0007 | 0.3563 | 0.27 | 0.07 | 0.25 |
| NX Q12805-1 | EFEMP1 | 7 | 5 | 0.1200 | 0.0028 | 0.1035 | 0.0039 | 0.0070 | 0.0068 | 13.60 | 0.20 | 0.0392 | 0.0014 | 0.24 | 0.01 | 0.0040 | 0.0060 | 0.0071 | 0.2675 | 0.0020 | 0.0081 | 33.64 | 3.63 | 158.44 |
| NX Q13822-1 | ENPP2 | 14 | 9 | 0.2032 | 0.0036 | 0.1183 | 0.0041 | -0.0051 | 0.0045 | 10.80 | 0.00 | 0.0700 | 0.0019 | 0.14 | 0.00 | 0.0055 | 0.0057 | 0.0048 | 0.1339 | 0.0027 | 0.0039 | 55.33 | 2.77 | 36.55 |
| NX P00734-1 | F2 | 2 | 2 | 0.0358 | 0.0078 | 0.0527 | 0.0141 | 0.0003 | 0.0004 | 24.47 | 0.00 | 0.0059 | 0.0003 | 0.81 | 0.17 | 0.0109 | 0.0410 | 0.0006 | 0.5906 | 0.0005 | 0.3488 | 0.34 | 628.30 | 0.17 |
| NX P12259-1 | F5 | 3 | 2 | 0.1704 | 0.0065 | 0.1413 | 0.0090 | 0.0056 | 0.0031 | 11.40 | 0.20 | 0.0727 | 0.0040 | 0.17 | 0.01 | 0.0095 | 0.0134 | 0.0048 | 0.2846 | 0.0057 | 0.0093 | 0.45 | 34.63 | 1.51 |
| NX Q92520-1 | FAM3C | 1 | 1 | 0.0833 | 0.0075 | 0.4071 | 0.1115 | 0.0546 | 0.0598 | 17.00 | 1.00 | 0.1097 | 0.0361 | 0.35 | 0.03 | 0.0215 | 0.1792 | 0.0747 | 2.6990 | 0.0490 | 732.0127 | 29.98 | 15.30 | 33.40 |
| NX Q9NYQ8-1 | FAT2 | 1 | 1 | 0.1168 | 0.0031 | 0.0885 | 0.0032 | 0.0010 | 0.0010 | 13.80 | 0.20 | 0.0327 | 0.0011 | 0.25 | 0.01 | 0.0042 | 0.0049 | 0.0015 | 0.2919 | 0.0017 | 0.0089 | 30.26 | 0.04 | 4.05 |
| NX P23142-1 | FBLN1 | 8 | 6 | 0.1685 | 0.0044 | 0.0857 | 0.0042 | -0.0056 | 0.0056 | 11.60 | 0.20 | 0.0437 | 0.0018 | 0.17 | 0.00 | 0.0067 | 0.0065 | 0.0060 | 0.2127 | 0.0028 | 0.0068 | 10.85 | 2.83 | 251.09 |
| NX P23142-4 | FBLN1 | 10 | 7 | 0.1717 | 0.0039 | 0.0802 | 0.0031 | -0.0057 | 0.0059 | 11.40 | 0.20 | 0.0415 | 0.0013 | 0.17 | 0.00 | 0.0060 | 0.0049 | 0.0061 | 0.1792 | 0.0021 | 0.0059 | 10.85 | 2.83 | 251.09 |
| NX Q5XX13-1 | FBXW10 | 1 | 1 | 0.0545 | 0.0100 | 0.0783 | 0.0144 | -0.0012 | 0.0014 | 23.20 | 1.27 | 0.0140 | 0.0012 | 0.53 | 0.09 | 0.0148 | 0.0483 | 0.0021 | 2.4682 | 0.0018 | 0.2297 | 0.35 | 0.28 | 0.52 |
| NX Q9Y6R7-1 | FCGBP | 25 | 18 | 0.0861 | 0.0020 | 0.0931 | 0.0027 | 0.0026 | 0.0024 | 16.60 | 0.20 | 0.0259 | 0.0007 | 0.34 | 0.01 | 0.0031 | 0.0040 | 0.0025 | 0.4238 | 0.0010 | 0.0121 | 0.79 | 0.38 | 14.94 |
| NX Q9UGM5-1 | FETUB | 2 | 1 | 0.0274 | 0.0038 | 0.1672 | 0.0343 | 0.0022 | 0.0022 | 24.47 | 0.00 | 0.0130 | 0.0006 | 1.05 | 0.15 | 0.0069 | 0.0719 | 0.0023 | 0.0720 | 0.0008 | 0.2760 | 0.06 | 40.77 | 0.36 |
| NX P02671-1 | FGA | 1 | 1 | 0.0980 | 0.0878 | 0.2224 | 0.2496 | 0.8360 | 0.8736 | 15.40 | 9.07 | 0.0699 | 0.0371 | 0.29 | 0.27 | 1.1700 | 0.9065 | 0.8640 | 9.3283 | 0.0433 | 4688.4616 | 0.06 | 5717.00 | 0.98 |
| NX P02679-1 | FGG | 1 | 1 | 0.0538 | 0.0047 | 0.0053 | 0.0006 | 0.0002 | 0.0001 | 23.40 | 1.07 | 0.0009 | 0.0001 | 0.54 | 0.05 | 0.0087 | 0.0012 | 0.0002 | 1.8075 | 0.0001 | 0.0963 | 0.12 | 6623.00 | 1.24 |
| NX P02751-1 | FN1 | 16 | 11 | 0.0680 | 0.0028 | 0.0760 | 0.0032 | -0.0010 | 0.0010 | 19.60 | 0.60 | 0.0168 | 0.0005 | 0.42 | 0.02 | 0.0043 | 0.0049 | 0.0012 | 0.9364 | 0.0008 | 0.0276 | 18.78 | 237.40 | 943.58 |
| NX P14207-1 | FOLR2 | 1 | 1 | 0.0138 | 0.0007 | 0.4744 | 0.0212 | 0.0006 | 0.0009 | 24.47 | 0.00 | 0.0122 | 0.0012 | 2.10 | 0.11 | 0.0013 | 0.0538 | 0.0013 | 0.0000 | 0.0019 | 0.2494 | 3.20 | 2.52 | 16.35 |
| NX Q5SYB0-1 | FRMPD1 | 1 | 1 | 0.1519 | 0.0048 | 0.1778 | 0.0095 | -0.0296 | 0.0300 | 12.00 | 0.20 | 0.0830 | 0.0039 | 0.19 | 0.01 | 0.0076 | 0.0144 | 0.0304 | 0.2987 | 0.0058 | 0.0094 | 1.19 | 0.21 | 0.77 |
| NX Q12841-1 | FSTL1 | 2 | 2 | 0.1400 | 0.0068 | 0.0975 | 0.0083 | -0.0022 | 0.0042 | 12.60 | 0.30 | 0.0424 | 0.0028 | 0.21 | 0.01 | 0.0093 | 0.0119 | 0.0051 | 0.4210 | 0.0042 | 0.0137 | 10.32 | 9.14 | 165.47 |
| NX P04066-1 | FUCA1 | 1 | 1 | 0.1039 | 0.0024 | 0.0740 | 0.0028 | 0.0028 | 0.0028 | 14.80 | 0.20 | 0.0246 | 0.0009 | 0.28 | 0.01 | 0.0036 | 0.0042 | 0.0030 | 0.3193 | 0.0013 | 0.0098 | 10.93 | 24.74 | 36.47 |
| NX P02774-1 | GC | 7 | 7 | 0.0238 | 0.0022 | 0.1447 | 0.0162 | -0.0005 | 0.0008 | 24.47 | 0.00 | 0.0091 | 0.0005 | 1.21 | 0.12 | 0.0064 | 0.0678 | 0.0010 | 0.0378 | 0.0007 | 0.3266 | 0.01 | 984.20 | 0.65 |
| NX P17900-1 | GM2A | 2 | 2 | 0.0726 | 0.0053 | 0.0946 | 0.0075 | 0.0029 | 0.0025 | 18.80 | 1.00 | 0.0224 | 0.0015 | 0.40 | 0.03 | 0.0082 | 0.0131 | 0.0031 | 1.6358 | 0.0021 | 0.0509 | 21.64 | 7.62 | 31.74 |
| NX Q8NBJ4-1 | GOLM1 | 1 | 1 | 0.0253 | 0.0088 | 0.2718 | 0.1363 | -0.0032 | 0.0047 | 24.47 | 0.00 | 0.0188 | 0.0016 | 1.14 | 0.41 | 0.0150 | 0.2488 | 0.0051 | 0.9704 | 0.0025 | 0.5588 | 61.32 | 7.31 | 36.61 |
| NX P17174-1 | GOT1 | 1 | 1 | 0.1759 | 0.0113 | 0.3327 | 0.0465 | 0.0128 | 0.0169 | 11.40 | 0.40 | 0.1756 | 0.0188 | 0.16 | 0.01 | 0.0178 | 0.0672 | 0.0241 | 0.4500 | 0.0285 | 0.0157 | 88.76 | 166.80 | 48.51 |
| NX P22352-1 | GPX3 | 1 | 1 | 0.1770 | 0.0086 | 0.0683 | 0.0057 | 0.0283 | 0.0282 | 11.20 | 0.20 | 0.0362 | 0.0025 | 0.16 | 0.01 | 0.0686 | 0.1015 | 0.0323 | 4.2962 | 0.0126 | 9395.5077 | 112.16 | 384.50 | 1406.70 |
| NX P06396-1 | GSN | 7 | 6 | 0.0719 | 0.0031 | 0.0873 | 0.0043 | 0.0024 | 0.0024 | 18.80 | 0.60 | 0.0204 | 0.0009 | 0.40 | 0.02 | 0.0046 | 0.0063 | 0.0026 | 0.9064 | 0.0012 | 0.0267 | 52.31 | 8.52 | 301.76 |
| NX P02042-1 | HBD | 1 | 1 | 0.0101 | 0.0009 | 0.6178 | 0.0725 | -0.0009 | 0.0011 | 24.47 | 0.00 | 0.0093 | 0.0009 | 2.85 | 0.28 | 0.0012 | 0.0966 | 0.0015 | 0.0000 | 0.0013 | 0.4186 | 2.01 | 3.55 | 427.78 |
| NX P02790-1 | HPX | 1 | 1 | 0.0106 | 0.0008 | 0.2343 | 0.0372 | 0.0036 | 0.0037 | 24.47 | 0.00 | 0.0038 | 0.0003 | 2.71 | 0.21 | 0.0028 | 0.0764 | 0.0037 | 0.0000 | 0.0004 | 0.8493 | 1.58 | 2039.00 | 1.78 |
| NX P00738-1 | HP | 1 | 1 | 0.0361 | 0.0115 | 0.0460 | 0.0151 | -0.0002 | 0.0007 | 24.47 | 0.00 | 0.0052 | 0.0006 | 0.80 | 0.24 | 0.0163 | 0.0538 | 0.0010 | 1.6720 | 0.0009 | 0.5396 | 1.44 | 12710.00 | 8.98 |
| NX P04196-1 | HRG | 1 | 1 | 0.0081 | 0.0003 | 0.6458 | 0.0341 | -0.0002 | 0.0003 | 24.47 | 0.00 | 0.0065 | 0.0004 | 3.57 | 0.12 | 0.0005 | 0.0517 | 0.0004 | 0.0000 | 0.0006 | 0.2769 | 0.02 | 582.90 | 0.19 |
| NX Q92743-1 | HTRA1 | 1 | 1 | 0.2133 | 0.0106 | 0.0535 | 0.0045 | -0.0021 | 0.0027 | 10.60 | 0.20 | 0.0329 | 0.0023 | 0.14 | 0.01 | 0.0159 | 0.0067 | 0.0036 | 0.2657 | 0.0035 | 0.0099 | 231.63 | 88.63 | 194.16 |
| NX P01344-1 | IGF2 | 1 | 1 | 0.1757 | 0.0251 | 0.0325 | 0.0094 | -0.0267 | 0.0257 | 11.40 | 0.60 | 0.0172 | 0.0037 | 0.16 | 0.02 | 0.1229 | 0.2107 | 0.0237 | 6.9489 | 0.0105 | 4268.0335 | 4.91 | 82.23 | 29.06 |
| NX P35858-1 | IGFALS | 2 | 2 | 0.0089 | 0.0004 | 0.6778 | 0.0277 | -0.0018 | 0.0018 | 24.47 | 0.00 | 0.0080 | 0.0007 | 3.25 | 0.15 | 0.0007 | 0.0510 | 0.0019 | 0.0000 | 0.0010 | 0.2808 | 1.22 | 65.17 | 1.02 |
| NX P18065-1 | IGFBP2 | 8 | 5 | 0.1535 | 0.0024 | 0.0884 | 0.0025 | -0.0035 | 0.0035 | 12.00 | 0.00 | 0.0416 | 0.0010 | 0.19 | 0.00 | 0.0036 | 0.0036 | 0.0037 | 0.1460 | 0.0014 | 0.0044 | 10.09 | 166.30 | 161.33 |
| NX P22692-1 | IGFBP4 | 2 | 2 | 0.1119 | 0.0069 | 0.1047 | 0.0096 | 0.0001 | 0.0027 | 14.20 | 0.60 | 0.0372 | 0.0032 | 0.26 | 0.02 | 0.0103 | 0.0146 | 0.0040 | 0.8108 | 0.0050 | 0.0252 | 41.29 | 989.30 | 716.72 |
| NX P24592-1 | IGFBP6 | 3 | 3 | 0.1035 | 0.0089 | 0.0574 | 0.0068 | -0.0097 | 0.0103 | 14.80 | 0.80 | 0.0190 | 0.0020 | 0.28 | 0.02 | 0.0132 | 0.0100 | 0.0106 | 1.2871 | 0.0030 | 0.0393 | 30.51 | 5.51 | 275.61 |
| NX Q16270-1 | IGFBP7 | 11 | 5 | 0.2559 | 0.0063 | 0.1233 | 0.0047 | -0.0050 | 0.0067 | 10.00 | 0.00 | 0.0859 | 0.0027 | 0.11 | 0.00 | 0.0098 | 0.0072 | 0.0073 | 0.1103 | 0.0040 | 0.0043 | 95.31 | 109.30 | 827.40 |
| NX P01876-1 | IGHA1 | 7 | 3 | 0.0907 | 0.0022 | 0.0751 | 0.0021 | 0.0009 | 0.0009 | 16.20 | 0.20 | 0.0219 | 0.0006 | 0.32 | 0.01 | 0.0031 | 0.0032 | 0.0012 | 0.3671 | 0.0008 | 0.0110 | 3.32 | 120.60 | 927.71 |
| NX P01857-1 | IGHG1 | 4 | 2 | 0.1062 | 0.0027 | 0.0621 | 0.0029 | 0.0022 | 0.0021 | 14.60 | 0.20 | 0.0210 | 0.0009 | 0.27 | 0.01 | 0.0041 | 0.0044 | 0.0024 | 0.3494 | 0.0014 | 0.0106 | 1.03 | 36.82 | 155.78 |
| NX P01859-1 | IGHG2 | 4 | 3 | 0.1093 | 0.0033 | 0.0785 | 0.0035 | 0.0047 | 0.0042 | 14.40 | 0.20 | 0.0273 | 0.0010 | 0.26 | 0.01 | 0.0045 | 0.0051 | 0.0043 | 0.3589 | 0.0016 | 0.0111 | 0.72 | 27.10 | 91.29 |
| NX P01860-1 | IGHG3 | 1 | 1 | 0.1295 | 0.0077 | 0.0623 | 0.0078 | 0.0065 | 0.0059 | 13.00 | 0.40 | 0.0253 | 0.0029 | 0.22 | 0.01 | 0.0117 | 0.0118 | 0.0067 | 0.6474 | 0.0044 | 0.0204 | 0.59 | 17.46 | 63.20 |
| NX P01871-1 | IGHM | 11 | 7 | 0.0997 | 0.0019 | 0.0719 | 0.0020 | 0.0029 | 0.0027 | 15.20 | 0.20 | 0.0230 | 0.0006 | 0.29 | 0.01 | 0.0029 | 0.0029 | 0.0028 | 0.2822 | 0.0009 | 0.0085 | 1.00 | 34.16 | 361.92 |
| NX A0A0B4J1X8-1 | IGHV3-43 | 1 | 1 | 0.1011 | 0.0025 | 0.1156 | 0.0034 | 0.0003 | 0.0008 | 15.00 | 0.20 | 0.0374 | 0.0010 | 0.29 | 0.01 | 0.0036 | 0.0053 | 0.0012 | 0.3359 | 0.0016 | 0.0101 | 0.00 | 0.49 | 3.70 |
| NX A0A0B4J1X5-1 | IGHV3-74 | 1 | 1 | 0.0518 | 0.0101 | 0.2045 | 0.0414 | 0.0087 | 0.0083 | 24.20 | 0.27 | 0.0348 | 0.0035 | 0.56 | 0.11 | 0.0155 | 0.1839 | 0.0093 | 2.5407 | 0.0050 | 0.3114 | 0.06 | 1.84 | 12.09 |
| NX P01834-1 | IGKC | 17 | 6 | 0.1101 | 0.0017 | 0.0740 | 0.0018 | 0.0021 | 0.0020 | 14.40 | 0.20 | 0.0259 | 0.0006 | 0.26 | 0.00 | 0.0025 | 0.0028 | 0.0021 | 0.2158 | 0.0009 | 0.0061 | 0.00 | 0.00 | 0.10 |
| NX P01599-1 | IGKV1-17 | 1 | 1 | 0.0980 | 0.0036 | 0.1159 | 0.0061 | 0.0007 | 0.0015 | 15.40 | 0.40 | 0.0365 | 0.0018 | 0.29 | 0.01 | 0.0053 | 0.0092 | 0.0022 | 0.5307 | 0.0027 | 0.0162 | 0.02 | 2.14 | 11.73 |
| NX P01619-1 | IGKV3-20 | 1 | 1 | 0.1060 | 0.0020 | 0.0848 | 0.0027 | 0.0019 | 0.0020 | 14.60 | 0.20 | 0.0287 | 0.0008 | 0.27 | 0.01 | 0.0031 | 0.0039 | 0.0022 | 0.2708 | 0.0012 | 0.0080 | 0.52 | 20.39 | 130.92 |
| NX A0A087WSY6-1 | IGKV3D-15 | 1 | 1 | 0.1347 | 0.0048 | 0.0675 | 0.0069 | 0.0053 | 0.0047 | 12.80 | 0.20 | 0.0284 | 0.0025 | 0.21 | 0.01 | 0.0074 | 0.0102 | 0.0056 | 0.3622 | 0.0037 | 0.0117 | 0.00 | 0.39 | 3.10 |
| NX P06312-1 | IGKV4-1 | 2 | 2 | 0.1242 | 0.0059 | 0.0834 | 0.0061 | 0.0060 | 0.0048 | 13.40 | 0.40 | 0.0326 | 0.0021 | 0.23 | 0.01 | 0.0084 | 0.0094 | 0.0054 | 0.5063 | 0.0033 | 0.0160 | 0.21 | 8.65 | 69.69 |
| NX B9A064-1 | IGLL5 | 2 | 1 | 0.1103 | 0.0014 | 0.0697 | 0.0013 | 0.0030 | 0.0031 | 14.40 | 0.20 | 0.0245 | 0.0004 | 0.26 | 0.00 | 0.0021 | 0.0020 | 0.0032 | 0.1894 | 0.0007 | 0.0050 | 0.00 | 0.26 | 2.13 |
| NX P01700-1 | IGLV1-47 | 2 | 1 | 0.1082 | 0.0031 | 0.0580 | 0.0027 | 0.0031 | 0.0030 | 14.40 | 0.20 | 0.0200 | 0.0009 | 0.27 | 0.01 | 0.0046 | 0.0040 | 0.0031 | 0.3781 | 0.0013 | 0.0115 | 0.08 | 3.85 | 23.83 |
| NX P19827-1 | ITIH1 | 1 | 1 | 0.0110 | 0.0057 | 0.5042 | 0.1083 | -0.0023 | 0.0037 | 24.47 | 0.00 | 0.0088 | 0.0066 | 2.62 | 1.84 | 0.0094 | 0.1749 | 0.0103 | 2.4482 | 0.0106 | 2734815.0134 | 0.04 | 541.60 | 0.24 |
| NX P19823-1 | ITIH2 | 4 | 4 | 0.0801 | 0.0040 | 0.0784 | 0.0047 | 0.0096 | 0.0090 | 17.60 | 0.60 | 0.0204 | 0.0012 | 0.36 | 0.02 | 0.0071 | 0.0108 | 0.0092 | 0.9424 | 0.0018 | 0.1188 | 1.46 | 267.50 | 0.20 |
| NX Q06033-1 | ITIH3 | 2 | 2 | 0.0108 | 0.0011 | 0.8432 | 0.0710 | 0.0048 | 0.0057 | 24.47 | 0.00 | 0.0142 | 0.0019 | 2.67 | 0.30 | 0.0014 | 0.1570 | 0.0060 | 0.0000 | 0.0027 | 0.4372 | 0.39 | 702.30 | 9.72 |
| NX Q14624-1 | ITIH4 | 2 | 1 | 0.0084 | 0.0007 | 0.6572 | 0.0731 | 0.0008 | 0.0010 | 24.47 | 0.00 | 0.0070 | 0.0004 | 3.44 | 0.32 | 0.0009 | 0.1072 | 0.0011 | 0.0000 | 0.0005 | 0.4074 | 5.93 | 1570.00 | 18.69 |
| NX Q86UX2-1 | ITIH5 | 1 | 1 | 0.1870 | 0.0065 | 0.2852 | 0.0169 | 0.0182 | 0.0178 | 11.00 | 0.20 | 0.1582 | 0.0078 | 0.15 | 0.01 | 0.0100 | 0.0247 | 0.0191 | 0.2389 | 0.0112 | 0.0082 | 6.69 | 0.41 | 21.21 |
| NX P01591-1 | JCHAIN | 5 | 4 | 0.0977 | 0.0040 | 0.1033 | 0.0066 | -0.0002 | 0.0017 | 15.40 | 0.40 | 0.0324 | 0.0019 | 0.30 | 0.01 | 0.0060 | 0.0102 | 0.0026 | 0.6071 | 0.0030 | 0.0186 | NA | NA | NA |
| NX Q92876-1 | KLK6 | 4 | 2 | 0.0258 | 0.0013 | 0.2439 | 0.0201 | 0.0009 | 0.0008 | 24.47 | 0.00 | 0.0174 | 0.0004 | 1.12 | 0.06 | 0.0033 | 0.0571 | 0.0009 | 0.0000 | 0.0005 | 0.1611 | 45.74 | 0.02 | 2.46 |
| NX P01042-1 | KNG1 | 3 | 2 | 0.0084 | 0.0010 | 0.7492 | 0.1581 | 0.0013 | 0.0012 | 24.47 | 0.00 | 0.0081 | 0.0005 | 3.43 | 0.45 | 0.0013 | 0.2453 | 0.0013 | 0.0000 | 0.0007 | 0.6853 | 0.05 | 410.40 | 1.78 |
| NX P13645-1 | KRT10 | 1 | 1 | 0.0114 | 0.0009 | 0.9560 | 0.0803 | 0.0066 | 0.0063 | 24.47 | 0.00 | 0.0178 | 0.0018 | 2.53 | 0.20 | 0.0015 | 0.1622 | 0.0065 | 0.0000 | 0.0027 | 0.4521 | 21.34 | 14.21 | 1202.16 |
| NX P13646-1 | KRT13 | 1 | 1 | 0.0030 | 0.0001 | 0.6879 | 0.0000 | 0.0001 | 0.0001 | 24.47 | 0.00 | 0.0011 | 0.0000 | 9.56 | 0.20 | 0.0002 | 0.0087 | 0.0001 | 0.0000 | 0.0002 | 0.9496 | 1.65 | 3.97 | 1255.03 |
| NX P04264-1 | KRT1 | 2 | 1 | 0.2113 | 0.0095 | 0.1405 | 0.0104 | -0.0242 | 0.0238 | 10.60 | 0.20 | 0.0856 | 0.0050 | 0.14 | 0.01 | 0.0145 | 0.0159 | 0.0246 | 0.2446 | 0.0079 | 0.0092 | 1.86 | 1.67 | 772.99 |
| NX Q6VAB6-1 | KSR2 | 1 | 1 | 0.0437 | 0.0125 | 0.0933 | 0.0217 | 0.0007 | 0.0016 | 24.47 | 0.00 | 0.0133 | 0.0017 | 0.66 | 0.18 | 0.0165 | 0.1026 | 0.0025 | 2.0062 | 0.0026 | 0.4708 | 4.57 | 0.01 | 0.17 |
| NX Q08380-1 | LGALS3BP | 3 | 3 | 0.1262 | 0.0048 | 0.1217 | 0.0086 | 0.0112 | 0.0101 | 13.20 | 0.20 | 0.0483 | 0.0032 | 0.23 | 0.01 | 0.0072 | 0.0126 | 0.0108 | 0.4245 | 0.0047 | 0.0132 | 64.19 | 44.65 | 181.02 |
| NX P49257-1 | LMAN1 | 1 | 1 | 0.0957 | 0.0060 | 0.1229 | 0.0103 | 0.0072 | 0.0074 | 15.60 | 0.60 | 0.0378 | 0.0027 | 0.30 | 0.02 | 0.0094 | 0.0147 | 0.0080 | 1.0355 | 0.0042 | 0.0317 | 18.03 | 63.65 | 43.26 |
| NX P28300-1 | LOX | 2 | 1 | 0.0640 | 0.0073 | 0.1202 | 0.0149 | 0.0068 | 0.0068 | 20.60 | 1.80 | 0.0251 | 0.0030 | 0.45 | 0.05 | 0.0119 | 0.0316 | 0.0081 | 2.3276 | 0.0044 | 0.1259 | 0.96 | 0.59 | 20.34 |
| NX P02750-1 | LRG1 | 4 | 4 | 0.0091 | 0.0003 | 0.6368 | 0.0245 | 0.0006 | 0.0003 | 24.47 | 0.00 | 0.0078 | 0.0004 | 3.18 | 0.09 | 0.0005 | 0.0526 | 0.0004 | 0.0000 | 0.0006 | 0.2053 | 0.27 | 647.40 | 10.58 |
| NX Q14767-1 | LTBP2 | 1 | 1 | 0.0999 | 0.0035 | 0.0471 | 0.0036 | -0.0004 | 0.0011 | 15.20 | 0.40 | 0.0151 | 0.0011 | 0.29 | 0.01 | 0.0052 | 0.0050 | 0.0015 | 0.4938 | 0.0016 | 0.0151 | 2.03 | 1.14 | 66.80 |
| NX Q9Y5Y7-1 | LYVE1 | 2 | 1 | 0.0350 | 0.0012 | 0.1014 | 0.0046 | 0.0008 | 0.0008 | 24.47 | 0.00 | 0.0110 | 0.0002 | 0.83 | 0.03 | 0.0024 | 0.0096 | 0.0008 | 0.0000 | 0.0002 | 0.0596 | 2.78 | 6.73 | 36.30 |
| NX P61626-1 | LYZ | 1 | 1 | 0.1372 | 0.0091 | 0.0473 | 0.0063 | 0.0006 | 0.0017 | 12.60 | 0.40 | 0.0202 | 0.0023 | 0.21 | 0.01 | 0.0143 | 0.0091 | 0.0026 | 0.6535 | 0.0034 | 0.0212 | 2.42 | 6.44 | 323.74 |
| NX Q9UHG2-1 | Little-SAAS | 1 | 1 | 0.1889 | 0.0089 | 0.0984 | 0.0093 | -0.0011 | 0.0037 | 11.00 | 0.20 | 0.0550 | 0.0041 | 0.15 | 0.01 | 0.0139 | 0.0132 | 0.0053 | 0.2996 | 0.0058 | 0.0109 | NA | NA | NA |
| NX Q7Z7M0-1 | MEGF8 | 3 | 2 | 0.1207 | 0.0060 | 0.0753 | 0.0058 | 0.0057 | 0.0050 | 13.60 | 0.40 | 0.0287 | 0.0020 | 0.24 | 0.01 | 0.0093 | 0.0085 | 0.0054 | 0.6029 | 0.0029 | 0.0190 | 28.09 | 3.79 | 14.69 |
| NX Q9UNW1-1 | MINPP1 | 1 | 1 | 0.0947 | 0.0067 | 0.0756 | 0.0073 | 0.0048 | 0.0045 | 15.60 | 0.60 | 0.0230 | 0.0019 | 0.30 | 0.02 | 0.0106 | 0.0111 | 0.0052 | 1.2683 | 0.0030 | 0.0382 | 6.68 | 13.51 | 10.85 |
| NX P14780-1 | MMP9 | 1 | 1 | 0.2874 | 0.0186 | 0.0483 | 0.0050 | -0.0053 | 0.0080 | 9.80 | 0.20 | 0.0362 | 0.0032 | 0.10 | 0.01 | 0.0268 | 0.0076 | 0.0088 | 0.2085 | 0.0048 | 0.0093 | 0.39 | 1.17 | 20.07 |
| NX P41271-1 | NBL1 | 2 | 1 | 0.0839 | 0.0018 | 0.1029 | 0.0035 | 0.0036 | 0.0035 | 17.00 | 0.20 | 0.0279 | 0.0010 | 0.34 | 0.01 | 0.0028 | 0.0050 | 0.0037 | 0.3819 | 0.0015 | 0.0114 | 64.71 | 1.90 | 196.03 |
| NX Q99435-1 | NELL2 | 1 | 1 | 0.1906 | 0.0456 | 0.2445 | 0.0593 | 0.1212 | 0.1353 | 11.00 | 1.00 | 0.1377 | 0.0456 | 0.15 | 0.03 | 0.1574 | 1.2424 | 0.1667 | 8.1893 | 0.0926 | 145023.1626 | 31.22 | 0.09 | 1.90 |
| NX P48745-1 | NOV | 1 | 1 | 0.1567 | 0.0098 | 0.1101 | 0.0116 | 0.0070 | 0.0084 | 12.00 | 0.40 | 0.0528 | 0.0047 | 0.18 | 0.01 | 0.0146 | 0.0174 | 0.0098 | 0.5092 | 0.0070 | 0.0170 | 7.86 | 0.59 | 79.60 |

| | | | | | | | | | | | | | | | | | | | | | | | | | |
|---|---|---|---|---|---|---|---|---|---|---|---|---|---|---|---|---|---|---|---|---|---|---|---|---|---|
| NX | Q86UD1-1 | OAF | 1 | 1 | 0.0240 | 0.0050 | 0.3207 | 0.1039 | -0.0034 | 0.0062 | 24.47 | 0.00 | 0.0204 | 0.0037 | 1.20 | 0.29 | 0.0154 | 0.1878 | 0.0093 | 1.0430 | 0.0063 | 0.5042 | 17.09 | 187.20 | 42.77 |
| NX | P20774-1 | OGN | 2 | 2 | 0.1153 | 0.0050 | 0.0551 | 0.0040 | 0.0046 | 0.0046 | 14.00 | 0.40 | 0.0201 | 0.0013 | 0.25 | 0.01 | 0.0078 | 0.0059 | 0.0048 | 0.5446 | 0.0020 | 0.0170 | 2.23 | 0.40 | 57.87 |
| NX | P02763-1 | ORM1 | 1 | 1 | 0.0148 | 0.0010 | 0.1227 | 0.0061 | 0.0005 | 0.0006 | 24.47 | 0.00 | 0.0036 | 0.0003 | 1.95 | 0.13 | 0.0047 | 0.0295 | 0.0006 | 0.1639 | 0.0005 | 0.3050 | 0.17 | 5384.00 | 1.75 |
| NX | P19021-1 | PAM | 1 | 1 | 0.1423 | 0.0078 | 0.1448 | 0.0186 | 0.0061 | 0.0076 | 12.40 | 0.40 | 0.0639 | 0.0070 | 0.20 | 0.01 | 0.0125 | 0.0256 | 0.0101 | 0.5441 | 0.0100 | 0.0177 | 28.70 | 2.72 | 79.80 |
| NX | Q15113-1 | PCOLCE | 2 | 2 | 0.1033 | 0.0045 | 0.0833 | 0.0062 | 0.0044 | 0.0035 | 14.80 | 0.40 | 0.0275 | 0.0020 | 0.28 | 0.01 | 0.0067 | 0.0091 | 0.0041 | 0.6167 | 0.0028 | 0.0187 | 12.57 | 35.88 | 94.94 |
| NX | Q9UHG2-1 | PCSK1N | 5 | 4 | 0.0991 | 0.0055 | 0.0545 | 0.0039 | -0.0100 | 0.0104 | 15.22 | 0.58 | 0.0173 | 0.0011 | 0.29 | 0.02 | 0.0081 | 0.0058 | 0.0105 | 0.7924 | 0.0016 | 0.0242 | 432.06 | 0.32 | 10.13 |
| NX | P30086-1 | PEBP1 | 2 | 2 | 0.1560 | 0.0093 | 0.0587 | 0.0070 | -0.0074 | 0.0087 | 12.00 | 0.40 | 0.0280 | 0.0029 | 0.19 | 0.01 | 0.0150 | 0.0103 | 0.0092 | 0.5102 | 0.0042 | 0.0172 | 910.30 | 1000.00 | 459.39 |
| NX | P01210-1 | PENK | 2 | 1 | 0.0105 | 0.0004 | 0.5613 | 0.0254 | -0.0001 | 0.0003 | 24.47 | 0.00 | 0.0091 | 0.0005 | 2.74 | 0.10 | 0.0007 | 0.0456 | 0.0005 | 0.0000 | 0.0007 | 0.1971 | 41.57 | 0.02 | 4.41 |
| NX | Q9UHG2-1 | PEN | 1 | 1 | 0.0869 | 0.0108 | 0.1205 | 0.0198 | 0.0074 | 0.0081 | 16.60 | 1.38 | 0.0338 | 0.0055 | 0.33 | 0.04 | 0.0175 | 0.0585 | 0.0102 | 2.6520 | 0.0084 | 0.1458 | NA | NA | NA |
| NX | P00747-1 | PLG | 3 | 3 | 0.0100 | 0.0008 | 0.4340 | 0.0402 | -0.0030 | 0.0031 | 24.47 | 0.00 | 0.0064 | 0.0005 | 2.88 | 0.25 | 0.0013 | 0.0672 | 0.0031 | 0.0000 | 0.0007 | 0.3633 | 0.04 | 324.50 | 0.33 |
| NX | P55058-1 | PLTP | 6 | 3 | 0.1547 | 0.0054 | 0.0954 | 0.0066 | 0.0108 | 0.0111 | 12.00 | 0.20 | 0.0453 | 0.0026 | 0.19 | 0.01 | 0.0083 | 0.0100 | 0.0116 | 0.2990 | 0.0039 | 0.0100 | 46.61 | 8.62 | 109.61 |
| NX | P07225-1 | PROS1 | 2 | 1 | 0.1230 | 0.0027 | 0.0712 | 0.0023 | -0.0005 | 0.0007 | 13.40 | 0.20 | 0.0276 | 0.0008 | 0.23 | 0.01 | 0.0039 | 0.0036 | 0.0011 | 0.2412 | 0.0013 | 0.0074 | 4.99 | 83.70 | 35.29 |
| NX | P07602-1 | PSAP | 3 | 3 | 0.2677 | 0.0186 | 0.0477 | 0.0051 | -0.0292 | 0.0303 | 9.80 | 0.20 | 0.0342 | 0.0029 | 0.11 | 0.01 | 0.0276 | 0.0073 | 0.0303 | 0.2858 | 0.0046 | 0.0113 | 661.32 | 360.70 | 921.19 |
| NX | P41222-1 | PTGDS | 12 | 4 | 0.0928 | 0.0025 | 0.0891 | 0.0031 | 0.0004 | 0.0007 | 15.80 | 0.20 | 0.0266 | 0.0008 | 0.31 | 0.01 | 0.0038 | 0.0046 | 0.0010 | 0.4511 | 0.0012 | 0.0131 | 807.06 | 14.48 | 326.35 |
| NX | Q9UHG2-1 | ProSAAS | 3 | 3 | 0.0958 | 0.0061 | 0.0555 | 0.0039 | -0.0072 | 0.0075 | 15.60 | 0.60 | 0.0171 | 0.0010 | 0.30 | 0.02 | 0.0087 | 0.0059 | 0.0076 | 0.9336 | 0.0015 | 0.0285 | NA | NA | NA |
| NX | P07602-1 | Prosaposin | 2 | 2 | 0.2768 | 0.0105 | 0.0858 | 0.0053 | -0.0080 | 0.0086 | 9.80 | 0.00 | 0.0629 | 0.0031 | 0.10 | 0.00 | 0.0157 | 0.0077 | 0.0091 | 0.1393 | 0.0045 | 0.0059 | NA | NA | NA |
| NX | O00391-1 | QSOX1 | 2 | 1 | 0.1112 | 0.0059 | 0.0669 | 0.0036 | 0.0072 | 0.0063 | 14.20 | 0.40 | 0.0237 | 0.0012 | 0.26 | 0.01 | 0.0109 | 0.0203 | 0.0066 | 0.8855 | 0.0019 | 0.0897 | 12.55 | 11.58 | 49.97 |
| NX | P02753-1 | RBP4 | 2 | 2 | 0.0524 | 0.0063 | 0.1747 | 0.0207 | 0.0020 | 0.0020 | 24.00 | 0.47 | 0.0300 | 0.0018 | 0.55 | 0.06 | 0.0097 | 0.0562 | 0.0029 | 1.8064 | 0.0026 | 0.1412 | 49.19 | 4148.00 | 36.06 |
| NX | P78509-1 | RELN | 2 | 2 | 0.0562 | 0.0123 | 0.2310 | 0.0524 | 0.0209 | 0.0240 | 22.60 | 1.87 | 0.0425 | 0.0088 | 0.51 | 0.10 | 0.0189 | 0.2328 | 0.0269 | 2.5656 | 0.0132 | 0.4198 | 22.39 | 3.60 | 2.11 |
| NX | P35249-1 | RFC4 | 1 | 1 | 0.0146 | 0.0014 | 0.3902 | 0.0453 | 0.0036 | 0.0046 | 24.47 | 0.00 | 0.0112 | 0.0011 | 1.97 | 0.21 | 0.0020 | 0.0575 | 0.0048 | 0.0000 | 0.0017 | 0.3190 | 7.93 | 3.80 | 11.44 |
| NX | O00584-1 | RNASET2 | 1 | 1 | 0.1679 | 0.0138 | 0.0565 | 0.0170 | 0.0076 | 0.0079 | 11.60 | 0.40 | 0.0287 | 0.0076 | 0.17 | 0.01 | 0.0421 | 0.0230 | 0.0111 | 0.7730 | 0.0103 | 0.0247 | 20.65 | 8.32 | 39.55 |
| NX | P62266-1 | RPS23 | 1 | 1 | 0.1237 | 0.0022 | 0.2053 | 0.0119 | 0.0181 | 0.0195 | 13.40 | 0.20 | 0.0800 | 0.0040 | 0.23 | 0.00 | 0.0033 | 0.0172 | 0.0202 | 0.2038 | 0.0059 | 0.0063 | 112.88 | 53.54 | 254.09 |
| NX | P13521-1 | SCG2 | 2 | 2 | 0.2235 | 0.0198 | 0.0267 | 0.0037 | 0.0008 | 0.0016 | 10.40 | 0.23 | 0.0169 | 0.0019 | 0.13 | 0.01 | 0.0333 | 0.0054 | 0.0023 | 0.4537 | 0.0028 | 0.0181 | 132.55 | 0.06 | 2.05 |
| NX | Q8WXD2-1 | SCG3 | 8 | 6 | 0.1350 | 0.0050 | 0.2147 | 0.0181 | 0.0319 | 0.0329 | 12.80 | 0.20 | 0.0904 | 0.0061 | 0.21 | 0.01 | 0.0073 | 0.0262 | 0.0339 | 0.3572 | 0.0093 | 0.0115 | 84.02 | 0.04 | 0.55 |
| NX | P01009-1 | SERPINA1 | 5 | 4 | 0.0092 | 0.0006 | 0.4309 | 0.0342 | 0.0005 | 0.0004 | 24.47 | 0.00 | 0.0054 | 0.0002 | 3.14 | 0.21 | 0.0007 | 0.0592 | 0.0005 | 0.0000 | 0.0003 | 0.2765 | 2.31 | 6242.00 | 46.15 |
| NX | P01011-1 | SERPINA3 | 12 | 8 | 0.1038 | 0.0020 | 0.0406 | 0.0011 | 0.0020 | 0.0026 | 14.80 | 0.20 | 0.0135 | 0.0003 | 0.28 | 0.01 | 0.0030 | 0.0016 | 0.0026 | 0.2702 | 0.0005 | 0.0081 | 13.03 | 1722.00 | 83.69 |
| NX | P05543-1 | SERPINA7 | 1 | 1 | 0.0461 | 0.0144 | 0.0274 | 0.0071 | 0.0016 | 0.0019 | 24.47 | 0.00 | 0.0041 | 0.0008 | 0.63 | 0.18 | 0.0214 | 0.0278 | 0.0022 | 2.0693 | 0.0012 | 0.8094 | 0.00 | 67.86 | 0.02 |
| NX | P01008-1 | SERPINC1 | 5 | 4 | 0.0078 | 0.0004 | 0.7142 | 0.0609 | -0.0014 | 0.0017 | 24.47 | 0.00 | 0.0067 | 0.0004 | 3.68 | 0.18 | 0.0008 | 0.1265 | 0.0018 | 0.0000 | 0.0005 | 0.3619 | 0.18 | 1045.00 | 0.30 |
| NX | P05546-1 | SERPIND1 | 6 | 6 | 0.0876 | 0.0035 | 0.0508 | 0.0024 | 0.0002 | 0.0006 | 16.60 | 0.40 | 0.0144 | 0.0006 | 0.33 | 0.01 | 0.0050 | 0.0036 | 0.0008 | 0.6344 | 0.0009 | 0.0191 | 0.41 | 196.20 | 0.44 |
| NX | P36955-1 | SERPINF1 | 18 | 8 | 0.2182 | 0.0029 | 0.0658 | 0.0014 | -0.0142 | 0.0145 | 10.40 | 0.00 | 0.0411 | 0.0007 | 0.13 | 0.00 | 0.0045 | 0.0022 | 0.0145 | 0.1151 | 0.0011 | 0.0027 | 14.45 | 285.70 | 259.09 |
| NX | P08697-1 | SERPINF2 | 3 | 2 | 0.0483 | 0.0093 | 0.0784 | 0.0164 | -0.0010 | 0.0022 | 24.47 | 0.00 | 0.0124 | 0.0009 | 0.60 | 0.11 | 0.0135 | 0.0574 | 0.0025 | 2.1823 | 0.0014 | 0.2693 | 1.45 | 1145.00 | 11.79 |
| NX | P05155-1 | SERPING1 | 8 | 5 | 0.0947 | 0.0029 | 0.0774 | 0.0033 | -0.0006 | 0.0009 | 15.60 | 0.29 | 0.0236 | 0.0009 | 0.30 | 0.01 | 0.0043 | 0.0047 | 0.0013 | 0.4635 | 0.0013 | 0.0138 | 28.96 | 1148.00 | 499.81 |
| NX | Q9BRG2-1 | SH2D3A | 1 | 1 | 0.0109 | 0.0007 | 0.6842 | 0.0356 | 0.0012 | 0.0015 | 24.47 | 0.00 | 0.0117 | 0.0016 | 2.66 | 0.18 | 0.0015 | 0.1026 | 0.0021 | 0.0000 | 0.0024 | 0.5253 | 0.47 | 1.59 | 12.26 |
| NX | Q99963-1 | SH3GL3 | 1 | 1 | 0.1000 | 0.0020 | 0.0914 | 0.0025 | 0.0015 | 0.0017 | 15.20 | 0.20 | 0.0293 | 0.0008 | 0.29 | 0.01 | 0.0030 | 0.0037 | 0.0019 | 0.2914 | 0.0011 | 0.0088 | 35.92 | 0.01 | 3.08 |
| NX | Q9H0K1-1 | SIK2 | 1 | 1 | 0.0167 | 0.0020 | 0.0566 | 0.0081 | 0.0006 | 0.0007 | 24.47 | 0.00 | 0.0020 | 0.0002 | 1.73 | 0.21 | 0.0035 | 0.0165 | 0.0007 | 0.0000 | 0.0003 | 0.4314 | 13.94 | 16.68 | 17.17 |
| NX | O95219-1 | SNX4 | 1 | 1 | 0.0123 | 0.0009 | 0.2204 | 0.0180 | 0.0000 | 0.0004 | 24.47 | 0.00 | 0.0047 | 0.0005 | 2.35 | 0.18 | 0.0016 | 0.0348 | 0.0005 | 0.0000 | 0.0008 | 0.3822 | 19.36 | 18.85 | 23.44 |
| NX | P08294-1 | SOD3 | 3 | 3 | 0.1690 | 0.0073 | 0.0909 | 0.0078 | -0.0037 | 0.0067 | 11.60 | 0.20 | 0.0464 | 0.0032 | 0.17 | 0.01 | 0.0115 | 0.0117 | 0.0078 | 0.3391 | 0.0049 | 0.0113 | 11.04 | 7.67 | 249.26 |
| NX | Q14515-1 | SPARCL1 | 5 | 3 | 0.1739 | 0.0054 | 0.0789 | 0.0043 | -0.0105 | 0.0107 | 11.40 | 0.20 | 0.0413 | 0.0021 | 0.17 | 0.01 | 0.0083 | 0.0071 | 0.0110 | 0.2272 | 0.0031 | 0.0078 | 646.66 | 7.23 | 698.20 |
| NX | P09486-1 | SPARC | 1 | 1 | 0.2626 | 0.0094 | 0.1360 | 0.0064 | -0.0093 | 0.0104 | 10.00 | 0.00 | 0.0964 | 0.0038 | 0.11 | 0.00 | 0.0136 | 0.0099 | 0.0110 | 0.1474 | 0.0057 | 0.0057 | 385.02 | 58.13 | 651.26 |
| NX | Q96T58-1 | SPEN | 1 | 1 | 0.0902 | 0.0046 | 0.2164 | 0.0151 | 0.0388 | 0.0395 | 16.20 | 0.60 | 0.0629 | 0.0042 | 0.32 | 0.02 | 0.0070 | 0.0233 | 0.0396 | 0.8479 | 0.0064 | 0.0258 | 16.95 | 8.52 | 26.67 |
| NX | P10451-1 | SPP1 | 11 | 7 | 0.0950 | 0.0031 | 0.0719 | 0.0029 | 0.0007 | 0.0008 | 15.60 | 0.38 | 0.0220 | 0.0008 | 0.30 | 0.01 | 0.0046 | 0.0042 | 0.0011 | 0.4980 | 0.0011 | 0.0149 | 271.11 | 7.49 | 76.36 |
| NX | O00186-1 | STXBP3 | 1 | 1 | 0.0086 | 0.0008 | 0.7361 | 0.1103 | 0.0035 | 0.0042 | 24.47 | 0.00 | 0.0082 | 0.0008 | 3.37 | 0.32 | 0.0012 | 0.2274 | 0.0043 | 0.0000 | 0.0013 | 0.6104 | 19.07 | 7.45 | 21.73 |
| NX | P07602-1 | Saposin-C | 1 | 1 | 0.1061 | 0.0072 | 0.0452 | 0.0044 | -0.0086 | 0.0083 | 14.60 | 0.60 | 0.0153 | 0.0013 | 0.27 | 0.02 | 0.0151 | 0.0411 | 0.0084 | 1.3038 | 0.0021 | 0.3454 | NA | NA | NA |
| NX | P02787-1 | TF | 19 | 11 | 0.0081 | 0.0004 | 0.6092 | 0.0363 | 0.0014 | 0.0013 | 24.47 | 0.00 | 0.0062 | 0.0002 | 3.55 | 0.17 | 0.0013 | 0.1834 | 0.0013 | 0.0000 | 0.0003 | 1.1773 | 251.89 | 1601.00 | 6.06 |
| NX | Q15582-1 | TGFBI | 1 | 1 | 0.1225 | 0.0034 | 0.0189 | 0.0008 | 0.0051 | 0.0051 | 13.40 | 0.20 | 0.0073 | 0.0003 | 0.24 | 0.01 | 0.0170 | 0.0314 | 0.0052 | 1.4366 | 0.0008 | 0.4973 | 7.95 | 32.86 | 120.01 |
| NX | P01033-1 | TIMP1 | 4 | 3 | 0.1421 | 0.0038 | 0.0918 | 0.0053 | -0.0002 | 0.0018 | 12.40 | 0.20 | 0.0404 | 0.0020 | 0.20 | 0.01 | 0.0057 | 0.0078 | 0.0027 | 0.2698 | 0.0029 | 0.0082 | 39.61 | 69.63 | 346.16 |
| NX | A6NML5-1 | TMEM212 | 1 | 1 | 0.0694 | 0.0089 | 0.0712 | 0.0099 | -0.0057 | 0.0060 | 19.40 | 1.80 | 0.0161 | 0.0016 | 0.42 | 0.05 | 0.0135 | 0.0281 | 0.0063 | 2.5224 | 0.0024 | 0.1305 | 0.01 | 0.00 | 0.03 |
| NX | Q13641-1 | TPBG | 1 | 1 | 0.1489 | 0.0148 | 0.2480 | 0.0597 | 0.0446 | 0.0438 | 12.20 | 0.60 | 0.1138 | 0.0267 | 0.19 | 0.02 | 0.0368 | 0.1372 | 0.0562 | 2.0418 | 0.0393 | 0.2432 | 3.68 | 0.47 | 15.12 |
| NX | P02766-1 | TTR | 1 | 1 | 0.1856 | 0.0195 | 0.0874 | 0.0195 | -0.0116 | 0.0128 | 11.00 | 0.40 | 0.0482 | 0.0083 | 0.16 | 0.02 | 0.0309 | 0.0271 | 0.0158 | 0.6424 | 0.0119 | 0.0232 | 11.90 | 2726.00 | 6.98 |
| NX | Q6EMK4-1 | VASN | 1 | 1 | 0.0875 | 0.0085 | 0.0439 | 0.0077 | 0.0024 | 0.0027 | 16.60 | 1.00 | 0.0124 | 0.0023 | 0.33 | 0.03 | 0.0144 | 0.0124 | 0.0037 | 2.2156 | 0.0034 | 0.0931 | 6.20 | 20.53 | 61.06 |
| NX | P19320-1 | VCAM1 | 1 | 1 | 0.1147 | 0.0074 | 0.1097 | 0.0096 | -0.0046 | 0.0059 | 14.00 | 0.60 | 0.0399 | 0.0033 | 0.25 | 0.02 | 0.0119 | 0.0143 | 0.0067 | 0.8803 | 0.0049 | 0.0277 | 2.73 | 2.83 | 20.39 |
| NX | P04004-1 | VTN | 4 | 4 | 0.0197 | 0.0010 | 0.4260 | 0.0247 | 0.0051 | 0.0045 | 24.47 | 0.00 | 0.0199 | 0.0008 | 1.47 | 0.07 | 0.0042 | 0.1304 | 0.0046 | 0.0000 | 0.0012 | 0.3388 | 4.89 | 2164.00 | 5.24 |
