## Supplementary material for "*In Vivo* Large Scale Mapping Of Protein Turnover In The Human Cerebrospinal Fluid": TQ protein table

| patient | protein | gene | lambda | k.d | apex | r.max | shift |
|---|---|---|---|---|---|---|---|
| Pat1 | A1BG_HUMAN | A1BG | 0.6895 | 0.0052 | 24.47 | 0.0030 | 0 |
| Pat1 | A2MG_HUMAN | A2M | 0.0292 | 0.0840 | 17.00 | 0.0079 | 0 |
| Pat1 | ALBU_HUMAN | ALB | 0.0954 | 0.0064 | 24.47 | 0.0006 | 0 |
| Pat1 | AMBP_HUMAN | AMBP | 0.1144 | 0.0495 | 24.47 | 0.0186 | 0 |
| Pat1 | APOA1_HUMAN | APOA1 | 0.2138 | 0.0098 | 24.47 | 0.0030 | 0 |
| Pat1 | APOA4_HUMAN | APOA4 | 0.3584 | 0.0226 | 24.47 | 0.0209 | 0 |
| Pat1 | APOD_HUMAN | APOD | 0.0258 | 0.0862 | 16.60 | 0.0072 | 0 |
| Pat1 | APOE_HUMAN | APOE | 0.0348 | 0.0554 | 23.00 | 0.0063 | 0 |
| Pat1 | CO4B_HUMAN | C4B | 0.0671 | 0.0622 | 21.00 | 0.0136 | 0 |
| Pat1 | CO9_HUMAN | C9 | 0.6578 | 0.0036 | 24.47 | 0.0014 | 0 |
| Pat1 | CFAB_HUMAN | CFB | 0.0437 | 0.0484 | 24.47 | 0.0069 | 0 |
| Pat1 | CLUS_HUMAN | CLU | 0.0586 | 0.1559 | 12.00 | 0.0280 | 0 |
| Pat1 | CYTC_HUMAN | CST3 | 0.1033 | 0.0661 | 20.00 | 0.0223 | 0 |
| Pat1 | FA12_HUMAN | F12 | 0.4229 | 0.0076 | 24.47 | 0.0038 | 0 |
| Pat1 | VTDB_HUMAN | GC | 0.5583 | 0.0043 | 24.47 | 0.0017 | 0 |
| Pat1 | GELS_HUMAN | GSN | 0.0721 | 0.0528 | 23.80 | 0.0125 | 0 |
| Pat1 | NRP2_HUMAN | NRP2 | 0.1046 | 0.0651 | 20.38 | 0.0222 | 0 |
| Pat1 | RET4_HUMAN | RBP4 | 0.2017 | 0.0275 | 24.47 | 0.0157 | 0 |
| Pat1 | AACT_HUMAN | SERPINA3 | 0.0300 | 0.0728 | 18.80 | 0.0071 | 0 |
| Pat1 | PEDF_HUMAN | SERPINF1 | 0.0855 | 0.1934 | 11.00 | 0.0487 | 0 |
| Pat1 | TTHY_HUMAN | TTR | 0.1033 | 0.2079 | 10.60 | 0.0622 | 0 |
| Pat2 | A2MG_HUMAN | A2M | 0.0962 | 0.0529 | 23.80 | 0.0167 | 0 |
| Pat2 | ALBU_HUMAN | ALB | 0.1042 | 0.0077 | 41.50 | 0.0016 | 0 |
| Pat2 | APOA1_HUMAN | APOA1 | 0.7304 | 0.0051 | 41.50 | 0.0053 | 0 |
| Pat2 | APOD_HUMAN | APOD | 0.0470 | 0.0676 | 19.80 | 0.0104 | 0 |
| Pat2 | APOE_HUMAN | APOE | 0.1004 | 0.0299 | 38.20 | 0.0099 | 0 |
| Pat2 | B2MG_HUMAN | B2M | 0.1323 | 0.0646 | 20.40 | 0.0279 | 0 |
| Pat2 | CO4B_HUMAN | C4B | 0.1089 | 0.0559 | 22.80 | 0.0200 | 0 |
| Pat2 | CO9_HUMAN | C9 | 0.2245 | 0.0126 | 41.50 | 0.0075 | 0 |
| Pat2 | CLUS_HUMAN | CLU | 0.1142 | 0.1131 | 14.00 | 0.0410 | 0 |

| | | | | | | | |
|---|---|---|---|---|---|---|---|
| Pat2 | CNDP1_HUMAN | CNDP1 | 0.1174 | 0.0237 | 41.50 | 0.0091 | 0 |
| Pat2 | CYTC_HUMAN | CST3 | 0.1862 | 0.0585 | 22.00 | 0.0356 | 0 |
| Pat2 | FA12_HUMAN | F12 | 0.6591 | 0.0078 | 41.50 | 0.0101 | 0 |
| Pat2 | VTDB_HUMAN | GC | 0.0278 | 0.0294 | 38.80 | 0.0027 | 0 |
| Pat2 | GELS_HUMAN | GSN | 0.1111 | 0.0548 | 23.20 | 0.0200 | 0 |
| Pat2 | NRP2_HUMAN | NRP2 | 0.1787 | 0.0604 | 21.40 | 0.0353 | 0 |
| Pat2 | RET4_HUMAN | RBP4 | 0.1724 | 0.0397 | 30.00 | 0.0225 | 0 |
| Pat2 | AACT_HUMAN | SERPINA3 | 0.0480 | 0.0560 | 22.80 | 0.0088 | 0 |
| Pat2 | ANT3_HUMAN | SERPINC1 | 0.3265 | 0.0923 | 16.00 | 0.0970 | 0 |
| Pat2 | PEDF_HUMAN | SERPINF1 | 0.2276 | 0.1695 | 11.60 | 0.1165 | 0 |
| Pat2 | TTHY_HUMAN | TTR | 0.2311 | 0.1637 | 11.60 | 0.1149 | 0 |
| Pat4 | A1BG_HUMAN | A1BG | 0.5596 | 0.0069 | 23.88 | 0.0041 | 0 |
| Pat4 | A2MG_HUMAN | A2M | 0.0576 | 0.0822 | 17.20 | 0.0153 | 0 |
| Pat4 | ALBU_HUMAN | ALB | 0.0938 | 0.0078 | 23.88 | 0.0009 | 0 |
| Pat4 | APOA1_HUMAN | APOA1 | 0.2418 | 0.0107 | 23.88 | 0.0039 | 0 |
| Pat4 | APOA2_HUMAN | APOA2 | 0.0168 | 0.0600 | 21.60 | 0.0033 | 0 |
| Pat4 | APOD_HUMAN | APOD | 0.0244 | 0.0893 | 16.40 | 0.0070 | 0 |
| Pat4 | APOE_HUMAN | APOE | 0.0726 | 0.0440 | 23.88 | 0.0104 | 0 |
| Pat4 | B2MG_HUMAN | B2M | 0.1836 | 0.0700 | 19.20 | 0.0419 | 0 |
| Pat4 | CO3_HUMAN | C3 | 0.0761 | 0.1140 | 14.00 | 0.0275 | 0 |
| Pat4 | CO4B_HUMAN | C4B | 0.0964 | 0.0502 | 23.88 | 0.0159 | 0 |
| Pat4 | CO9_HUMAN | C9 | 0.5663 | 0.0085 | 23.88 | 0.0060 | 0 |
| Pat4 | CFAB_HUMAN | CFB | 0.0810 | 0.0803 | 17.40 | 0.0211 | 0 |
| Pat4 | CLUS_HUMAN | CLU | 0.0688 | 0.1342 | 12.80 | 0.0288 | 0 |
| Pat4 | CYTC_HUMAN | CST3 | 0.1575 | 0.0574 | 22.40 | 0.0296 | 0 |
| Pat4 | FA12_HUMAN | F12 | 0.5535 | 0.0071 | 23.88 | 0.0042 | 0 |
| Pat4 | VTDB_HUMAN | GC | 0.5269 | 0.0047 | 23.88 | 0.0018 | 0 |
| Pat4 | GELS_HUMAN | GSN | 0.0561 | 0.0745 | 18.40 | 0.0136 | 0 |
| Pat4 | NRP2_HUMAN | NRP2 | 0.1687 | 0.0537 | 23.40 | 0.0297 | 0 |
| Pat4 | RET4_HUMAN | RBP4 | 0.1660 | 0.0326 | 23.88 | 0.0162 | 0 |
| Pat4 | AACT_HUMAN | SERPINA3 | 0.0307 | 0.0748 | 18.40 | 0.0075 | 0 |
| Pat4 | PEDF_HUMAN | SERPINF1 | 0.1099 | 0.1648 | 11.60 | 0.0550 | 0 |
| Pat4 | TTHY_HUMAN | TTR | 0.1210 | 0.1826 | 11.20 | 0.0659 | 0 |
