## Supplementary material for "*In Vivo* Large Scale Mapping Of Protein Turnover In The Human Cerebrospinal Fluid": GTEx expression

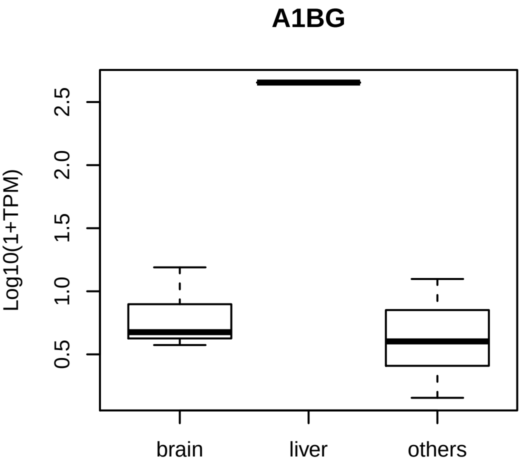

A1BG
Log10(1+TPM)
0.5
1.0
1.5
2.0
2.5
brain
liver
others

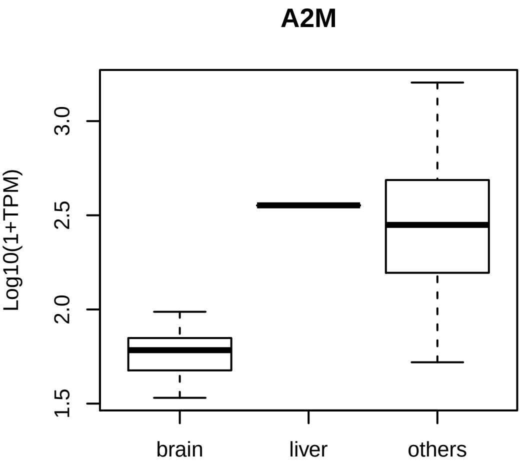
A2M
Log10(1+TPM)
1.5
2.0
2.5
3.0
brain
liver
others

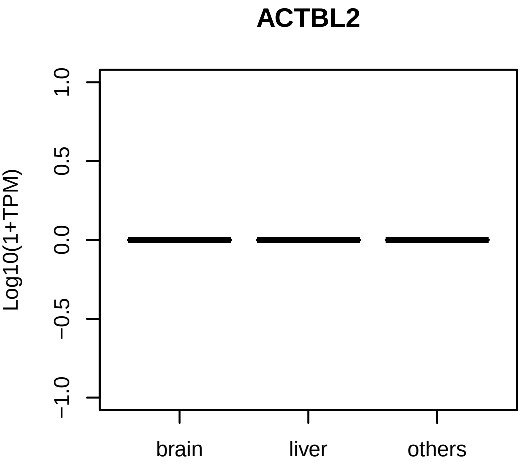

ACTBL2
Log10(1+TPM)
1.0
0.5
0.0
−0.5
−1.0
brain
liver
others

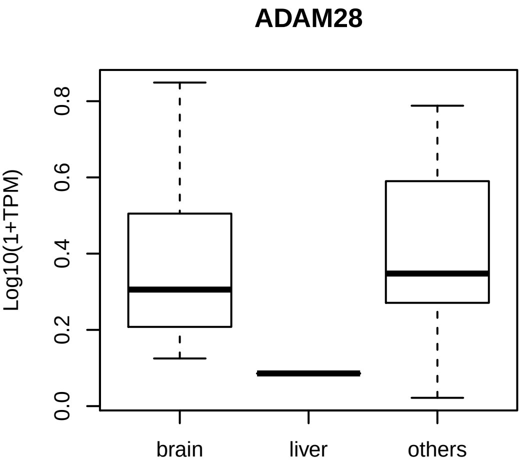
ADAM28
Log10(1+TPM)
0.0
0.2
0.4
0.6
0.8
brain
liver
others

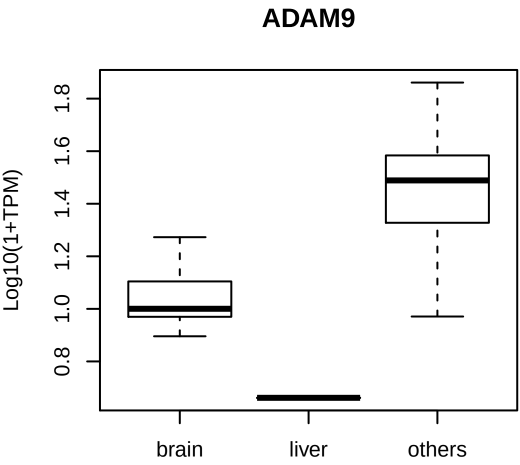
ADAM9
Log10(1+TPM)
0.8
1.0
1.2
1.4
1.6
1.8
brain
liver
others

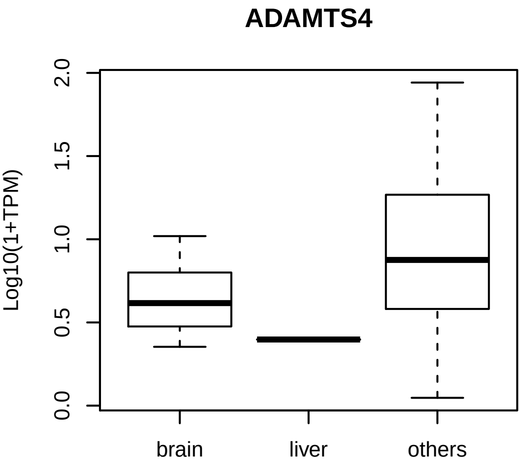
ADAMTS4
Log10(1+TPM)
0.0
0.5
1.0
1.5
2.0
brain
liver
others

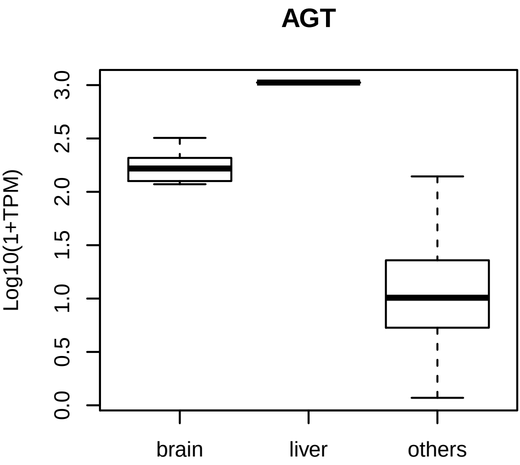
AGT
Log10(1+TPM)
0.0
0.5
1.0
1.5
2.0
2.5
3.0
brain
liver
others

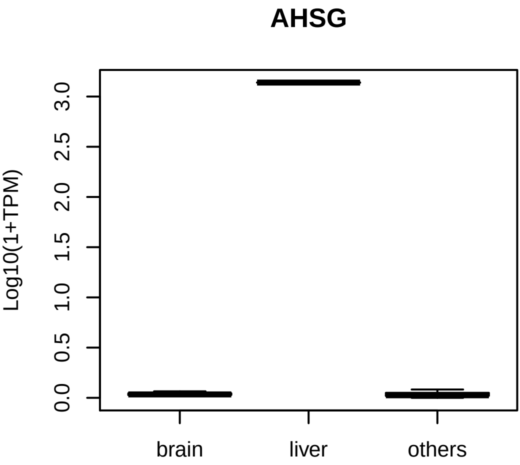
AHSG
Log10(1+TPM)
0.0
0.5
1.0
1.5
2.0
2.5
3.0
brain
liver
others

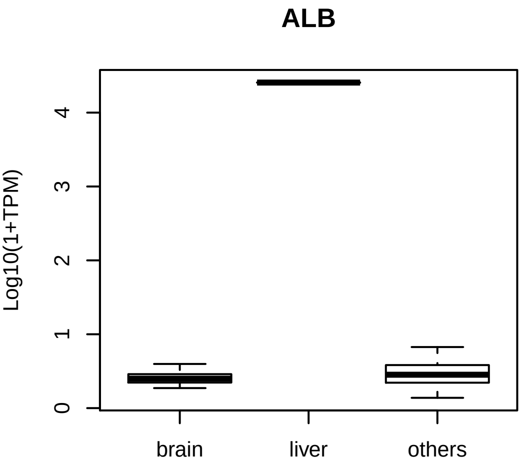

ALB
Log10(1+TPM)
0
1
2
3
4
brain
liver
others

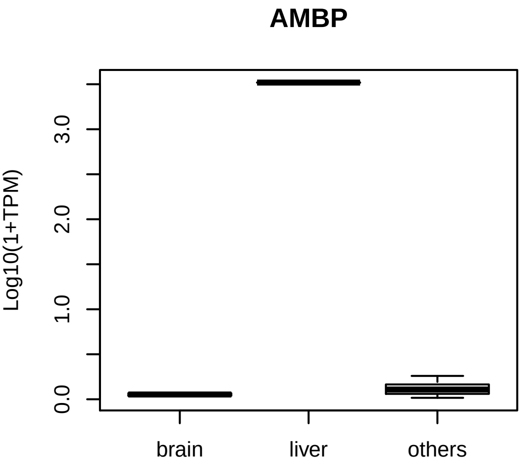
AMBP
Log10(1+TPM)
0.0
1.0
2.0
3.0
brain
liver
others

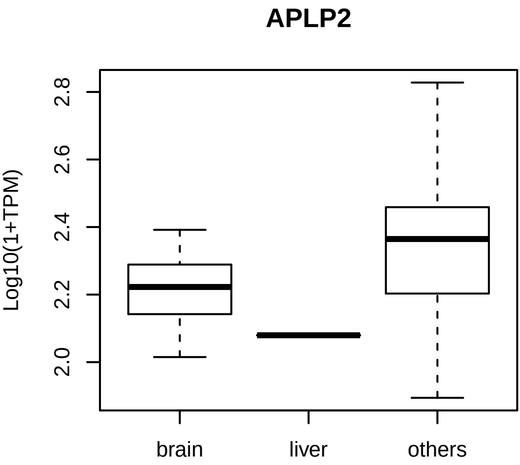
APLP2
Log10(1+TPM)
2.0
2.2
2.4
2.6
2.8
brain
liver
others

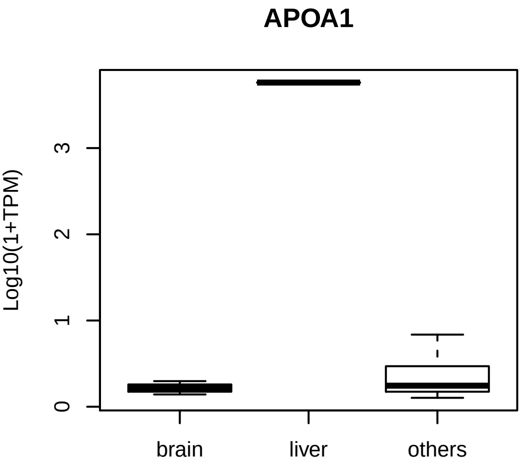
APOA1
Log10(1+TPM)
0
1
2
3
brain
liver
others

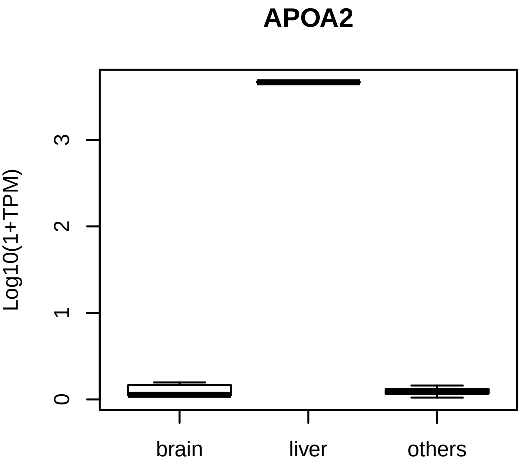
APOA2
Log10(1+TPM)
0
1
2
3
brain
liver
others

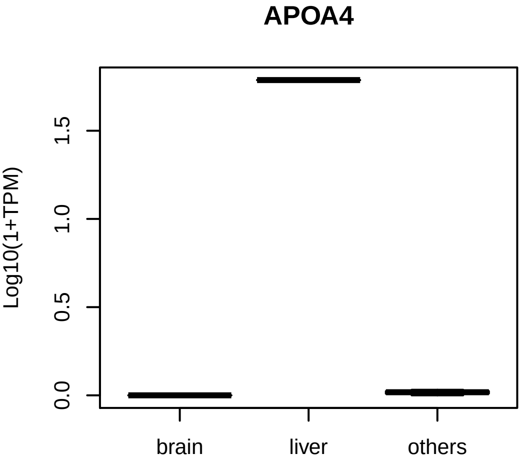
APOA4
Log10(1+TPM)
0.0
0.5
1.0
1.5
brain
liver
others

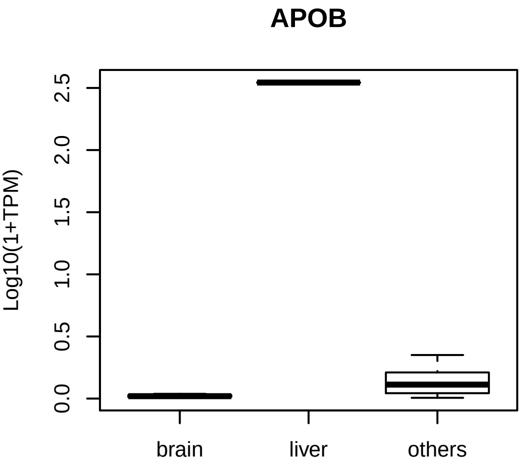
APOB
Log10(1+TPM)
0.0
0.5
1.0
1.5
2.0
2.5
brain
liver
others

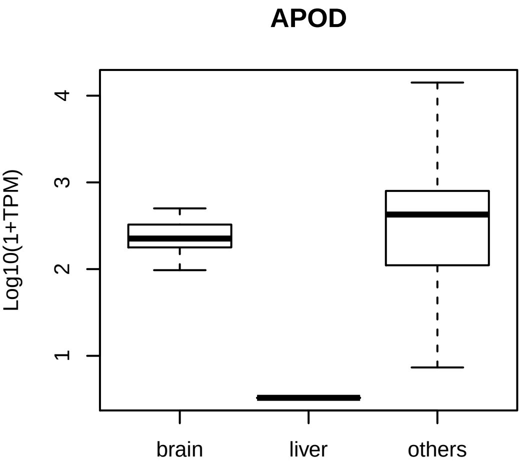
APOD
Log10(1+TPM)
1
2
3
4
brain
liver
others

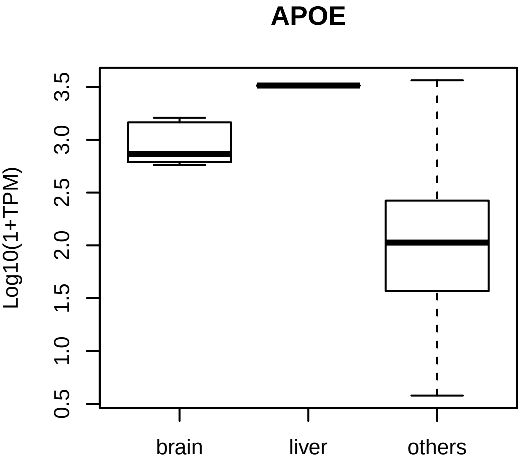

APOE
Log10(1+TPM)
0.5
1.0
1.5
2.0
2.5
3.0
3.5
brain
liver
others

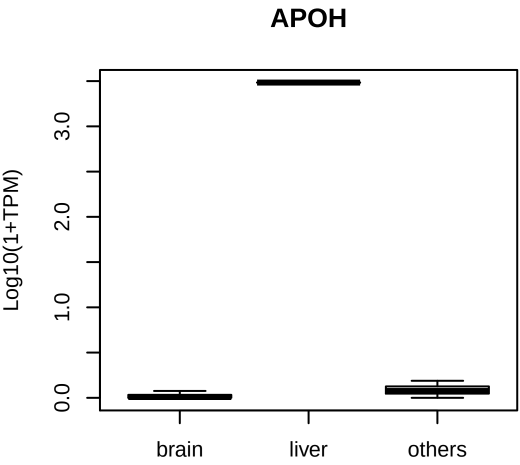
APOH
Log10(1+TPM)
0.0
1.0
2.0
3.0
brain
liver
others

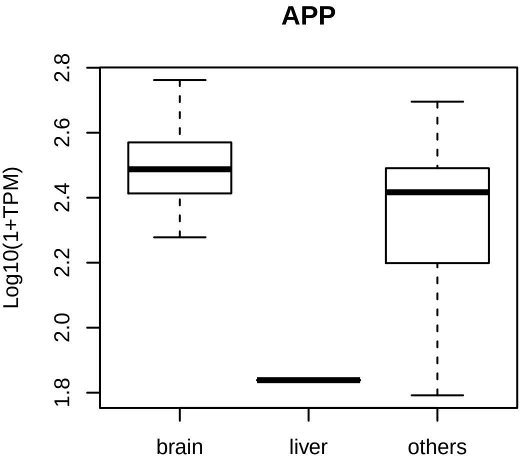
APP
Log10(1+TPM)
2.8
2.6
2.4
2.2
2.0
1.8
brain
liver
others

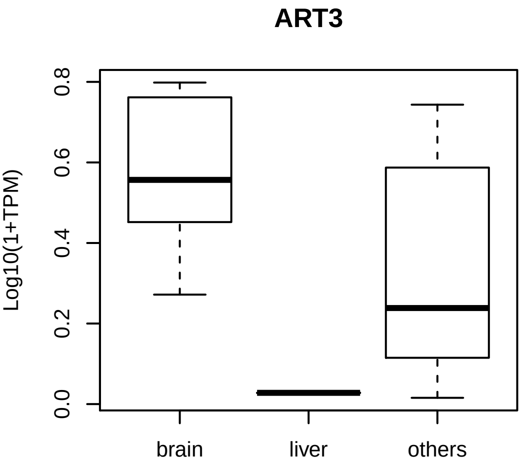
ART3
Log10(1+TPM)
0.0
0.2
0.4
0.6
0.8
brain
liver
others

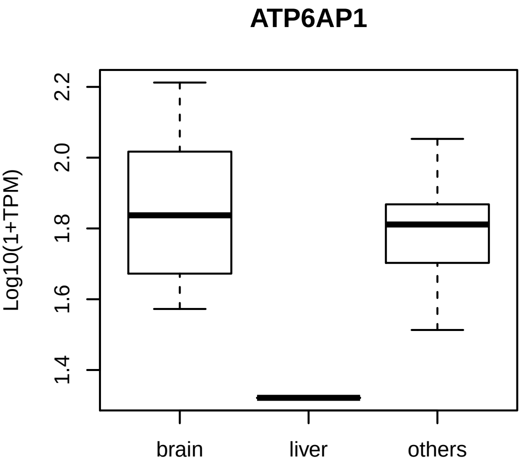
ATP6AP1
Log10(1+TPM)
2.2
2.0
1.8
1.6
1.4
brain
liver
others

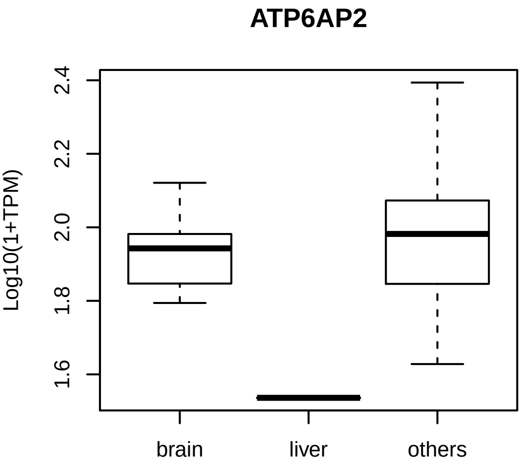
ATP6AP2
Log10(1+TPM)
2.4
2.2
2.0
1.8
1.6
brain
liver
others

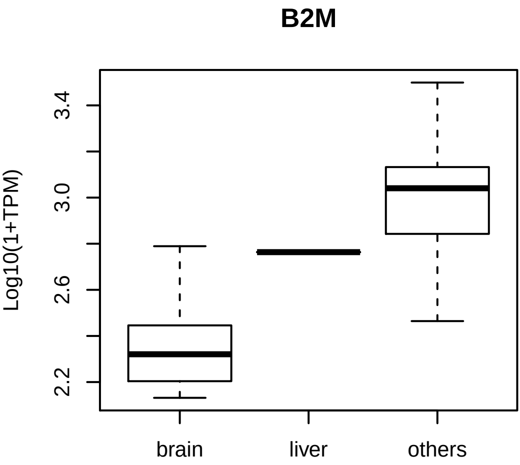
B2M
Log10(1+TPM)
2.2
2.6
3.0
3.4
brain
liver
others

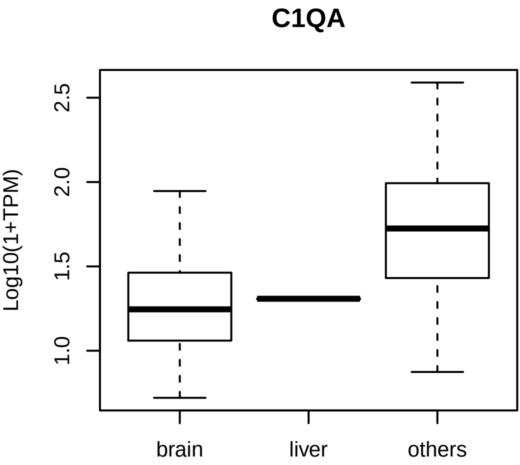
C1QA
Log10(1+TPM)
1.0
1.5
2.0
2.5
brain
liver
others

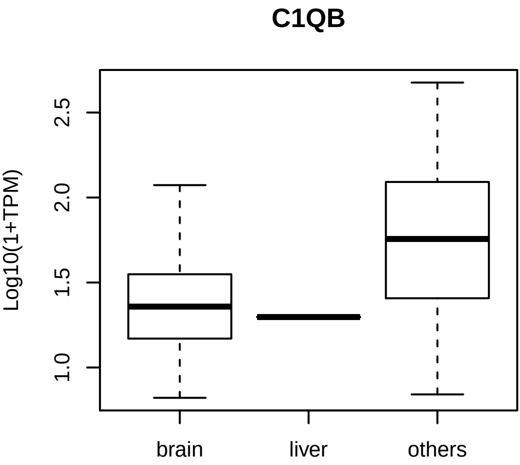
C1QB
Log10(1+TPM)
1.0
1.5
2.0
2.5
brain
liver
others

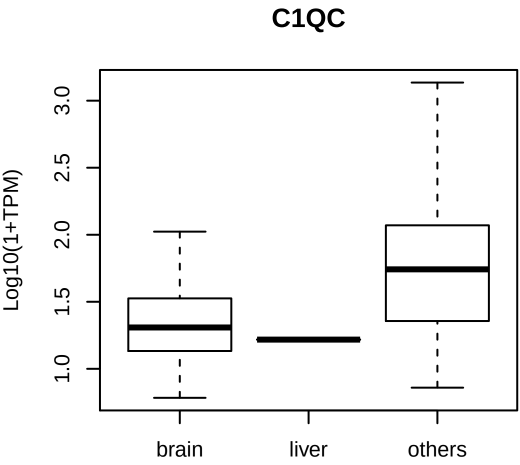
C1QC
Log10(1+TPM)
1.0
1.5
2.0
2.5
3.0
brain
liver
others

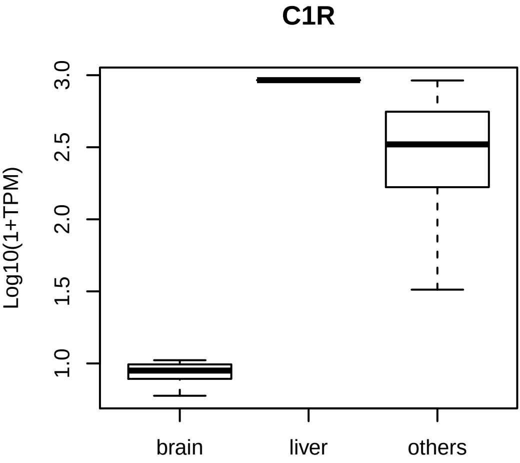
C1R
Log10(1+TPM)
3.0
2.5
2.0
1.5
1.0
brain
liver
others

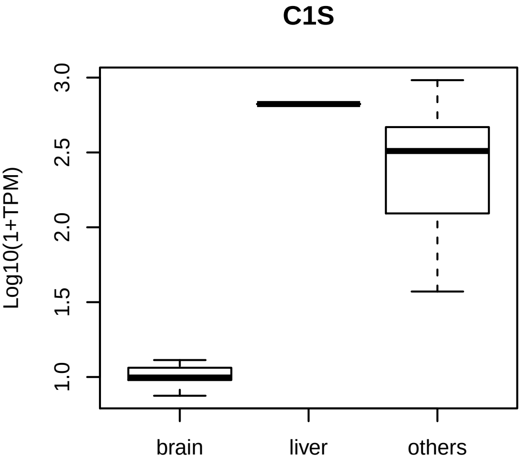

C1S
Log10(1+TPM)
3.0
2.5
2.0
1.5
1.0
brain
liver
others

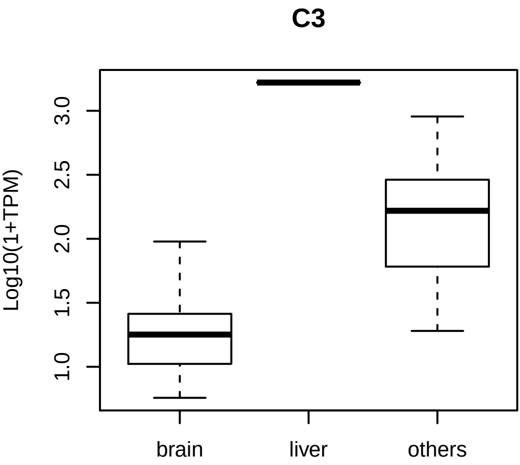
C3
Log10(1+TPM)
1.0
1.5
2.0
2.5
3.0
brain
liver
others

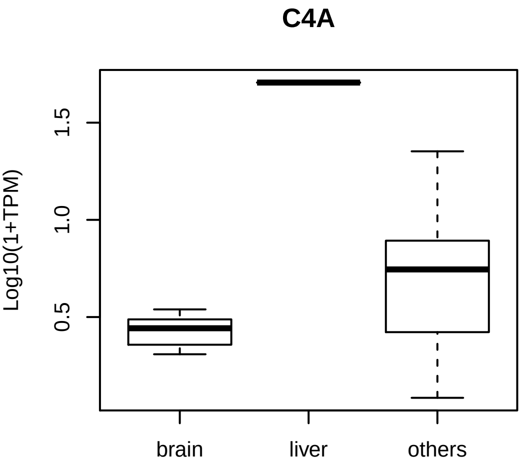
C4A
Log10(1+TPM)
0.5
1.0
1.5
brain
liver
others

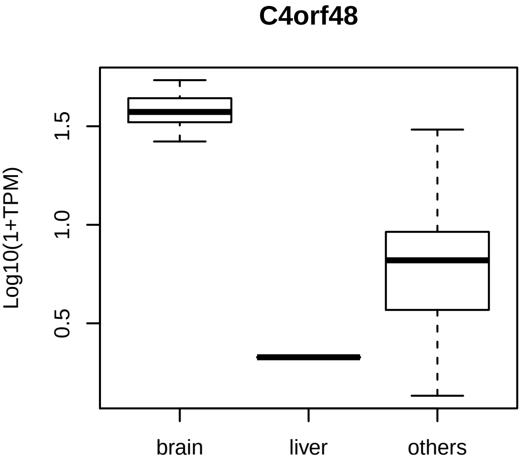
C4orf48
Log10(1+TPM)
1.5
1.0
0.5
brain
liver
others

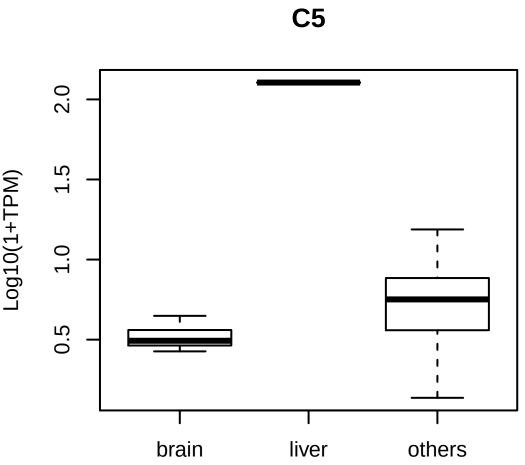
C5
Log10(1+TPM)
0.5
1.0
1.5
2.0
brain
liver
others

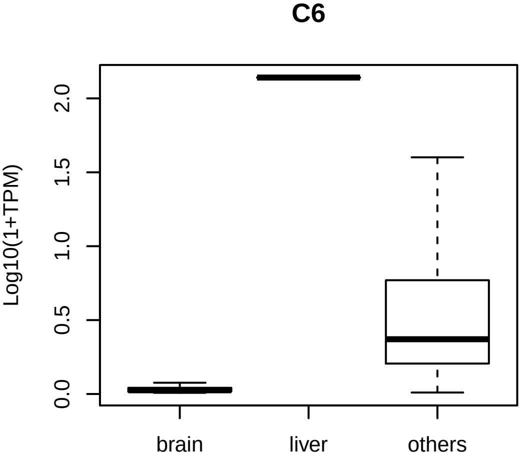

C6
Log10(1+TPM)
0.0
0.5
1.0
1.5
2.0
brain
liver
others

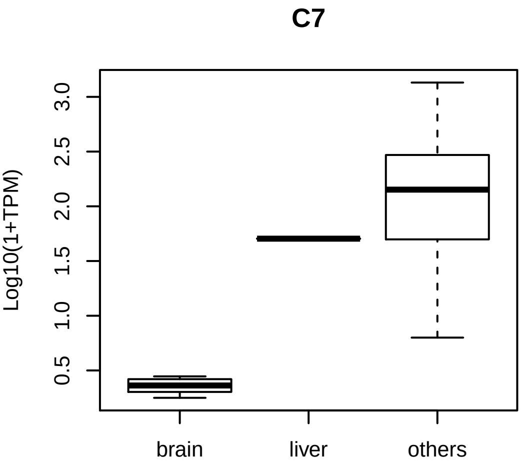
C7
Log10(1+TPM)
0.5
1.0
1.5
2.0
2.5
3.0
brain
liver
others

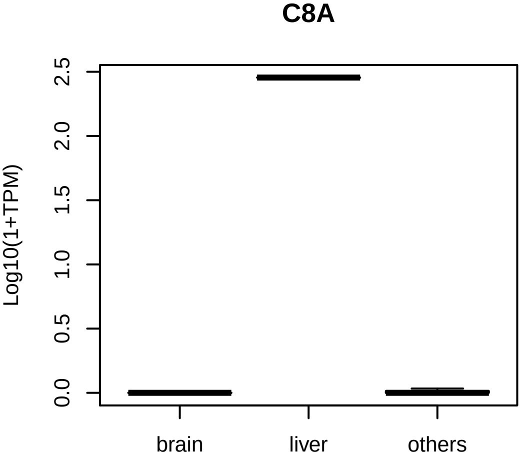
C8A
Log10(1+TPM)
0.0
0.5
1.0
1.5
2.0
2.5
brain
liver
others

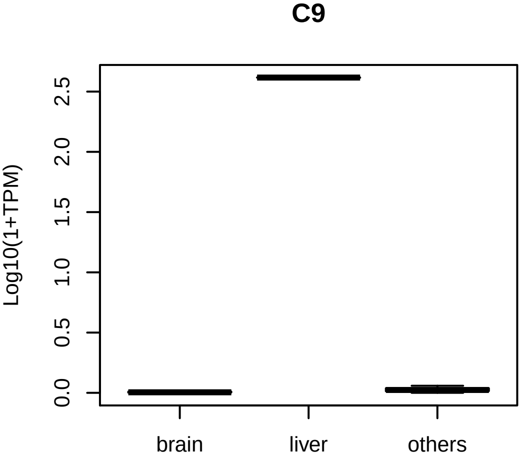

C9
Log10(1+TPM)
0.0
0.5
1.0
1.5
2.0
2.5
brain
liver
others

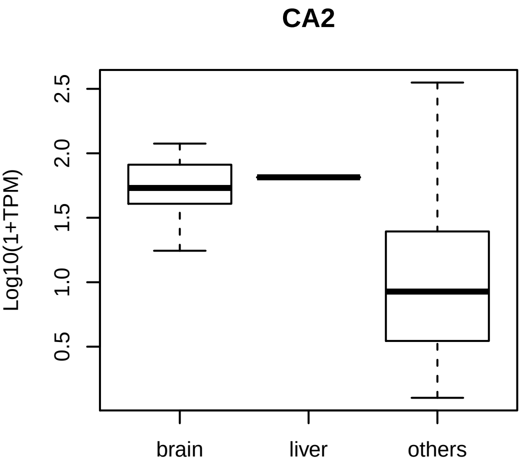
CA2
Log10(1+TPM)
0.5
1.0
1.5
2.0
2.5
brain
liver
others

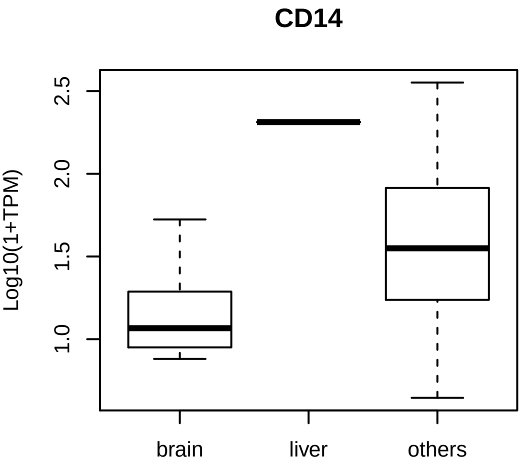
CD14
Log10(1+TPM)
1.0
1.5
2.0
2.5
brain
liver
others

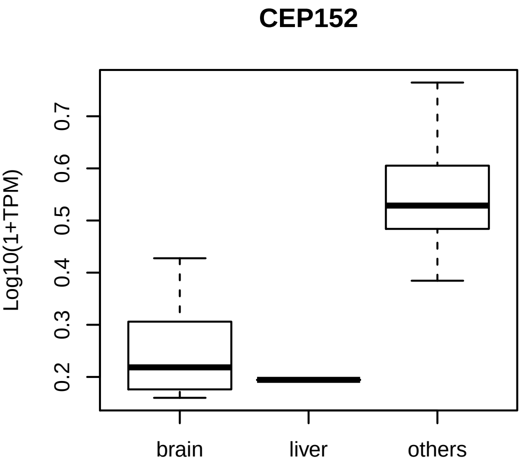
CEP152
Log10(1+TPM)
0.2
0.3
0.4
0.5
0.6
0.7
brain
liver
others

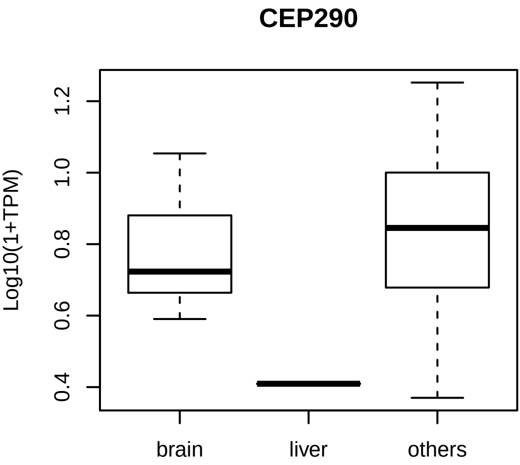

CEP290
Log10(1+TPM)
0.4
0.6
0.8
1.0
1.2
brain
liver
others

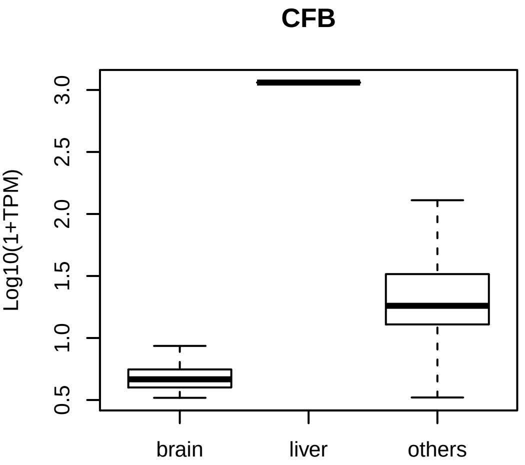
CFB
Log10(1+TPM)
0.5
1.0
1.5
2.0
2.5
3.0
brain
liver
others

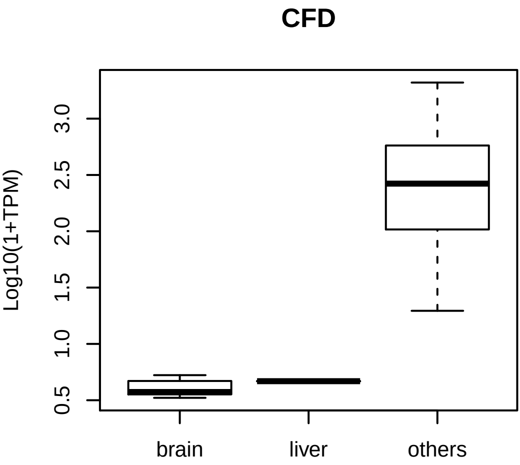
CFD
Log10(1+TPM)
0.5
1.0
1.5
2.0
2.5
3.0
brain
liver
others

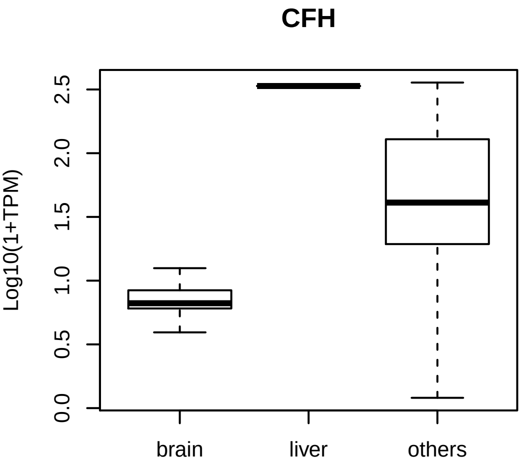

CFH
Log10(1+TPM)
0.0
0.5
1.0
1.5
2.0
2.5
brain
liver
others

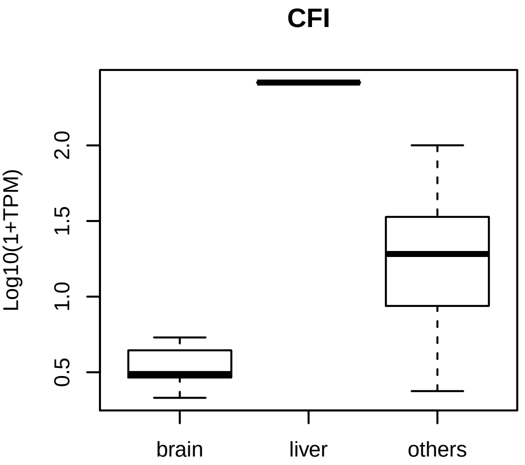
CFI
Log10(1+TPM)
0.5
1.0
1.5
2.0
brain
liver
others

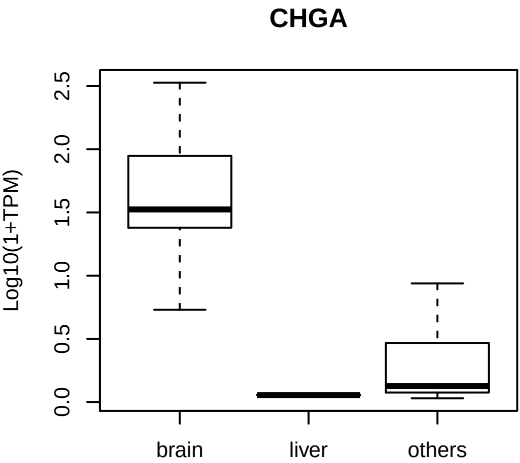
CHGA
Log10(1+TPM)
0.0
0.5
1.0
1.5
2.0
2.5
brain
liver
others

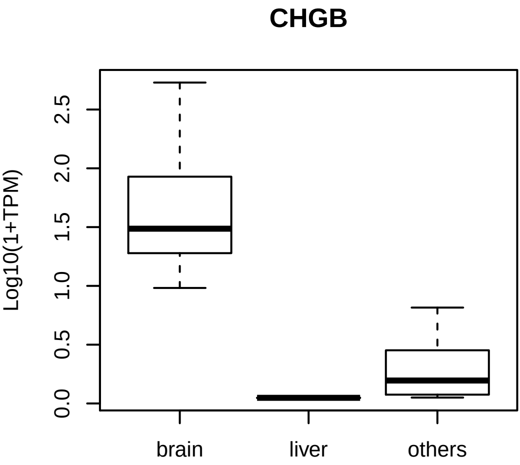
CHGB
Log10(1+TPM)
0.0
0.5
1.0
1.5
2.0
2.5
brain
liver
others

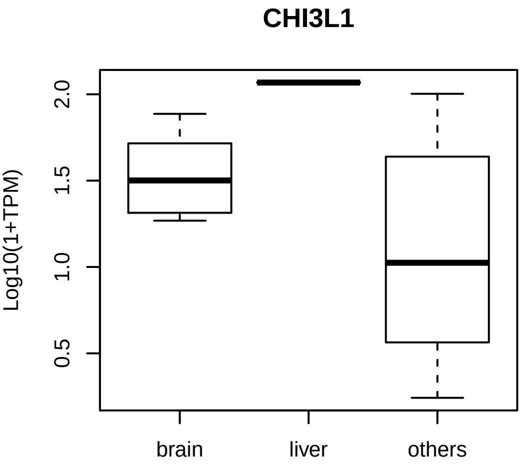
CHI3L1
Log10(1+TPM)
2.0
1.5
1.0
0.5
brain
liver
others

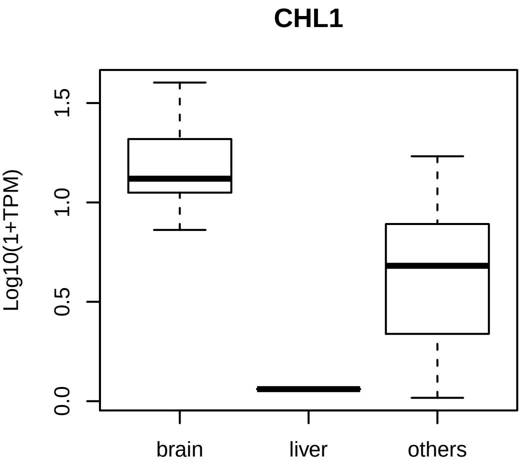
CHL1
Log10(1+TPM)
0.0
0.5
1.0
1.5
brain
liver
others

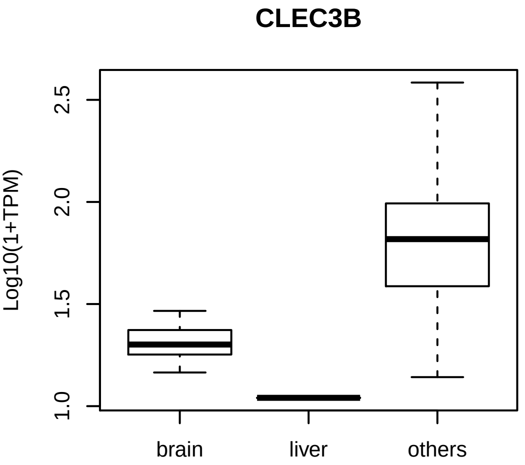
CLEC3B
Log10(1+TPM)
2.5
2.0
1.5
1.0
brain
liver
others

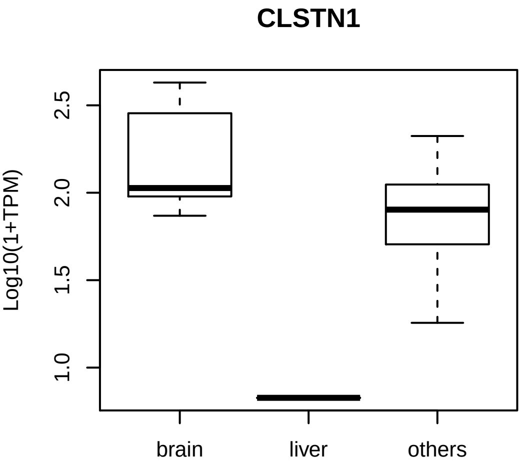

CLSTN1
Log10(1+TPM)
2.5
2.0
1.5
1.0
brain
liver
others

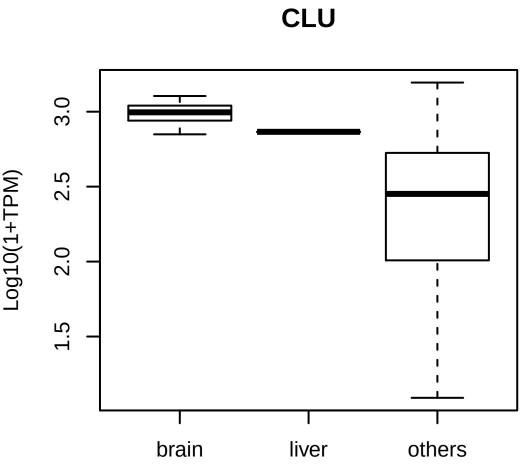

CLU
Log10(1+TPM)
3.0
2.5
2.0
1.5
brain
liver
others

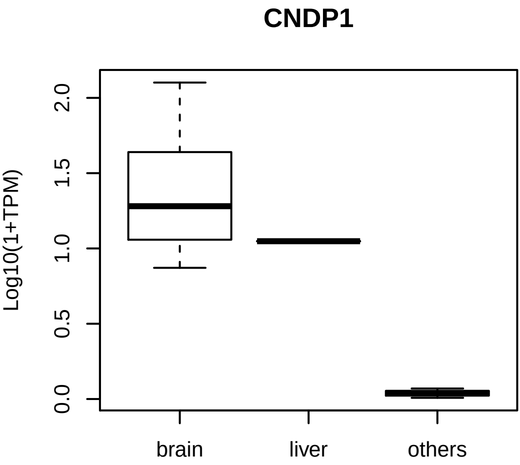
CNDP1
Log10(1+TPM)
0.0
0.5
1.0
1.5
2.0
brain
liver
others

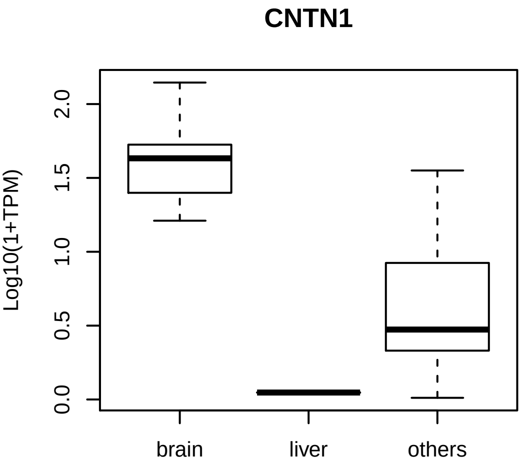

CNTN1
Log10(1+TPM)
0.0
0.5
1.0
1.5
2.0
brain
liver
others

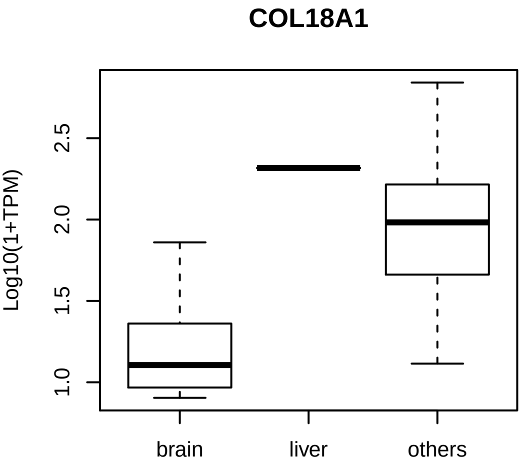
COL18A1
Log10(1+TPM)
1.0
1.5
2.0
2.5
brain
liver
others

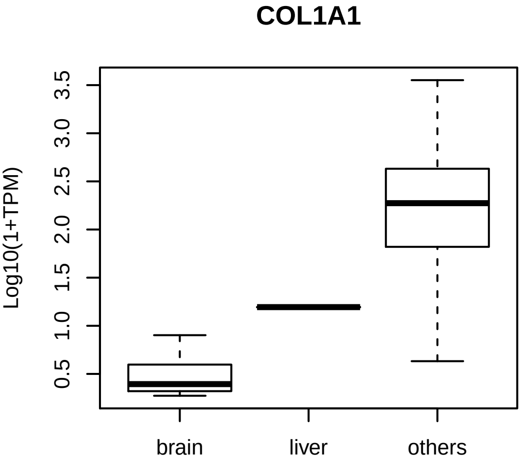

COL1A1
Log10(1+TPM)
0.5
1.0
1.5
2.0
2.5
3.0
3.5
brain
liver
others

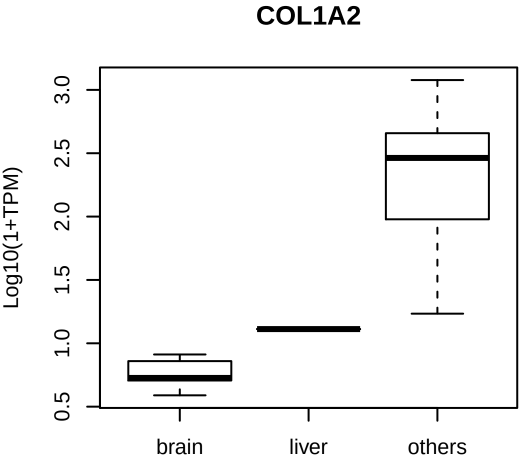
COL1A2
Log10(1+TPM)
0.5
1.0
1.5
2.0
2.5
3.0
brain
liver
others

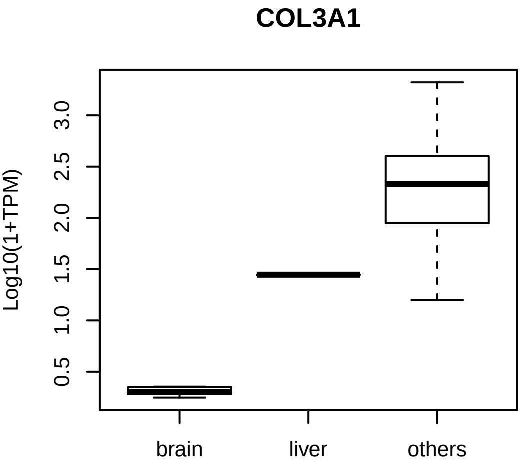
COL3A1
Log10(1+TPM)
0.5
1.0
1.5
2.0
2.5
3.0
brain
liver
others

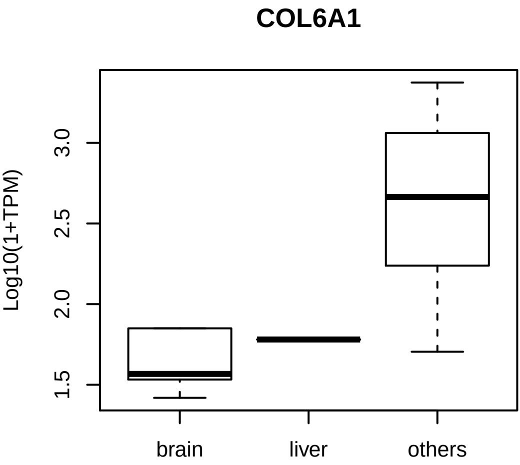
COL6A1
Log10(1+TPM)
1.5
2.0
2.5
3.0
brain
liver
others

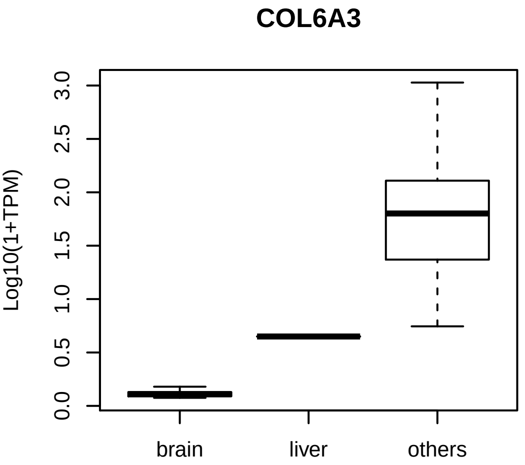
COL6A3
Log10(1+TPM)
0.0
0.5
1.0
1.5
2.0
2.5
3.0
brain
liver
others

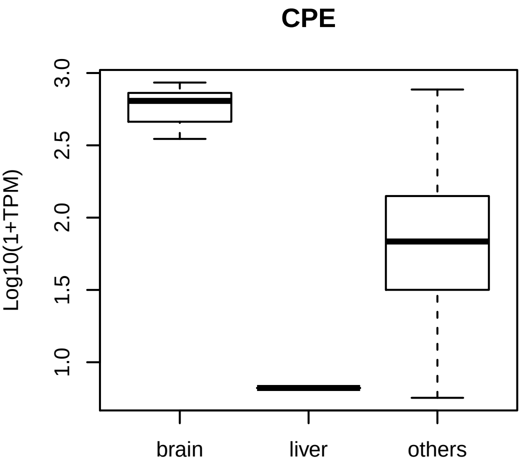

CPE
Log10(1+TPM)
3.0
2.5
2.0
1.5
1.0
brain
liver
others

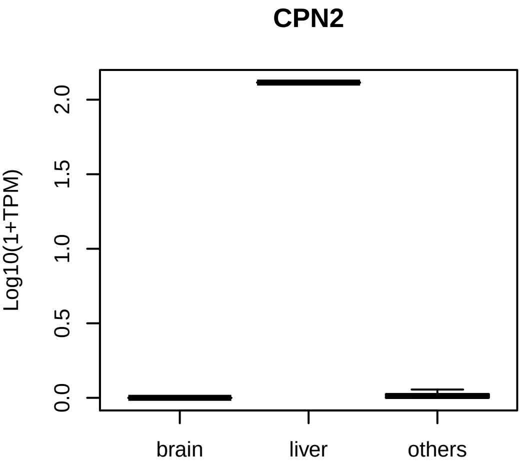
CPN2
Log10(1+TPM)
0.0
0.5
1.0
1.5
2.0
brain
liver
others

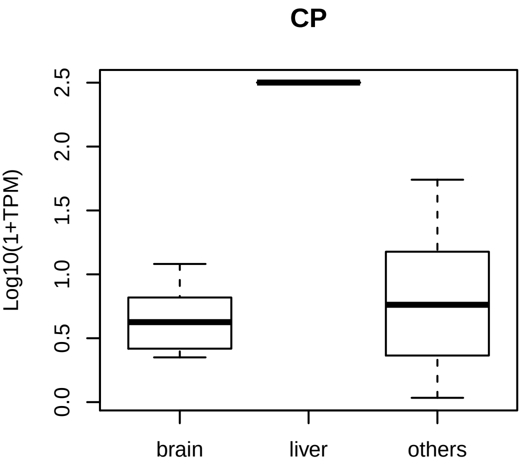
CP
Log10(1+TPM)
0.0
0.5
1.0
1.5
2.0
2.5
brain
liver
others

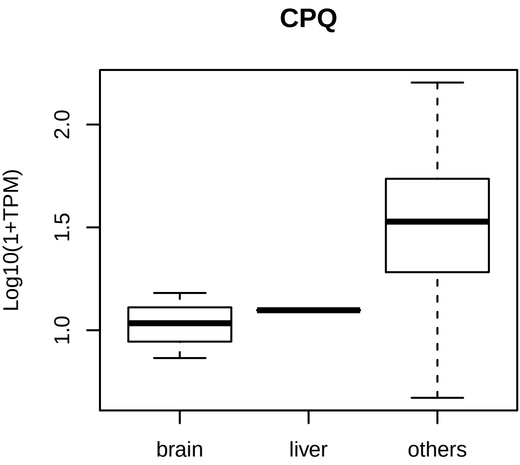
CPQ
Log10(1+TPM)
1.0
1.5
2.0
brain
liver
others

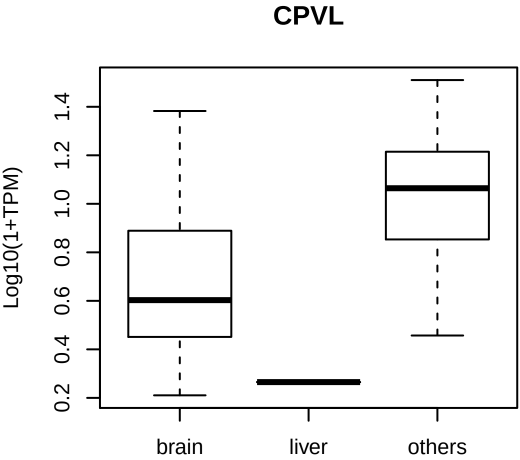
CPVL
Log10(1+TPM)
0.2
0.4
0.6
0.8
1.0
1.2
1.4
brain
liver
others

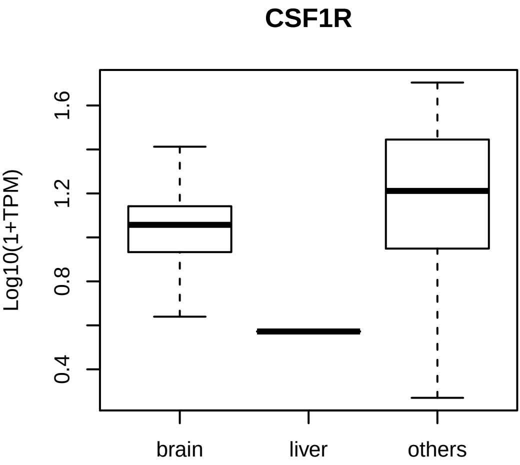

CSF1R
Log10(1+TPM)
1.6
1.2
0.8
0.4
brain
liver
others

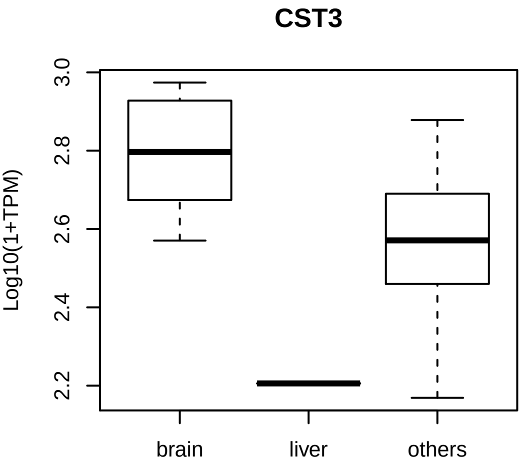
CST3
Log10(1+TPM)
2.2
2.4
2.6
2.8
3.0
brain
liver
others

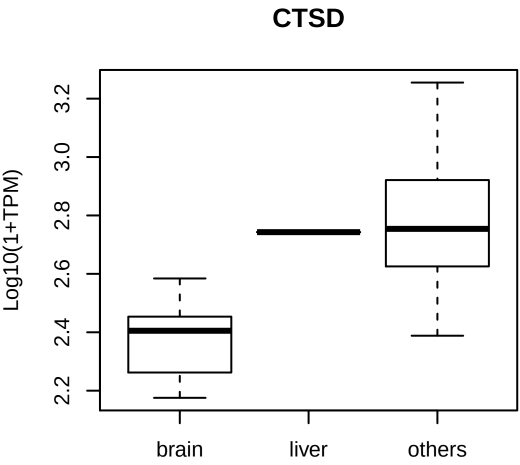
CTSD
Log10(1+TPM)
3.2
3.0
2.8
2.6
2.4
2.2
brain
liver
others

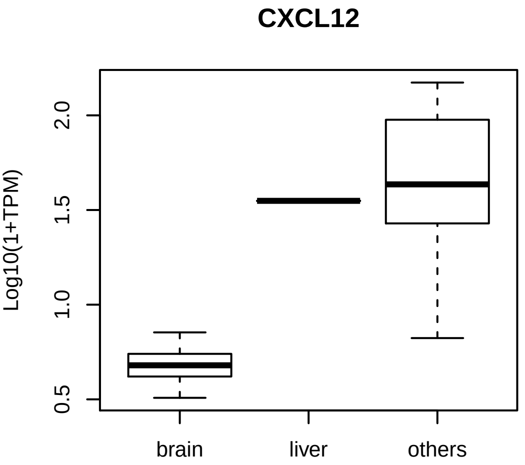
CXCL12
Log10(1+TPM)
0.5
1.0
1.5
2.0
brain
liver
others

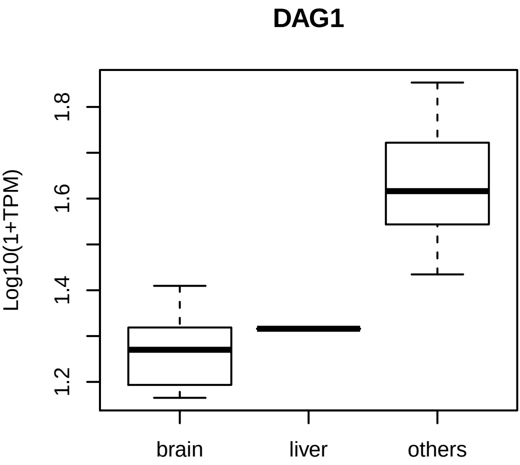
DAG1
Log10(1+TPM)
1.2
1.4
1.6
1.8
brain
liver
others

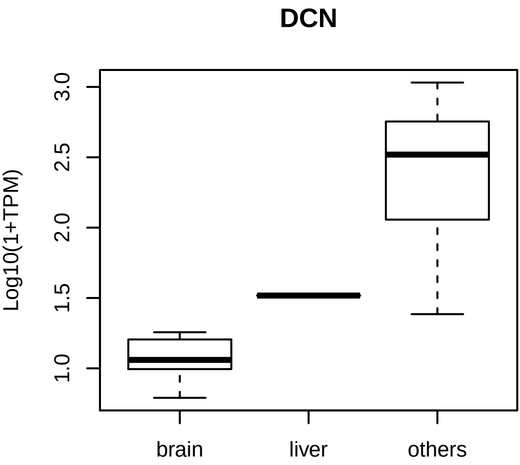
DCN
Log10(1+TPM)
3.0
2.5
2.0
1.5
1.0
brain
liver
others

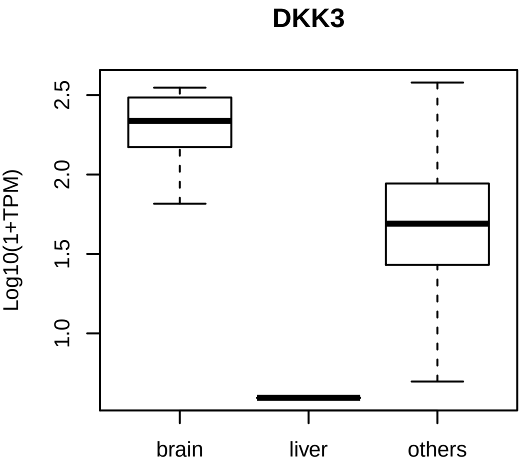
DKK3
Log10(1+TPM)
2.5
2.0
1.5
1.0
brain
liver
others

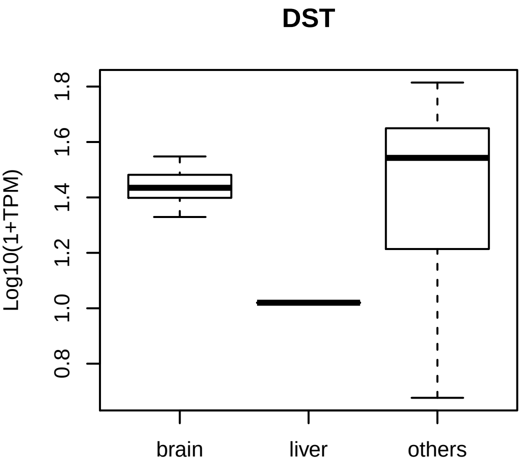
DST
Log10(1+TPM)
0.8
1.0
1.2
1.4
1.6
1.8
brain
liver
others

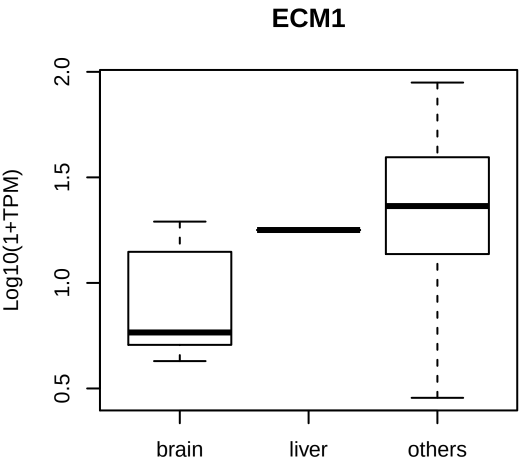
ECM1
Log10(1+TPM)
0.5
1.0
1.5
2.0
brain
liver
others

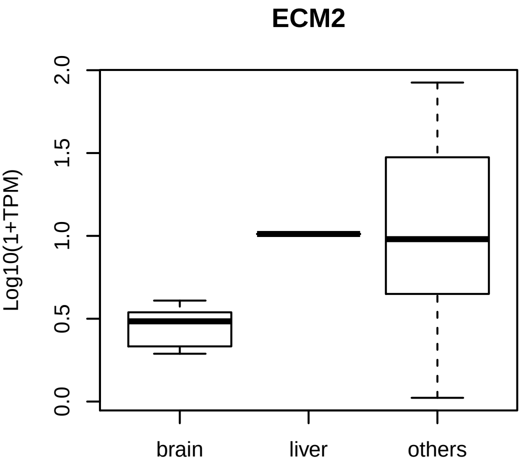
ECM2
Log10(1+TPM)
0.0
0.5
1.0
1.5
2.0
brain
liver
others

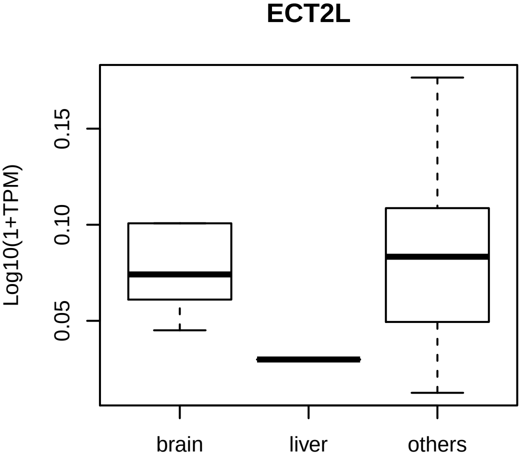
ECT2L
Log10(1+TPM)
0.05
0.10
0.15
brain
liver
others

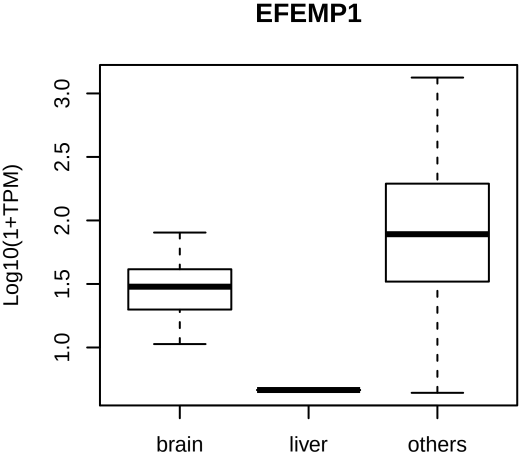
EFEMP1
Log10(1+TPM)
1.0
1.5
2.0
2.5
3.0
brain
liver
others

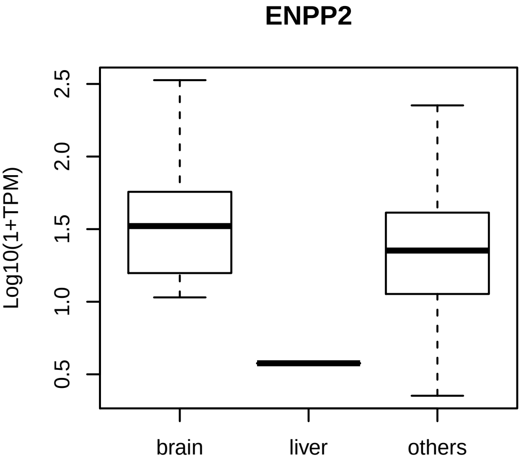
ENPP2
Log10(1+TPM)
0.5
1.0
1.5
2.0
2.5
brain
liver
others

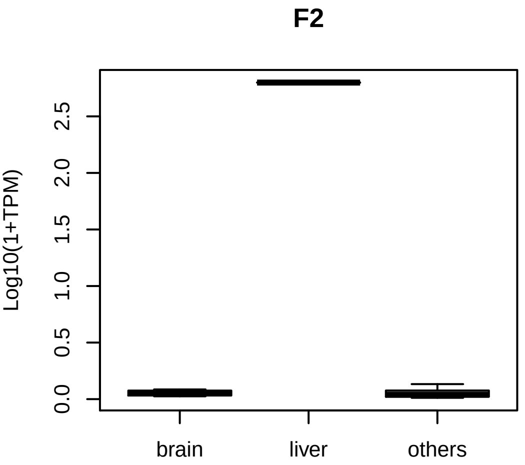
F2
Log10(1+TPM)
0.0
0.5
1.0
1.5
2.0
2.5
brain
liver
others

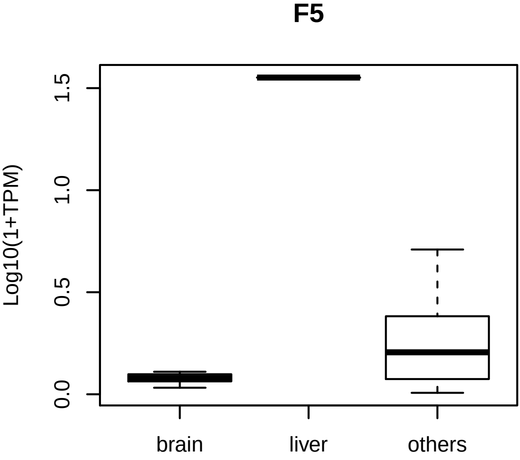
F5
Log10(1+TPM)
0.0
0.5
1.0
1.5
brain
liver
others

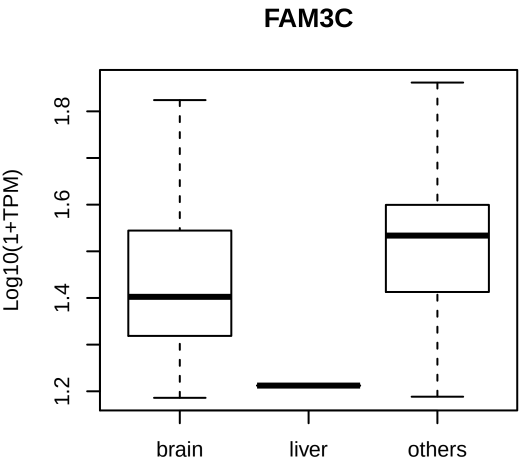
FAM3C
Log10(1+TPM)
1.2
1.4
1.6
1.8
brain
liver
others

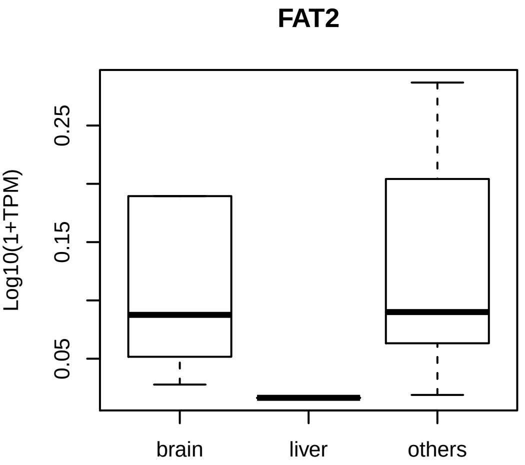
FAT2
Log10(1+TPM)
0.05
0.15
0.25
brain
liver
others

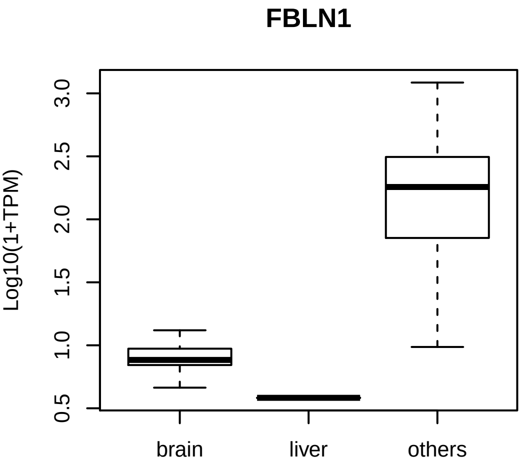
FBLN1
Log10(1+TPM)
0.5
1.0
1.5
2.0
2.5
3.0
brain
liver
others

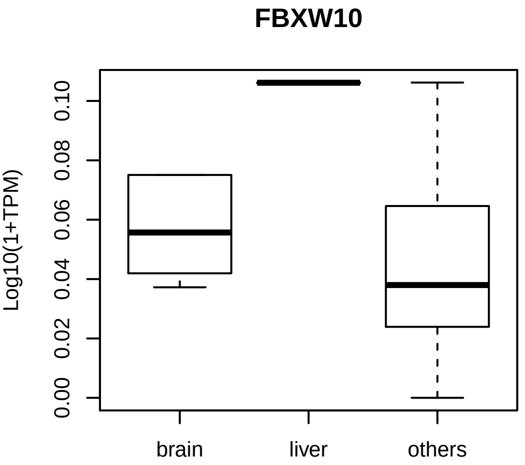
FBXW10
Log10(1+TPM)
0.00
0.02
0.04
0.06
0.08
0.10
brain
liver
others

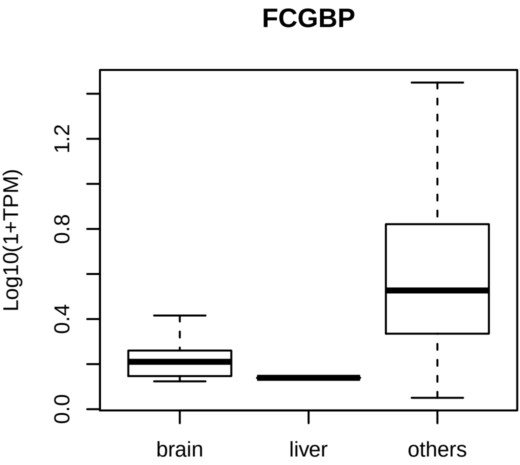
FCGBP
Log10(1+TPM)
0.0
0.4
0.8
1.2
brain
liver
others

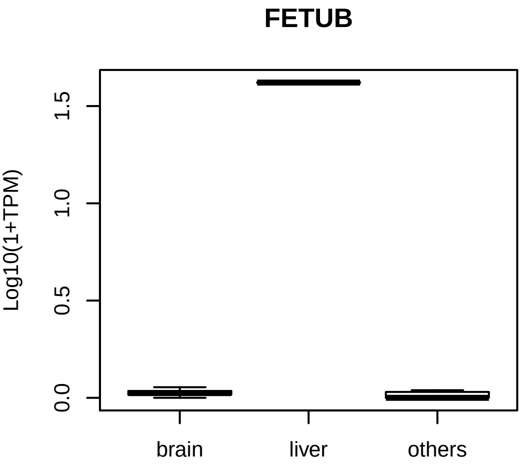
FETUB
Log10(1+TPM)
0.0
0.5
1.0
1.5
brain
liver
others

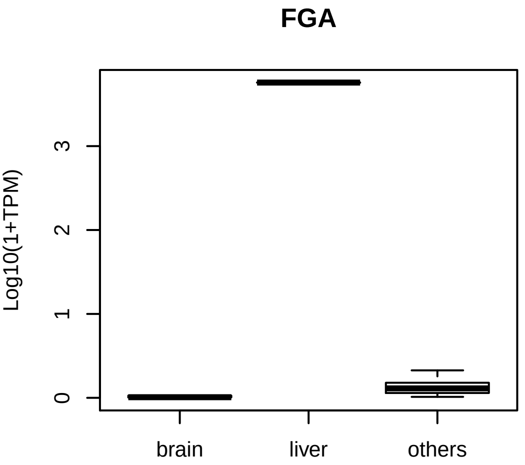
FGA
Log10(1+TPM)
0
1
2
3
brain
liver
others

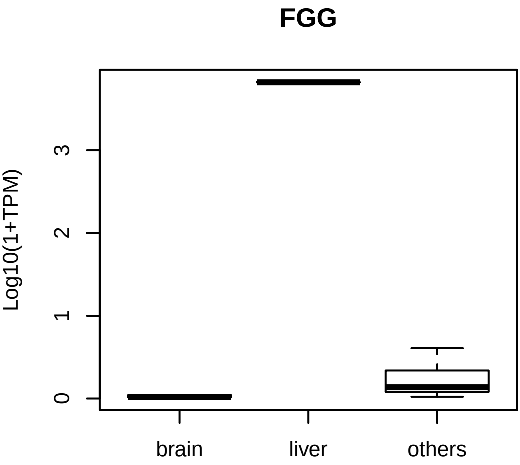
FGG
Log10(1+TPM)
0
1
2
3
brain
liver
others

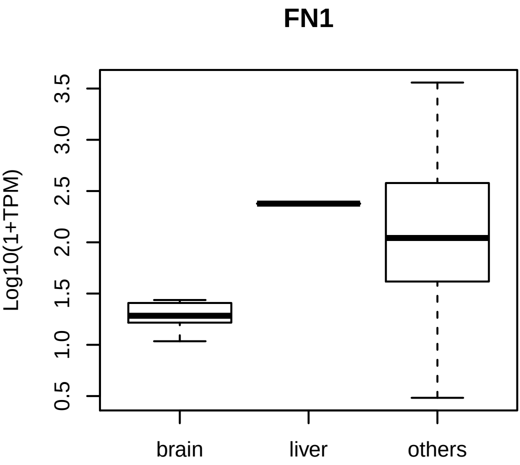

FN1
Log10(1+TPM)
0.5
1.0
1.5
2.0
2.5
3.0
3.5
brain
liver
others

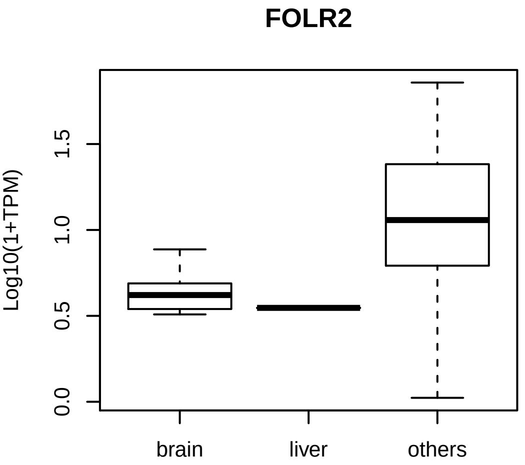
FOLR2
Log10(1+TPM)
0.0
0.5
1.0
1.5
brain
liver
others

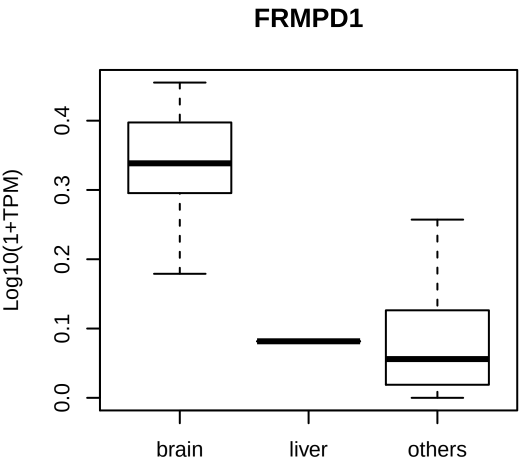
FRMPD1
Log10(1+TPM)
0.0
0.1
0.2
0.3
0.4
brain
liver
others

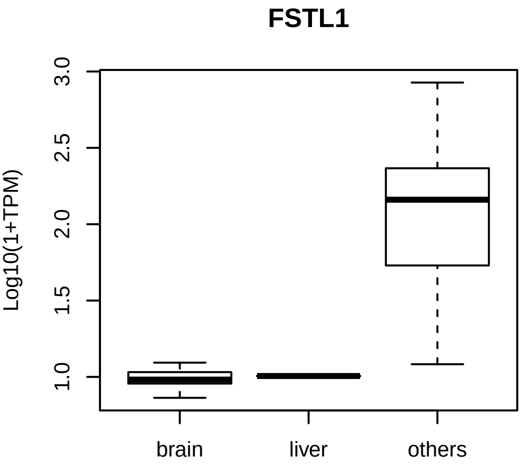
FSTL1
Log10(1+TPM)
3.0
2.5
2.0
1.5
1.0
brain
liver
others

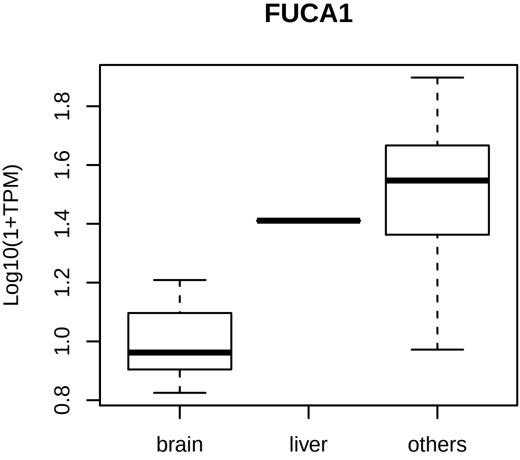
FUCA1
Log10(1+TPM)
0.8
1.0
1.2
1.4
1.6
1.8
brain
liver
others

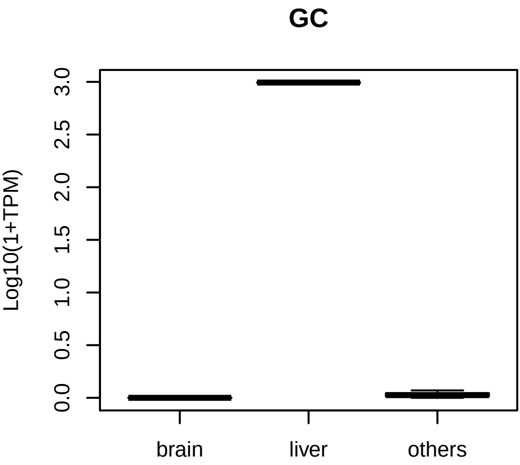
GC
Log10(1+TPM)
0.0
0.5
1.0
1.5
2.0
2.5
3.0
brain
liver
others

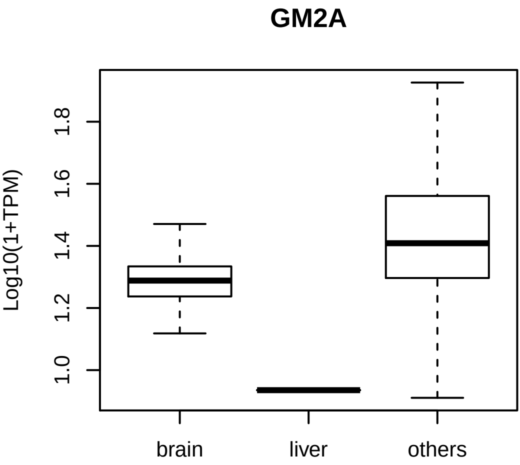
GM2A
Log10(1+TPM)
1.0
1.2
1.4
1.6
1.8
brain
liver
others

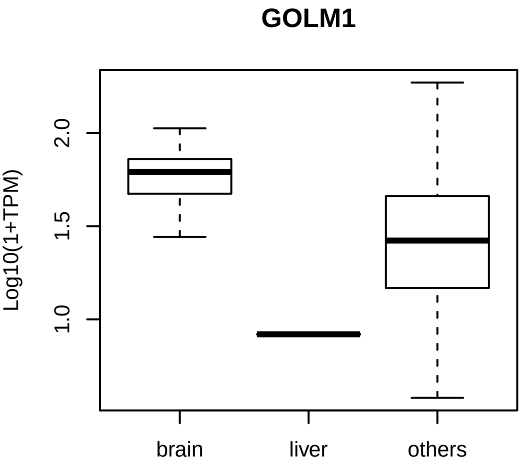

GOLM1
Log10(1+TPM)
2.0
1.5
1.0
brain
liver
others

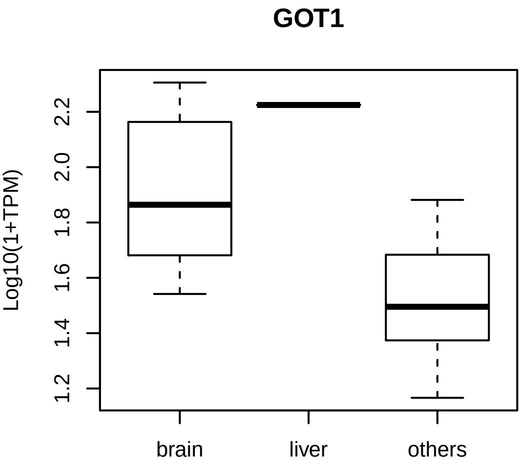
GOT1
Log10(1+TPM)
1.2
1.4
1.6
1.8
2.0
2.2
brain
liver
others

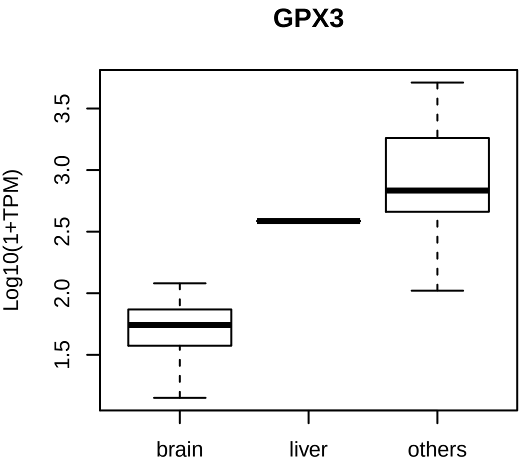
GPX3
Log10(1+TPM)
1.5
2.0
2.5
3.0
3.5
brain
liver
others

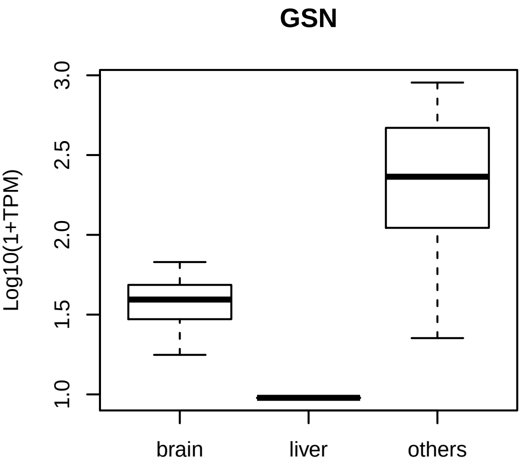
GSN
Log10(1+TPM)
3.0
2.5
2.0
1.5
1.0
brain
liver
others

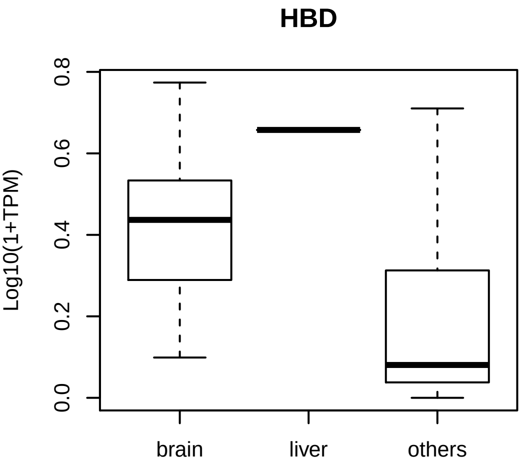
HBD
Log10(1+TPM)
0.0
0.2
0.4
0.6
0.8
brain
liver
others

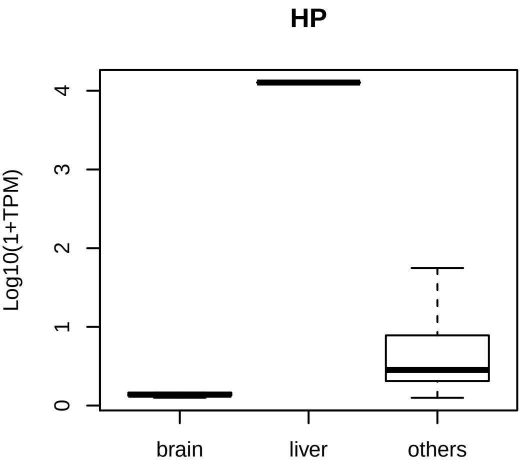
HP
Log10(1+TPM)
0
1
2
3
4
brain
liver
others

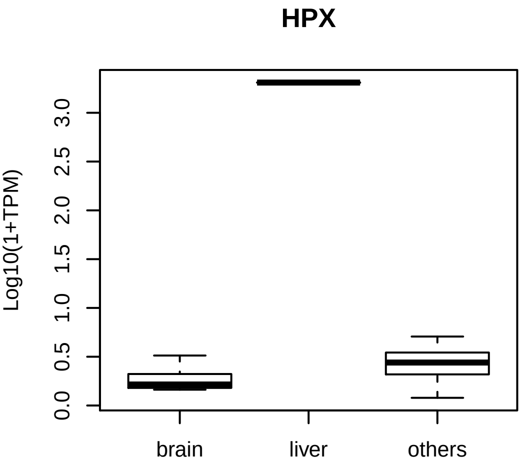

HPX
Log10(1+TPM)
0.0
0.5
1.0
1.5
2.0
2.5
3.0
brain
liver
others

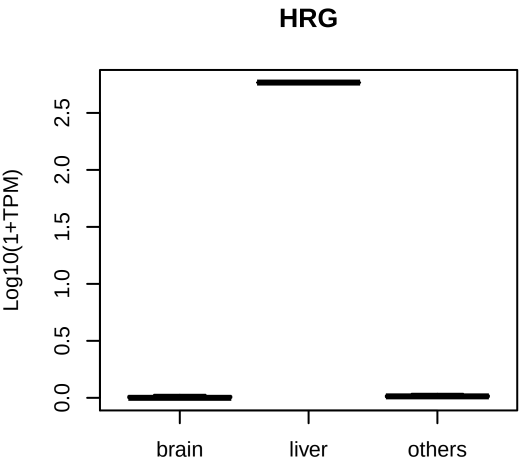
HRG
Log10(1+TPM)
0.0
0.5
1.0
1.5
2.0
2.5
brain
liver
others

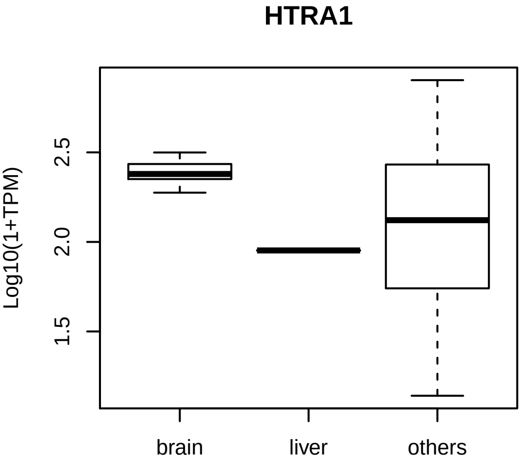

HTRA1
Log10(1+TPM)
1.5
2.0
2.5
brain
liver
others

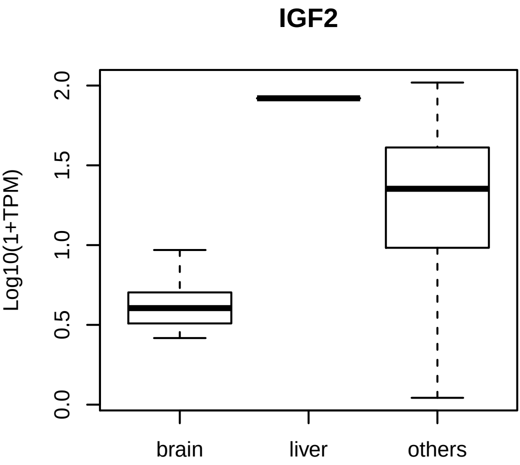
IGF2
Log10(1+TPM)
0.0
0.5
1.0
1.5
2.0
brain
liver
others

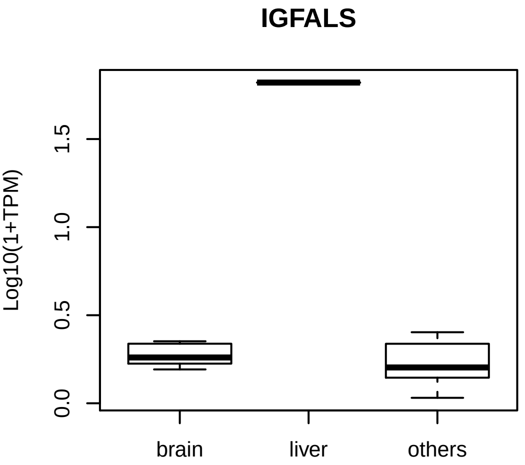
IGFALS
Log10(1+TPM)
0.0
0.5
1.0
1.5
brain
liver
others

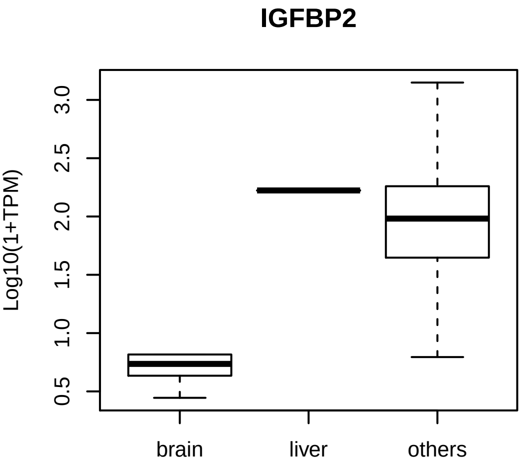
IGFBP2
Log10(1+TPM)
0.5
1.0
1.5
2.0
2.5
3.0
brain
liver
others

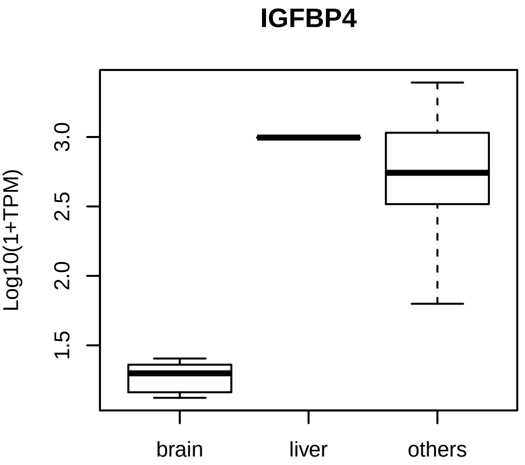
IGFBP4
Log10(1+TPM)
3.0
2.5
2.0
1.5
brain
liver
others

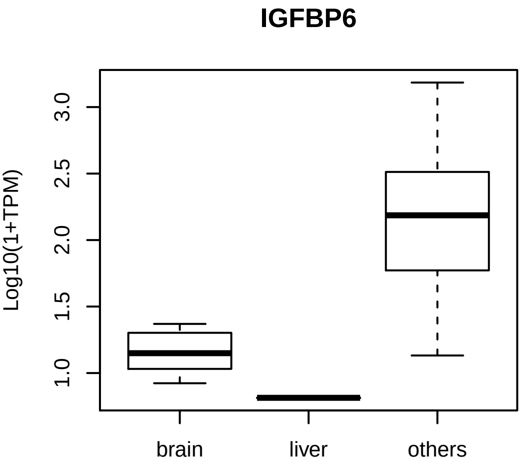
IGFBP6
Log10(1+TPM)
1.0
1.5
2.0
2.5
3.0
brain
liver
others

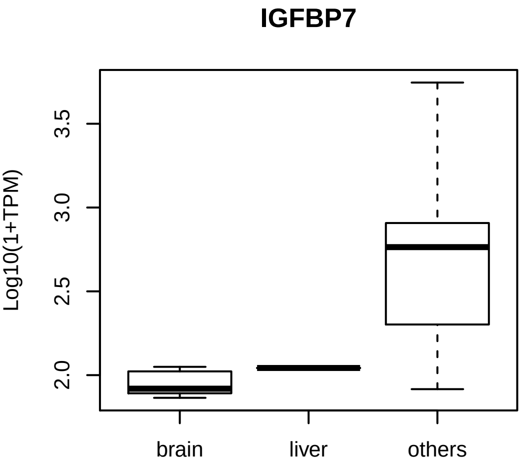
IGFBP7
Log10(1+TPM)
2.0
2.5
3.0
3.5
brain
liver
others

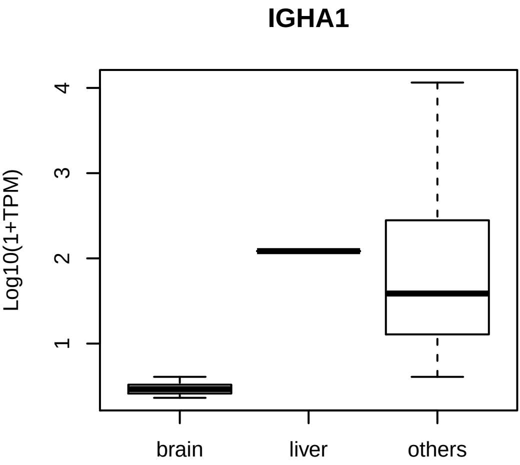
IGHA1
Log10(1+TPM)
1
2
3
4
brain
liver
others

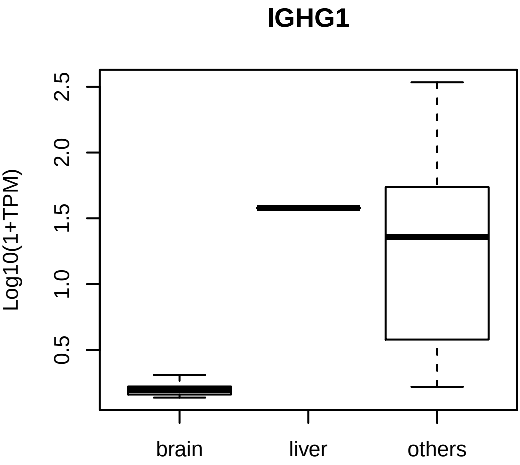
IGHG1
Log10(1+TPM)
0.5
1.0
1.5
2.0
2.5
brain
liver
others

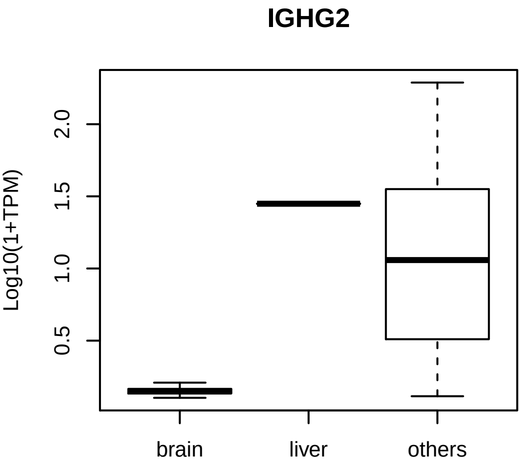
IGHG2
Log10(1+TPM)
2.0
1.5
1.0
0.5
brain
liver
others

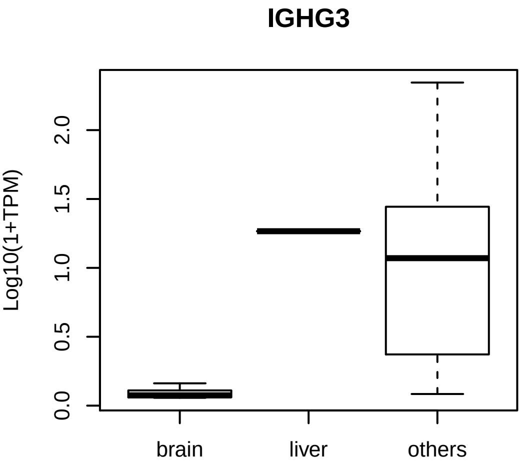
IGHG3
Log10(1+TPM)
0.0
0.5
1.0
1.5
2.0
brain
liver
others

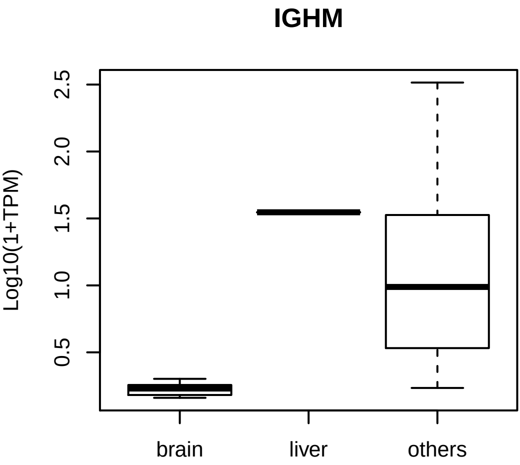
IGHM
Log10(1+TPM)
0.5
1.0
1.5
2.0
2.5
brain
liver
others

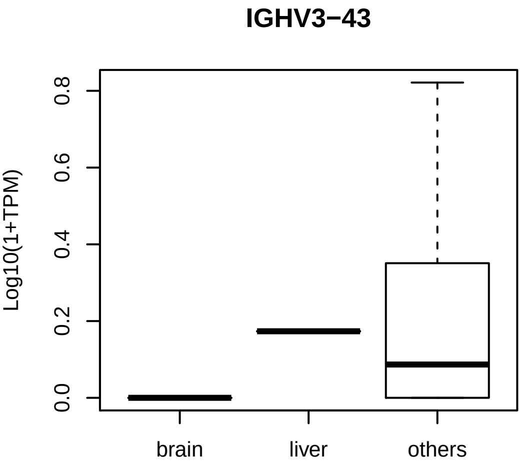
IGHV3−43
Log10(1+TPM)
0.0
0.2
0.4
0.6
0.8
brain
liver
others

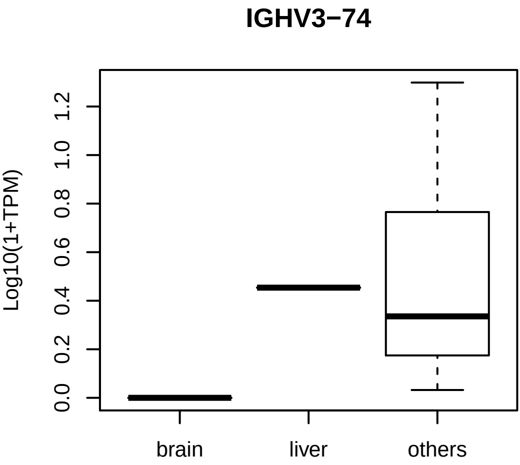
IGHV3−74
Log10(1+TPM)
0.0
0.2
0.4
0.6
0.8
1.0
1.2
brain
liver
others

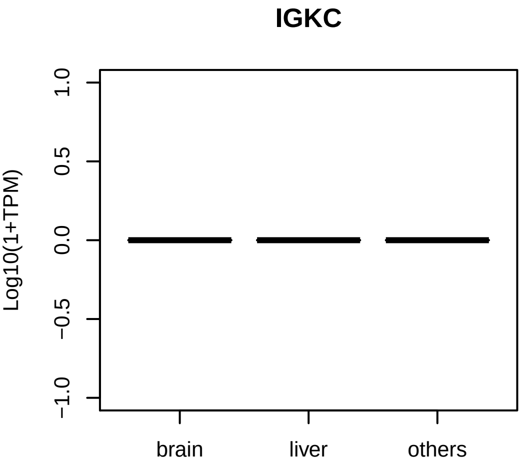

IGKC
Log10(1+TPM)
1.0
0.5
0.0
−0.5
−1.0
brain
liver
others

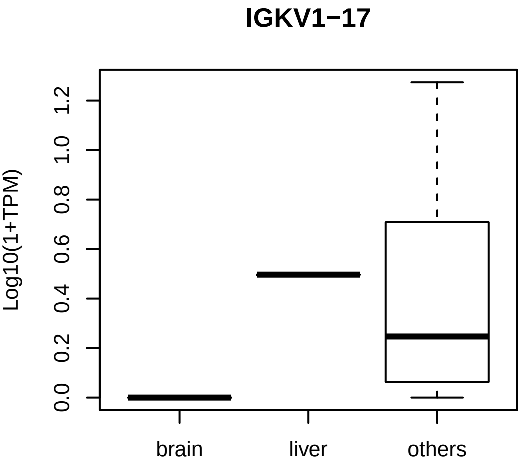
IGKV1−17
Log10(1+TPM)
0.0
0.2
0.4
0.6
0.8
1.0
1.2
brain
liver
others

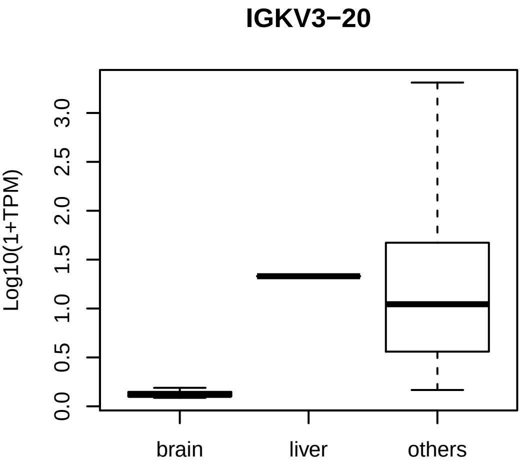
IGKV3−20
Log10(1+TPM)
0.0
0.5
1.0
1.5
2.0
2.5
3.0
brain
liver
others

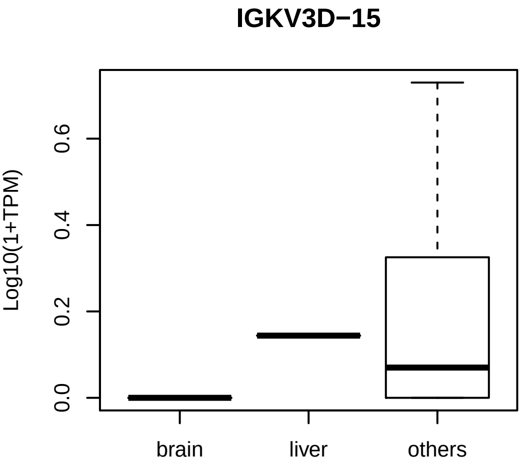
IGKV3D−15
Log10(1+TPM)
0.0
0.2
0.4
0.6
brain
liver
others

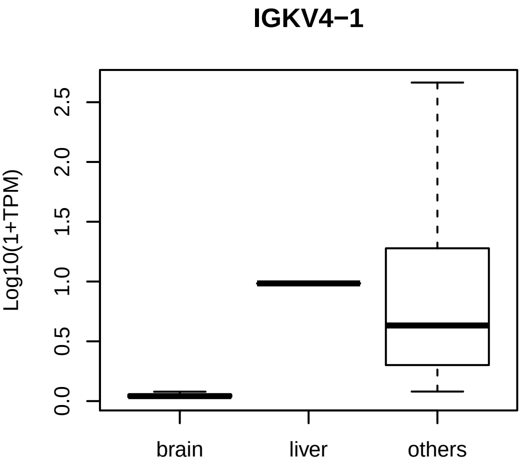

IGKV4−1
Log10(1+TPM)
0.0
0.5
1.0
1.5
2.0
2.5
brain
liver
others

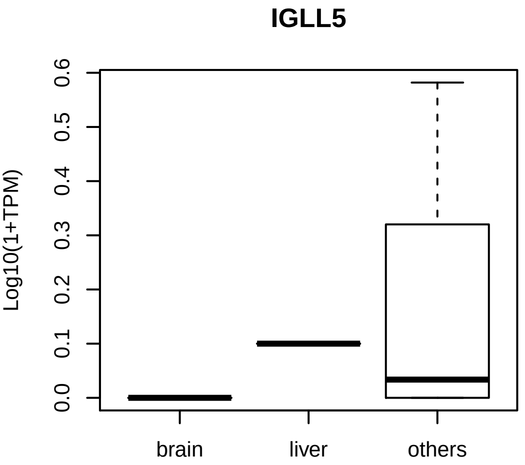

IGLL5
Log10(1+TPM)
0.0
0.1
0.2
0.3
0.4
0.5
0.6
brain
liver
others

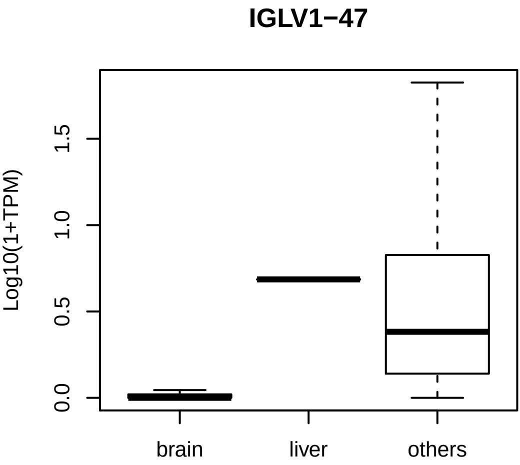
IGLV1−47
Log10(1+TPM)
0.0
0.5
1.0
1.5
brain
liver
others

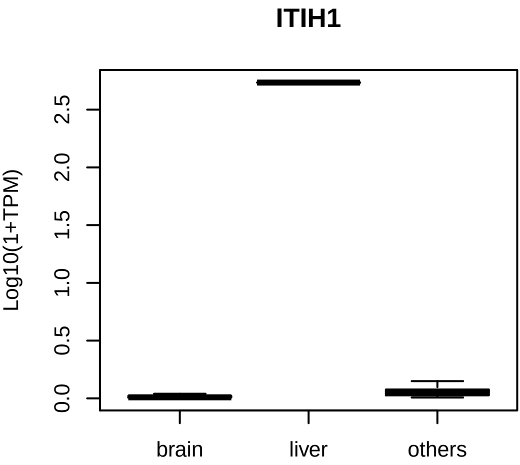
ITIH1
Log10(1+TPM)
0.0
0.5
1.0
1.5
2.0
2.5
brain
liver
others

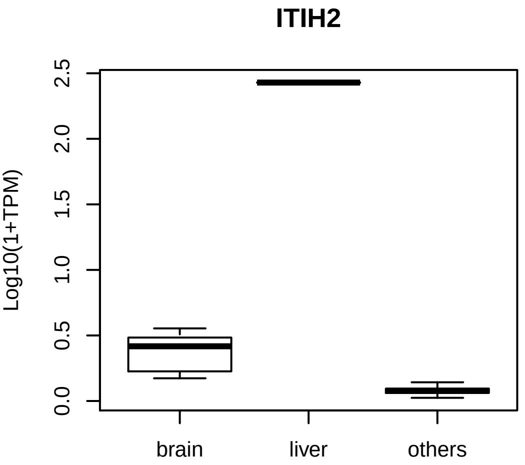
ITIH2
Log10(1+TPM)
0.0
0.5
1.0
1.5
2.0
2.5
brain
liver
others

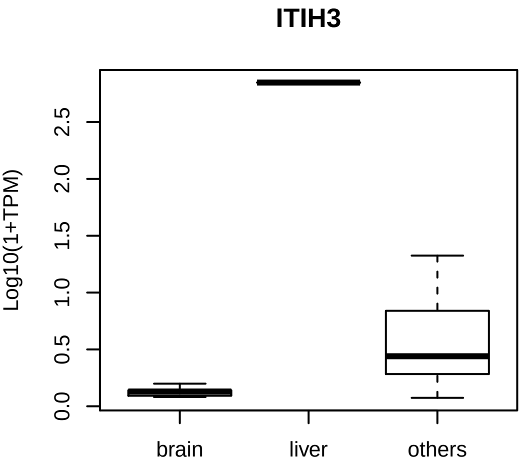
ITIH3
Log10(1+TPM)
0.0
0.5
1.0
1.5
2.0
2.5
brain
liver
others

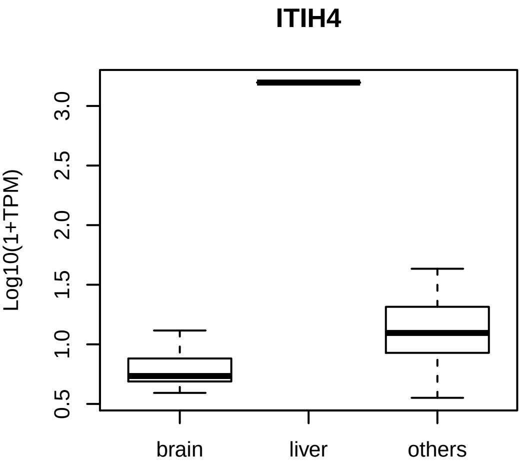
ITIH4
Log10(1+TPM)
0.5
1.0
1.5
2.0
2.5
3.0
brain
liver
others

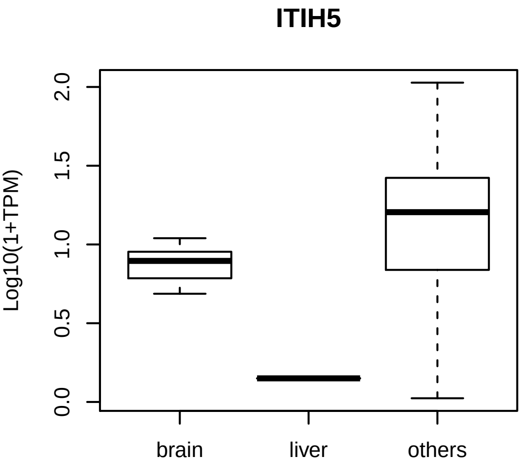
ITIH5
Log10(1+TPM)
0.0
0.5
1.0
1.5
2.0
brain
liver
others

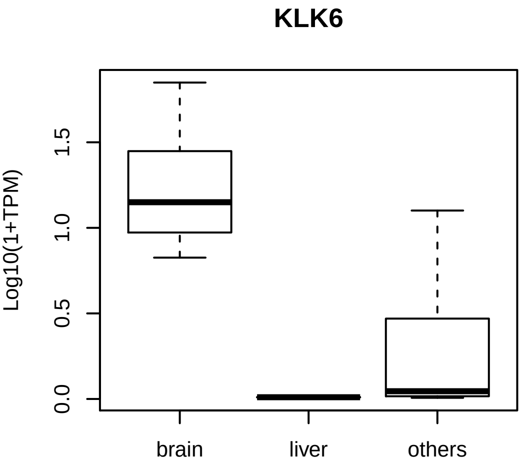
KLK6
Log10(1+TPM)
0.0
0.5
1.0
1.5
brain
liver
others

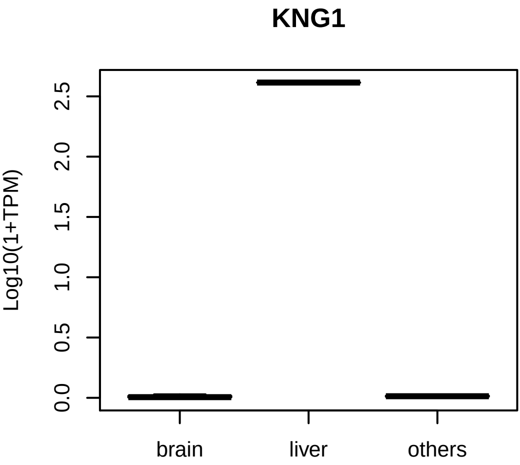
KNG1
Log10(1+TPM)
0.0
0.5
1.0
1.5
2.0
2.5
brain
liver
others

KRT10

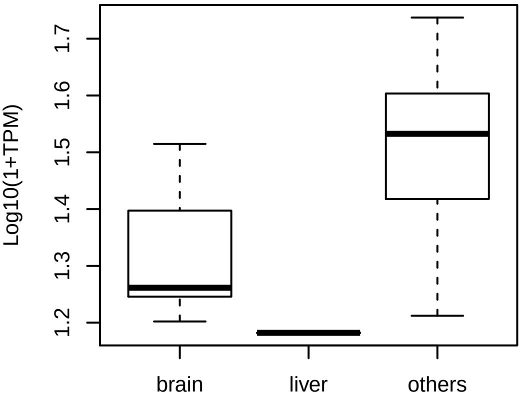

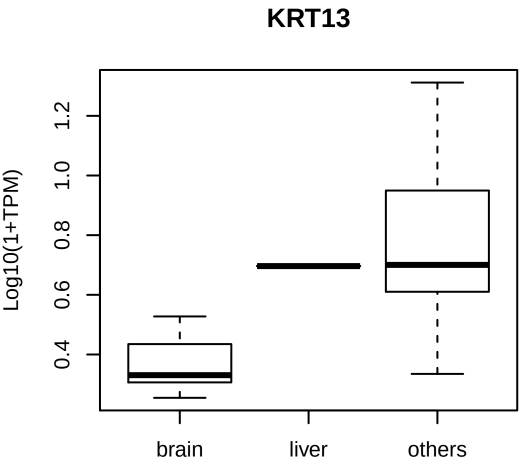
KRT13
Log10(1+TPM)
0.4
0.6
0.8
1.0
1.2
brain
liver
others

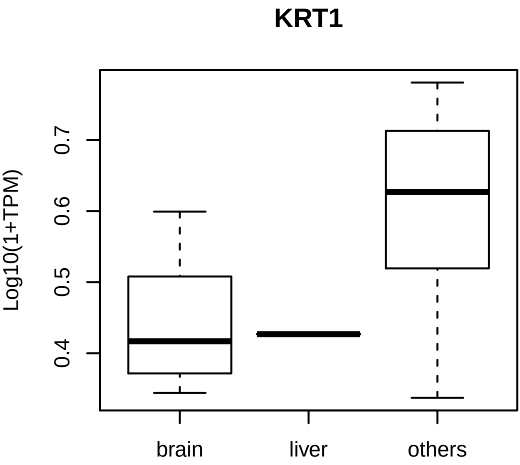
KRT1
Log10(1+TPM)
0.4
0.5
0.6
0.7
brain
liver
others

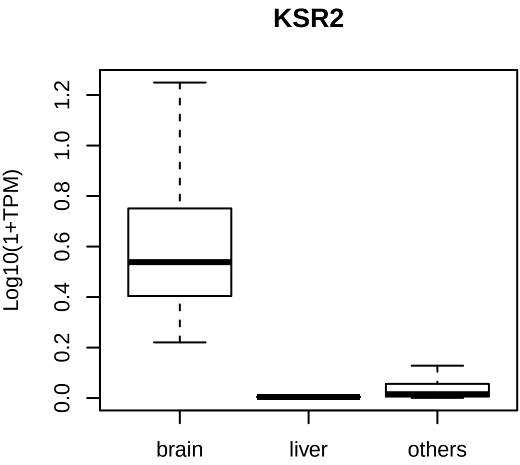

KSR2
Log10(1+TPM)
0.0
0.2
0.4
0.6
0.8
1.0
1.2
brain
liver
others

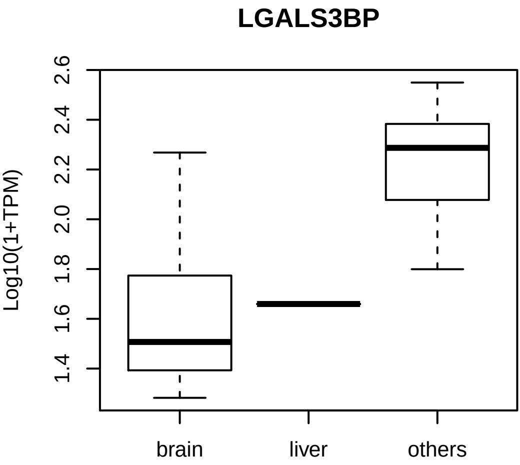
LGALS3BP
Log10(1+TPM)
1.4
1.6
1.8
2.0
2.2
2.4
2.6
brain
liver
others

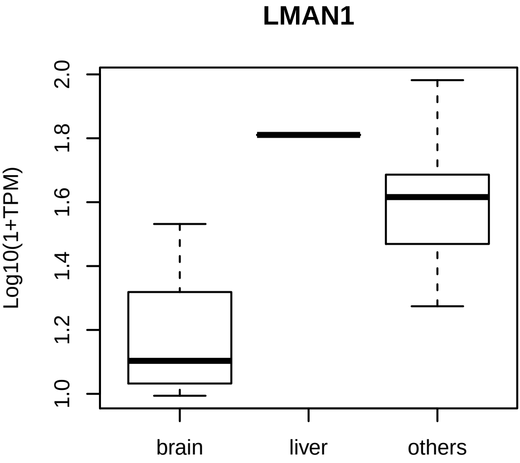
LMAN1
Log10(1+TPM)
1.0
1.2
1.4
1.6
1.8
2.0
brain
liver
others

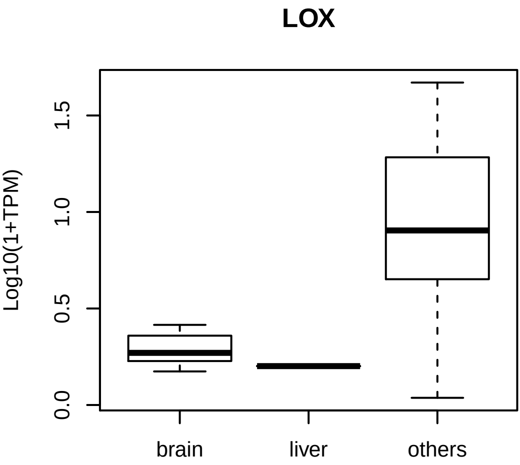
LOX
Log10(1+TPM)
0.0
0.5
1.0
1.5
brain
liver
others

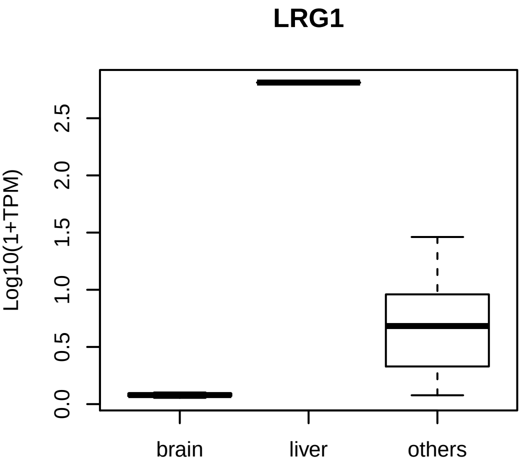
LRG1
Log10(1+TPM)
0.0
0.5
1.0
1.5
2.0
2.5
brain
liver
others

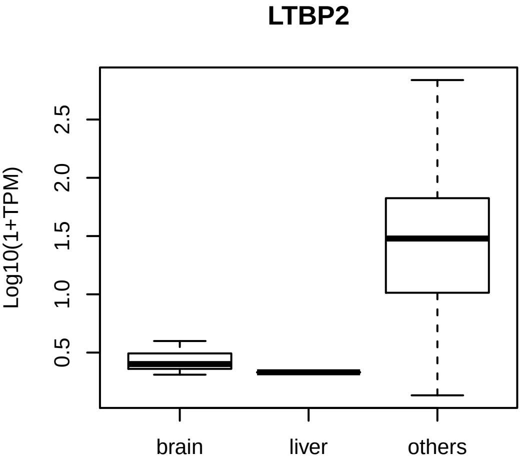
LTBP2
Log10(1+TPM)
0.5
1.0
1.5
2.0
2.5
brain
liver
others

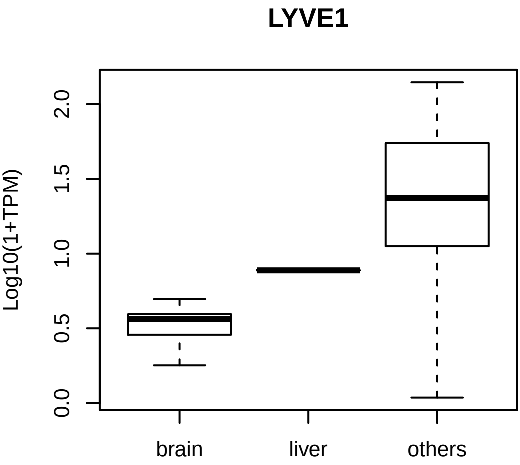

LYVE1
Log10(1+TPM)
0.0
0.5
1.0
1.5
2.0
brain
liver
others

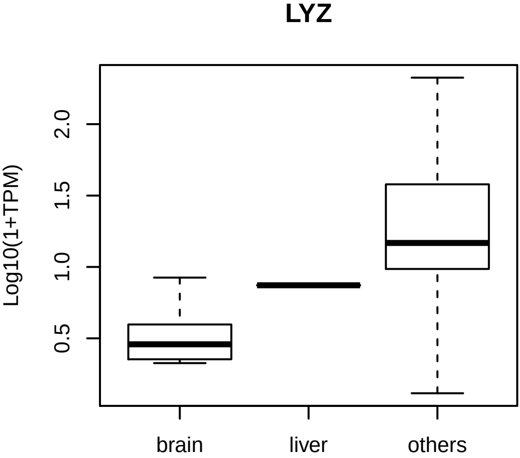
LYZ
Log10(1+TPM)
0.5
1.0
1.5
2.0
brain
liver
others

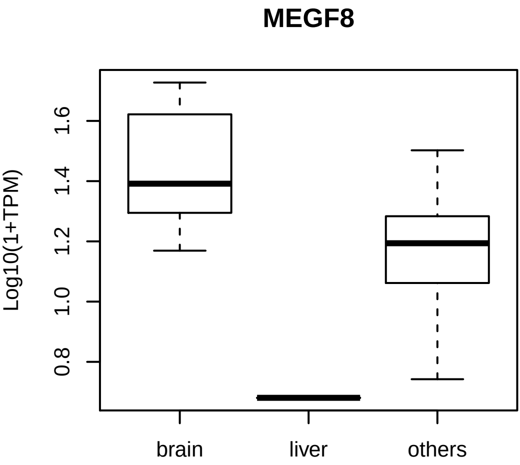
MEGF8
Log10(1+TPM)
0.8
1.0
1.2
1.4
1.6
brain
liver
others

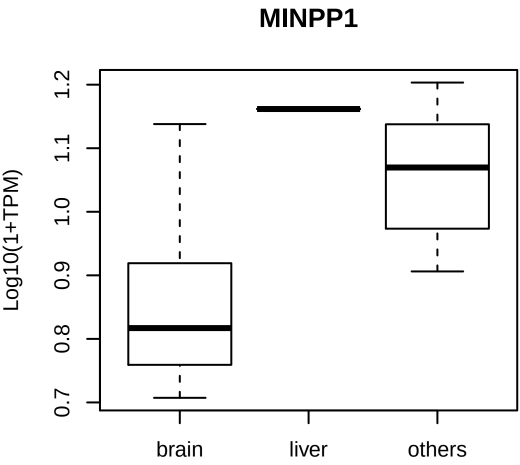
MINPP1
Log10(1+TPM)
0.7
0.8
0.9
1.0
1.1
1.2
brain
liver
others

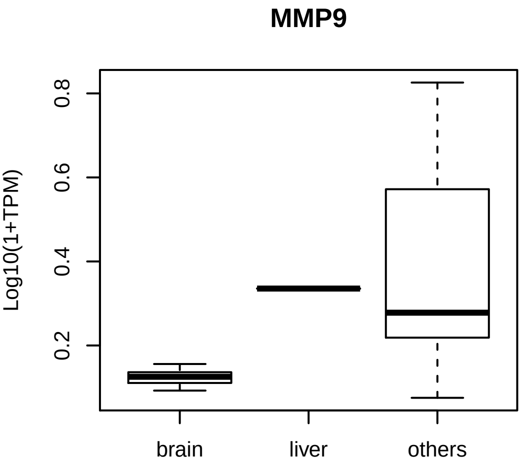
MMP9
Log10(1+TPM)
0.2
0.4
0.6
0.8
brain
liver
others

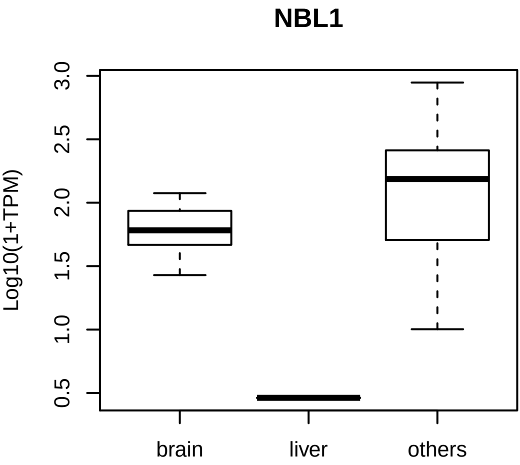
NBL1
Log10(1+TPM)
0.5
1.0
1.5
2.0
2.5
3.0
brain
liver
others

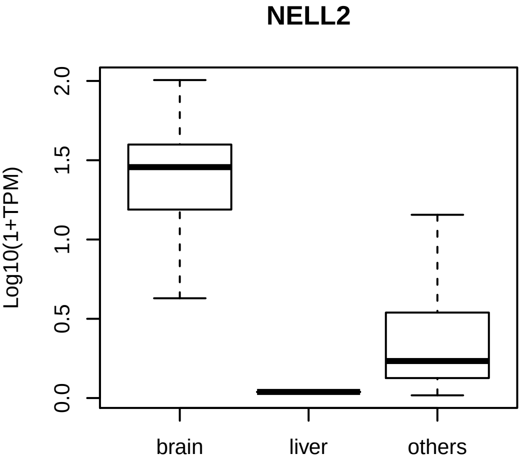
NELL2
Log10(1+TPM)
0.0
0.5
1.0
1.5
2.0
brain
liver
others

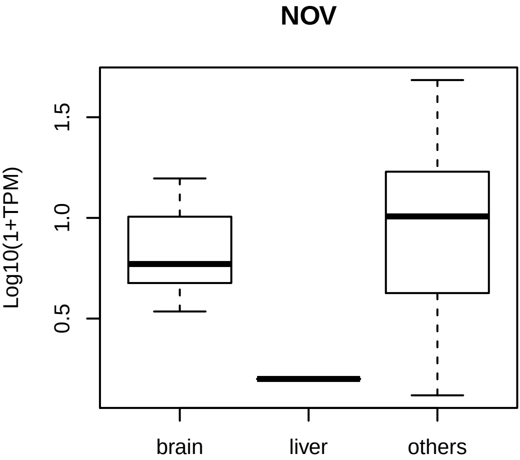
NOV
Log10(1+TPM)
0.5
1.0
1.5
brain
liver
others

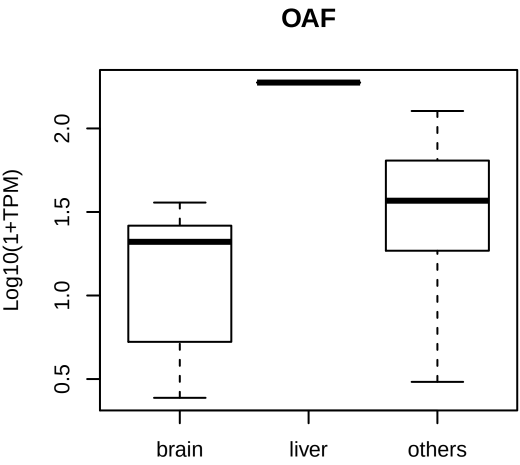
OAF
Log10(1+TPM)
0.5
1.0
1.5
2.0
brain
liver
others

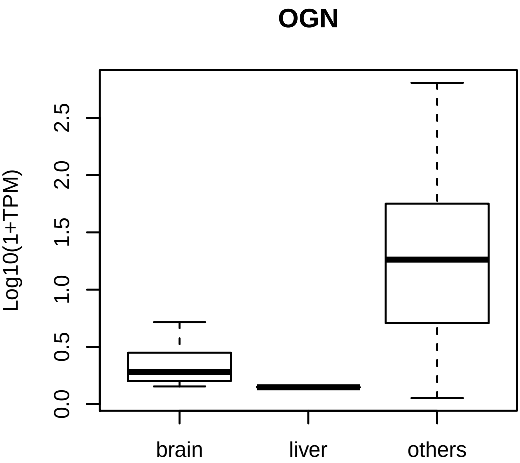
OGN
Log10(1+TPM)
0.0
0.5
1.0
1.5
2.0
2.5
brain
liver
others

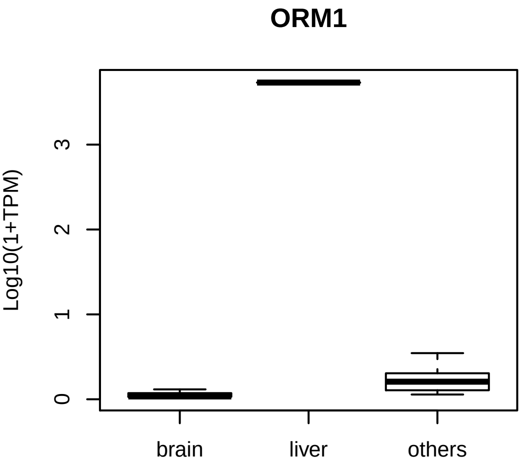
ORM1
Log10(1+TPM)
0
1
2
3
brain
liver
others

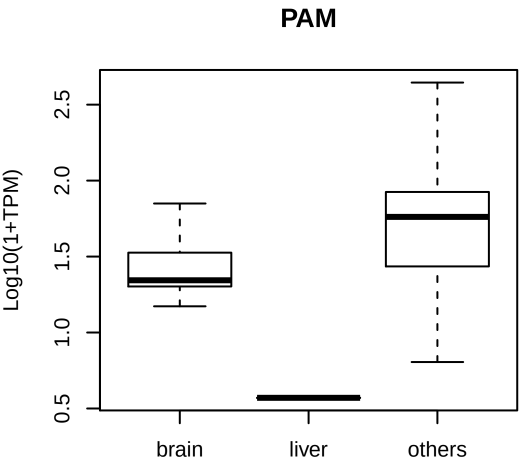

PAM
Log10(1+TPM)
0.5
1.0
1.5
2.0
2.5
brain
liver
others

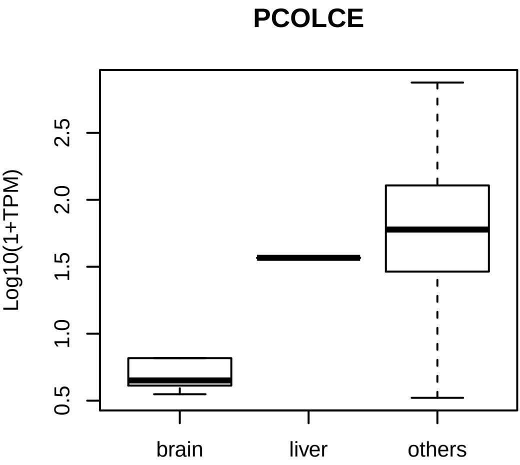
PCOLCE
Log10(1+TPM)
0.5
1.0
1.5
2.0
2.5
brain
liver
others

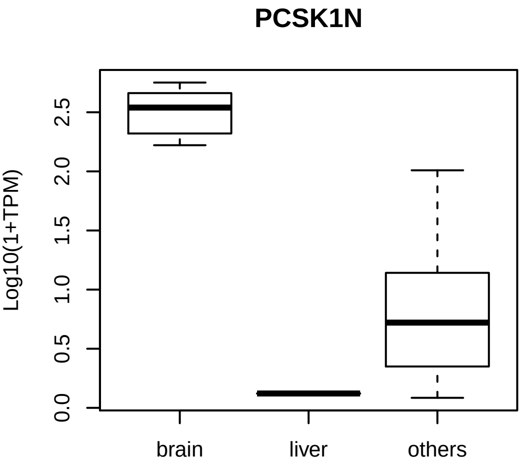
PCSK1N
Log10(1+TPM)
0.0
0.5
1.0
1.5
2.0
2.5
brain
liver
others

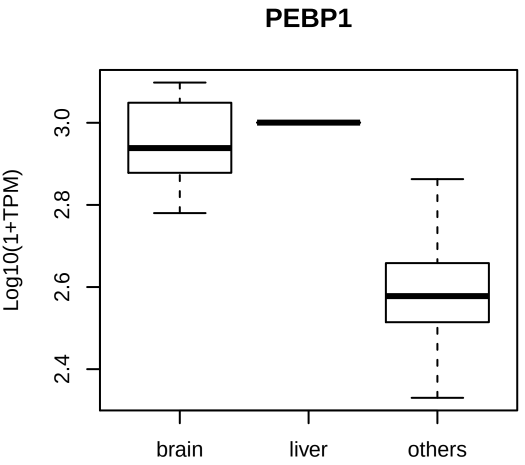
PEBP1
Log10(1+TPM)
3.0
2.8
2.6
2.4
brain
liver
others

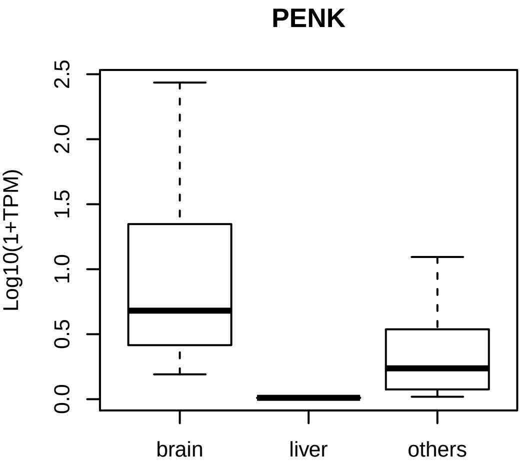
PENK
Log10(1+TPM)
0.0
0.5
1.0
1.5
2.0
2.5
brain
liver
others

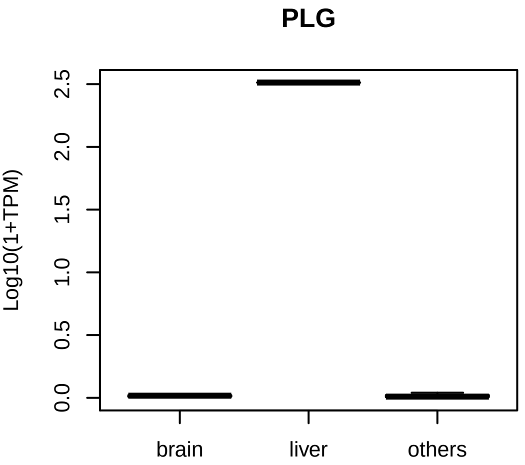
PLG
Log10(1+TPM)
0.0
0.5
1.0
1.5
2.0
2.5
brain
liver
others

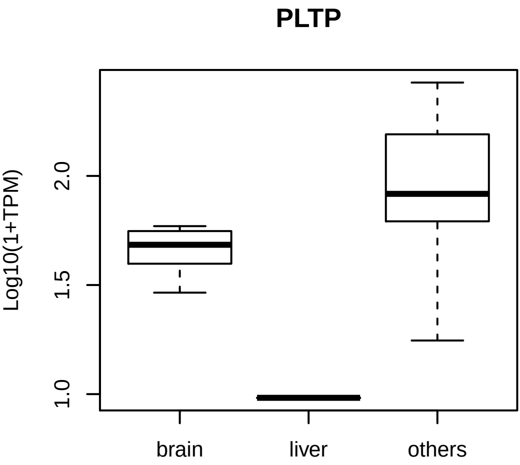
PLTP
Log10(1+TPM)
1.0
1.5
2.0
brain
liver
others

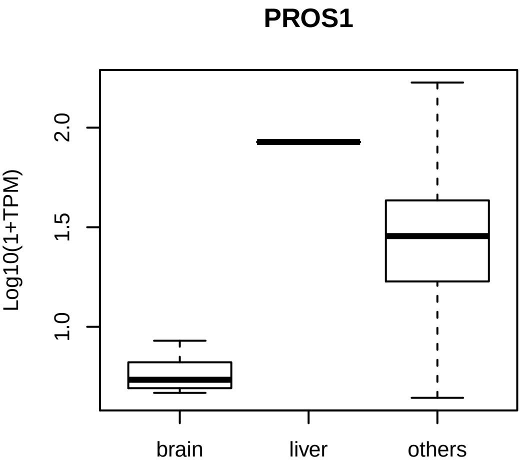

PROS1
Log10(1+TPM)
2.0
1.5
1.0
brain
liver
others

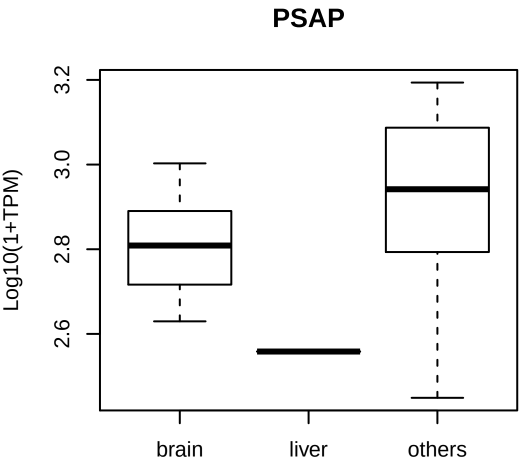
PSAP
Log10(1+TPM)
3.2
3.0
2.8
2.6
brain
liver
others

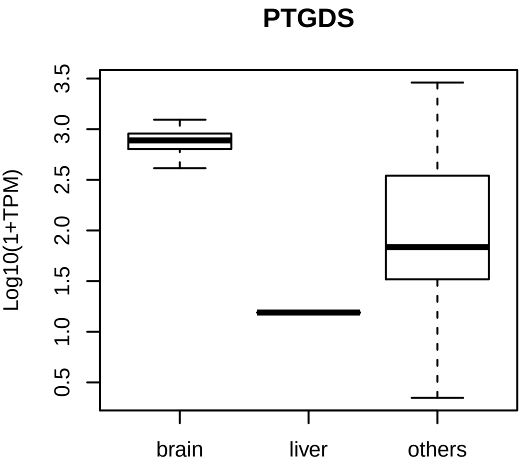

PTGDS
Log10(1+TPM)
0.5
1.0
1.5
2.0
2.5
3.0
3.5
brain
liver
others

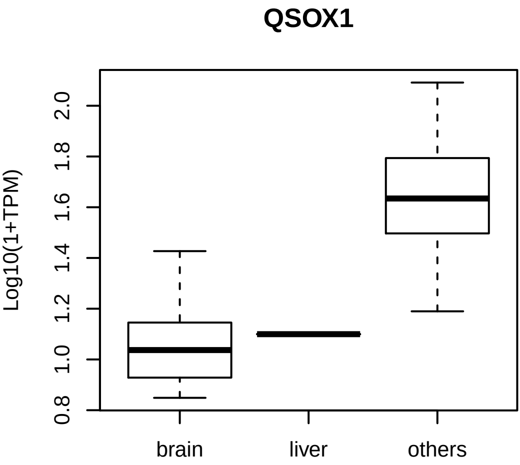
QSOX1
Log10(1+TPM)
0.8
1.0
1.2
1.4
1.6
1.8
2.0
brain
liver
others

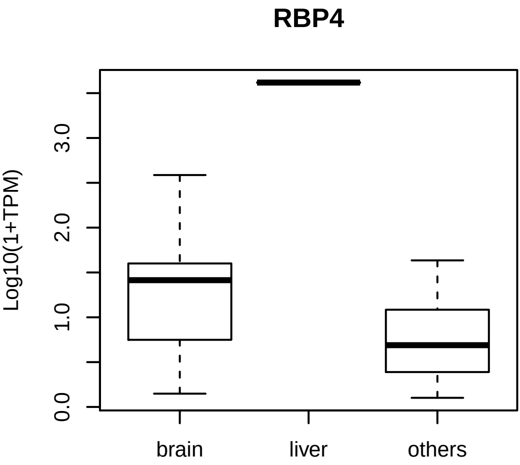
RBP4
Log10(1+TPM)
0.0
1.0
2.0
3.0
brain
liver
others

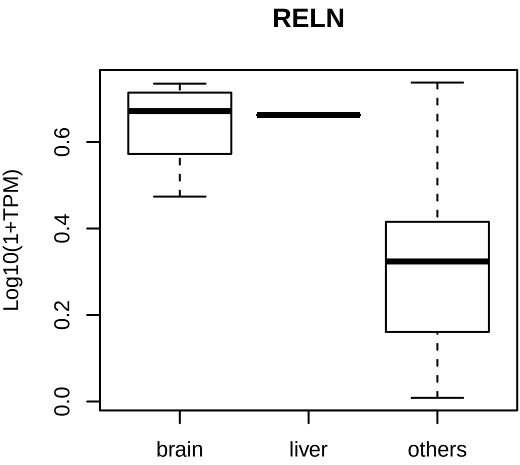
RELN
Log10(1+TPM)
0.0
0.2
0.4
0.6
brain
liver
others

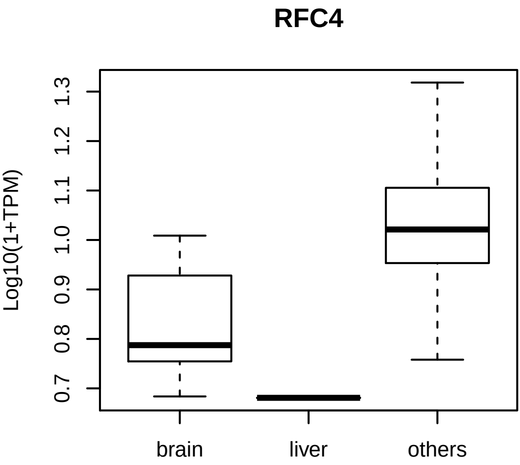
RFC4
Log10(1+TPM)
0.7
0.8
0.9
1.0
1.1
1.2
1.3
brain
liver
others

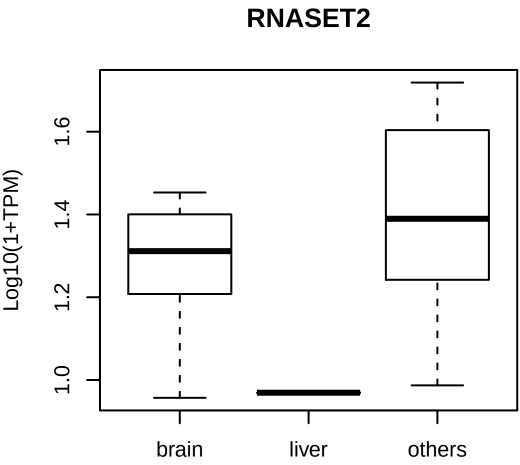
RNASET2
Log10(1+TPM)
1.0
1.2
1.4
1.6
brain
liver
others

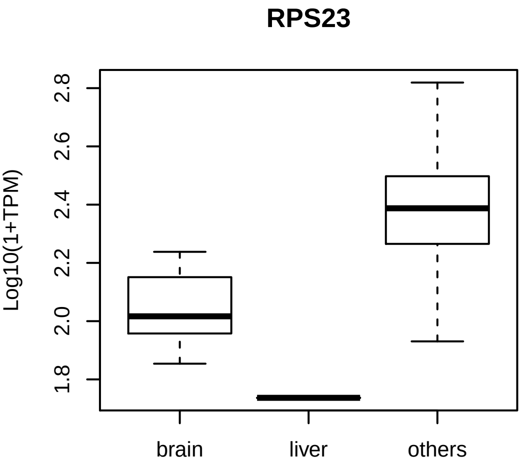
RPS23
Log10(1+TPM)
2.8
2.6
2.4
2.2
2.0
1.8
brain
liver
others

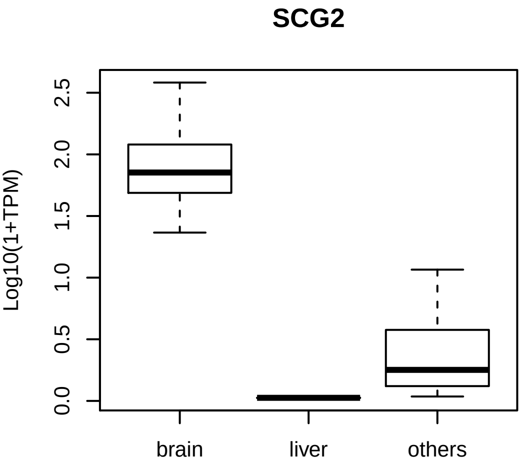
SCG2
Log10(1+TPM)
0.0
0.5
1.0
1.5
2.0
2.5
brain
liver
others

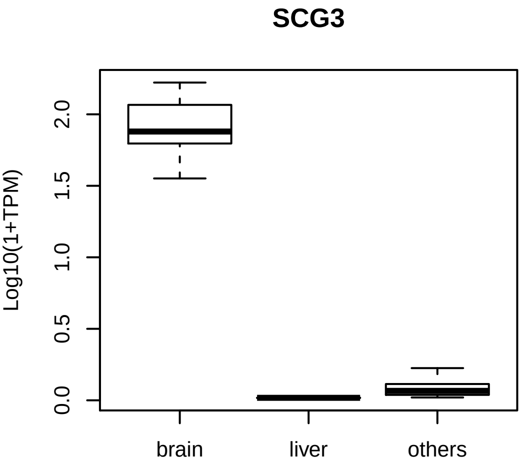
SCG3
Log10(1+TPM)
0.0
0.5
1.0
1.5
2.0
brain
liver
others

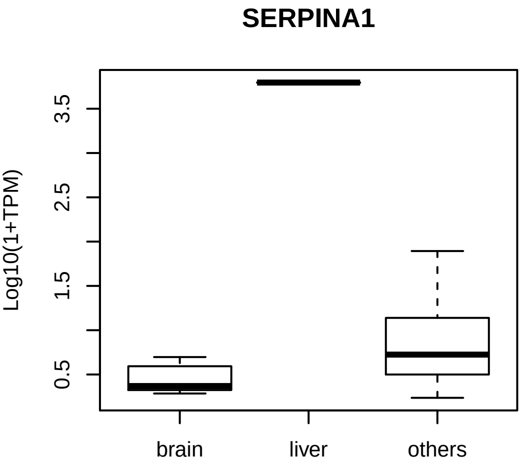
SERPINA1
Log10(1+TPM)
0.5
1.5
2.5
3.5
brain
liver
others

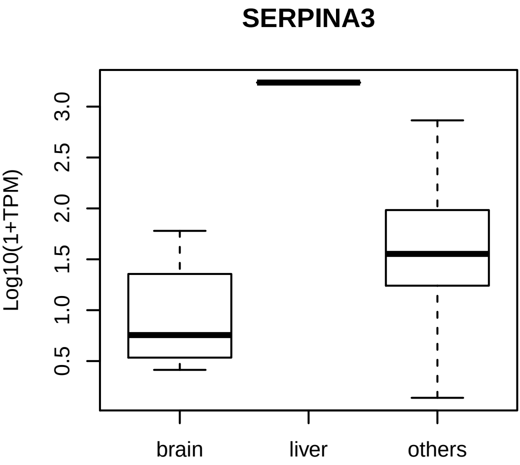
SERPINA3
Log10(1+TPM)
0.5
1.0
1.5
2.0
2.5
3.0
brain
liver
others

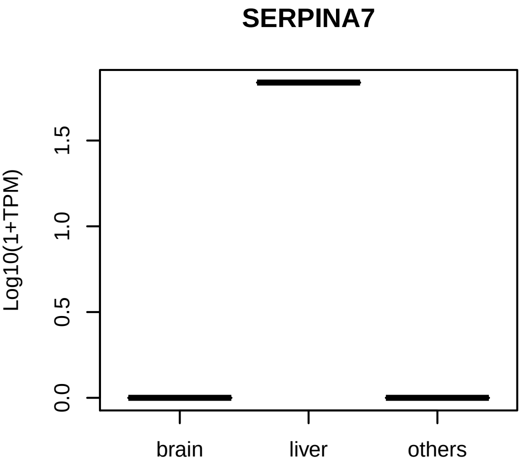

SERPINA7
Log10(1+TPM)
0.0
0.5
1.0
1.5
brain
liver
others

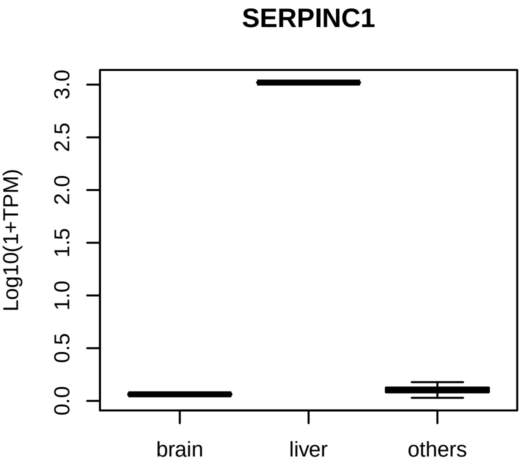

SERPINC1
Log10(1+TPM)
0.0
0.5
1.0
1.5
2.0
2.5
3.0
brain
liver
others

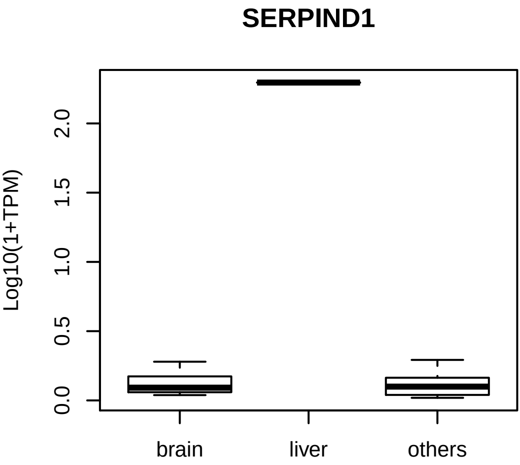
SERPIND1
Log10(1+TPM)
0.0
0.5
1.0
1.5
2.0
brain
liver
others

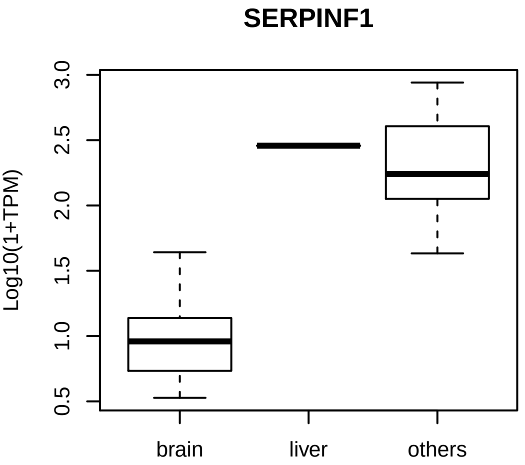
SERPINF1
Log10(1+TPM)
0.5
1.0
1.5
2.0
2.5
3.0
brain
liver
others

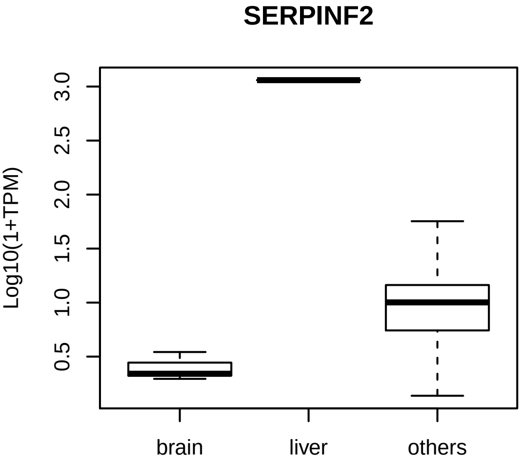
SERPINF2
Log10(1+TPM)
0.5
1.0
1.5
2.0
2.5
3.0
brain
liver
others

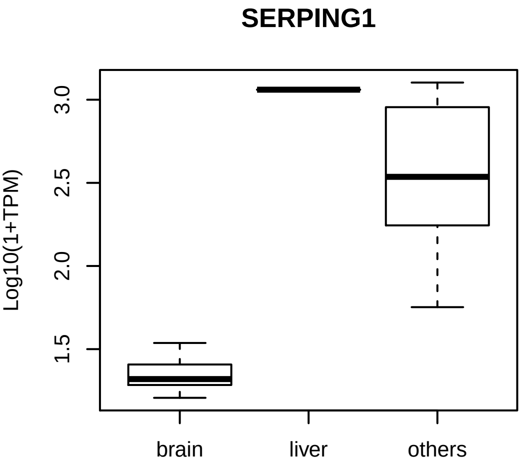
SERPING1
Log10(1+TPM)
1.5
2.0
2.5
3.0
brain
liver
others

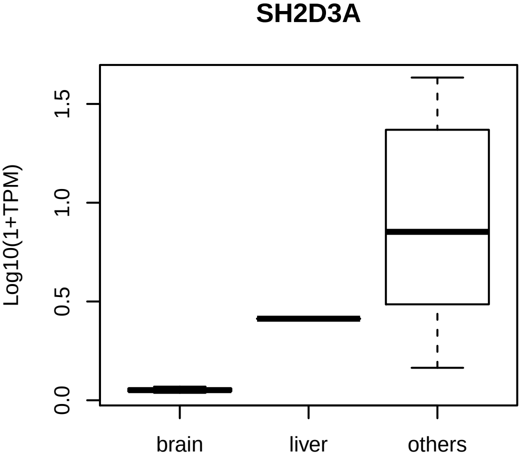
SH2D3A
Log10(1+TPM)
0.0
0.5
1.0
1.5
brain
liver
others

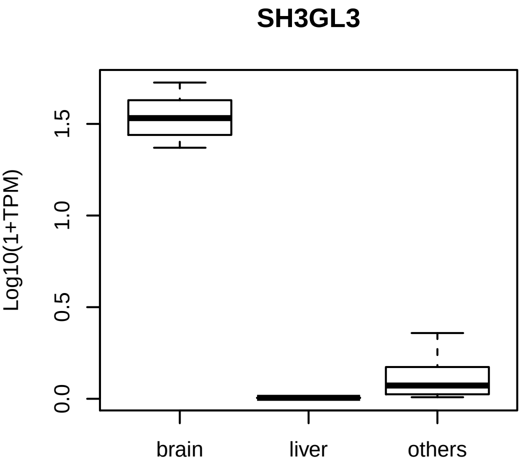
SH3GL3
Log10(1+TPM)
0.0
0.5
1.0
1.5
brain
liver
others

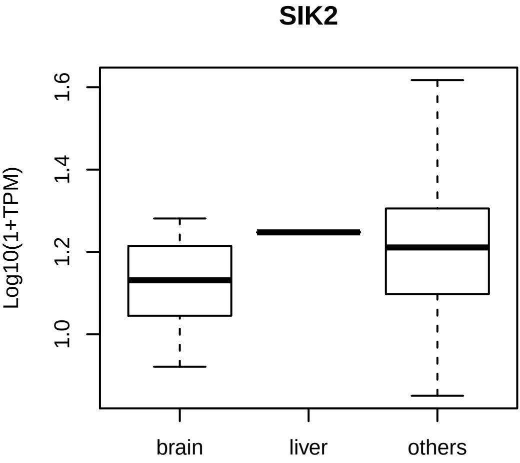
SIK2
Log10(1+TPM)
1.0
1.2
1.4
1.6
brain
liver
others

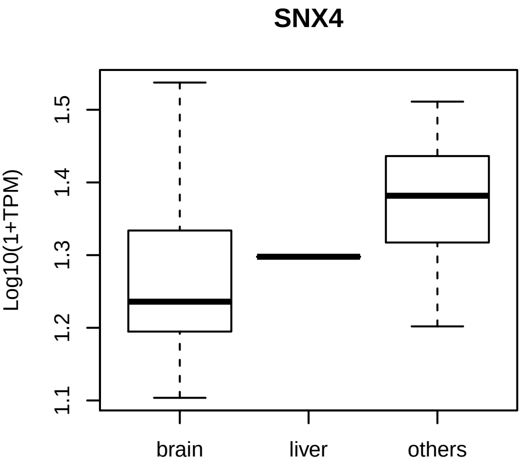
SNX4
Log10(1+TPM)
1.1
1.2
1.3
1.4
1.5
brain
liver
others

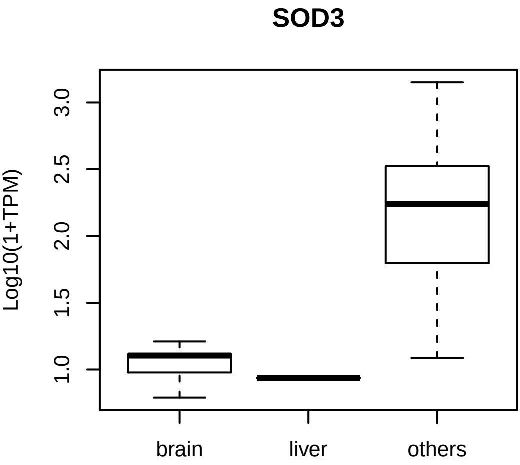
SOD3
Log10(1+TPM)
1.0
1.5
2.0
2.5
3.0
brain
liver
others

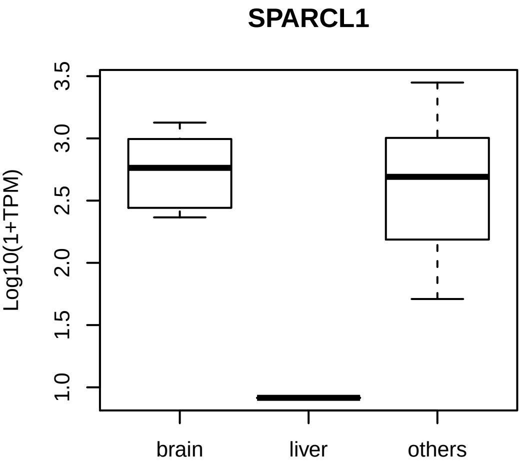
SPARCL1
Log10(1+TPM)
1.0
1.5
2.0
2.5
3.0
3.5
brain
liver
others

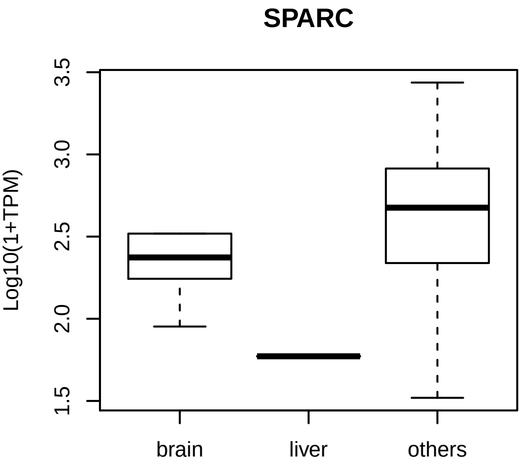
SPARC
Log10(1+TPM)
1.5
2.0
2.5
3.0
3.5
brain
liver
others

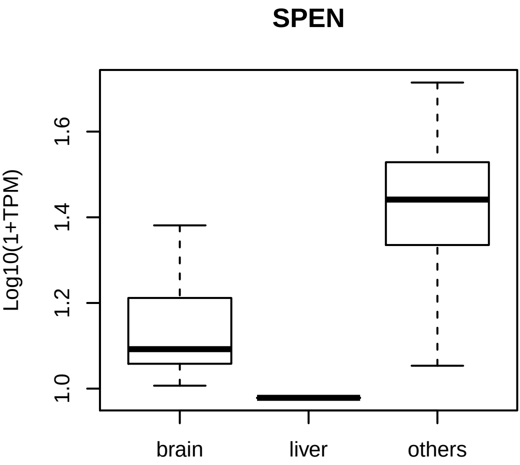
SPEN
Log10(1+TPM)
1.0
1.2
1.4
1.6
brain
liver
others

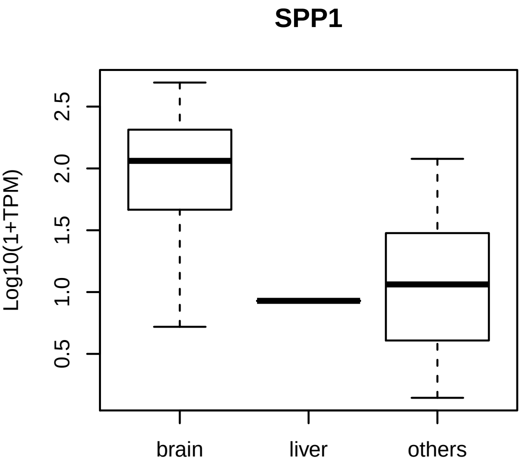
SPP1
Log10(1+TPM)
0.5
1.0
1.5
2.0
2.5
brain
liver
others

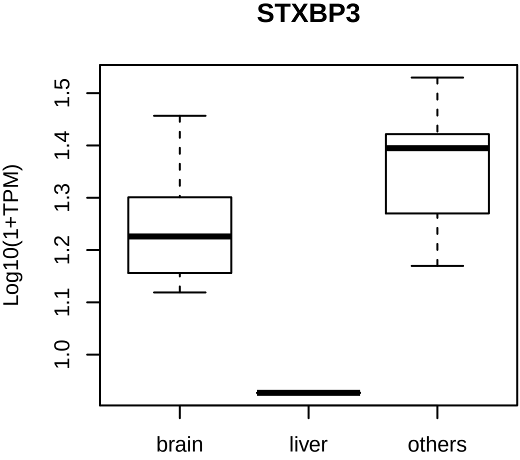
STXBP3
Log10(1+TPM)
1.0
1.1
1.2
1.3
1.4
1.5
brain
liver
others

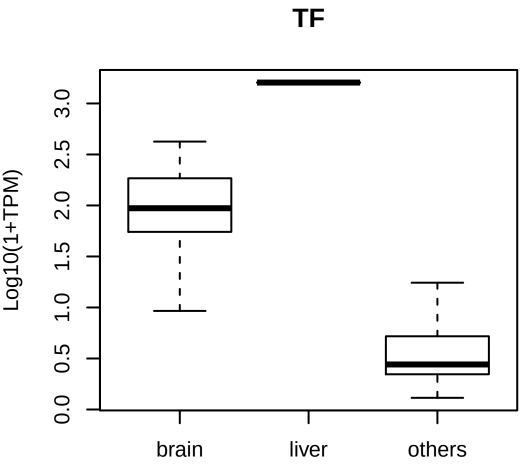
TF
Log10(1+TPM)
0.0
0.5
1.0
1.5
2.0
2.5
3.0
brain
liver
others

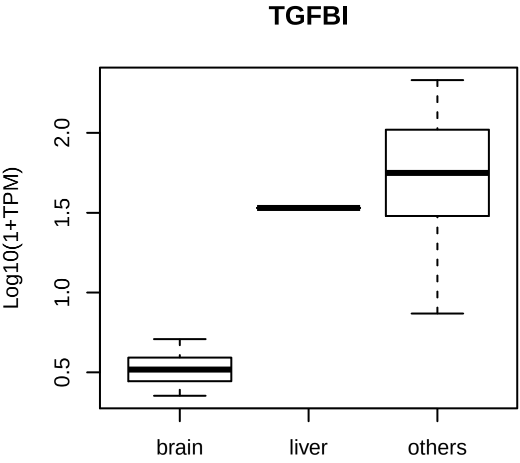
TGFBI
Log10(1+TPM)
0.5
1.0
1.5
2.0
brain
liver
others

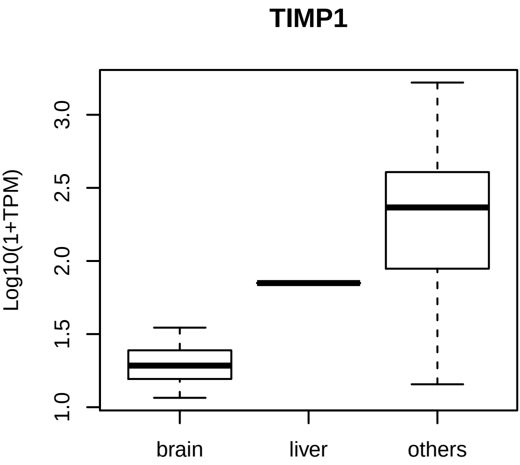
TIMP1
Log10(1+TPM)
1.0
1.5
2.0
2.5
3.0
brain
liver
others

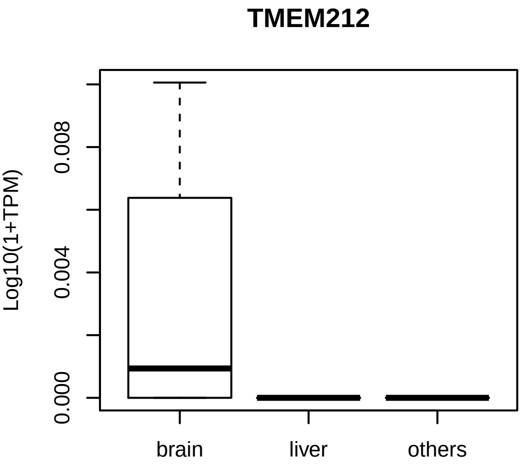
TMEM212
Log10(1+TPM)
0.000
0.004
0.008
brain
liver
others

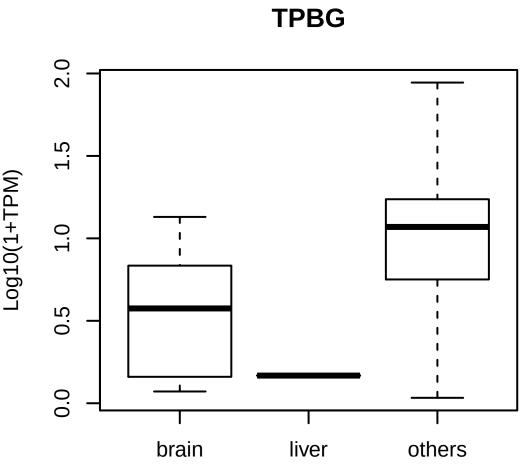
TPBG
Log10(1+TPM)
2.0
1.5
1.0
0.5
0.0
brain
liver
others

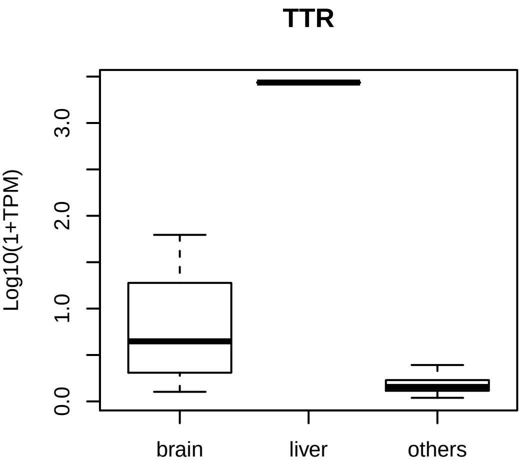
TTR
Log10(1+TPM)
0.0
1.0
2.0
3.0
brain
liver
others

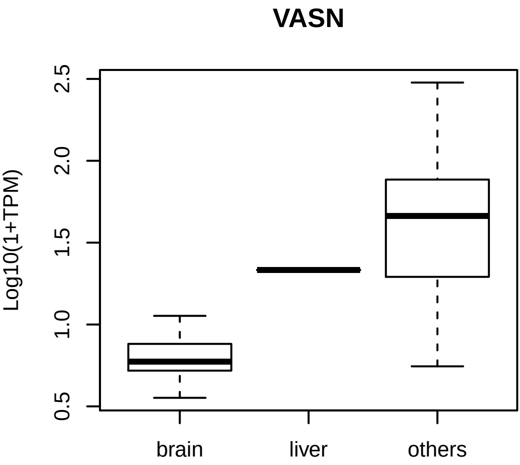
VASN
Log10(1+TPM)
0.5
1.0
1.5
2.0
2.5
brain
liver
others

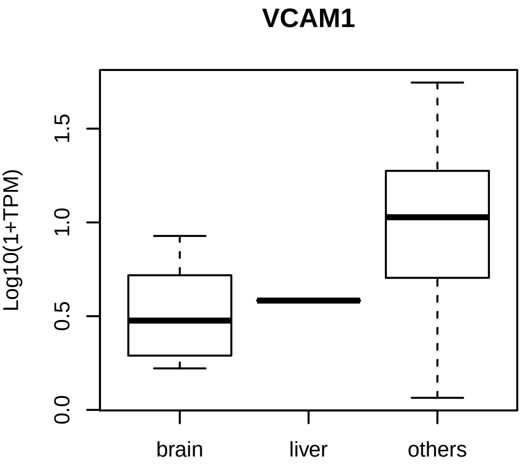
VCAM1
Log10(1+TPM)
0.0
0.5
1.0
1.5
brain
liver
others

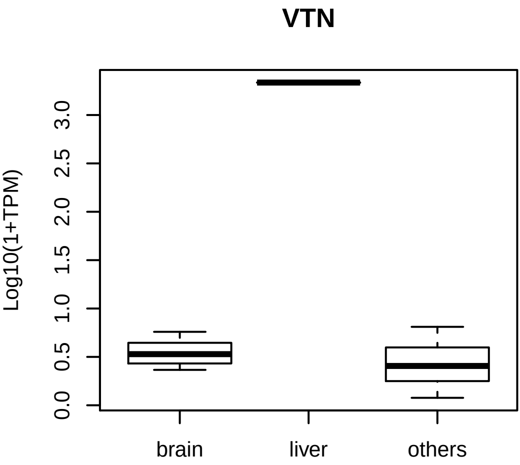
VTN
Log10(1+TPM)
0.0
0.5
1.0
1.5
2.0
2.5
3.0
brain
liver
others
