## Supplementary material for "*In Vivo* Large Scale Mapping Of Protein Turnover In The Human Cerebrospinal Fluid": HRMS protein models

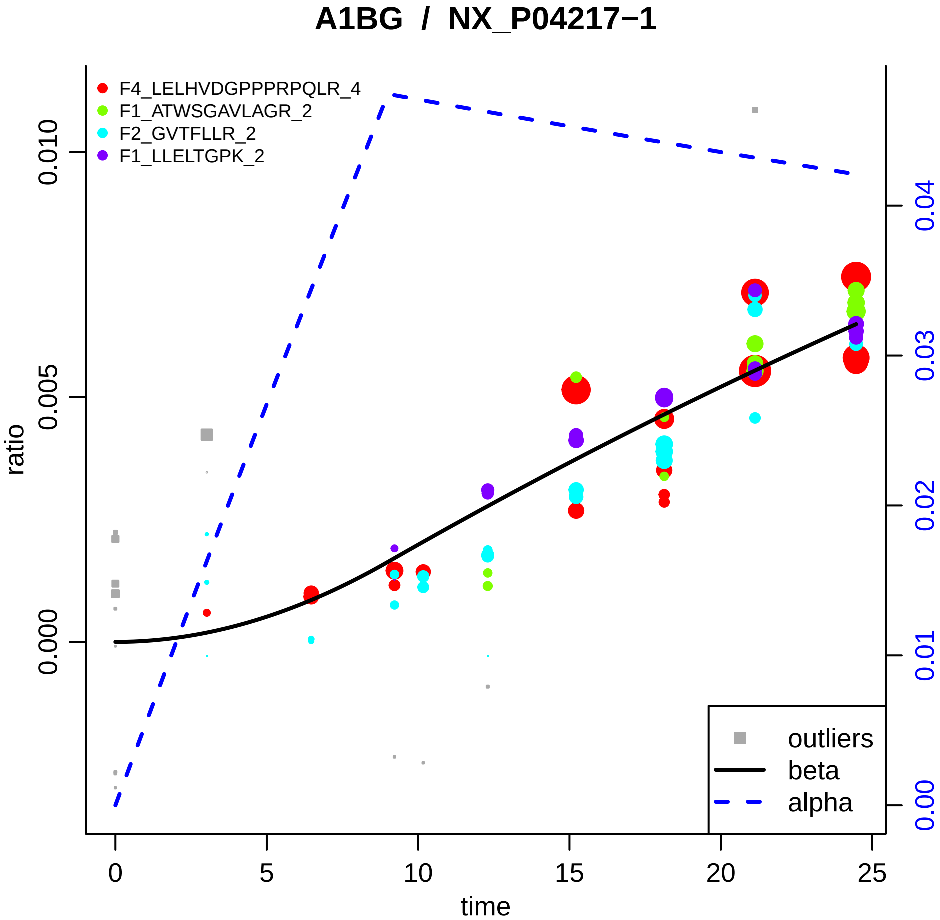

A1BG / NX_P04217−1
F4_LELHVDGPPPRPQLR_4
F1_ATWSGAVLAGR_2
F2_GVTFLLR_2
F1_LLELTGPK_2
outliers
beta
alpha
ratio
time
0.000
0.005
0.010
0
5
10
15
20
25
0.00
0.01
0.02
0.03
0.04

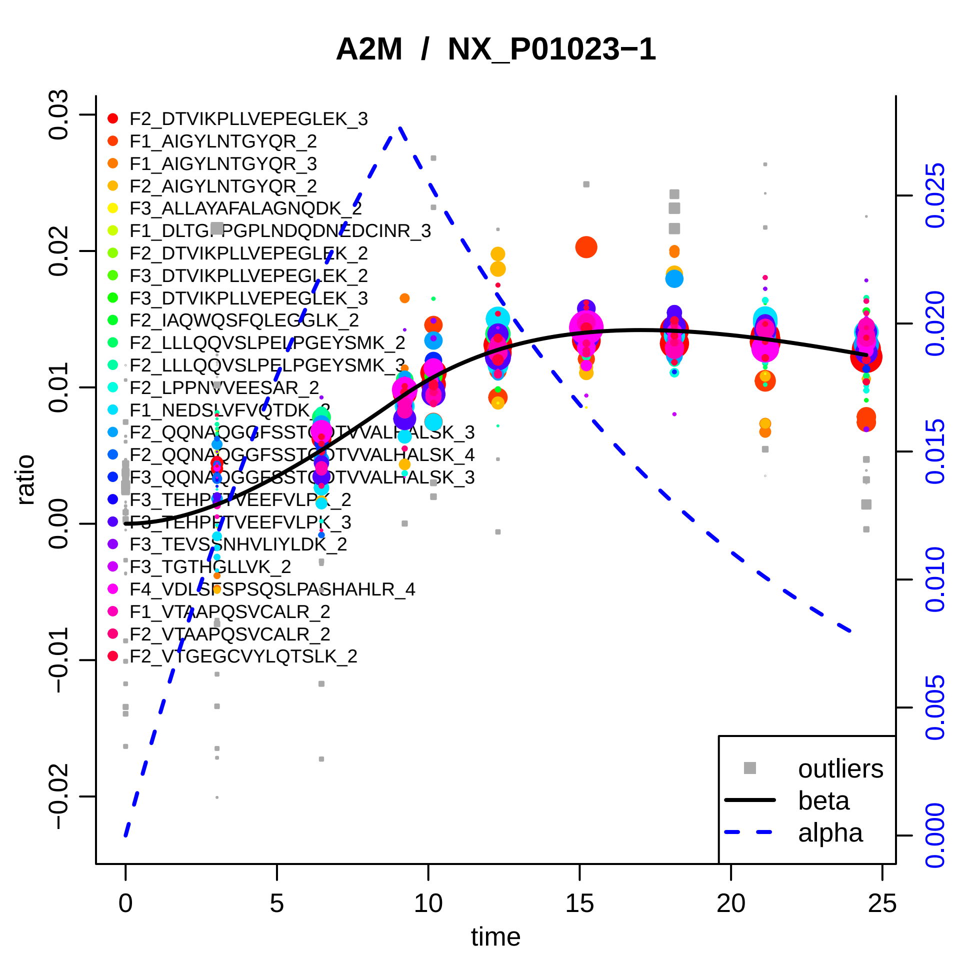

A2M / NX_P01023−1
F2_DTVIKPLLVEPEGLEK_3
F1_AIGYLNTGYQR_2
F1_AIGYLNTGYQR_3
F2_AIGYLNTGYQR_2
F3_ALLAYAFALAGNQDK_2
F1_DLTGFPGPLNDQDNEDCINR_3
F2_DTVIKPLLVEPEGLEK_2
F3_DTVIKPLLVEPEGLEK_2
F3_DTVIKPLLVEPEGLEK_3
F2_IAQWQSFQLEGGLK_2
F2_LLLQQVSLPELPGEYSMK_2
F2_LLLQQVSLPELPGEYSMK_3
F2_LPPNVVEESAR_2
F1_NEDSLVFVQTDK_2
F2_QQNAQGGFSSTQDTVVALHALSK_3
F2_QQNAQGGFSSTQDTVVALHALSK_4
F3_QQNAQGGFSSTQDTVVALHALSK_3
F3_TEHPFTVEEFVLPK_2
F3_TEHPFTVEEFVLPK_3
F3_TEVSSNHVLIYLDK_2
F3_TGTHGLLVK_2
F4_VDLSFSPSQSLPASHAHLR_4
F1_VTAAPQSVCALR_2
F2_VTAAPQSVCALR_2
F2_VTGEGCVYLQTSLK_2
outliers
beta
alpha
ratio
time
0.03
0.02
0.01
0.00
−0.01
−0.02
0
5
10
15
20
25
0.025
0.020
0.015
0.010
0.005
0.000

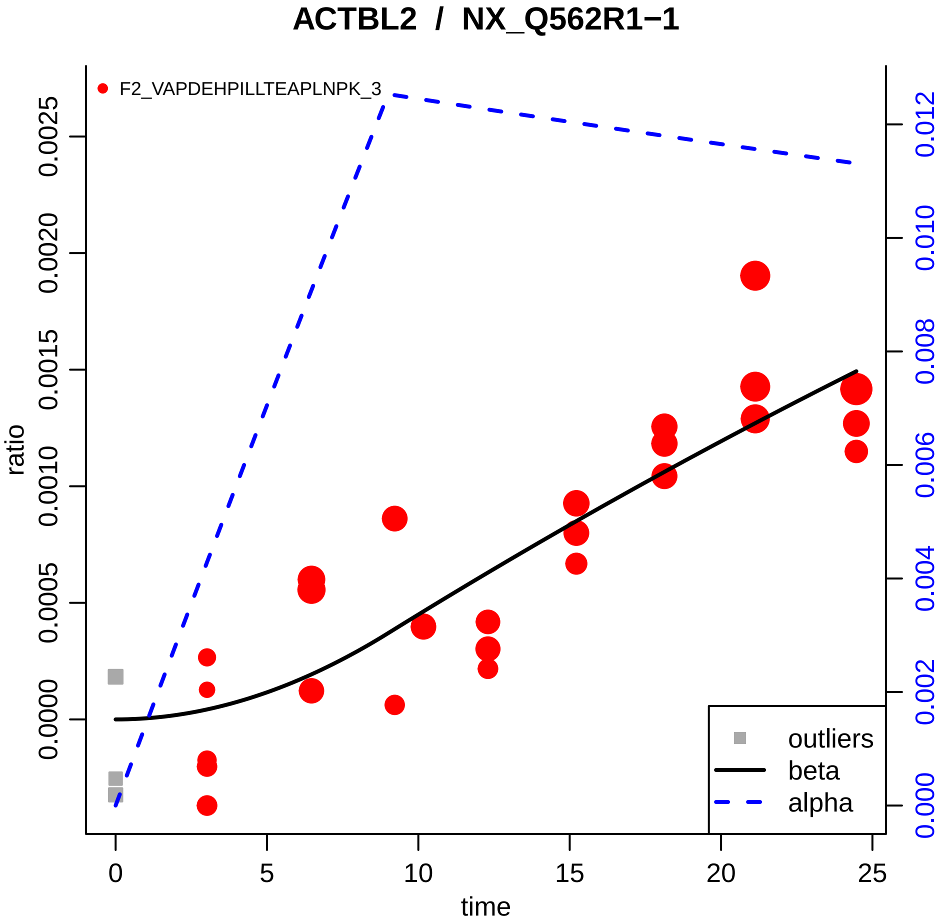

ACTBL2 / NX_Q562R1−1
F2_VAPDEHPILLTEAPLNPK_3
ratio
time
0.0000
0.0005
0.0010
0.0015
0.0020
0.0025
0
5
10
15
20
25
0.000
0.002
0.004
0.006
0.008
0.010
0.012
outliers
beta
alpha

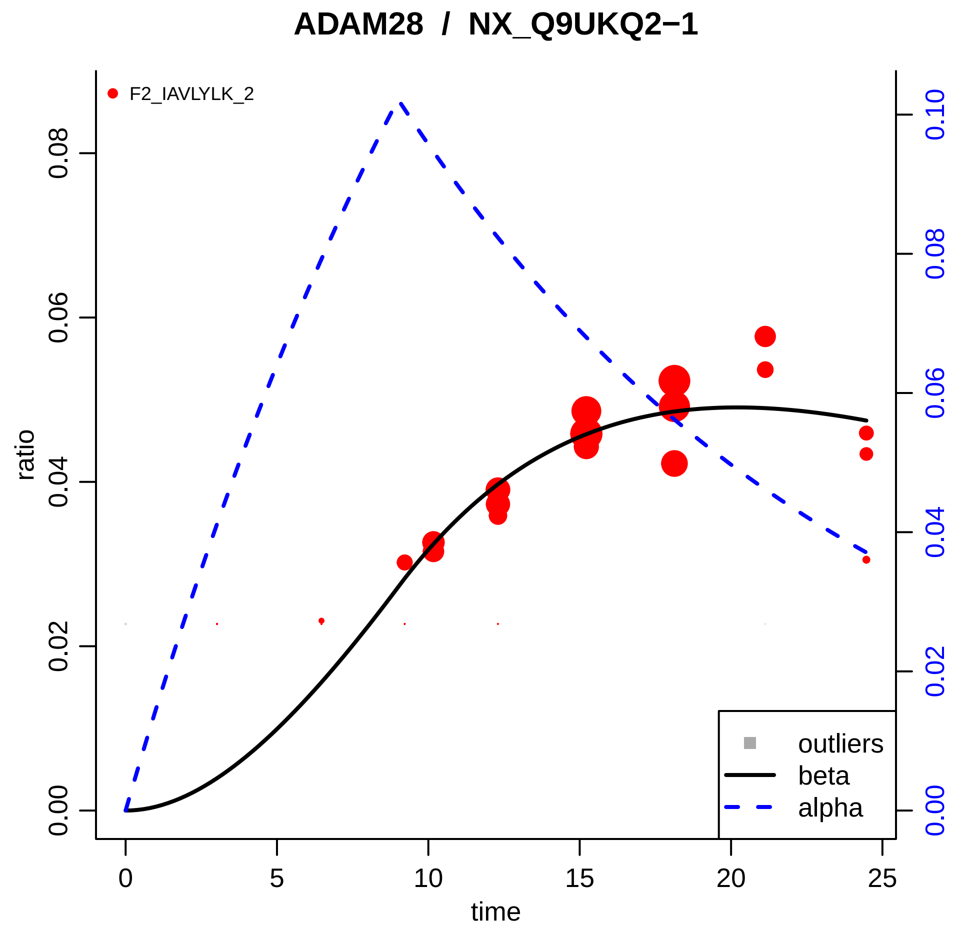

ADAM28 / NX_Q9UKQ2−1
F2_IAVLYLK_2
ratio
time
0.00
0.02
0.04
0.06
0.08
0.10
0
5
10
15
20
25
outliers
beta
alpha

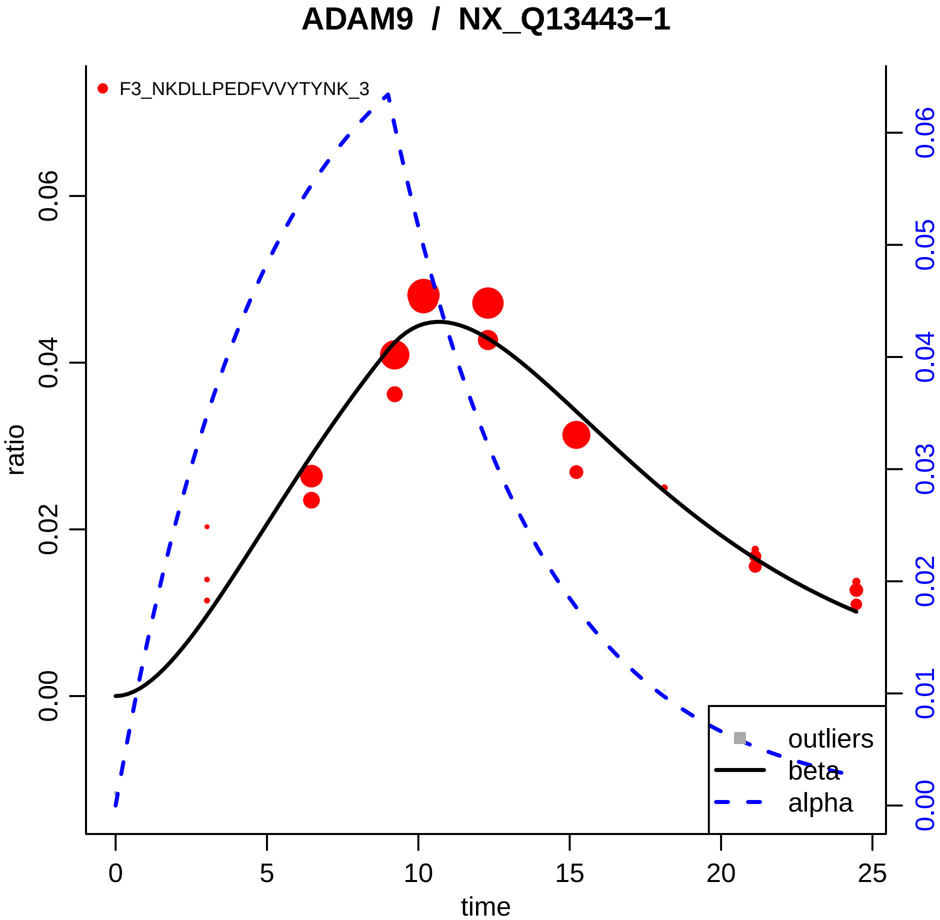

ADAM9 / NX_Q13443−1
F3_NKDLLPEDFVVYTYNK_3
ratio
time
0.00
0.02
0.04
0.06
0
5
10
15
20
25
0.00
0.01
0.02
0.03
0.04
0.05
0.06
outliers
beta
alpha

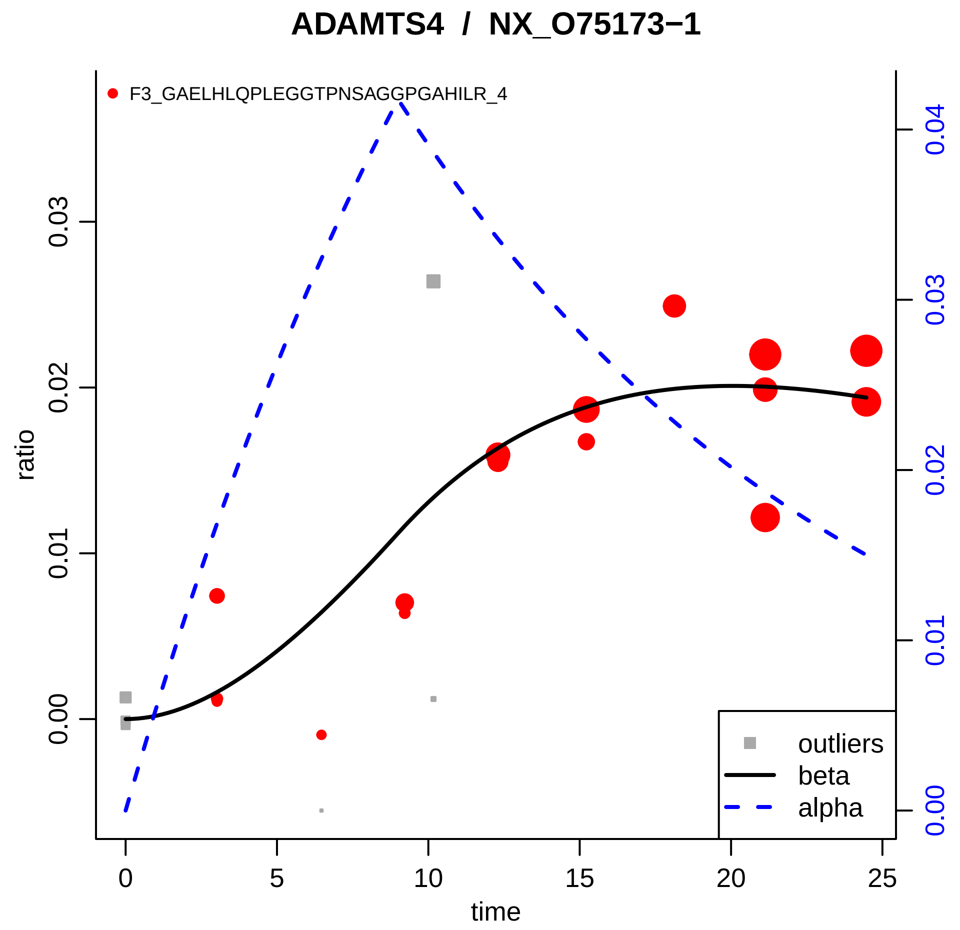
ADAMTS4 / NX_O75173−1
F3_GAELHLQPLEGGTPNSAGGPGAHILR_4
ratio
time
0.00
0.01
0.02
0.03
0.04
0
5
10
15
20
25
outliers
beta
alpha

### AGT / NX_P01019−1

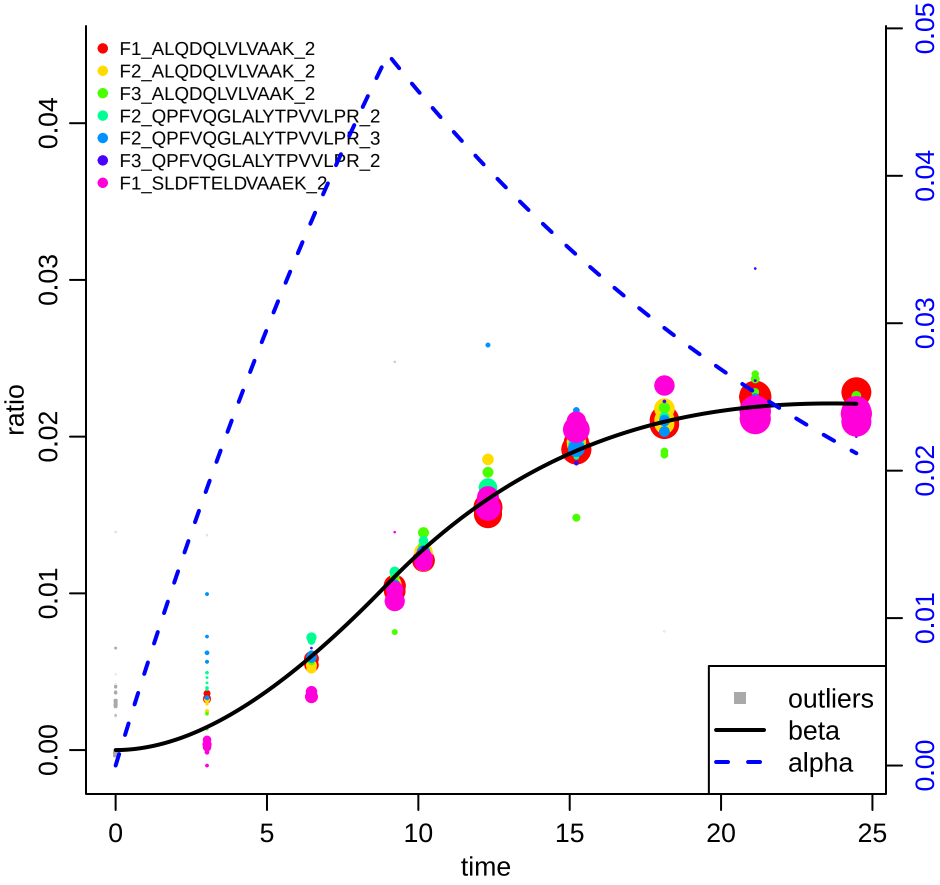

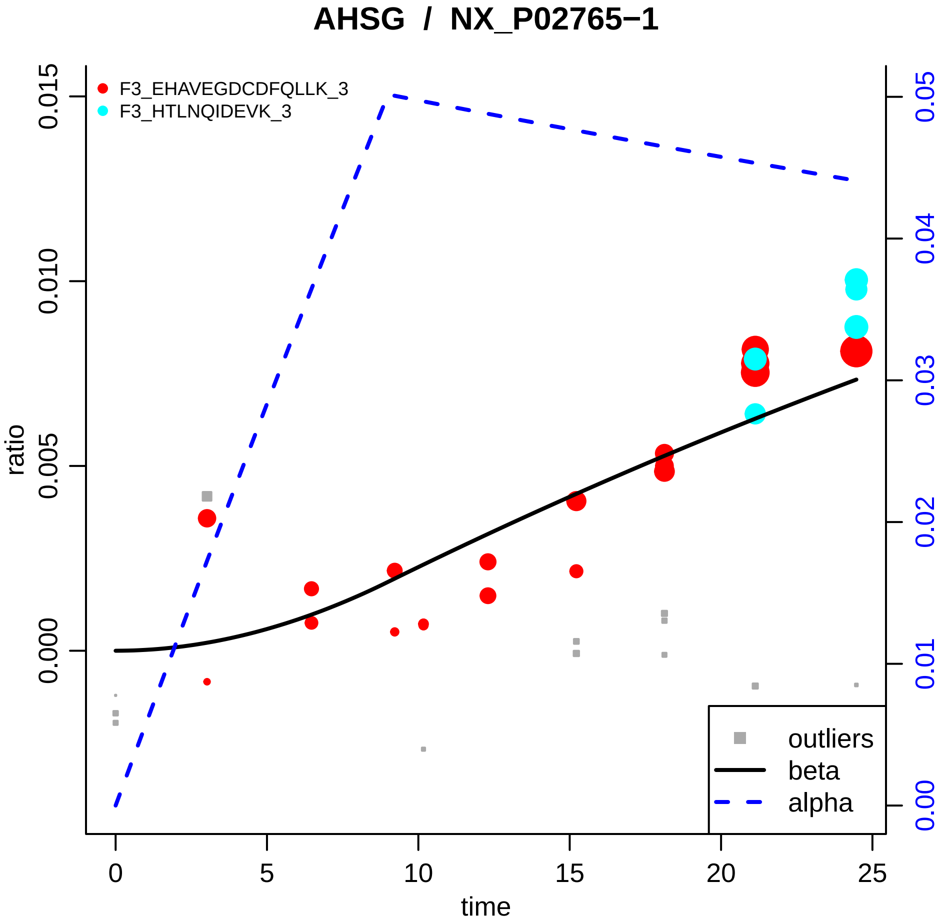
AHSG / NX_P02765−1
F3_EHAVEGDCDFQLLK_3
F3_HTLNQIDEVK_3
ratio
time
0.000
0.005
0.010
0.015
0.00
0.01
0.02
0.03
0.04
0.05
0
5
10
15
20
25
outliers
beta
alpha

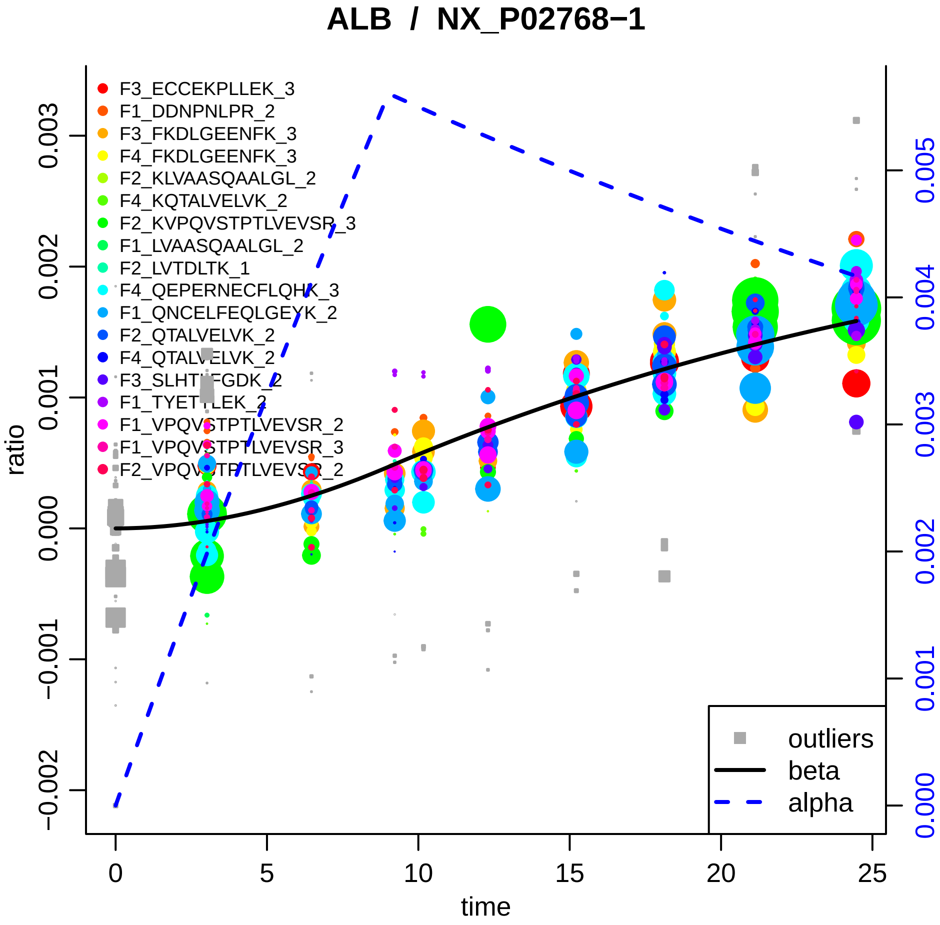

ALB / NX_P02768−1
F3_ECCEKPLLEK_3
F1_DDNPNLPR_2
F3_FKDLGEENFK_3
F4_FKDLGEENFK_3
F2_KLVAASQAALGL_2
F4_KQTALVELVK_2
F2_KVPQVSTPTLVEVSR_3
F1_LVAASQAALGL_2
F2_LVTDLTK_1
F4_QEPERNECFLQHK_3
F1_QNCELFEQLGEYK_2
F2_QTALVELVK_2
F4_QTALVELVK_2
F3_SLHTLFGDK_2
F1_TYETTLEK_2
F1_VPQVSTPTLVEVSR_2
F1_VPQVSTPTLVEVSR_3
F2_VPQVSTPTLVEVSR_2
outliers
beta
alpha
ratio
time
0.003
0.002
0.001
0.000
−0.001
−0.002
0
5
10
15
20
25
0.005
0.004
0.003
0.002
0.001
0.000

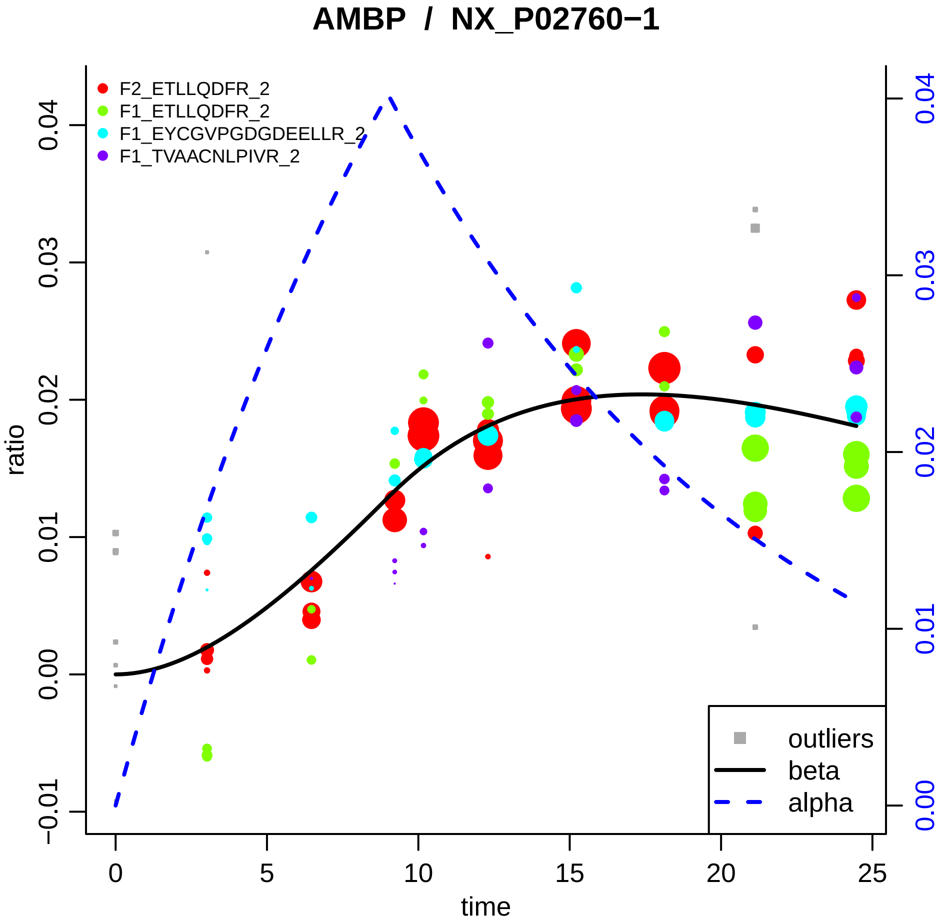

AMBP / NX_P02760−1
F2_ETLLQDFR_2
F1_ETLLQDFR_2
F1_EYCGVPGDGDEELLR_2
F1_TVAACNLPIVR_2
outliers
beta
alpha
ratio
time
−0.01
0.00
0.01
0.02
0.03
0.04
0
5
10
15
20
25

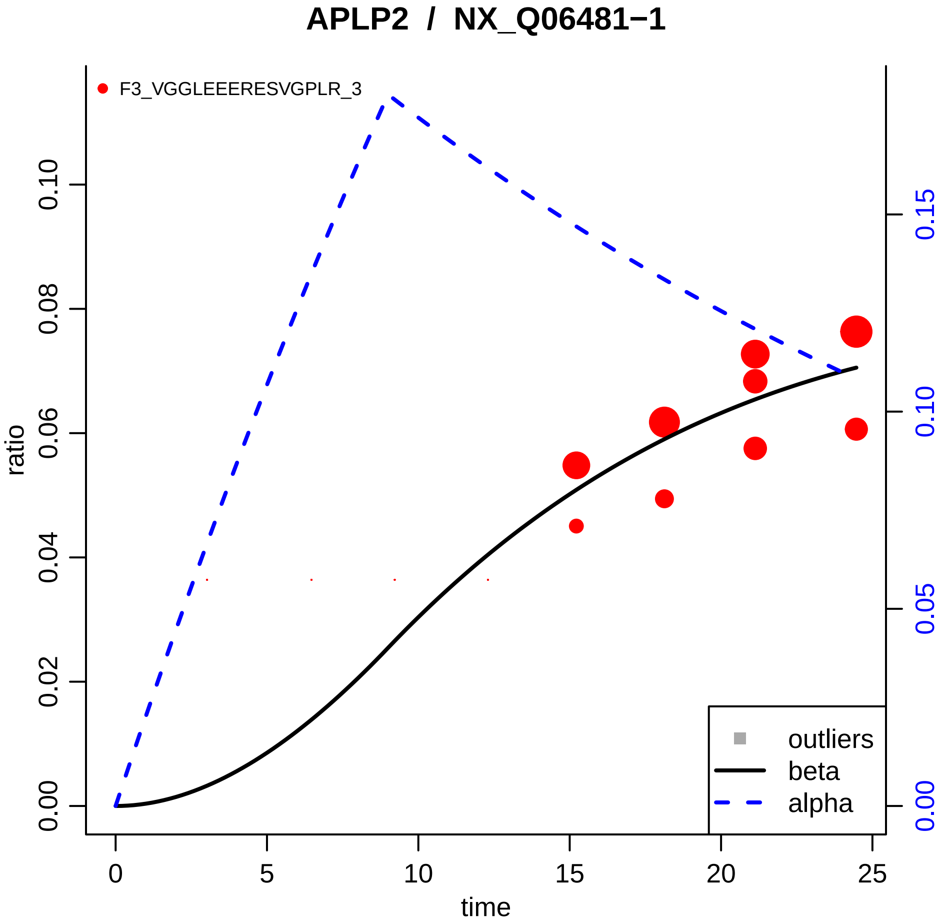

APLP2 / NX_Q06481−1
F3_VGGLEEERESVGPLR_3
ratio
0.00
0.02
0.04
0.06
0.08
0.10
0.00
0.05
0.10
0.15
0
5
10
15
20
25
time
outliers
beta
alpha

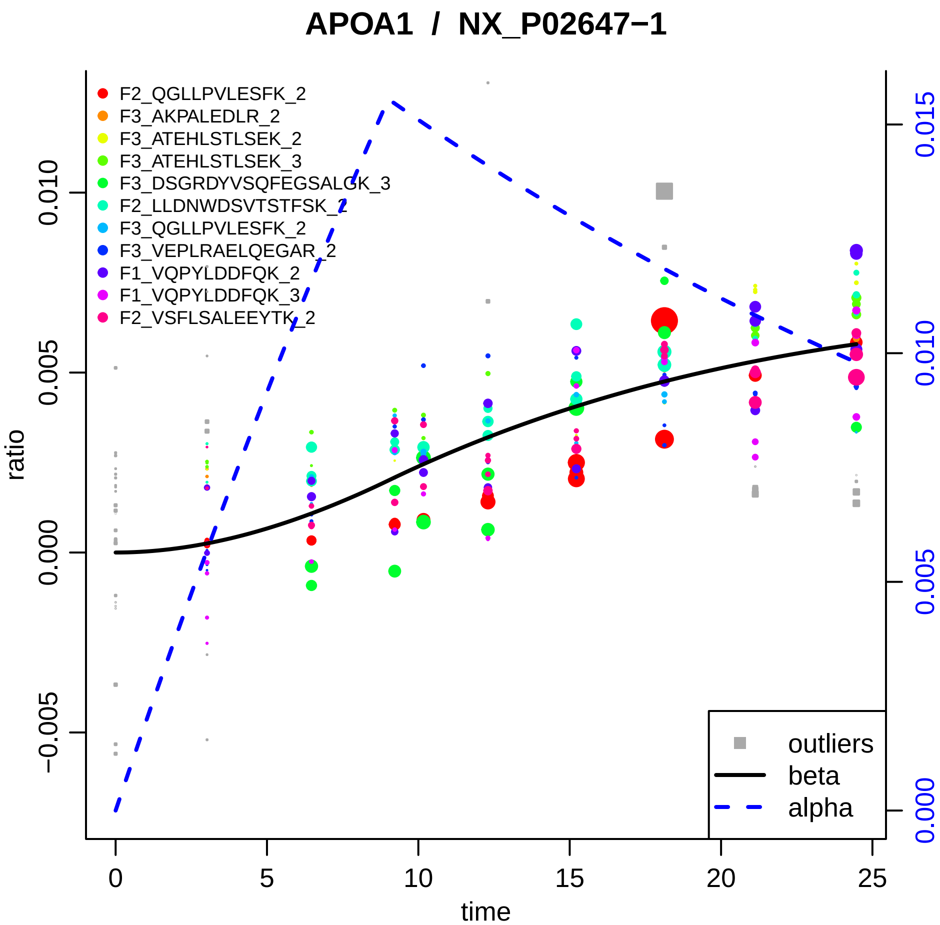

APOA1 / NX_P02647−1
F2_QGLLPVLESFK_2
F3_AKPALEDLR_2
F3_ATEHLSTLSEK_2
F3_ATEHLSTLSEK_3
F3_DSGRDYVSQFEGSALGK_3
F2_LLDNWDSVTSTFSK_2
F3_QGLLPVLESFK_2
F3_VEPLRAELQEGAR_2
F1_VQPYLDDFQK_2
F1_VQPYLDDFQK_3
F2_VSFLSALEEYTK_2
outliers
beta
alpha
ratio
time
−0.005
0.000
0.005
0.010
0.000
0.005
0.010
0.015
0
5
10
15
20
25

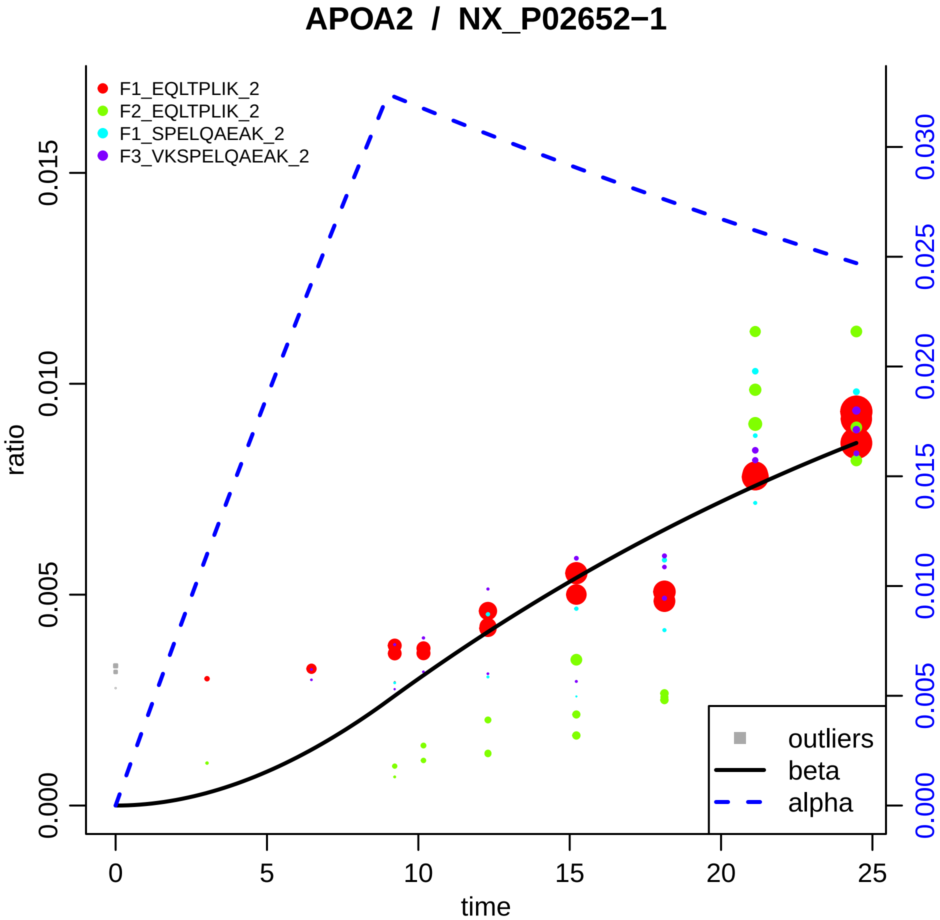
APOA2 / NX_P02652−1
F1_EQLTPLIK_2
F2_EQLTPLIK_2
F1_SPELQAEAK_2
F3_VKSPELQAEAK_2
outliers
beta
alpha
ratio
time
0.000
0.005
0.010
0.015
0
5
10
15
20
25
0.000
0.005
0.010
0.015
0.020
0.025
0.030

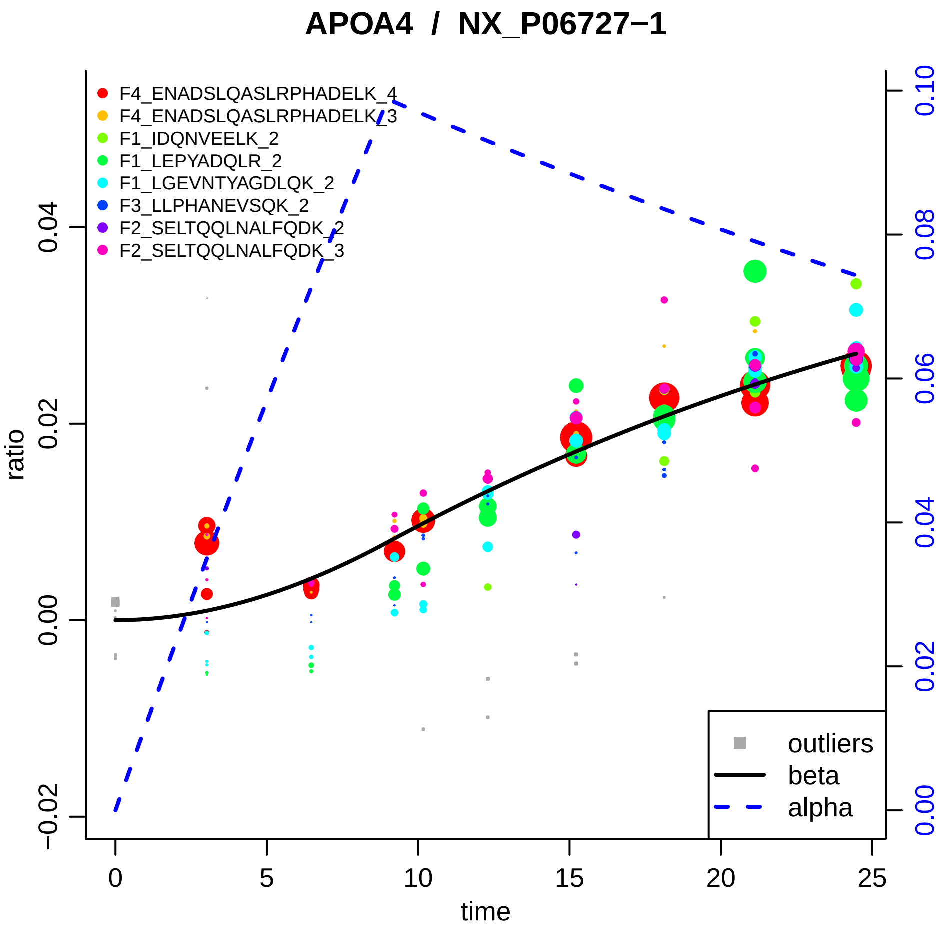
APOA4 / NX_P06727−1
F4_ENADSLQASLRPHADELK_4
F4_ENADSLQASLRPHADELK_3
F1_IDQNVEELK_2
F1_LEPYADQLR_2
F1_LGEVNTYAGDLQK_2
F3_LLPHANEVSQK_2
F2_SELTQQLNALFQDK_2
F2_SELTQQLNALFQDK_3
outliers
beta
alpha
ratio
time
−0.02
0.00
0.02
0.04
0
5
10
15
20
25
0.00
0.02
0.04
0.06
0.08
0.10

### APOB / NX_P04114−1

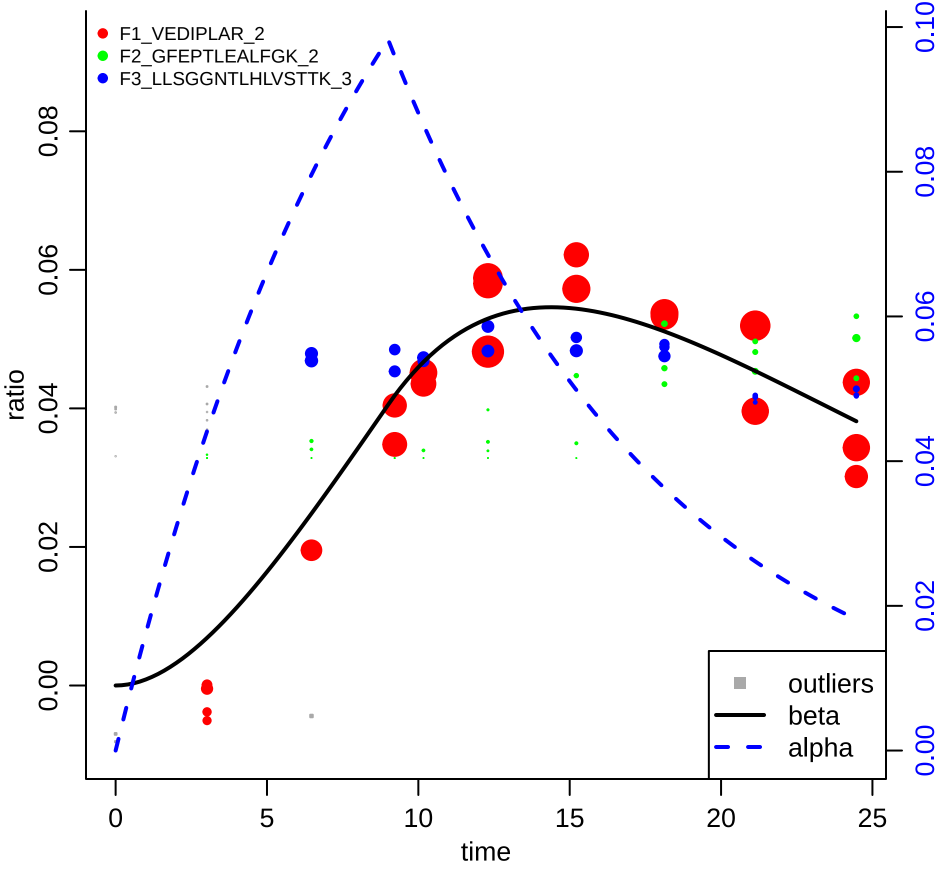

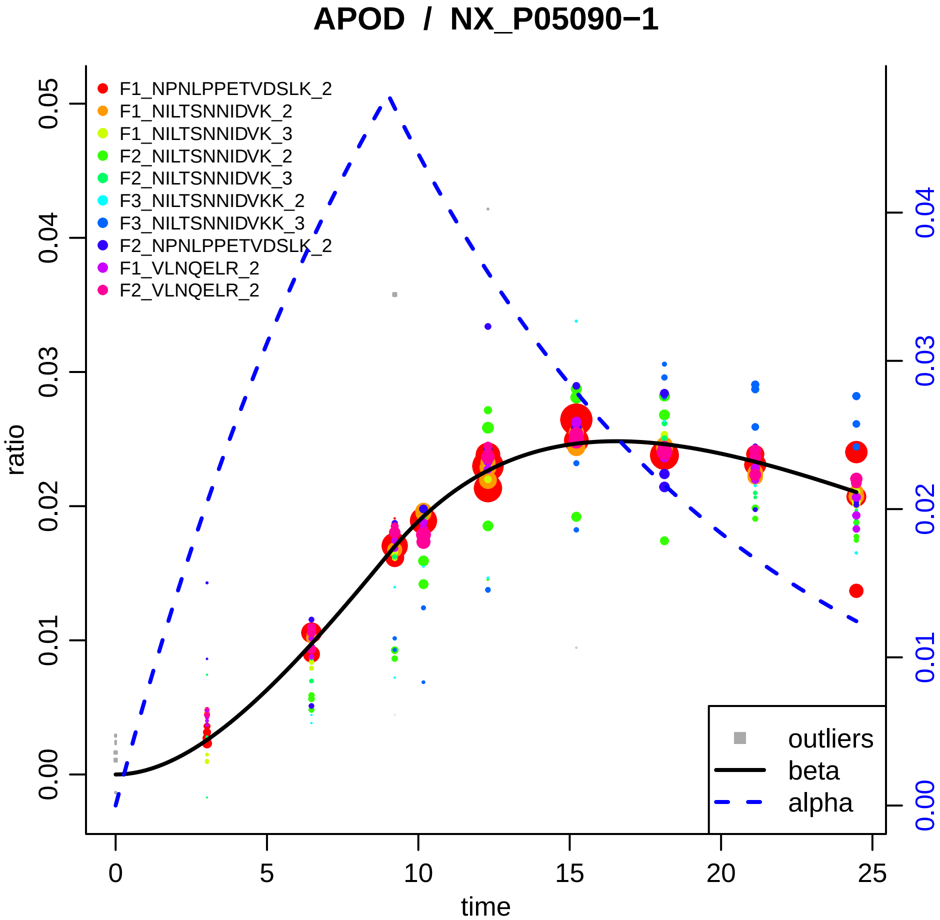

APOD / NX_P05090−1
F1_NPNLPPETVDSLK_2
F1_NILTSNNIDVK_2
F1_NILTSNNIDVK_3
F2_NILTSNNIDVK_2
F2_NILTSNNIDVK_3
F3_NILTSNNIDVKK_2
F3_NILTSNNIDVKK_3
F2_NPNLPPETVDSLK_2
F1_VLNQELR_2
F2_VLNQELR_2
outliers
beta
alpha
ratio
time
0.00
0.01
0.02
0.03
0.04
0.05
0
5
10
15
20
25

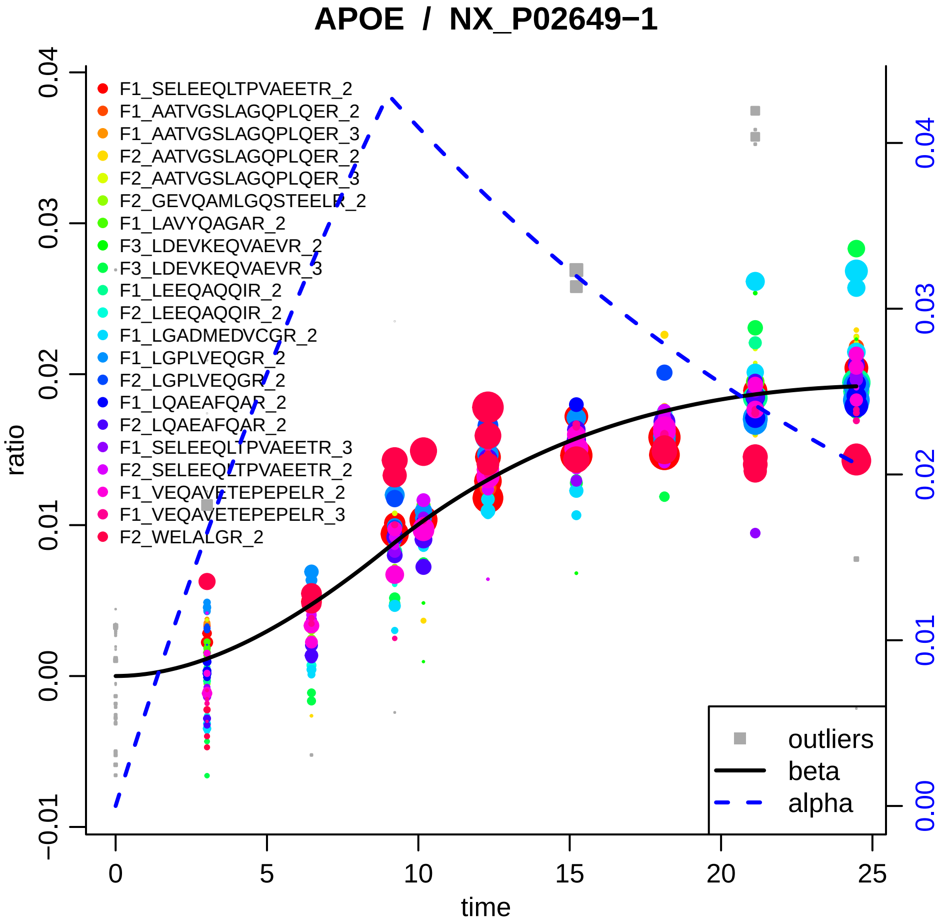
APOE / NX_P02649−1
F1_SELEEQLTPVAEETR_2
F1_AATVGSLAGQPLQER_2
F1_AATVGSLAGQPLQER_3
F2_AATVGSLAGQPLQER_2
F2_AATVGSLAGQPLQER_3
F2_GEVQAMLGQSTEELR_2
F1_LAVYQAGAR_2
F3_LDEVKEQVAEVR_2
F3_LDEVKEQVAEVR_3
F1_LEEQAQQIR_2
F2_LEEQAQQIR_2
F1_LGADMEDVCGR_2
F1_LGPLVEQGR_2
F2_LGPLVEQGR_2
F1_LQAEAFQAR_2
F2_LQAEAFQAR_2
F1_SELEEQLTPVAEETR_3
F2_SELEEQLTPVAEETR_2
F1_VEQAVETEPEPELR_2
F1_VEQAVETEPEPELR_3
F2_WELALGR_2
outliers
beta
alpha
ratio
time
−0.01
0.00
0.01
0.02
0.03
0.04
0
5
10
15
20
25

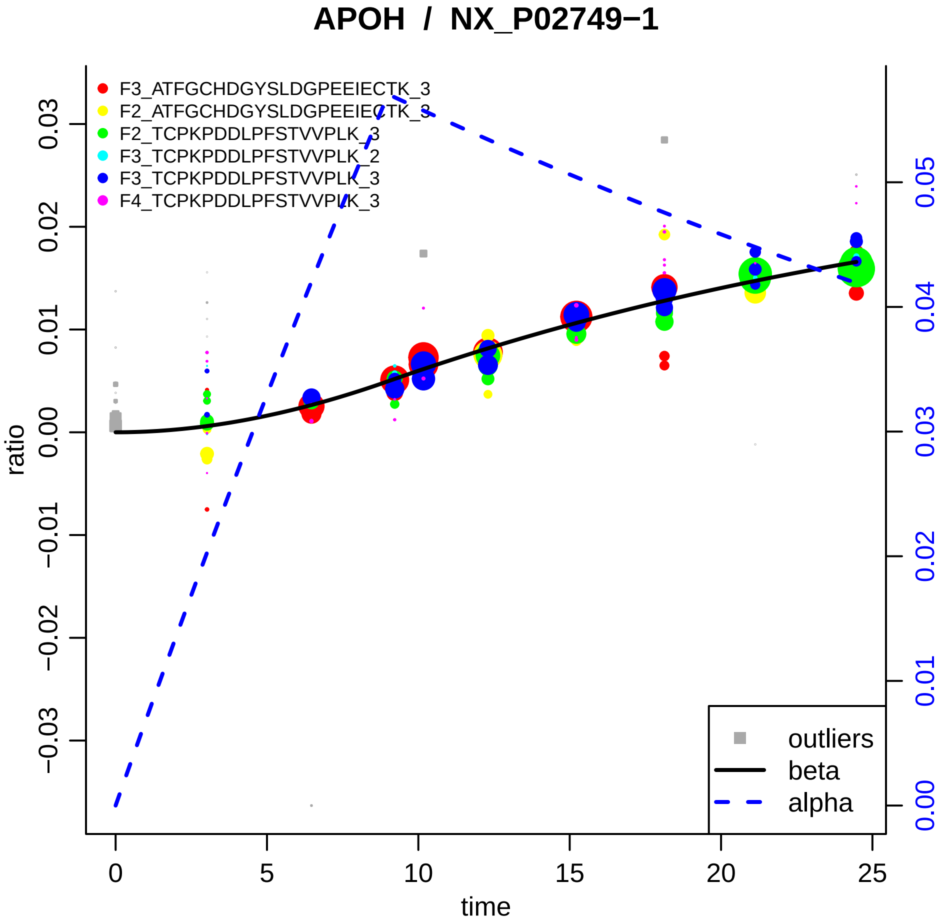

APOH / NX_P02749−1
F3_ATFGCHDGYSLDGPEEIECTK_3
F2_ATFGCHDGYSLDGPEEIECTK_3
F2_TCPKPDDLPFSTVVPLK_3
F3_TCPKPDDLPFSTVVPLK_2
F3_TCPKPDDLPFSTVVPLK_3
F4_TCPKPDDLPFSTVVPLK_3
outliers
beta
alpha
ratio
time
0
5
10
15
20
25
0.03
0.02
0.01
0.00
−0.01
−0.02
−0.03
0.05
0.04
0.03
0.02
0.01
0.00

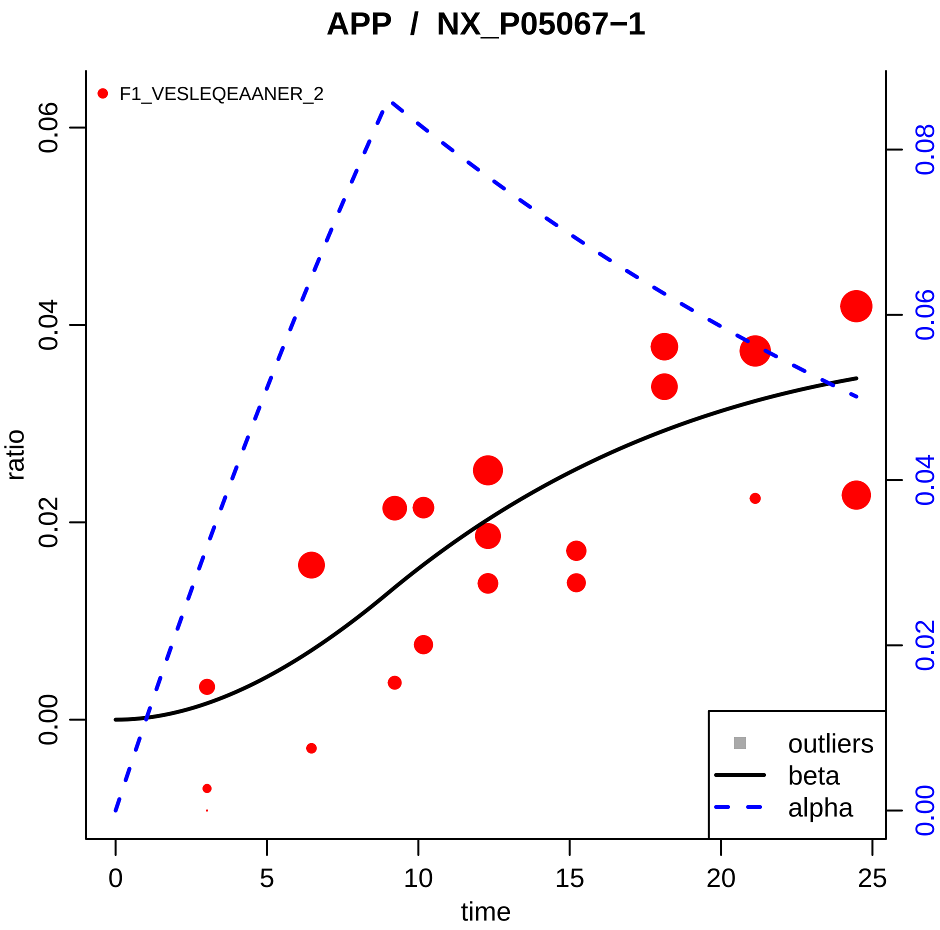

APP / NX_P05067−1
F1_VESLEQEAANER_2
ratio
0.00
0.02
0.04
0.06
time
0
5
10
15
20
25
0.00
0.02
0.04
0.06
0.08
outliers
beta
alpha

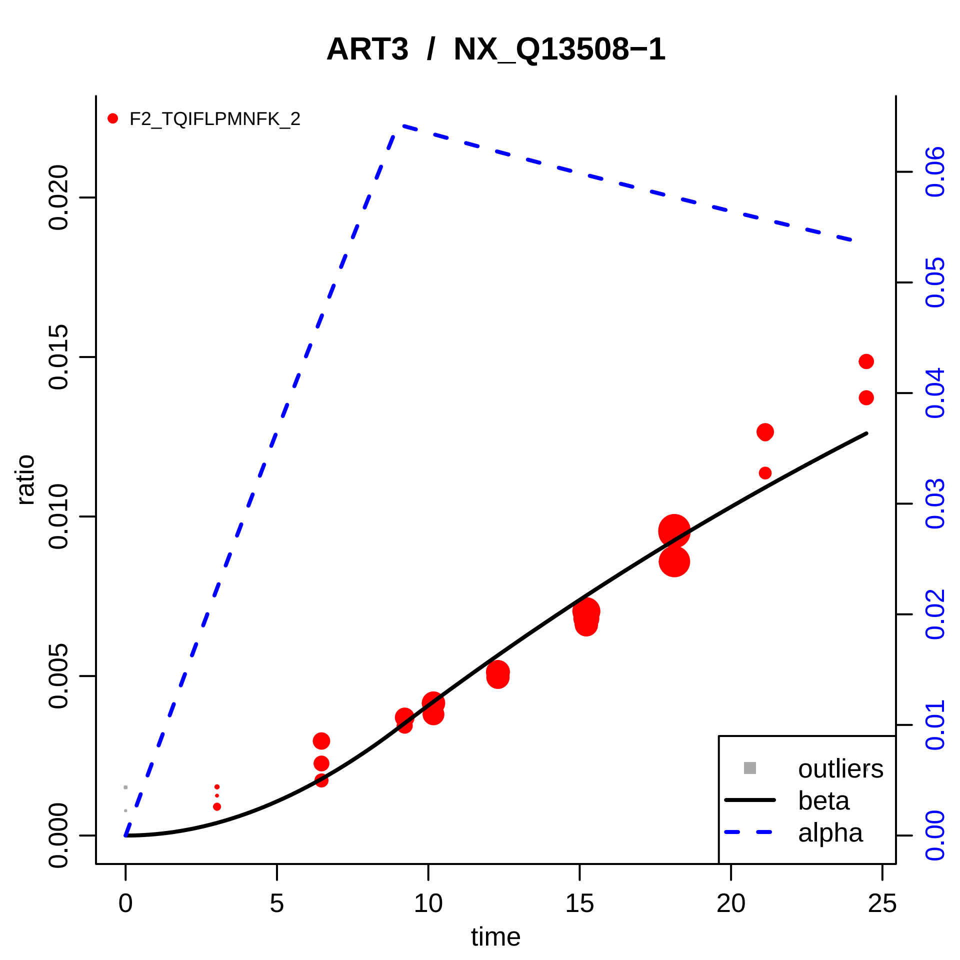

ART3 / NX_Q13508−1
F2_TQIFLPMNFK_2
ratio
0.000
0.005
0.010
0.015
0.020
time
0
5
10
15
20
25
0.00
0.01
0.02
0.03
0.04
0.05
0.06
outliers
beta
alpha

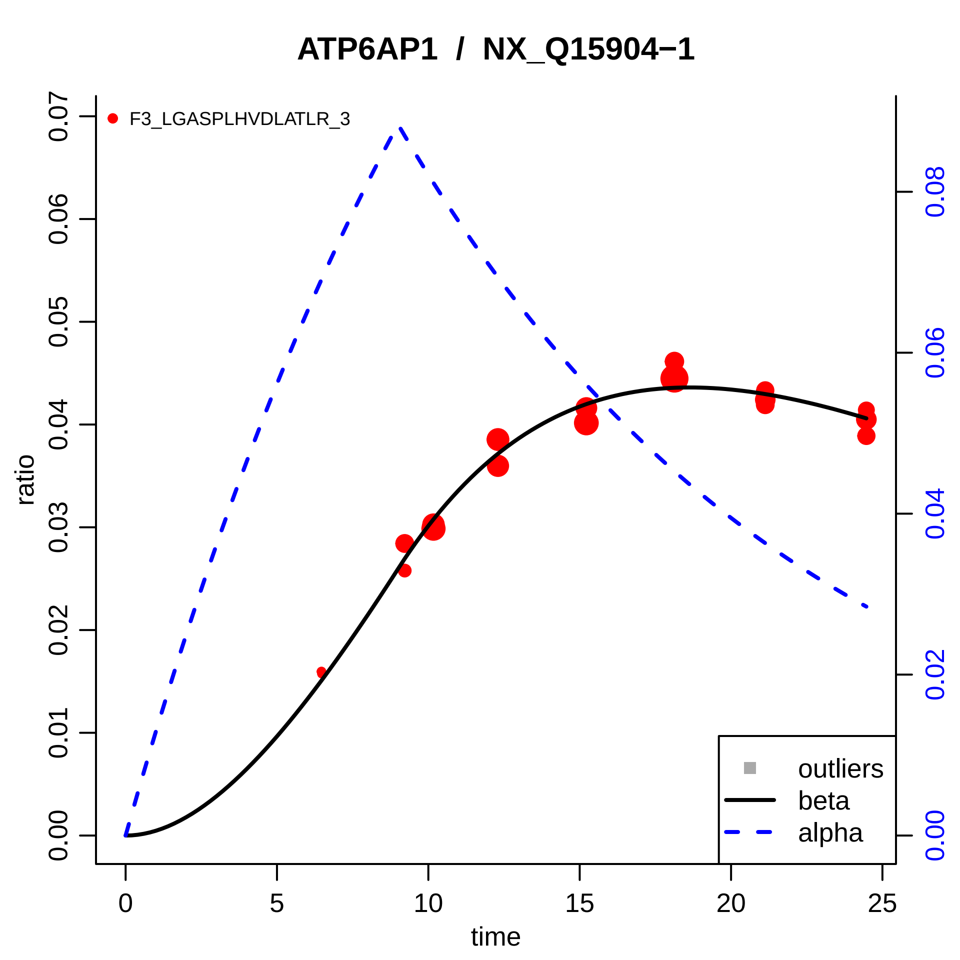

ATP6AP1 / NX_Q15904−1
F3_LGASPLHVDLATLR_3
ratio
time
0.00
0.01
0.02
0.03
0.04
0.05
0.06
0.07
0
5
10
15
20
25
0.00
0.02
0.04
0.06
0.08
outliers
beta
alpha

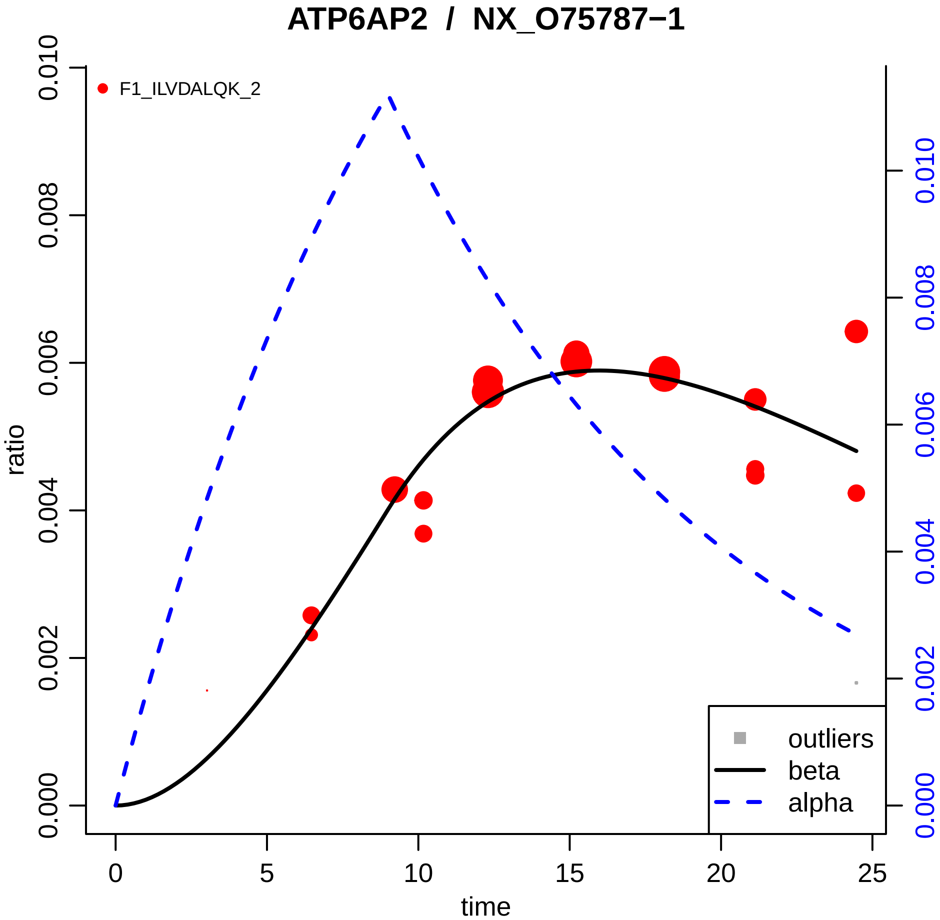
ATP6AP2 / NX_O75787−1
F1_ILVDALQK_2
ratio
time
outliers
beta
alpha
0.000
0.002
0.004
0.006
0.008
0.010
0
5
10
15
20
25

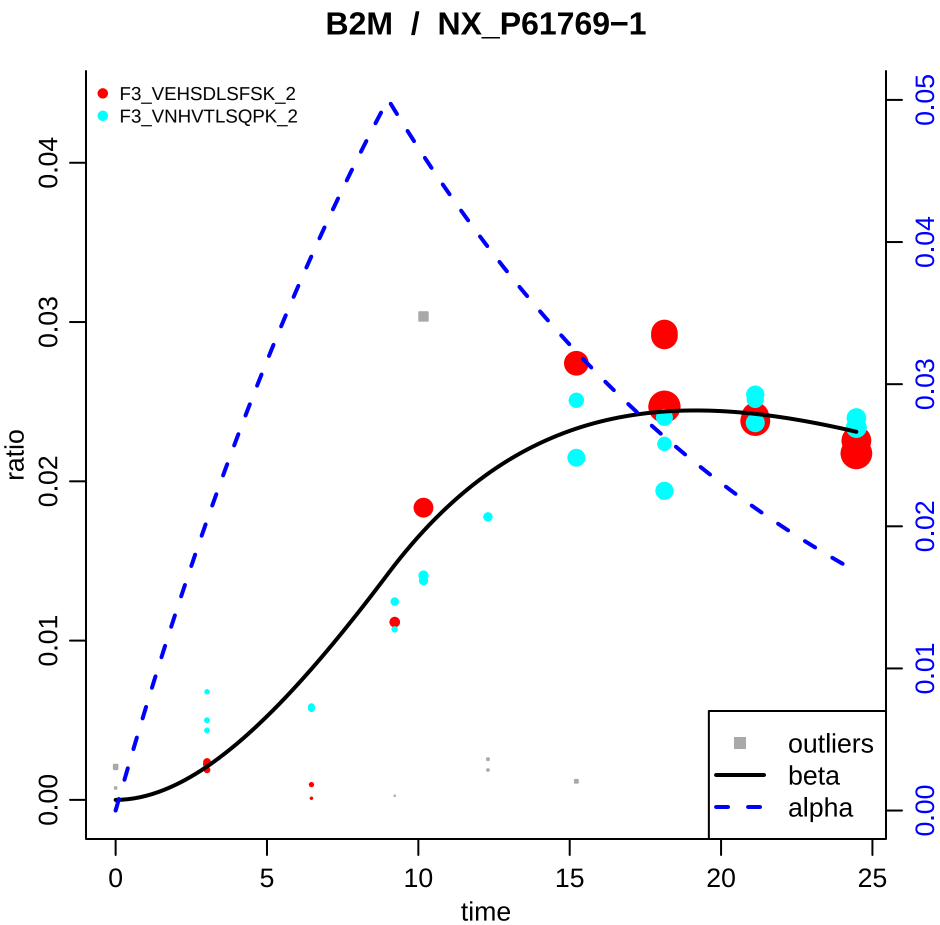

B2M / NX_P61769−1
F3_VEHSDLSFSK_2
F3_VNHVTLSQPK_2
ratio
time
0.00
0.01
0.02
0.03
0.04
0.05
0
5
10
15
20
25
outliers
beta
alpha

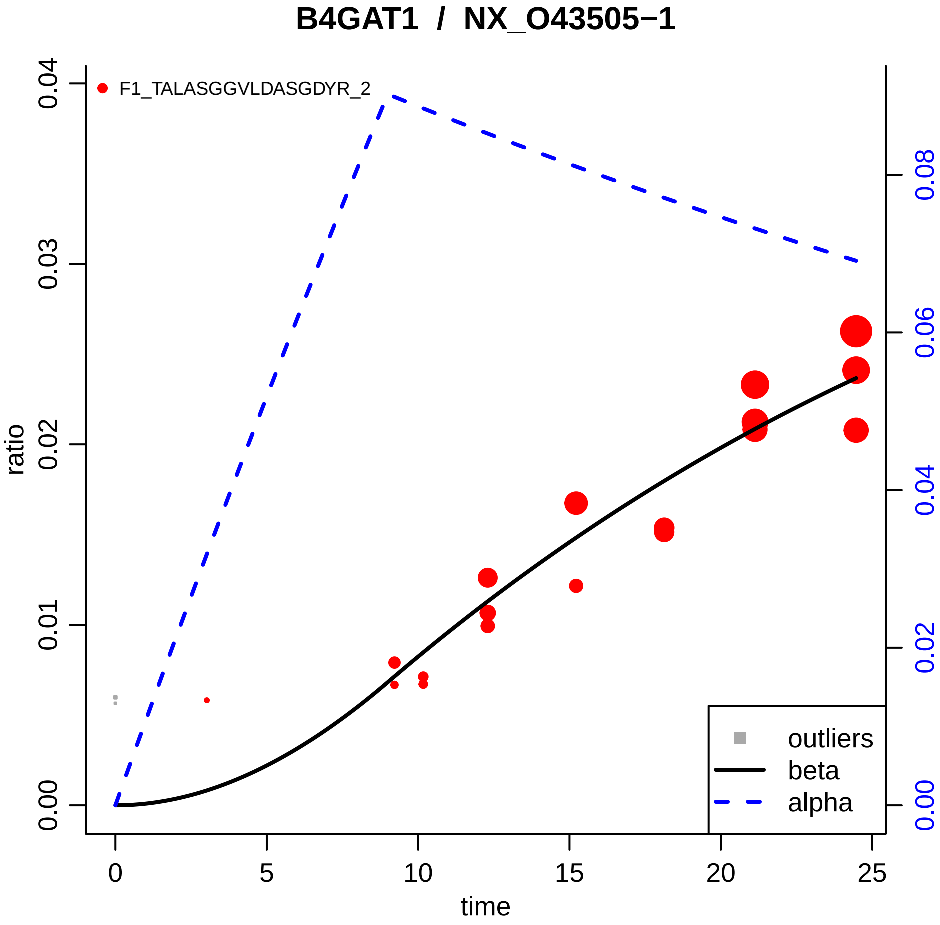

B4GAT1 / NX_O43505−1
F1_TALASGGVLDASGDYR_2
ratio
0.00
0.01
0.02
0.03
0.04
time
0
5
10
15
20
25
0.00
0.02
0.04
0.06
0.08
outliers
beta
alpha

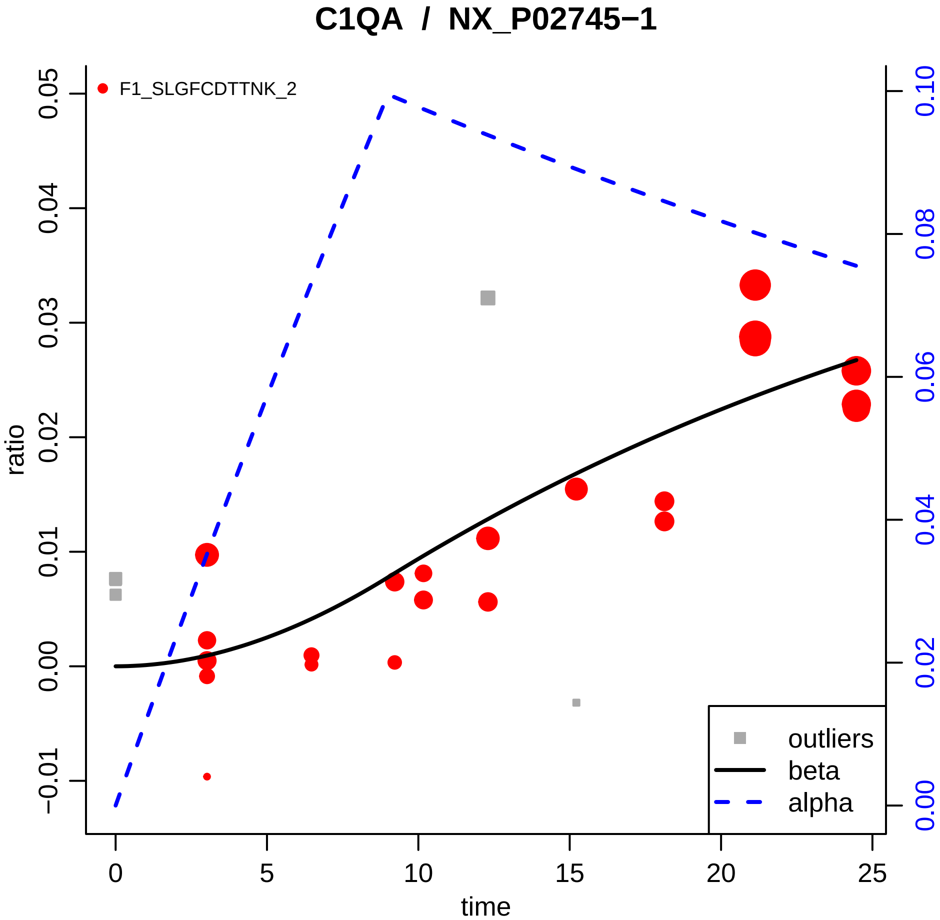

C1QA / NX_P02745−1
F1_SLGFCDTTNK_2
ratio
time
0.05
0.04
0.03
0.02
0.01
0.00
−0.01
0.10
0.08
0.06
0.04
0.02
0.00
0
5
10
15
20
25
outliers
beta
alpha

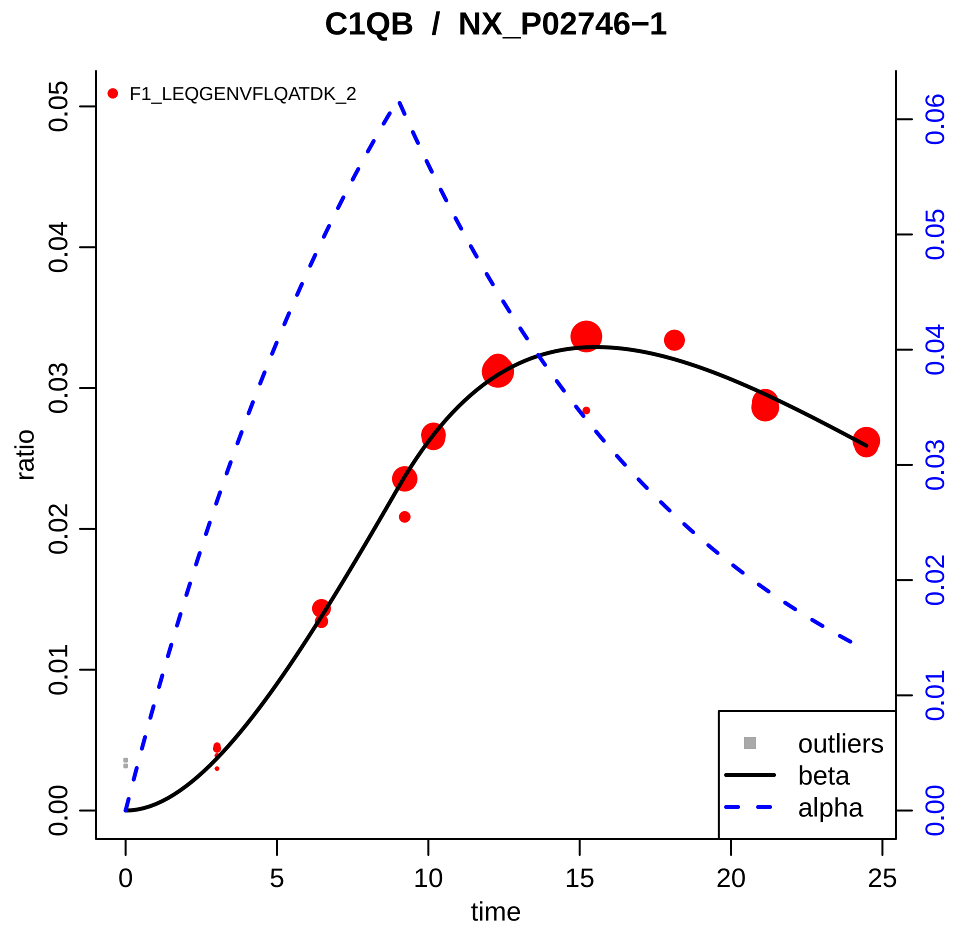

C1QB / NX_P02746−1
F1_LEQGENVFLQATDK_2
ratio
time
0.00
0.01
0.02
0.03
0.04
0.05
0.06
0
5
10
15
20
25
outliers
beta
alpha

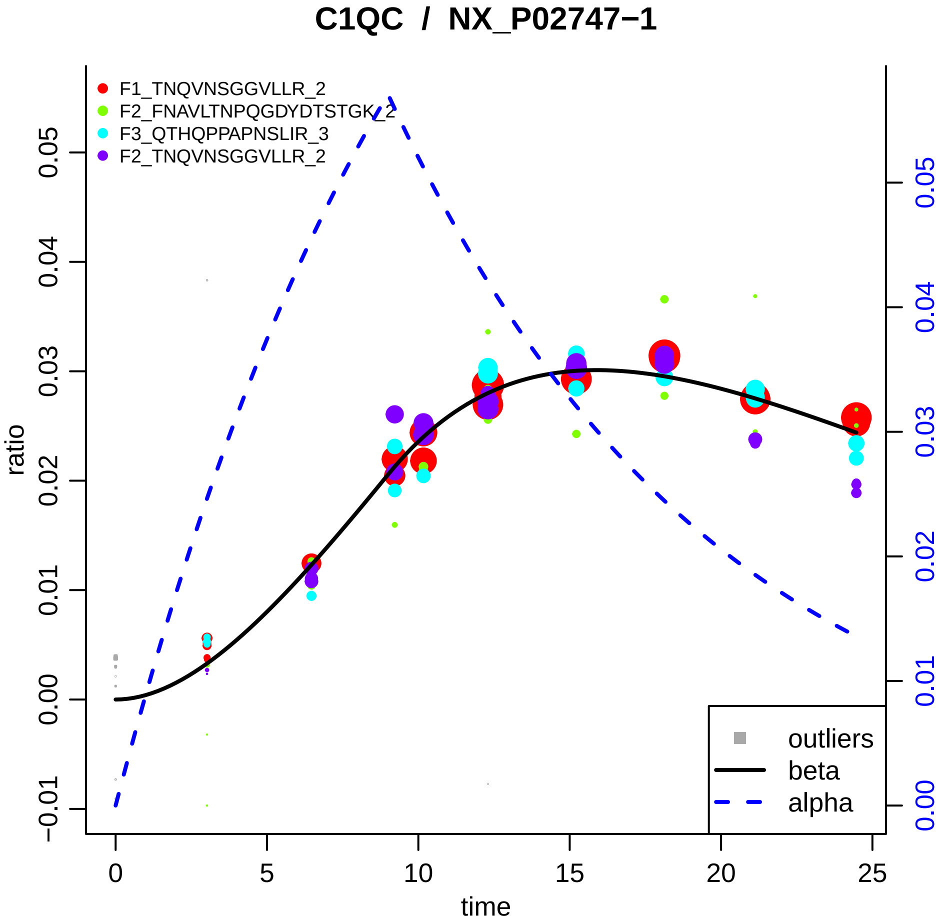
C1QC / NX_P02747−1
F1_TNQVNSGGVLLR_2
F2_FNAVLTNPQGDYDTSTGK_2
F3_QTHQPPAPNSLIR_3
F2_TNQVNSGGVLLR_2
outliers
beta
alpha
ratio
time
0
5
10
15
20
25
−0.01
0.00
0.01
0.02
0.03
0.04
0.05
0.00
0.01
0.02
0.03
0.04
0.05

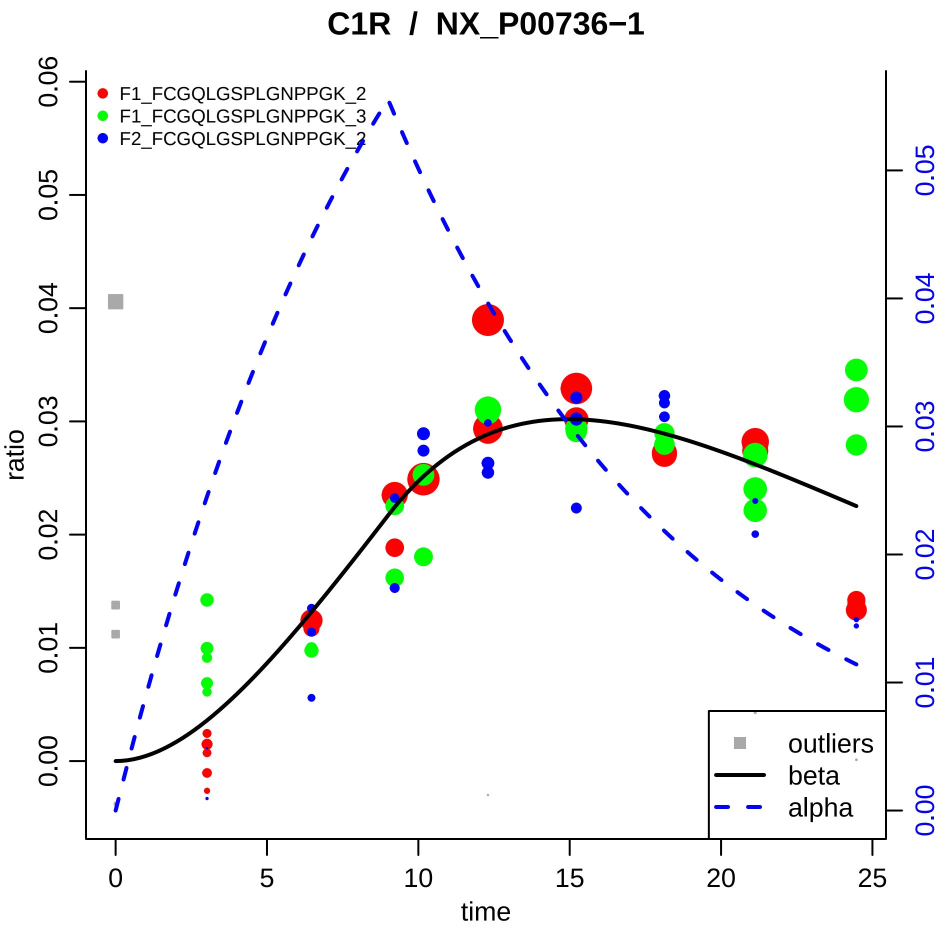

C1R / NX_P00736−1
F1_FCGQLGSPLGNPPGK_2
F1_FCGQLGSPLGNPPGK_3
F2_FCGQLGSPLGNPPGK_2
ratio
time
0.00
0.01
0.02
0.03
0.04
0.05
0.06
0
5
10
15
20
25
outliers
beta
alpha

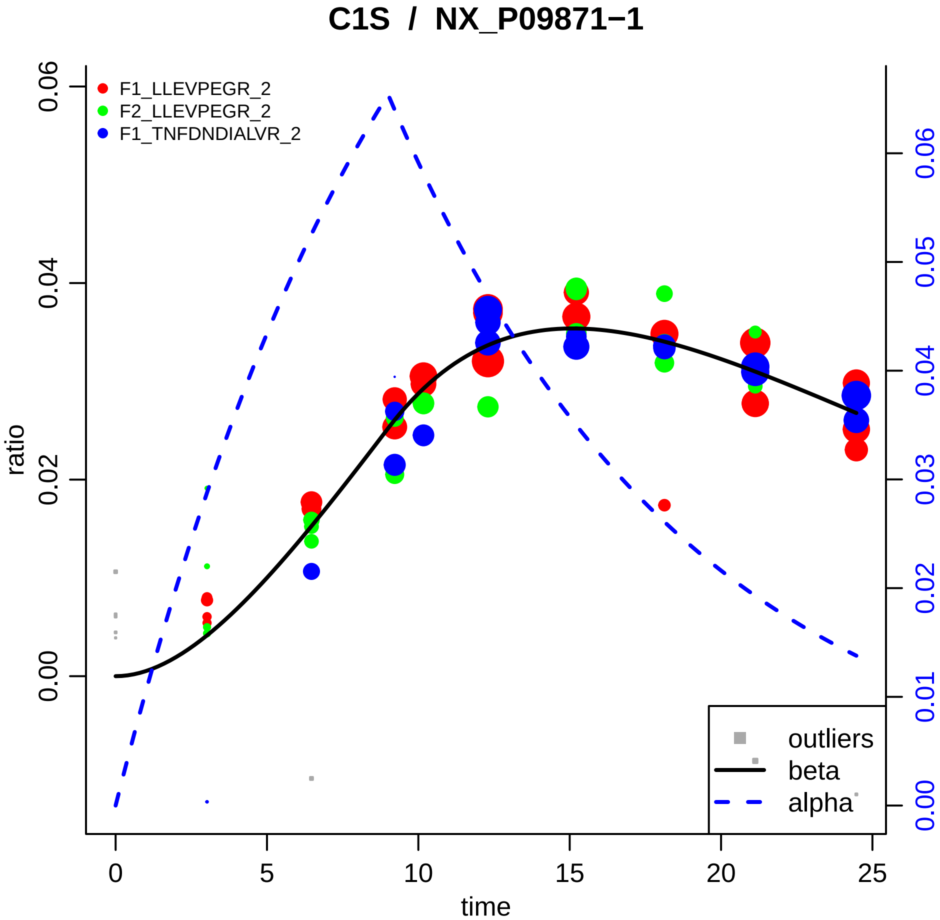

C1S / NX_P09871−1
F1_LLEVPEGR_2
F2_LLEVPEGR_2
F1_TNFDNDIALVR_2
ratio
0.00
0.02
0.04
0.06
time
0
5
10
15
20
25
0.00
0.01
0.02
0.03
0.04
0.05
0.06
outliers
beta
alpha

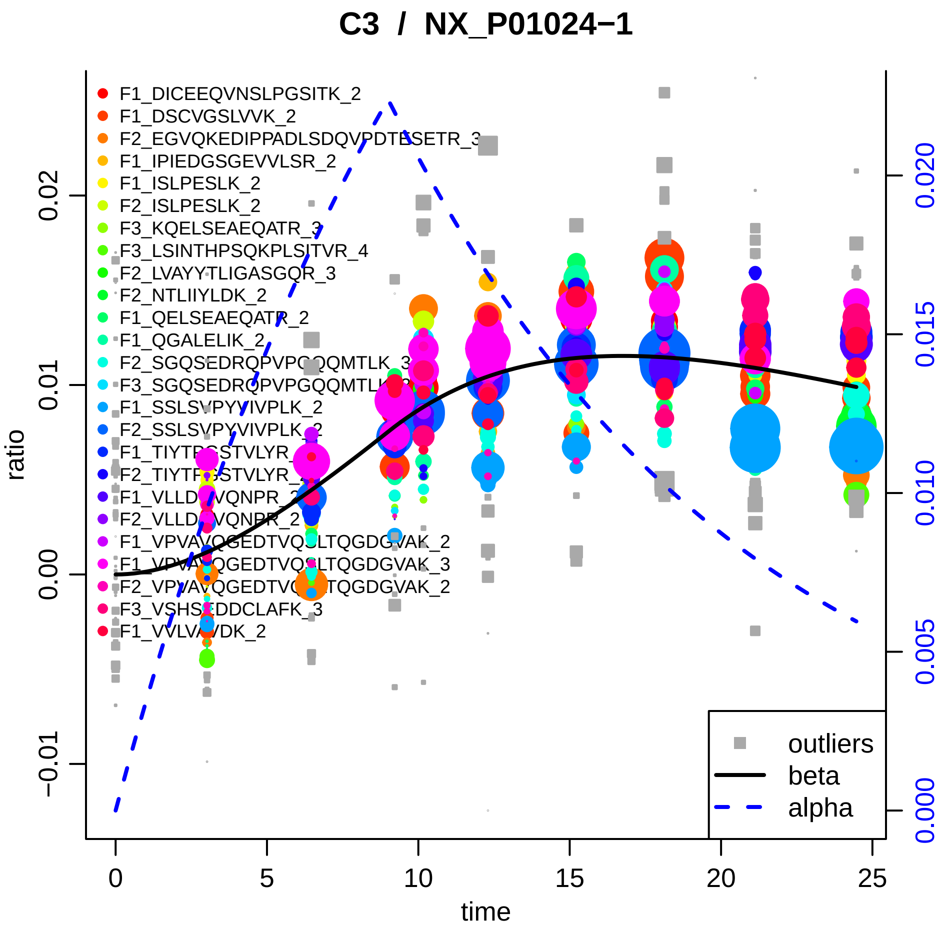
C3 / NX_P01024−1
F1_DICEEQVNSLPGSITK_2
F1_DSCVGSLVVK_2
F2_EGVQKEDIPPADLSDQVPDTESETR_3
F1_IPIEDGSGEVVLSR_2
F1_ISLPESLK_2
F2_ISLPESLK_2
F3_KQELSEAEQATR_3
F3_LSINTHPSQKPLSITVR_4
F2_LVAYYTLIGASGQR_3
F2_NTLIIYLDK_2
F1_QELSEAEQATR_2
F1_QGALELIK_2
F2_SGQSEDRQPVPGQQMTLK_3
F1_SSLSVPYVIVPLK_2
F2_SSLSVPYVIVPLK_2
F2_VLLDGVQNPR_2
F1_VPVAVQGEDTVQSLTQGDGVAK_2
F1_VPVAVQGEDTVQSLTQGDGVAK_3
F2_VPVAVQGEDTVQSLTQGDGVAK_2
F3_VSHSEDDCLAFK_3
F1_VVLVAVDK_2
outliers
beta
alpha
ratio
time
0
5
10
15
20
25
−0.01
0.00
0.01
0.02
0.000
0.005
0.010
0.015
0.020

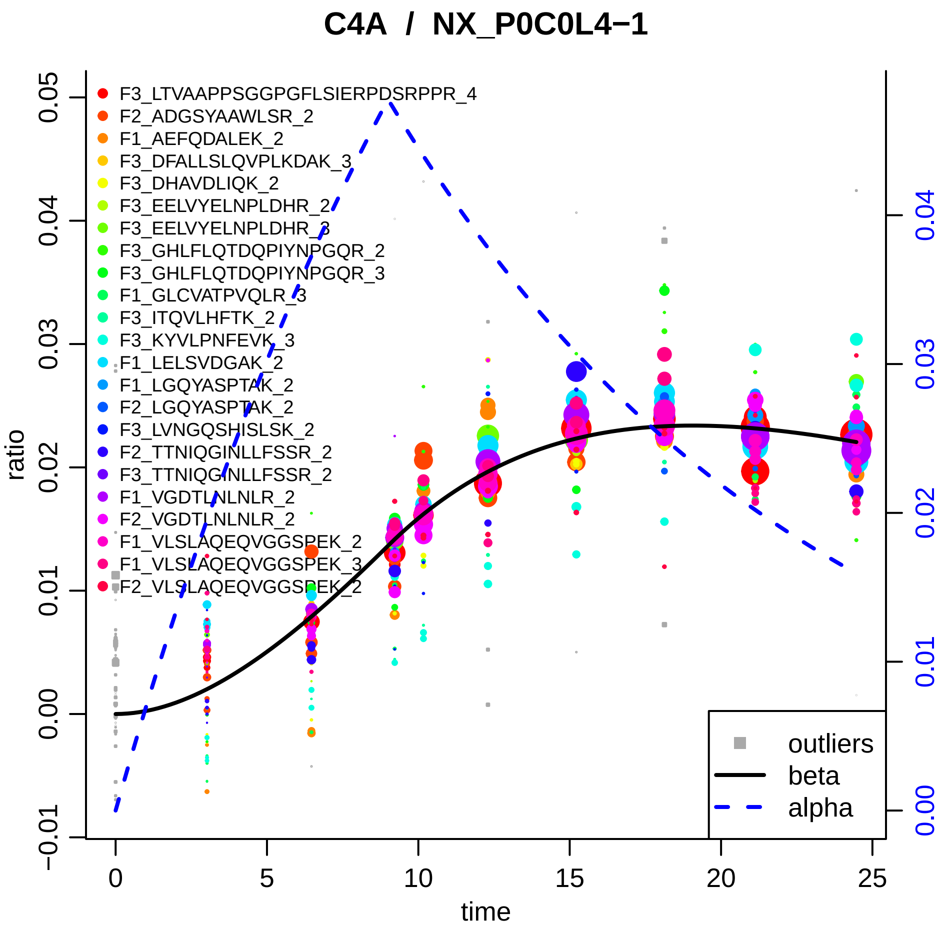

C4A / NX_P0C0L4−1
F3_LTVAAPPSGGPGFLSIERPDSRPPR_4
F2_ADGSYAAWLSR_2
F1_AEFQDALEK_2
F3_DFALLSLQVPLKDAK_3
F3_DHAVDLIQK_2
F3_EELVYELNPLDHR_2
F3_EELVYELNPLDHR_3
F3_GHLFLQTDQPIYNPGQR_2
F3_GHLFLQTDQPIYNPGQR_3
F1_GLCVATPVQLR_3
F3_ITQVLHFTK_2
F3_KYVLPNFEVK_3
F1_LELSVDGAK_2
F1_LGQYASPTAK_2
F2_LGQYASPTAK_2
F3_LVNGQSHISLSK_2
F2_TTNIQGINLLFSSR_2
F3_TTNIQGINLLFSSR_2
F1_VGDTLNLNLR_2
F2_VGDTLNLNLR_2
F1_VLSLAQEQVGGSPEK_2
F1_VLSLAQEQVGGSPEK_3
F2_VLSLAQEQVGGSPEK_2
outliers
beta
alpha
ratio
time
0.05
0.04
0.03
0.02
0.01
0.00
−0.01
0
5
10
15
20
25
0.04
0.03
0.02
0.01
0.00

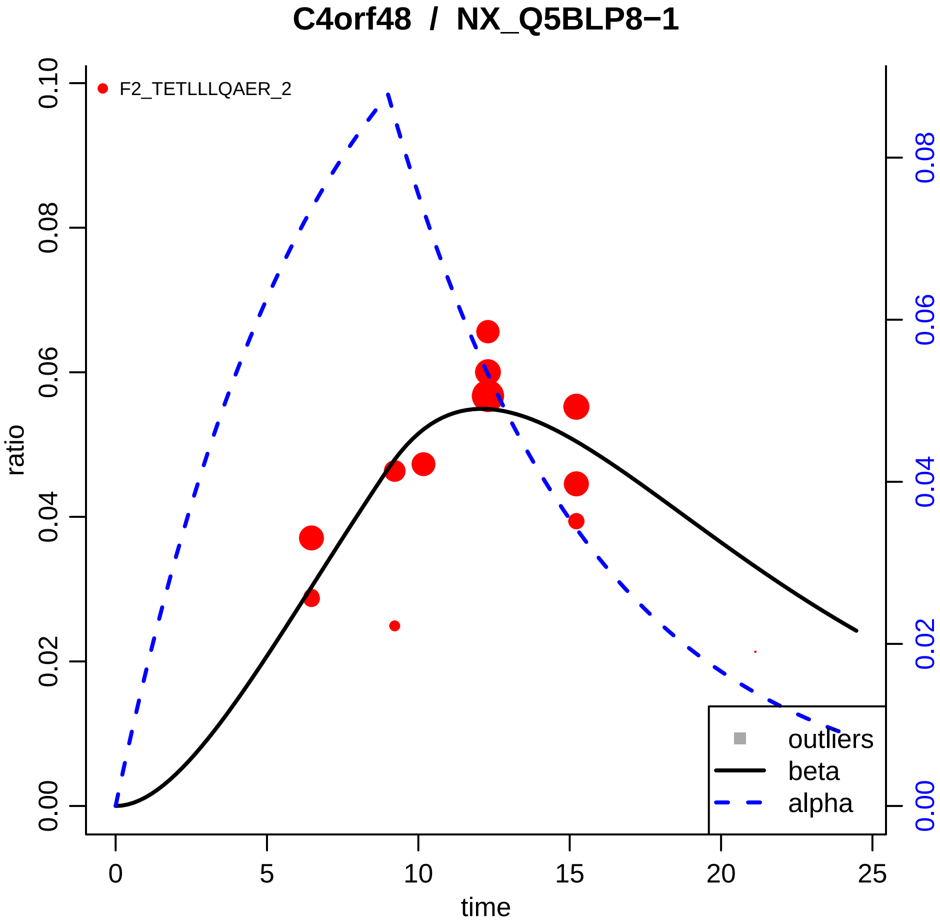

C4orf48 / NX_Q5BLP8−1
F2_TETLLLQAER_2
ratio
time
0.00
0.02
0.04
0.06
0.08
0.10
0
5
10
15
20
25
outliers
beta
alpha

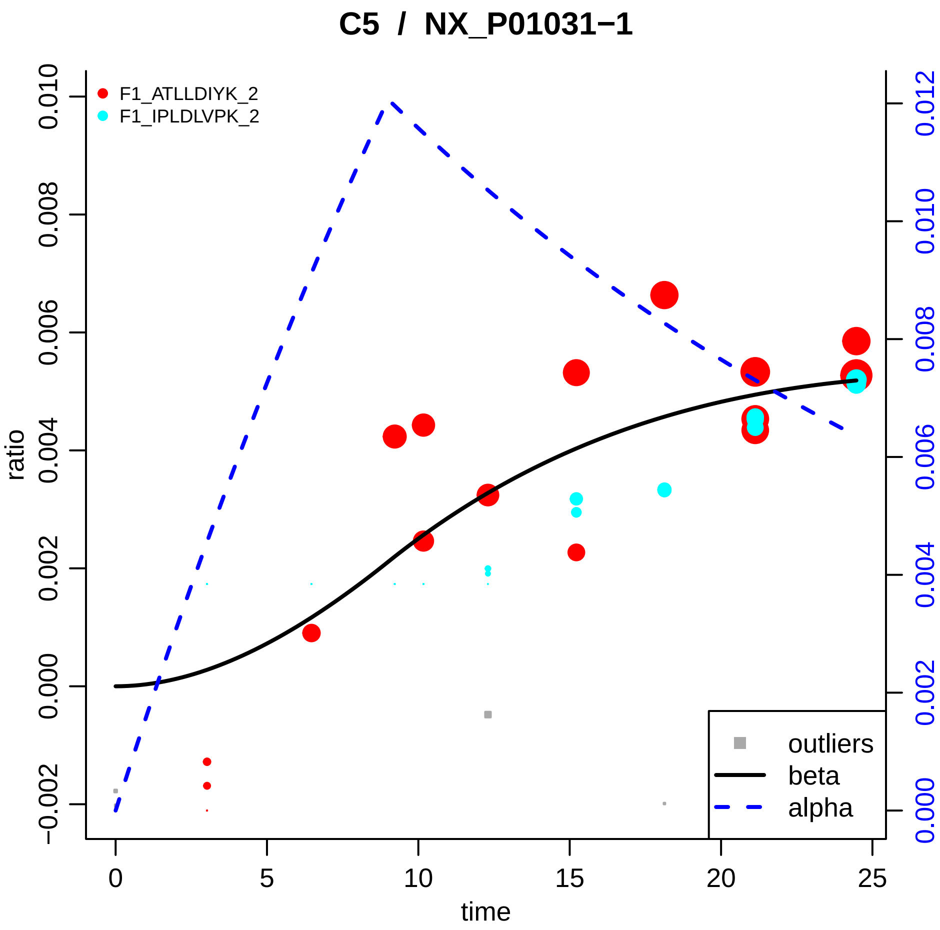
C5 / NX_P01031−1
F1_ATLLDIYK_2
F1_IPLDLVPK_2
ratio
time
0.010
0.008
0.006
0.004
0.002
0.000
−0.002
0.012
0.010
0.008
0.006
0.004
0.002
0.000
0
5
10
15
20
25
outliers
beta
alpha

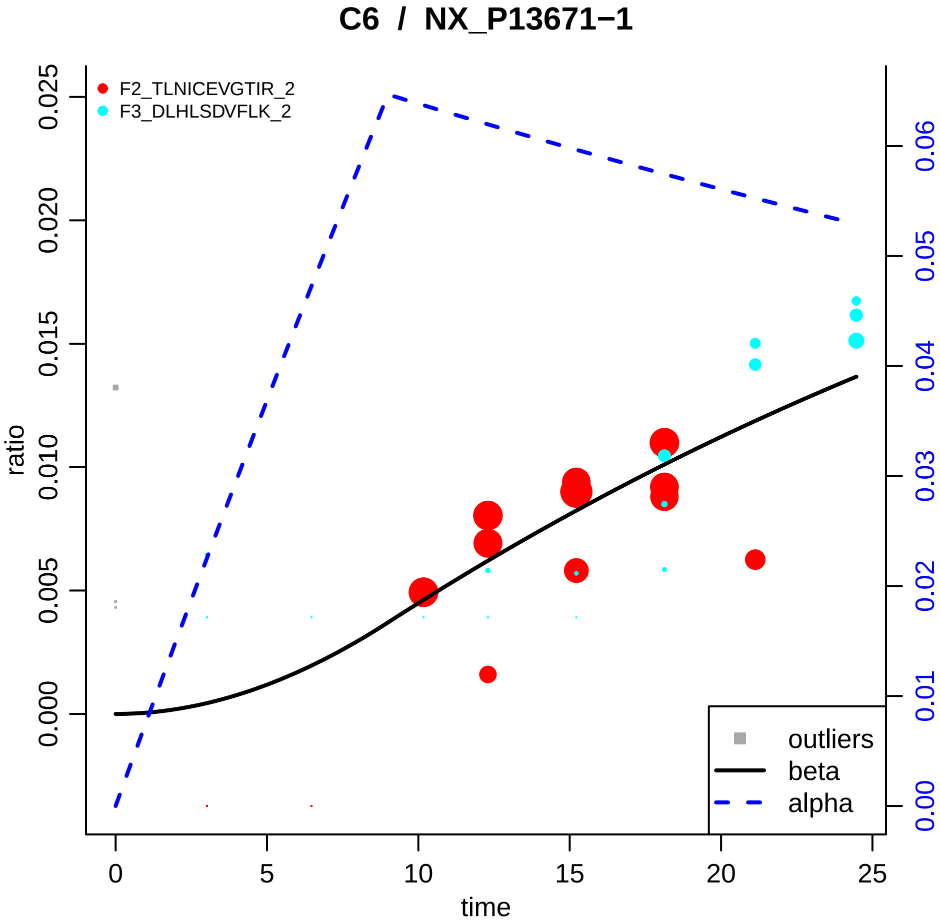
C6 / NX_P13671−1
F2_TLNICEVGTIR_2
F3_DLHLSDVFLK_2
ratio
0.000
0.005
0.010
0.015
0.020
0.025
0.00
0.01
0.02
0.03
0.04
0.05
0.06
time
0
5
10
15
20
25
outliers
beta
alpha

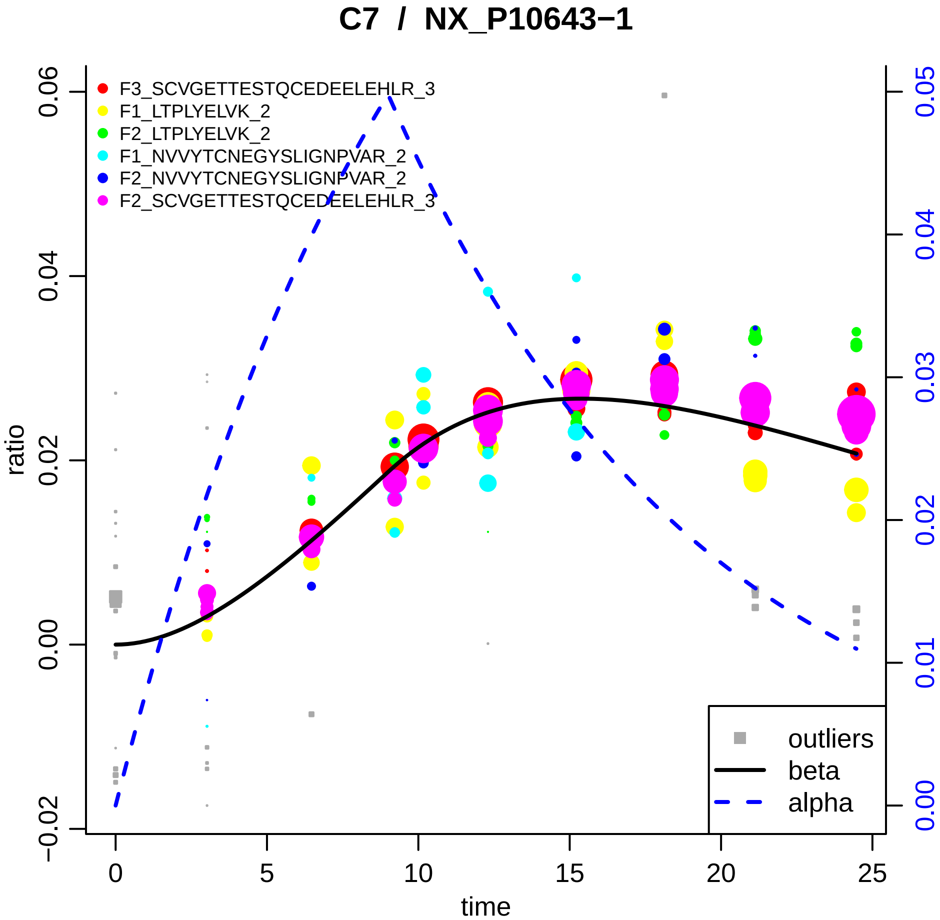

C7 / NX_P10643−1
F3_SCVGETTESTQCEDEELEHLR_3
F1_LTPLYELVK_2
F2_LTPLYELVK_2
F1_NVVYTCNEGYSLIGNPVAR_2
F2_NVVYTCNEGYSLIGNPVAR_2
F2_SCVGETTESTQCEDEELEHLR_3
outliers
beta
alpha
ratio
time
−0.02
0.00
0.02
0.04
0.06
0.00
0.01
0.02
0.03
0.04
0.05
0
5
10
15
20
25

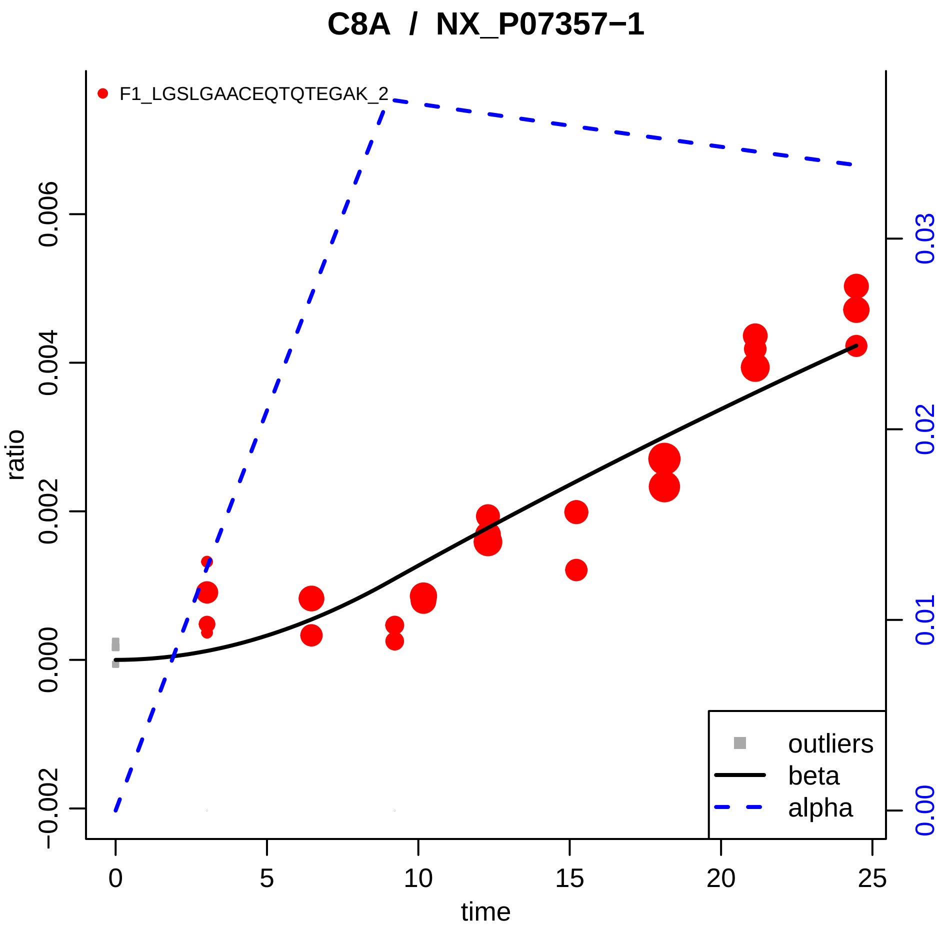
C8A / NX_P07357−1
F1_LGSLGAACEQTQTEGAK_2
ratio
time
0.006
0.004
0.002
0.000
−0.002
0.03
0.02
0.01
0.00
0
5
10
15
20
25
outliers
beta
alpha

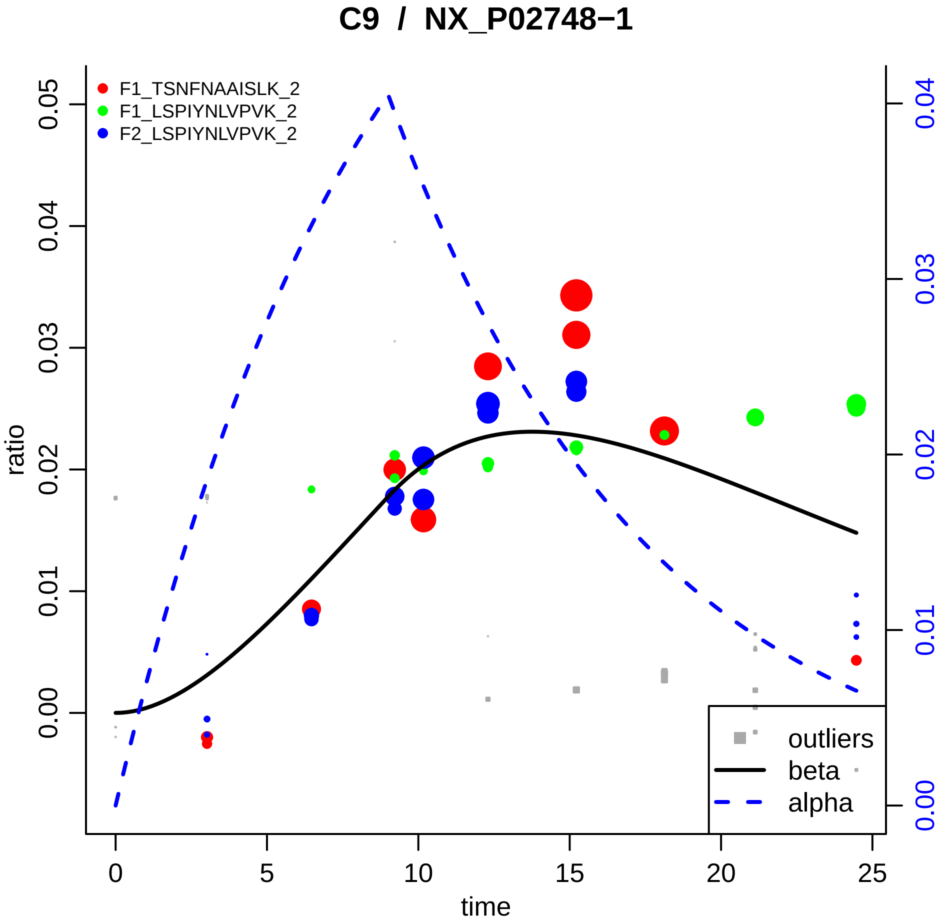

C9 / NX_P02748−1
F1_TSNFNAAISLK_2
F1_LSPIYNLVPVK_2
F2_LSPIYNLVPVK_2
ratio
time
0.00
0.01
0.02
0.03
0.04
0.05
0
5
10
15
20
25
0.00
0.01
0.02
0.03
0.04
outliers
beta
alpha

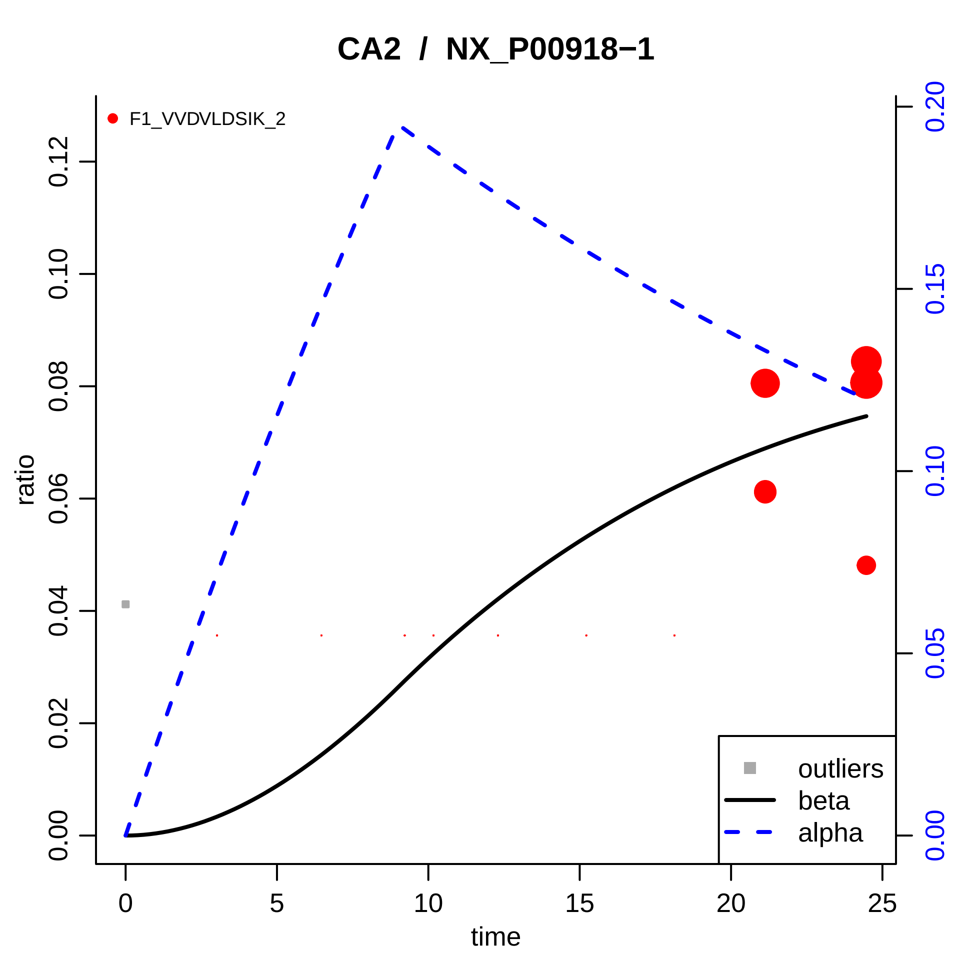

CA2 / NX_P00918−1
F1_VVDVLDSIK_2
ratio
0.00
0.02
0.04
0.06
0.08
0.10
0.12
0.00
0.05
0.10
0.15
0.20
time
0
5
10
15
20
25
outliers
beta
alpha

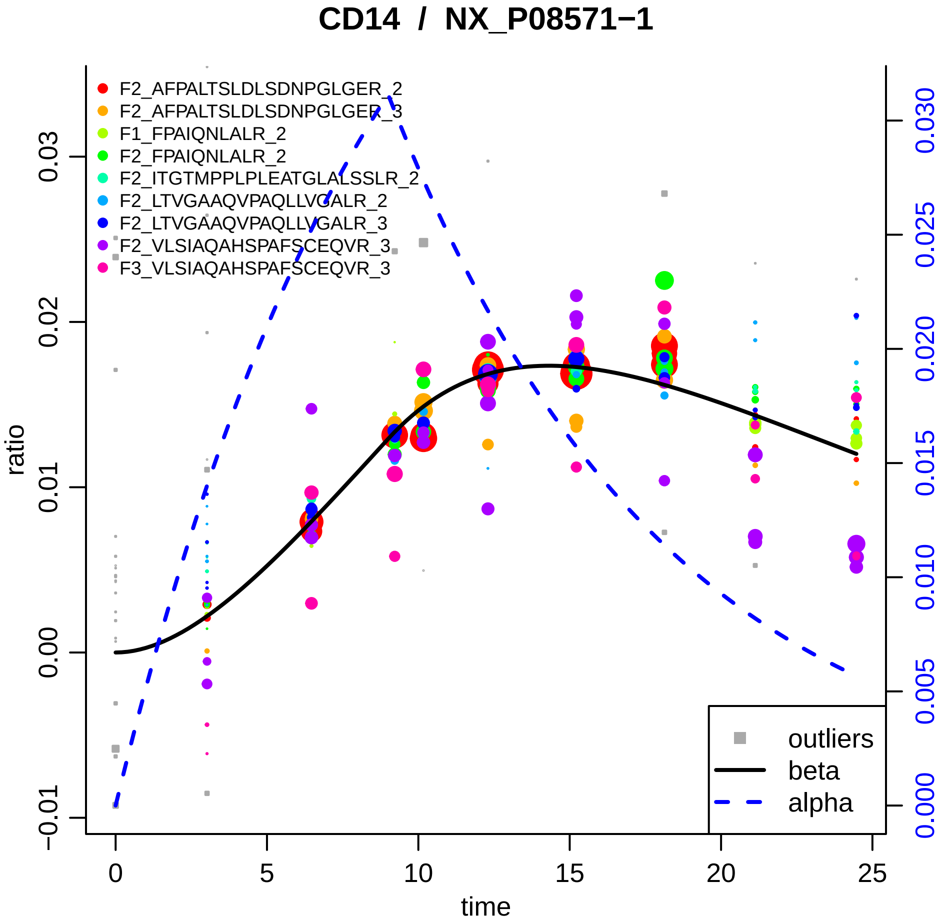

CD14 / NX_P08571−1
F2_AFPALTSLDLSDNPGLGER_2
F2_AFPALTSLDLSDNPGLGER_3
F1_FPAIQNLALR_2
F2_FPAIQNLALR_2
F2_ITGTMPPLPLEATGLALSSLR_2
F2_LTVGAAQVPAQLLVGALR_2
F2_LTVGAAQVPAQLLVGALR_3
F2_VLSIAQAHSPAFSCEQVR_3
F3_VLSIAQAHSPAFSCEQVR_3
outliers
beta
alpha
ratio
time
0
5
10
15
20
25
−0.01
0.00
0.01
0.02
0.03
0.000
0.005
0.010
0.015
0.020
0.025
0.030

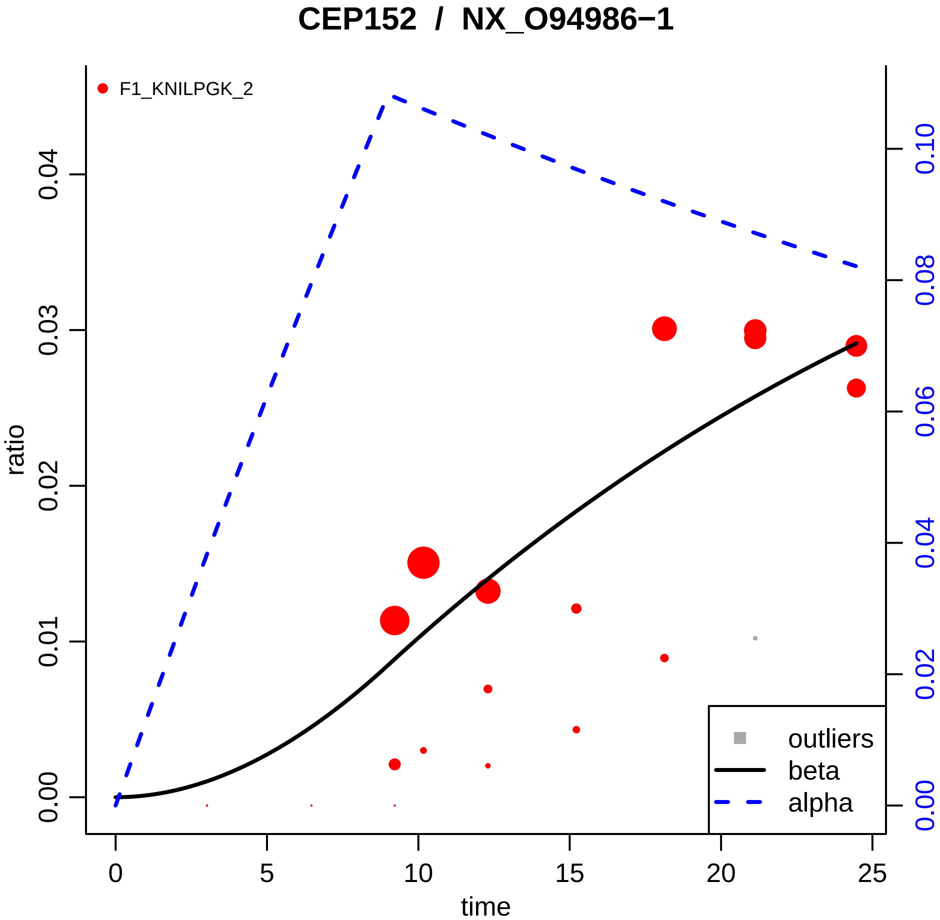

CEP152 / NX_O94986−1
F1_KNILPGK_2
ratio
time
0.00
0.01
0.02
0.03
0.04
0
5
10
15
20
25
0.00
0.02
0.04
0.06
0.08
0.10
outliers
beta
alpha

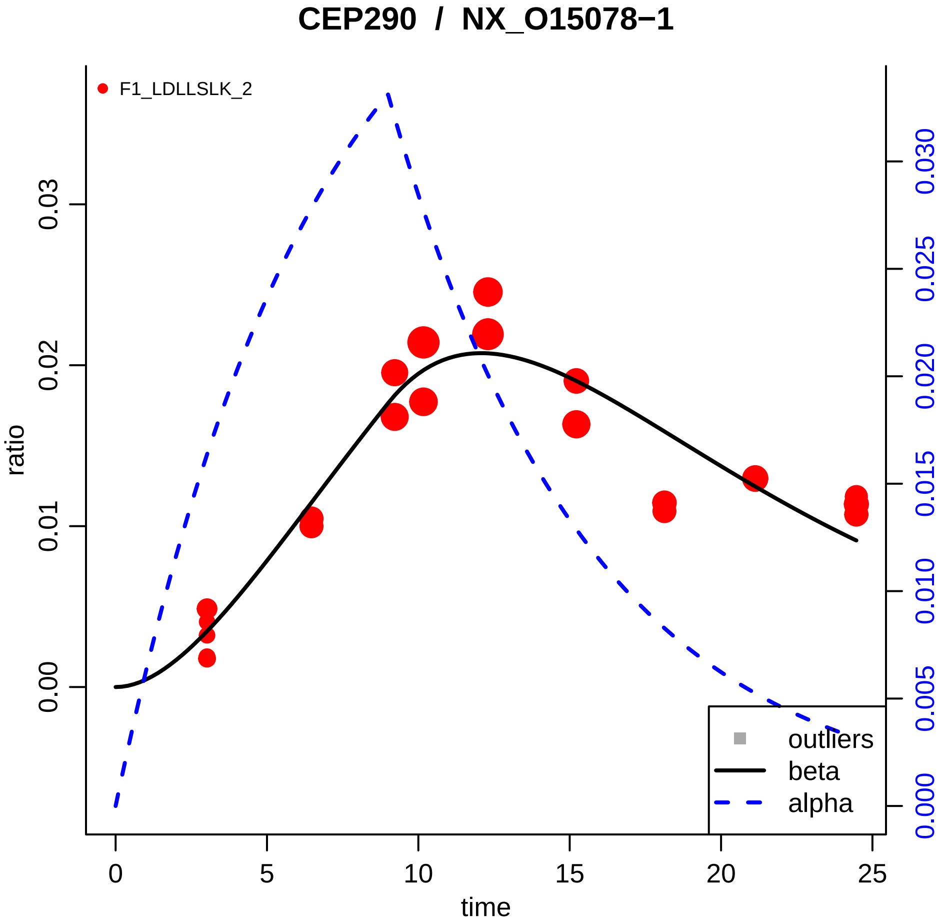
CEP290 / NX_O15078−1
F1_LDLLSLK_2
ratio
time
0.00
0.01
0.02
0.03
0
5
10
15
20
25
0.000
0.005
0.010
0.015
0.020
0.025
0.030
outliers
beta
alpha

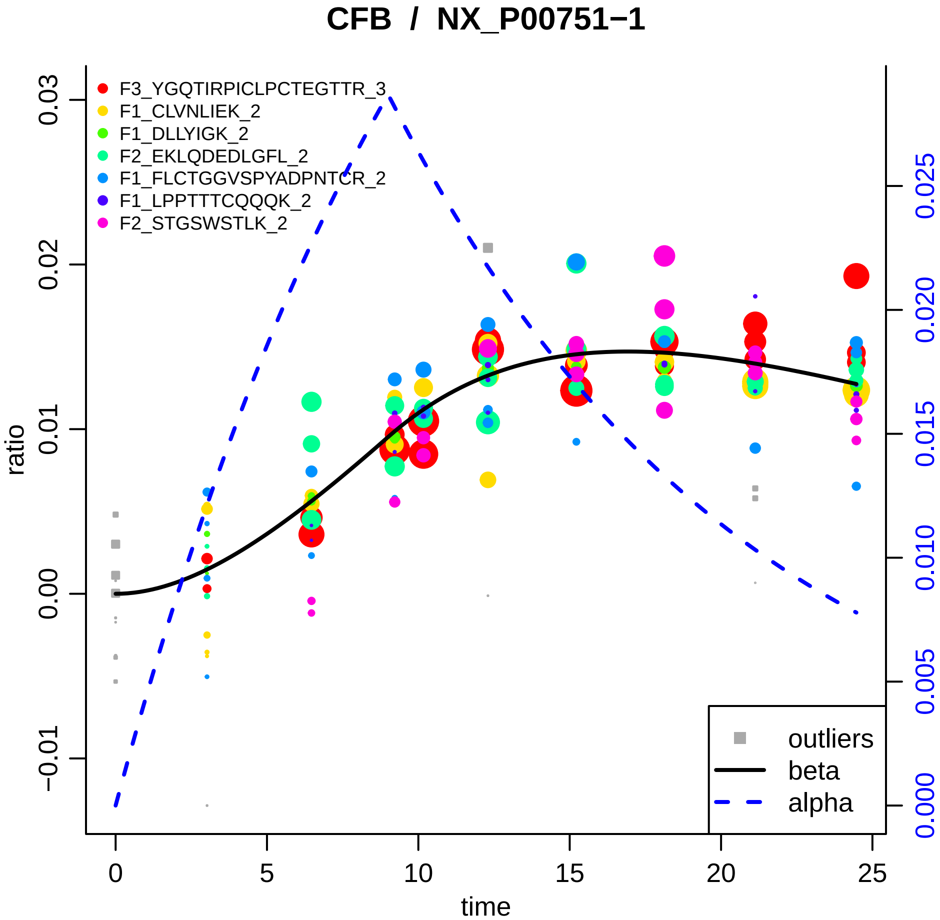
CFB / NX_P00751−1
F3_YGQTIRPICLPCTEGTTR_3
F1_CLVNLIEK_2
F1_DLLYIGK_2
F2_EKLQDEDLGFL_2
F1_FLCTGGVSPYADPNTCR_2
F1_LPPTTTCQQQK_2
F2_STGSWSTLK_2
outliers
beta
alpha
ratio
time
0.03
0.02
0.01
0.00
−0.01
0
5
10
15
20
25
0.025
0.020
0.015
0.010
0.005
0.000

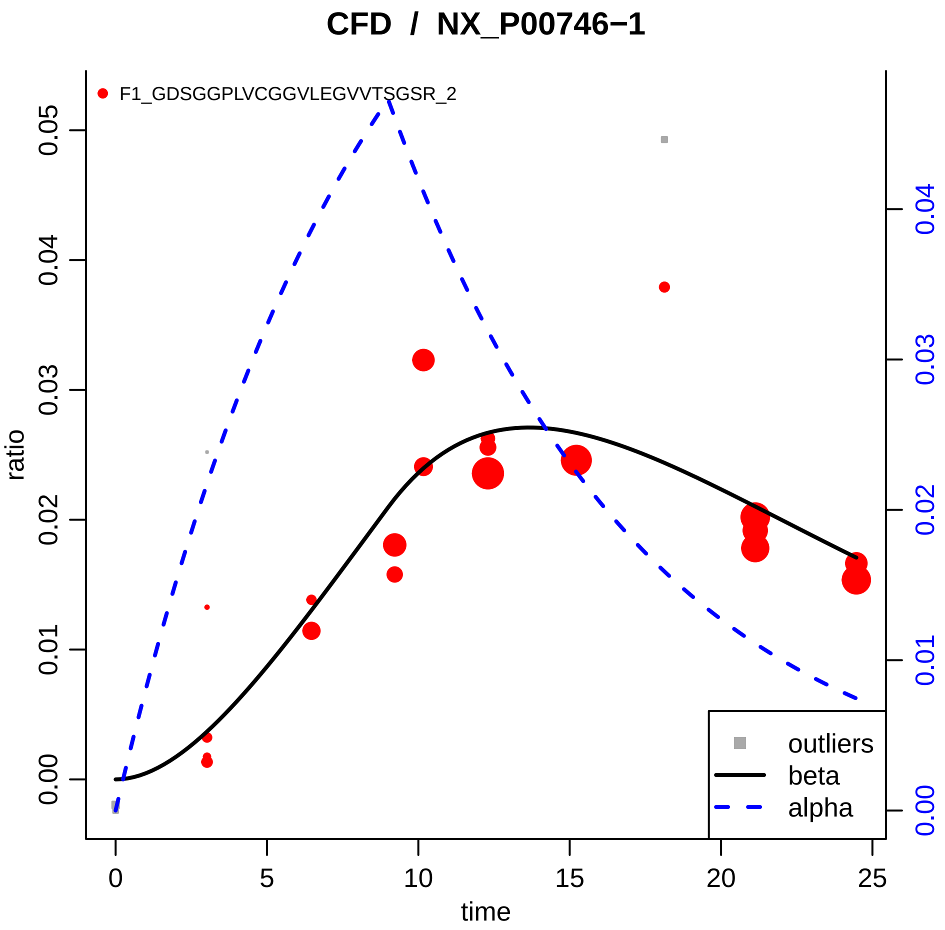

CFD / NX_P00746−1
F1_GDSGGPLVCGGVLEGVVTSGSR_2
ratio
time
0.00
0.01
0.02
0.03
0.04
0.05
0
5
10
15
20
25
0.00
0.01
0.02
0.03
0.04
outliers
beta
alpha

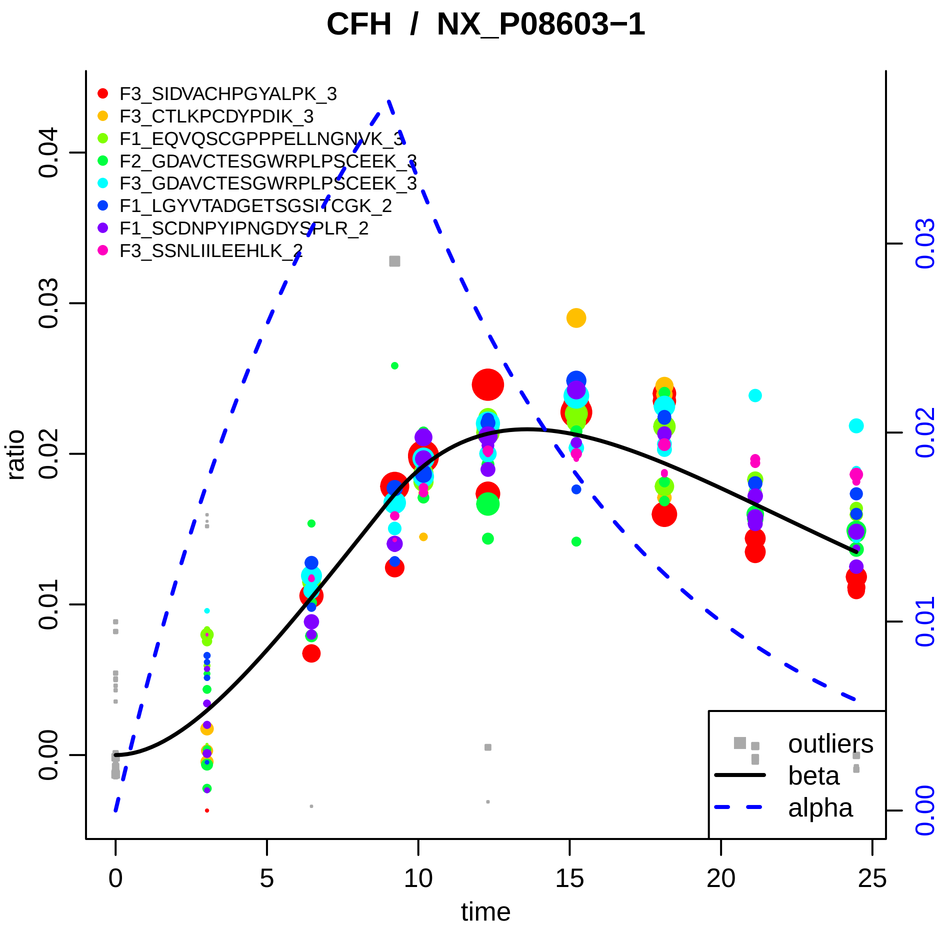

CFH / NX_P08603−1
F3_SIDVACHPGYALPK_3
F3_CTLKPCDYPDIK_3
F1_EQVQSCGPPPELLNGNVK_3
F2_GDAVCTESGWRPLPSCEEK_3
F3_GDAVCTESGWRPLPSCEEK_3
F1_LGYVTADGETSGSITCGK_2
F1_SCDNPYIPNGDYSPLR_2
F3_SSNLIILEEHLK_2
ratio
0.00
0.01
0.02
0.03
0.04
time
0
5
10
15
20
25
outliers
beta
alpha

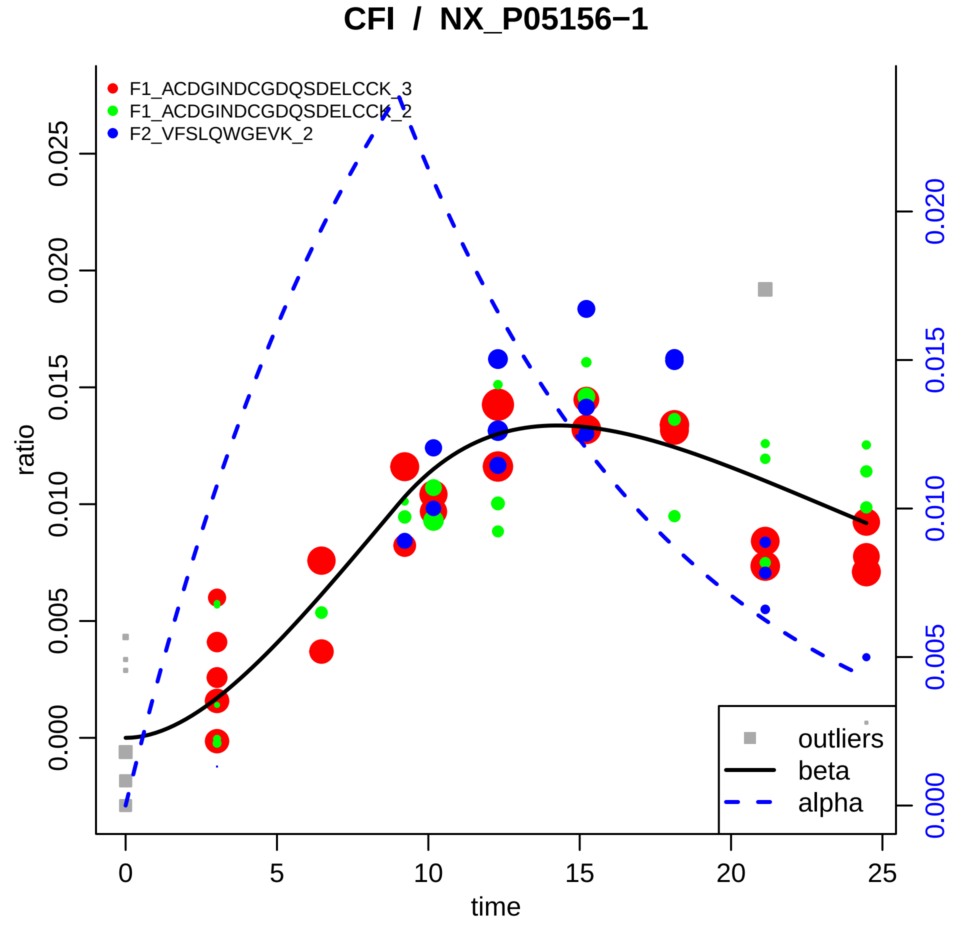

CFI / NX_P05156−1
F1_ACDGINDCGDQSDELCCK_3
F1_ACDGINDCGDQSDELCCK_2
F2_VFSLQWGEVK_2
ratio
0.000
0.005
0.010
0.015
0.020
0.025
time
0
5
10
15
20
25
0.000
0.005
0.010
0.015
0.020
outliers
beta
alpha

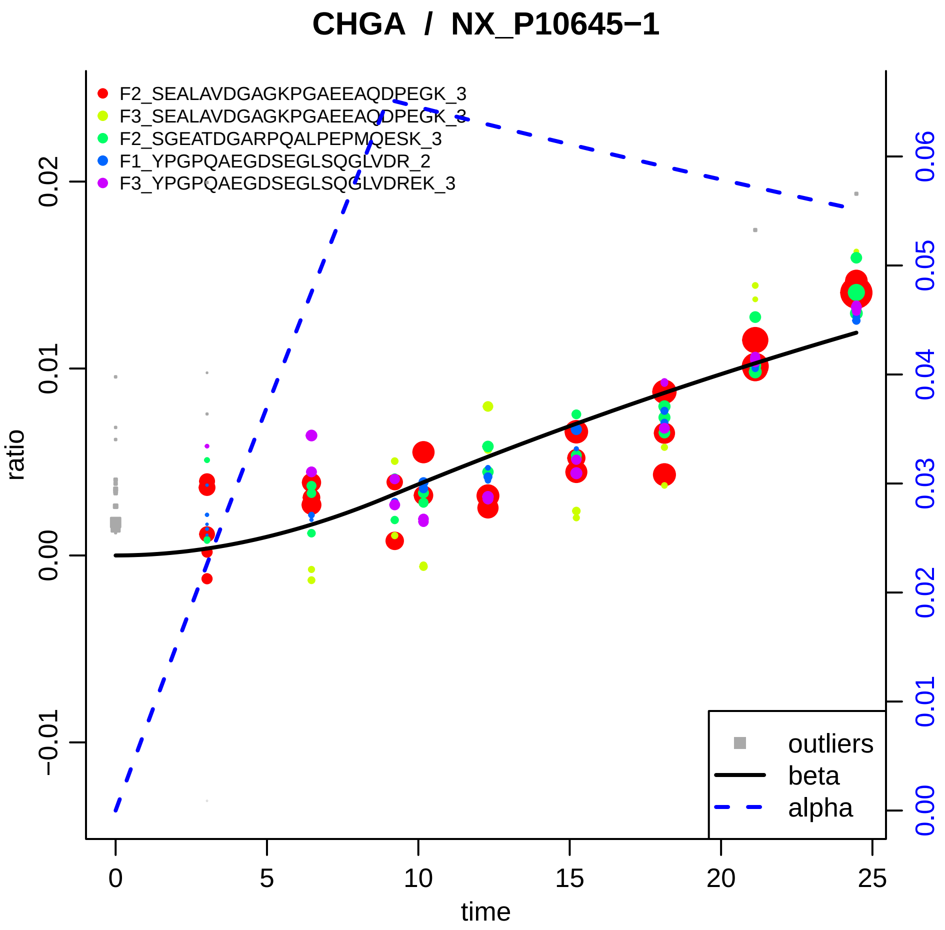

CHGA / NX_P10645−1
F2_SEALAVDGAGKPGAEEAQDPEGK_3
F3_SEALAVDGAGKPGAEEAQDPEGK_3
F2_SGEATDGARPQALPEPMQESK_3
F1_YPGPQAEGDSEGLSQGLVDR_2
F3_YPGPQAEGDSEGLSQGLVDREK_3
ratio
time
outliers
beta
alpha
−0.01
0.00
0.01
0.02
0
5
10
15
20
25
0.00
0.01
0.02
0.03
0.04
0.05
0.06

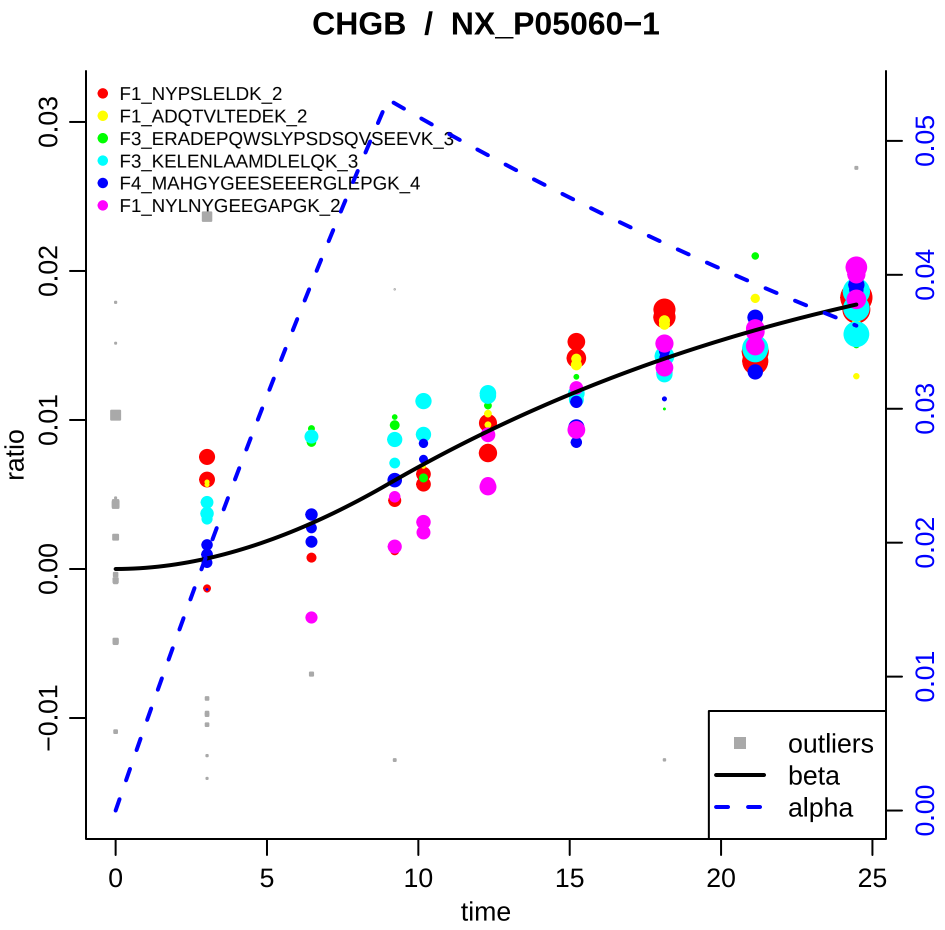

CHGB / NX_P05060−1
F1_NYPSLELDK_2
F1_ADQTVLTEDEK_2
F3_ERADEPQWSLYPSDSQVSEEVK_3
F3_KELENLAAMDLELQK_3
F4_MAHGYGEESEEERGLEPGK_4
F1_NYLNYGEEGAPGK_2
outliers
beta
alpha
ratio
time
0
5
10
15
20
25
−0.01
0.00
0.01
0.02
0.03
0.00
0.01
0.02
0.03
0.04
0.05

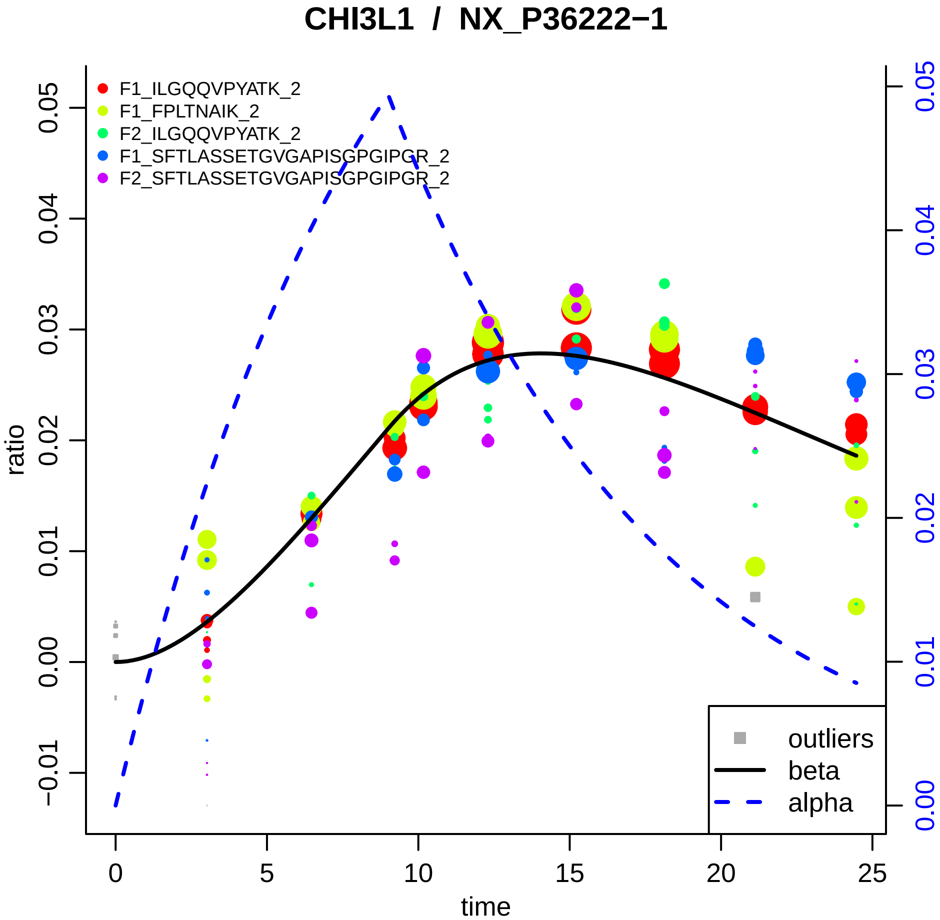

CHI3L1 / NX_P36222−1
F1_ILGQQVPYATK_2
F1_FPLTNAIK_2
F2_ILGQQVPYATK_2
F1_SFTLASSETGVGAPISGPGIPGR_2
F2_SFTLASSETGVGAPISGPGIPGR_2
outliers
beta
alpha
ratio
time
0.05
0.04
0.03
0.02
0.01
0.00
−0.01
0
5
10
15
20
25

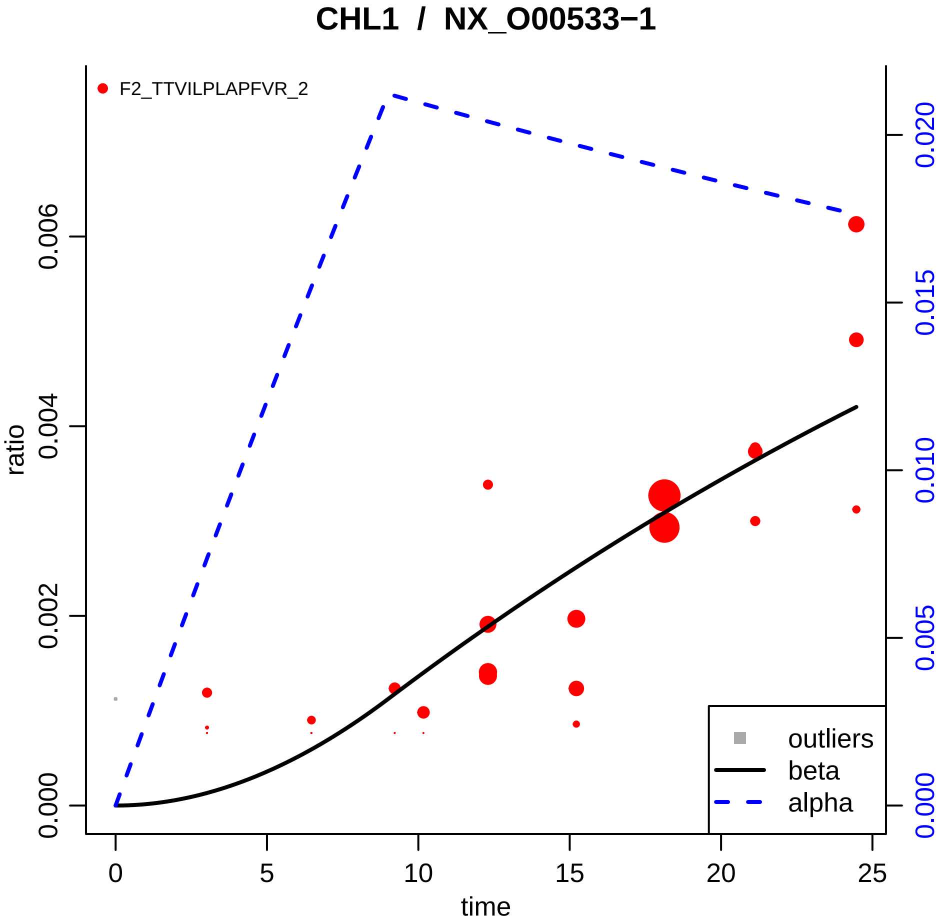

CHL1 / NX_O00533−1
F2_TTVILPLAPFVR_2
ratio
time
0.000
0.002
0.004
0.006
0
5
10
15
20
25
0.000
0.005
0.010
0.015
0.020
outliers
beta
alpha

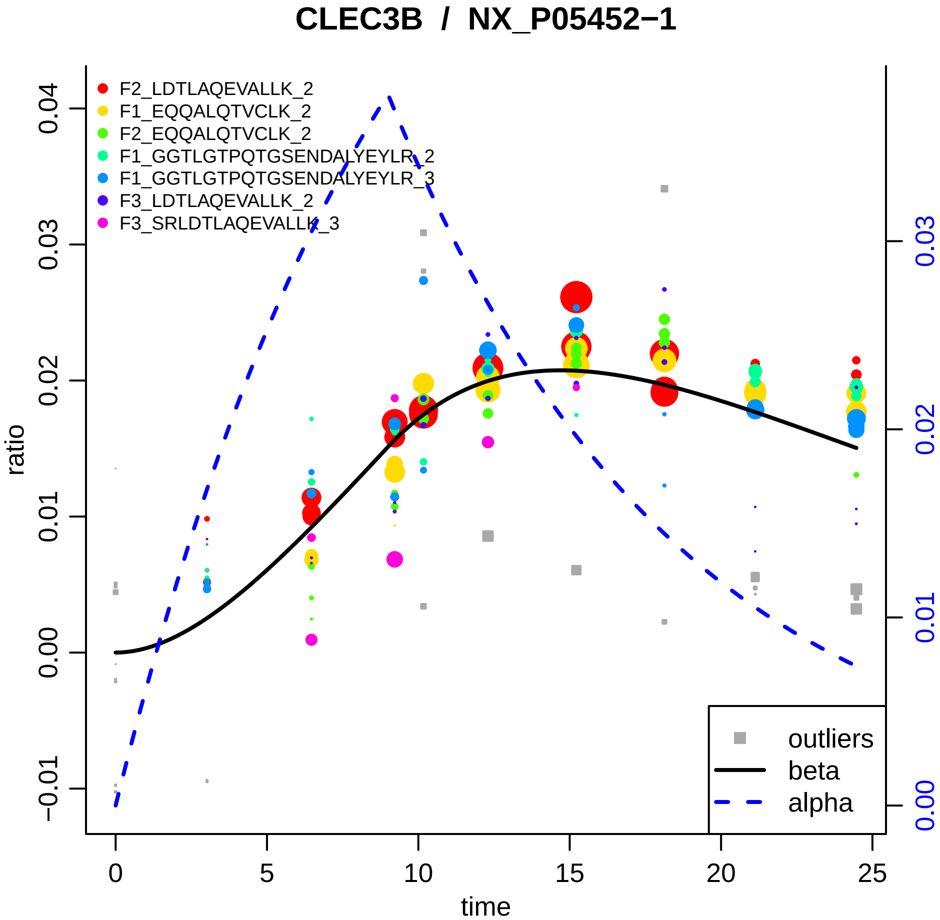

CLEC3B / NX_P05452−1
F2_LDTLAQEVALLK_2
F1_EQQALQTVCLK_2
F2_EQQALQTVCLK_2
F1_GGTLGTPQTGSENDALYEYLR_2
F1_GGTLGTPQTGSENDALYEYLR_3
F3_LDTLAQEVALLK_2
F3_SRLDTLAQEVALLK_3
ratio
time
−0.01
0.00
0.01
0.02
0.03
0.04
0
5
10
15
20
25
0.00
0.01
0.02
0.03
outliers
beta
alpha

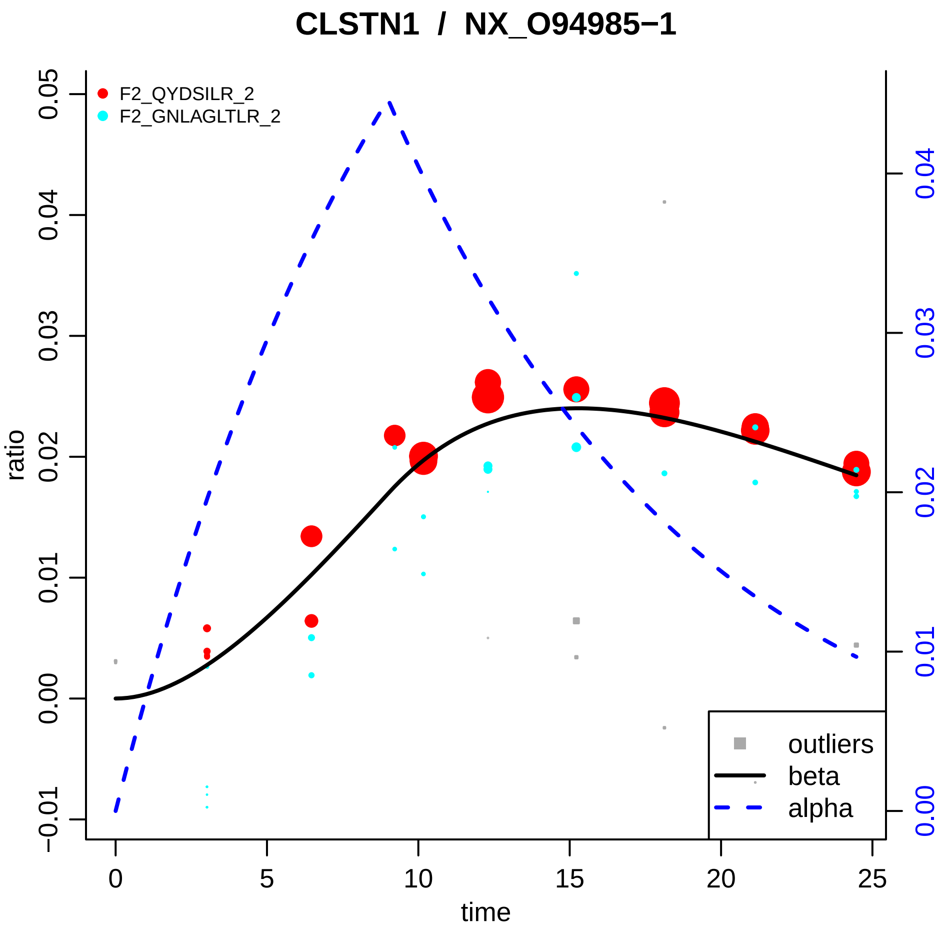

CLSTN1 / NX_O94985−1
F2_QYDSILR_2
F2_GNLAGLTLR_2
ratio
time
outliers
beta
alpha
0.05
0.04
0.03
0.02
0.01
0.00
−0.01
0
5
10
15
20
25
0.04
0.03
0.02
0.01
0.00

### CLU / NX_P10909−1

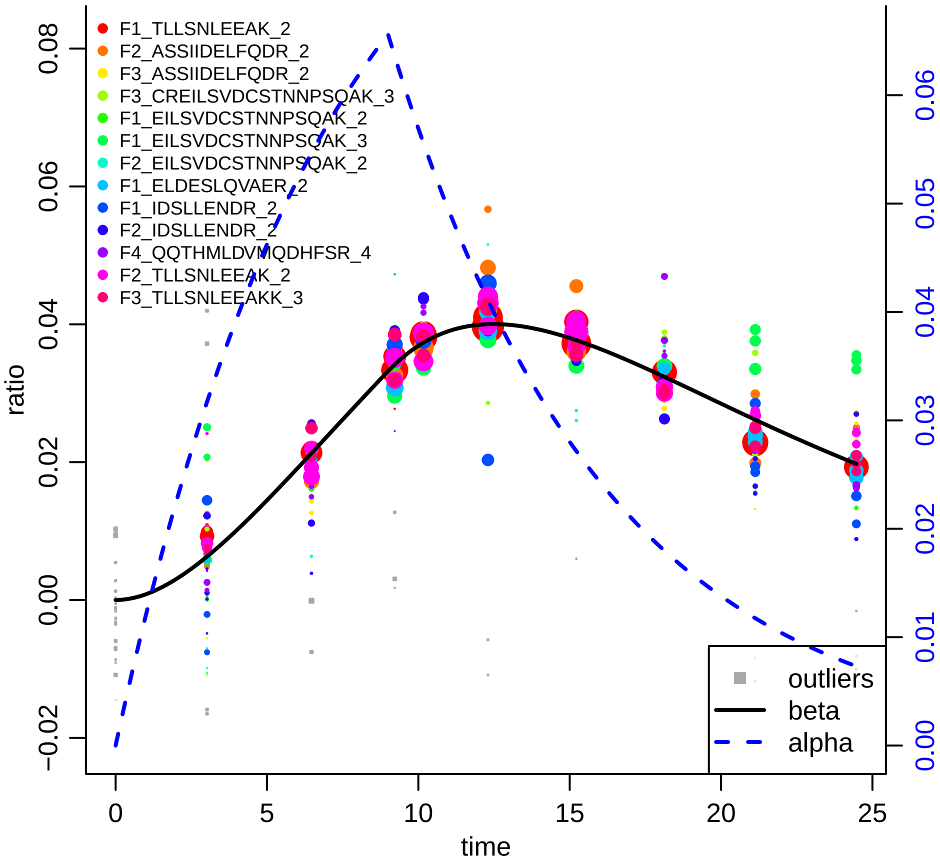

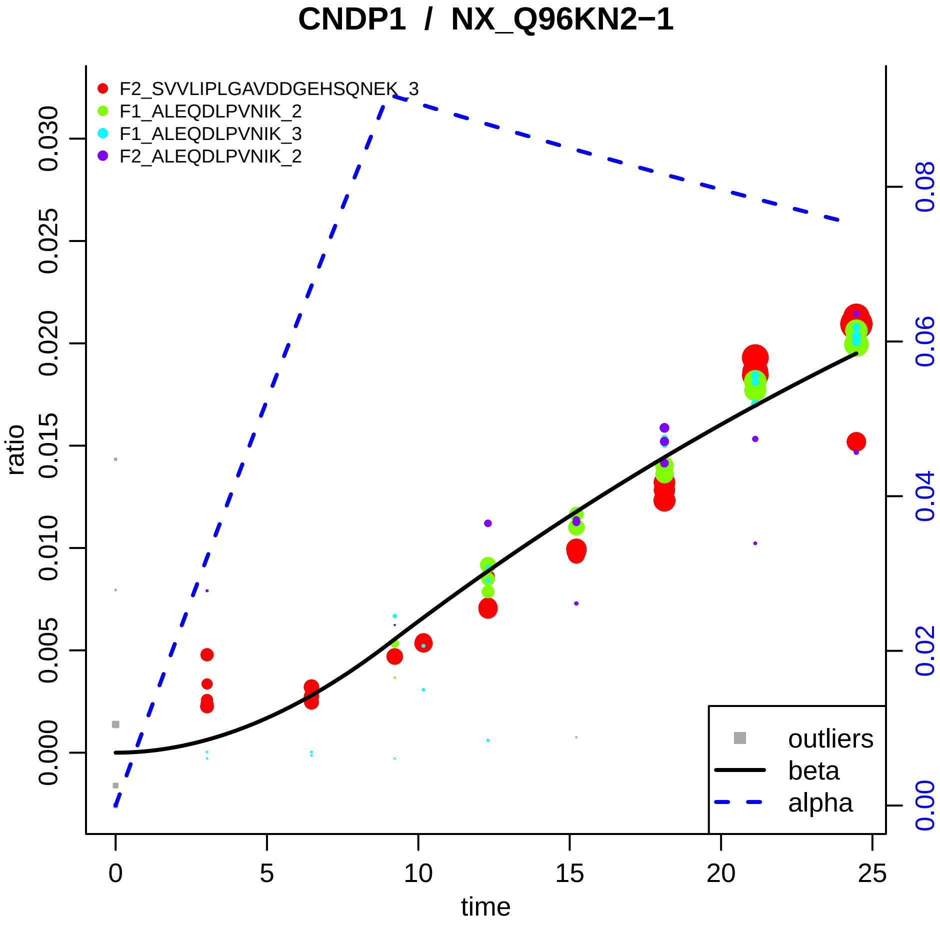
CNDP1 / NX_Q96KN2−1
F2_SVVLIPLGAVDDGEHSQNEK_3
F1_ALEQDLPVNIK_2
F1_ALEQDLPVNIK_3
F2_ALEQDLPVNIK_2
ratio
0.000
0.005
0.010
0.015
0.020
0.025
0.030
0.00
0.02
0.04
0.06
0.08
0
5
10
15
20
25
time
outliers
beta
alpha

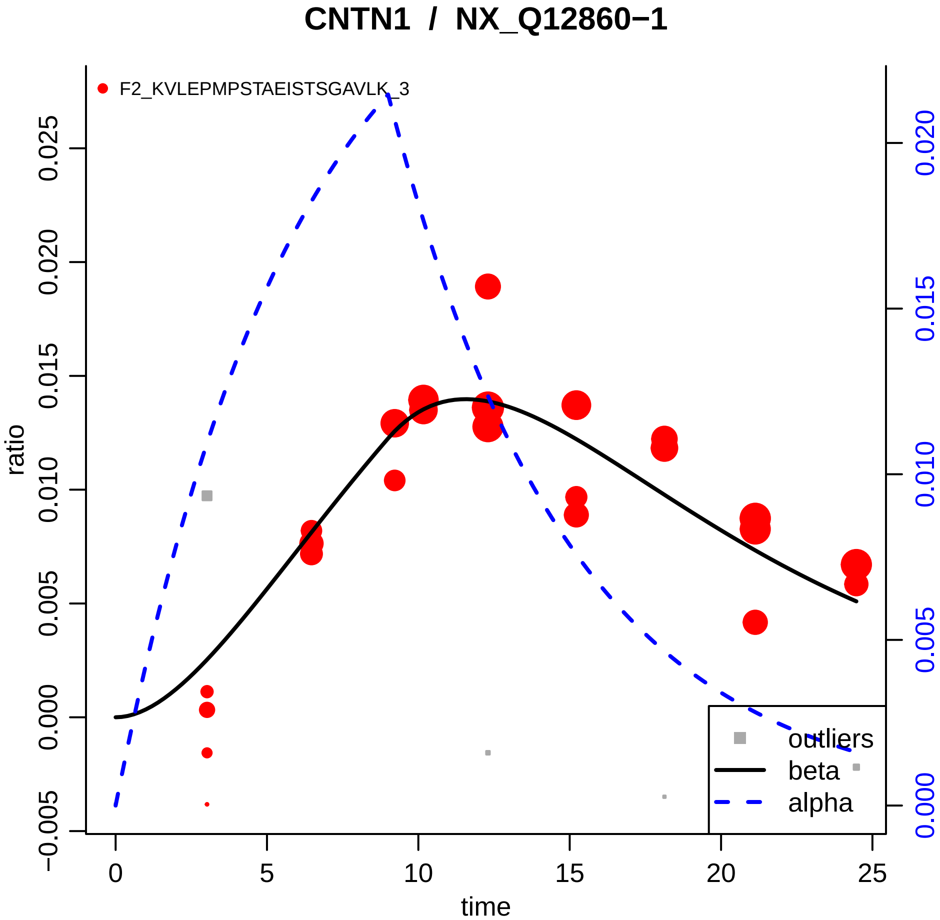

CNTN1 / NX_Q12860−1
F2_KVLEPMPSTAEISTSGAVLK_3
ratio
time
0
5
10
15
20
25
−0.005
0.000
0.005
0.010
0.015
0.020
0.025
0.000
0.005
0.010
0.015
0.020
outliers
beta
alpha

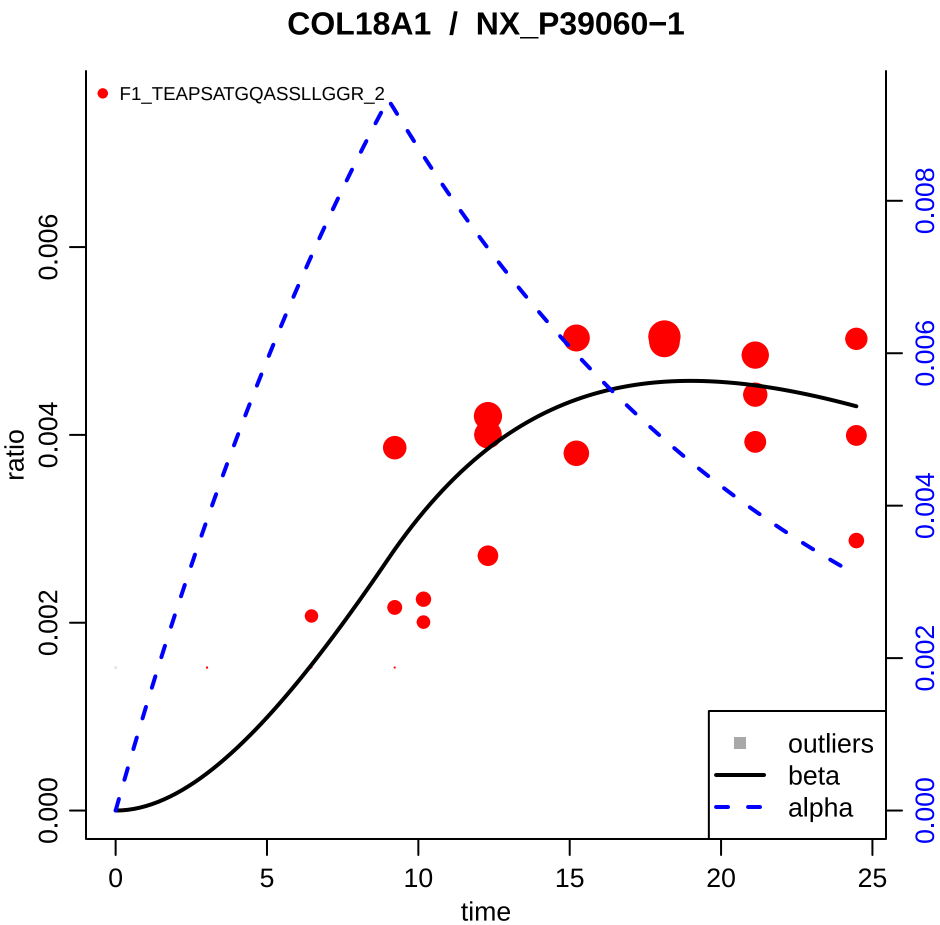
COL18A1 / NX_P39060−1
F1_TEAPSATGQASSLLGGR_2
ratio
0.000
0.002
0.004
0.006
0.000
0.002
0.004
0.006
0.008
0
5
10
15
20
25
time
outliers
beta
alpha

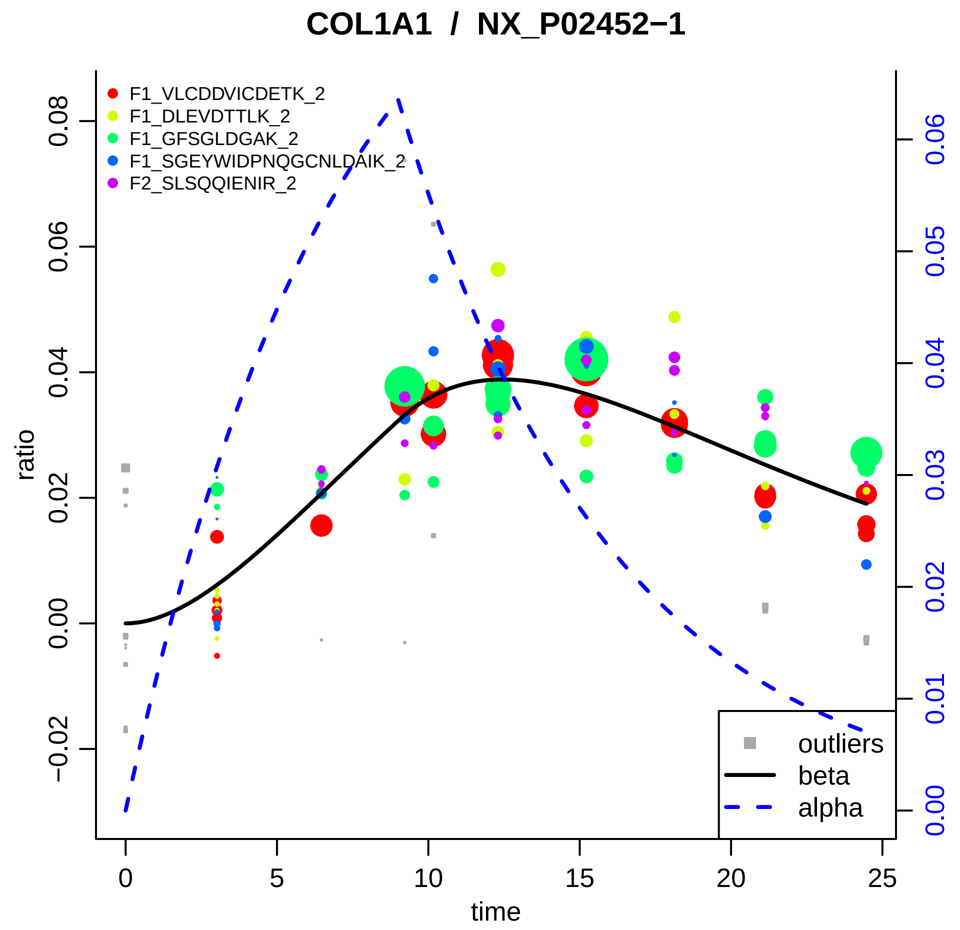

COLO1A1 / NX_P02452−1
F1_VLCDDVICDETK_2
F1_DLEVDTTLK_2
F1_GFSGLDGAK_2
F1_SGEYWIDPNQGCNLDAIK_2
F2_SLSQQIENIR_2
outliers
beta
alpha
ratio
time
−0.02
0.00
0.02
0.04
0.06
0.08
0
5
10
15
20
25
0.00
0.01
0.02
0.03
0.04
0.05
0.06

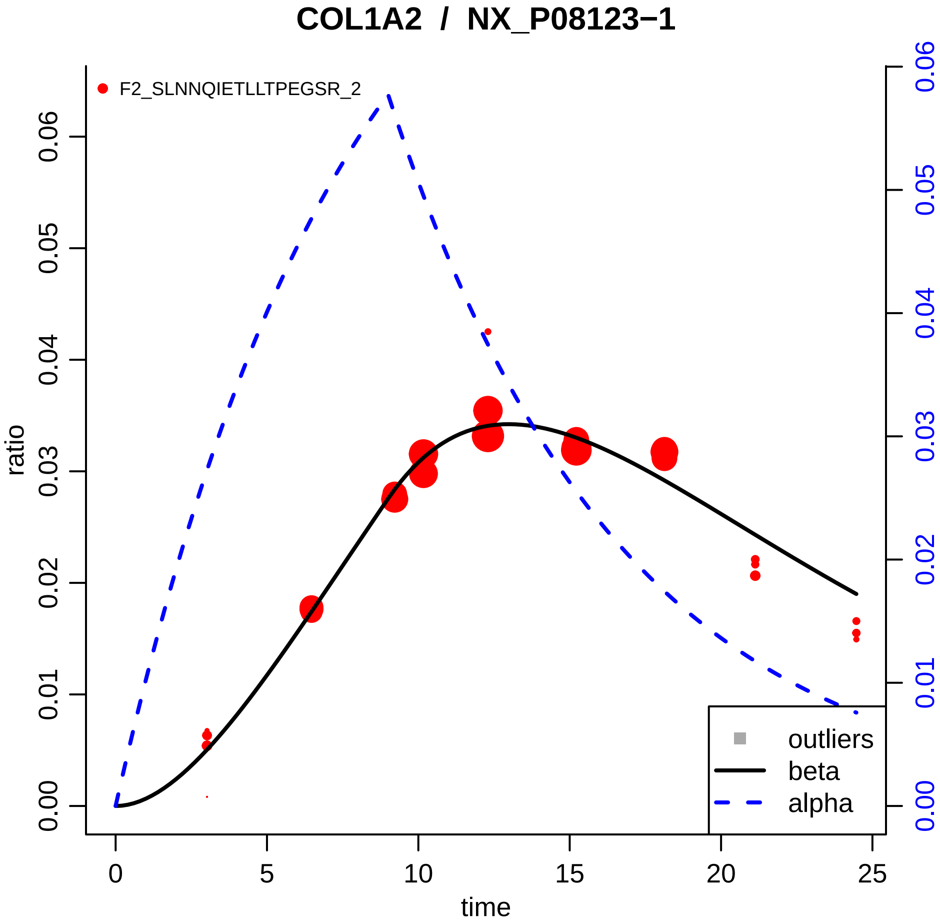

COL1A2 / NX_P08123−1
F2_SLNNQIETLLTPEGSR_2
ratio
time
0.00
0.01
0.02
0.03
0.04
0.05
0.06
0
5
10
15
20
25
outliers
beta
alpha

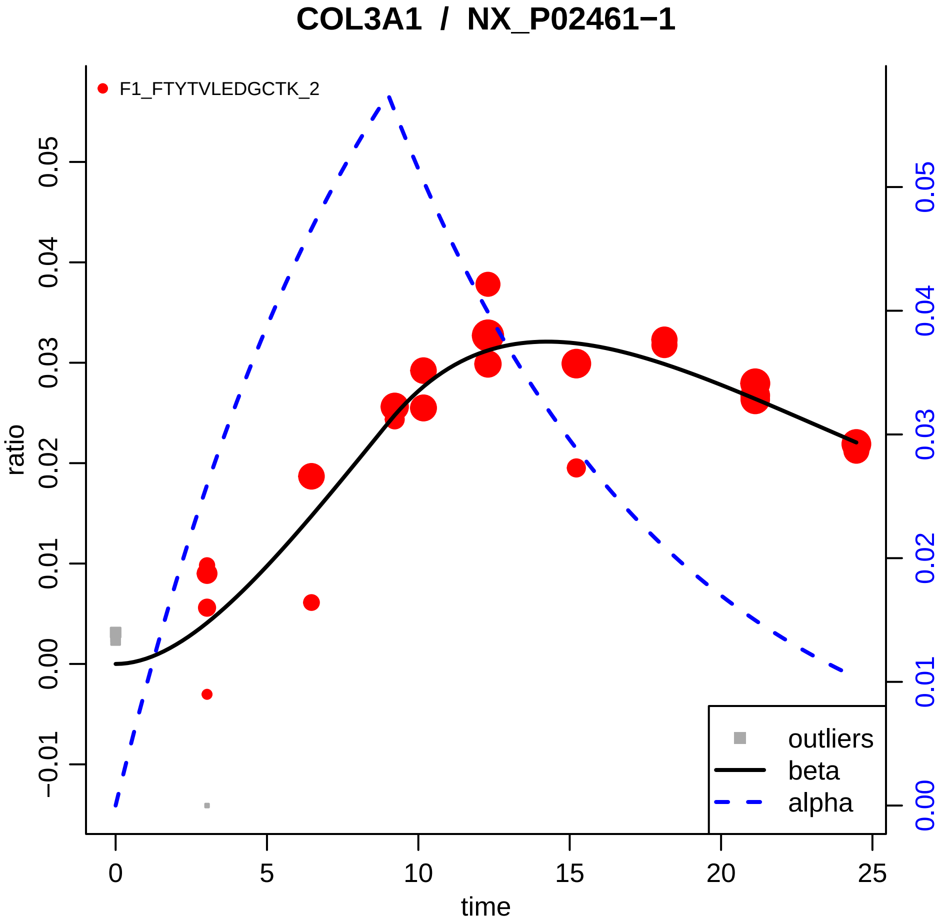

COL3A1 / NX_P02461−1
F1_FTYTVLEDGCTK_2
ratio
time
0
5
10
15
20
25
−0.01
0.00
0.01
0.02
0.03
0.04
0.05
outliers
beta
alpha

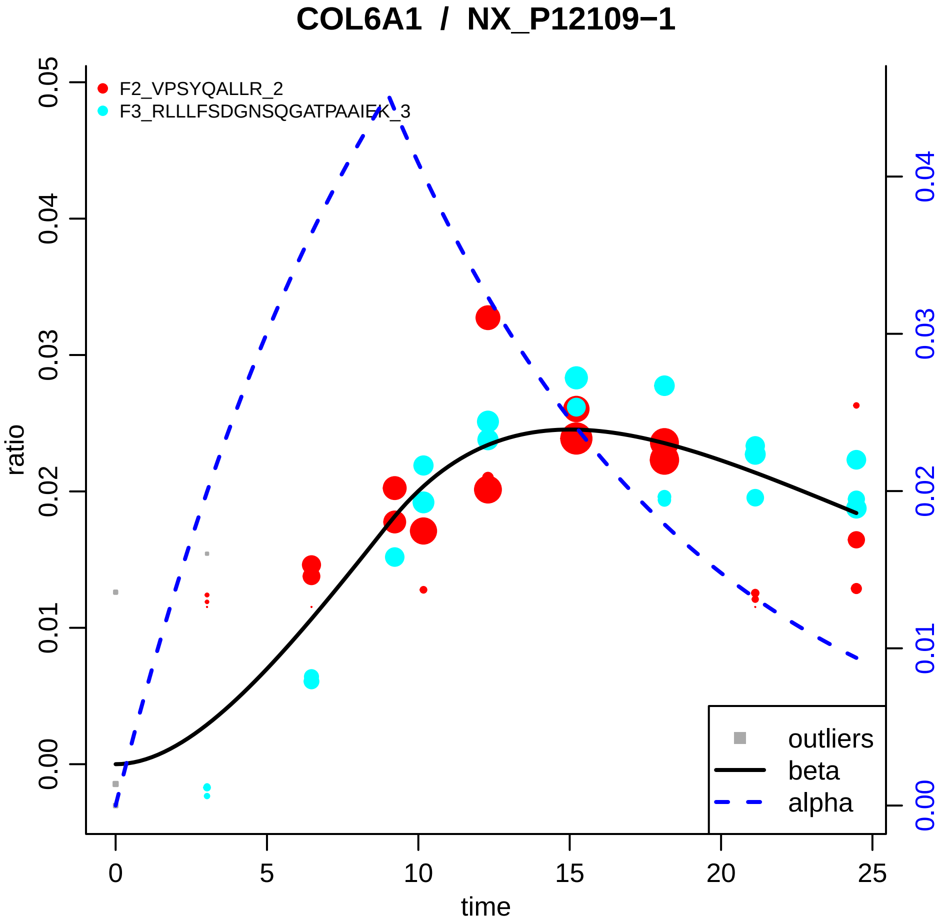

COL6A1 / NX_P12109−1
F2_VPSYQALLR_2
F3_RLLLFSDGNSQGATPAAIEK_3
ratio
time
0.00
0.01
0.02
0.03
0.04
0.05
0
5
10
15
20
25
outliers
beta
alpha

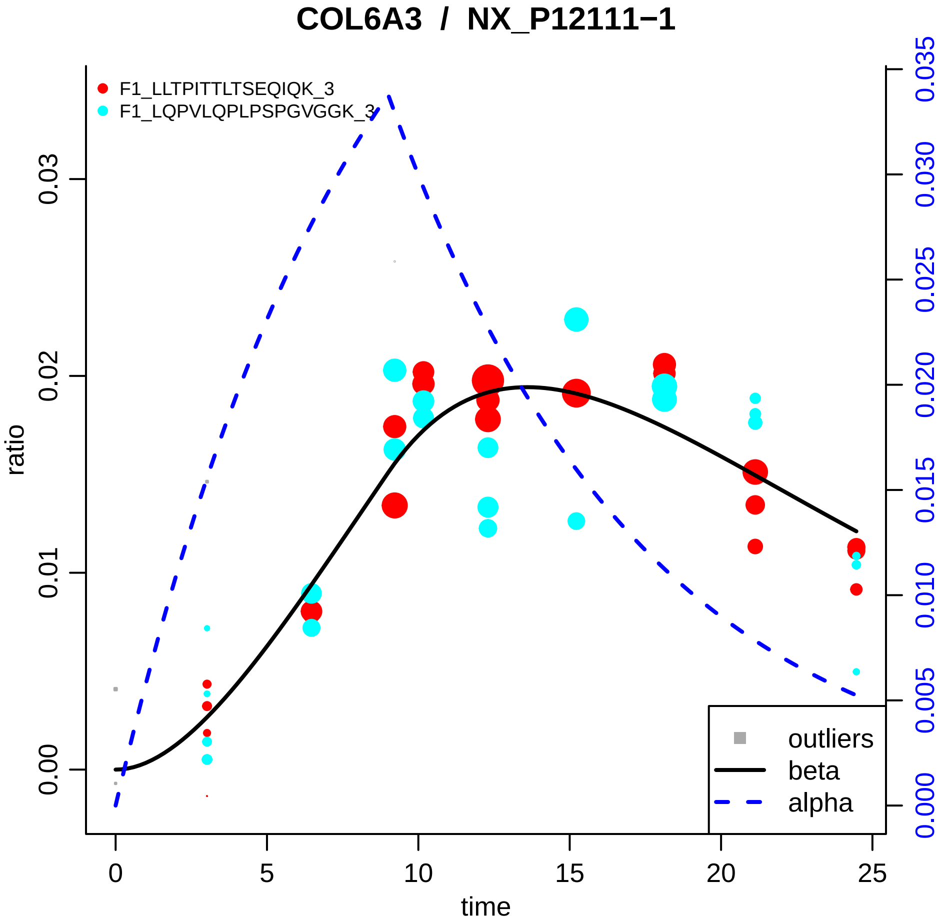

COL6A3 / NX_P12111−1
F1_LLTPITTLTSEQIQK_3
F1_LQPVLQPLPSPGVGGK_3
ratio
time
0.00
0.01
0.02
0.03
0
5
10
15
20
25
0.000
0.005
0.010
0.015
0.020
0.025
0.030
0.035
outliers
beta
alpha

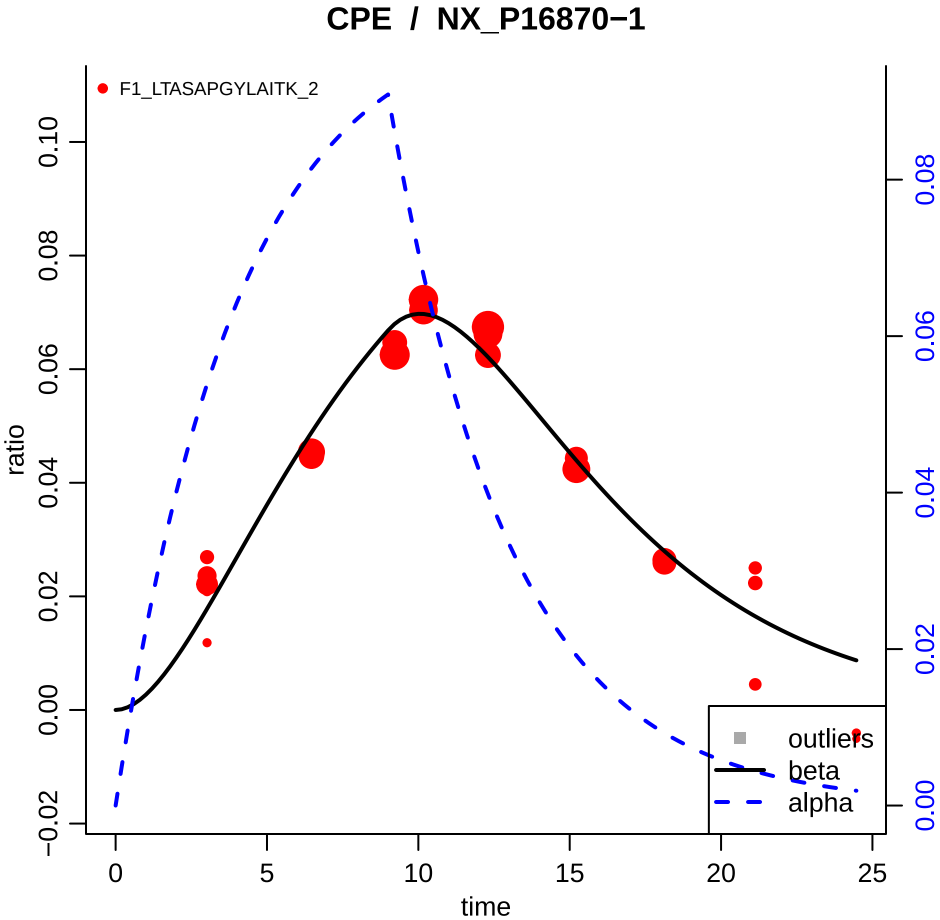

CPE / NX_P16870−1
F1_LTASAPGYLAITK_2
ratio
time
0.10
0.08
0.06
0.04
0.02
0.00
−0.02
0
5
10
15
20
25
0.08
0.06
0.04
0.02
0.00
outliers
beta
alpha

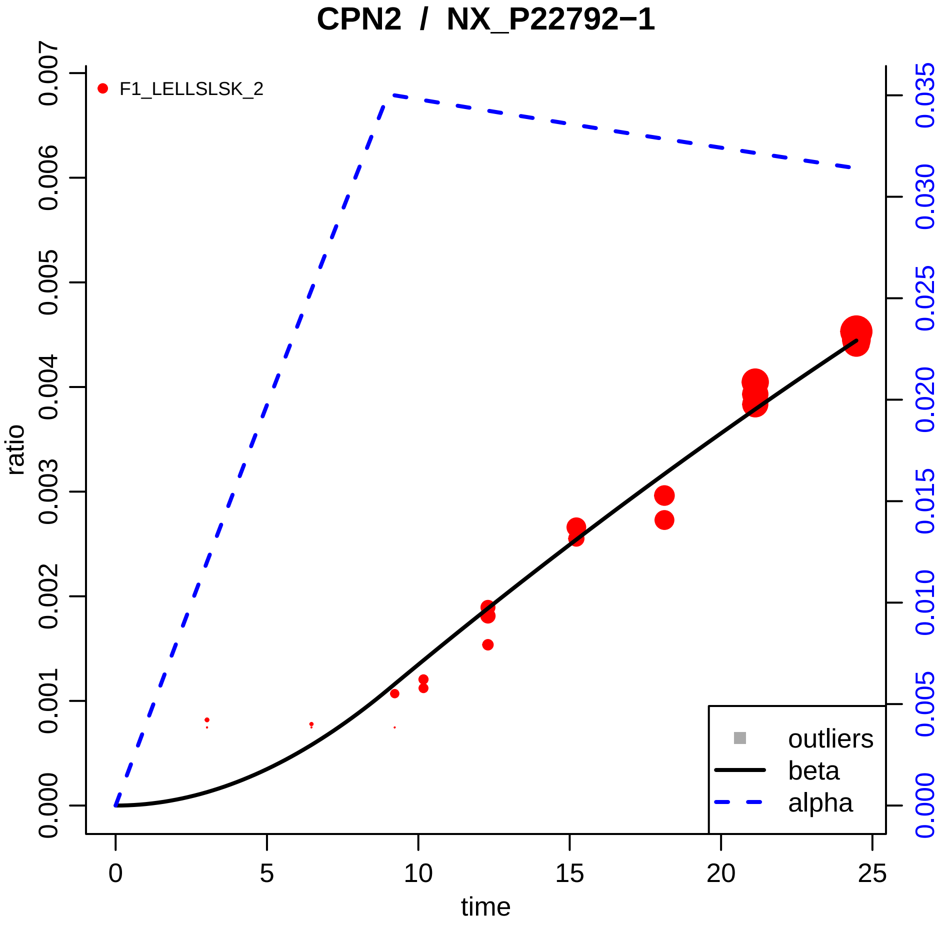

CPN2 / NX_P22792−1
F1_LELLSLSK_2
ratio
time
0.000
0.001
0.002
0.003
0.004
0.005
0.006
0.007
0
5
10
15
20
25
0.000
0.005
0.010
0.015
0.020
0.025
0.030
0.035
outliers
beta
alpha

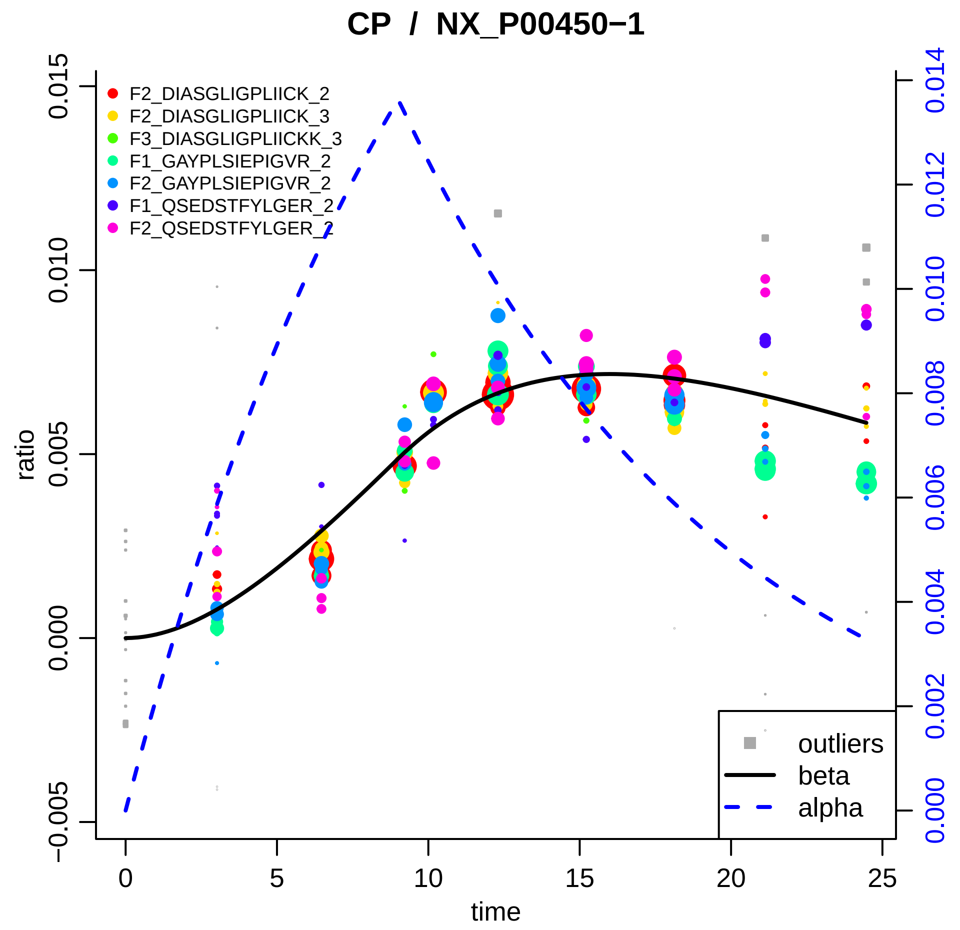

CP / NX_P00450−1
F2_DIASGLIGPLIICK_2
F2_DIASGLIGPLIICK_3
F3_DIASGLIGPLIICKK_3
F1_GAYPLSIEPIGVR_2
F2_GAYPLSIEPIGVR_2
F1_QSEDSTFYLGER_2
F2_QSEDSTFYLGER_2
outliers
beta
alpha
ratio
time
0.015
0.010
0.005
0.000
−0.005
0
5
10
15
20
25
0.014
0.012
0.010
0.008
0.006
0.004
0.002
0.000

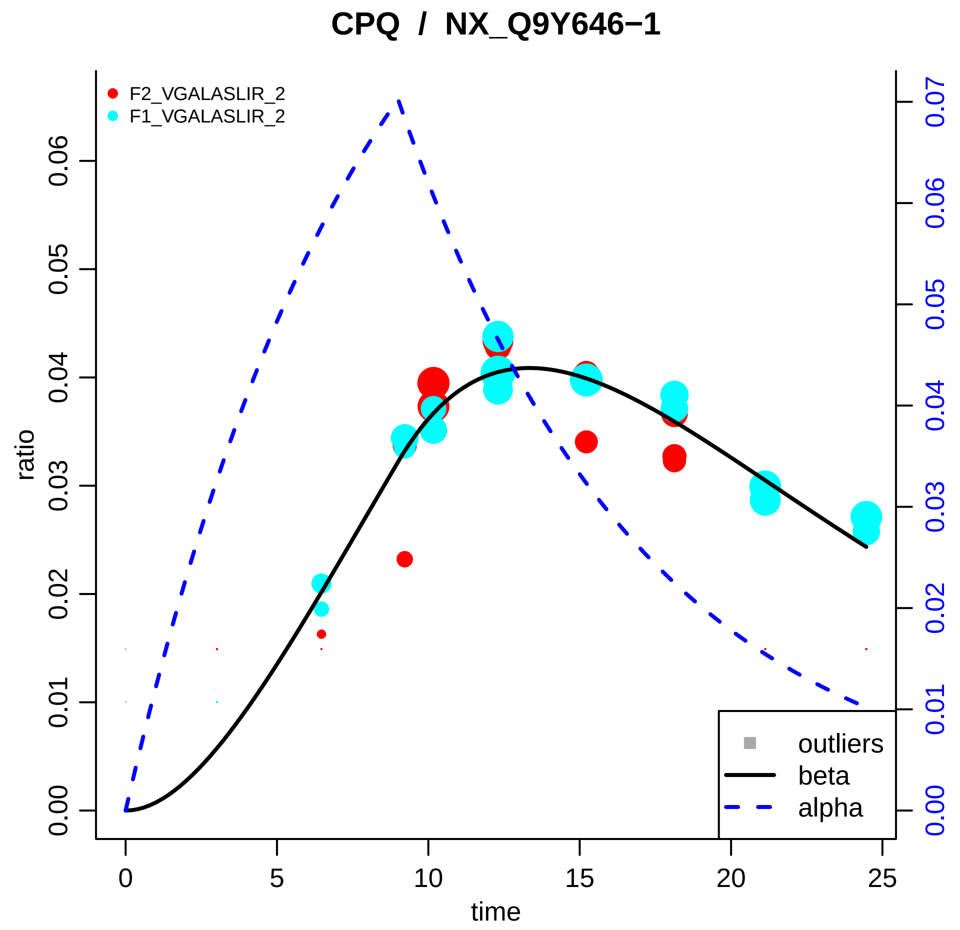
CPQ / NX_Q9Y646−1
F2_VGALASLIR_2
F1_VGALASLIR_2
outliers
beta
alpha
ratio
time
0.00
0.01
0.02
0.03
0.04
0.05
0.06
0.07
0
5
10
15
20
25

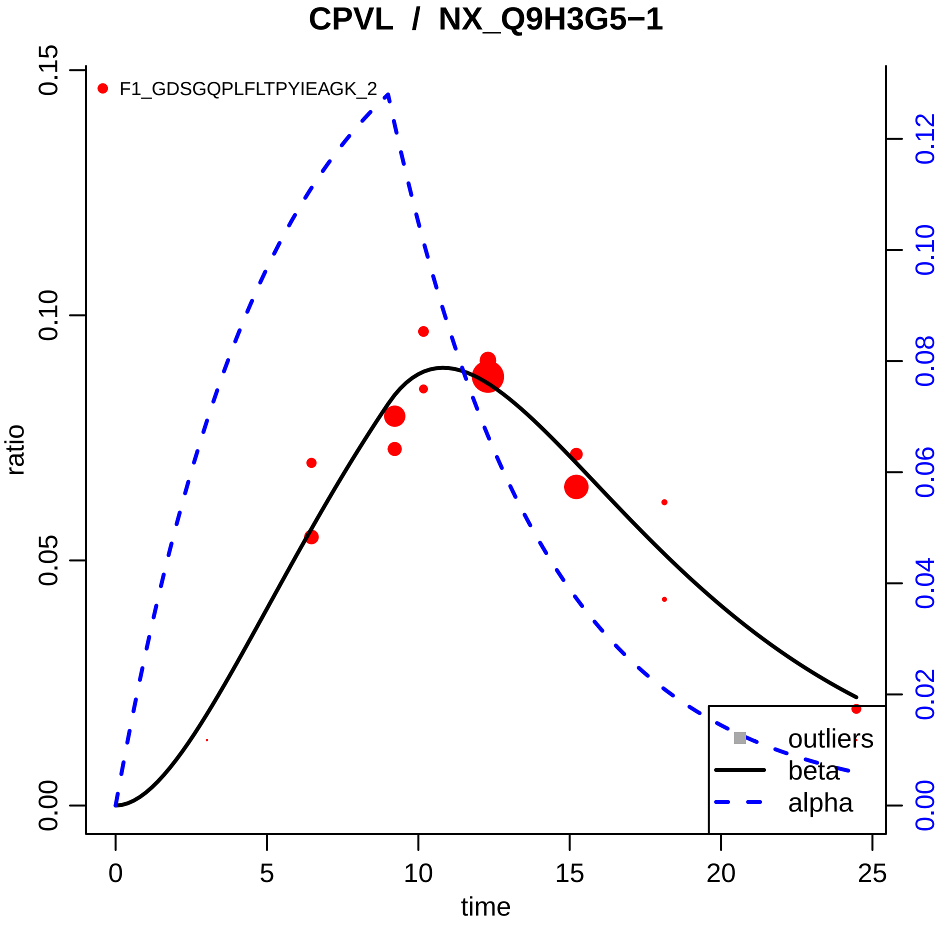

CPVL / NX_Q9H3G5−1
F1_GDSGQPLFLTPYIEAGK_2
ratio
time
0.00
0.05
0.10
0.15
0
5
10
15
20
25
0.00
0.02
0.04
0.06
0.08
0.10
0.12
outliers
beta
alpha

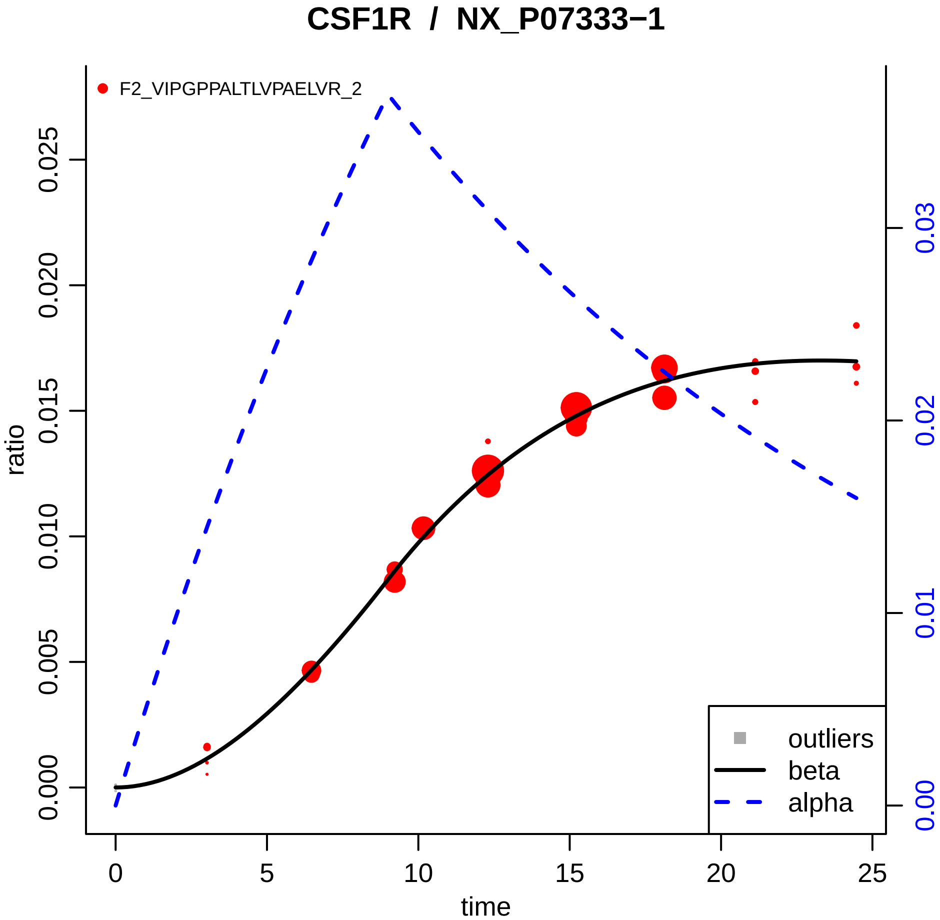

CSF1R / NX_P07333−1
F2_VIPGPPALTLVPAELVR_2
ratio
0.000
0.005
0.010
0.015
0.020
0.025
0.00
0.01
0.02
0.03
0
5
10
15
20
25
time
outliers
beta
alpha

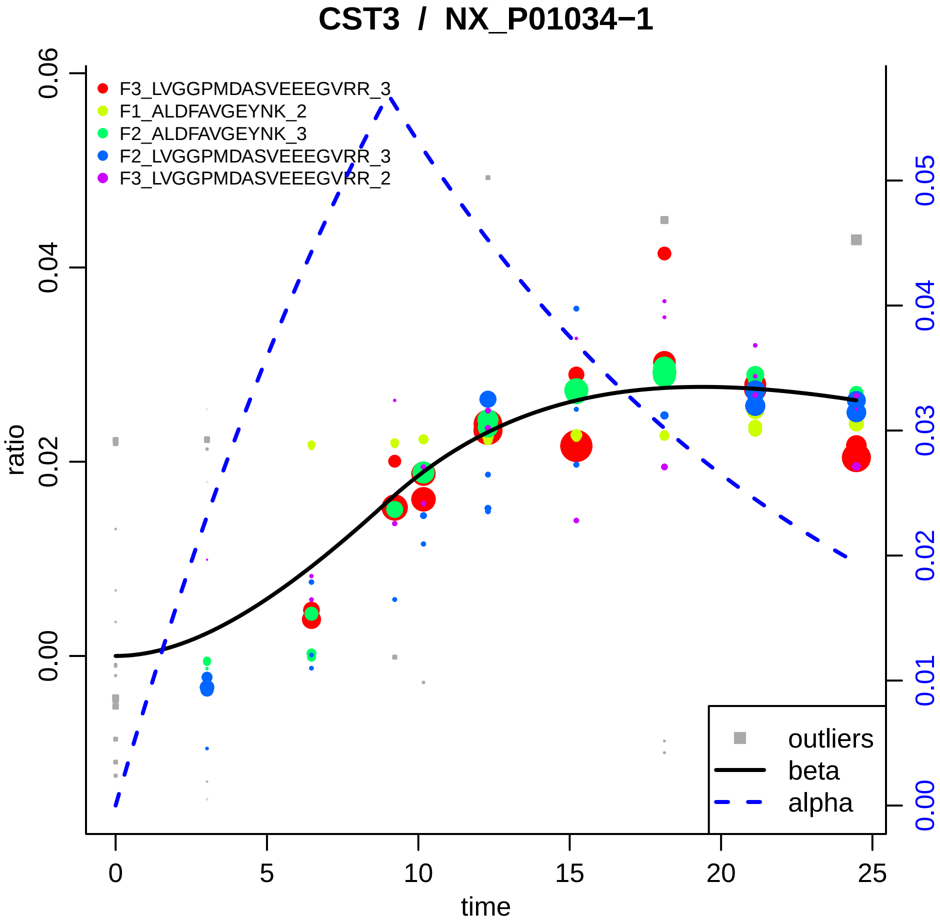

CST3 / NX_P01034−1
F3_LVGGPMDASVEEEGVRR_3
F1_ALDFAVGEYNK_2
F2_ALDFAVGEYNK_3
F2_LVGGPMDASVEEEGVRR_3
F3_LVGGPMDASVEEEGVRR_2
outliers
beta
alpha
ratio
time
0.00
0.02
0.04
0.06
0.00
0.01
0.02
0.03
0.04
0.05
0
5
10
15
20
25

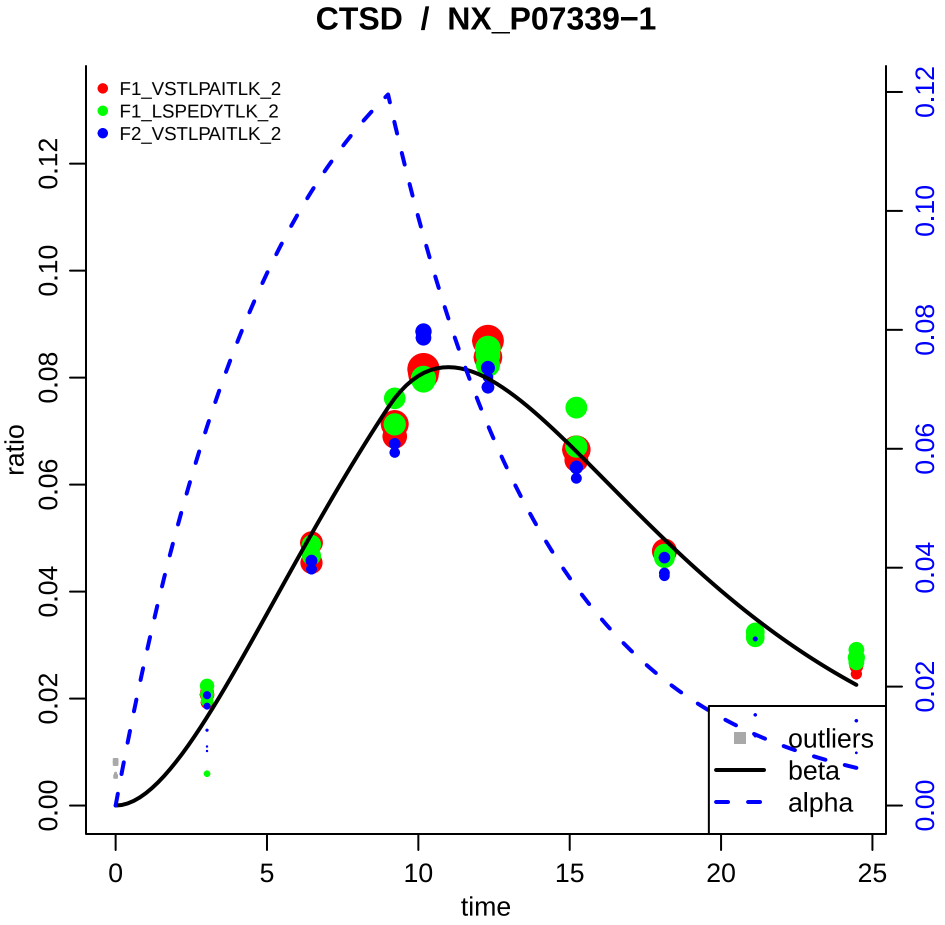
CTSD / NX_P07339−1
F1_VSTLPAITLK_2
F1_LSPEDYTLK_2
F2_VSTLPAITLK_2
ratio
time
0.00
0.02
0.04
0.06
0.08
0.10
0.12
0
5
10
15
20
25
outliers
beta
alpha

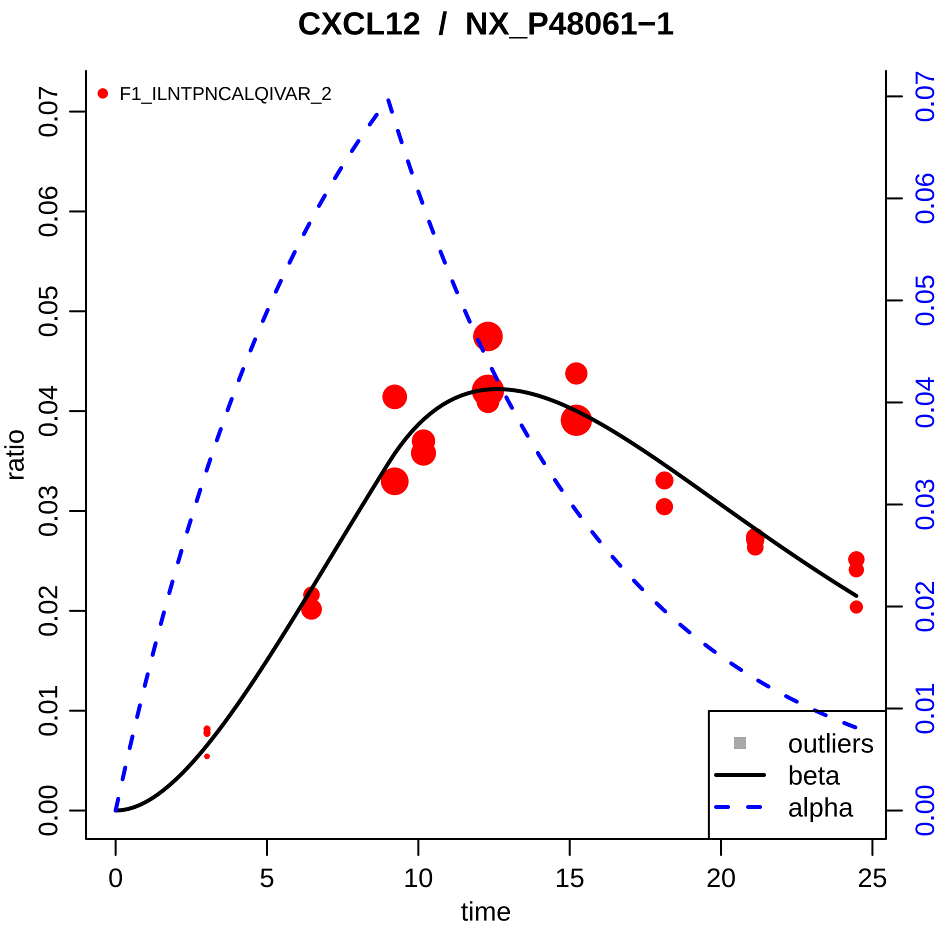
CXCL12 / NX_P48061−1
F1_ILNTPNCALQIVAR_2
ratio
time
0.00
0.01
0.02
0.03
0.04
0.05
0.06
0.07
0
5
10
15
20
25
outliers
beta
alpha

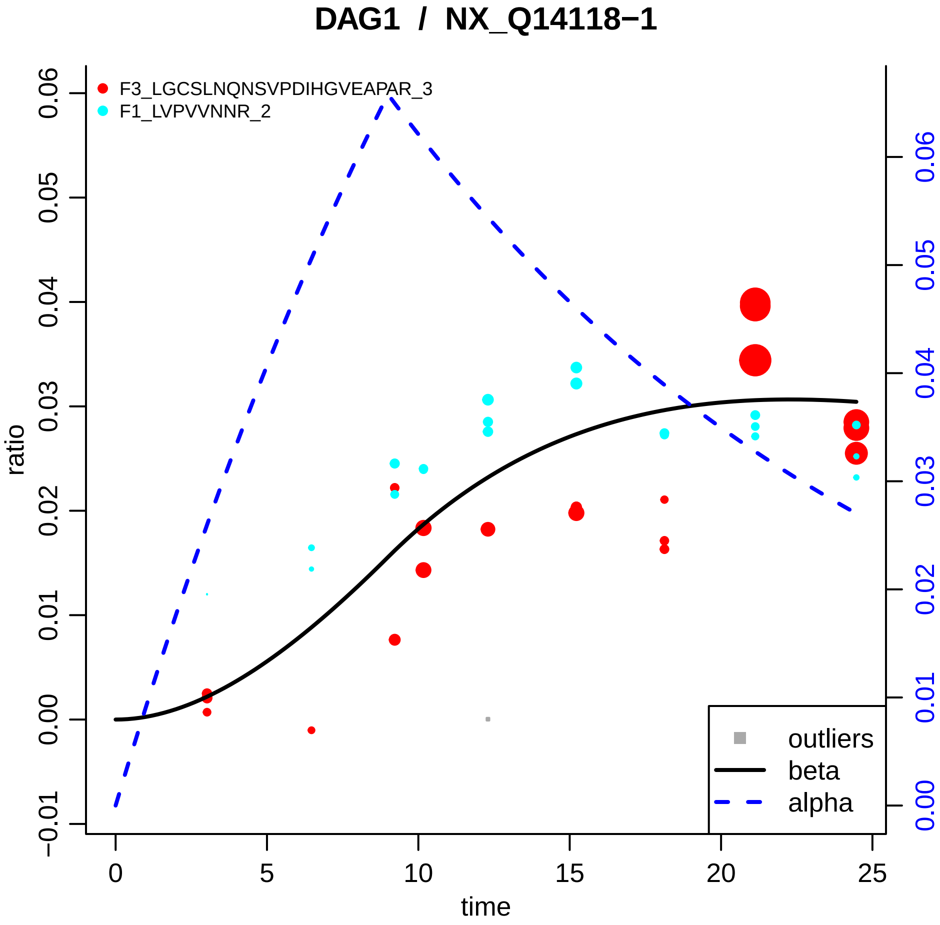

DAG1 / NX_Q14118−1
F3_LGCSLNQNSVPDIHGVEAPAR_3
F1_LVPVVNNR_2
ratio
time
outliers
beta
alpha
0.06
0.05
0.04
0.03
0.02
0.01
0.00
−0.01
0
5
10
15
20
25

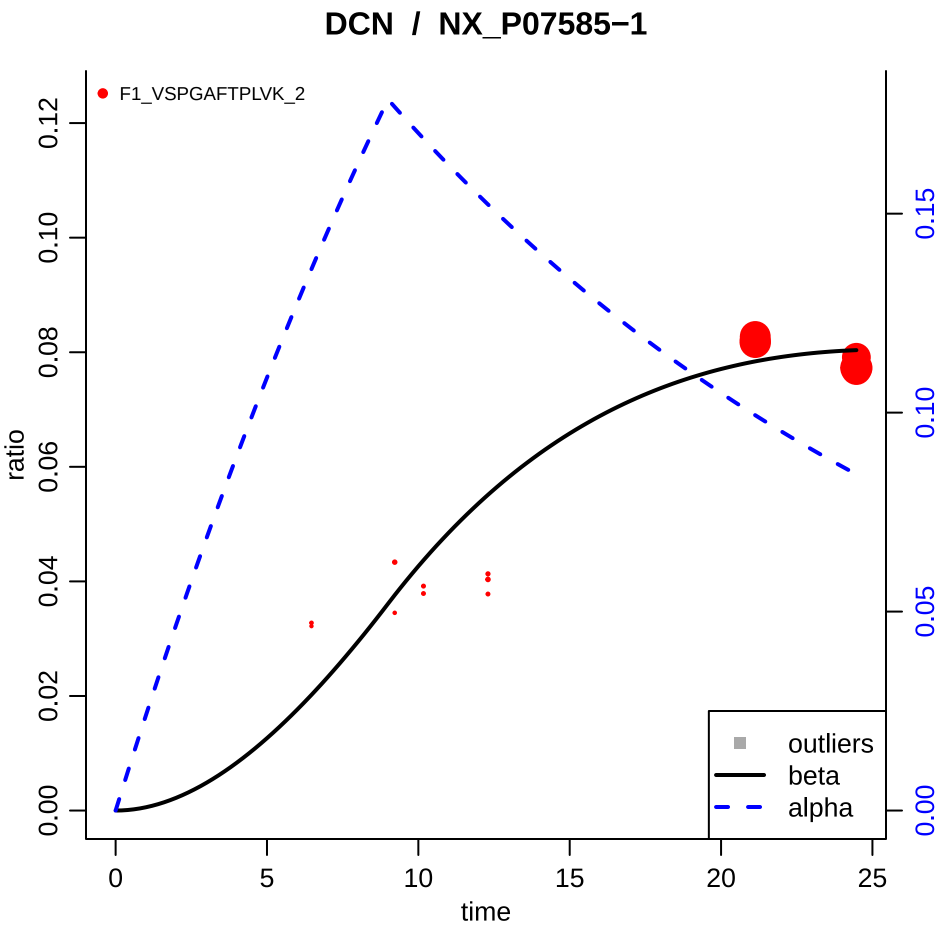

DCN / NX_P07585−1
F1_VSPGAFTPLVK_2
ratio
0.00
0.02
0.04
0.06
0.08
0.10
0.12
0.00
0.05
0.10
0.15
0
5
10
15
20
25
time
outliers
beta
alpha

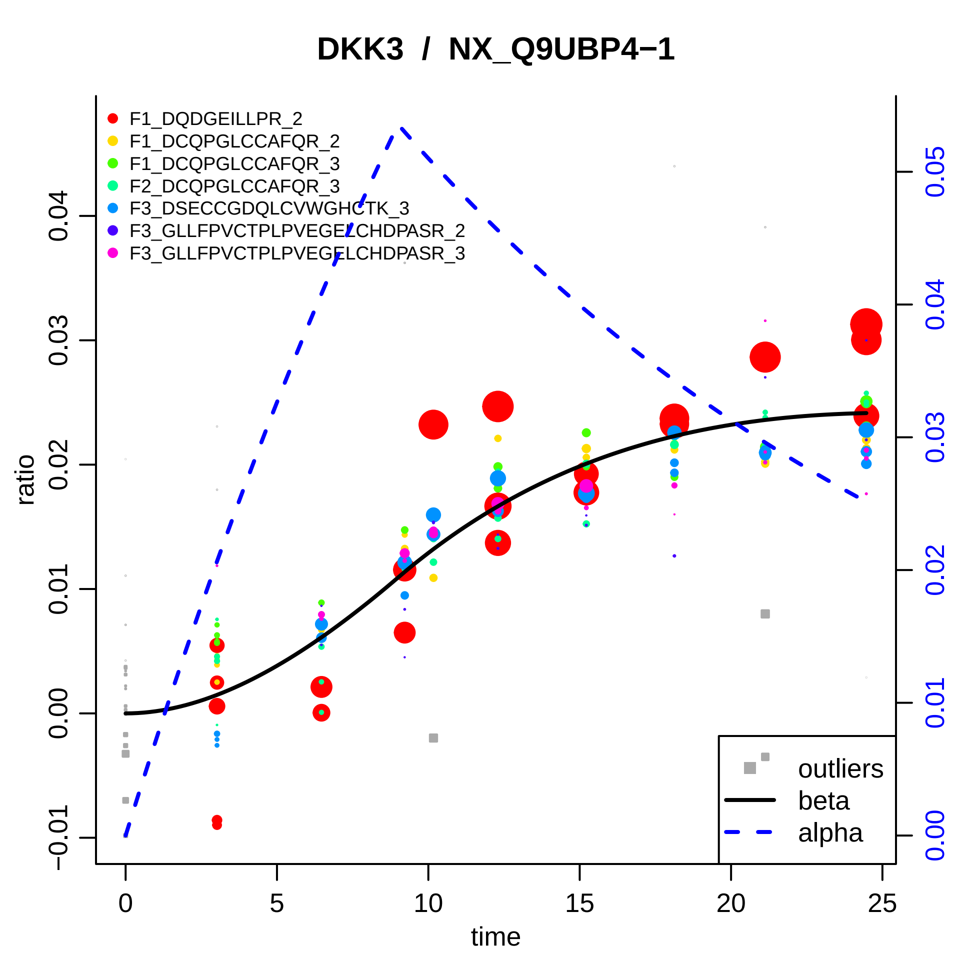
DKK3 / NX_Q9UBP4−1
F1_DQDGEILLPR_2
F1_DCQPGLCCAFQR_2
F1_DCQPGLCCAFQR_3
F2_DCQPGLCCAFQR_3
F3_DSECCGDQLCVWGHCTK_3
F3_GLLFPVCTPLPVEGELCHDPASR_2
F3_GLLFPVCTPLPVEGELCHDPASR_3
ratio
time
outliers
beta
alpha
0.04
0.03
0.02
0.01
0.00
−0.01
0.05
0
5
10
15
20
25

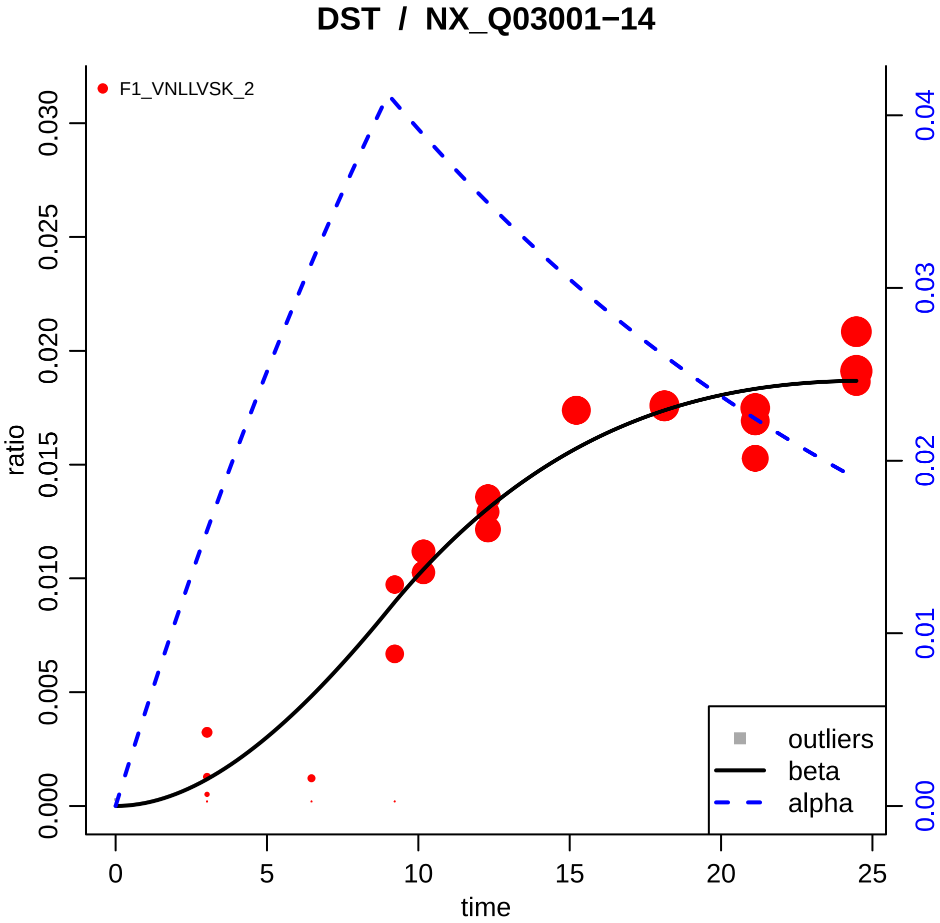

DST / NX_Q03001−14
F1_VNLLVSK_2
ratio
0.000
0.005
0.010
0.015
0.020
0.025
0.030
time
0
5
10
15
20
25
0.00
0.01
0.02
0.03
0.04
outliers
beta
alpha

### ECM1 / NX_Q16610−1

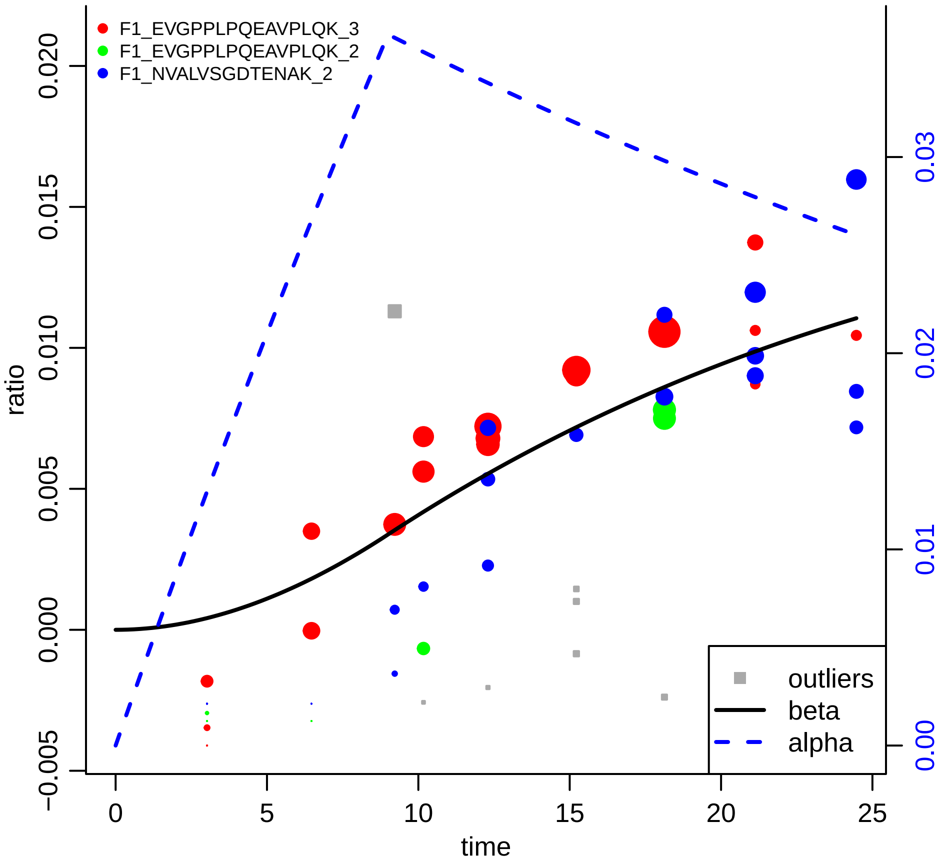

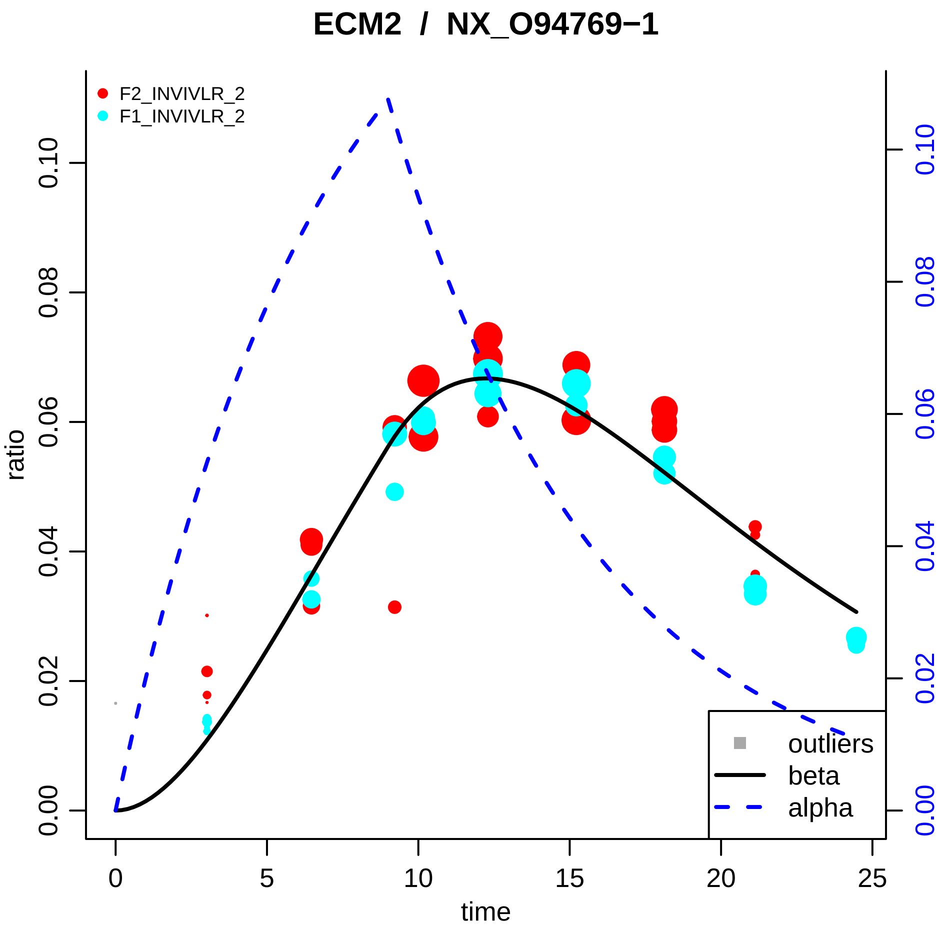
ECM2 / NX_O94769−1
F2_INVIVLR_2
F1_INVIVLR_2
ratio
0.00
0.02
0.04
0.06
0.08
0.10
time
0
5
10
15
20
25
outliers
beta
alpha

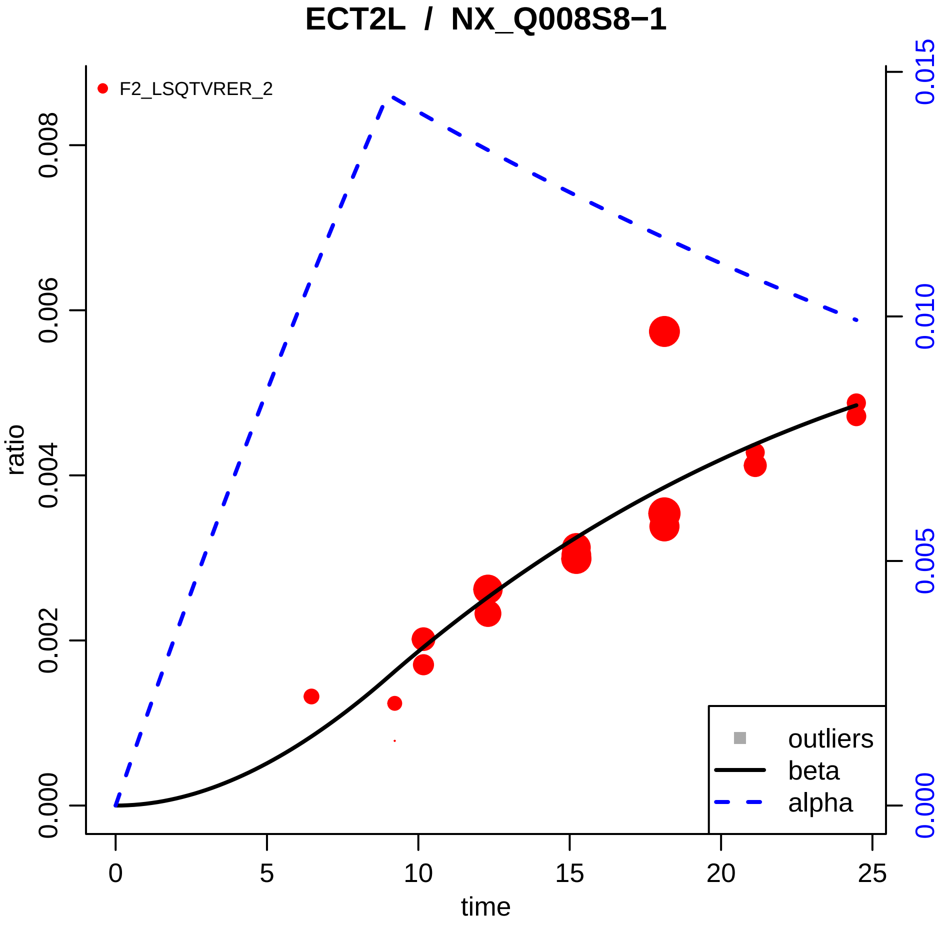

ECT2L / NX_Q008S8−1
F2_LSQTVRER_2
ratio
time
0.000
0.002
0.004
0.006
0.008
0.000
0.005
0.010
0.015
0
5
10
15
20
25
outliers
beta
alpha

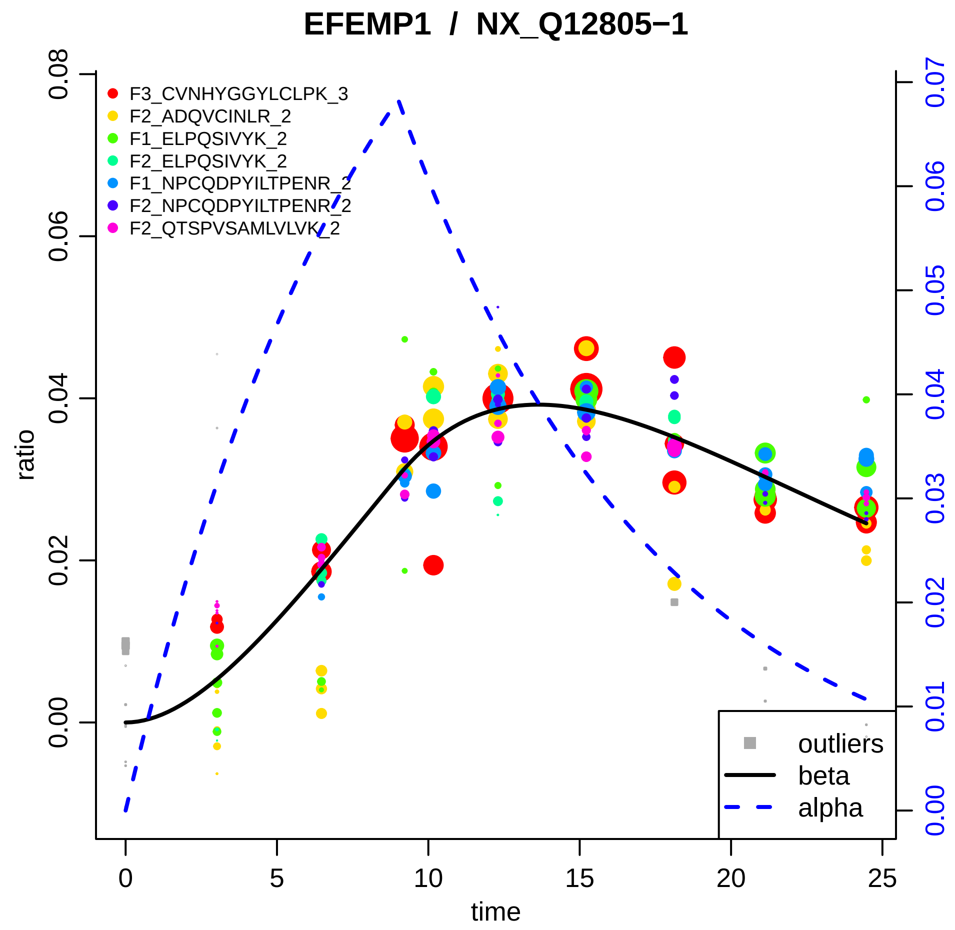

EFEMP1 / NX_Q12805−1
ratio
time
0.00
0.02
0.04
0.06
0.08
0
5
10
15
20
25
0.00
0.01
0.02
0.03
0.04
0.05
0.06
0.07
F3_CVNHYGGYLCLPK_3
F2_ADQVCINLR_2
F1_ELPQSIVYK_2
F2_ELPQSIVYK_2
F1_NPCQDPYILTPENR_2
F2_NPCQDPYILTPENR_2
F2_QTSPVSAMLVLVK_2
outliers
beta
alpha

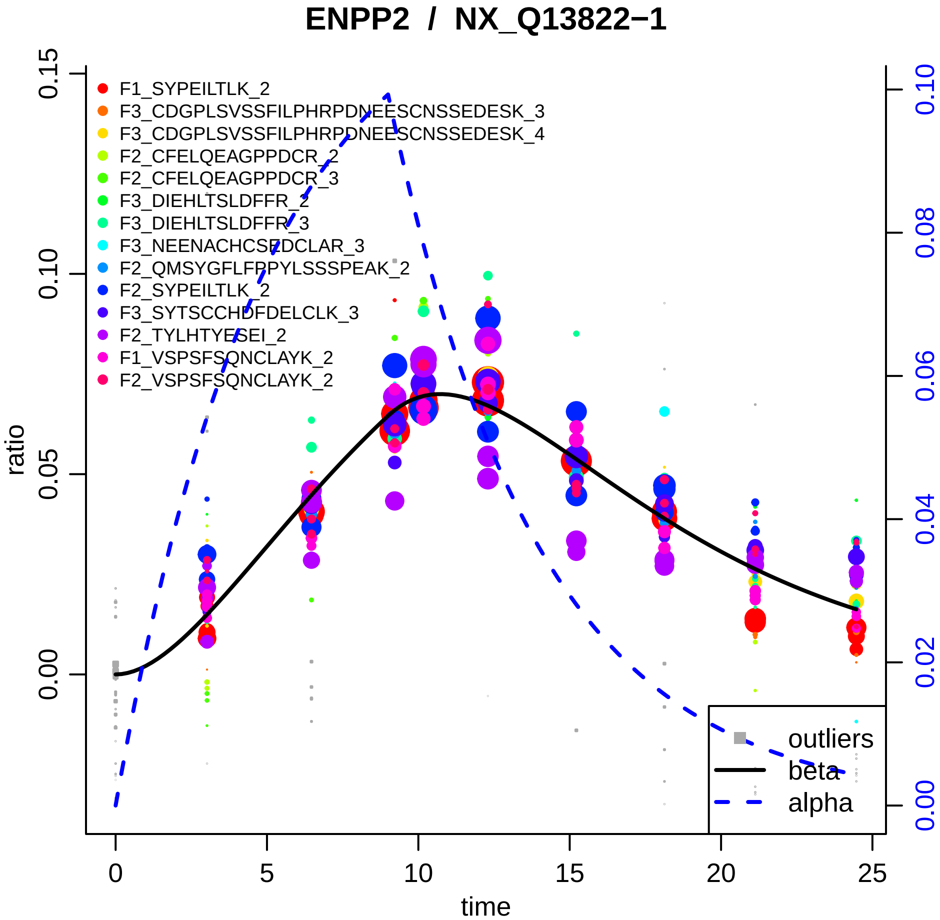

ENPP2 / NX_Q13822−1
F1_SYPEILTLK_2
F3_CDGPLSVSSFILPHRPDNEESCNSSEDESK_3
F3_CDGPLSVSSFILPHRPDNEESCNSSEDESK_4
F2_CFELQEAGPPDCR_2
F2_CFELQEAGPPDCR_3
F3_DIEHLTSLDFFR_2
F3_DIEHLTSLDFFR_3
F3_NEENACHCSEDCLAR_3
F2_QMSYGFLFPPYLSSSPEAK_2
F2_SYPEILTLK_2
F3_SYTSCCHDFDELCLK_3
F2_TYLHTYESEI_2
F1_VSPSFSQNCLAYK_2
F2_VSPSFSQNCLAYK_2
outliers
beta
alpha
ratio
time
0.00
0.05
0.10
0.15
0.00
0.02
0.04
0.06
0.08
0.10
0
5
10
15
20
25

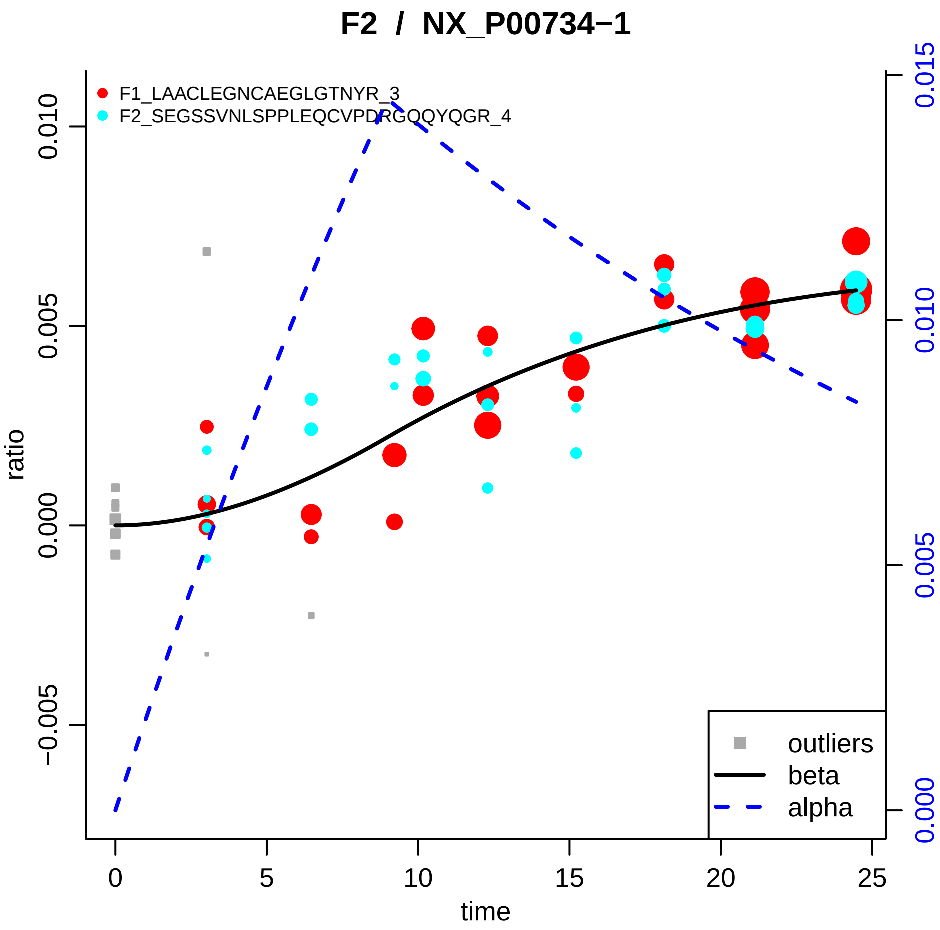

F2 / NX_P00734−1
F1_LAACLEGNCAEGLGTNYR_3
F2_SEGSSVNLSPPLEQCVPDRGQQYQGR_4
ratio
time
0.010
0.005
0.000
−0.005
0.015
0.010
0.005
0.000
0
5
10
15
20
25
outliers
beta
alpha

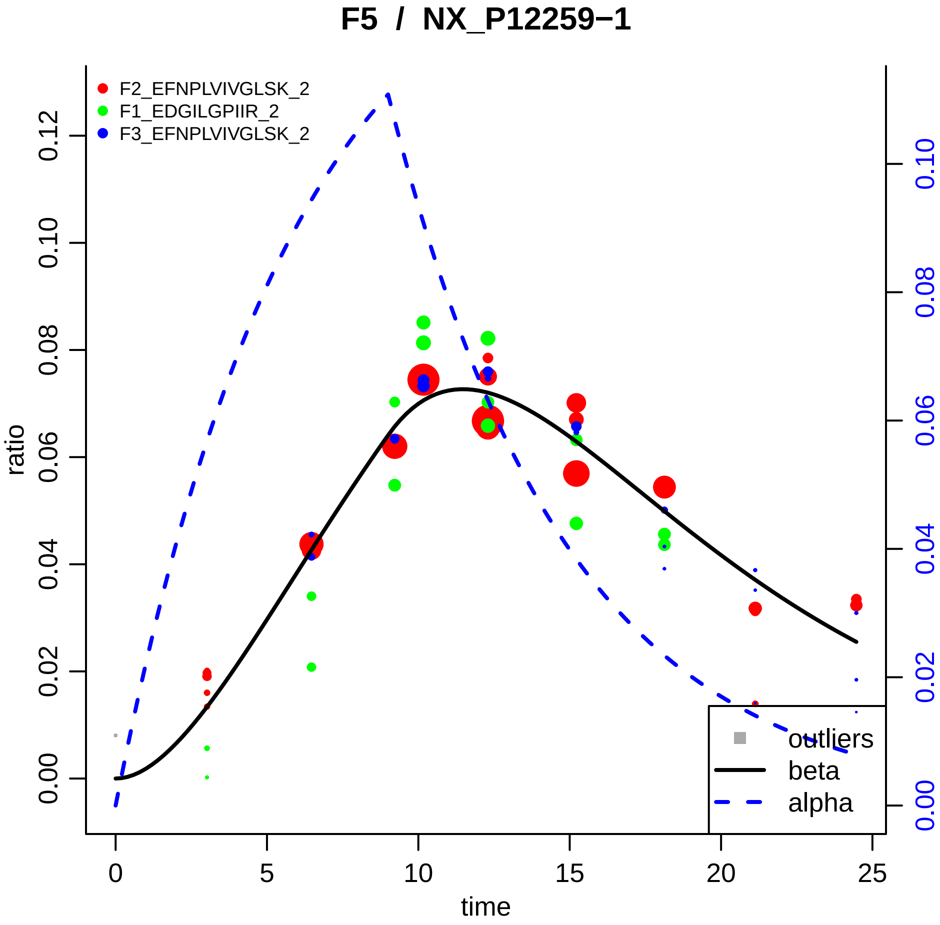
F5 / NX_P12259−1
F2_EFNPLVIVGLSK_2
F1_EDGILGPIIR_2
F3_EFNPLVIVGLSK_2
ratio
time
0.00
0.02
0.04
0.06
0.08
0.10
0.12
0
5
10
15
20
25
0.00
0.02
0.04
0.06
0.08
0.10
outliers
beta
alpha

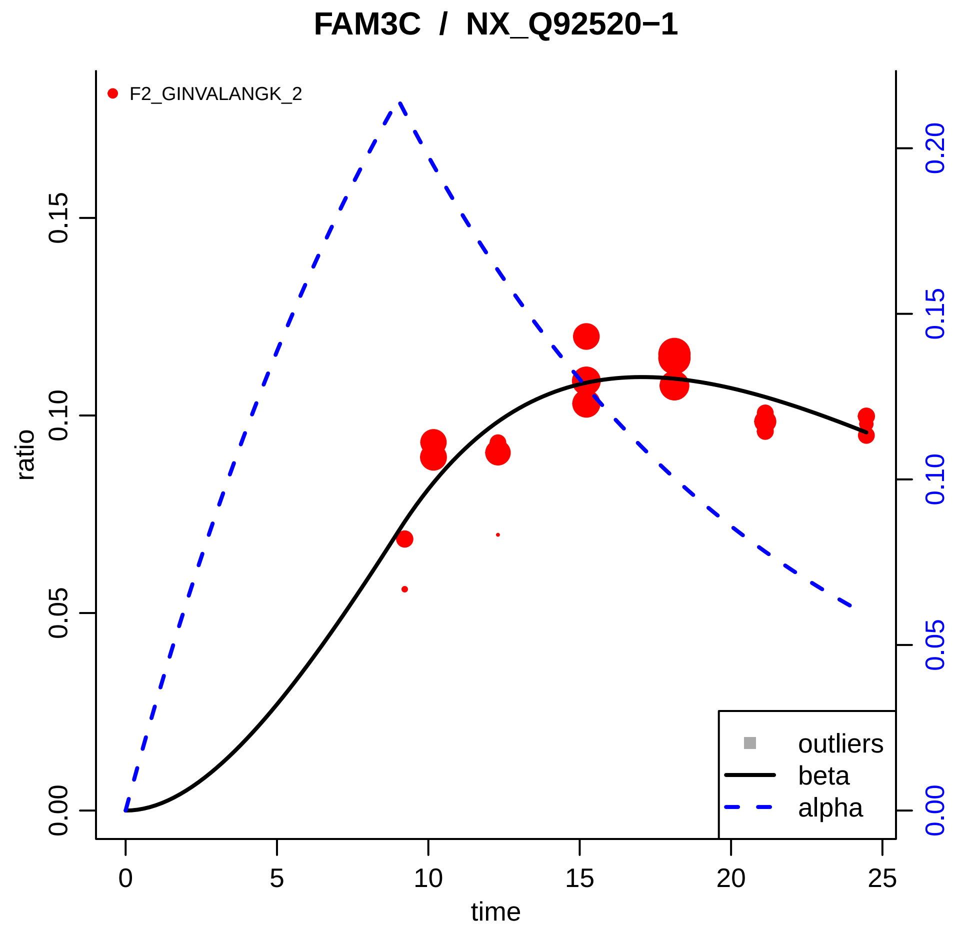

FAM3C / NX_Q92520−1
F2_GINVALANGK_2
ratio
time
0.00
0.05
0.10
0.15
0.20
0
5
10
15
20
25
outliers
beta
alpha

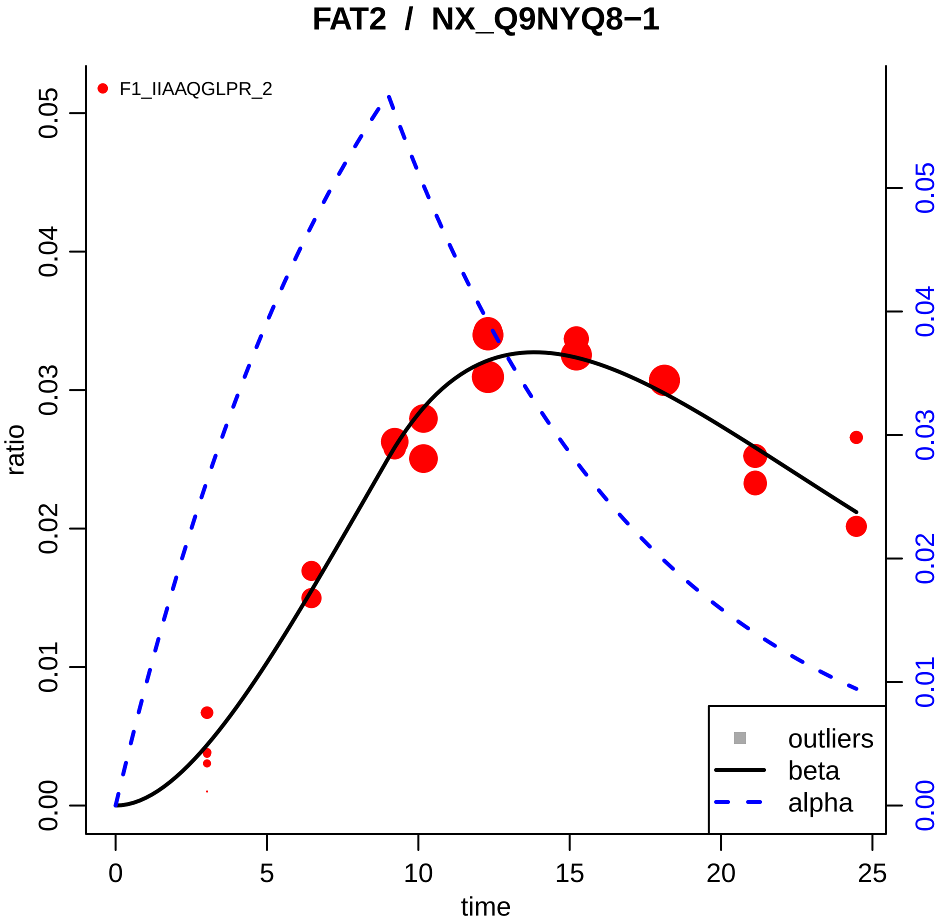
FAT2 / NX_Q9NYQ8−1
F1_IIAAQGLPR_2
ratio
time
0.00
0.01
0.02
0.03
0.04
0.05
0
5
10
15
20
25
outliers
beta
alpha

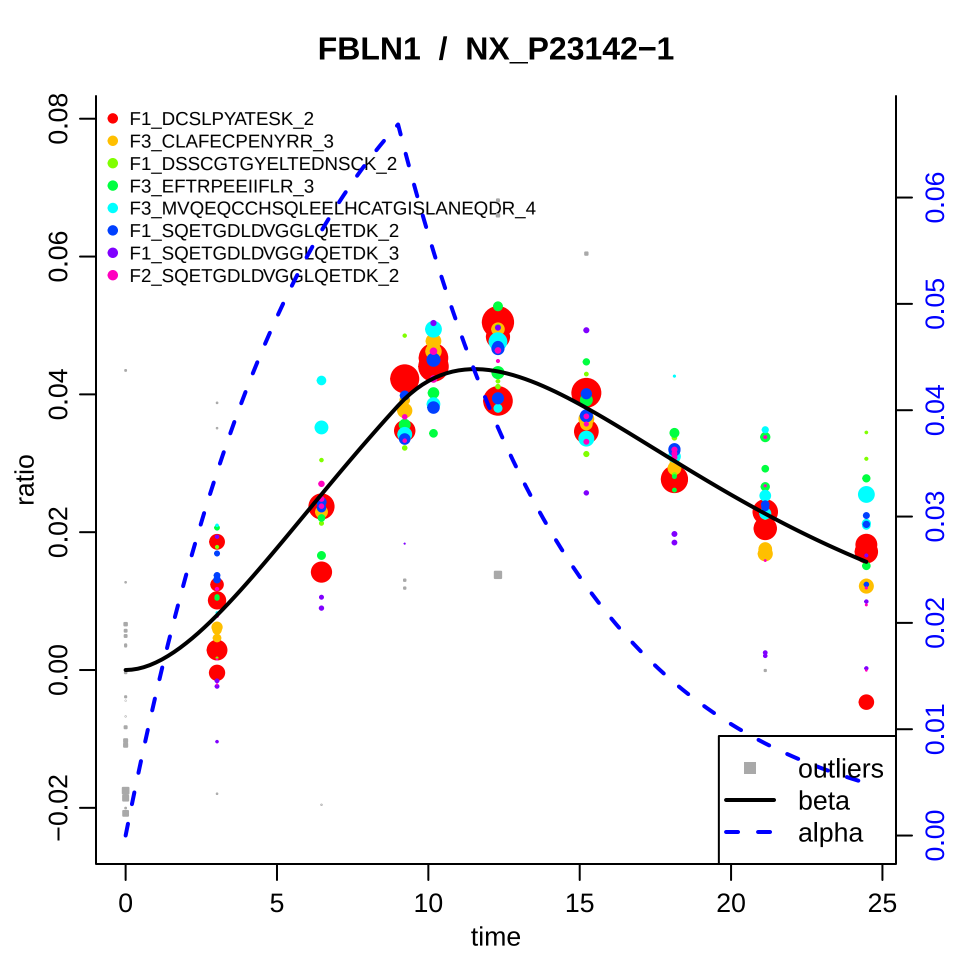

FBLN1 / NX_P23142−1
F1_DCSLPYATESK_2
F3_CLAFECPENYRR_3
F1_DSSCGTGYELTEDNSCK_2
F3_EFTRPEEIIFLR_3
F3_MVQEQCCHSQLEELHCATGISLANEQDR_4
F1_SQETGDLDVGGLQETDK_2
F1_SQETGDLDVGGLQETDK_3
F2_SQETGDLDVGGLQETDK_2
outliers
beta
alpha
ratio
time
0.08
0.06
0.04
0.02
0.00
−0.02
0
5
10
15
20
25
0.06
0.05
0.04
0.03
0.02
0.01
0.00

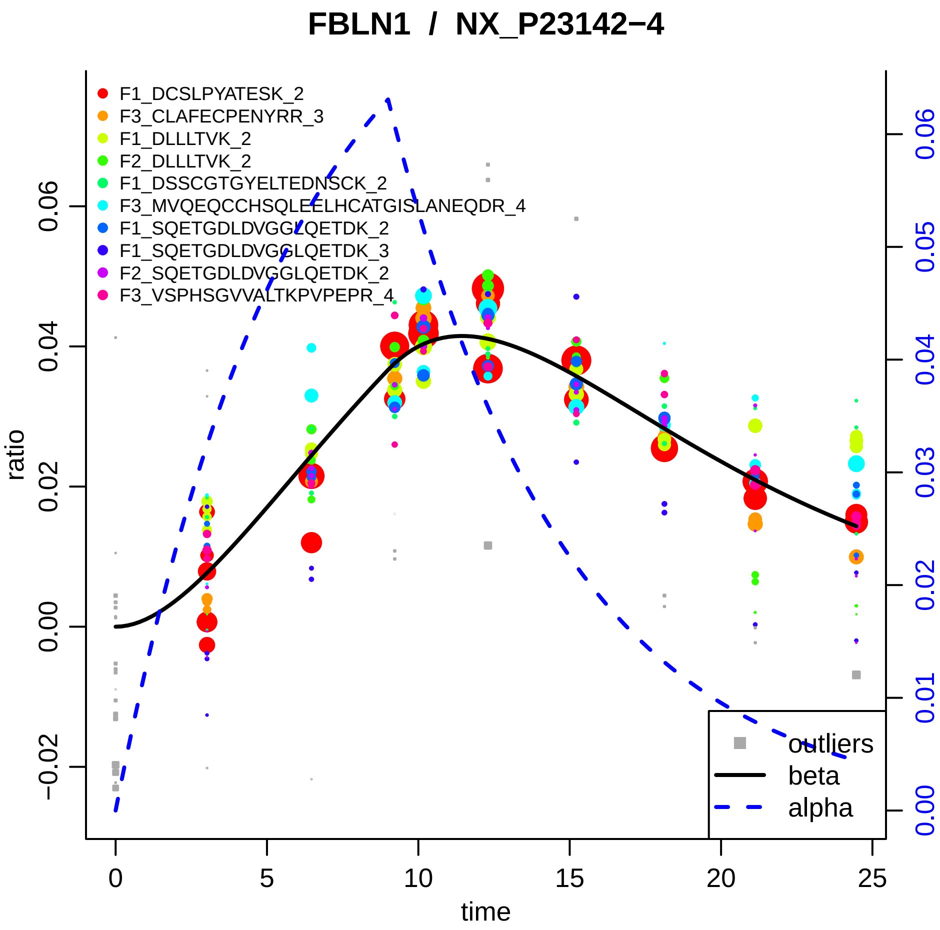

FBLN1 / NX_P23142−4
F1_DCSLPYATESK_2
F3_CLAFECPENYRR_3
F1_DLLLTVK_2
F2_DLLLTVK_2
F1_DSSCGTGYELTEDNSCK_2
F3_MVQEQCCHSQLEELHCATGISLANEQDR_4
F1_SQETGDLDVGGLQETDK_2
F1_SQETGDLDVGGLQETDK_3
F2_SQETGDLDVGGLQETDK_2
F3_VSPHSGVVALTKPVPEPR_4
outliers
beta
alpha
ratio
time
0.06
0.04
0.02
0.00
−0.02
0
5
10
15
20
25
0.00
0.01
0.02
0.03
0.04
0.05
0.06

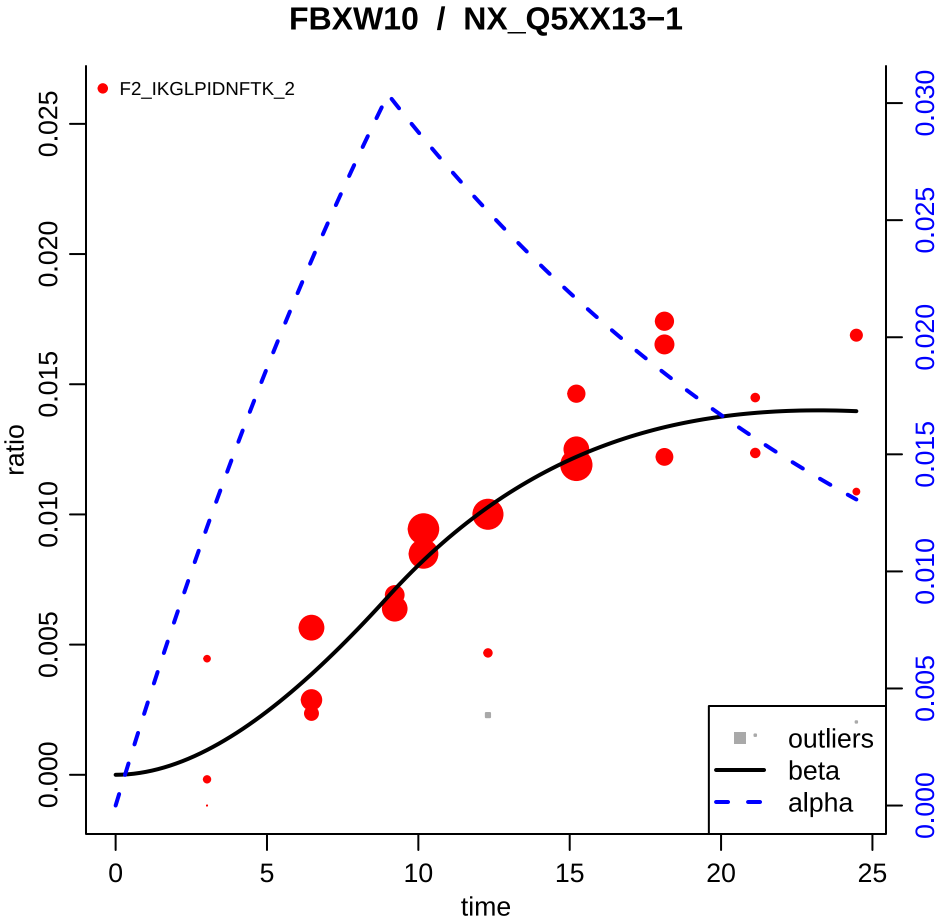

FBXW10 / NX_Q5XX13−1
F2_IKGLPIDNFTK_2
ratio
time
0.000
0.005
0.010
0.015
0.020
0.025
0.030
0
5
10
15
20
25
outliers
beta
alpha

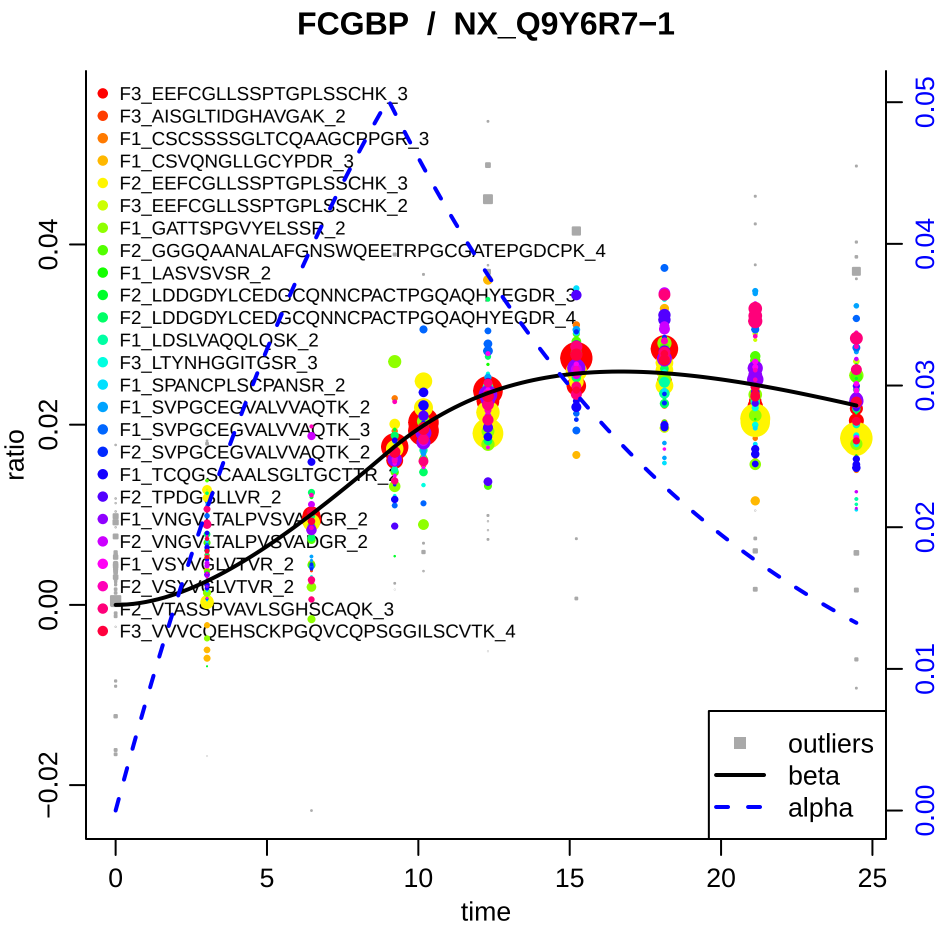

FCGBP / NX_Q9Y6R7−1
F3_EEFCGLLSSPTGPLSSCHK_3
F3_AISGLTIDGHAVGAK_2
F1_CSCSSSSGLTCQAAGCPPGR_3
F1_CSVQNGLLGCYPDR_3
F2_EEFCGLLSSPTGPLSSCHK_3
F3_EEFCGLLSSPTGPLSSCHK_2
F1_GATTSPGVYELSSR_2
F2_GGGQAANALAFGNSWQEETRPGCGATEPGDCPK_4
F1_LASVSVSR_2
F2_LDDGDYLCEDGCQNNCPACTPGQAQHYEGDR_3
F2_LDDGDYLCEDGCQNNCPACTPGQAQHYEGDR_4
F1_LDSLVAQQLQSK_2
F3_LTYNHGGITGSR_3
F1_SPANCPLSCPANSR_2
F1_SVPGCEGVALVVAQTK_2
F1_SVPGCEGVALVVAQTK_3
F2_SVPGCEGVALVVAQTK_2
F1_TCQGSCAALSGLTGCTTR_2
F2_TPDGGLLVR_2
F1_VNGVLTALPVSVADGR_2
F2_VNGVLTALPVSVADGR_2
F1_VSYVGLVTVR_2
F2_VSYVGLVTVR_2
F2_VTASSPVAVLSGHSCAQK_3
F3_VVVCQEHSCKPGQVCQPSGGILSCVTK_4
ratio
−0.02
0.00
0.02
0.04
time
0
5
10
15
20
25
0.00
0.01
0.02
0.03
0.04
0.05
outliers
beta
alpha

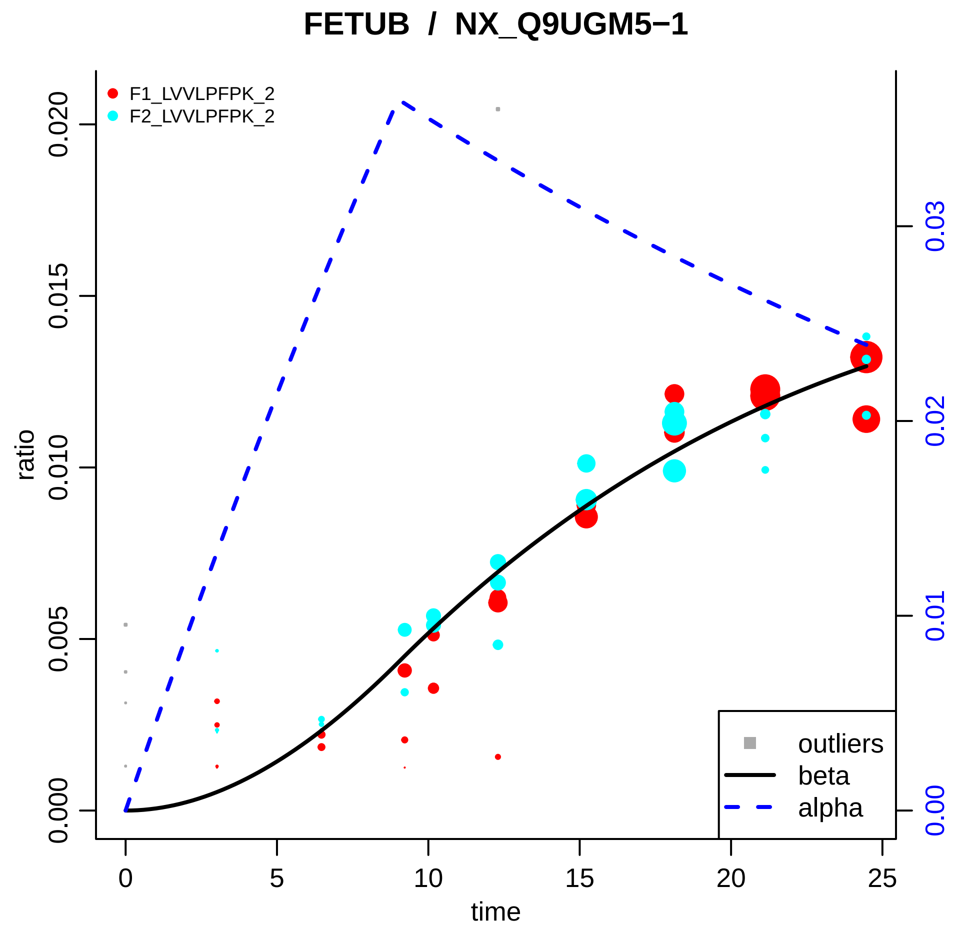
FETUB / NX_Q9UGM5−1
F1_LVVLPFPK_2
F2_LVVLPFPK_2
ratio
0.000
0.005
0.010
0.015
0.020
0.00
0.01
0.02
0.03
0
5
10
15
20
25
time
outliers
beta
alpha

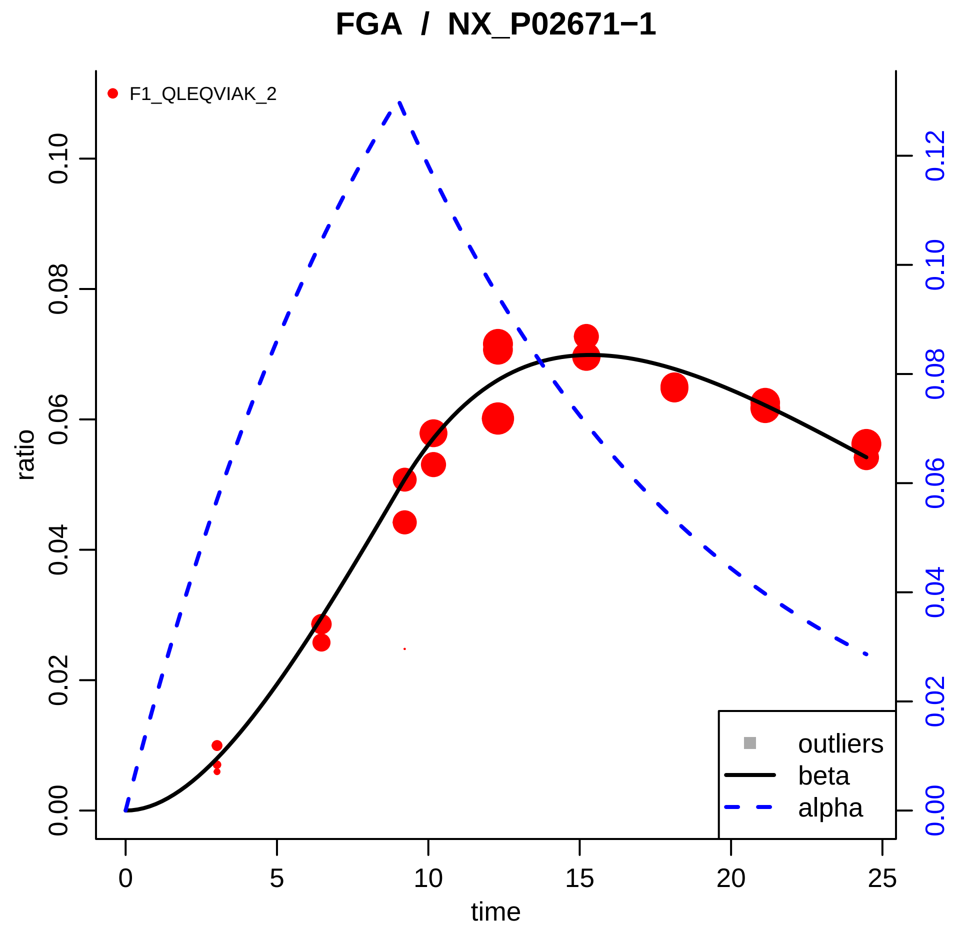

FGA / NX_P02671−1
F1_QLEQVIAK_2
ratio
0.00
0.02
0.04
0.06
0.08
0.10
0.12
time
0
5
10
15
20
25
outliers
beta
alpha

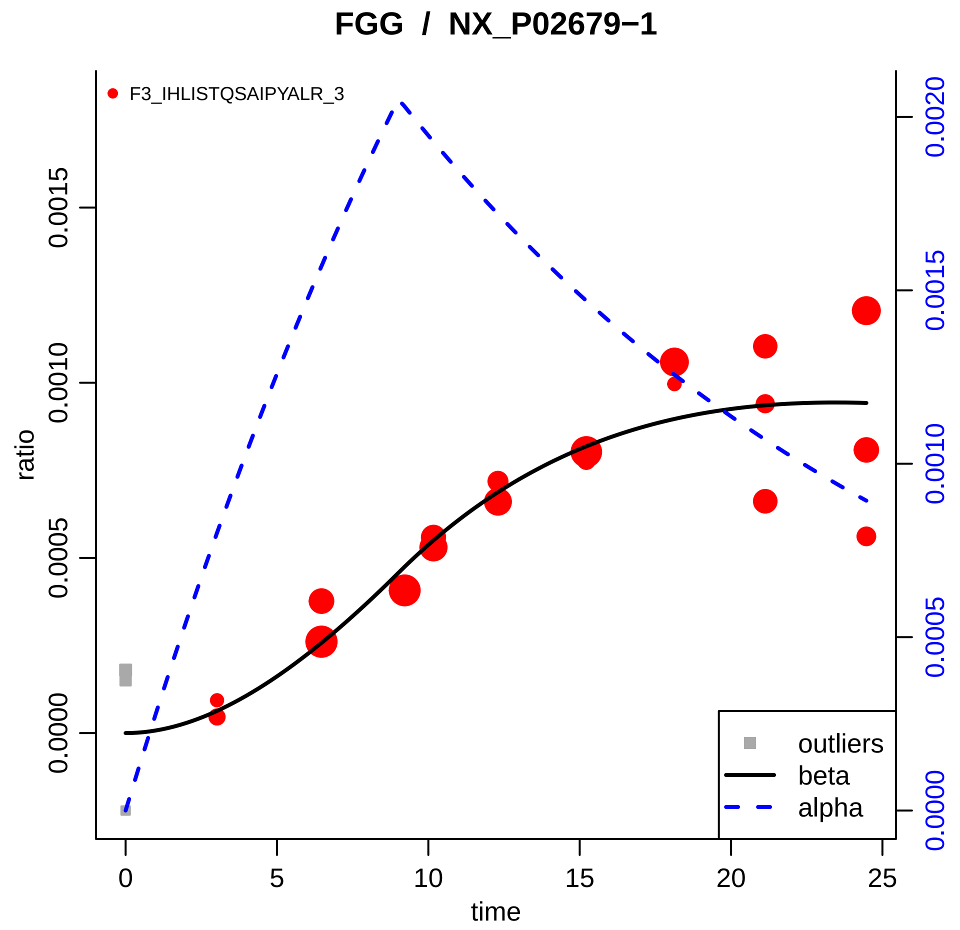

FGG / NX_P02679−1
F3_IHLISTQSAIPYALR_3
ratio
time
0.0000
0.0005
0.0010
0.0015
0.0020
0
5
10
15
20
25
outliers
beta
alpha

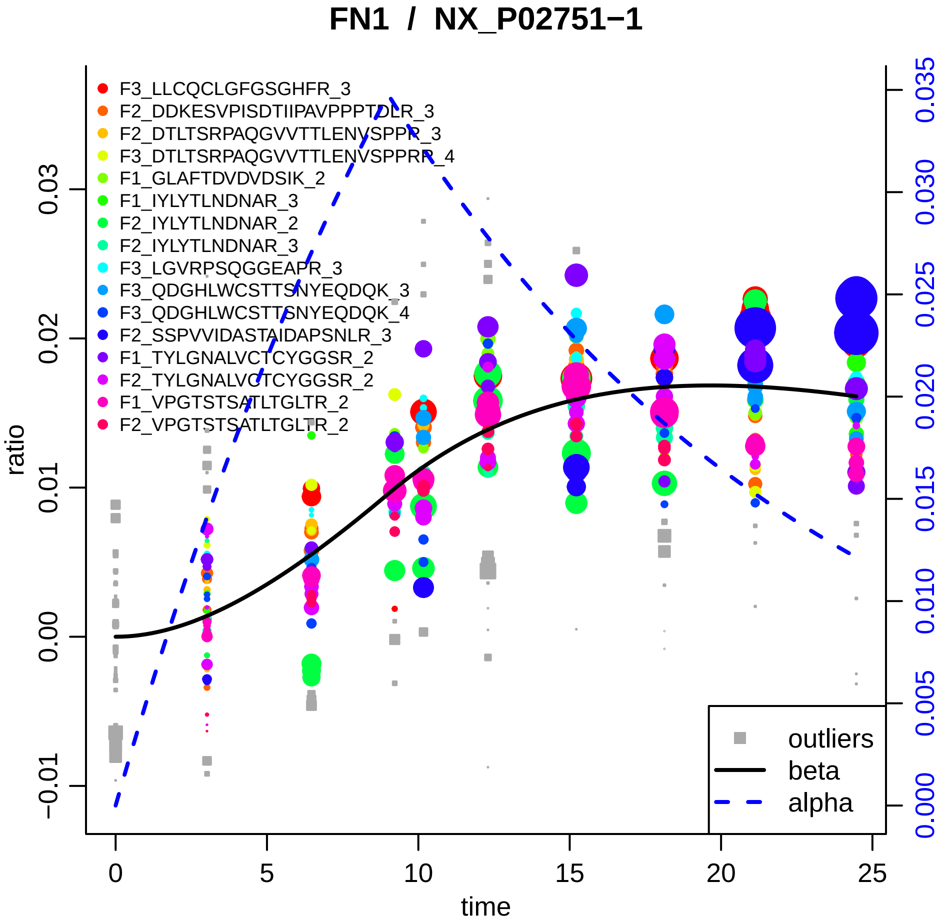

FN1 / NX_P02751−1
F3_LLCQCLGFGSGHFR_3
F2_DDKESVPISDTIIPAVPPPTDLR_3
F2_DTLTSRPAQGVVTTLENVSPPR_3
F3_DTLTSRPAQGVVTTLENVSPPRR_4
F1_GLAFTDVDVDSIK_2
F1_IYLYTLNDNAR_3
F2_IYLYTLNDNAR_2
F2_IYLYTLNDNAR_3
F3_LGVRPSQGGEAPR_3
F3_QDGHLWCSTTSNYEQDQK_3
F3_QDGHLWCSTTSNYEQDQK_4
F2_SSPVVIDASTAIDAPSNLR_3
F1_TYLGNALVCTCYGGSR_2
F2_TYLGNALVCTCYGGSR_2
F1_VPGTSTSATLTGLTR_2
F2_VPGTSTSATLTGLTR_2
outliers
beta
alpha
ratio
time
−0.01
0.00
0.01
0.02
0.03
0
5
10
15
20
25
0.000
0.005
0.010
0.015
0.020
0.025
0.030
0.035

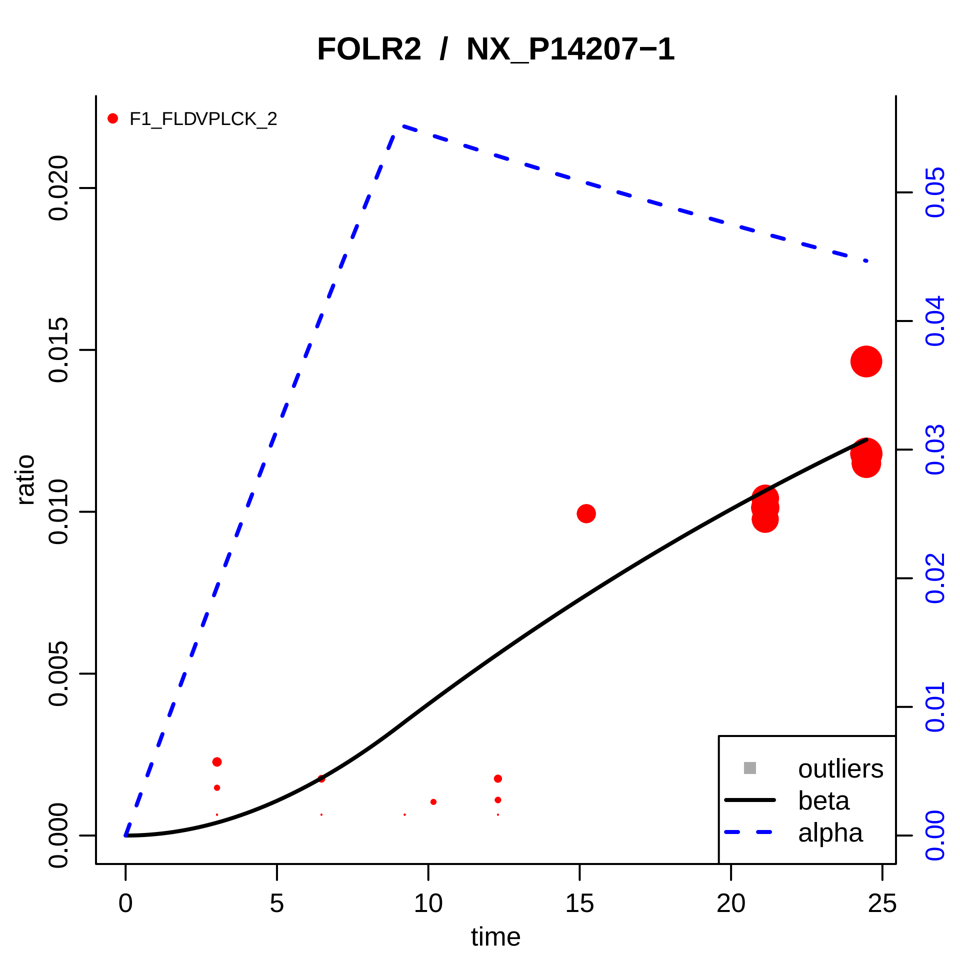

FOLR2 / NX_P14207−1
F1_FLDVPLCK_2
ratio
time
0.000
0.005
0.010
0.015
0.020
0
5
10
15
20
25
0.00
0.01
0.02
0.03
0.04
0.05
outliers
beta
alpha

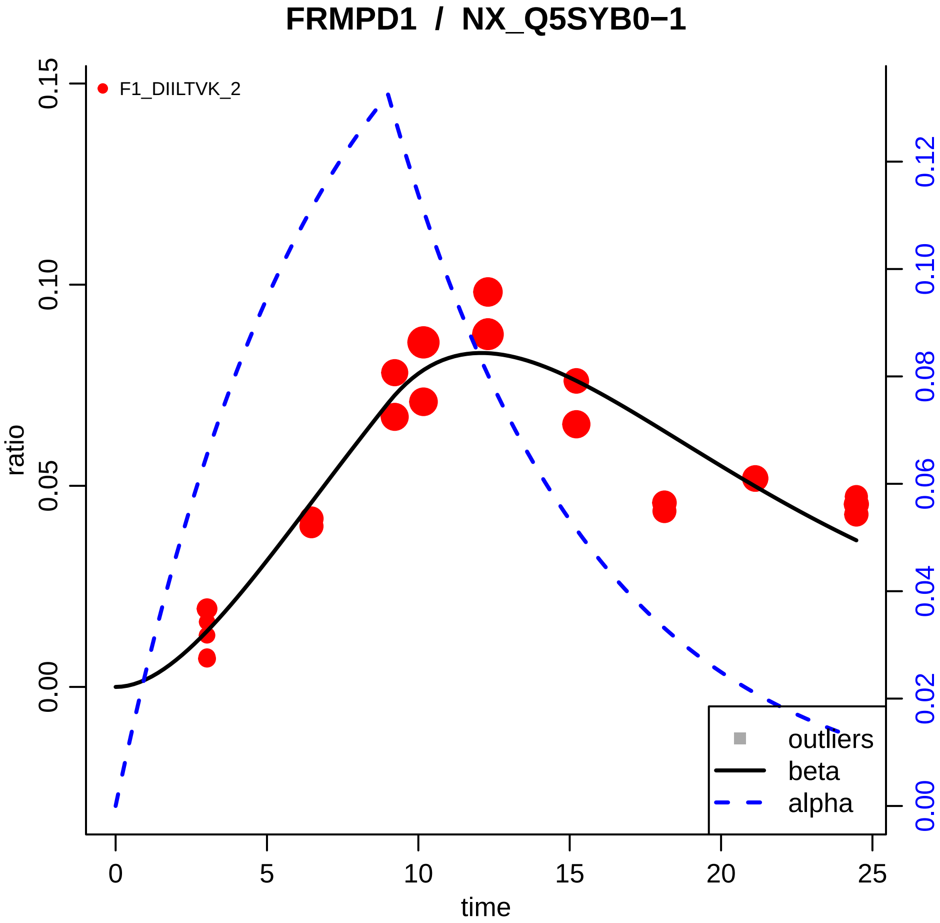

FRMPD1 / NX_Q5SYB0−1
F1_DIILTVK_2
ratio
time
outliers
beta
alpha
0.00
0.05
0.10
0.15
0
5
10
15
20
25
0.00
0.02
0.04
0.06
0.08
0.10
0.12

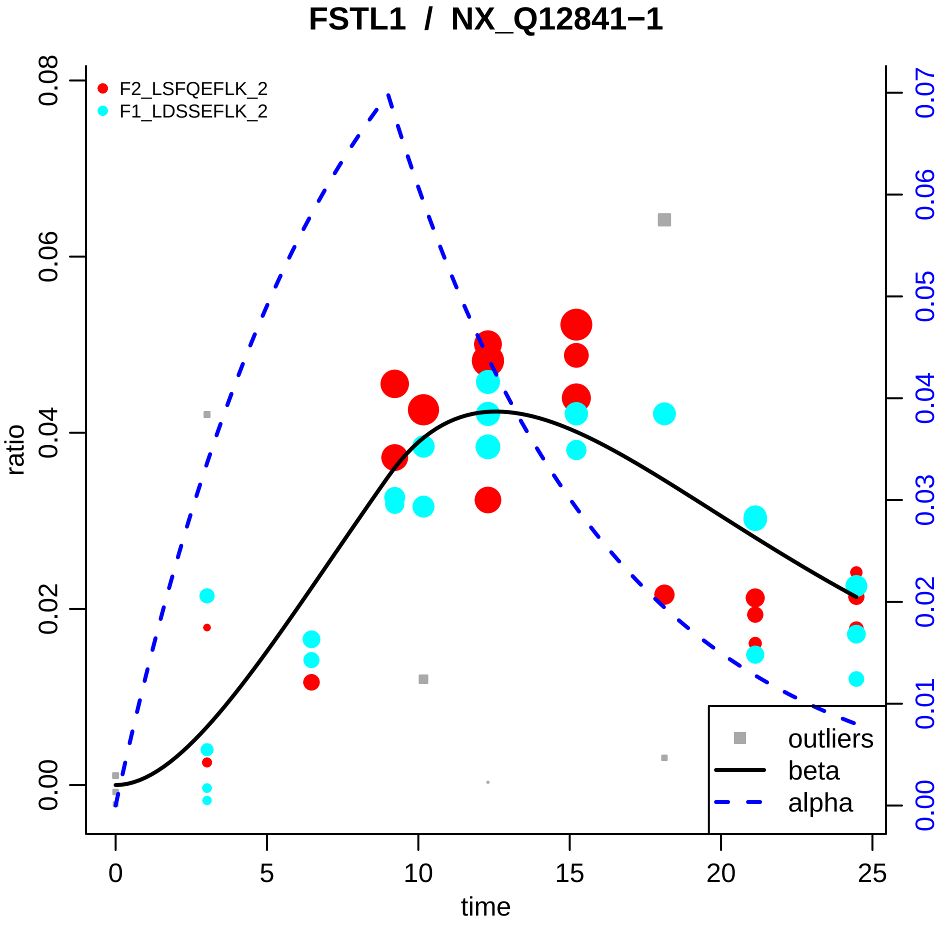
FSTL1 / NX_Q12841−1
F2_LSFQEFLK_2
F1_LDSSEFLK_2
outliers
beta
alpha
ratio
time
0.00
0.02
0.04
0.06
0.08
0
5
10
15
20
25
0.00
0.01
0.02
0.03
0.04
0.05
0.06
0.07

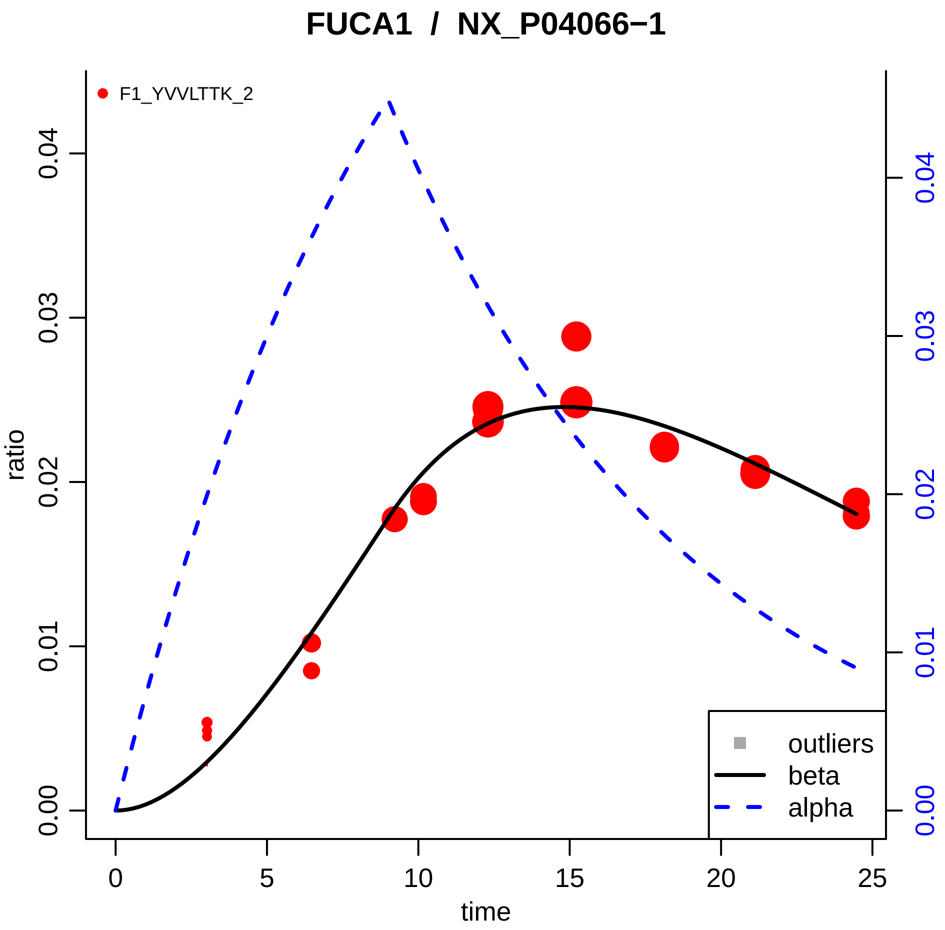

FUCA1 / NX_P04066−1
F1_YVVLTTK_2
ratio
time
0.00
0.01
0.02
0.03
0.04
0
5
10
15
20
25
outliers
beta
alpha

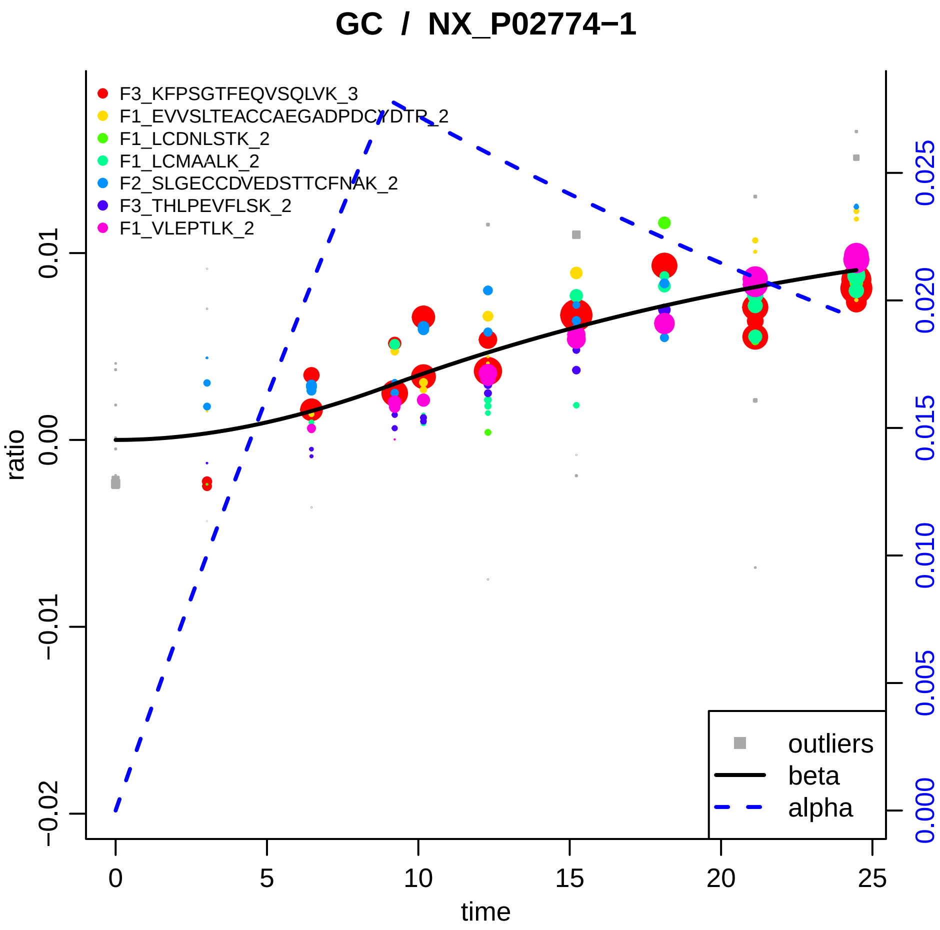

GC / NX_P02774−1
F3_KFPSGTFEQVSQLVK_3
F1_EVVSLTEACCAEGADPDCYDTR_2
F1_LCDNLSTK_2
F1_LCMAALK_2
F2_SLGECCDVEDSTTCFNAK_2
F3_THLPEVFLSK_2
F1_VLEPTLK_2
outliers
beta
alpha
ratio
time
0.01
0.00
−0.01
−0.02
0
5
10
15
20
25
0.000
0.005
0.010
0.015
0.020
0.025

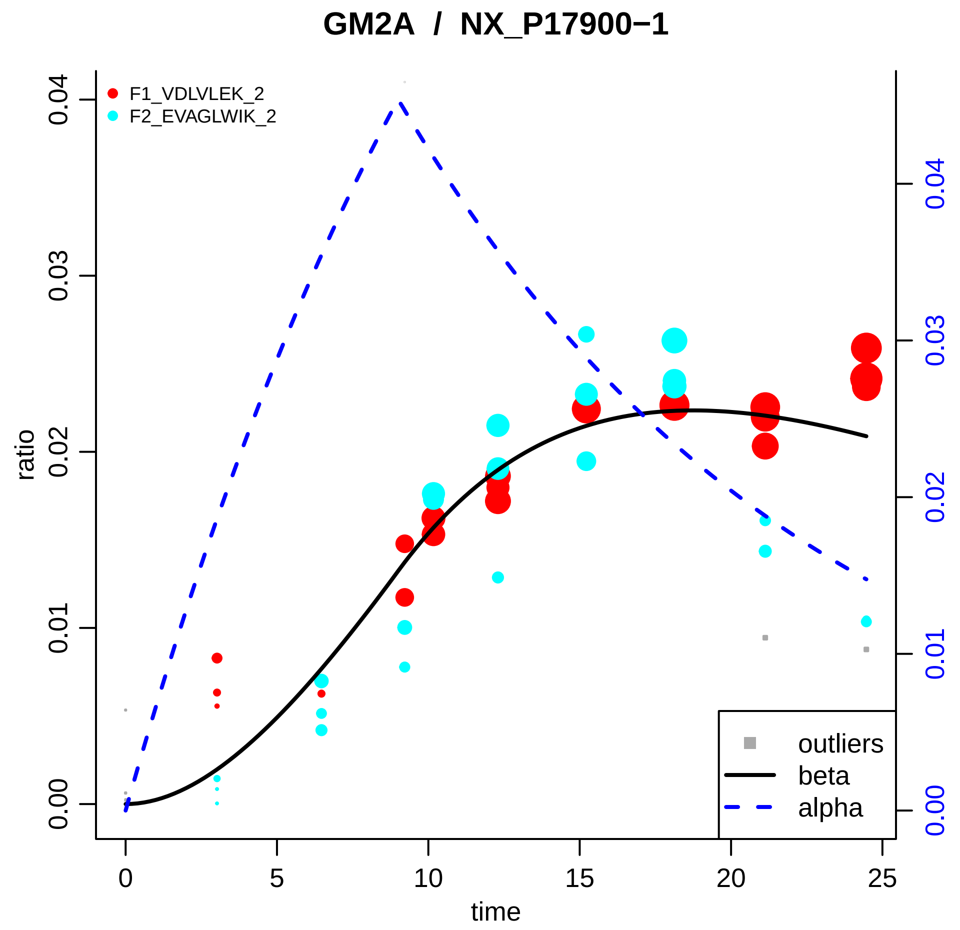

GM2A / NX_P17900−1
F1_VDLVLEK_2
F2_EVAGLWIK_2
ratio
0.00
0.01
0.02
0.03
0.04
time
0
5
10
15
20
25
outliers
beta
alpha

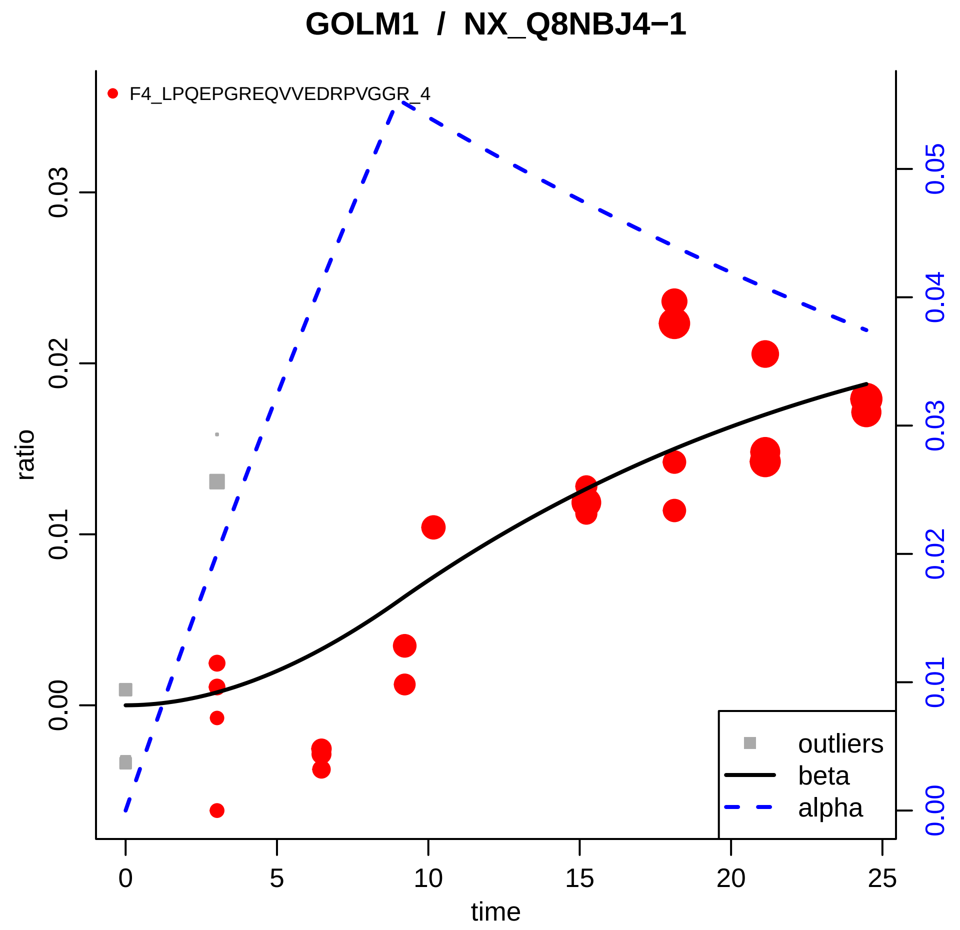

GOLM1 / NX_Q8NBJ4−1
F4_LPQEPGREQVVEDRPVGGR_4
ratio
time
0.00
0.01
0.02
0.03
0
5
10
15
20
25
0.00
0.01
0.02
0.03
0.04
0.05
outliers
beta
alpha

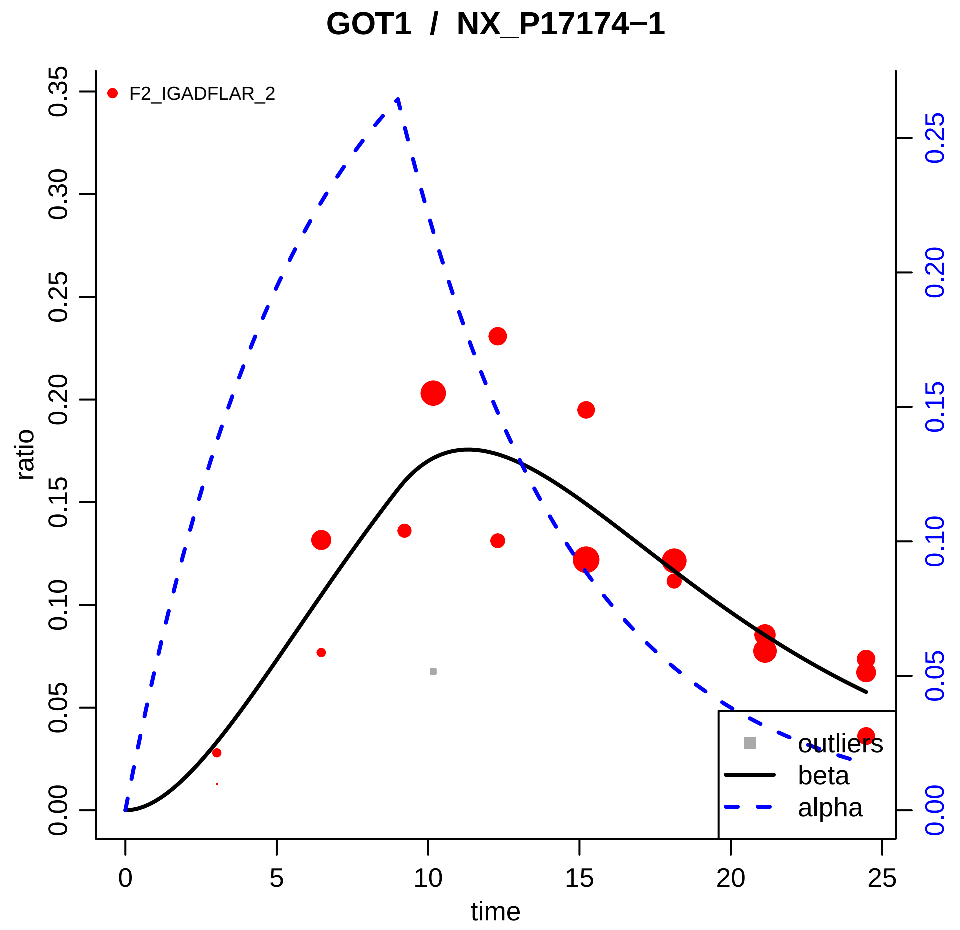
GOT1 / NX_P17174−1
F2_IGADFLAR_2
ratio
time
0.00
0.05
0.10
0.15
0.20
0.25
0.30
0.35
0
5
10
15
20
25
0.00
0.05
0.10
0.15
0.20
0.25
outliers
beta
alpha

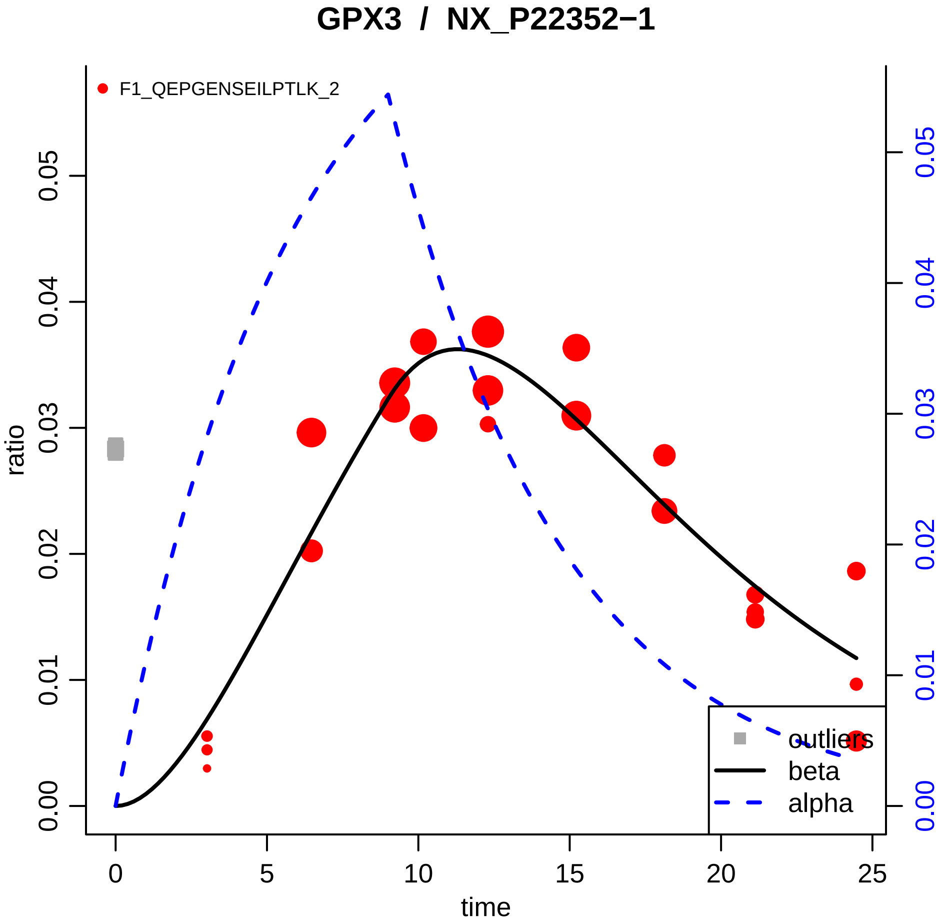
GPX3 / NX_P22352−1
F1_QEPGENSEILPTLK_2
ratio
time
0.00
0.01
0.02
0.03
0.04
0.05
0
5
10
15
20
25
outliers
beta
alpha

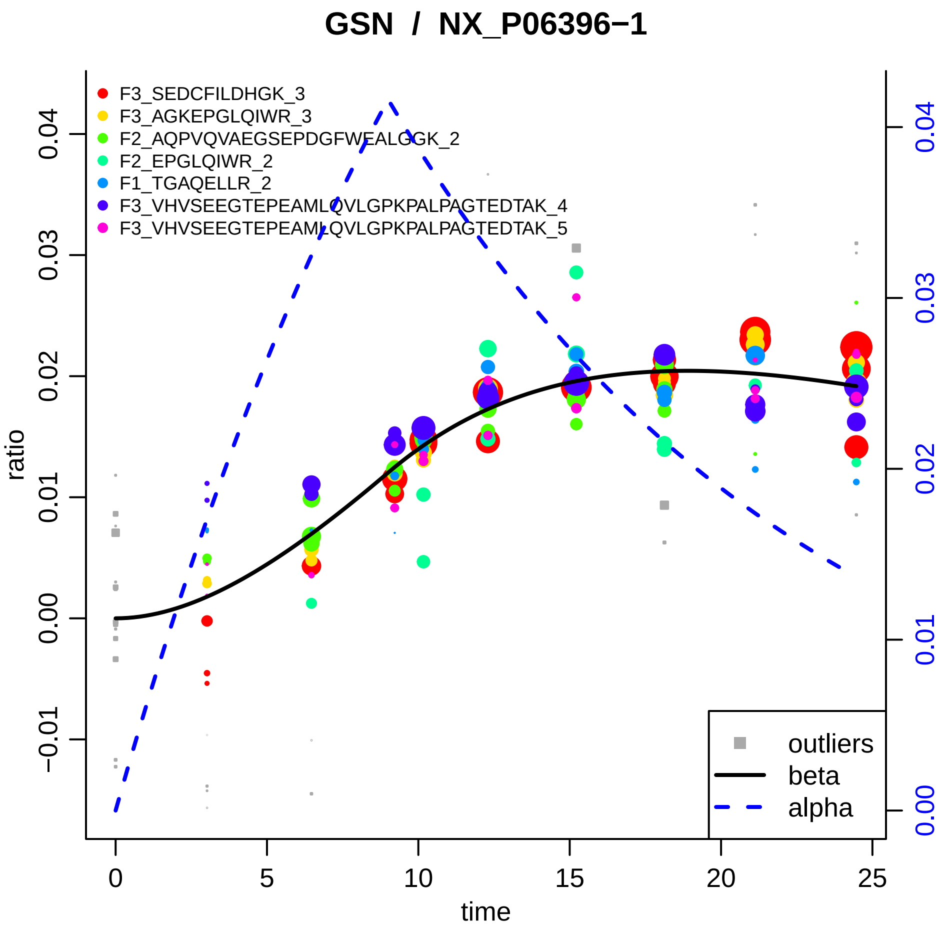

GSN / NX_P06396−1
F3_SEDCFILDHGK_3
F3_AGKEPGLQIWR_3
F2_AQPVQVAEGSEPDGFWEALGGK_2
F2_EPGLQIWR_2
F1_TGAQELLR_2
F3_VHVSEEGTEPEAMLQVLGPKPALPAGTEDTAK_4
F3_VHVSEEGTEPEAMLQVLGPKPALPAGTEDTAK_5
outliers
beta
alpha
ratio
time
0
5
10
15
20
25
−0.01
0.00
0.01
0.02
0.03
0.04

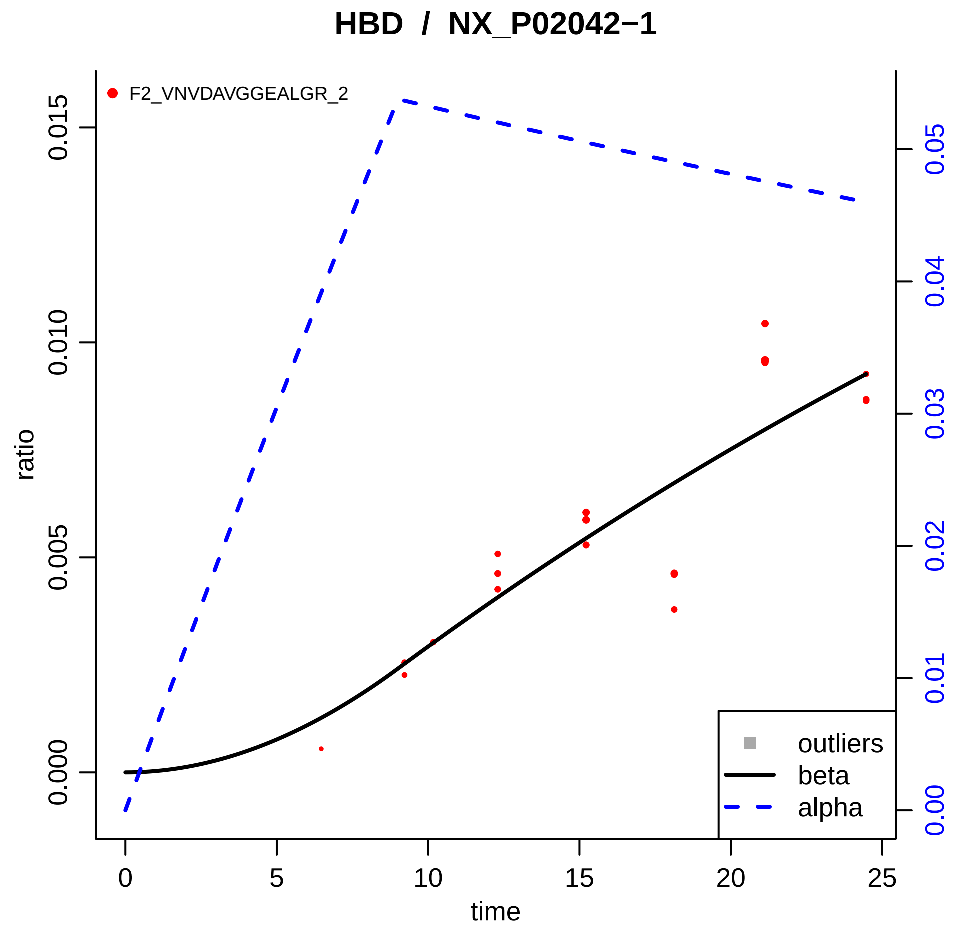

HBD / NX_P02042−1
F2_VNVDAVGGEALGR_2
ratio
time
0.000
0.005
0.010
0.015
0.00
0.01
0.02
0.03
0.04
0.05
0
5
10
15
20
25
outliers
beta
alpha

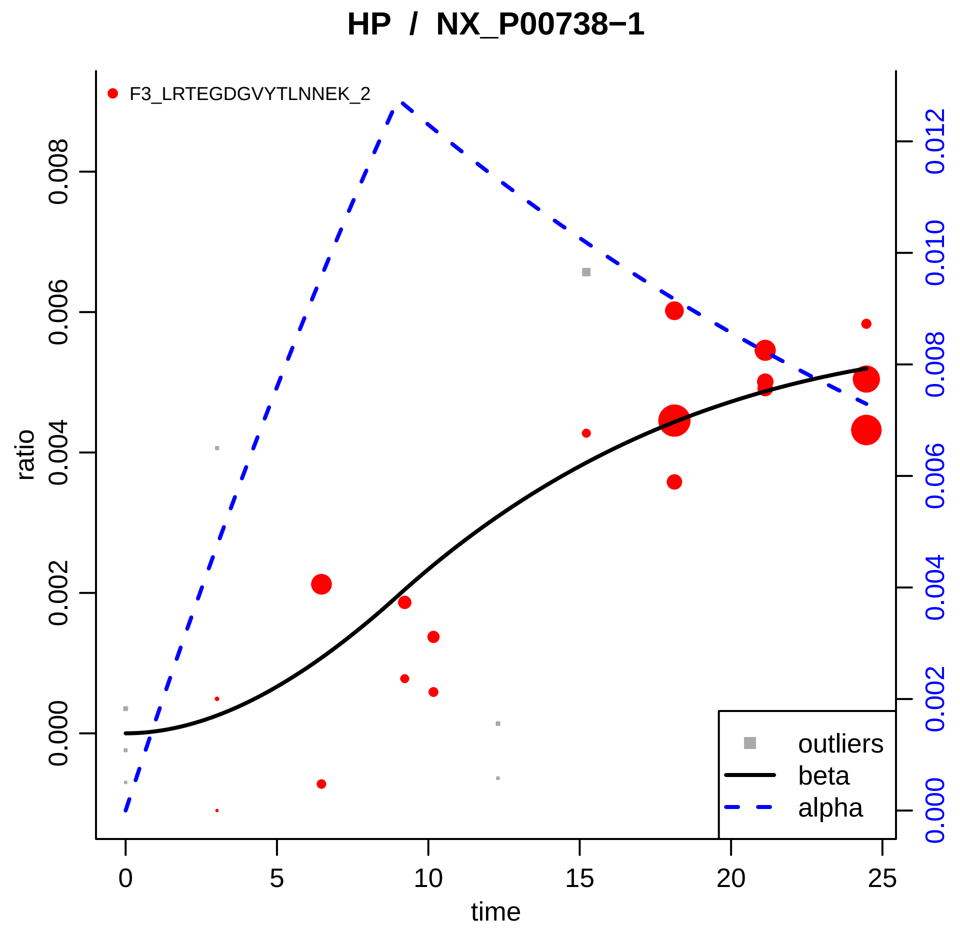

HP / NX_P00738−1
F3_LRTEGDGVYTLNNEK_2
ratio
0.000
0.002
0.004
0.006
0.008
time
0
5
10
15
20
25
0.000
0.002
0.004
0.006
0.008
0.010
0.012
outliers
beta
alpha

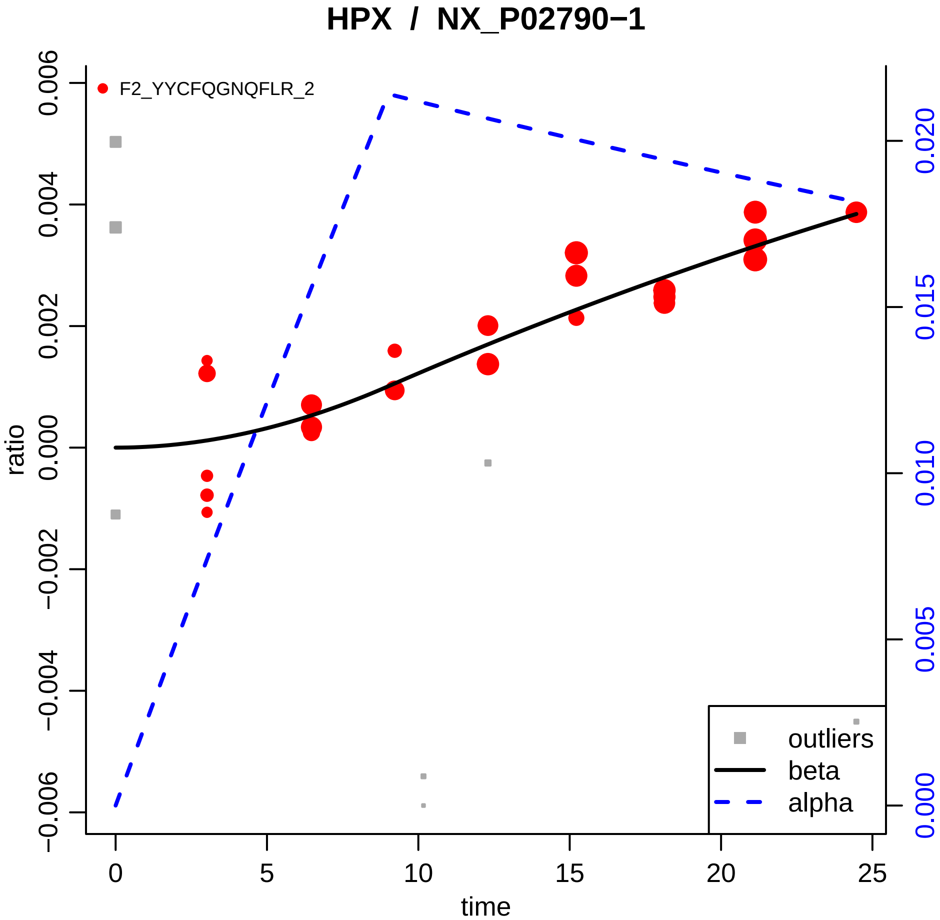

HPX / NX_P02790−1
F2_YYCFQGNQFLR_2
0.006
0.004
0.002
0.000
−0.002
−0.004
−0.006
ratio
0.020
0.015
0.010
0.005
0.000
0
5
10
15
20
25
time
outliers
beta
alpha

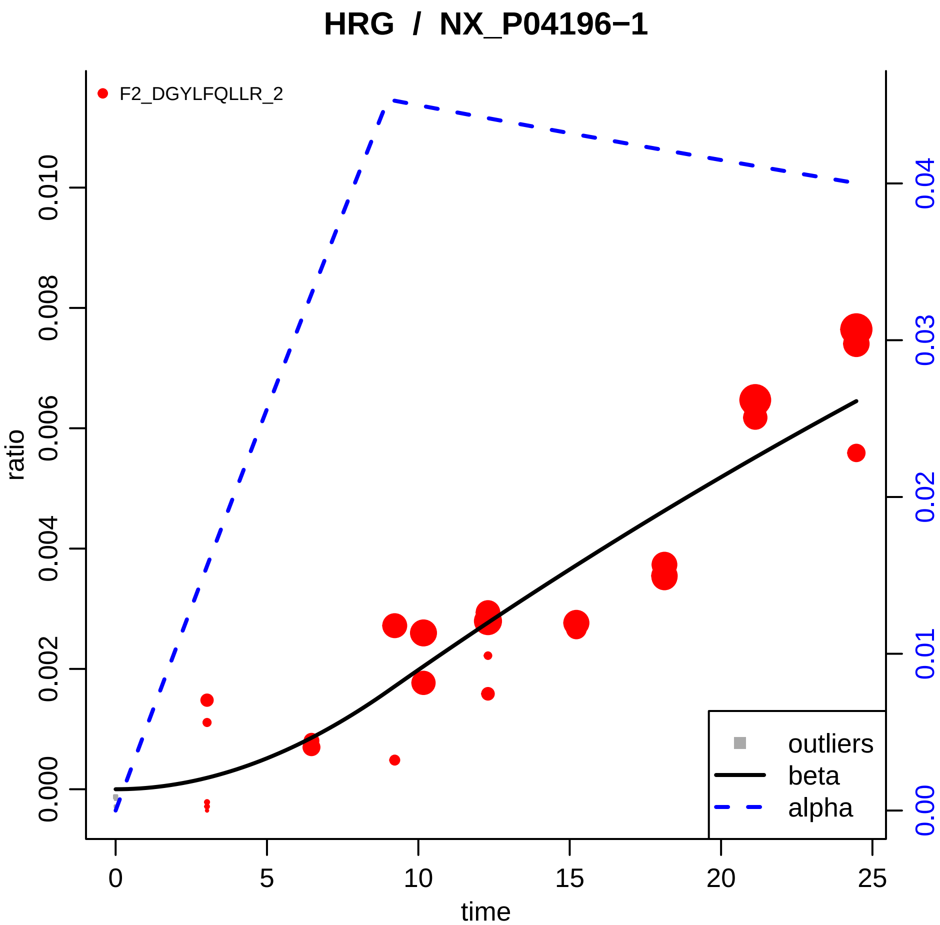

HRG / NX_P04196−1
F2_DGYLFQLLR_2
ratio
0.000
0.002
0.004
0.006
0.008
0.010
0.00
0.01
0.02
0.03
0.04
time
0
5
10
15
20
25
outliers
beta
alpha

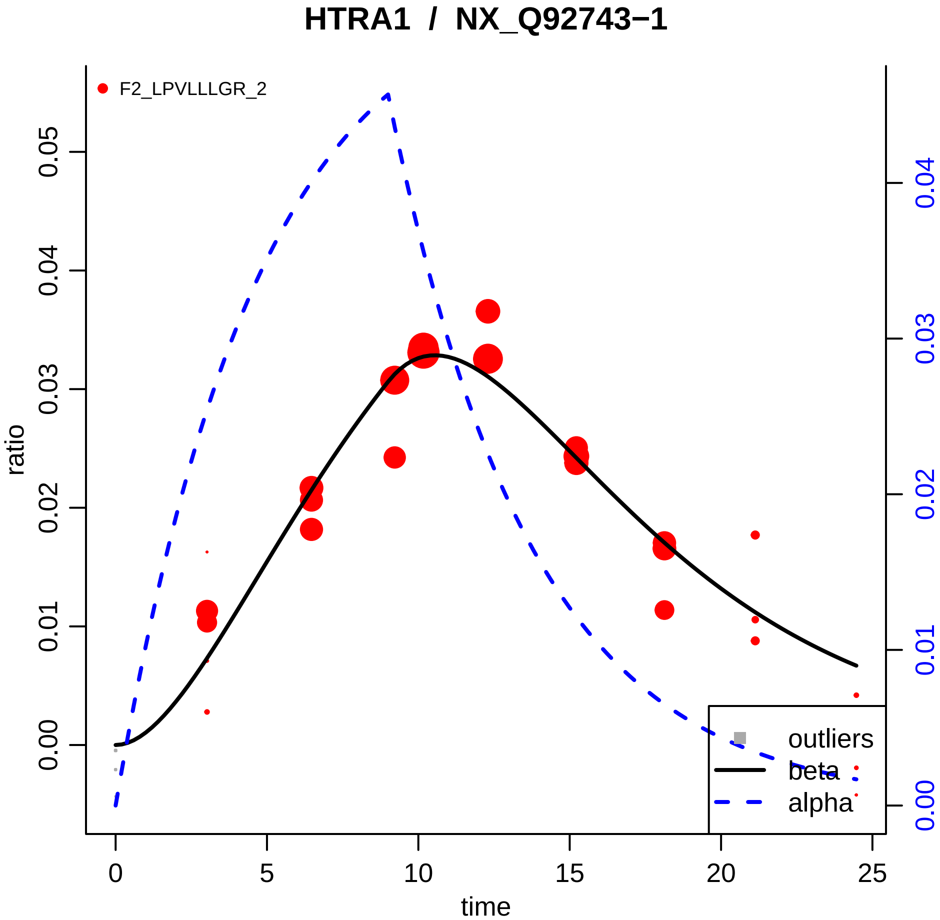
HTRA1 / NX_Q92743−1
F2_LPVLLLGR_2
ratio
time
0.00
0.01
0.02
0.03
0.04
0.05
0
5
10
15
20
25
0.00
0.01
0.02
0.03
0.04
outliers
beta
alpha

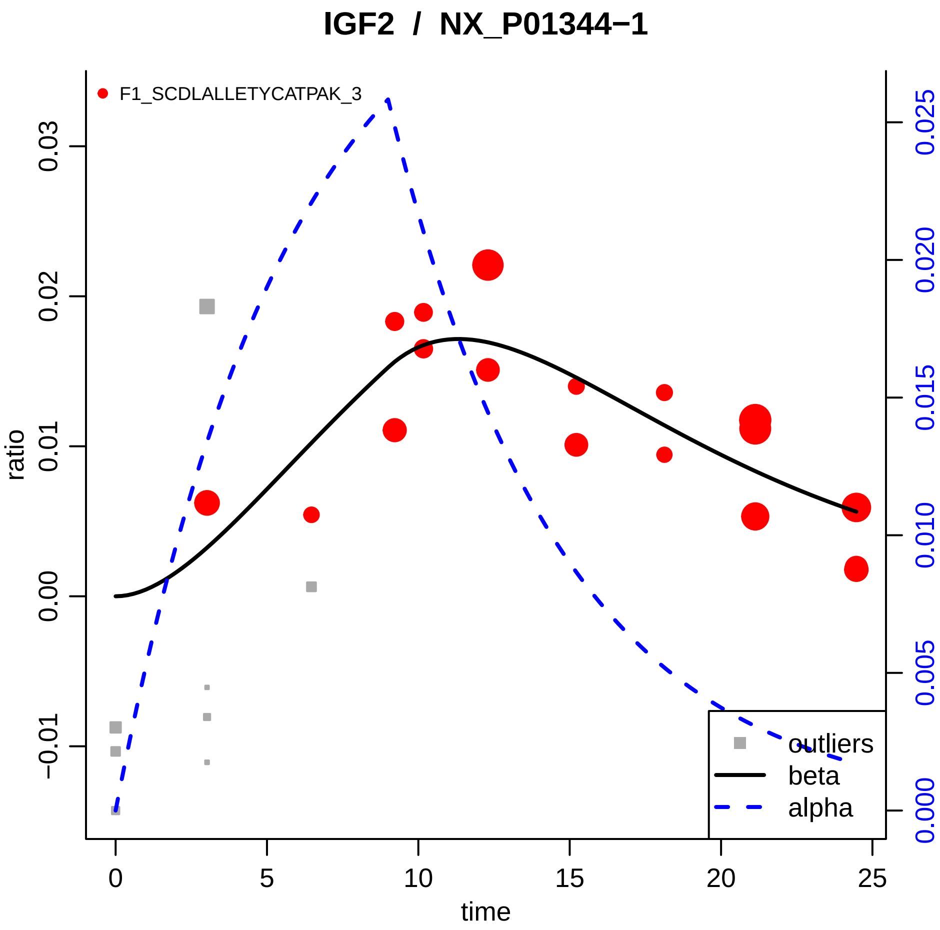

IGF2 / NX_P01344−1
F1_SCDLALLETYCATPAK_3
ratio
time
0.03
0.02
0.01
0.00
−0.01
0
5
10
15
20
25
0.025
0.020
0.015
0.010
0.005
0.000
outliers
beta
alpha

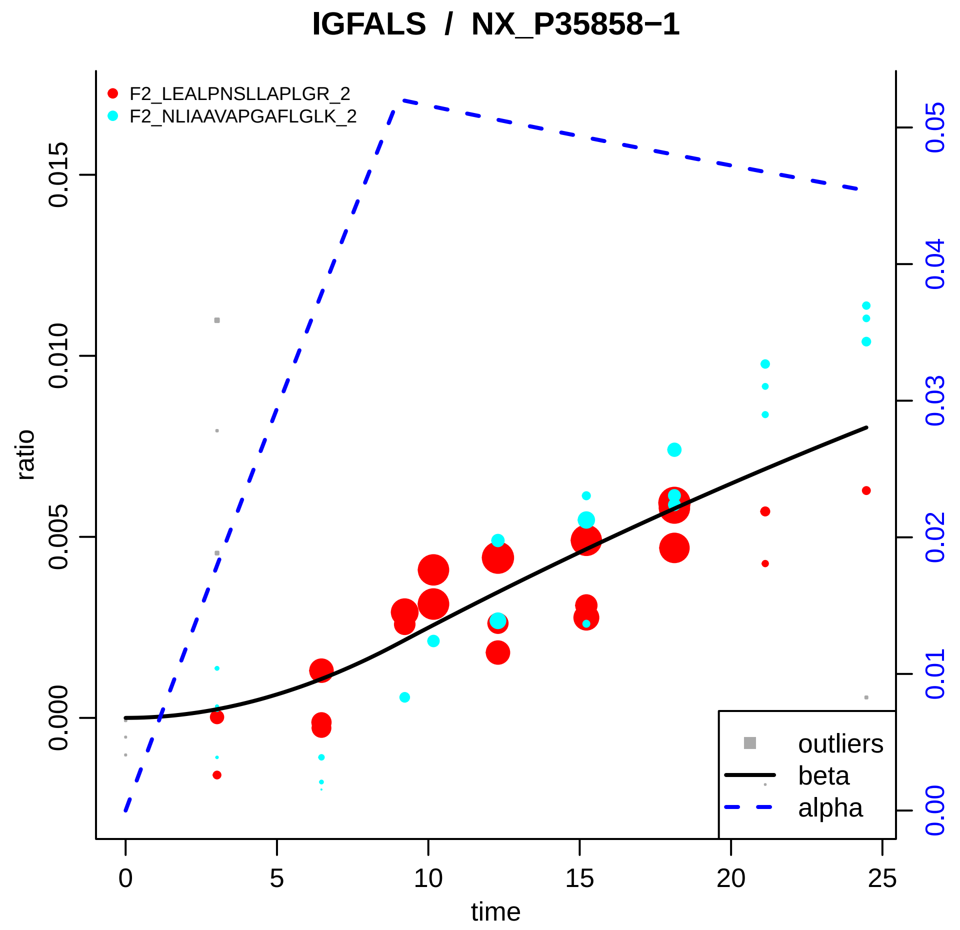

IGFALS / NX_P35858−1
F2_LEALPNSLLAPLGR_2
F2_NLIAAVAPGAFLGLK_2
ratio
0.000
0.005
0.010
0.015
0.00
0.01
0.02
0.03
0.04
0.05
0
5
10
15
20
25
time
outliers
beta
alpha

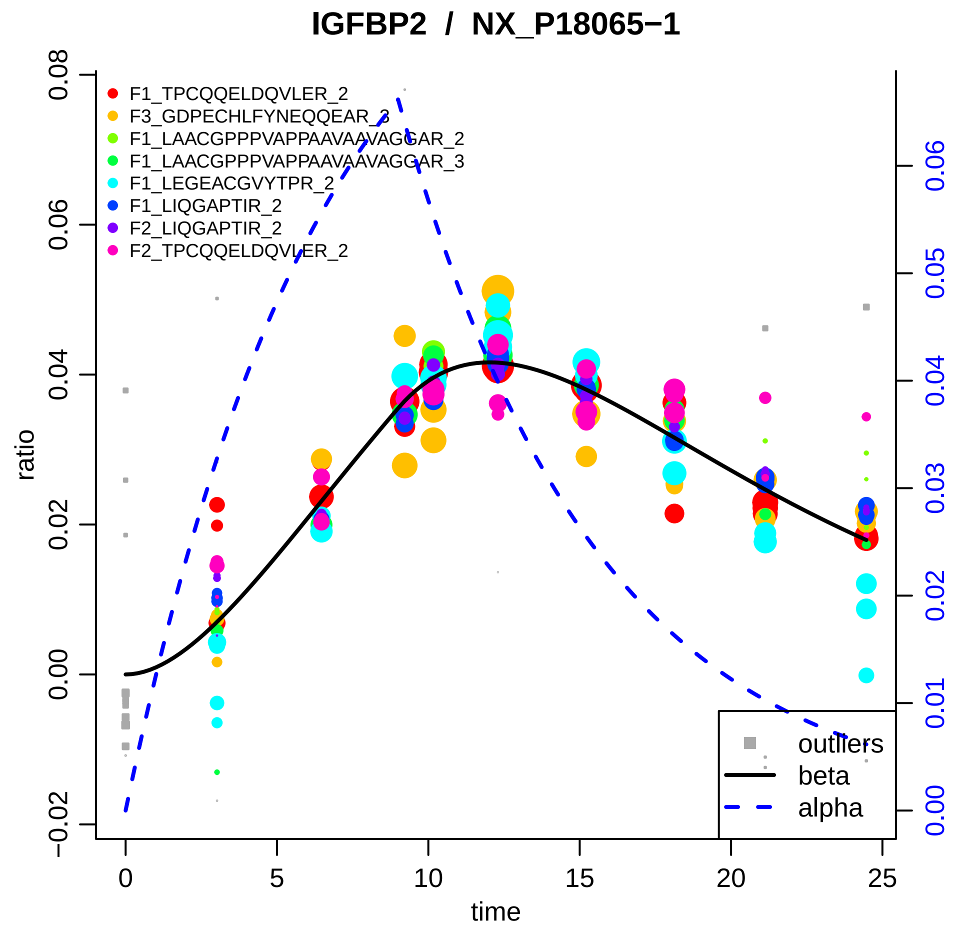

IGFBP2 / NX_P18065−1
F1_TPCQQELDQVLER_2
F3_GDPECHLFYNEQQEAR_3
F1_LAACGPPPVAPPAAVAAVAGGAR_2
F1_LAACGPPPVAPPAAVAAVAGGAR_3
F1_LEGEACGVYTPR_2
F1_LIQGAPTIR_2
F2_LIQGAPTIR_2
F2_TPCQQELDQVLER_2
outliers
beta
alpha
ratio
time
−0.02
0.00
0.02
0.04
0.06
0.08
0
5
10
15
20
25
0.00
0.01
0.02
0.03
0.04
0.05
0.06

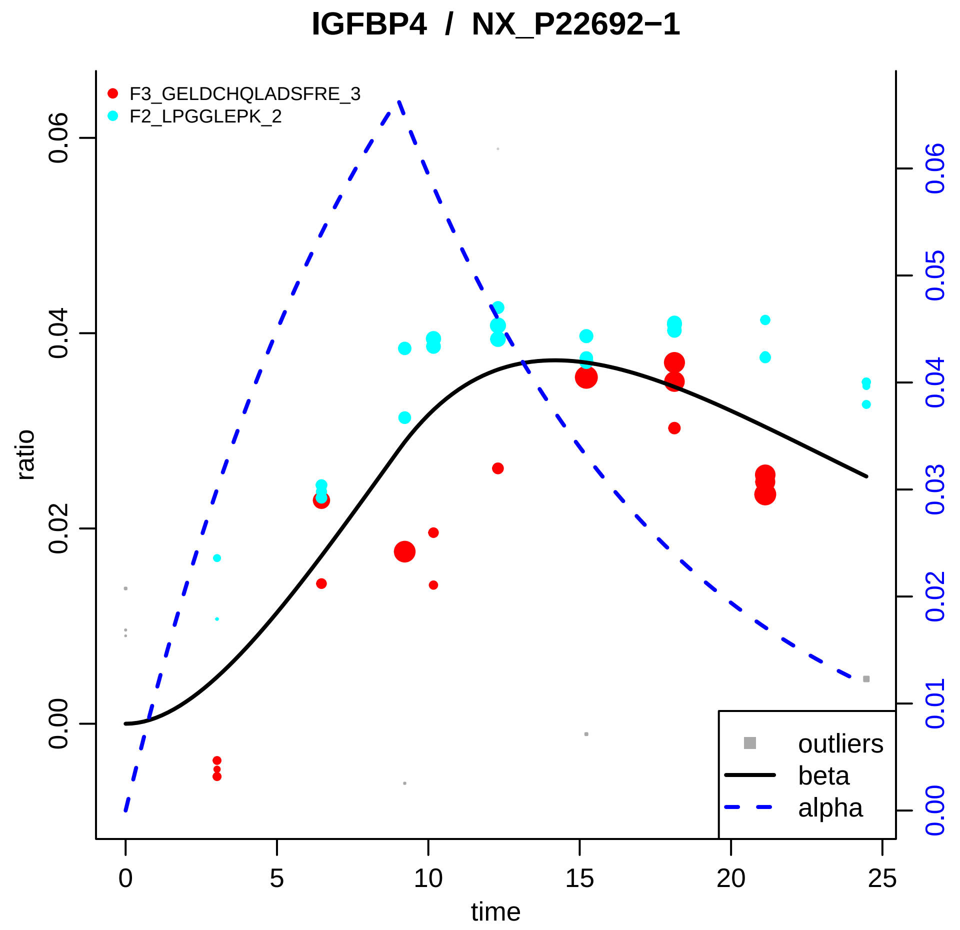

IGFBP4 / NX_P22692−1
F3_GELDCHQLADSFRE_3
F2_LPGGLEPK_2
ratio
0.00
0.02
0.04
0.06
time
0
5
10
15
20
25
0.00
0.01
0.02
0.03
0.04
0.05
0.06
outliers
beta
alpha

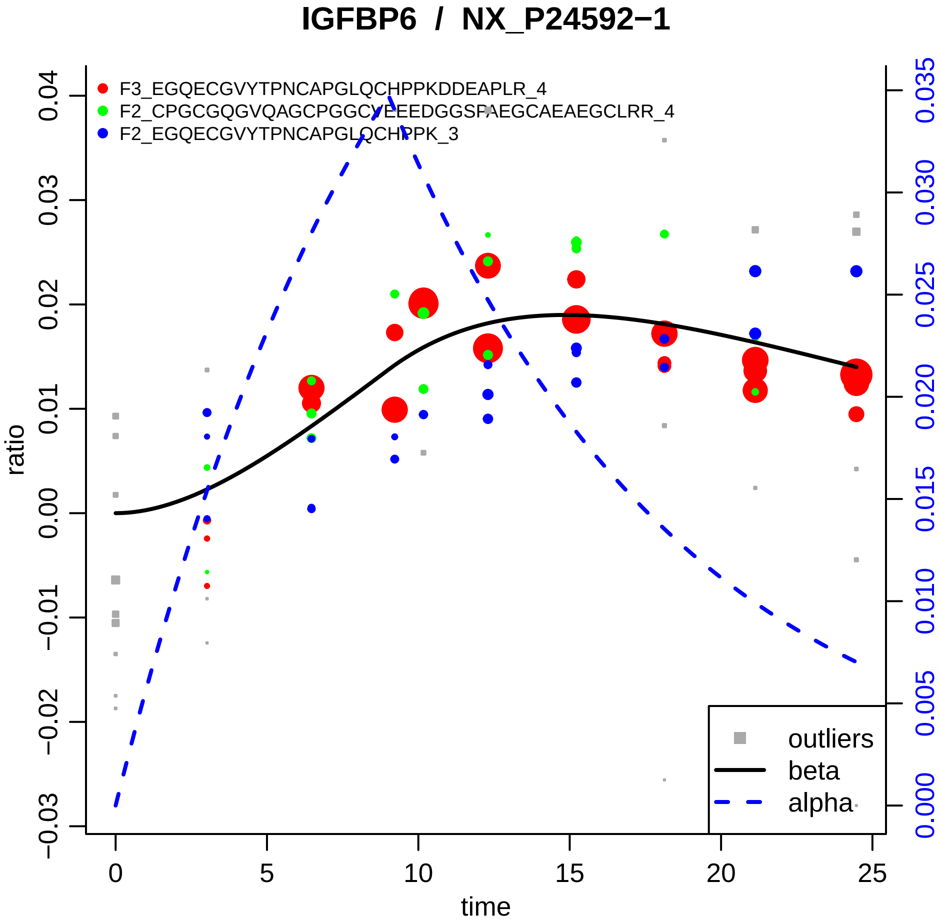
IGFBP6 / NX_P24592−1
F3_EGQECGVYTPNCAPGLQCHPPKDDEAPLR_4
F2_CPGCGQGVQAGCPGGCYEEEDGGSPAEGCAEAEGCLRR_4
F2_EGQECGVYTPNCAPGLQCHPPK_3
ratio
time
outliers
beta
alpha
0
5
10
15
20
25
−0.03
−0.02
−0.01
0.00
0.01
0.02
0.03
0.04
0.000
0.005
0.010
0.015
0.020
0.025
0.030
0.035

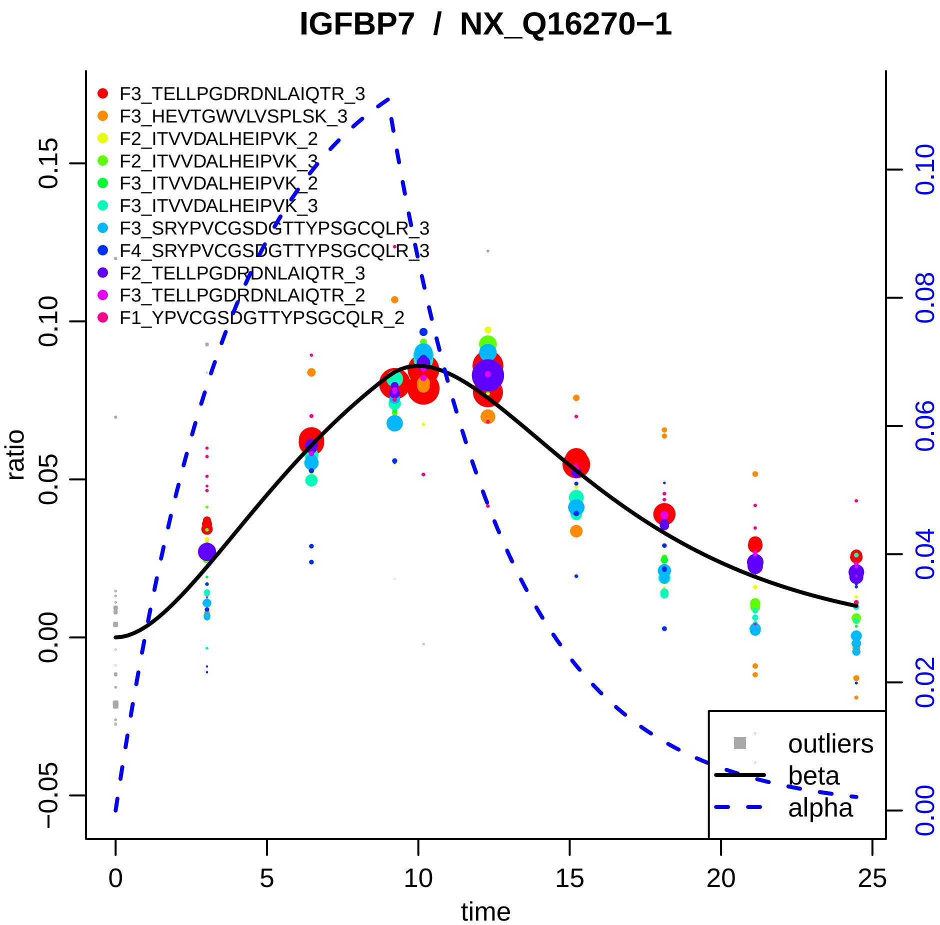

IGFBP7 / NX_Q16270−1
F3_TELLPGDRDNLAIQTR_3
F3_HEVTGWVLVSPLSK_3
F2_ITVVDALHEIPVK_2
F2_ITVVDALHEIPVK_3
F3_ITVVDALHEIPVK_2
F3_ITVVDALHEIPVK_3
F3_SRYPVCGSDGTTYPSGCQLR_3
F4_SRYPVCGSDGTTYPSGCQLR_3
F2_TELLPGDRDNLAIQTR_3
F3_TELLPGDRDNLAIQTR_2
F1_YPVCGSDGTTYPSGCQLR_2
outliers
beta
alpha
ratio
time
0.15
0.10
0.05
0.00
−0.05
0
5
10
15
20
25
0.10
0.08
0.06
0.04
0.02
0.00

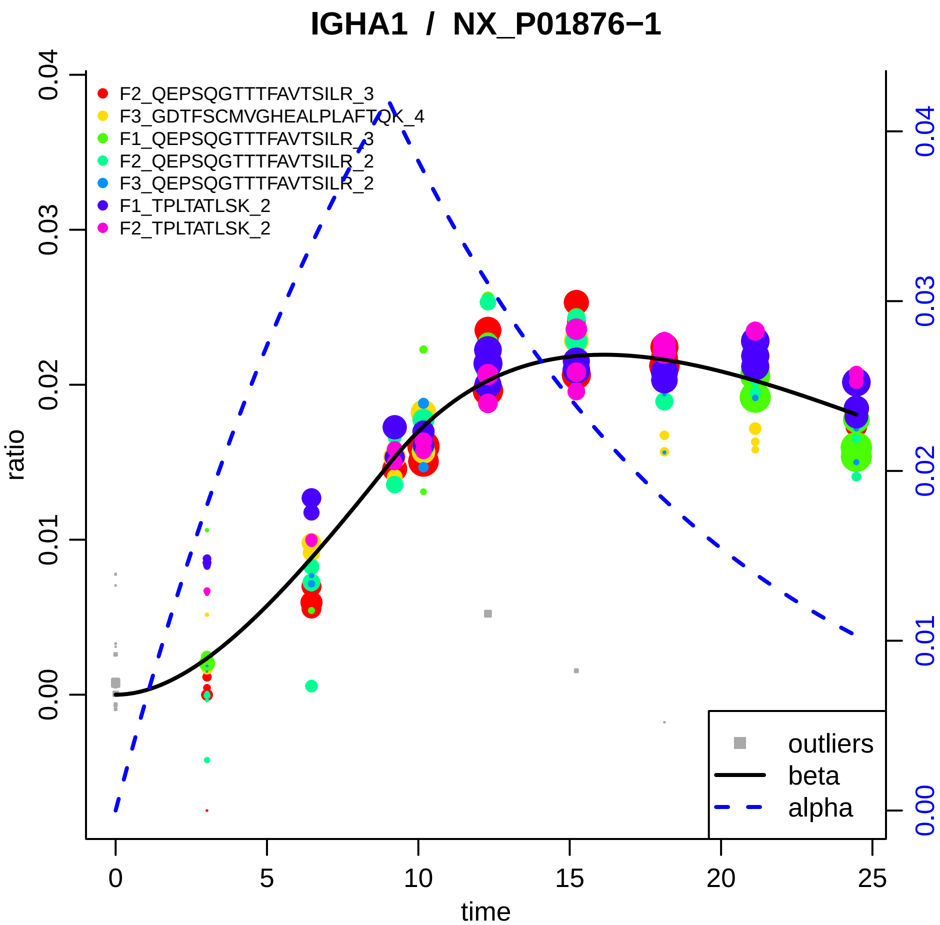
IGHA1 / NX_P01876−1
F2_QEPSQGTTTFAVTSILR_3
F3_GDTFSCMVGHEALPLAFTQK_4
F1_QEPSQGTTTFAVTSILR_3
F2_QEPSQGTTTFAVTSILR_2
F3_QEPSQGTTTFAVTSILR_2
F1_TPLTATLSK_2
F2_TPLTATLSK_2
outliers
beta
alpha
ratio
time
0.00
0.01
0.02
0.03
0.04
0
5
10
15
20
25

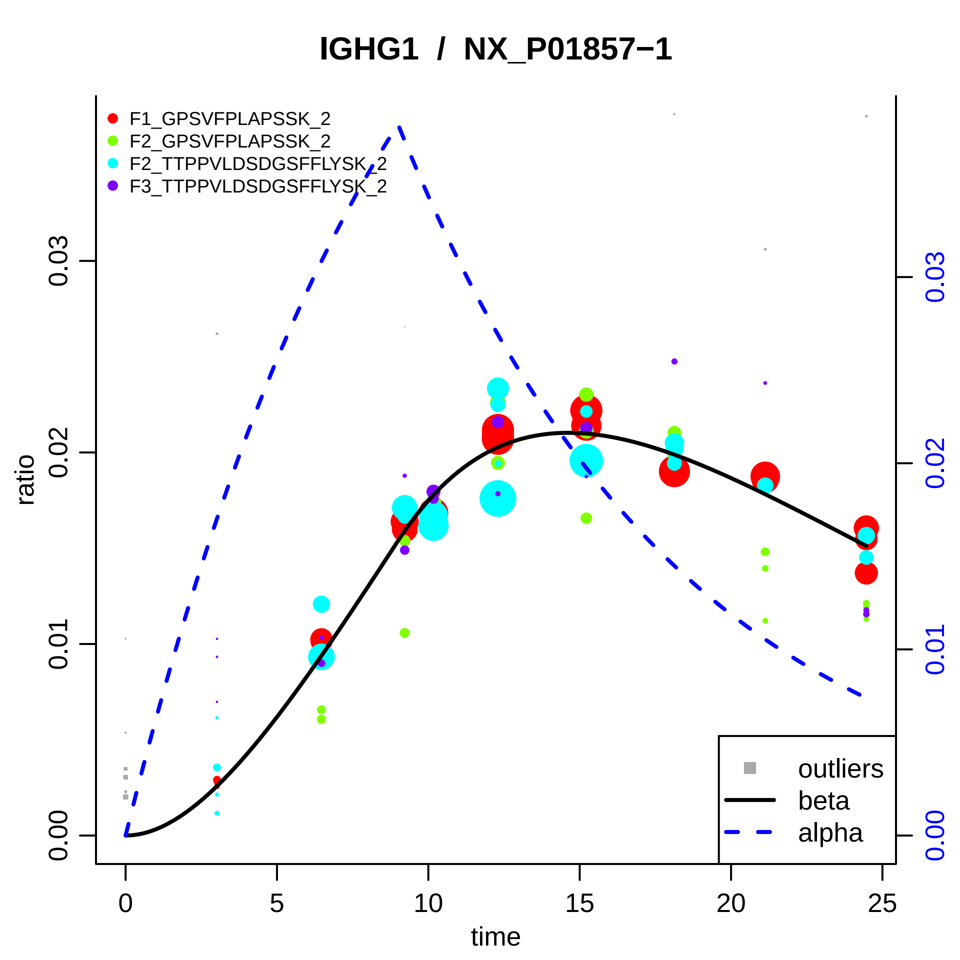

IGHG1 / NX_P01857−1
F1_GPSVFPLAPSSK_2
F2_GPSVFPLAPSSK_2
F2_TTPPVLDSDGSFFLYSK_2
F3_TTPPVLDSDGSFFLYSK_2
outliers
beta
alpha
ratio
time
0.00
0.01
0.02
0.03
0
5
10
15
20
25

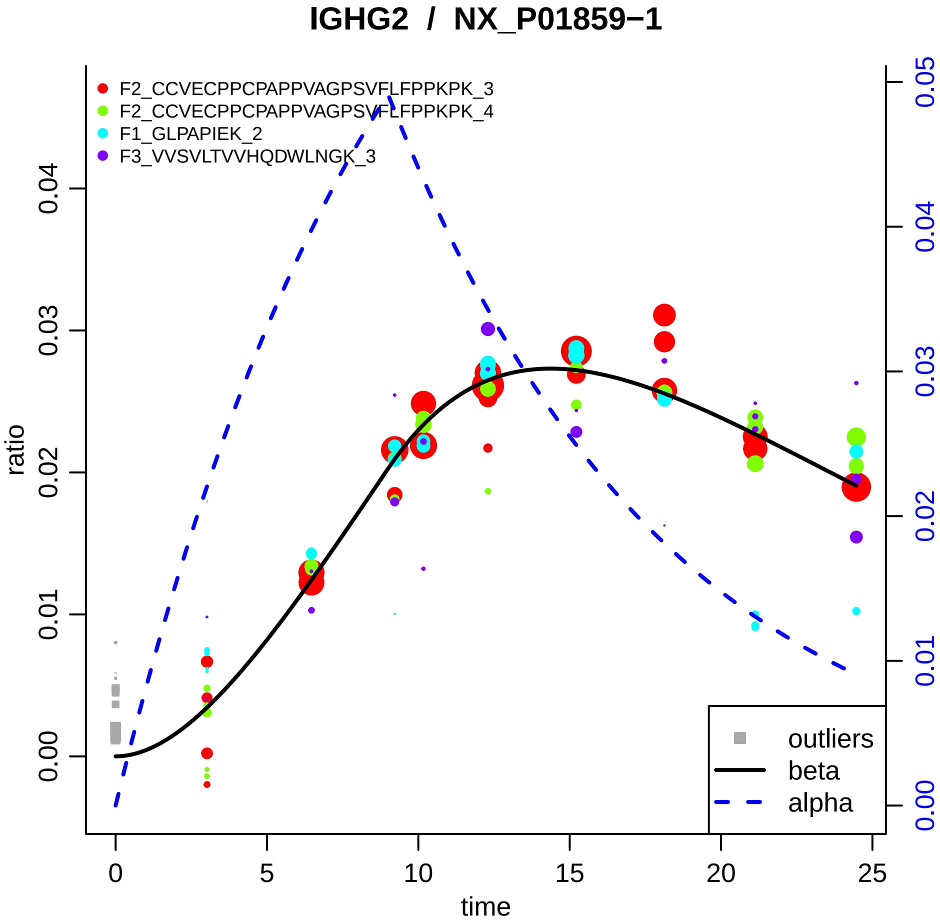

IGHG2 / NX_P01859−1
F2_CCVECPPCPAPPVAGPSVFLFPPKPK_3
F2_CCVECPPCPAPPVAGPSVFLFPPKPK_4
F1_GLPAPIEK_2
F3_VVSVLTVVHQDWLNGK_3
ratio
0.00
0.01
0.02
0.03
0.04
0.00
0.01
0.02
0.03
0.04
0.05
0
5
10
15
20
25
time
outliers
beta
alpha

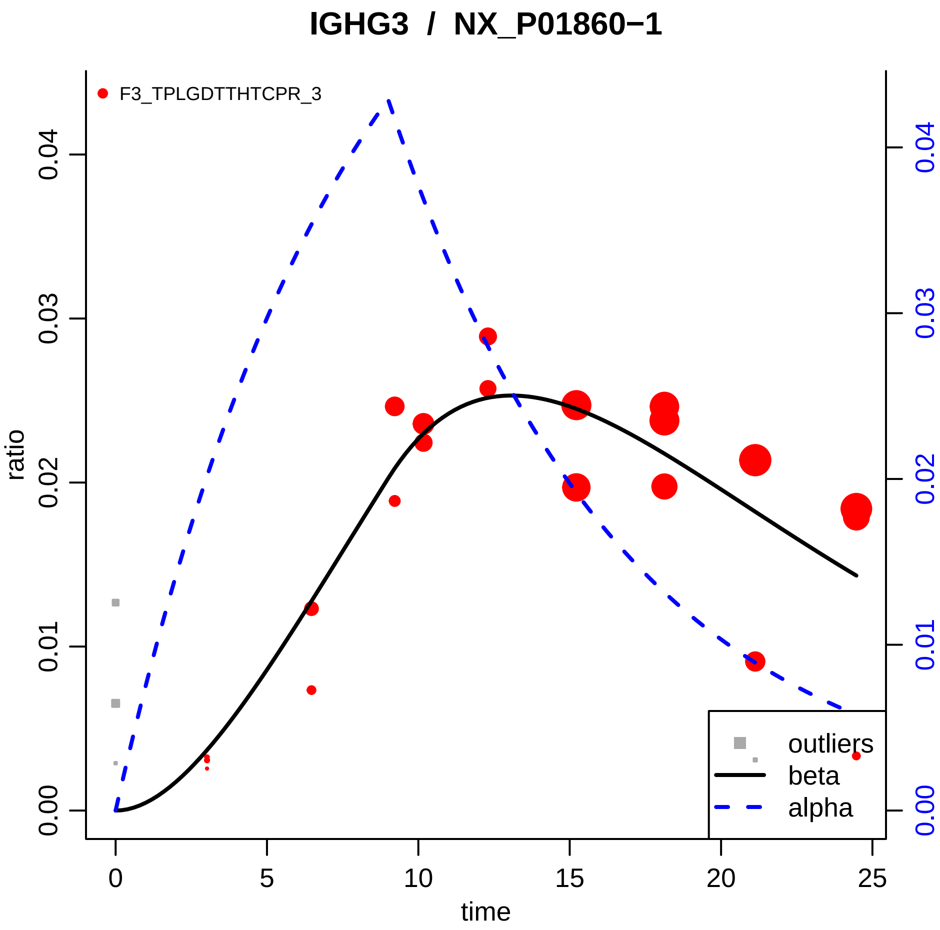

IGHG3 / NX_P01860−1
F3_TPLGDTTHTCPR_3
ratio
time
0.00
0.01
0.02
0.03
0.04
0
5
10
15
20
25
outliers
beta
alpha

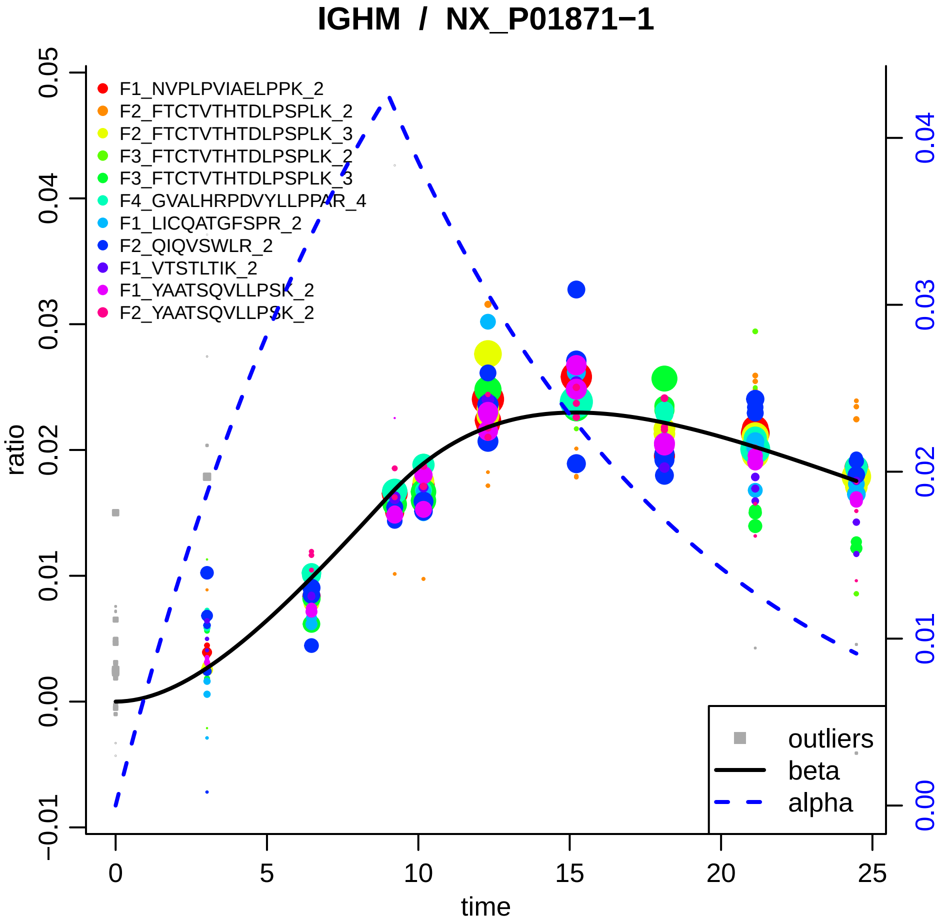
IGHM / NX_P01871−1
F1_NVPLPVIAELPPK_2
F2_FTCTVTHTDLPSPLK_2
F2_FTCTVTHTDLPSPLK_3
F3_FTCTVTHTDLPSPLK_2
F3_FTCTVTHTDLPSPLK_3
F4_GVALHRPDVYLLPPAR_4
F1_LICQATGFSPR_2
F2_QIQVSWLR_2
F1_VTSTLTIK_2
F1_YAATSQVLLPSK_2
F2_YAATSQVLLPSK_2
outliers
beta
alpha
ratio
time
−0.01
0.00
0.01
0.02
0.03
0.04
0.05
0
5
10
15
20
25

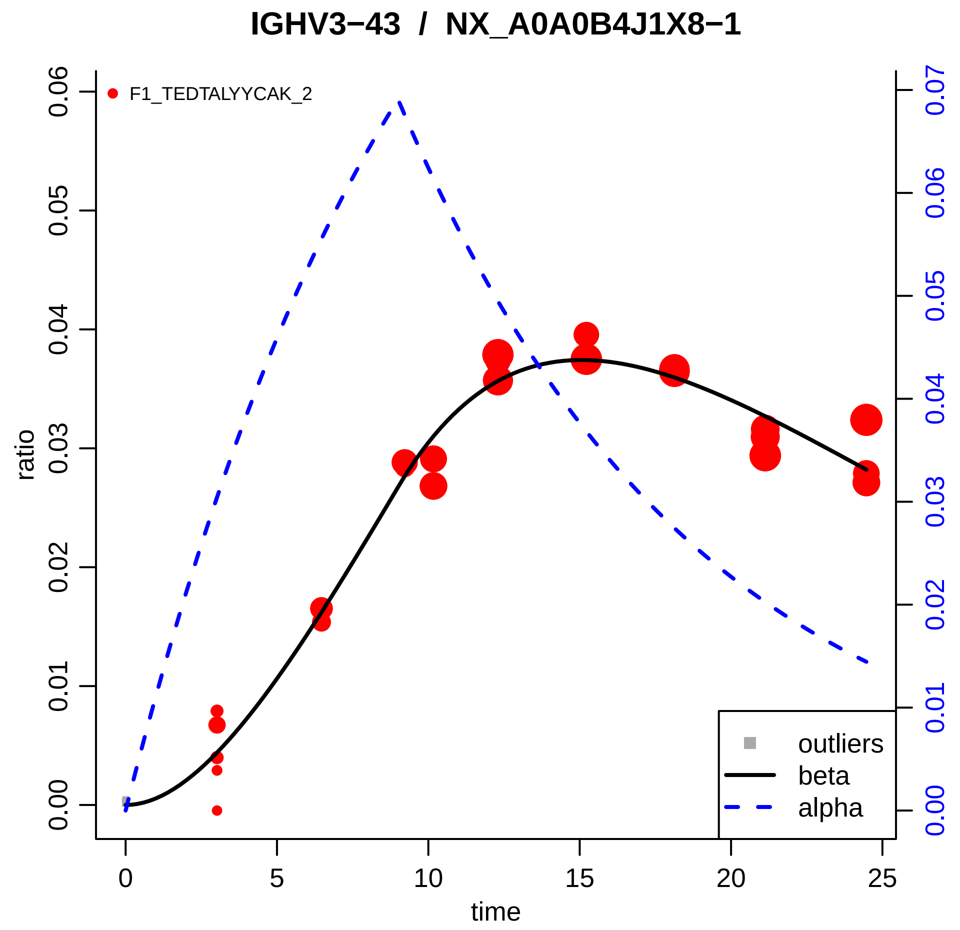

IGHV3−43 / NX_A0A0B4J1X8−1
F1_TEDTALYYCAK_2
ratio
time
0.00
0.01
0.02
0.03
0.04
0.05
0.06
0.07
0
5
10
15
20
25
outliers
beta
alpha

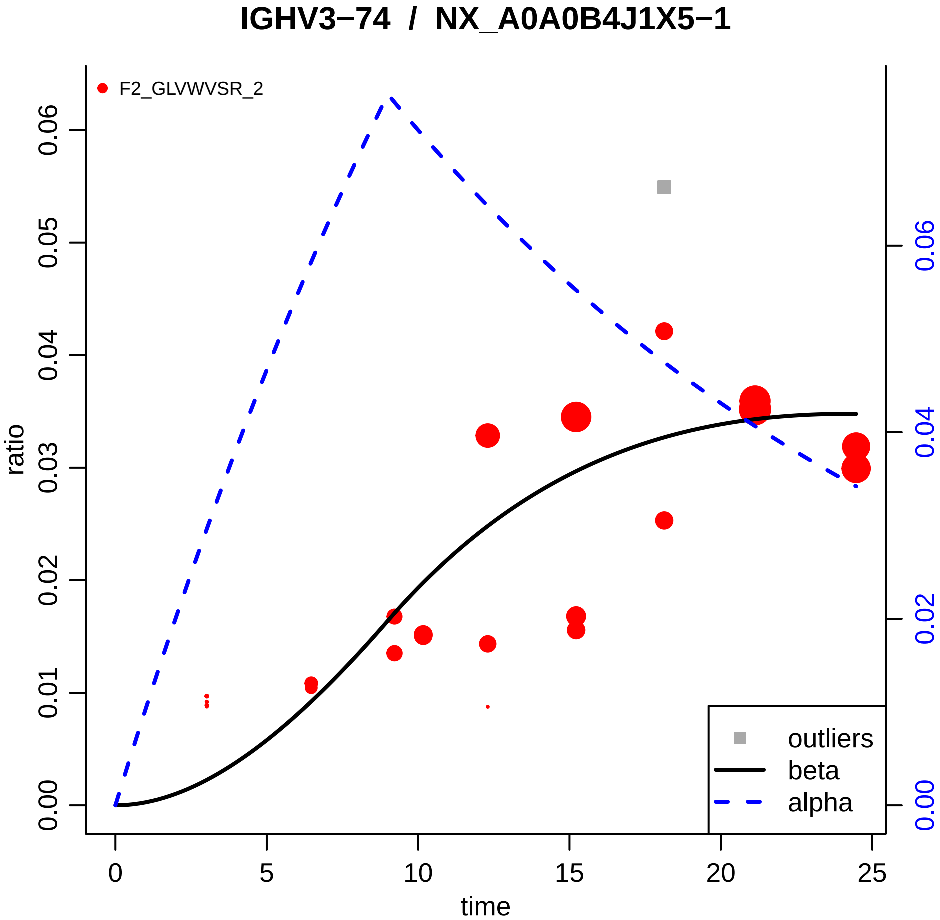

IGHV3−74 / NX_A0A0B4J1X5−1
F2_GLVWVSR_2
ratio
time
0.00
0.01
0.02
0.03
0.04
0.05
0.06
0
5
10
15
20
25
0.00
0.02
0.04
0.06
outliers
beta
alpha

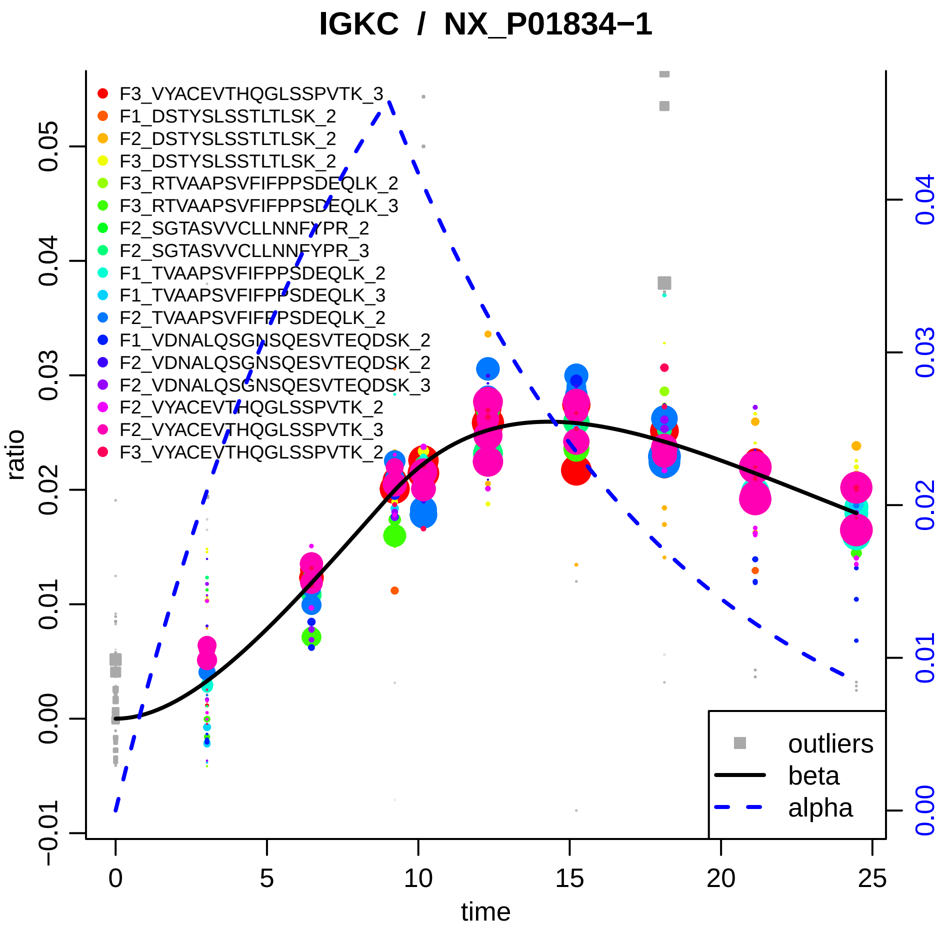

IGKC / NX_P01834−1
F3_VYACEVTHQGLSSPVTK_3
F1_DSTYSLSSTLTLSK_2
F2_DSTYSLSSTLTLSK_2
F3_DSTYSLSSTLTLSK_2
F3_RTVAAPSVFIFPPSDEQLK_2
F3_RTVAAPSVFIFPPSDEQLK_3
F2_SGTASVVCLLNNFYPR_2
F2_SGTASVVCLLNNFYPR_3
F1_TVAAPSVFIFPPSDEQLK_2
F1_TVAAPSVFIFPPSDEQLK_3
F2_TVAAPSVFIFPPSDEQLK_2
F1_VDNALQSGNSQESVTEQDSK_2
F2_VDNALQSGNSQESVTEQDSK_2
F2_VDNALQSGNSQESVTEQDSK_3
F2_VYACEVTHQGLSSPVTK_2
F2_VYACEVTHQGLSSPVTK_3
F3_VYACEVTHQGLSSPVTK_2
outliers
beta
alpha
ratio
time
0.05
0.04
0.03
0.02
0.01
0.00
−0.01
0
5
10
15
20
25
0.04
0.03
0.02
0.01
0.00

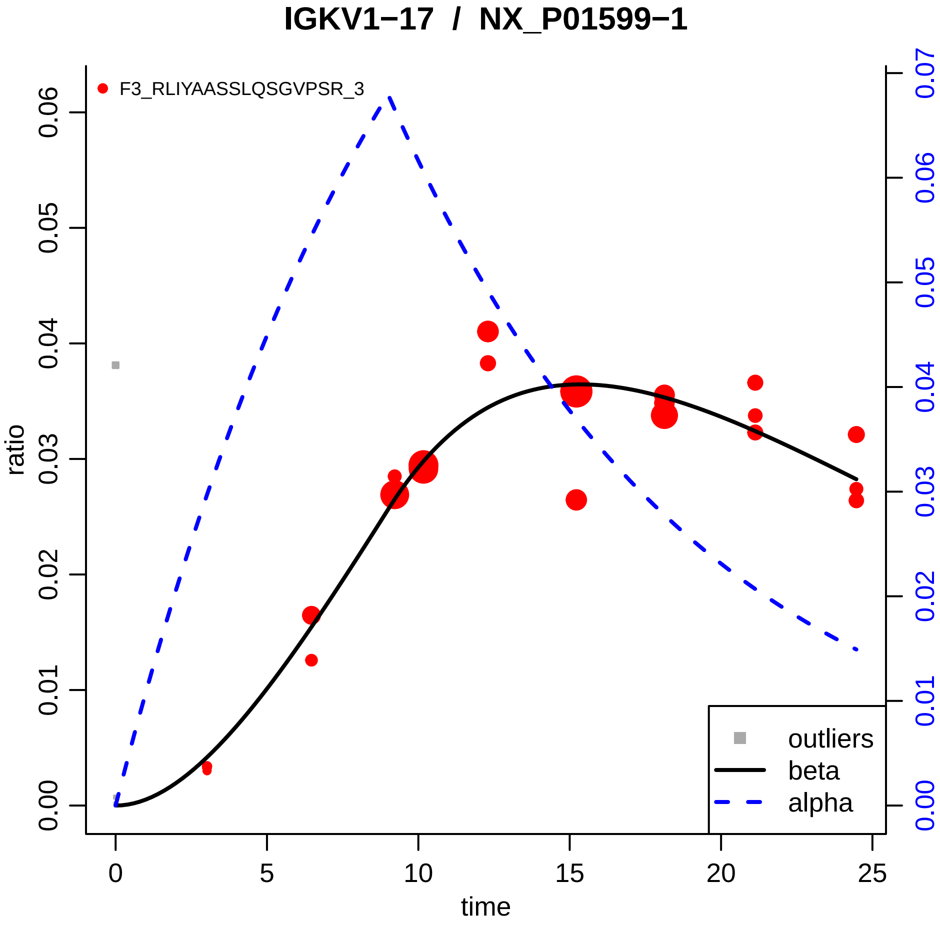

IGKV1−17 / NX_P01599−1
F3_RLIYAASSLQSGVPSR_3
ratio
time
0.00
0.01
0.02
0.03
0.04
0.05
0.06
0.07
0
5
10
15
20
25
outliers
beta
alpha

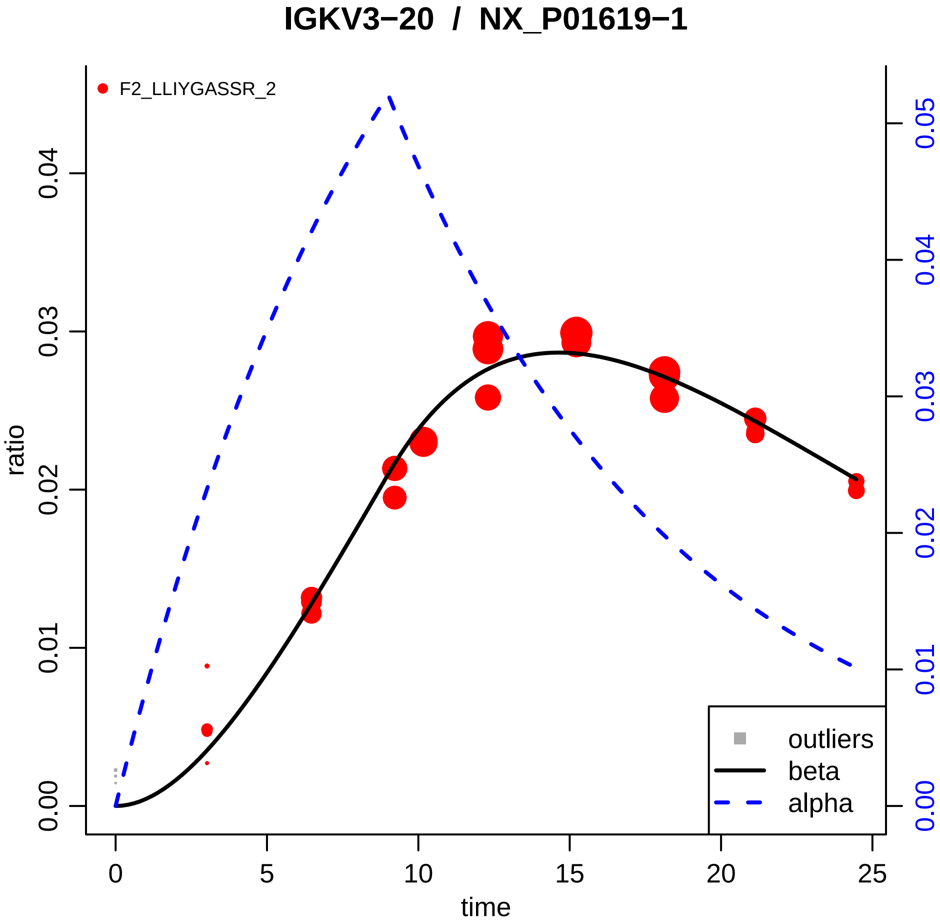

IGKV3−20 / NX_P01619−1
F2_LLIYGASSR_2
ratio
0.00
0.01
0.02
0.03
0.04
0.05
time
0
5
10
15
20
25
outliers
beta
alpha

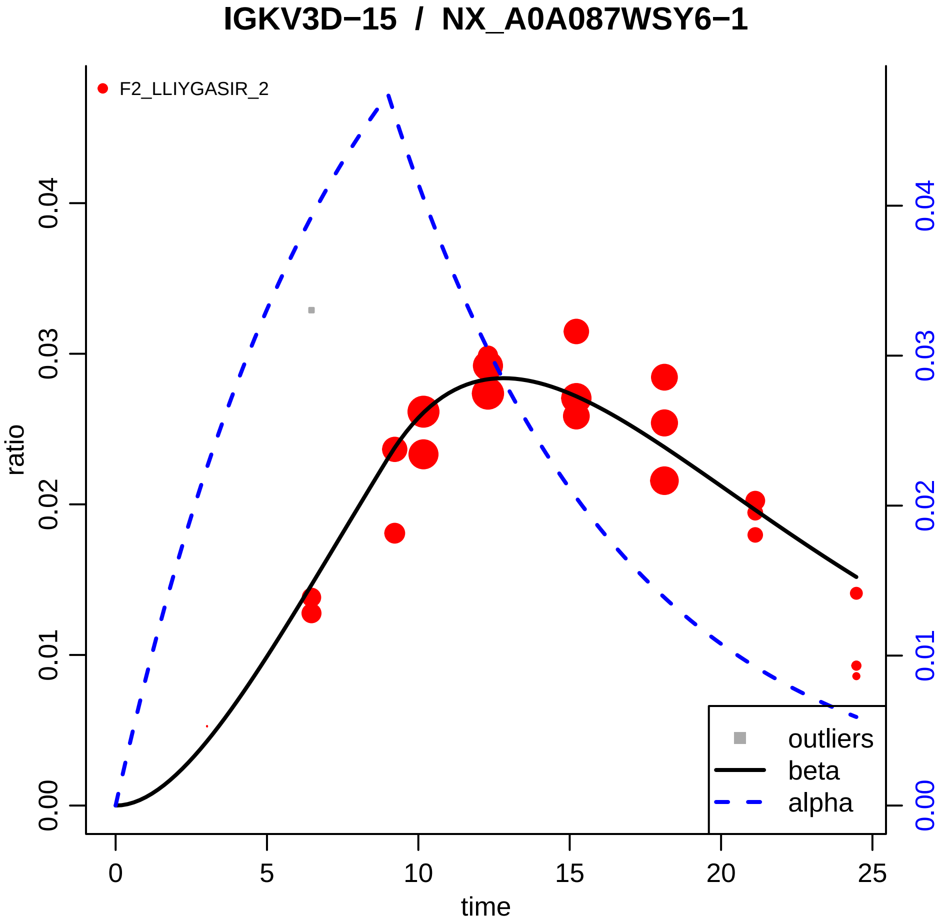

IGKV3D−15 / NX_A0A087WSY6−1
F2_LLIYGASIR_2
ratio
time
0.00
0.01
0.02
0.03
0.04
0
5
10
15
20
25
outliers
beta
alpha

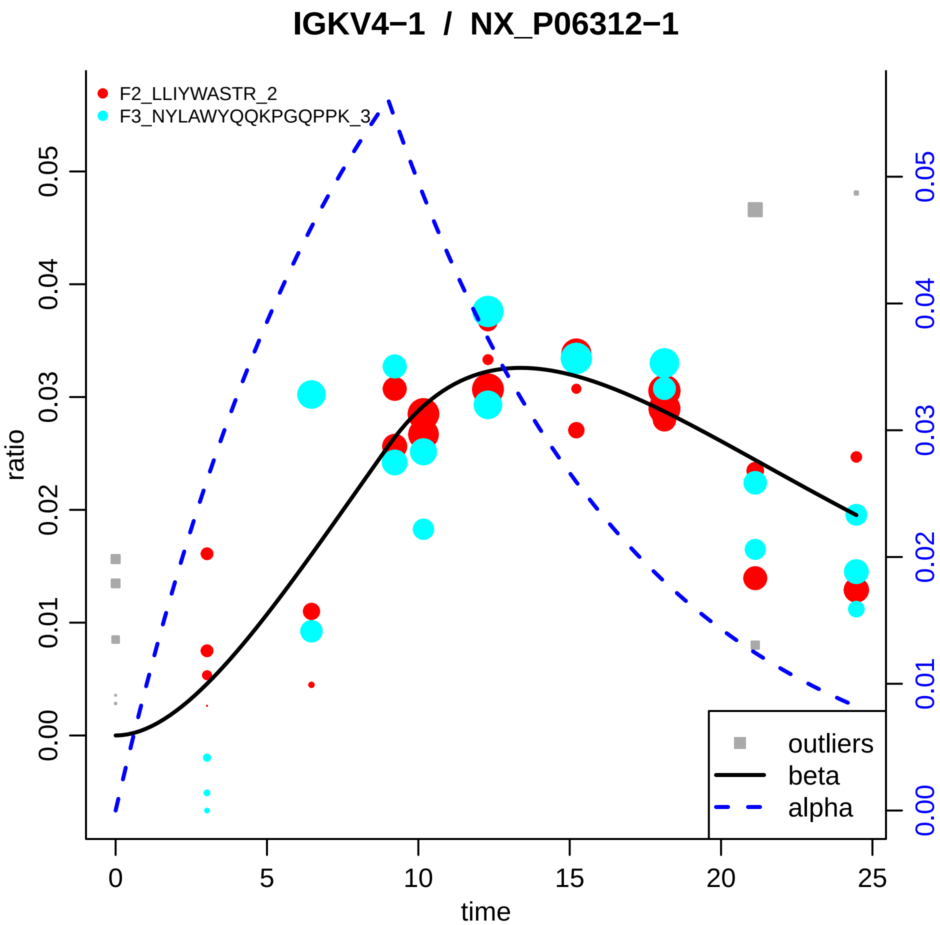

IGKV4−1 / NX_P06312−1
F2_LLIYWASTR_2
F3_NYLAWYQQKPGQPPK_3
ratio
0.00
0.01
0.02
0.03
0.04
0.05
time
0
5
10
15
20
25
outliers
beta
alpha

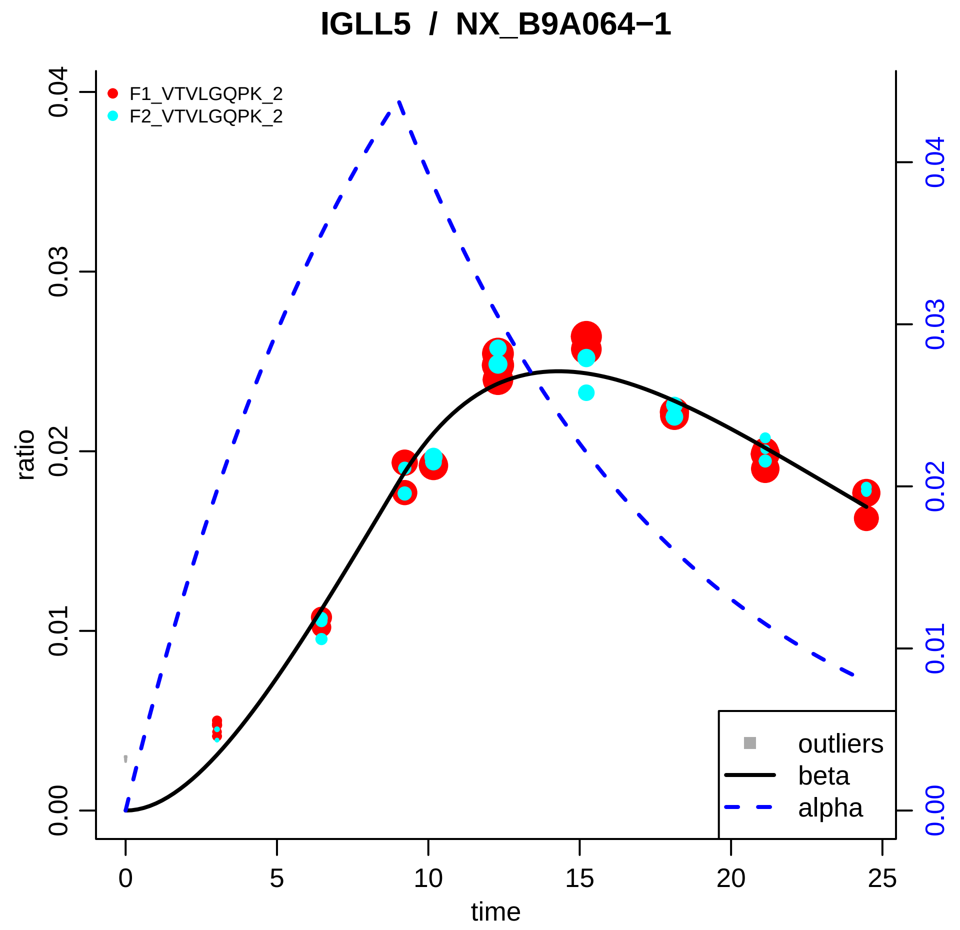

IGLL5 / NX_B9A064−1
F1_VTVLGQPK_2
F2_VTVLGQPK_2
ratio
0.00
0.01
0.02
0.03
0.04
time
0
5
10
15
20
25
outliers
beta
alpha

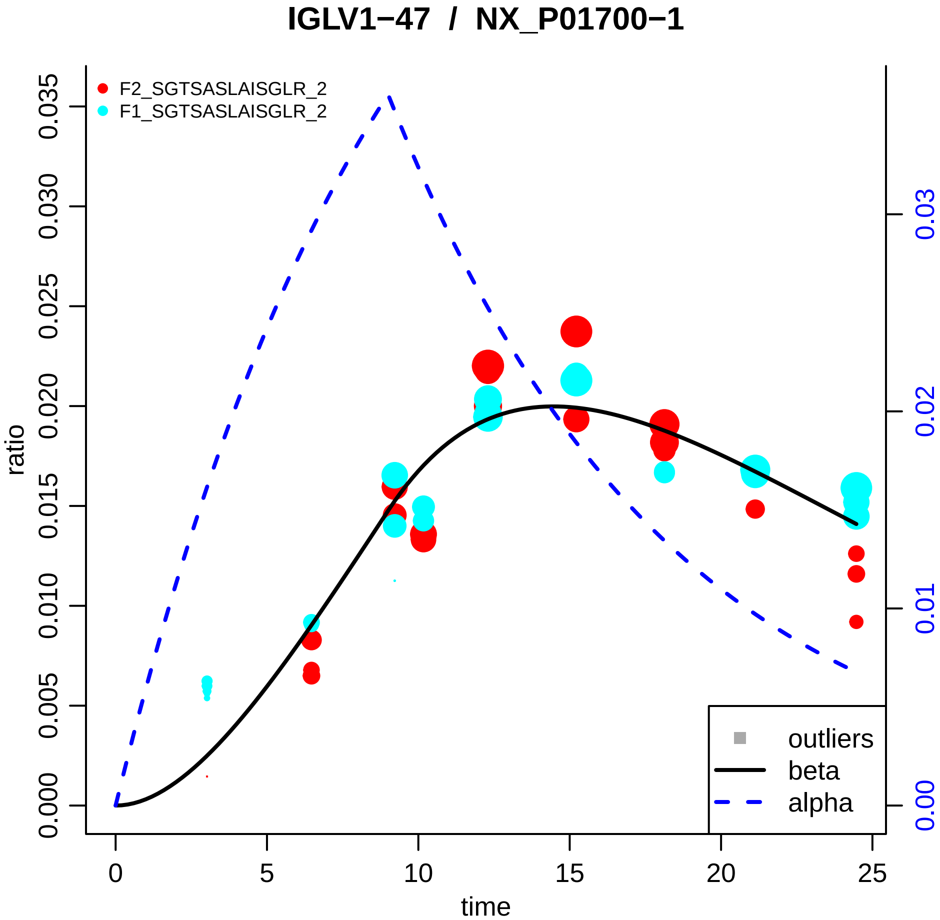

IGLV1−47 / NX_P01700−1
F2_SGTSASLAISGLR_2
F1_SGTSASLAISGLR_2
ratio
time
outliers
beta
alpha
0.000
0.005
0.010
0.015
0.020
0.025
0.030
0.035
0.00
0.01
0.02
0.03
0
5
10
15
20
25

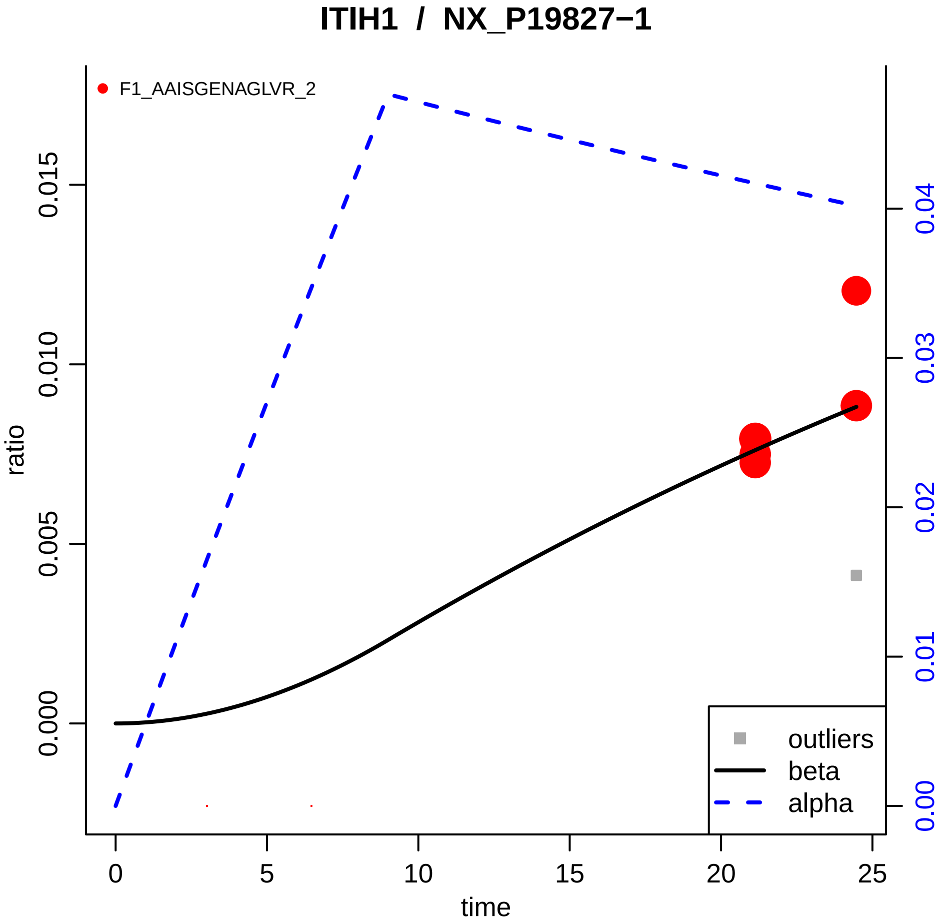

ITIH1 / NX_P19827−1
F1_AAISGENAGLVR_2
ratio
0.000
0.005
0.010
0.015
0.00
0.01
0.02
0.03
0.04
time
0
5
10
15
20
25
outliers
beta
alpha

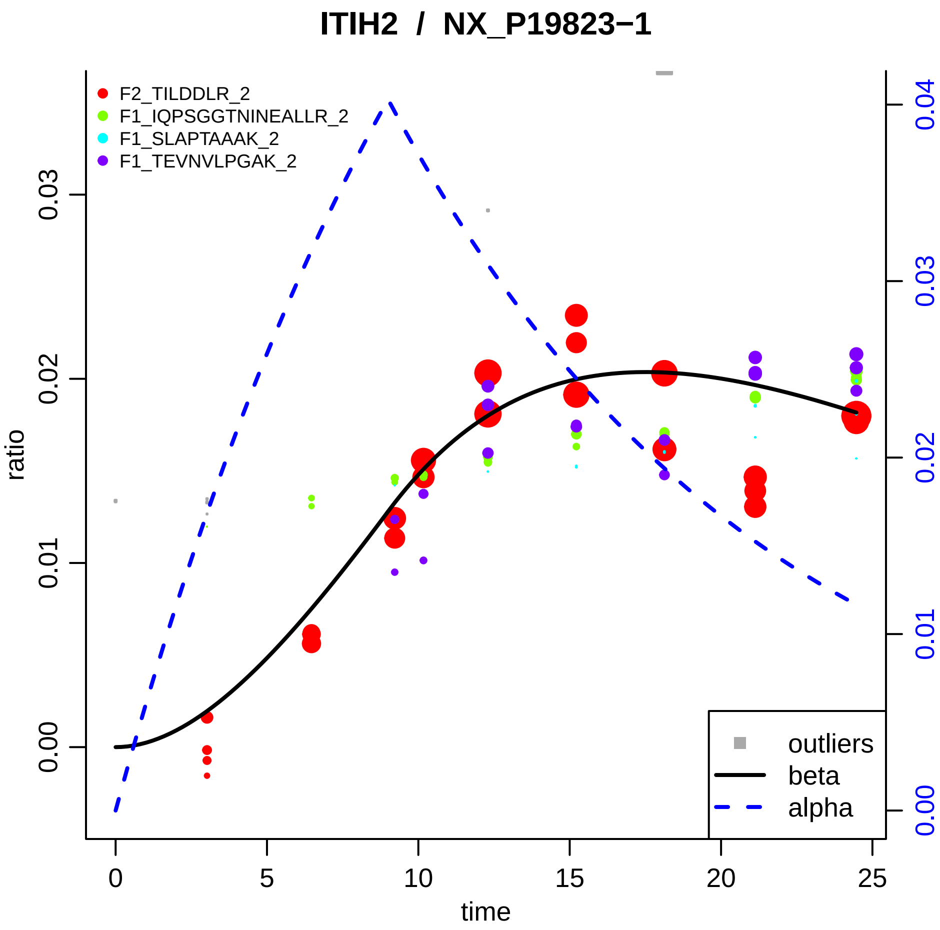

ITIH2 / NX_P19823−1
F2_TILDDLR_2
F1_IQPSGGTNINEALLR_2
F1_SLAPTAAAK_2
F1_TEVNVLPGAK_2
outliers
beta
alpha
ratio
time
0.00
0.01
0.02
0.03
0.04
0
5
10
15
20
25

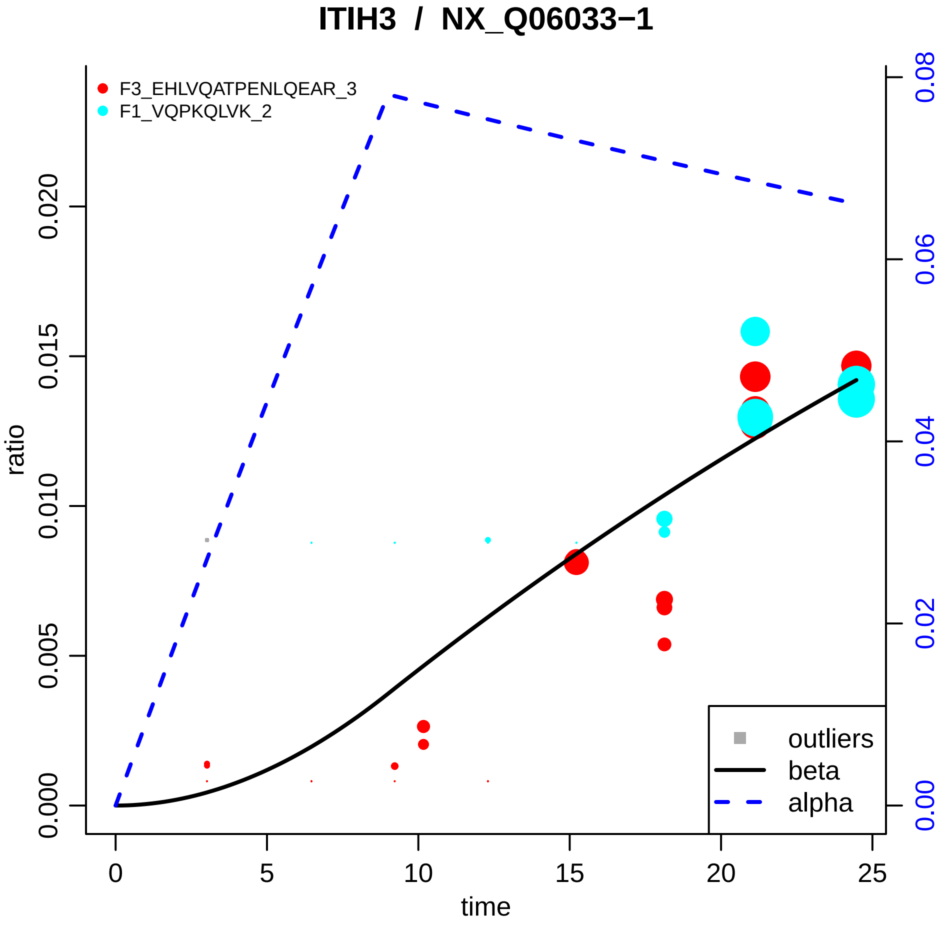

ITIH3 / NX_Q06033−1
F3_EHLVQATPENLQEAR_3
F1_VQPKQLVK_2
outliers
beta
alpha
ratio
time
0.000
0.005
0.010
0.015
0.020
0.00
0.02
0.04
0.06
0.08
0
5
10
15
20
25

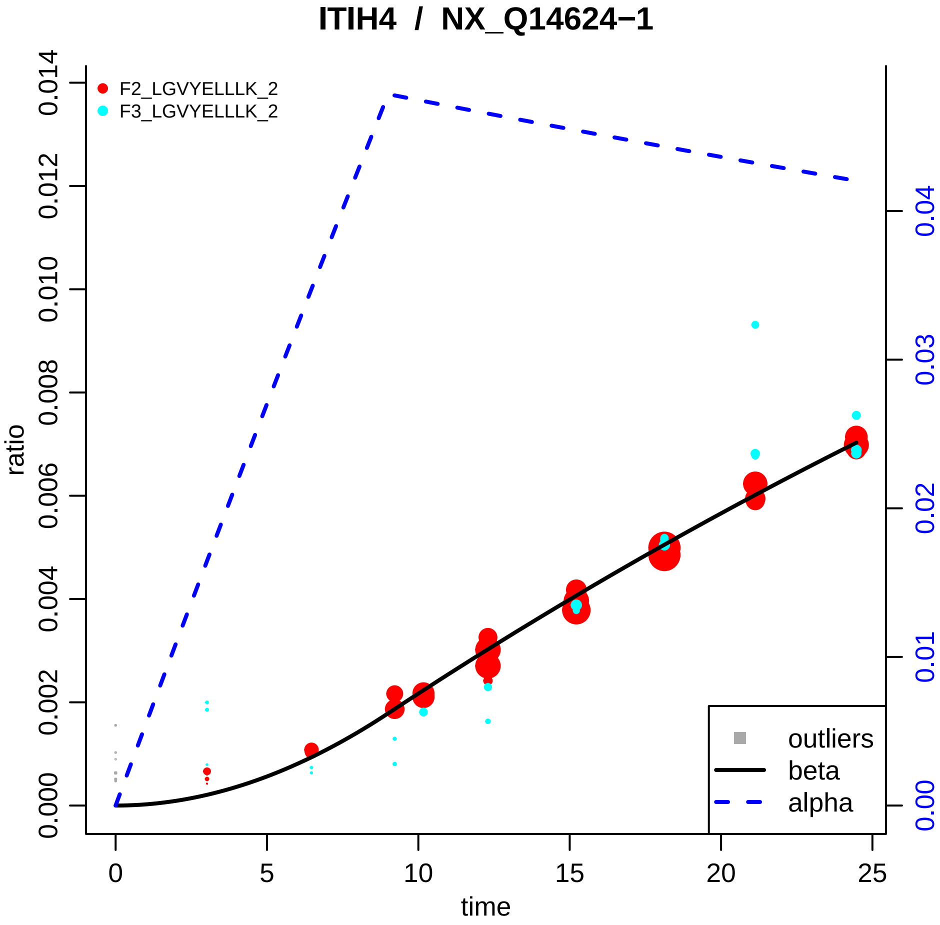

ITIH4 / NX_Q14624−1
F2_LGVYELLLK_2
F3_LGVYELLLK_2
ratio
time
0.000
0.002
0.004
0.006
0.008
0.010
0.012
0.014
0
5
10
15
20
25
0.00
0.01
0.02
0.03
0.04
outliers
beta
alpha

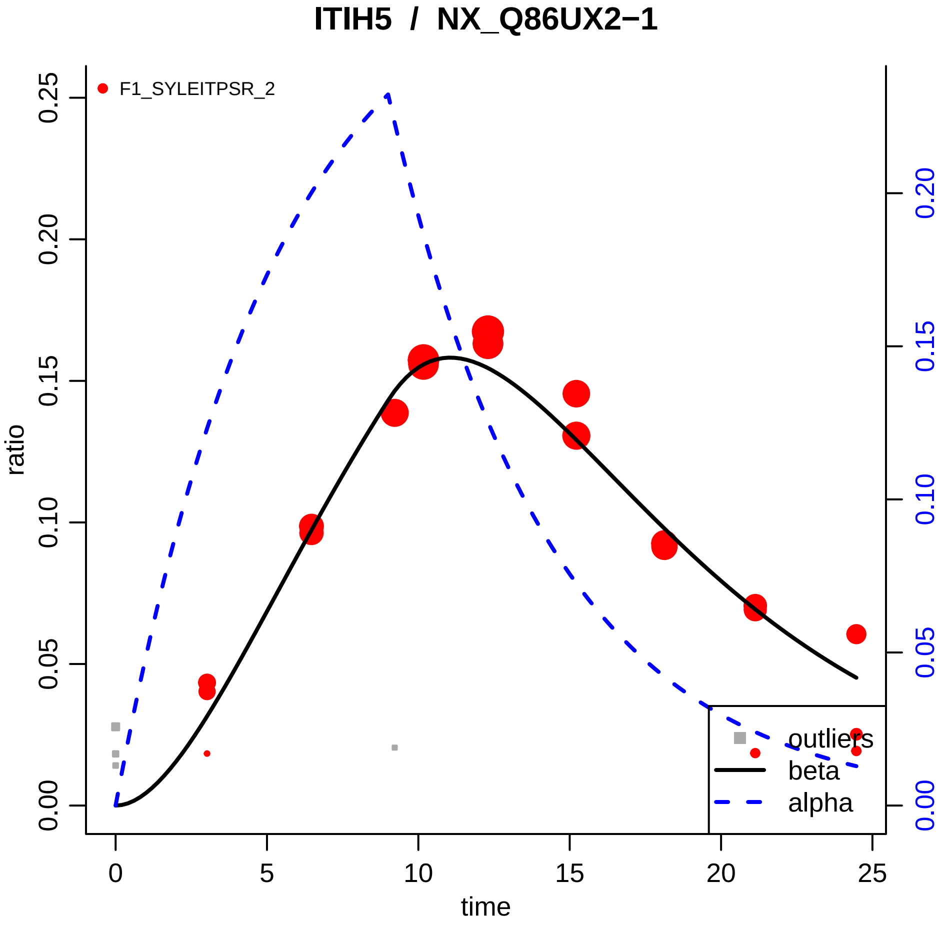

ITIH5 / NX_Q86UX2−1
F1_SYLEITPSR_2
ratio
time
0.00
0.05
0.10
0.15
0.20
0.25
0
5
10
15
20
25
outliers
beta
alpha

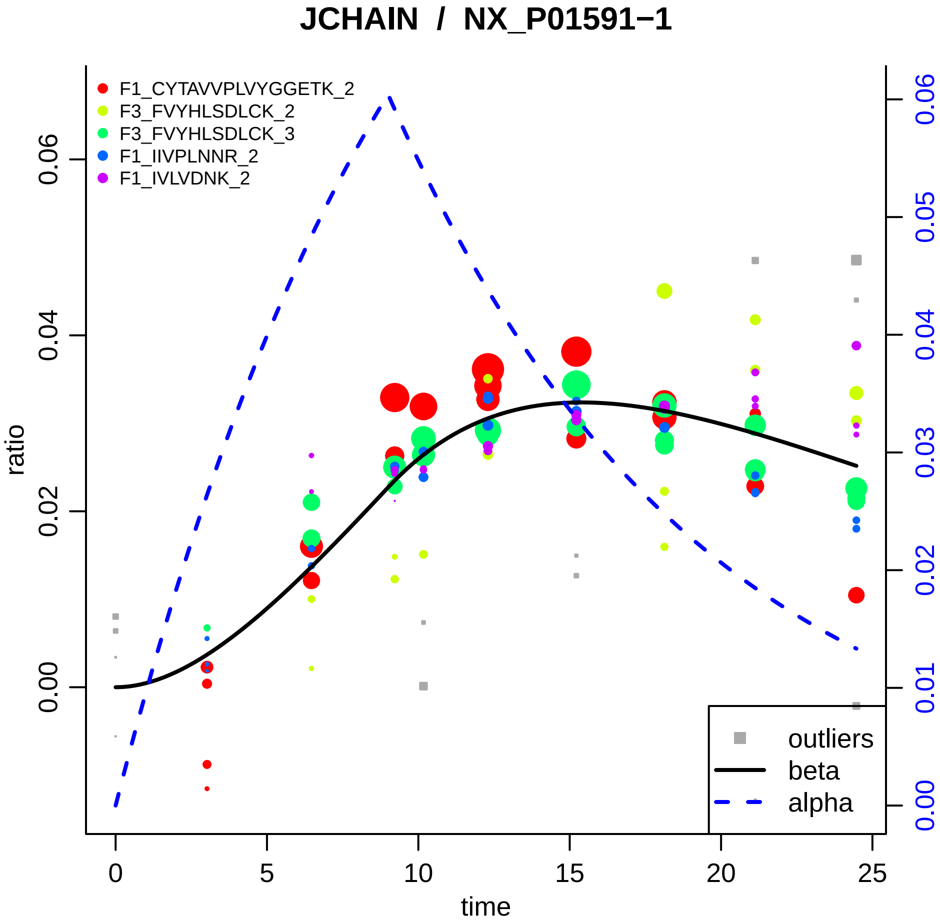

JCHAIN / NX_P01591−1
F1_CYTAVVPLVYGGETK_2
F3_FVYHLSDLCK_2
F3_FVYHLSDLCK_3
F1_IIVPLNNR_2
F1_IVLVDNK_2
outliers
beta
alpha
ratio
time
0.00
0.02
0.04
0.06
0
5
10
15
20
25
0.00
0.01
0.02
0.03
0.04
0.05
0.06

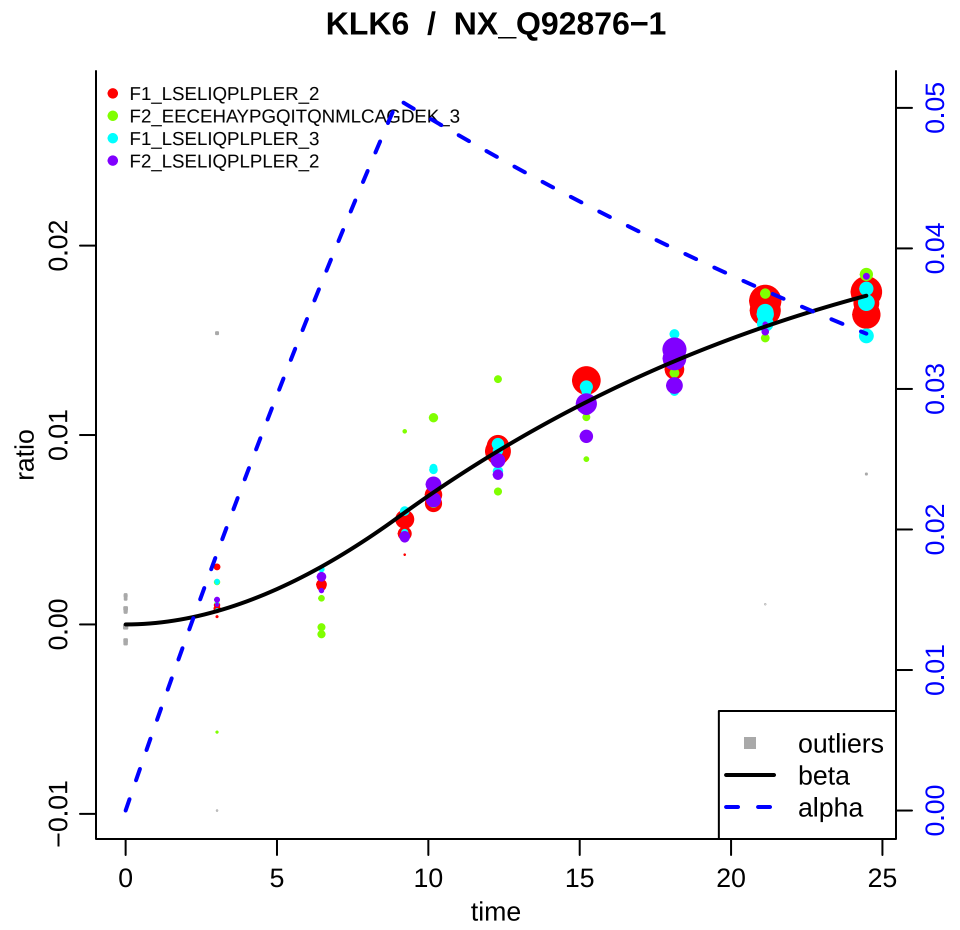

KLK6 / NX_Q92876−1
F1_LSELIQPLPLER_2
F2_EECEHAYPGQITQNMLCAGDEK_3
F1_LSELIQPLPLER_3
F2_LSELIQPLPLER_2
outliers
beta
alpha
ratio
time
−0.01
0.00
0.01
0.02
0
5
10
15
20
25
0.00
0.01
0.02
0.03
0.04
0.05

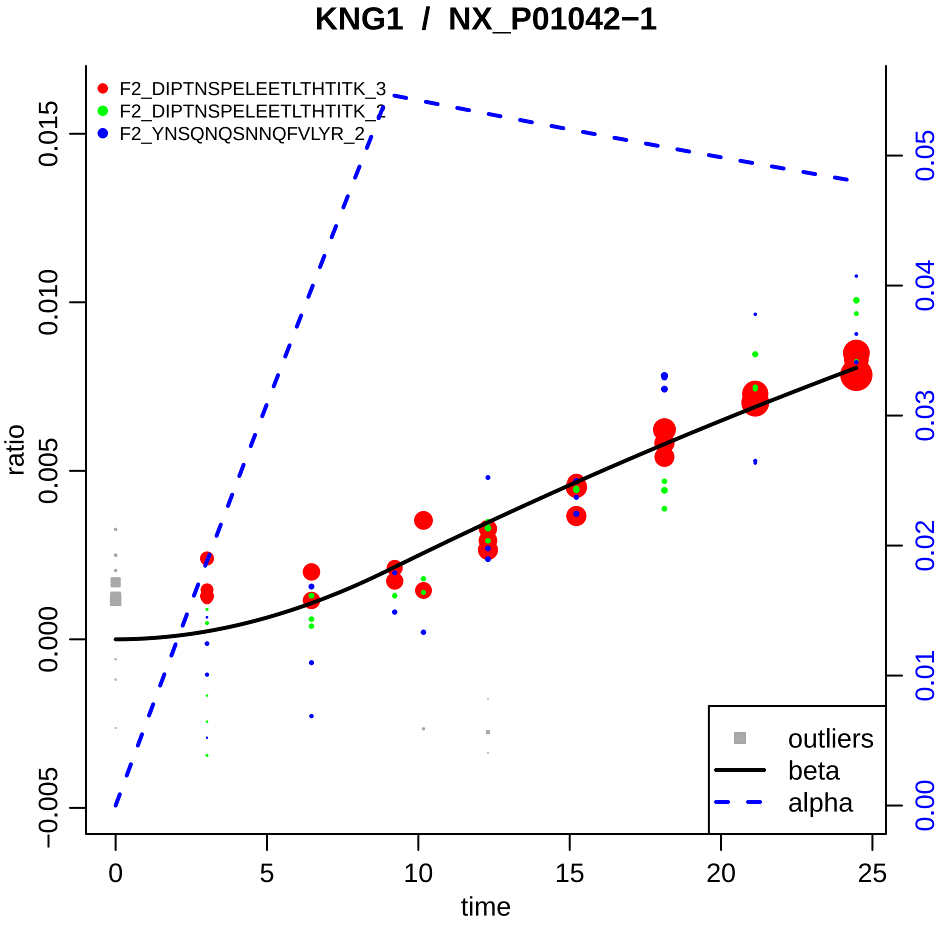

KNG1 / NX_P01042−1
F2_DIPTNSPELEETLTHTITK_3
F2_DIPTNSPELEETLTHTITK_2
F2_YNSQNQSNNQFVLYR_2
ratio
time
outliers
beta
alpha
−0.005
0.000
0.005
0.010
0.015
0
5
10
15
20
25
0.00
0.01
0.02
0.03
0.04
0.05

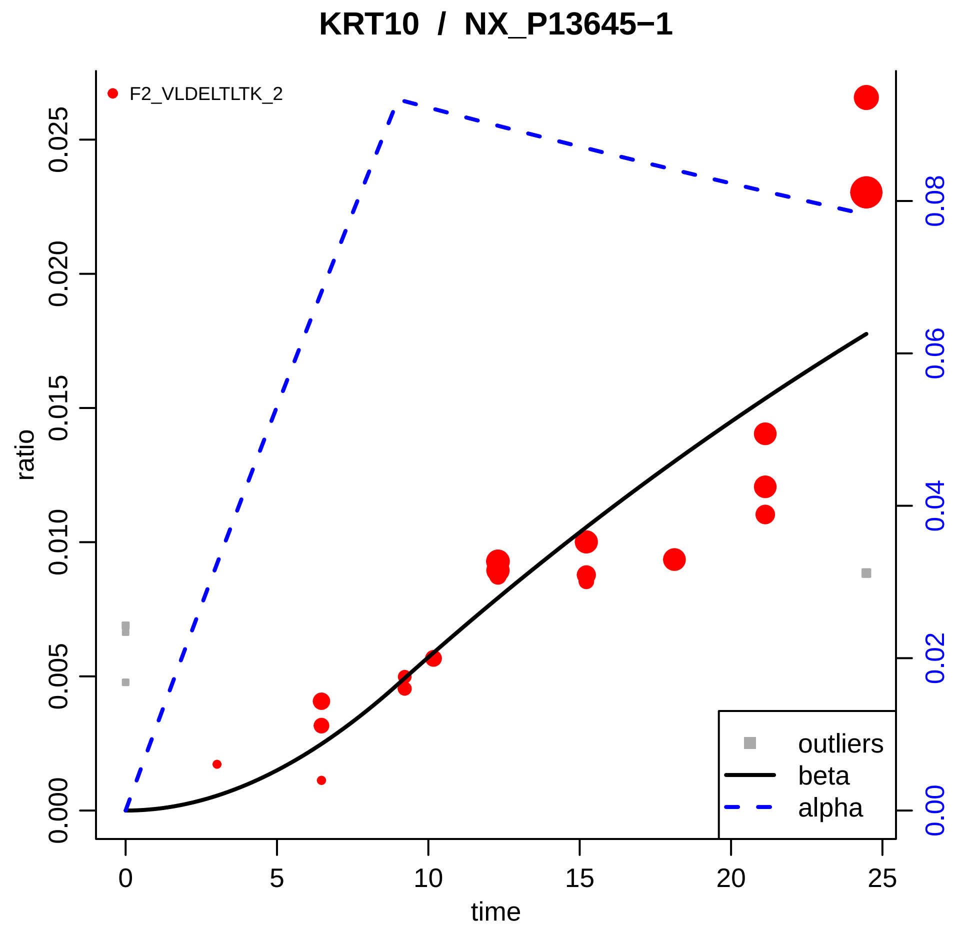

KRT10 / NX_P13645−1
F2_VLDELTLTK_2
ratio
0.000
0.005
0.010
0.015
0.020
0.025
0.00
0.02
0.04
0.06
0.08
0
5
10
15
20
25
time
outliers
beta
alpha

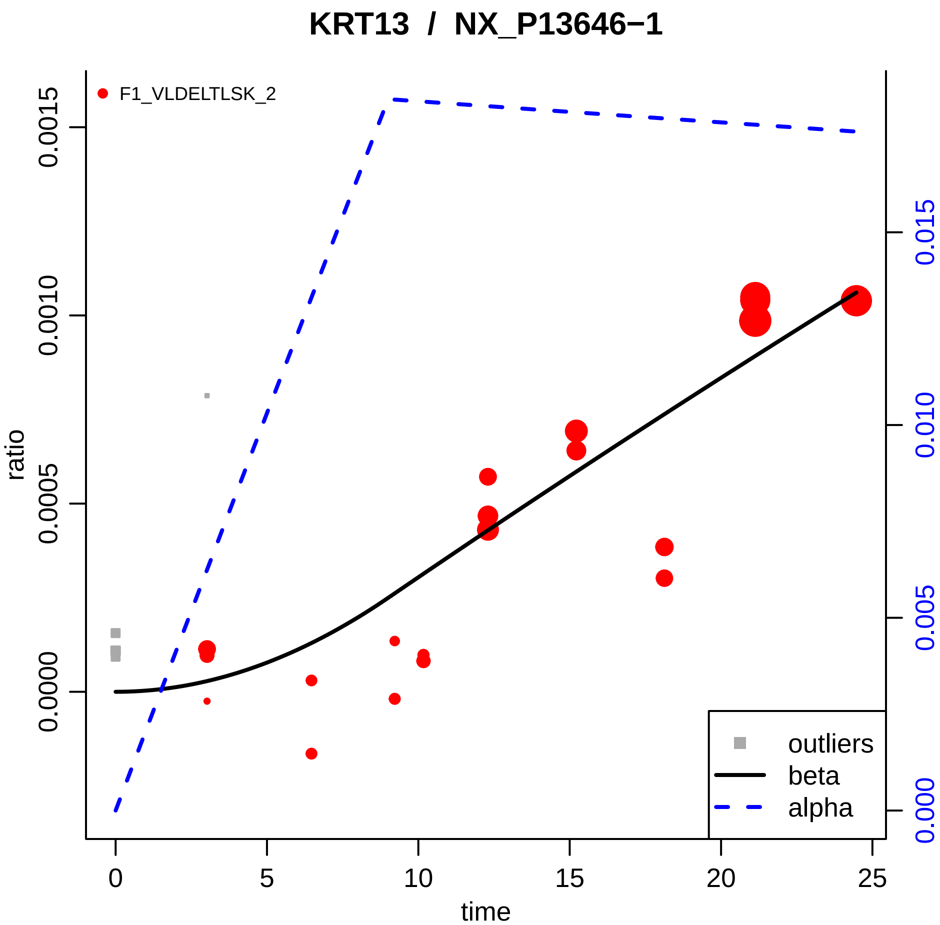

KRT13 / NX_P13646−1
F1_VLDELTLSK_2
ratio
0.0000
0.0005
0.0010
0.0015
time
0
5
10
15
20
25
0.000
0.005
0.010
0.015
outliers
beta
alpha

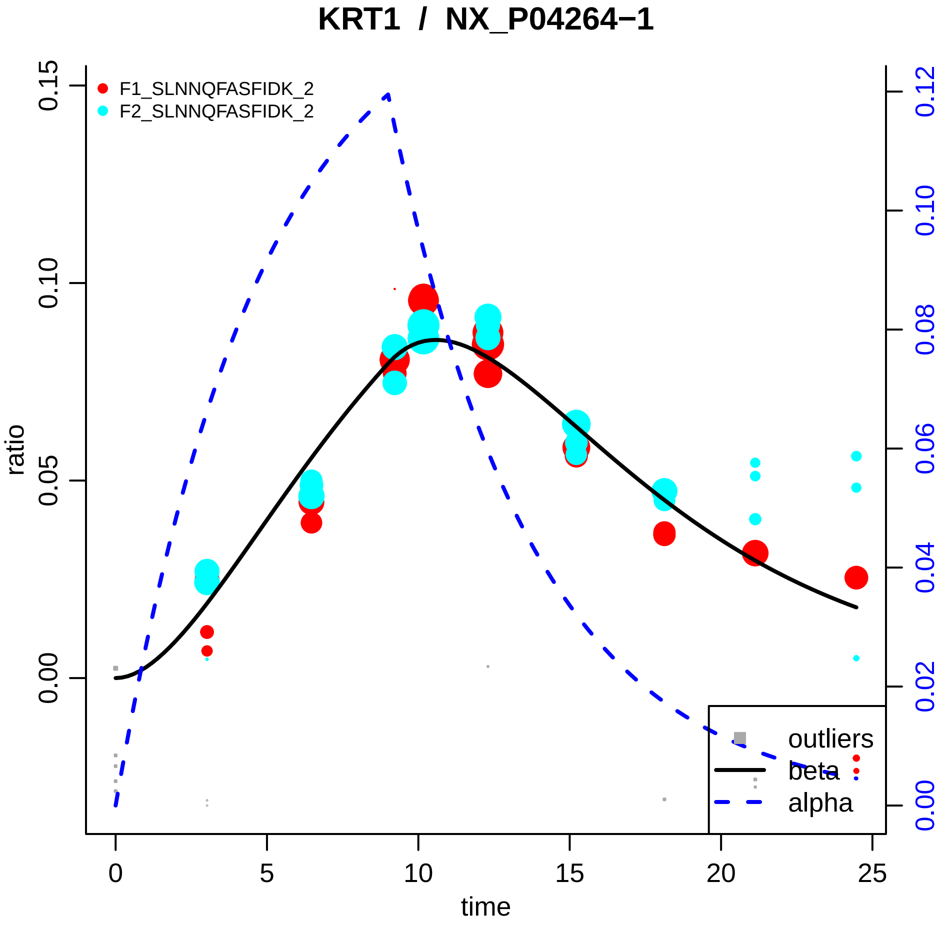

KRT1 / NX_P04264−1
F1_SLNNQFASFIDK_2
F2_SLNNQFASFIDK_2
ratio
0.00
0.05
0.10
0.15
time
0
5
10
15
20
25
0.00
0.02
0.04
0.06
0.08
0.10
0.12
outliers
beta
alpha

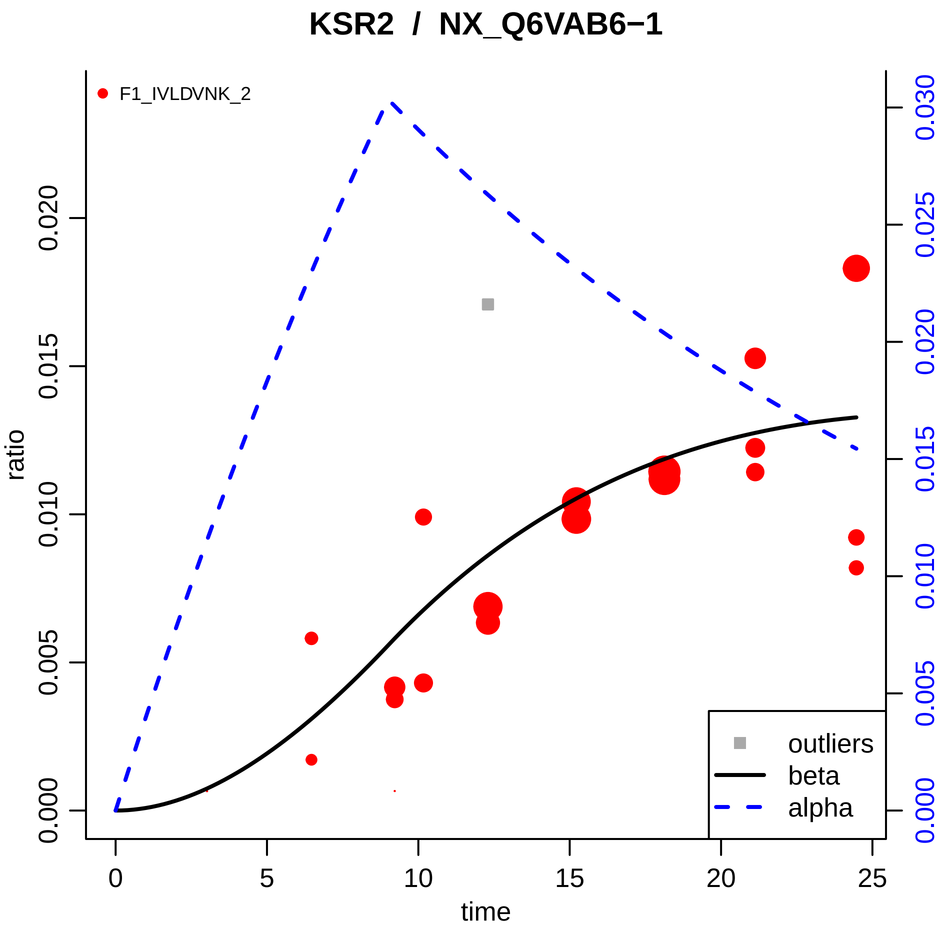

KSR2 / NX_Q6VAB6−1
F1_IVLDVNK_2
ratio
time
0.000
0.005
0.010
0.015
0.020
0.025
0.030
0
5
10
15
20
25
outliers
beta
alpha

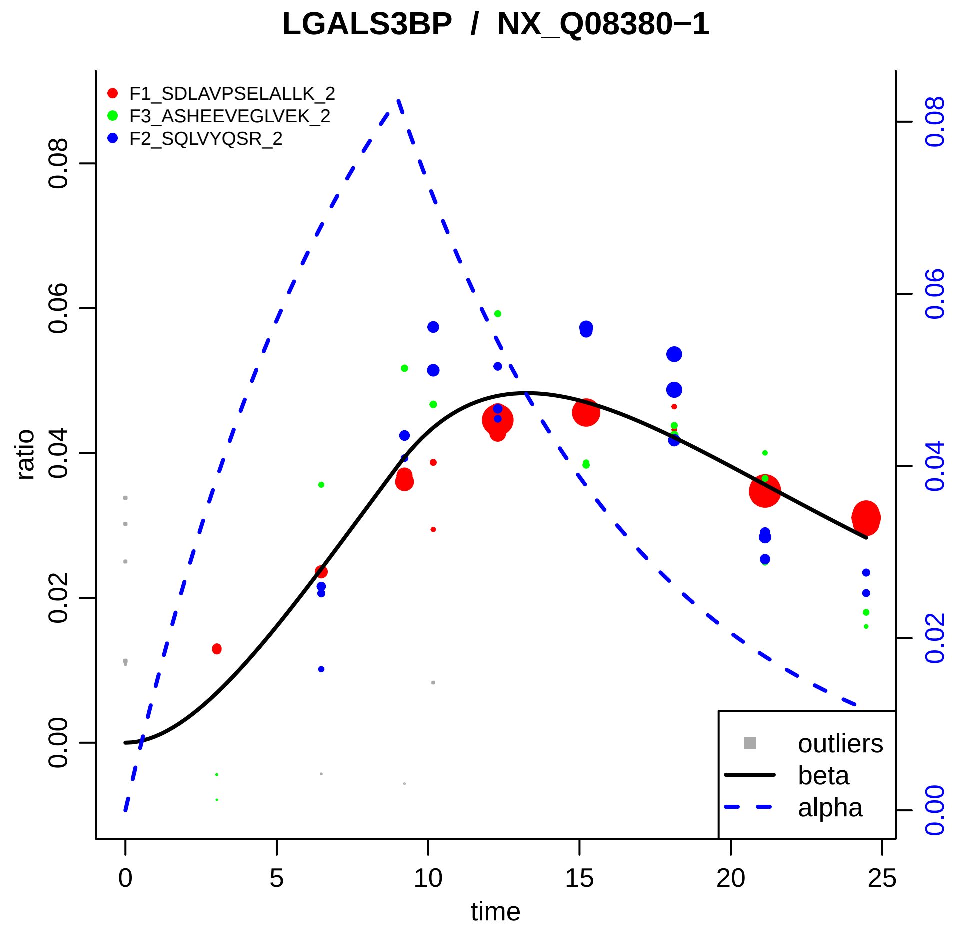
LGALS3BP / NX_Q08380−1
F1_SDLAVPSELALLK_2
F3_ASHEEVEGLVEK_2
F2_SQLVYQSR_2
ratio
0.00
0.02
0.04
0.06
0.08
time
0
5
10
15
20
25
outliers
beta
alpha

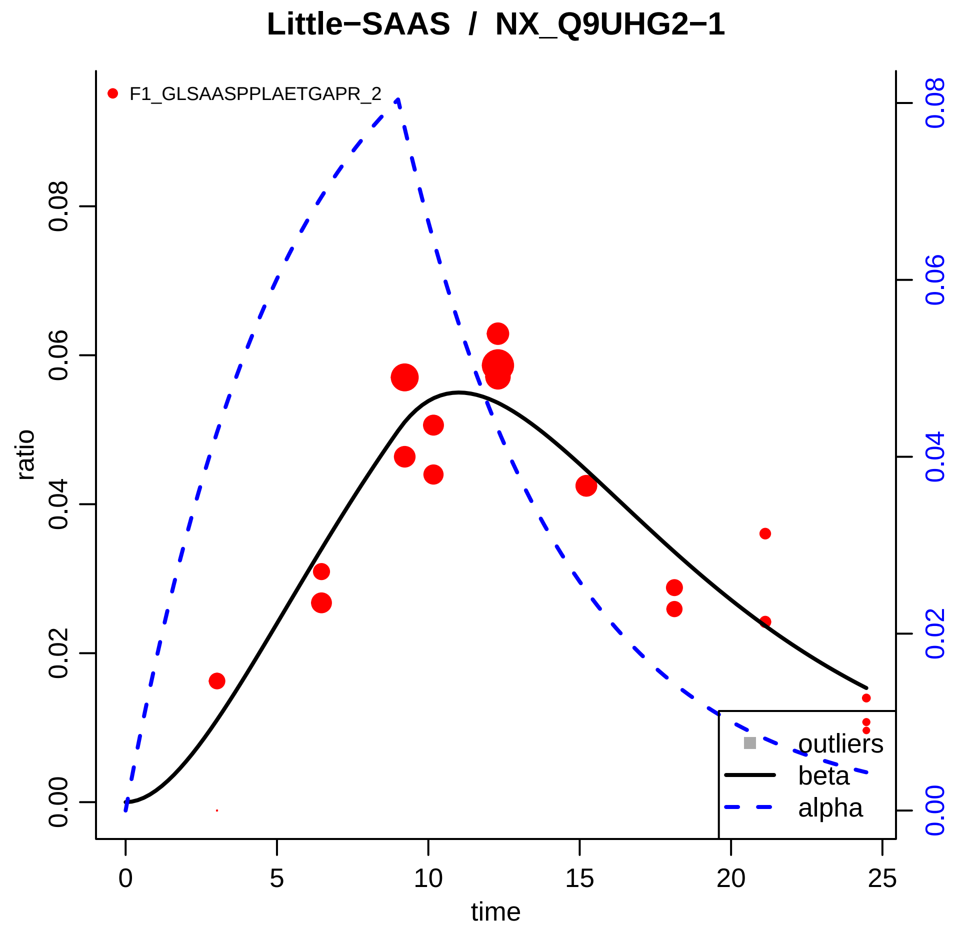

Little−SAAS / NX_Q9UHG2−1
F1_GLSAASPPLAETGAPR_2
ratio
0.00
0.02
0.04
0.06
0.08
time
0
5
10
15
20
25
outliers
beta
alpha

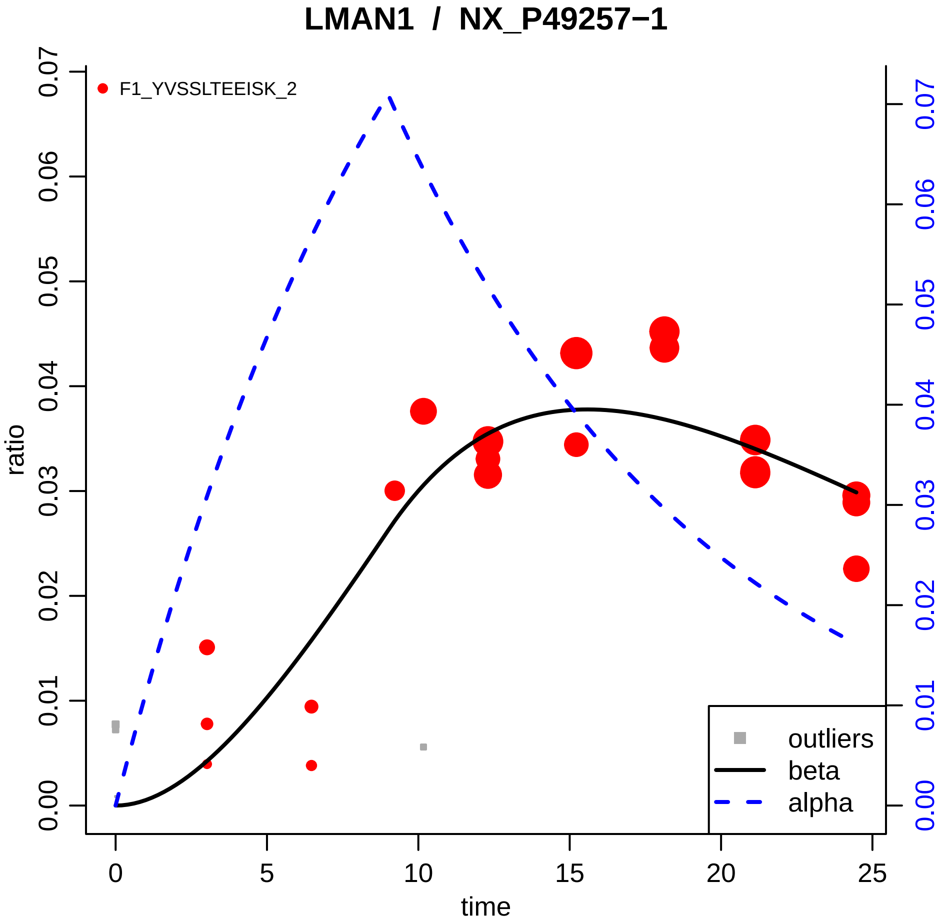

LMAN1 / NX_P49257−1
F1_YVSSLTEEISK_2
ratio
time
0.00
0.01
0.02
0.03
0.04
0.05
0.06
0.07
0
5
10
15
20
25
outliers
beta
alpha

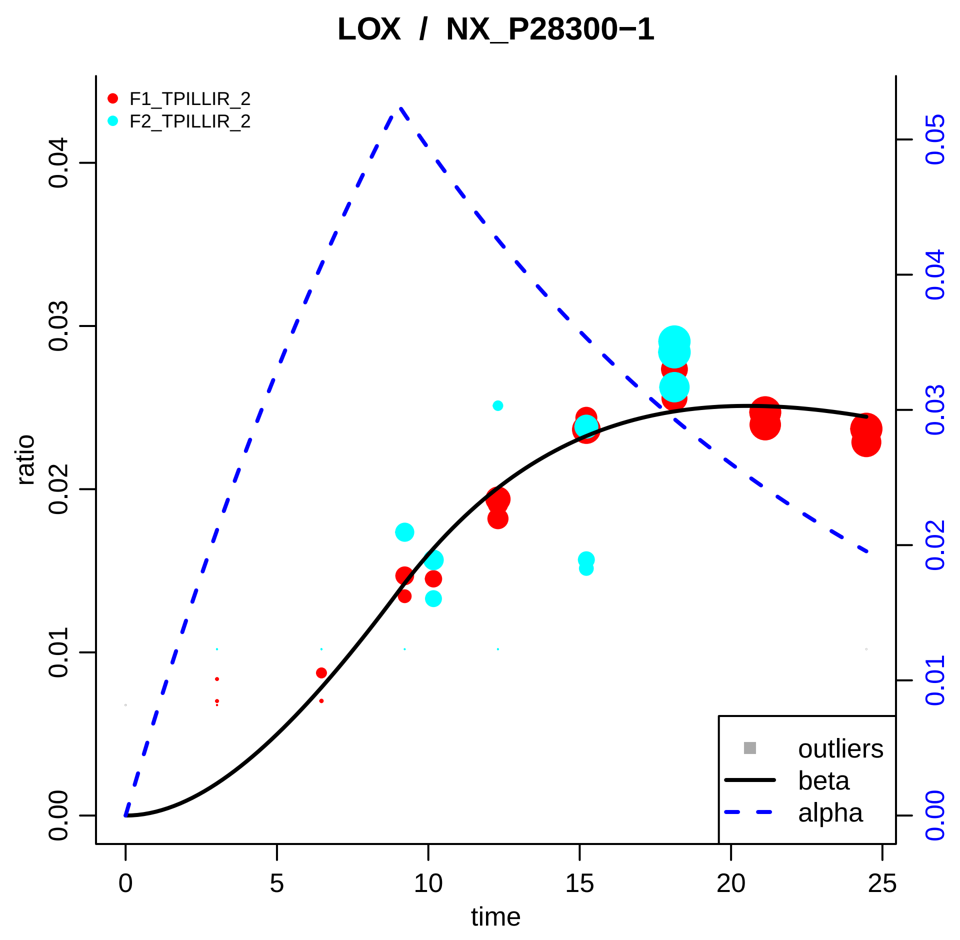

LOX / NX_P28300−1
F1_TPILLIR_2
F2_TPILLIR_2
ratio
0.00
0.01
0.02
0.03
0.04
0.05
time
0
5
10
15
20
25
outliers
beta
alpha

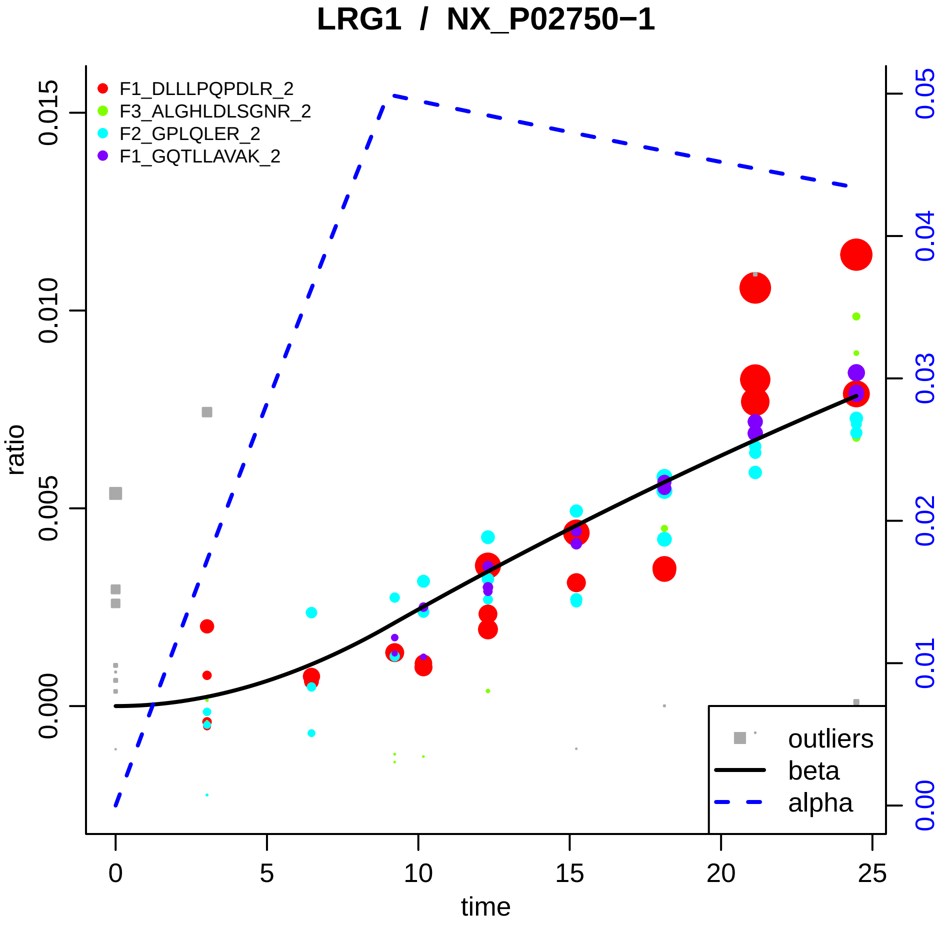

LRG1 / NX_P02750−1
F1_DLLLPQPDLR_2
F3_ALGHLDLSGNR_2
F2_GPLQLER_2
F1_GQTLLAVAK_2
ratio
0.000
0.005
0.010
0.015
time
0
5
10
15
20
25
0.00
0.01
0.02
0.03
0.04
0.05
outliers
beta
alpha

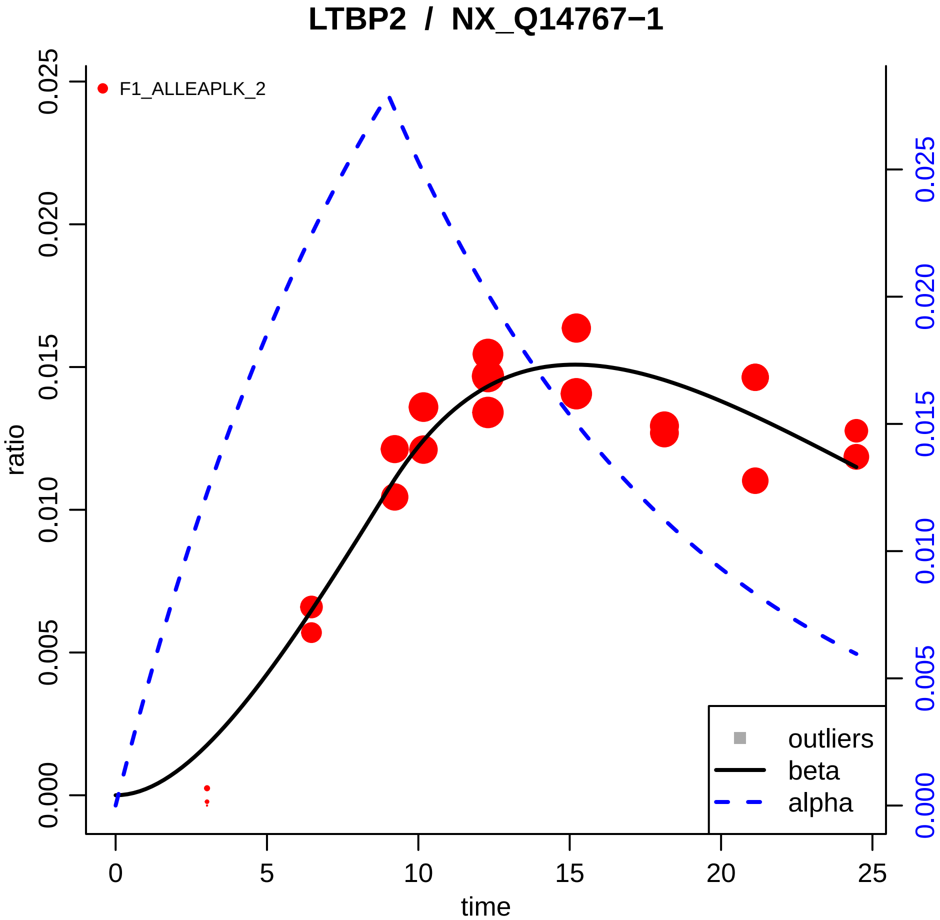

LTBP2 / NX_Q14767−1
F1_ALLEAPLK_2
ratio
time
outliers
beta
alpha
0.000
0.005
0.010
0.015
0.020
0.025
0
5
10
15
20
25

### LYVE1 / NX_Q9Y5Y7−1

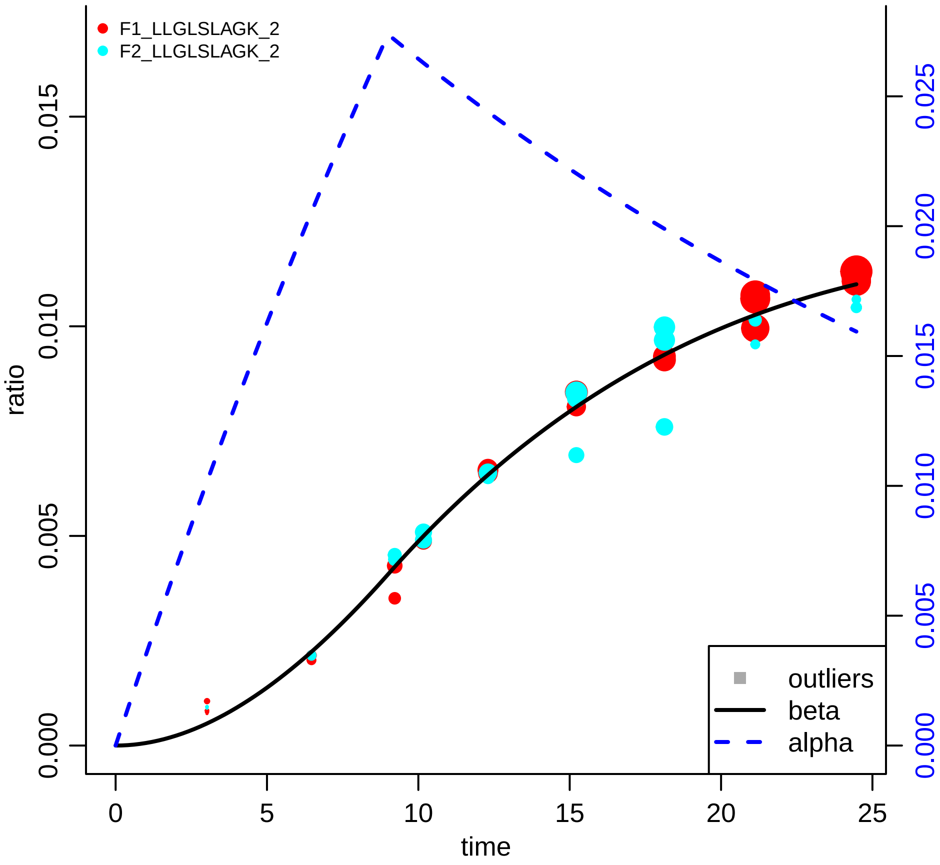

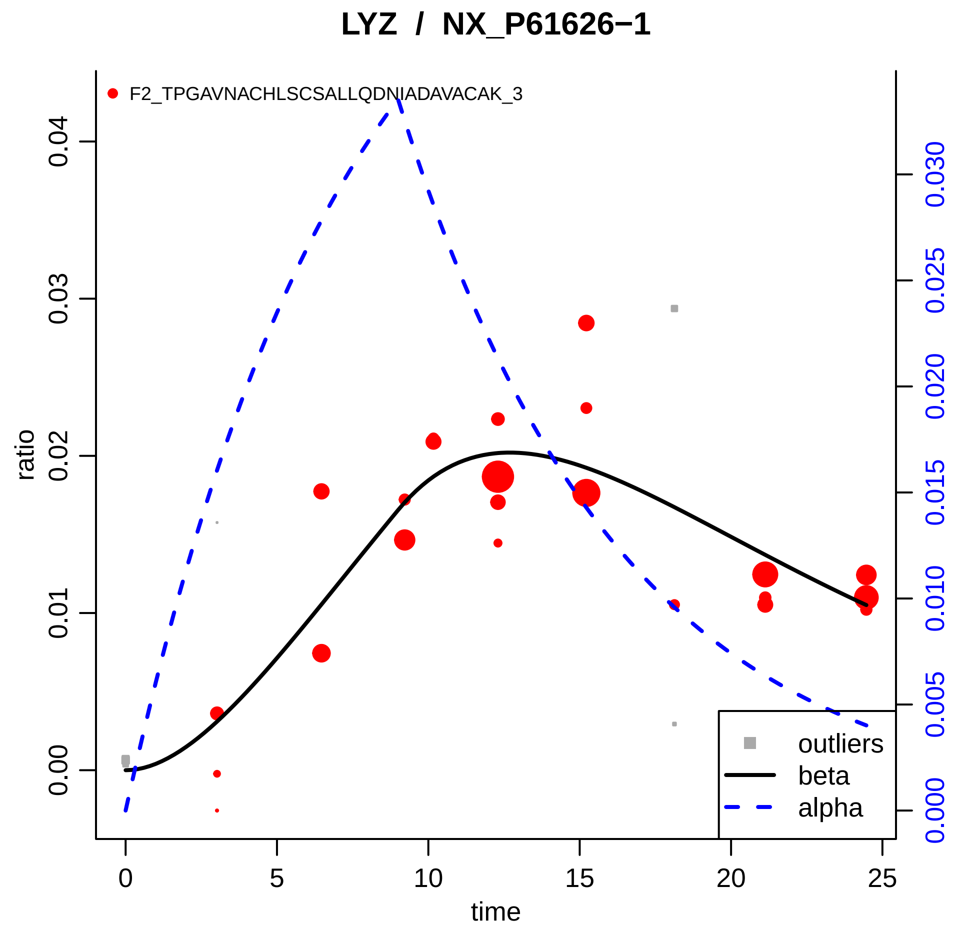

LYZ / NX_P61626−1
F2_TPGAVNACHLSCSALLQDNIADAVACAK_3
ratio
0.00
0.01
0.02
0.03
0.04
time
0
5
10
15
20
25
0.000
0.005
0.010
0.015
0.020
0.025
0.030
outliers
beta
alpha

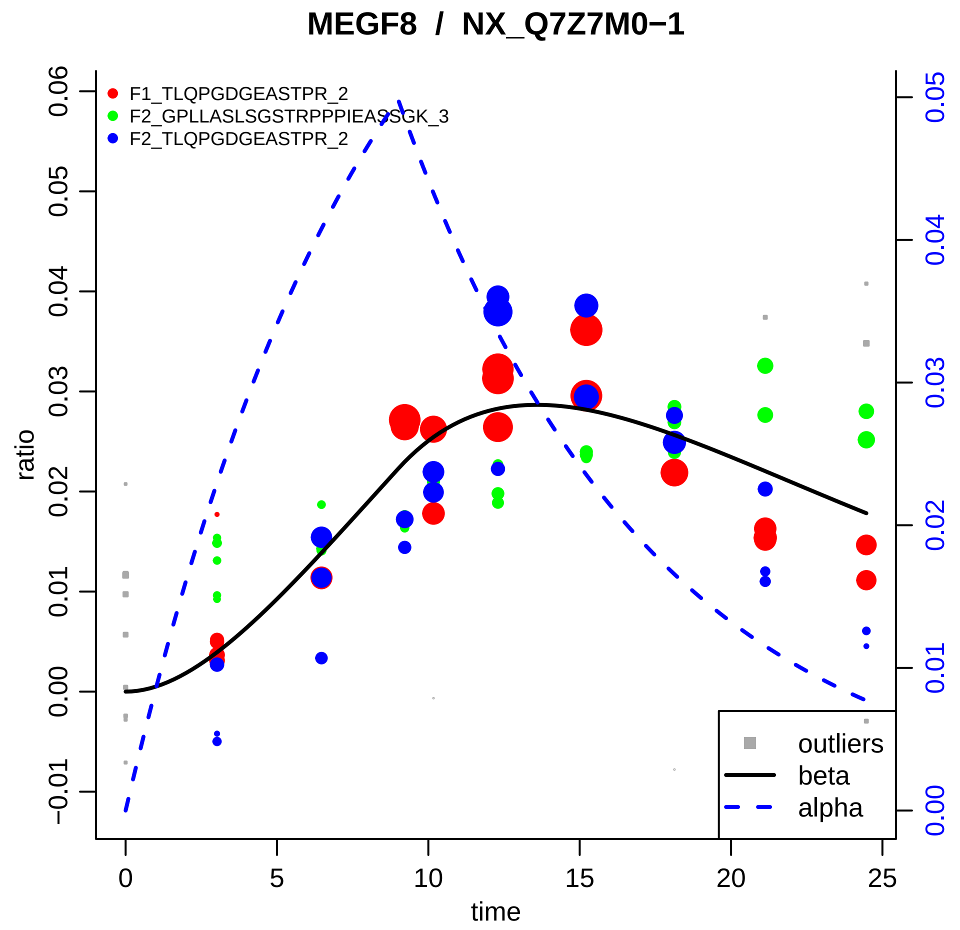

MEGF8 / NX_Q7Z7M0−1
F1_TLQPGDGEASTPR_2
F2_GPLLASLSGSTRPPPIEASSGK_3
F2_TLQPGDGEASTPR_2
ratio
time
outliers
beta
alpha
−0.01
0.00
0.01
0.02
0.03
0.04
0.05
0.06
0
5
10
15
20
25

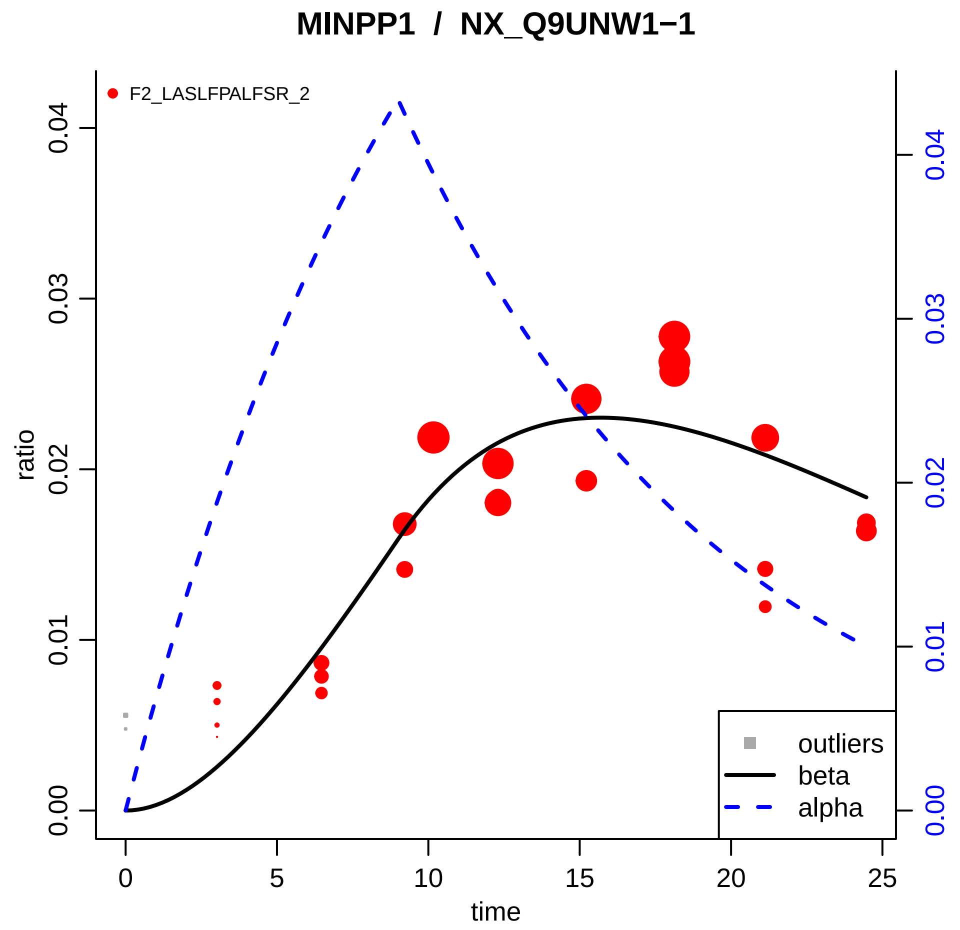

MINPP1 / NX_Q9UNW1−1
F2_LASLFPALFSR_2
ratio
time
0.00
0.01
0.02
0.03
0.04
0
5
10
15
20
25
outliers
beta
alpha

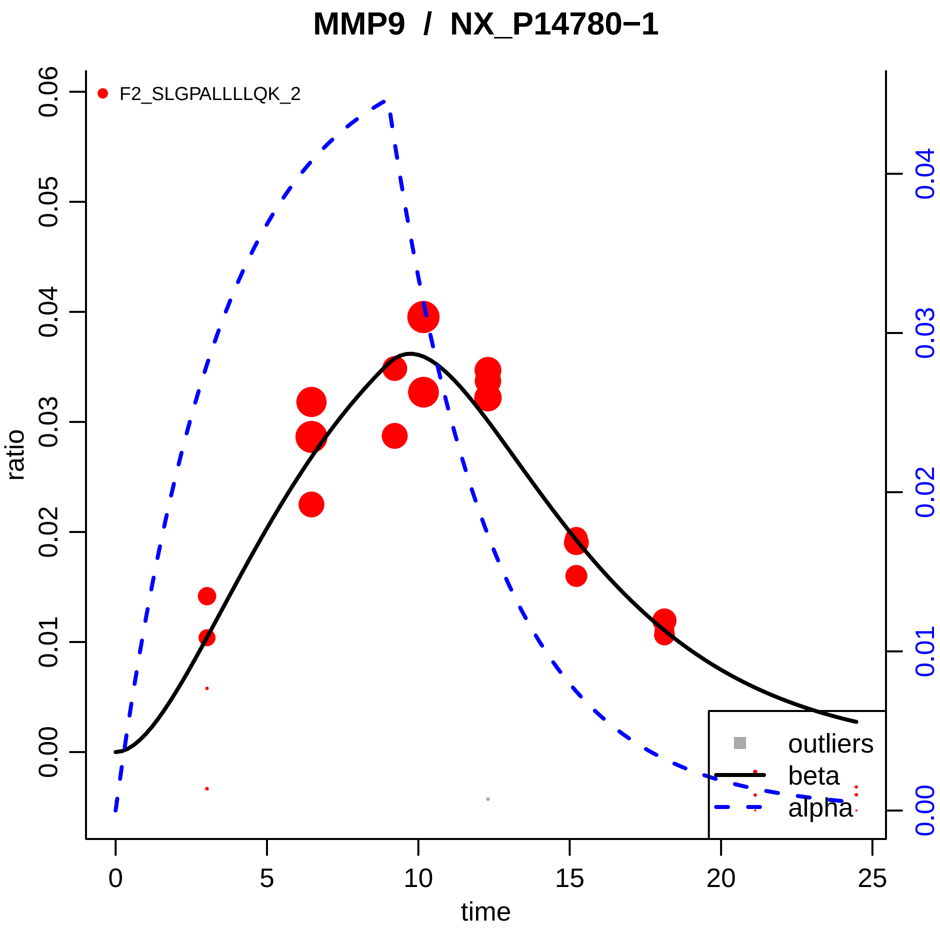

MMP9 / NX_P14780−1
F2_SLGPALLLLQK_2
ratio
time
0.00
0.01
0.02
0.03
0.04
0.05
0.06
0
5
10
15
20
25
0.00
0.01
0.02
0.03
0.04
outliers
beta
alpha

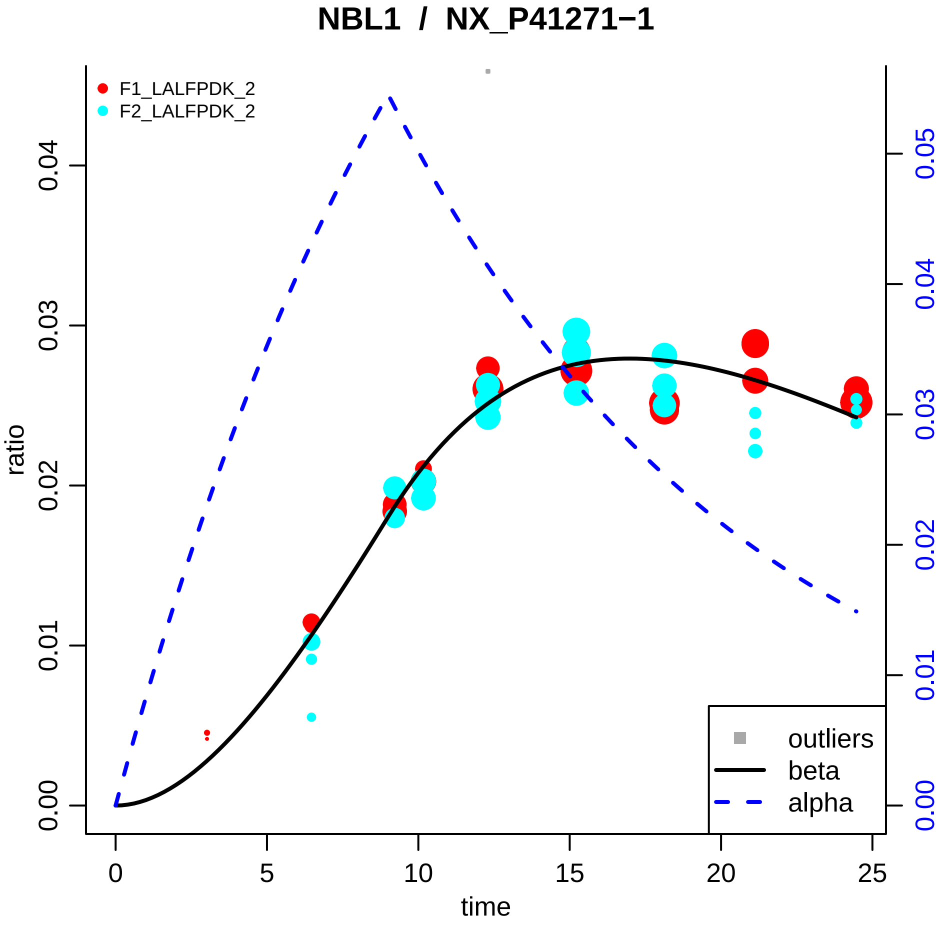

NBL1 / NX_P41271−1
F1_LALFPDK_2
F2_LALFPDK_2
ratio
time
0.00
0.01
0.02
0.03
0.04
0.05
0
5
10
15
20
25
outliers
beta
alpha

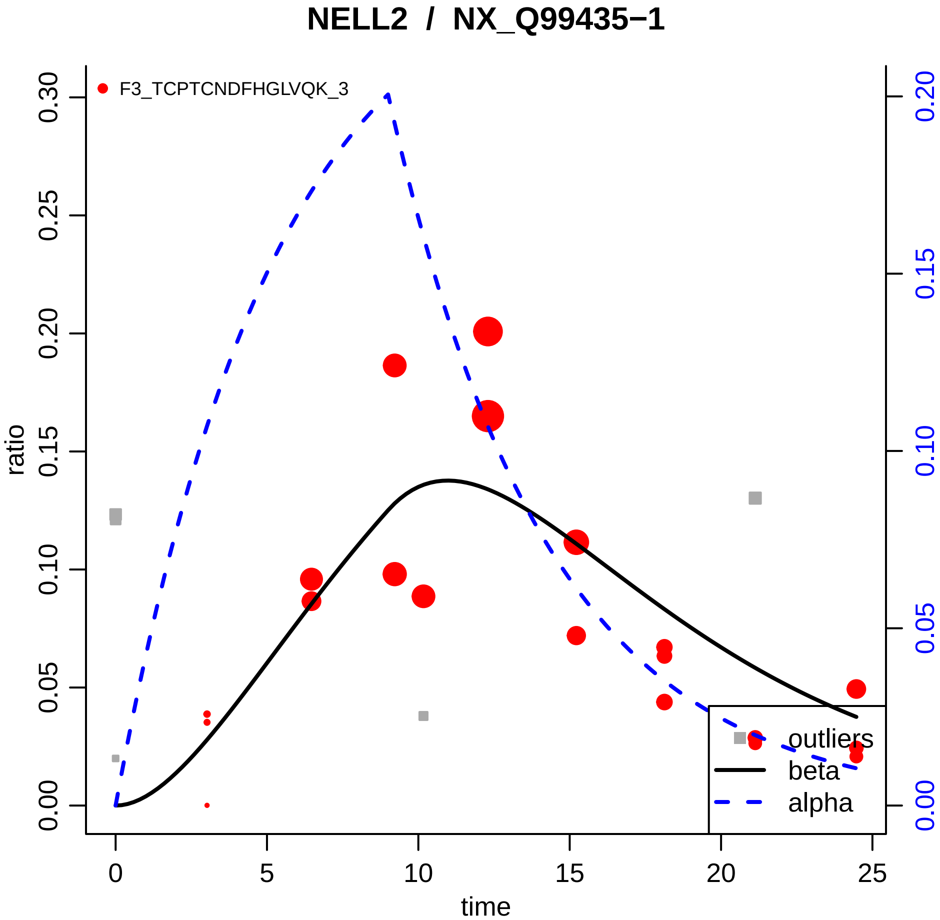

NELL2 / NX_Q99435−1
F3_TCPTCNDFHGLVQK_3
ratio
time
0.00
0.05
0.10
0.15
0.20
0.25
0.30
0
5
10
15
20
25
0.00
0.05
0.10
0.15
0.20
outliers
beta
alpha

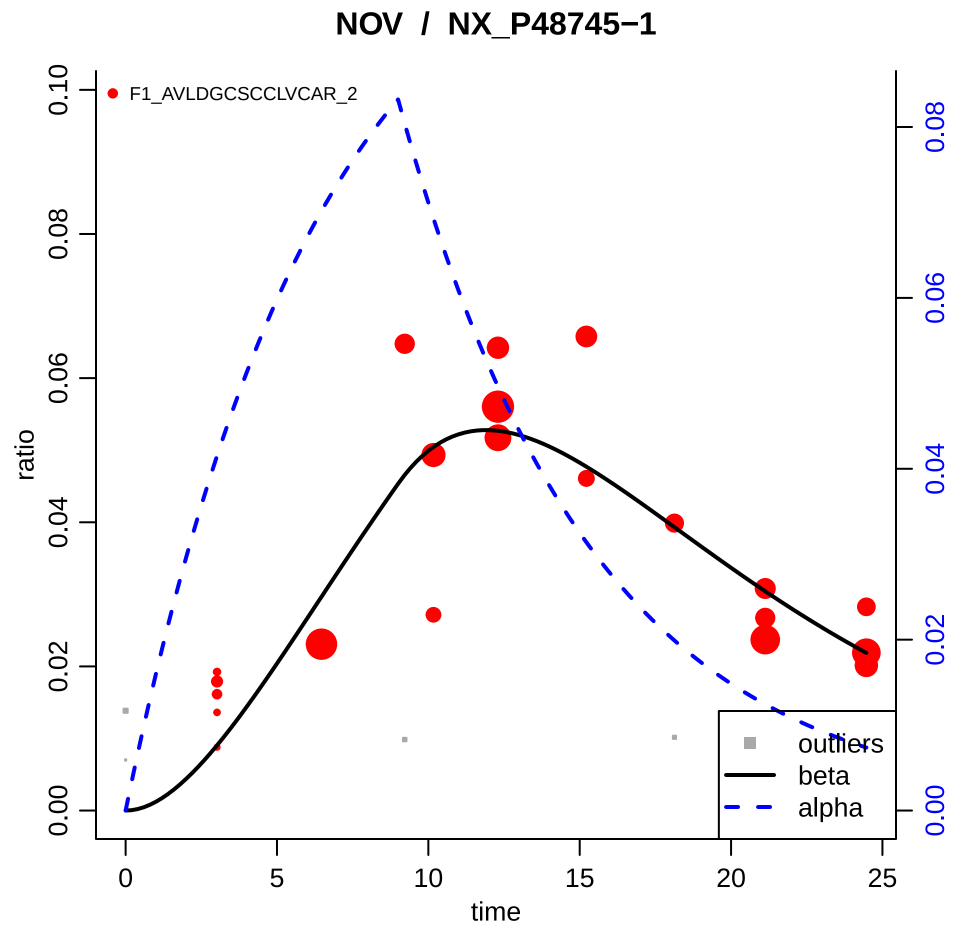

NOV / NX_P48745−1
F1_AVLDGCSCCLVCAR_2
ratio
time
0.00
0.02
0.04
0.06
0.08
0.10
0
5
10
15
20
25
outliers
beta
alpha

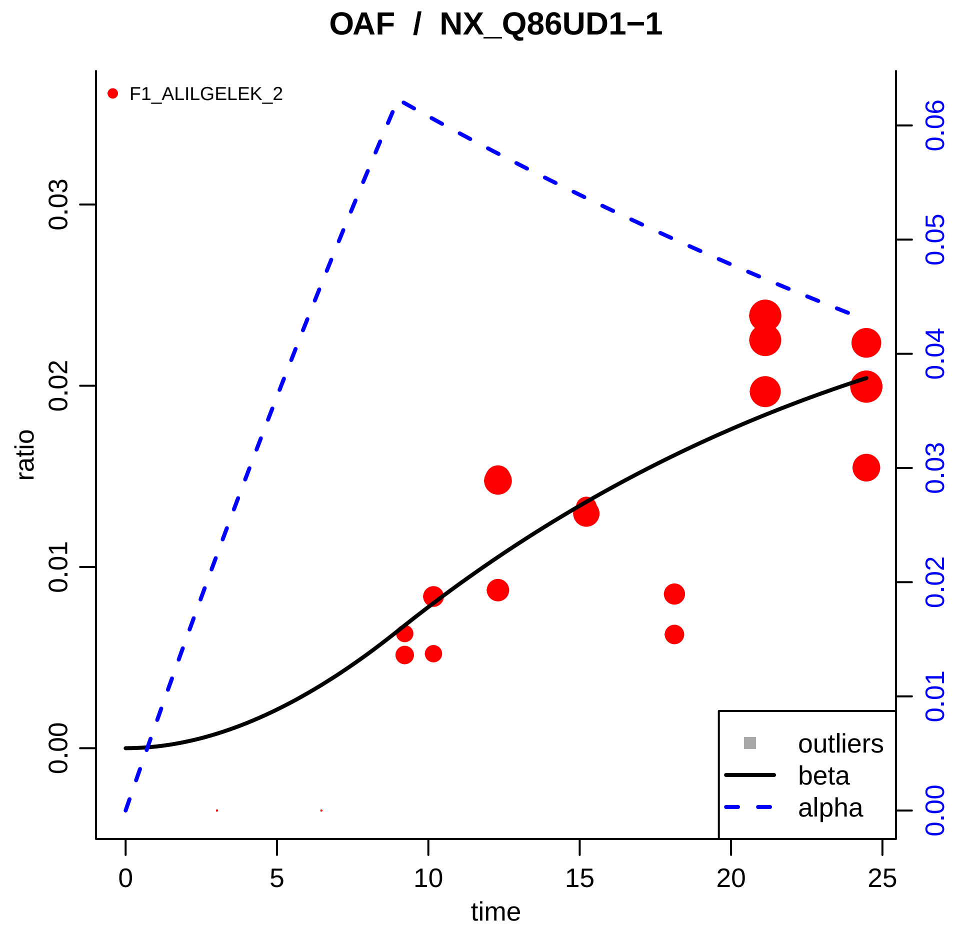

OAF / NX_Q86UD1−1
F1_ALILGELEK_2
ratio
time
0.00
0.01
0.02
0.03
0
5
10
15
20
25
0.00
0.01
0.02
0.03
0.04
0.05
0.06
outliers
beta
alpha

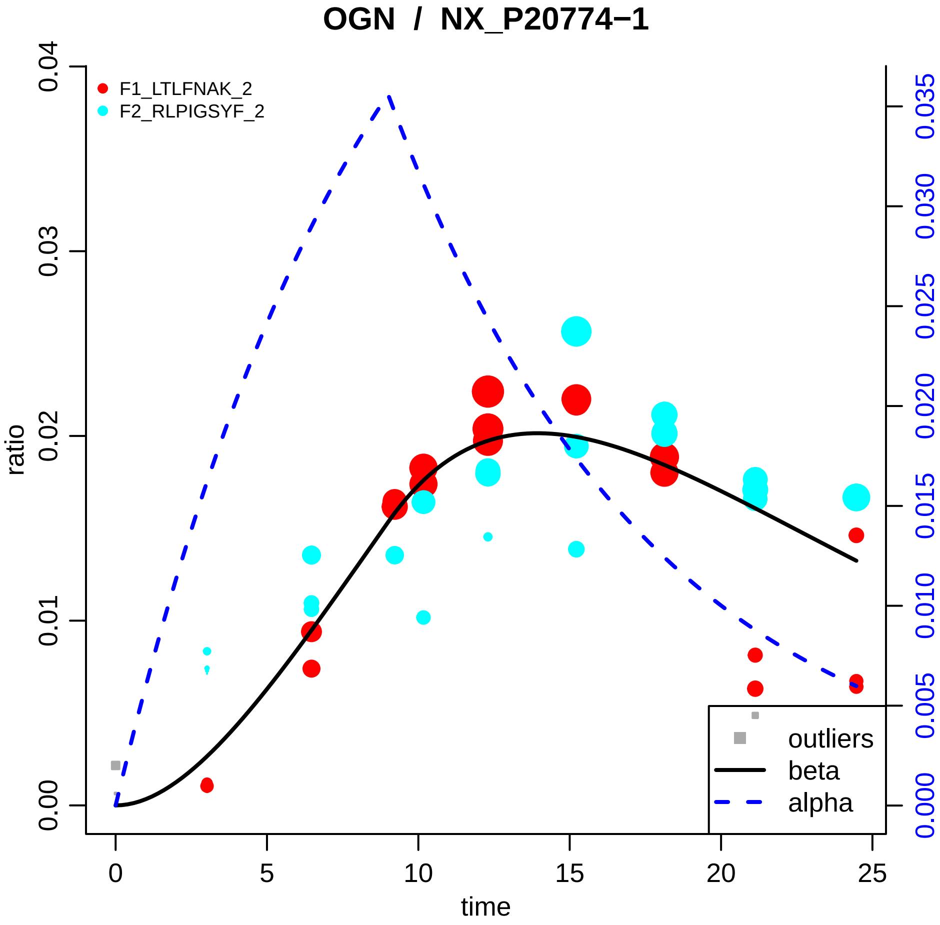

OGN / NX_P20774−1
F1_LTLFNAK_2
F2_RLPIGSYF_2
ratio
0.00
0.01
0.02
0.03
0.04
time
0
5
10
15
20
25
0.000
0.005
0.010
0.015
0.020
0.025
0.030
0.035
outliers
beta
alpha

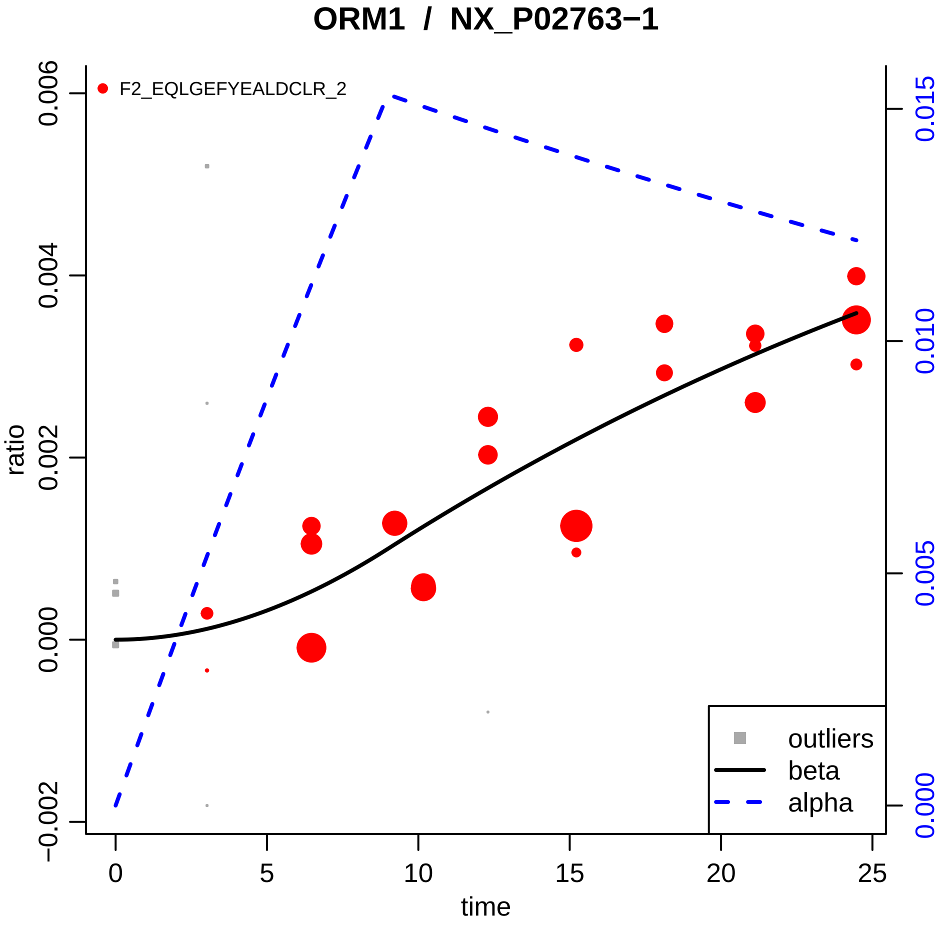

ORM1 / NX_P02763−1
F2_EQLGEFYEALDCLR_2
ratio
time
0.006
0.004
0.002
0.000
−0.002
0.015
0.010
0.005
0.000
0
5
10
15
20
25
outliers
beta
alpha

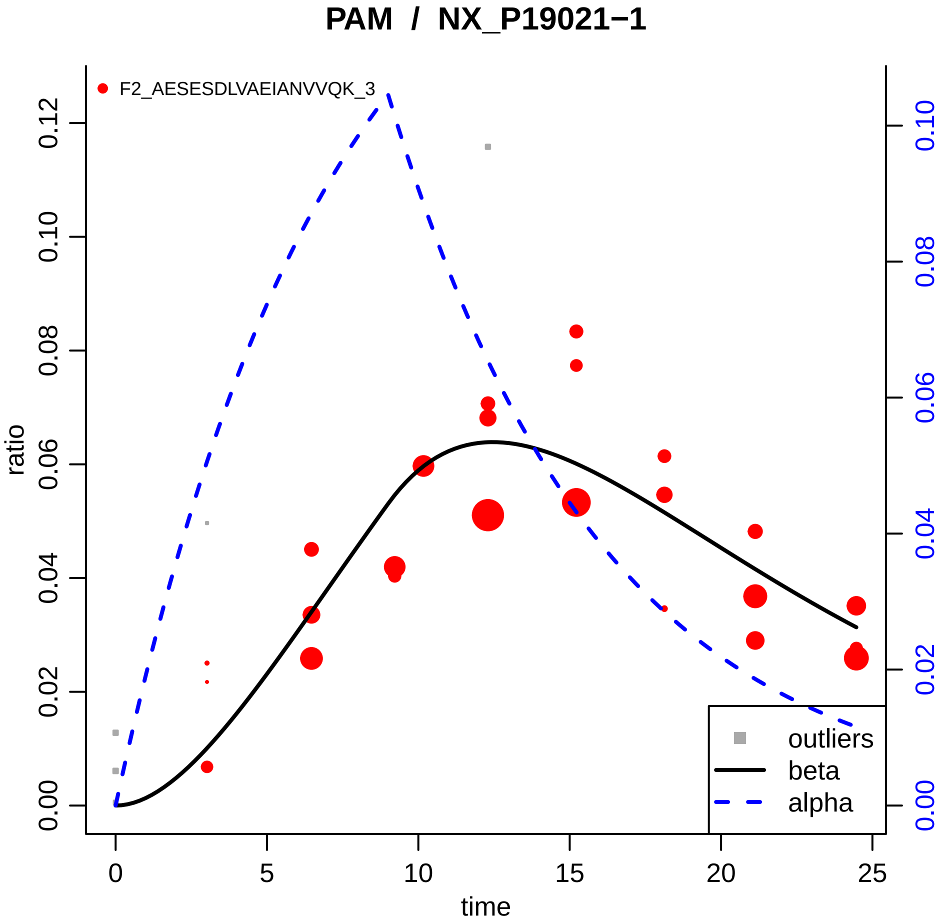

PAM / NX_P19021−1
F2_AESESDLVAEIANVVQK_3
ratio
time
0.00
0.02
0.04
0.06
0.08
0.10
0.12
0
5
10
15
20
25
0.00
0.02
0.04
0.06
0.08
0.10
outliers
beta
alpha

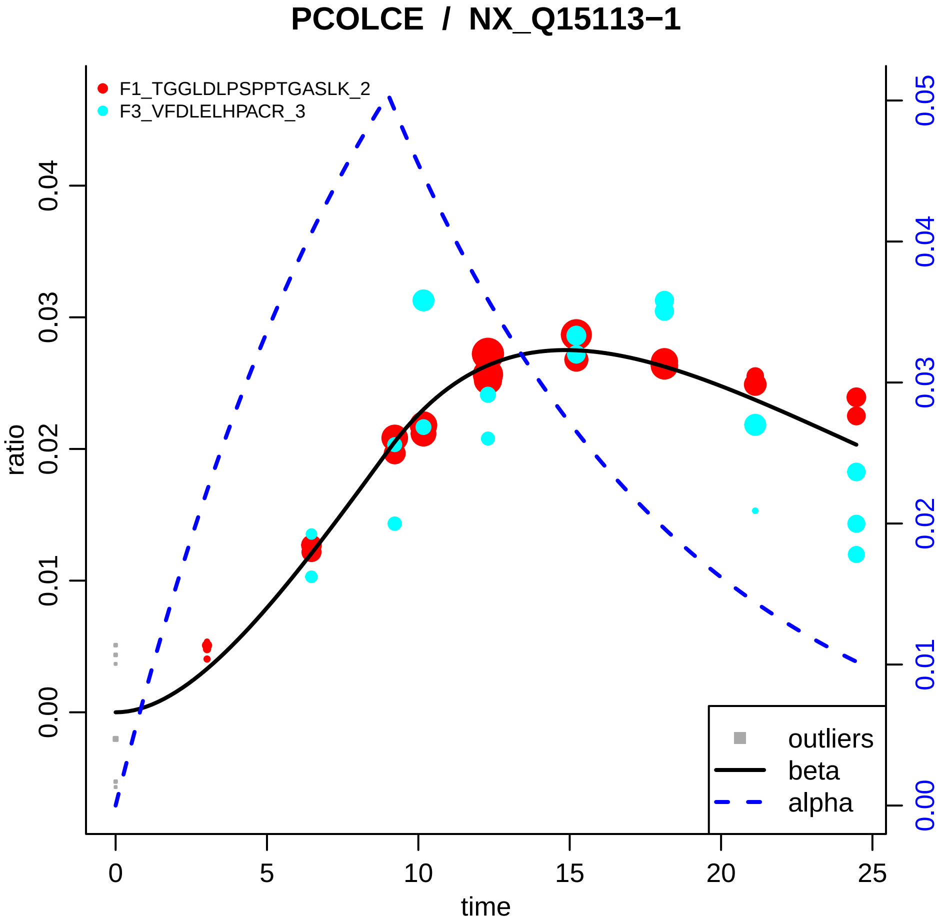

PCOLCE / NX_Q15113−1
F1_TGGLDLPSPPTGASLK_2
F3_VFDLELHPACR_3
ratio
0.00
0.01
0.02
0.03
0.04
0.05
time
0
5
10
15
20
25
outliers
beta
alpha

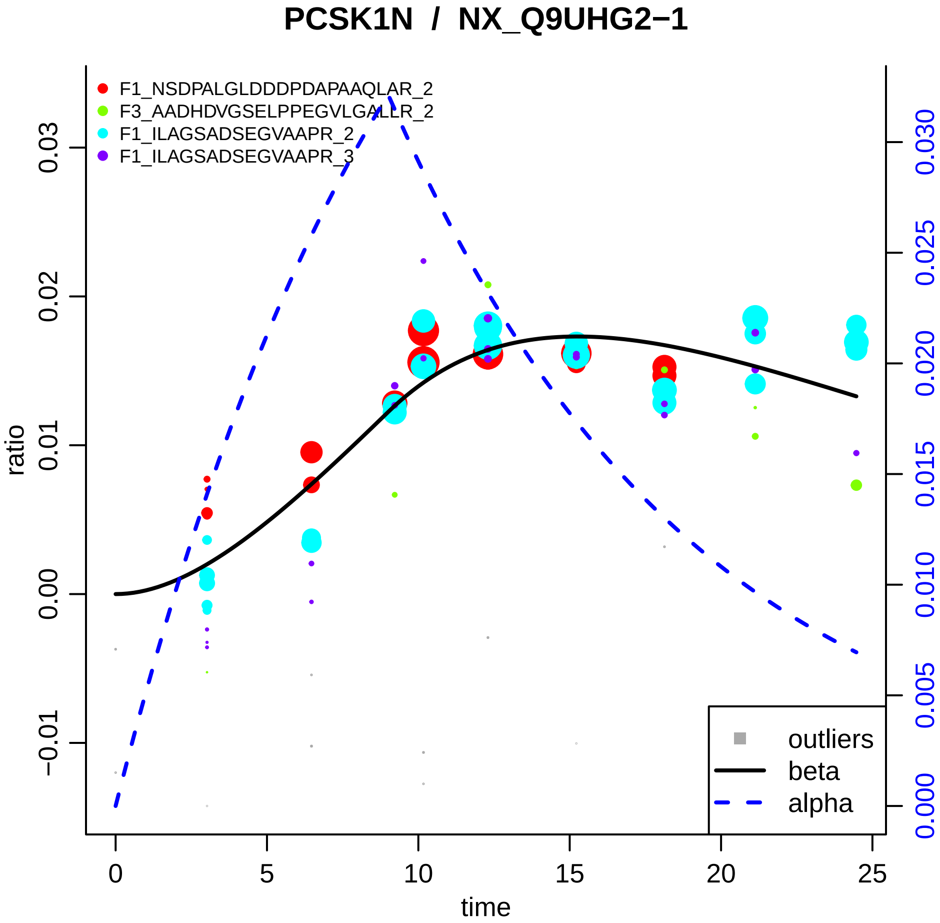

PCSK1N / NX_Q9UHG2−1
F1_NSDPALGLDDDPDAPAAQLAR_2
F3_AADHDVGSELPPEGVLGALLR_2
F1_ILAGSADSEGVAAPR_2
F1_ILAGSADSEGVAAPR_3
ratio
time
outliers
beta
alpha
−0.01
0.00
0.01
0.02
0.03
0
5
10
15
20
25
0.000
0.005
0.010
0.015
0.020
0.025
0.030

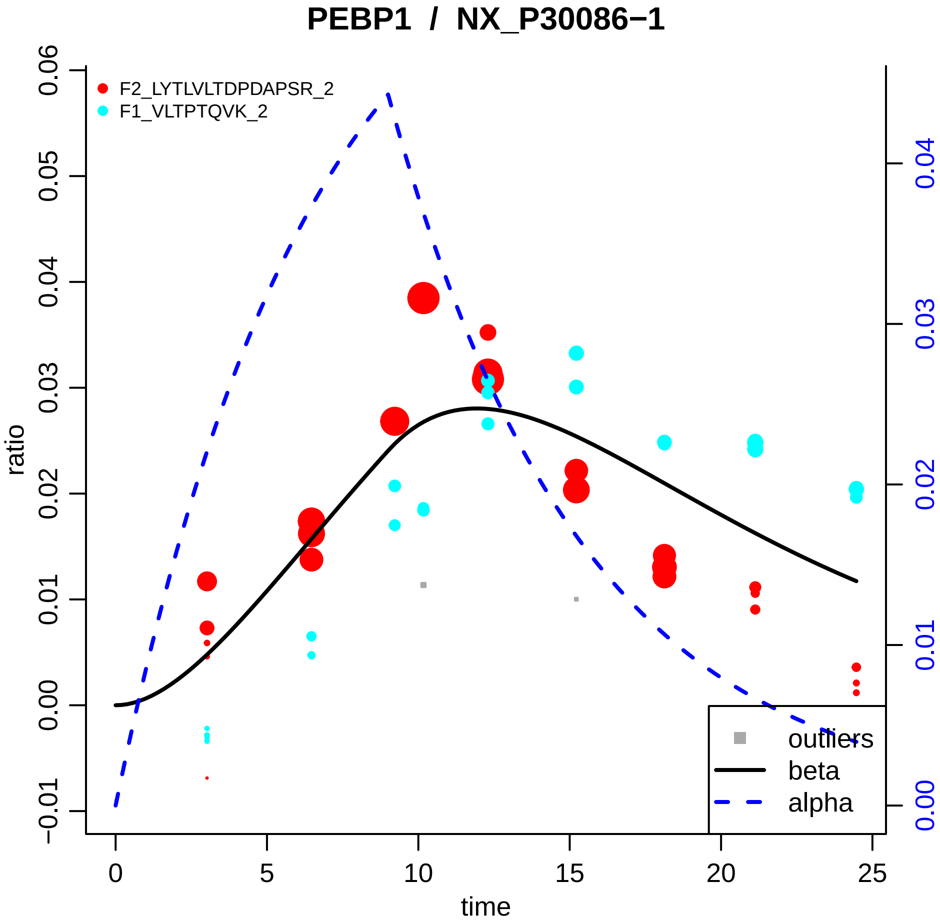

PEBP1 / NX_P30086−1
F2_LYTLVLTDPDAPSR_2
F1_VLTPTQVK_2
ratio
time
outliers
beta
alpha
0.06
0.05
0.04
0.03
0.02
0.01
0.00
−0.01
0
5
10
15
20
25
0.04
0.03
0.02
0.01
0.00

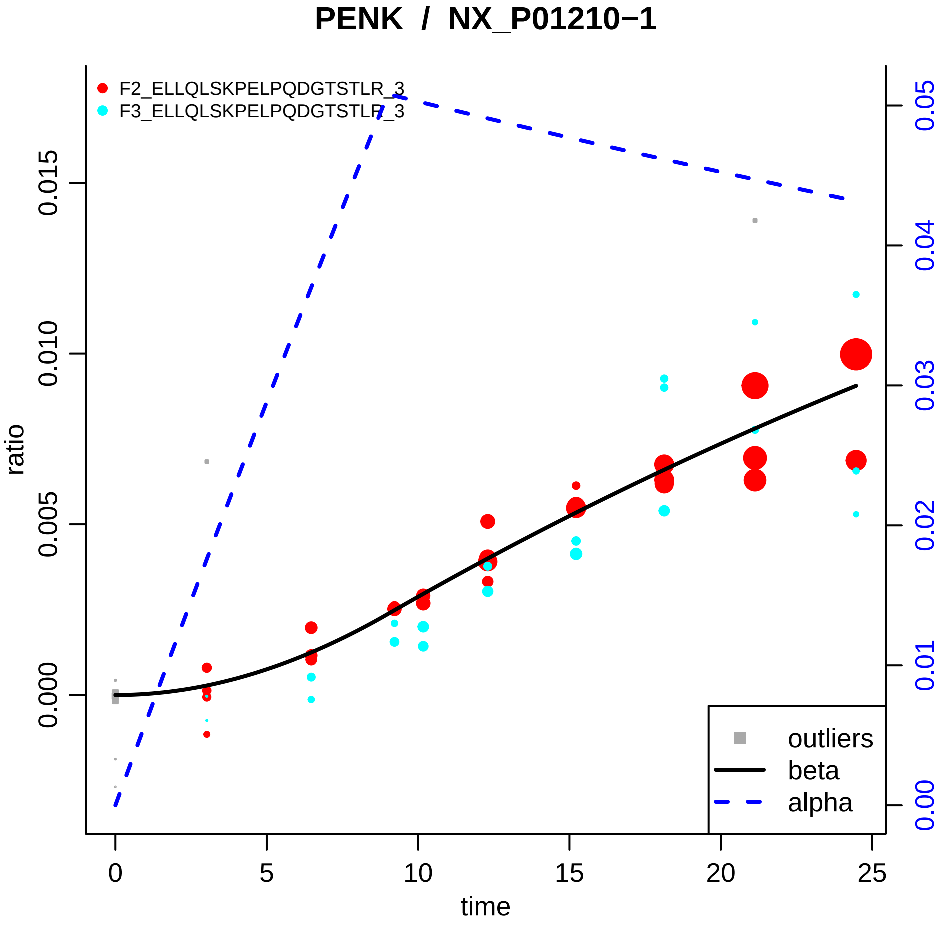

PENK / NX_P01210−1
F2_ELLQLSKPELPQDGTSTLR_3
F3_ELLQLSKPELPQDGTSTLR_3
ratio
time
0.000
0.005
0.010
0.015
0
5
10
15
20
25
0.00
0.01
0.02
0.03
0.04
0.05
outliers
beta
alpha

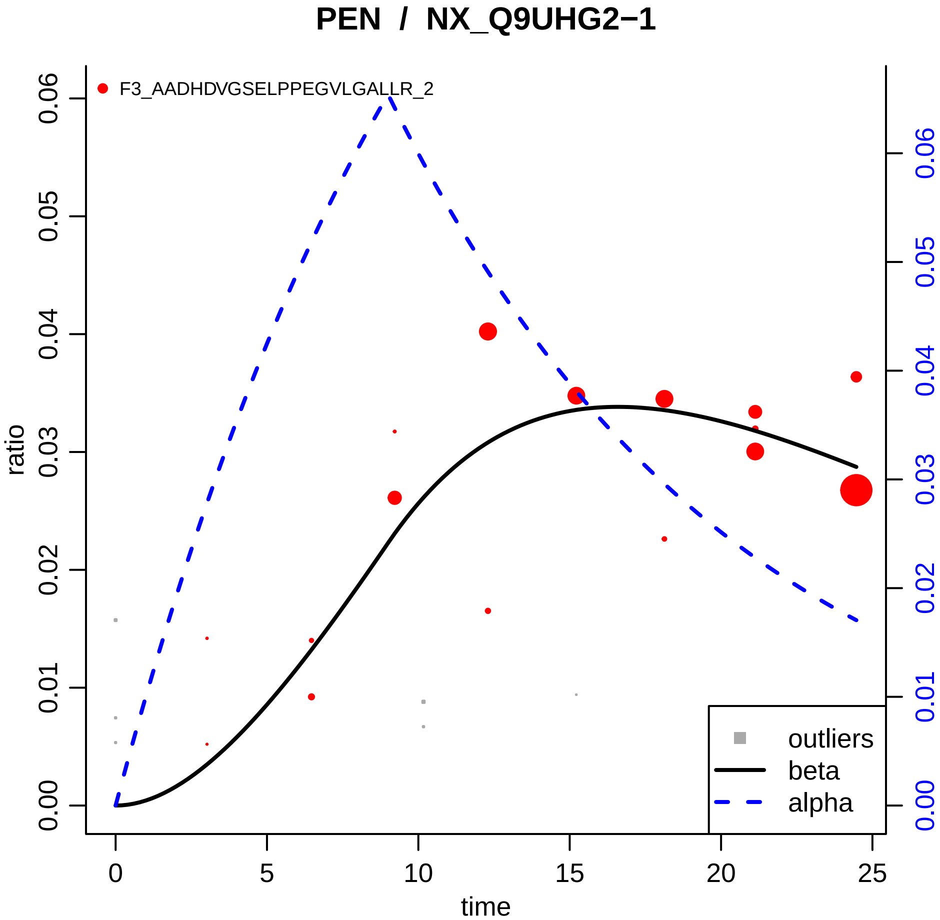

PEN / NX_Q9UHG2−1
F3_AADHDVGSELPPEGVLGALLR_2
ratio
time
0.00
0.01
0.02
0.03
0.04
0.05
0.06
0
5
10
15
20
25
outliers
beta
alpha

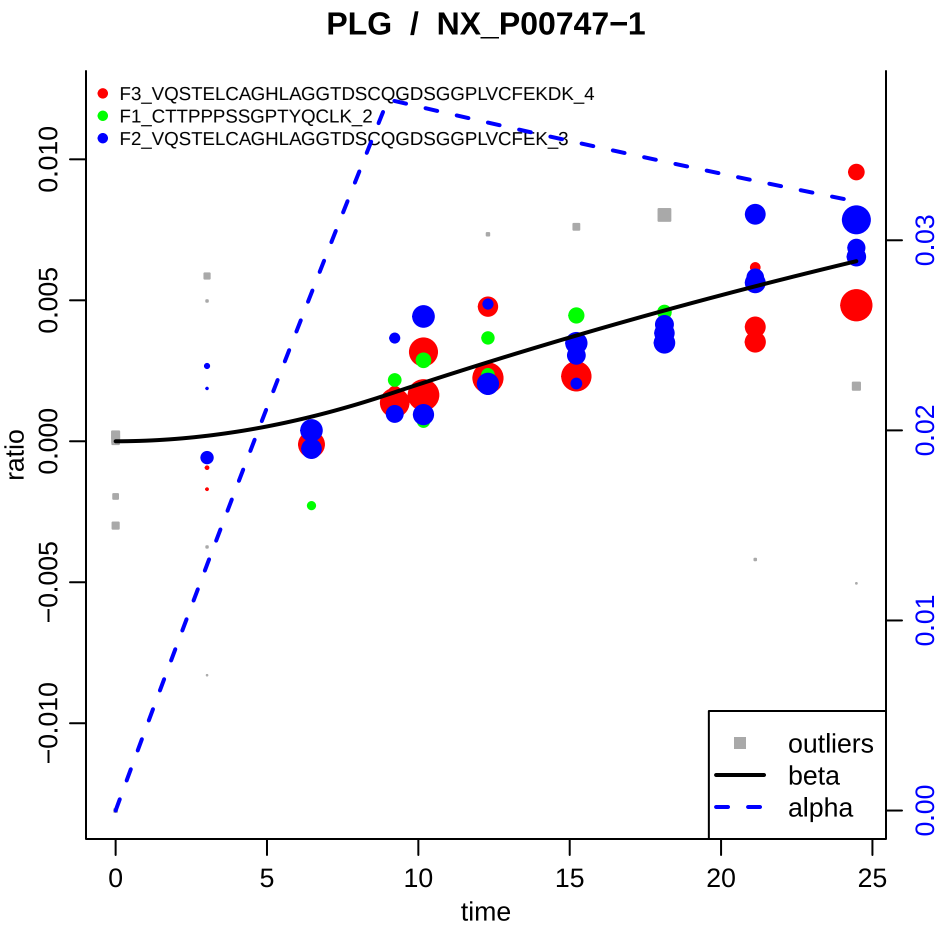

PLG / NX_P00747−1
F3_VQSTELCAGHLAGGTDSCQGDSGGPLVCFEKDK_4
F1_CTTPPPSSGPTYQCLK_2
F2_VQSTELCAGHLAGGTDSCQGDSGGPLVCFEK_3
ratio
time
outliers
beta
alpha
0.010
0.005
0.000
−0.005
−0.010
0
5
10
15
20
25
0.00
0.01
0.02
0.03

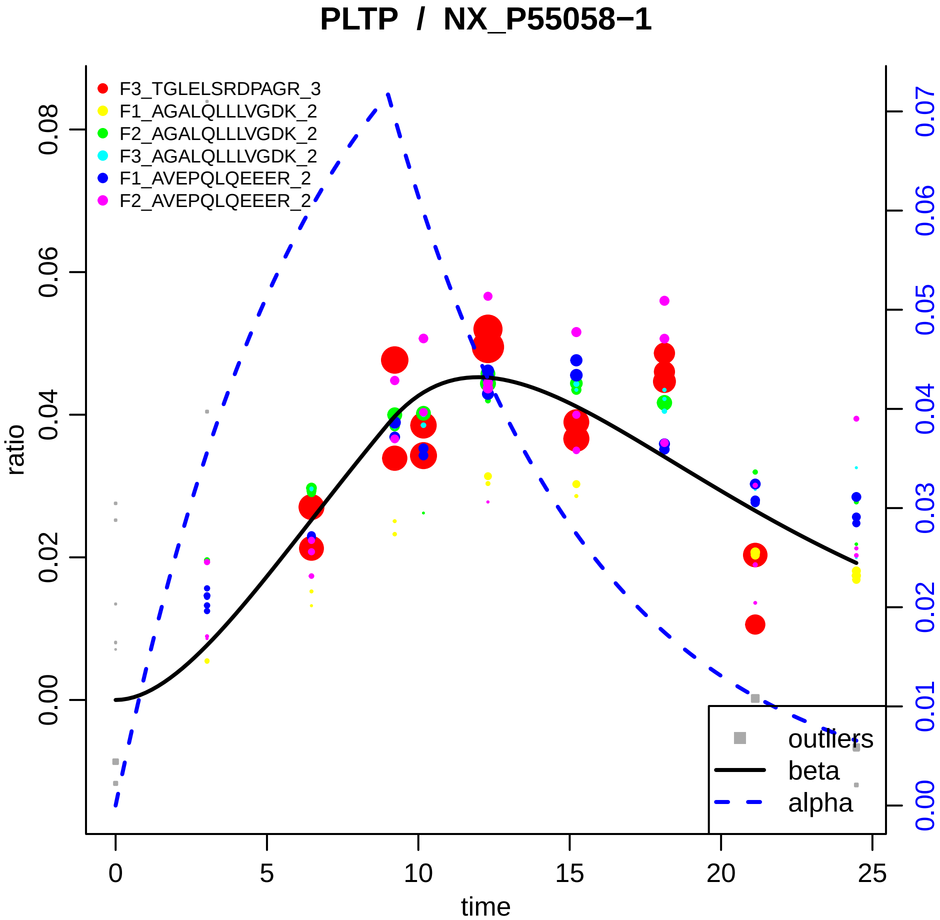

PLTP / NX_P55058−1
F3_TGLELSRDPAGR_3
F1_AGALQLLLVGDK_2
F2_AGALQLLLVGDK_2
F3_AGALQLLLVGDK_2
F1_AVEPQLQEEER_2
F2_AVEPQLQEEER_2
outliers
beta
alpha
ratio
time
0.00
0.02
0.04
0.06
0.08
0.00
0.01
0.02
0.03
0.04
0.05
0.06
0.07
0
5
10
15
20
25

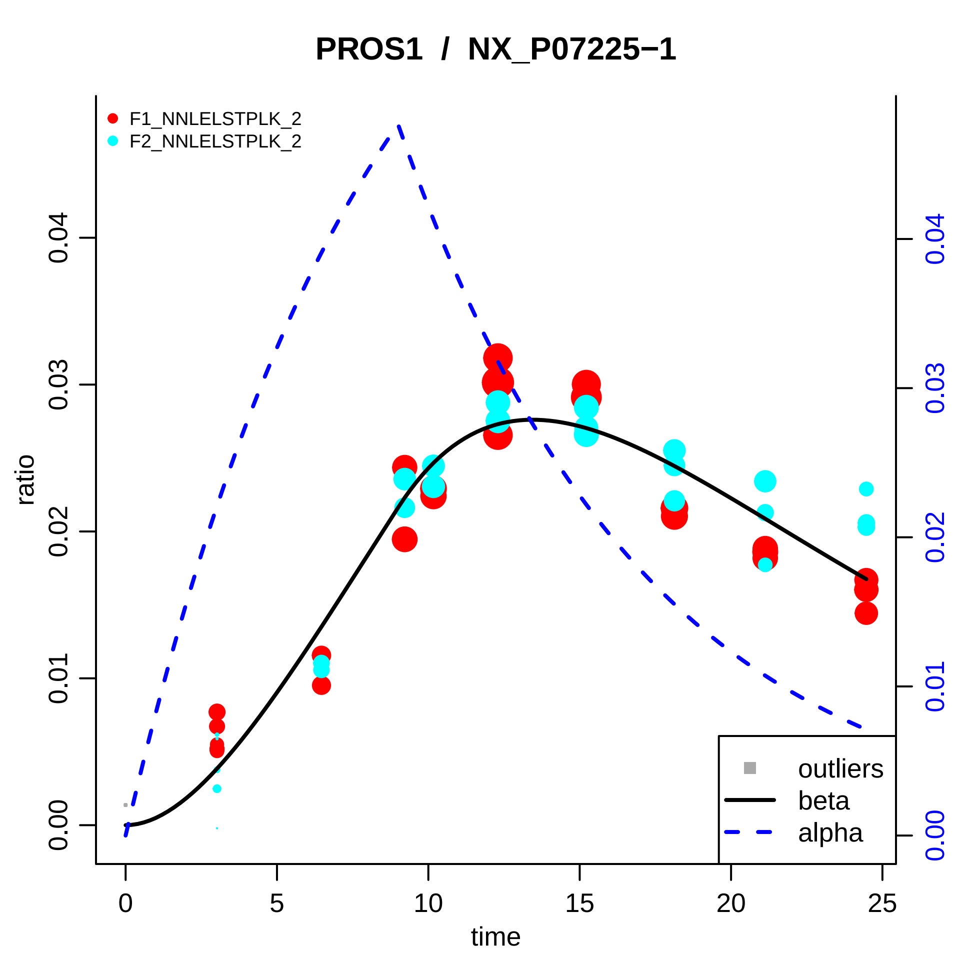

PROS1 / NX_P07225−1
F1_NNLELSTPLK_2
F2_NNLELSTPLK_2
outliers
beta
alpha
ratio
time
0.00
0.01
0.02
0.03
0.04
0
5
10
15
20
25

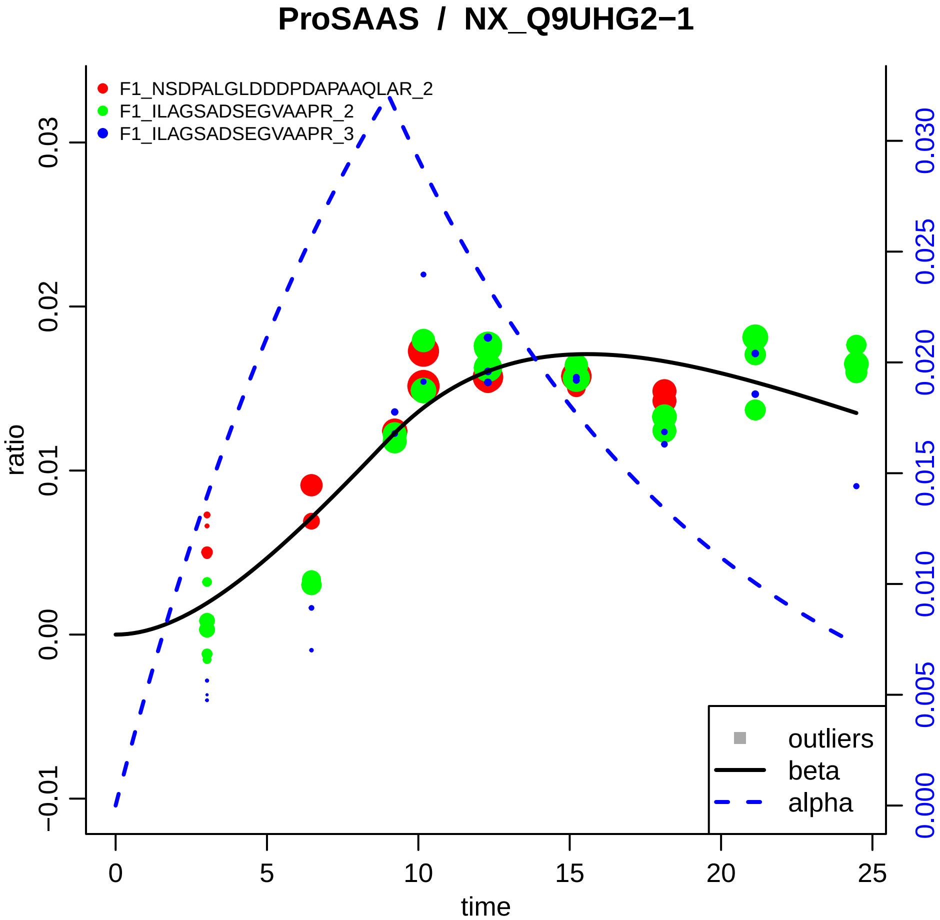

ProSAAS / NX_Q9UHG2−1
F1_NSDPALGLDDDPDAPAAQLAR_2
F1_ILAGSADSEGVAAPR_2
F1_ILAGSADSEGVAAPR_3
ratio
time
outliers
beta
alpha
−0.01
0.00
0.01
0.02
0.03
0
5
10
15
20
25
0.000
0.005
0.010
0.015
0.020
0.025
0.030

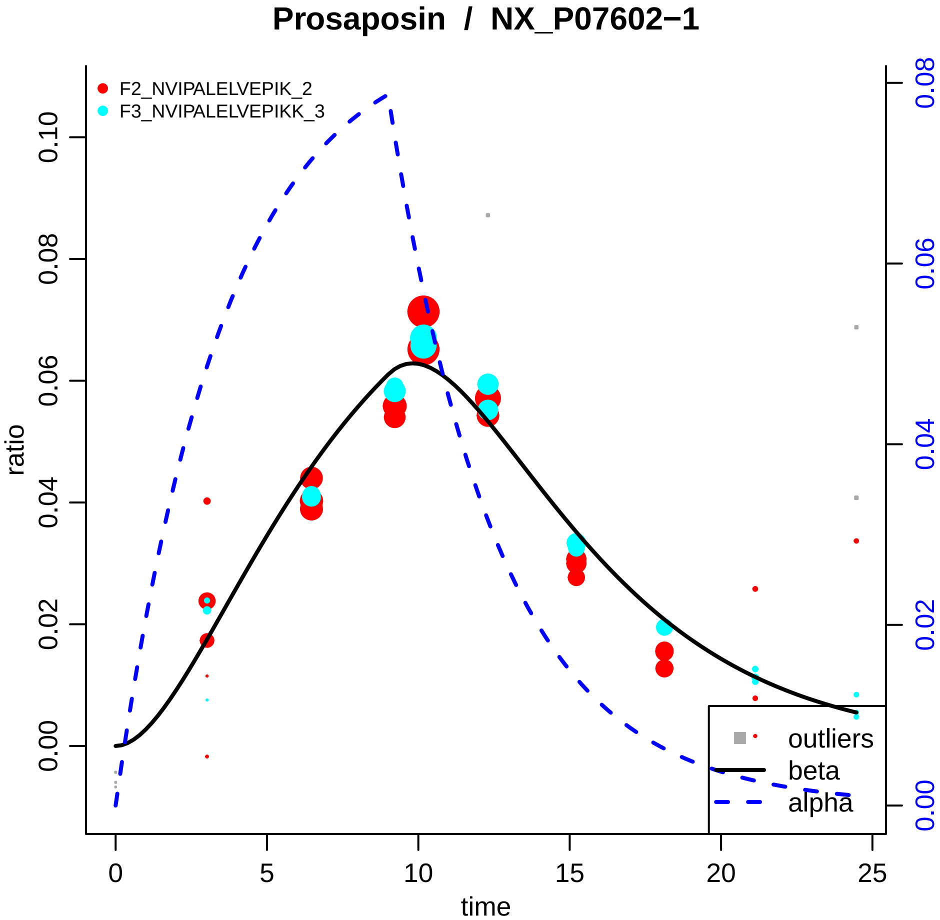

Prosaposin / NX_P07602−1
F2_NVIPALELVEPIK_2
F3_NVIPALELVEPIKK_3
ratio
time
0.00
0.02
0.04
0.06
0.08
0.10
0
5
10
15
20
25
0.00
0.02
0.04
0.06
0.08
outliers
beta
alpha

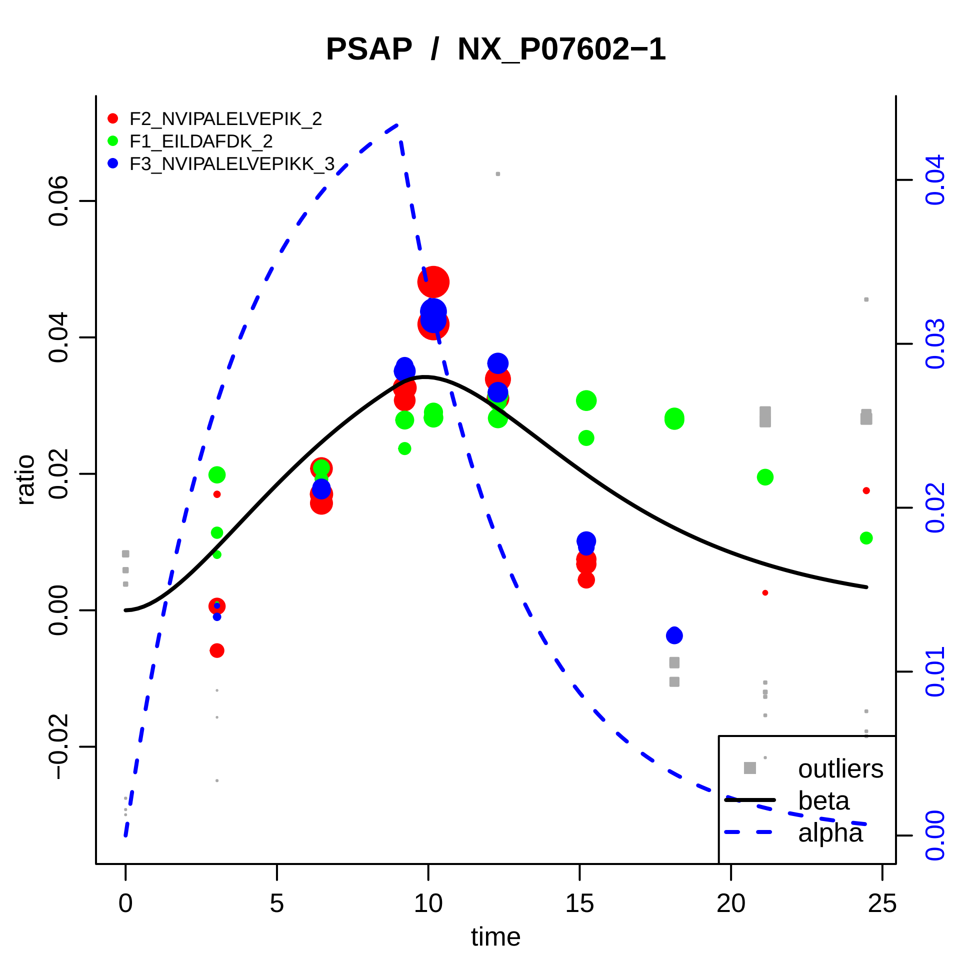

PSAP / NX_P07602−1
F2_NVIPALELVEPIK_2
F1_EILDAFDK_2
F3_NVIPALELVEPIKK_3
ratio
time
−0.02
0.00
0.02
0.04
0.06
0
5
10
15
20
25
0.00
0.01
0.02
0.03
0.04
outliers
beta
alpha

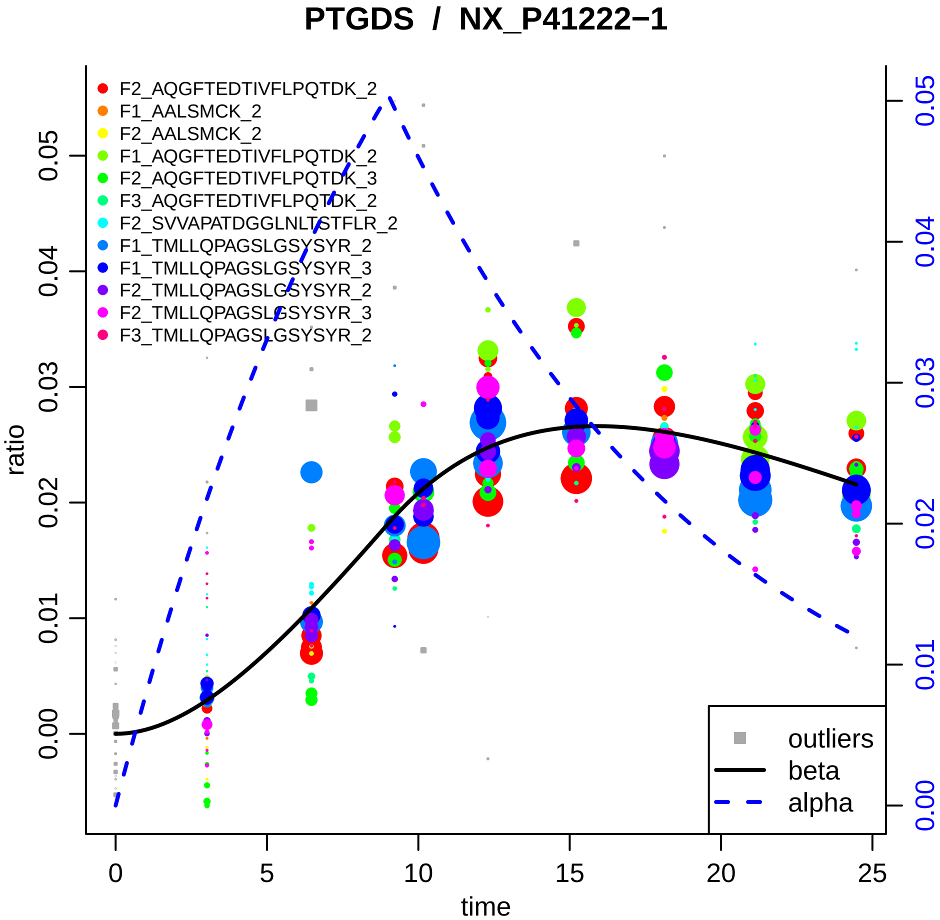

PTGDS / NX_P41222−1
F2_AQGFTEDTIVFLPQTDK_2
F1_AALSMCK_2
F2_AALSMCK_2
F1_AQGFTEDTIVFLPQTDK_2
F2_AQGFTEDTIVFLPQTDK_3
F3_AQGFTEDTIVFLPQTDK_2
F2_SVVAPATDGGLNLTSTFLR_2
F1_TMLLQPAGSLGSYSYR_2
F1_TMLLQPAGSLGSYSYR_3
F2_TMLLQPAGSLGSYSYR_2
F2_TMLLQPAGSLGSYSYR_3
F3_TMLLQPAGSLGSYSYR_2
outliers
beta
alpha
ratio
time
0.00
0.01
0.02
0.03
0.04
0.05
0
5
10
15
20
25

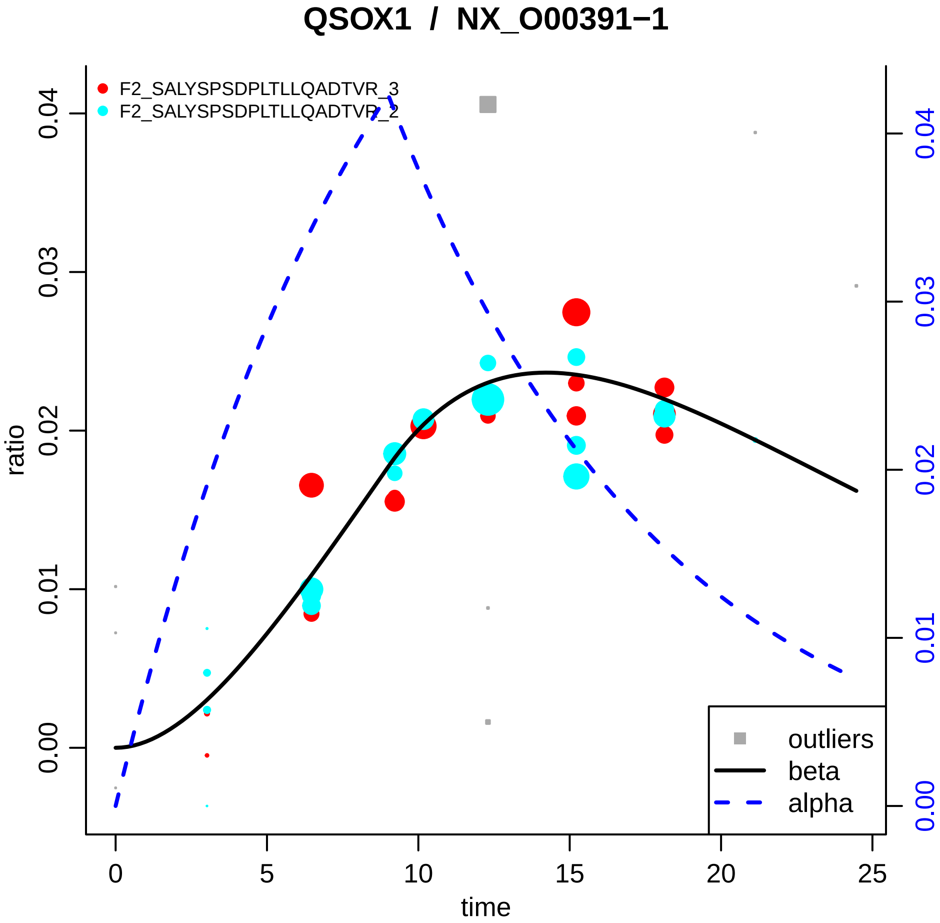

QSOX1 / NX_O00391−1
F2_SALYSPSDPLTLLQADTVR_3
F2_SALYSPSDPLTLLQADTVR_2
ratio
time
0.00
0.01
0.02
0.03
0.04
0
5
10
15
20
25
outliers
beta
alpha

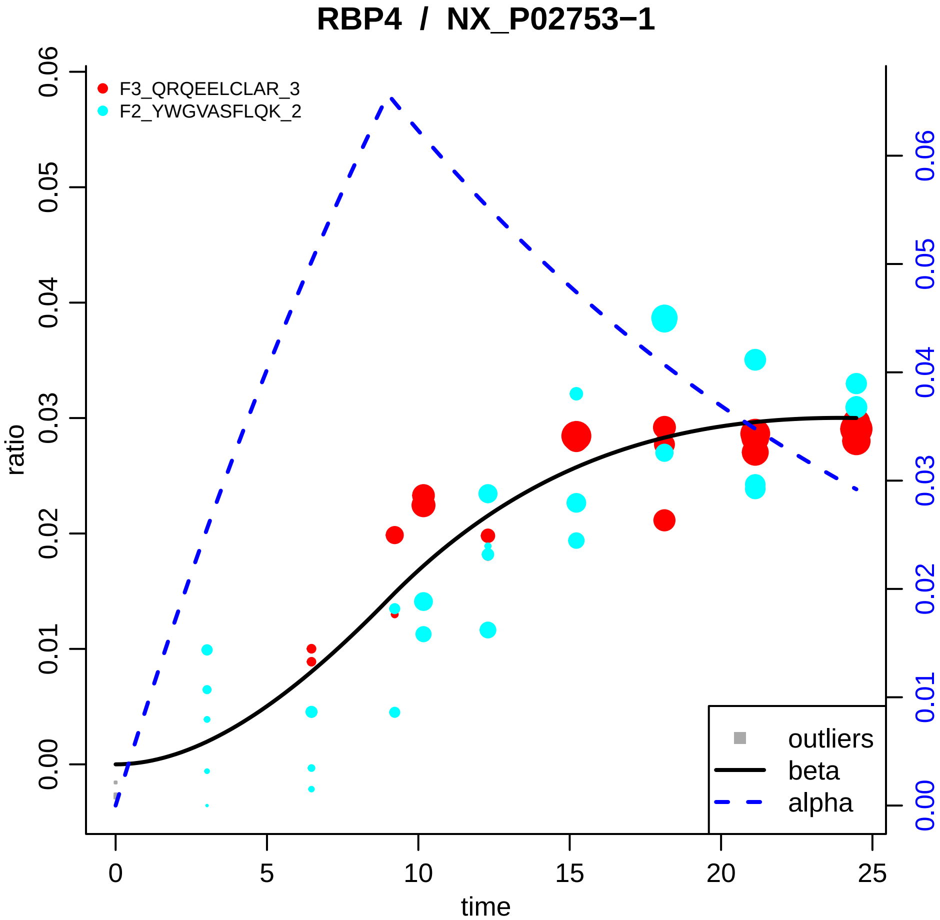

RBP4 / NX_P02753−1
F3_QRQEELCLAR_3
F2_YWGVASFLQK_2
ratio
time
outliers
beta
alpha
0.00
0.01
0.02
0.03
0.04
0.05
0.06
0
5
10
15
20
25

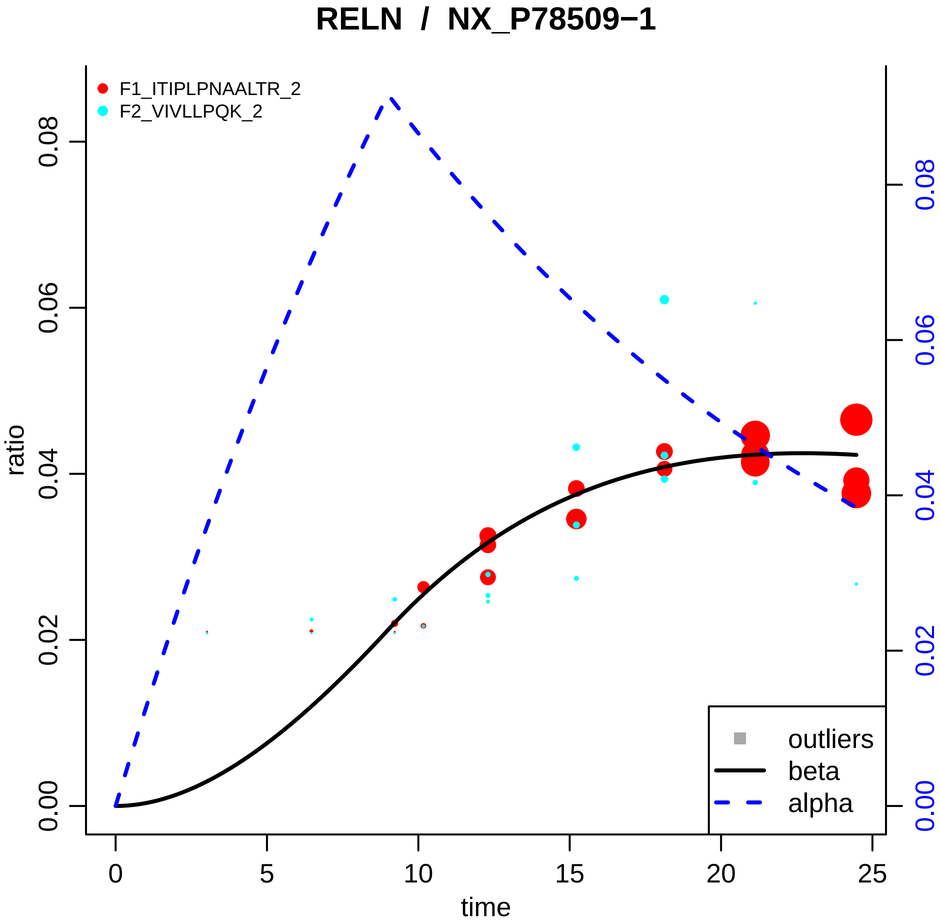
RELN / NX_P78509−1
F1_ITIPLPNAALTR_2
F2_VIVLLPQK_2
ratio
time
0.00
0.02
0.04
0.06
0.08
0
5
10
15
20
25
outliers
beta
alpha

### RFC4 / NX_P35249−1

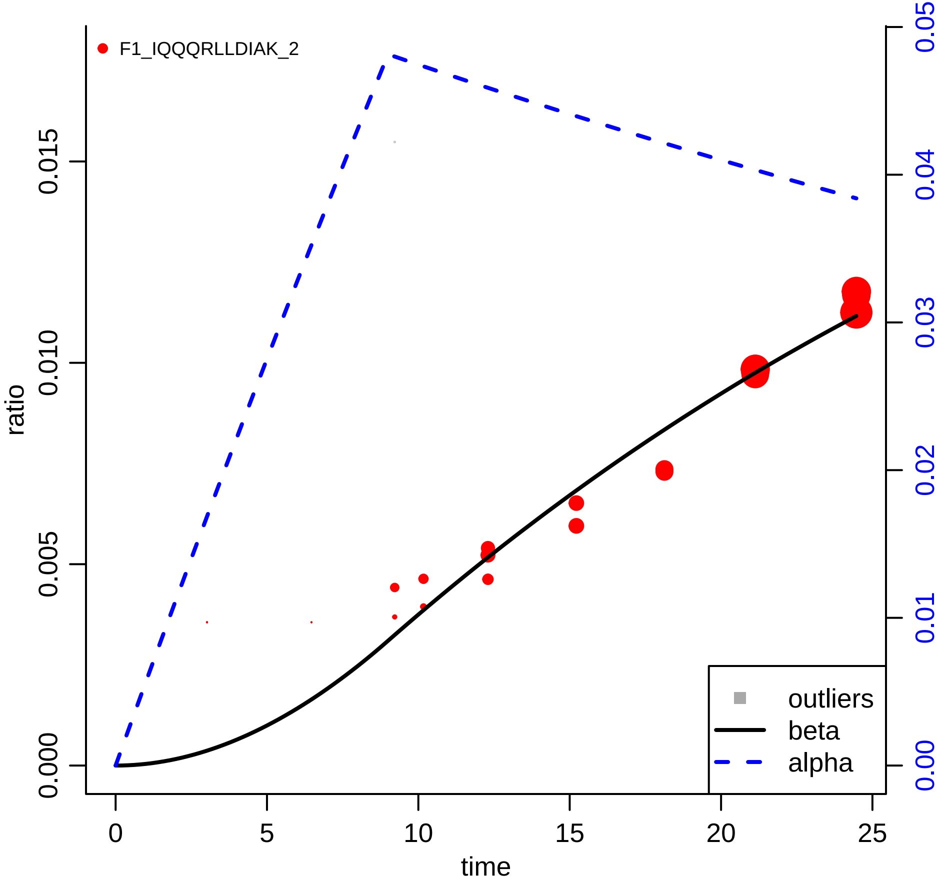

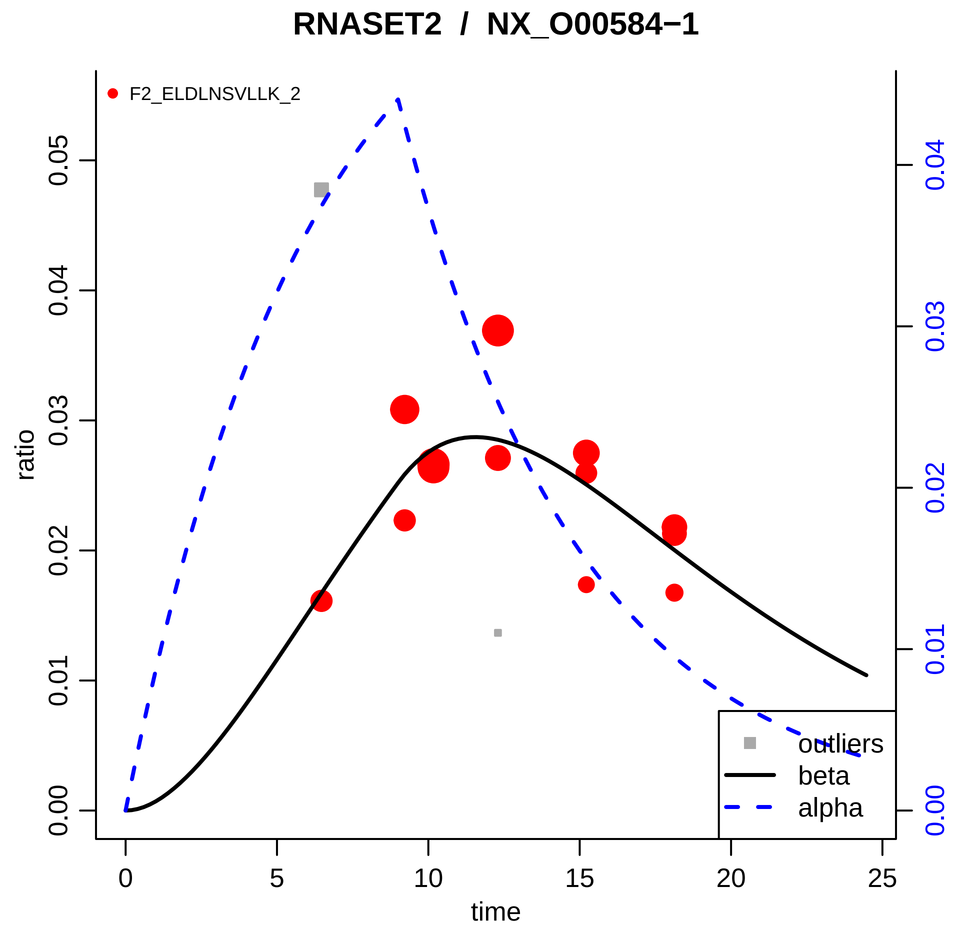
RNASET2 / NX_O00584−1
F2_ELDLNSVLLK_2
ratio
time
0.00
0.01
0.02
0.03
0.04
0.05
0
5
10
15
20
25
0.00
0.01
0.02
0.03
0.04
outliers
beta
alpha

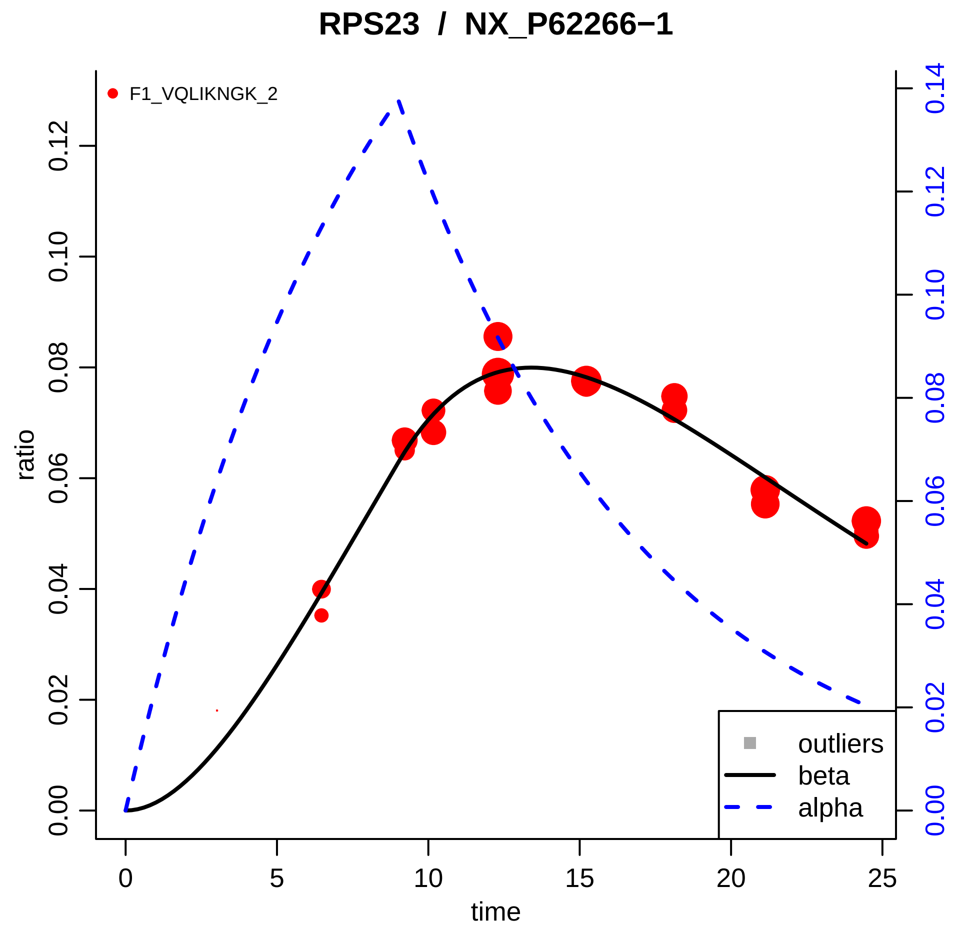

RPS23 / NX_P62266−1
F1_VQLIKNGK_2
ratio
time
0.00
0.02
0.04
0.06
0.08
0.10
0.12
0.14
0
5
10
15
20
25
outliers
beta
alpha

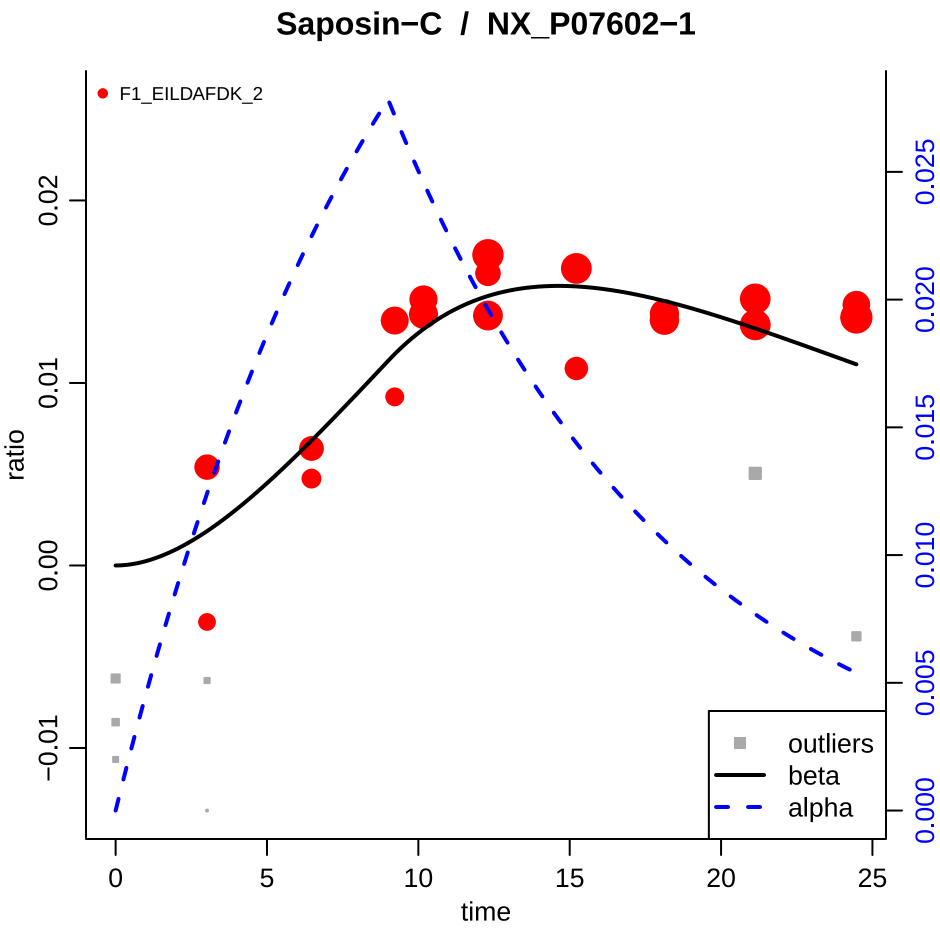

Saposin−C / NX_P07602−1
F1_EILDAFDK_2
ratio
time
0
5
10
15
20
25
−0.01
0.00
0.01
0.02
0.000
0.005
0.010
0.015
0.020
0.025
outliers
beta
alpha

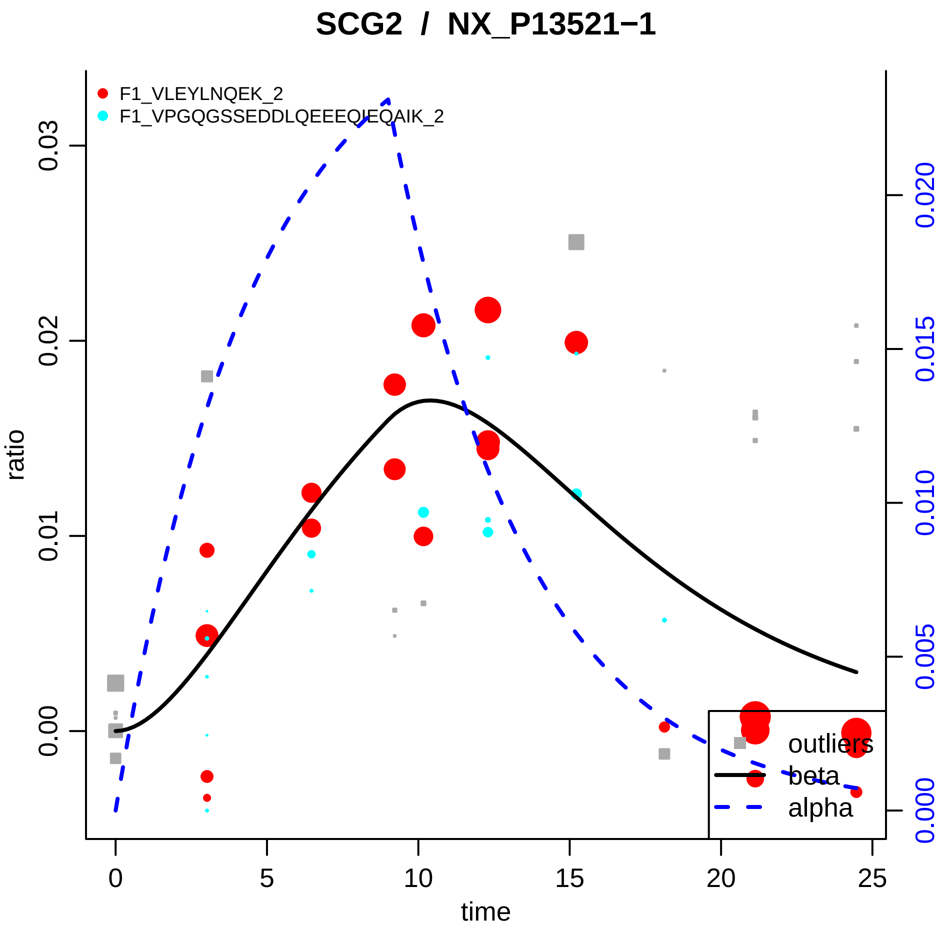

SCG2 / NX_P13521−1
F1_VLEYLNQEK_2
F1_VPGQGSSEDDLQEEEQIEQAIK_2
ratio
0.00
0.01
0.02
0.03
0.000
0.005
0.010
0.015
0.020
time
0
5
10
15
20
25
outliers
beta
alpha

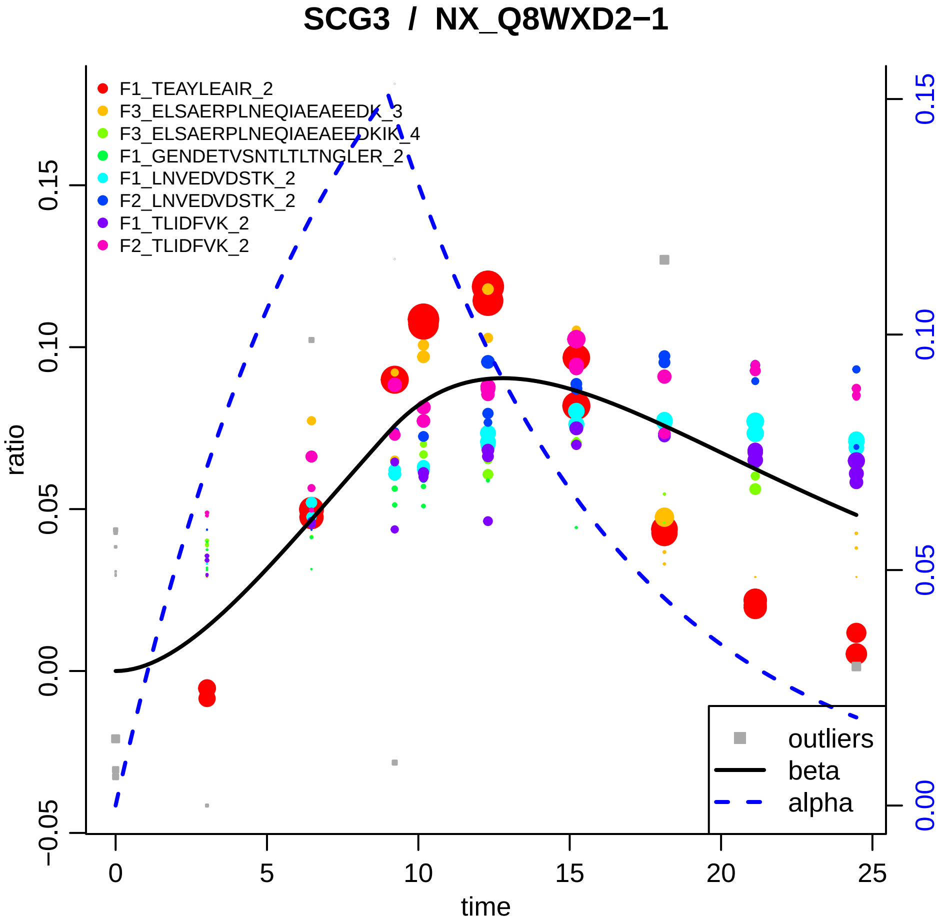

SCG3 / NX_Q8WXD2−1
F1_TEAYLEAIR_2
F3_ELSAERPLNEQIAEAEEDK_3
F3_ELSAERPLNEQIAEAEEDKIK_4
F1_GENDETVSNTLTLTNGLER_2
F1_LNVEDVDSTK_2
F2_LNVEDVDSTK_2
F1_TLIDFVK_2
F2_TLIDFVK_2
outliers
beta
alpha
ratio
time
−0.05
0.00
0.05
0.10
0.15
0
5
10
15
20
25
0.00
0.05
0.10
0.15

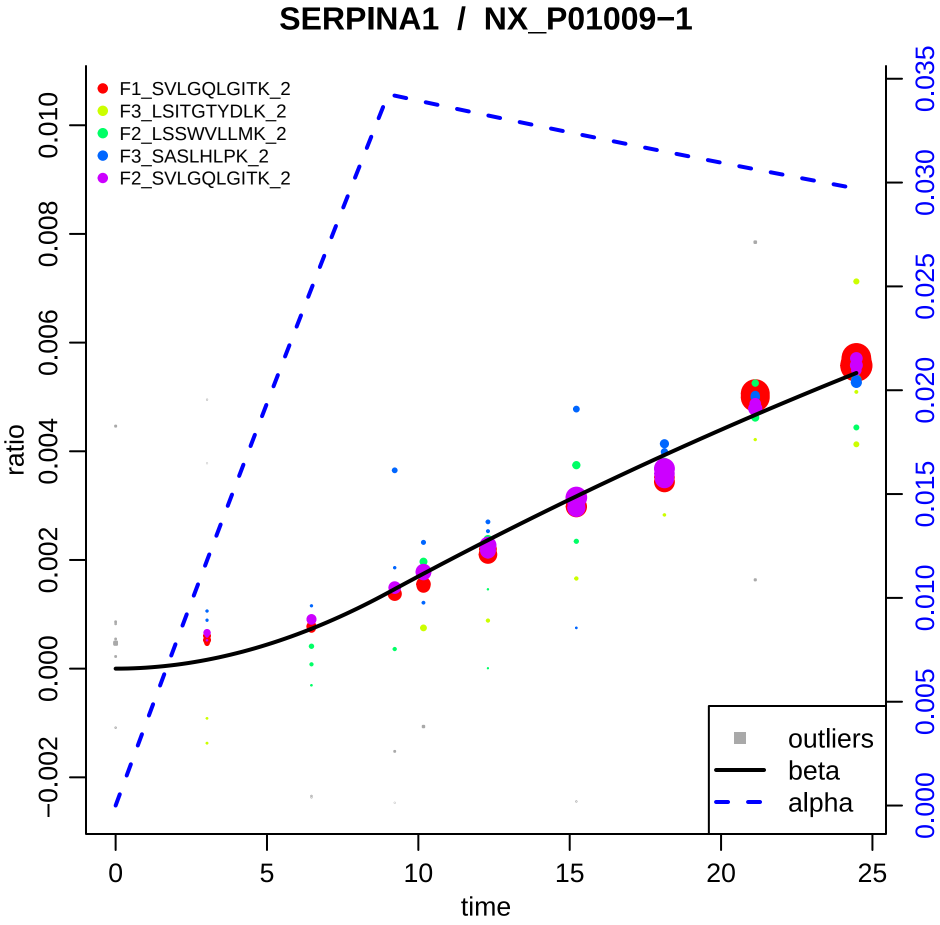

SERPINA1 / NX_P01009−1
F1_SVLGQLGITK_2
F3_LSITGTYDLK_2
F2_LSSWVLLMK_2
F3_SASLHLPK_2
F2_SVLGQLGITK_2
outliers
beta
alpha
ratio
time
0.010
0.008
0.006
0.004
0.002
0.000
−0.002
0
5
10
15
20
25
0.035
0.030
0.025
0.020
0.015
0.010
0.005
0.000

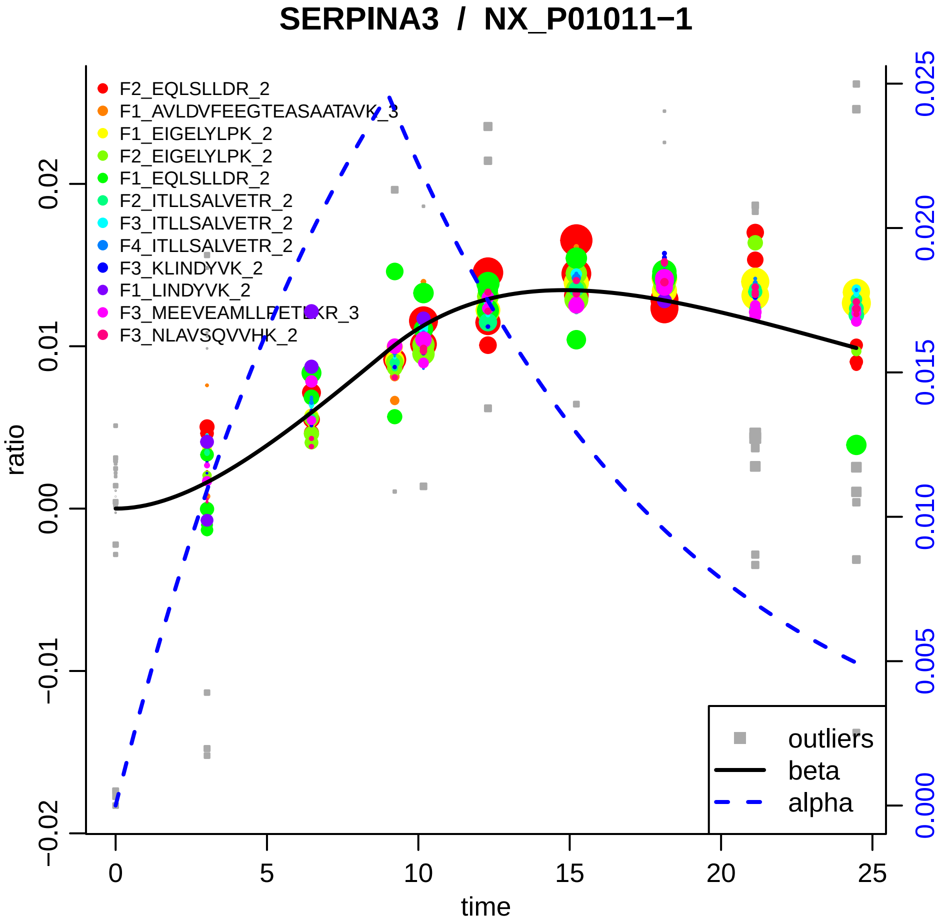
SERPINA3 / NX_P01011−1
F2_EQLSLLDR_2
F1_AVLDVFEEGTEASAATAVK_3
F1_EIGELYLPK_2
F2_EIGELYLPK_2
F1_EQLSLLDR_2
F2_ITLLSALVETR_2
F3_ITLLSALVETR_2
F4_ITLLSALVETR_2
F3_KLINDYVK_2
F1_LINDYVK_2
F3_MEEVEAMLLPETLKR_3
F3_NLAVSQVVHK_2
ratio
time
outliers
beta
alpha
0.02
0.01
0.00
−0.01
−0.02
0
5
10
15
20
25
0.025
0.020
0.015
0.010
0.005
0.000

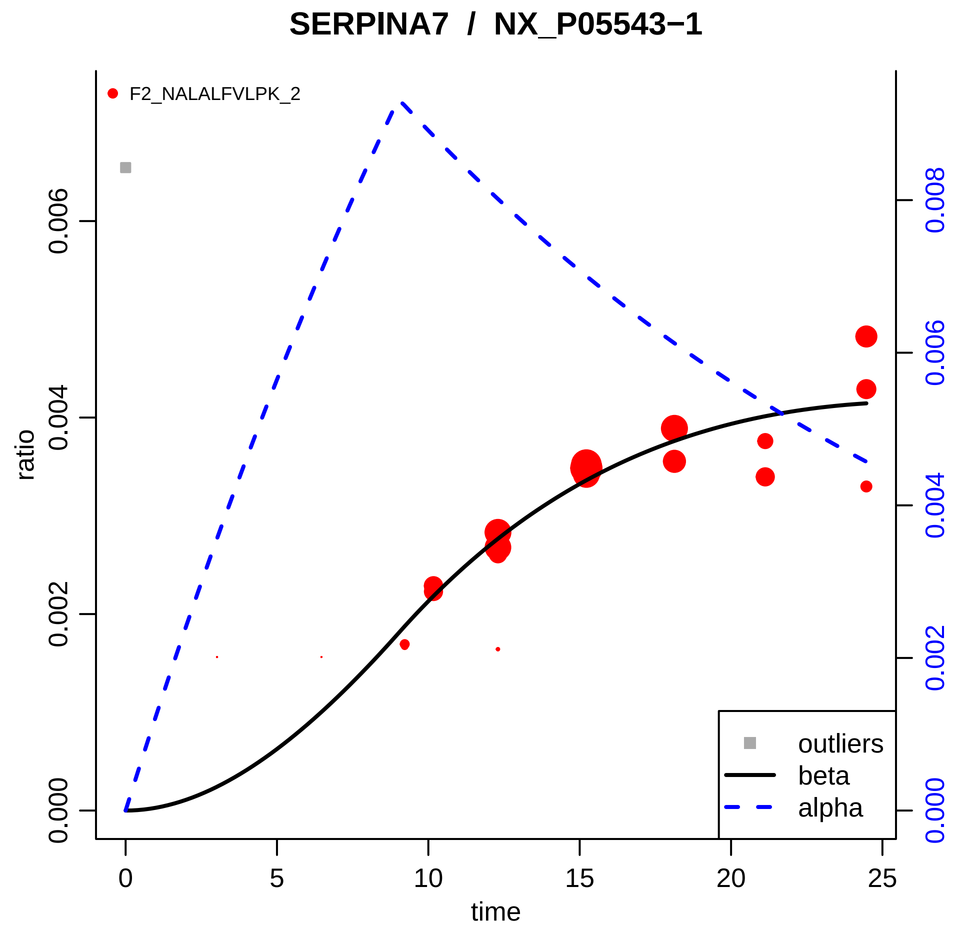

SERPINA7 / NX_P05543−1
F2_NALALFVLPK_2
ratio
0.000
0.002
0.004
0.006
time
0
5
10
15
20
25
0.000
0.002
0.004
0.006
0.008
outliers
beta
alpha

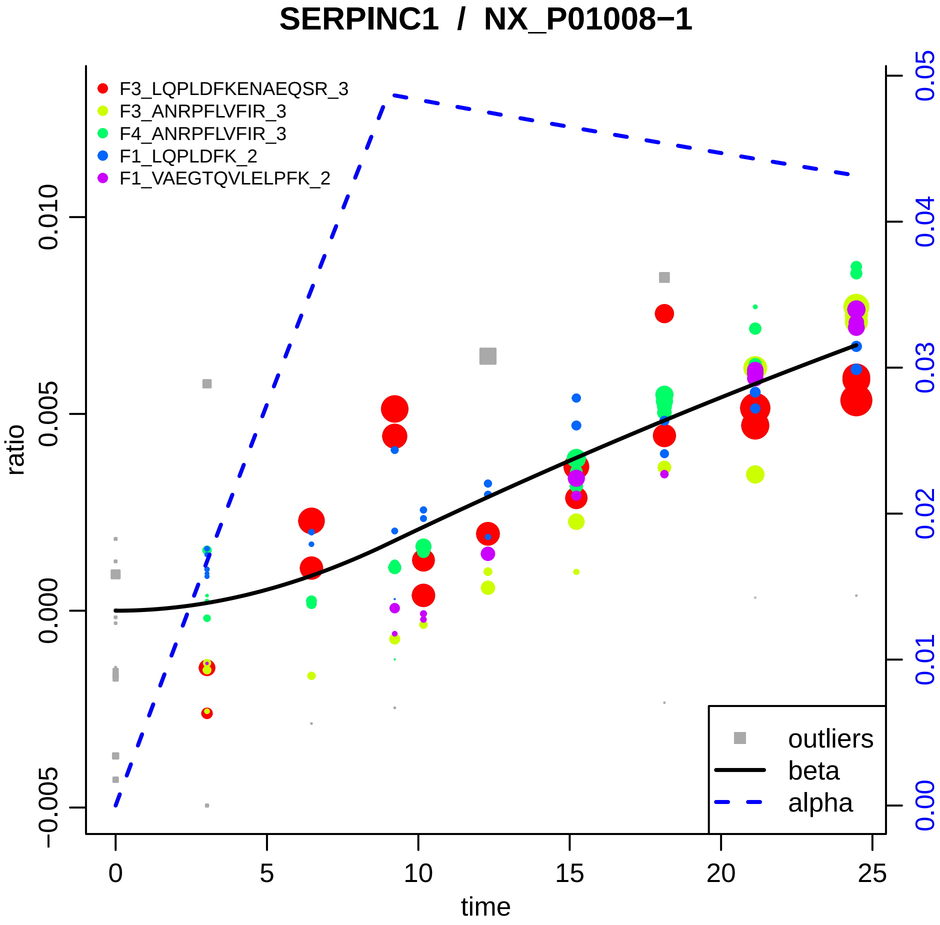

SERPINC1 / NX_P01008−1
F3_LQPLDFKENAEQSR_3
F3_ANRPFLVFIR_3
F4_ANRPFLVFIR_3
F1_LQPLDFK_2
F1_VAEGTQVLELPFK_2
outliers
beta
alpha
ratio
time
0.010
0.005
0.000
−0.005
0
5
10
15
20
25
0.05
0.04
0.03
0.02
0.01
0.00

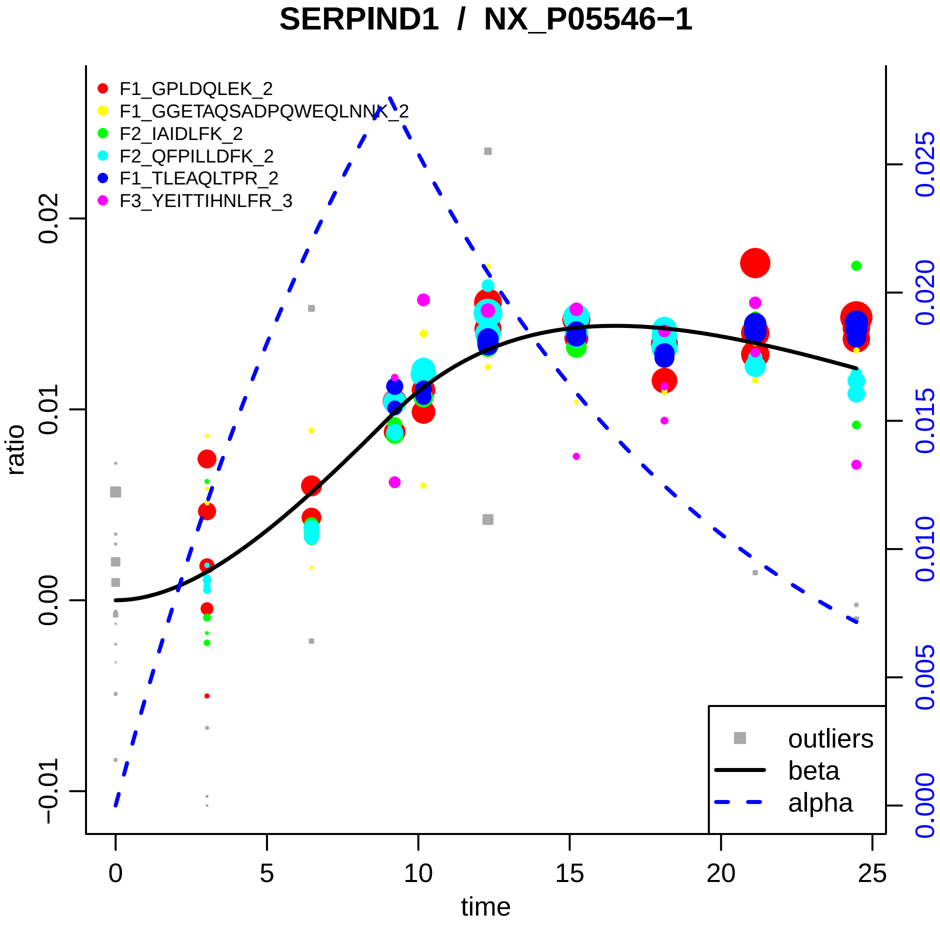

SERPIND1 / NX_P05546−1
F1_GPLDQLEK_2
F1_GGETAQSADPQWEQLNNK_2
F2_IAIDLFK_2
F2_QFPILLDFK_2
F1_TLEAQLTPR_2
F3_YEITTIHNLFR_3
ratio
time
outliers
beta
alpha
0
5
10
15
20
25
−0.01
0.00
0.01
0.02
0.000
0.005
0.010
0.015
0.020
0.025

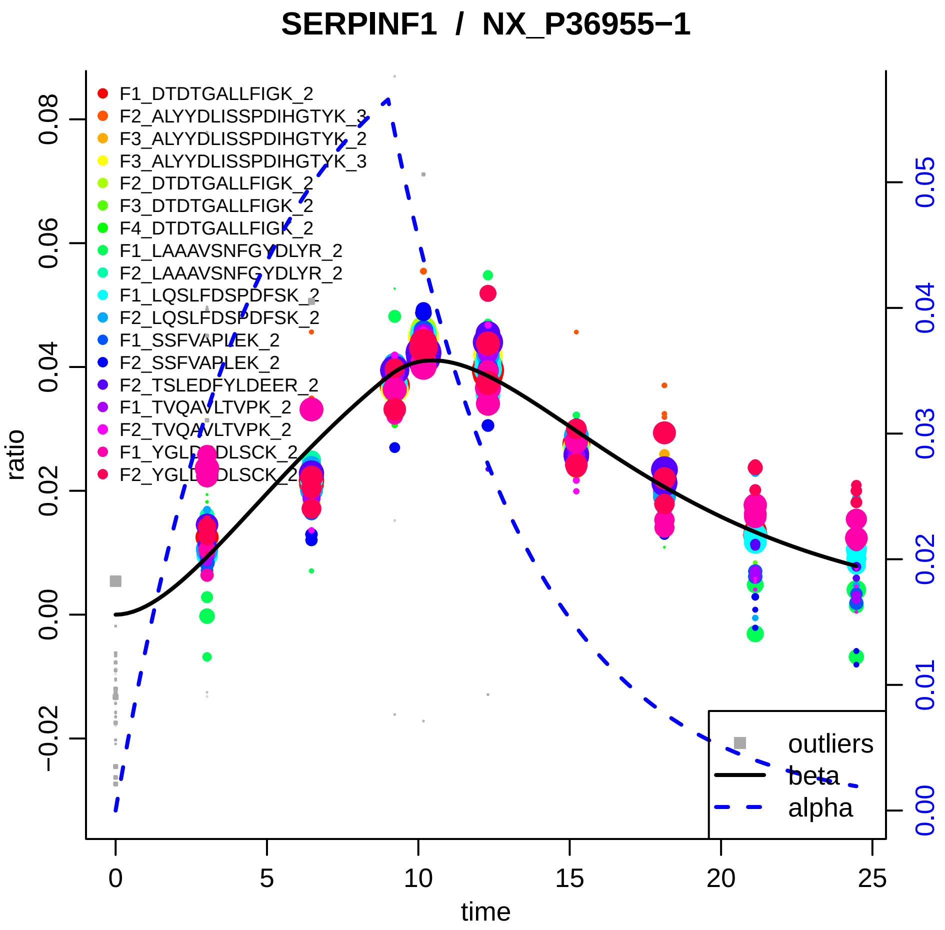
SERPINF1 / NX_P36955−1
F1_DTDTGALLFIGK_2
F2_ALYYDLISSPDIHGTYK_3
F3_ALYYDLISSPDIHGTYK_2
F3_ALYYDLISSPDIHGTYK_3
F2_DTDTGALLFIGK_2
F3_DTDTGALLFIGK_2
F4_DTDTGALLFIGK_2
F1_LAAAVSNFGYDLYR_2
F2_LAAAVSNFGYDLYR_2
F1_LQSLFDSPDFSK_2
F2_LQSLFDSPDFSK_2
F1_SSFVAPLEK_2
F2_SSFVAPLEK_2
F2_TSLEDFYLDEER_2
F1_TVQAVLTVPK_2
F2_TVQAVLTVPK_2
F1_YGLDSDLSCK_2
F2_YGLDSDLSCK_2
outliers
beta
alpha
ratio
time
−0.02
0.00
0.02
0.04
0.06
0.08
0.00
0.01
0.02
0.03
0.04
0.05
0
5
10
15
20
25

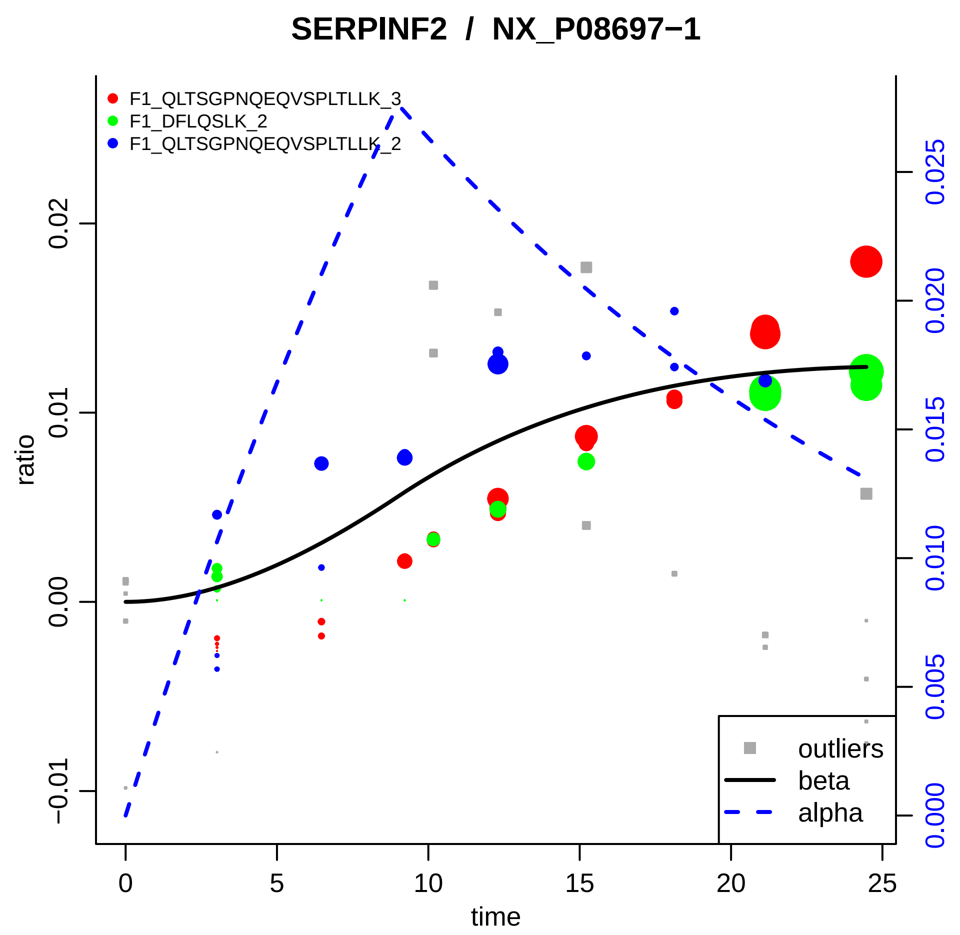
SERPINF2 / NX_P08697−1
F1_QLTSGPNQEQVSPLTLLK_3
F1_DFLQSLK_2
F1_QLTSGPNQEQVSPLTLLK_2
ratio
−0.01
0.00
0.01
0.02
0.000
0.005
0.010
0.015
0.020
0.025
time
0
5
10
15
20
25
outliers
beta
alpha

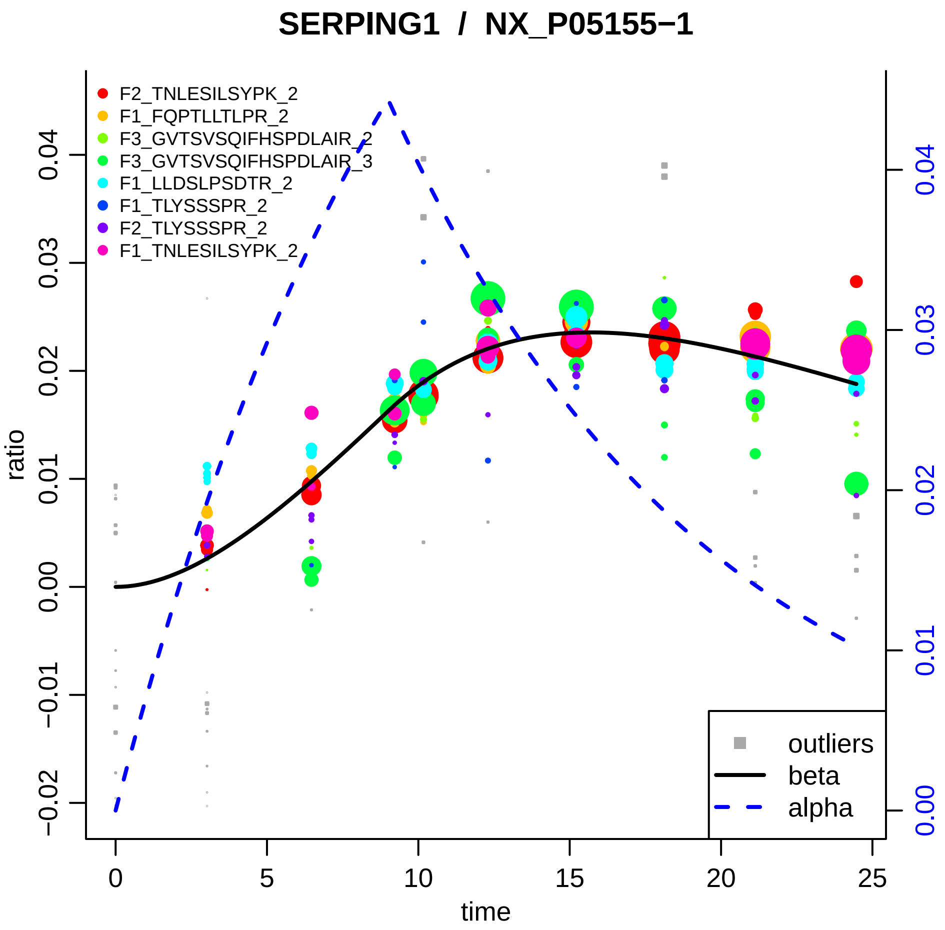
SERPING1 / NX_P05155−1
F2_TNLESILSYPK_2
F1_FQPTLLTLPR_2
F3_GVTSVSQIFHSPDLAIR_2
F3_GVTSVSQIFHSPDLAIR_3
F1_LLDSLPSDTR_2
F1_TLYSSSPR_2
F2_TLYSSSPR_2
F1_TNLESILSYPK_2
ratio
time
outliers
beta
alpha
0.04
0.03
0.02
0.01
0.00
−0.01
−0.02
0
5
10
15
20
25

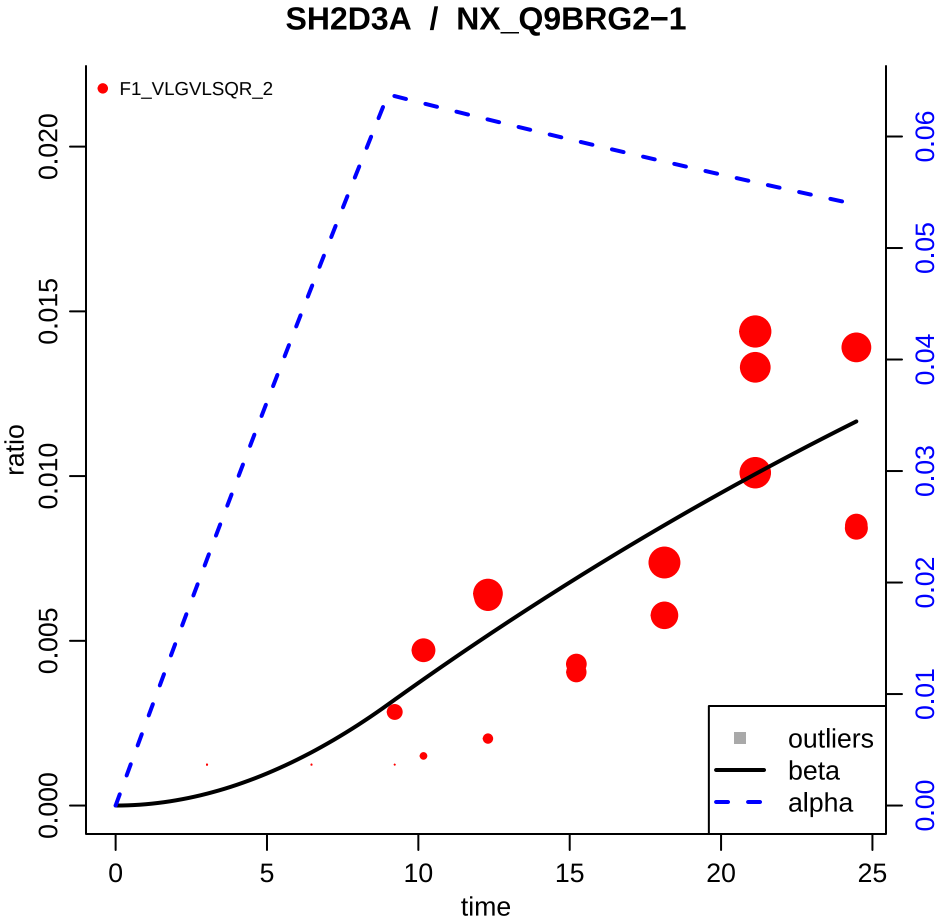

SH2D3A / NX_Q9BRG2−1
F1_VLGVLSQR_2
ratio
time
0.000
0.005
0.010
0.015
0.020
0
5
10
15
20
25
0.00
0.01
0.02
0.03
0.04
0.05
0.06
outliers
beta
alpha

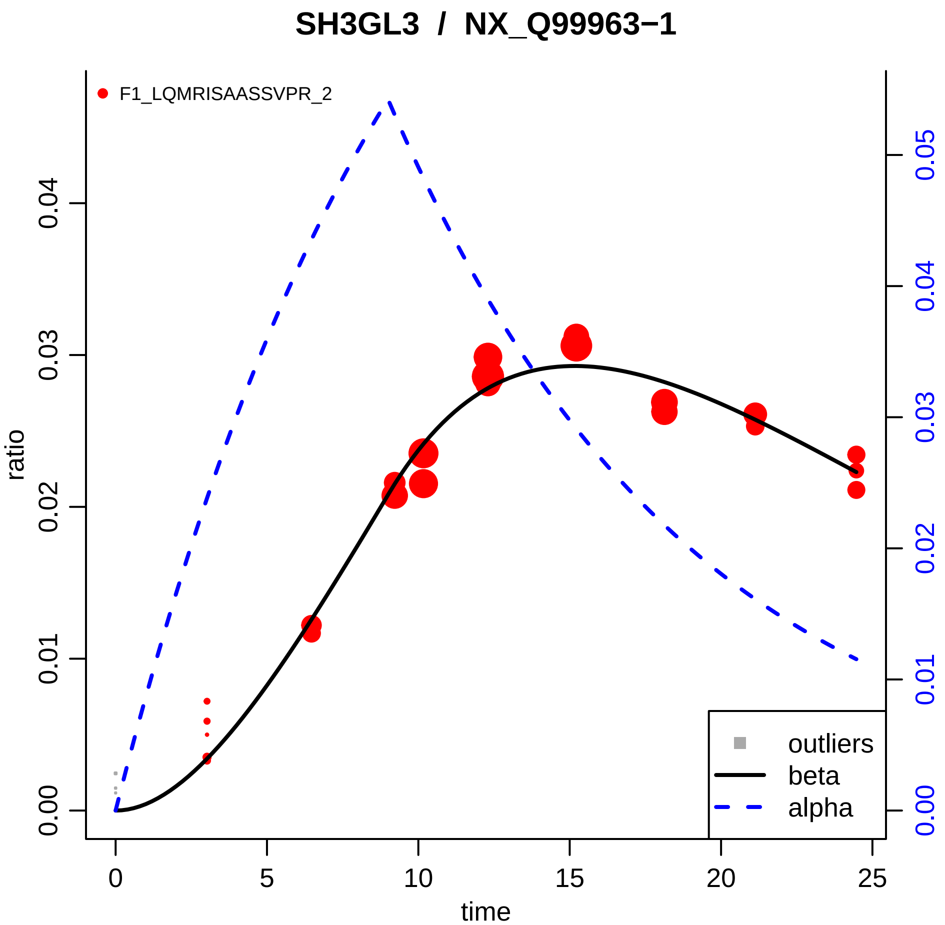

SH3GL3 / NX_Q99963−1
F1_LQMRISAASSVPR_2
ratio
time
0
5
10
15
20
25
0.00
0.01
0.02
0.03
0.04
0.05
outliers
beta
alpha

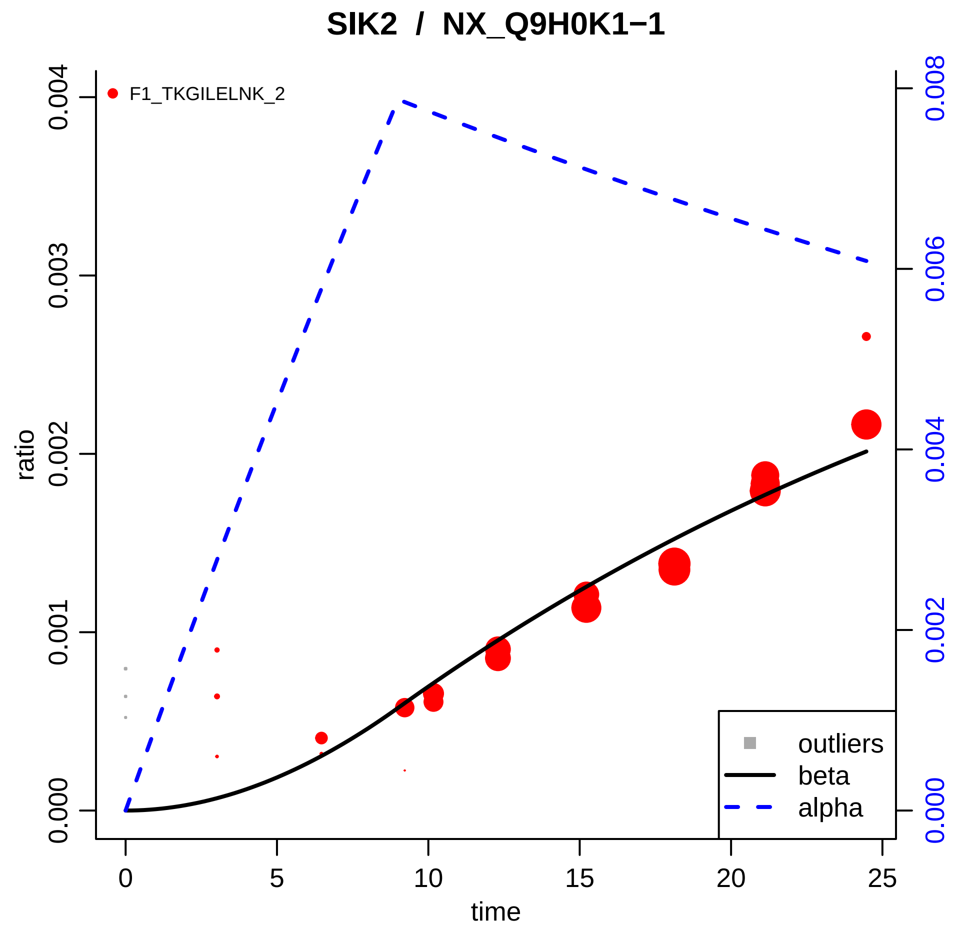

SIK2 / NX_Q9H0K1−1
F1_TKGILELNK_2
ratio
time
0.000
0.001
0.002
0.003
0.004
0
5
10
15
20
25
0.000
0.002
0.004
0.006
0.008
outliers
beta
alpha

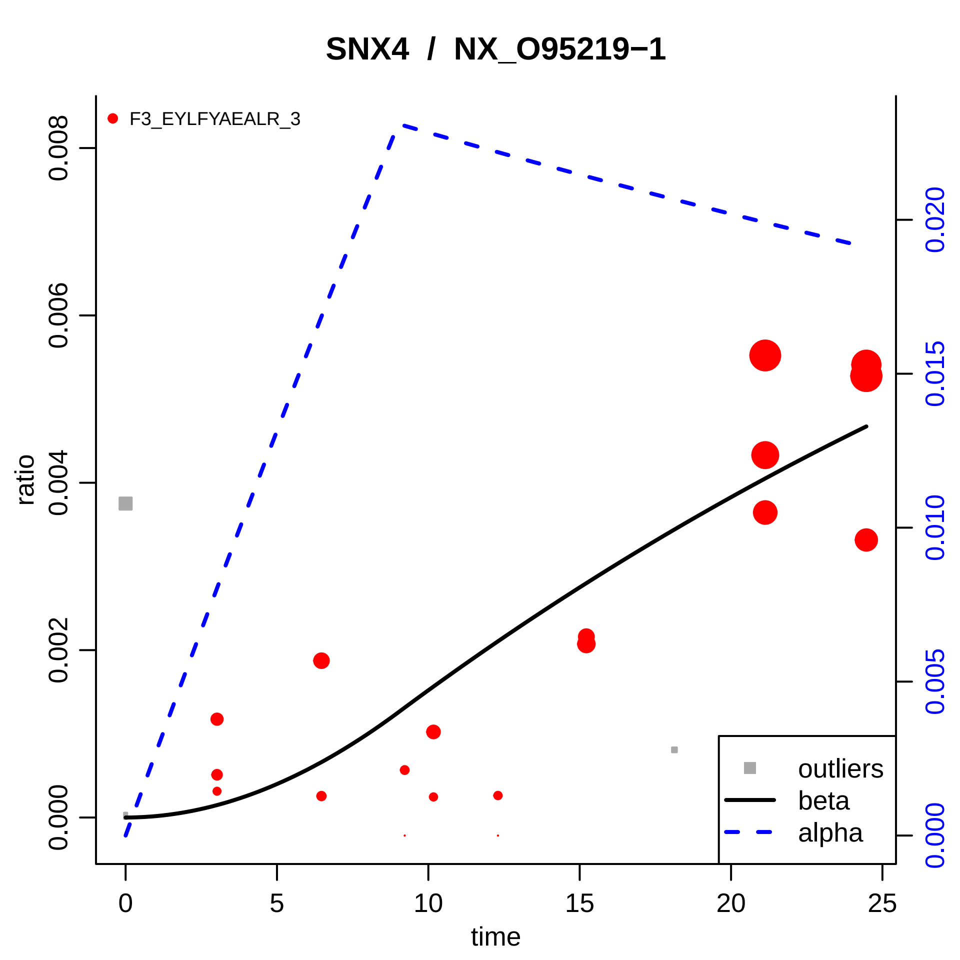
SNX4 / NX_O95219−1
F3_EYLFYAEALR_3
ratio
time
0.000
0.002
0.004
0.006
0.008
0
5
10
15
20
25
0.000
0.005
0.010
0.015
0.020
outliers
beta
alpha

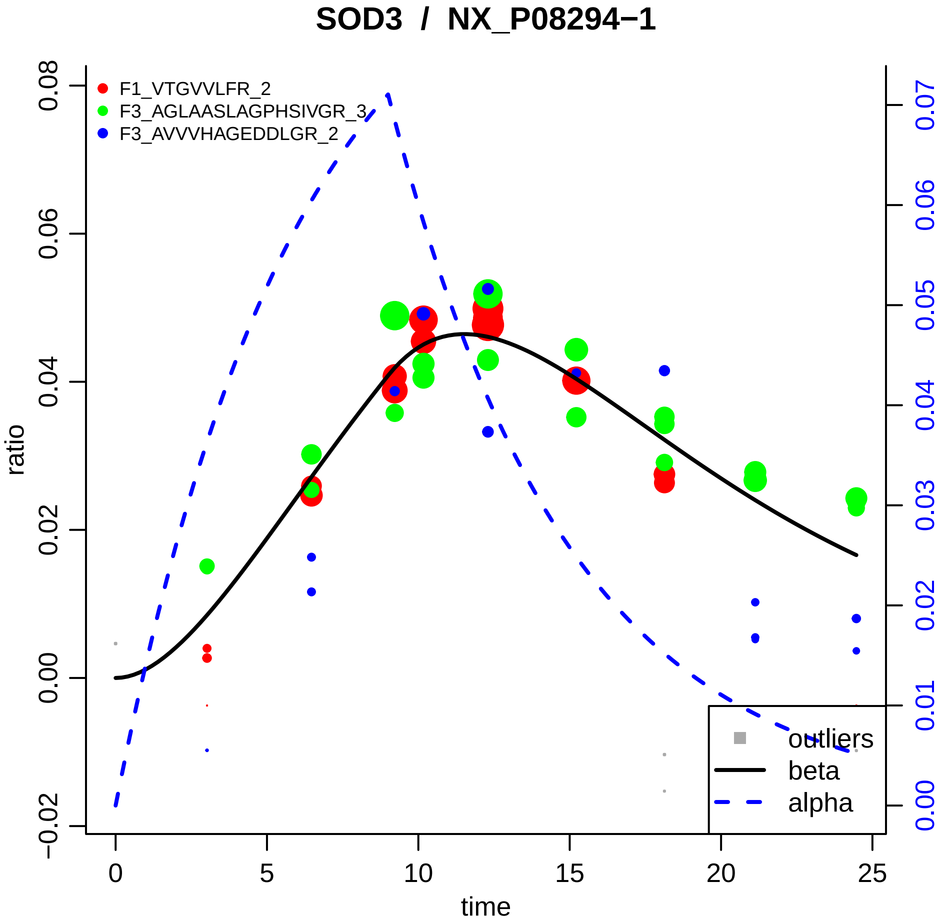

SOD3 / NX_P08294−1
F1_VTGVVLFR_2
F3_AGLAASLAGPHSIVGR_3
F3_AVVVHAGEDDLGR_2
outliers
beta
alpha
ratio
time
0.08
0.06
0.04
0.02
0.00
−0.02
0
5
10
15
20
25
0.07
0.06
0.05
0.04
0.03
0.02
0.01
0.00

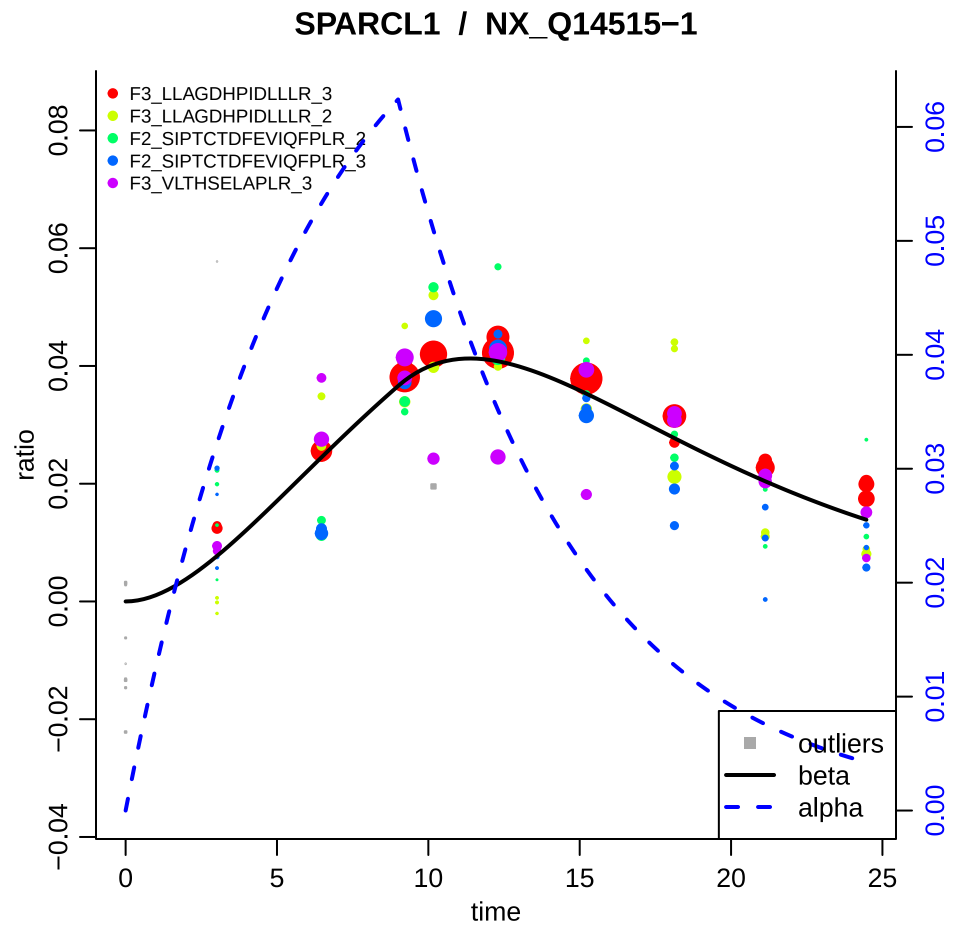
SPARCL1 / NX_Q14515−1
F3_LLAGDHPIDLLLR_3
F3_LLAGDHPIDLLLR_2
F2_SIPTCTDFEVIQFPLR_2
F2_SIPTCTDFEVIQFPLR_3
F3_VLTHSELAPLR_3
ratio
time
outliers
beta
alpha
0.08
0.06
0.04
0.02
0.00
−0.02
−0.04
0
5
10
15
20
25
0.06
0.05
0.04
0.03
0.02
0.01
0.00

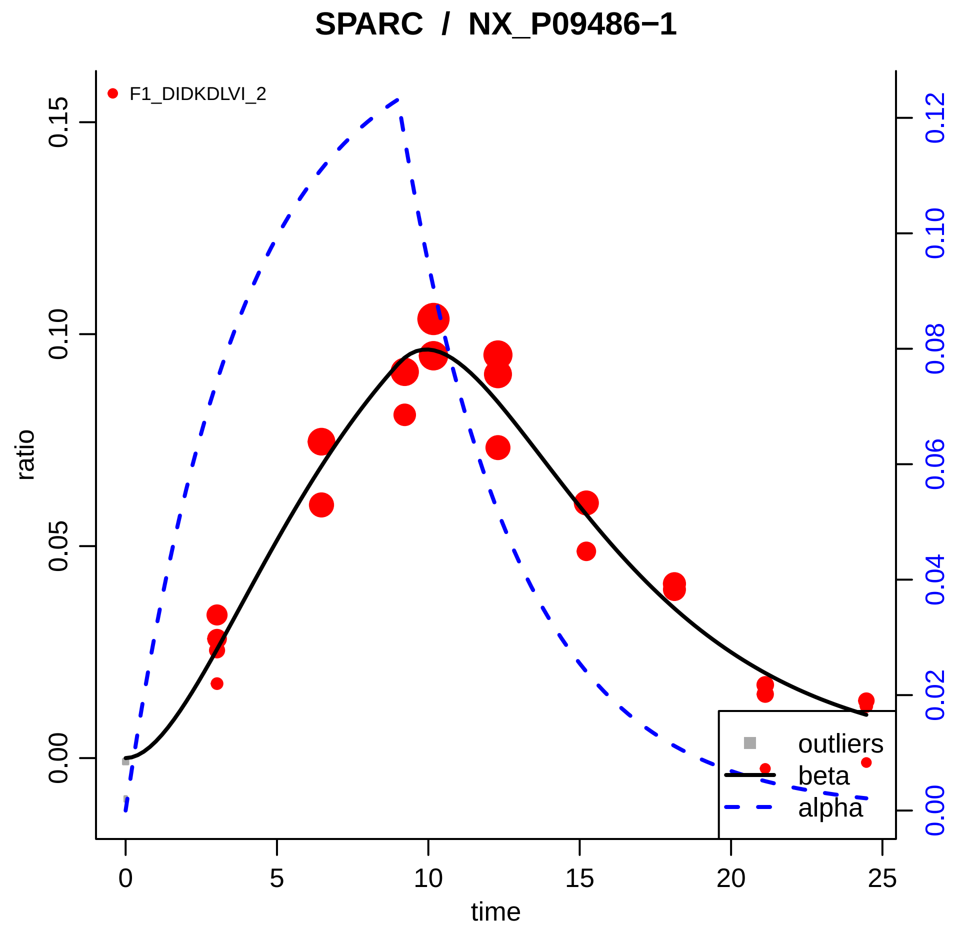

SPARC / NX_P09486−1
F1_DIDKDLVI_2
ratio
time
0.00
0.05
0.10
0.15
0
5
10
15
20
25
0.00
0.02
0.04
0.06
0.08
0.10
0.12
outliers
beta
alpha

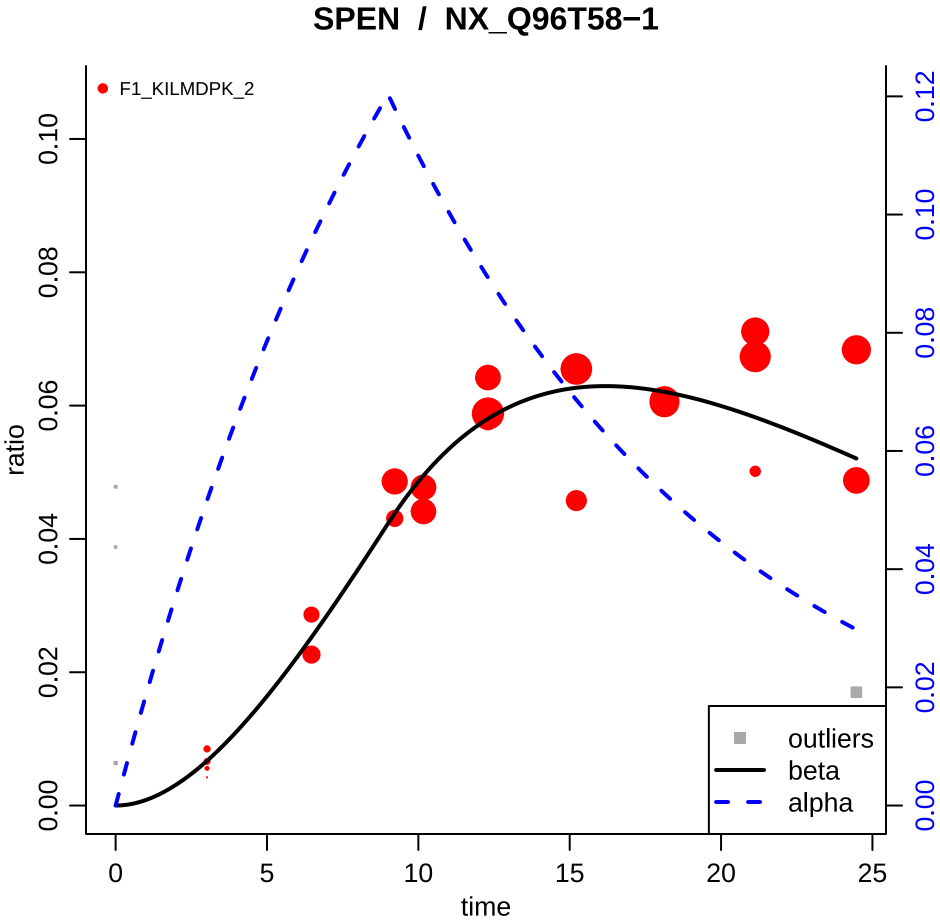

SPEN / NX_Q96T58−1
F1_KILMDPK_2
ratio
time
0.00
0.02
0.04
0.06
0.08
0.10
0.12
0
5
10
15
20
25
outliers
beta
alpha

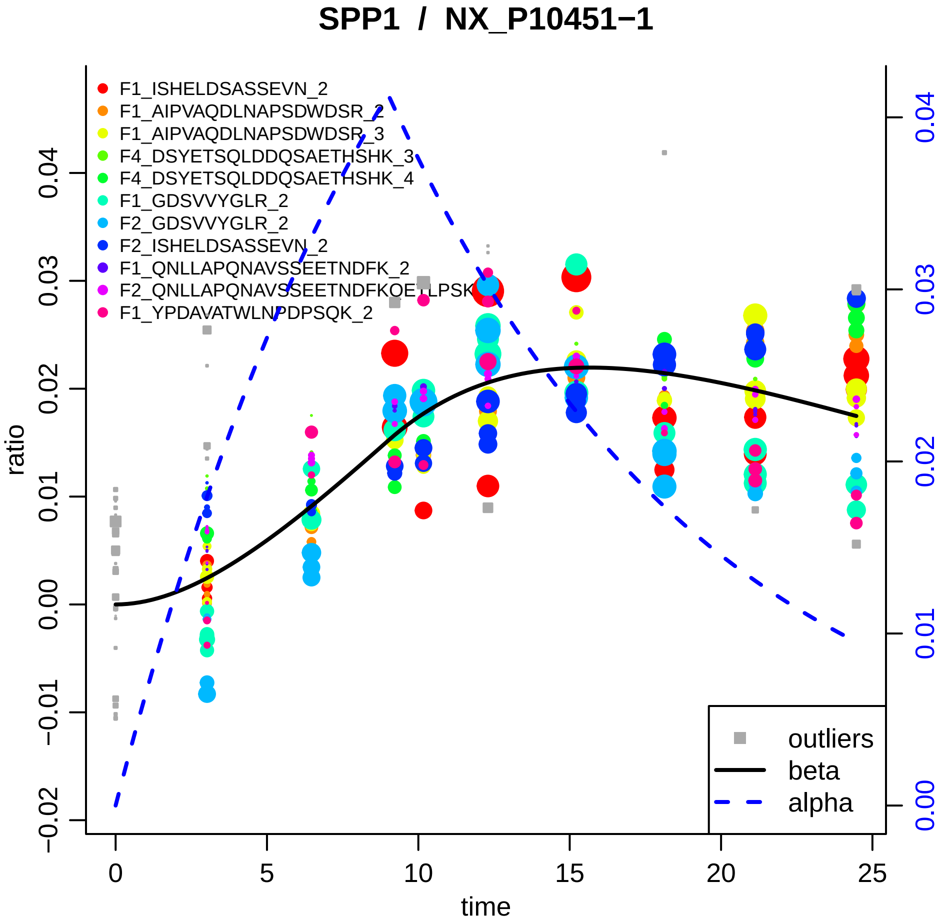

SPP1 / NX_P10451−1
F1_ISHELDSASSEVN_2
F1_AIPVAQDLNAPSDWDSR_2
F1_AIPVAQDLNAPSDWDSR_3
F4_DSYETSQLDDQSAETHSHK_3
F4_DSYETSQLDDQSAETHSHK_4
F1_GDSVVYGLR_2
F2_GDSVVYGLR_2
F2_ISHELDSASSEVN_2
F1_QNLLAPQNAVSSEETNDFK_2
F2_QNLLAPQNAVSSEETNDFKQETLPSK
F1_YPDAVATWLNPDPSQK_2
outliers
beta
alpha
ratio
time
−0.02
−0.01
0.00
0.01
0.02
0.03
0.04
0
5
10
15
20
25
0.00
0.01
0.02
0.03
0.04

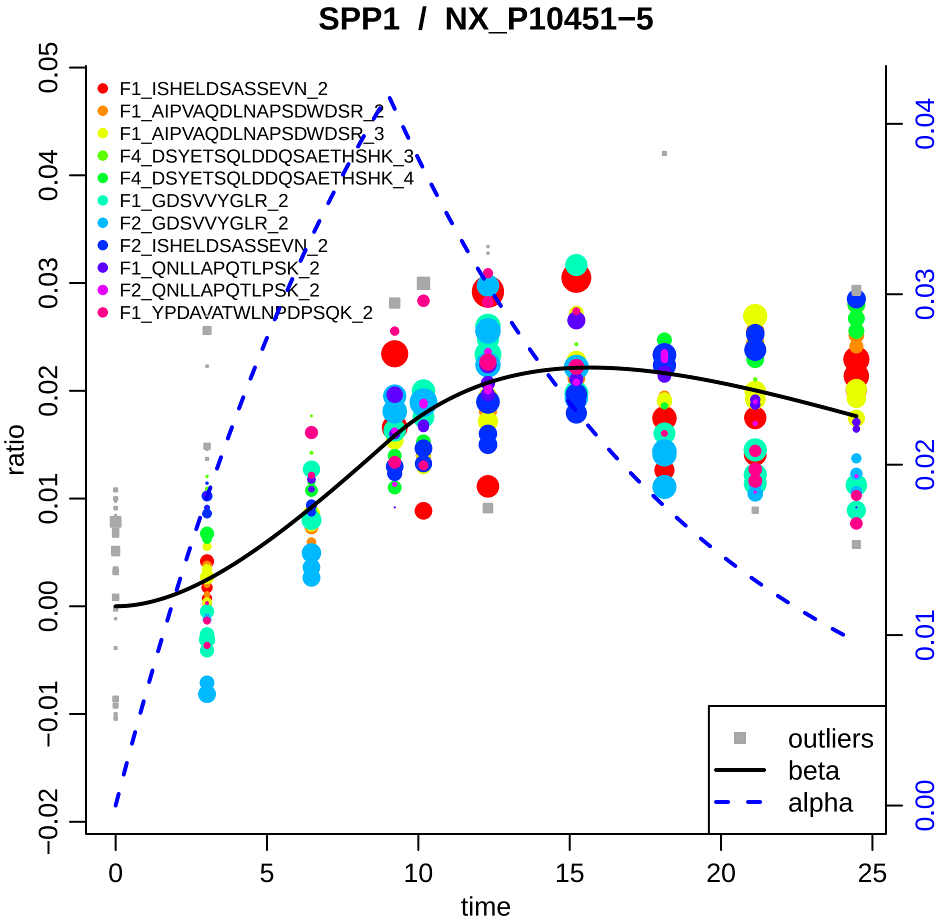

SPP1 / NX_P10451−5
F1_ISHELDSASSEVN_2
F1_AIPVAQDLNAPSDWDSR_2
F1_AIPVAQDLNAPSDWDSR_3
F4_DSYETSQLDDQSAETHSHK_3
F4_DSYETSQLDDQSAETHSHK_4
F1_GDSVVYGLR_2
F2_GDSVVYGLR_2
F2_ISHELDSASSEVN_2
F1_QNLLAPQTLPSK_2
F2_QNLLAPQTLPSK_2
F1_YPDAVATWLNPDPSQK_2
outliers
beta
alpha
ratio
time
0.05
0.04
0.03
0.02
0.01
0.00
−0.01
−0.02
0
5
10
15
20
25
0.04
0.03
0.02
0.01
0.00

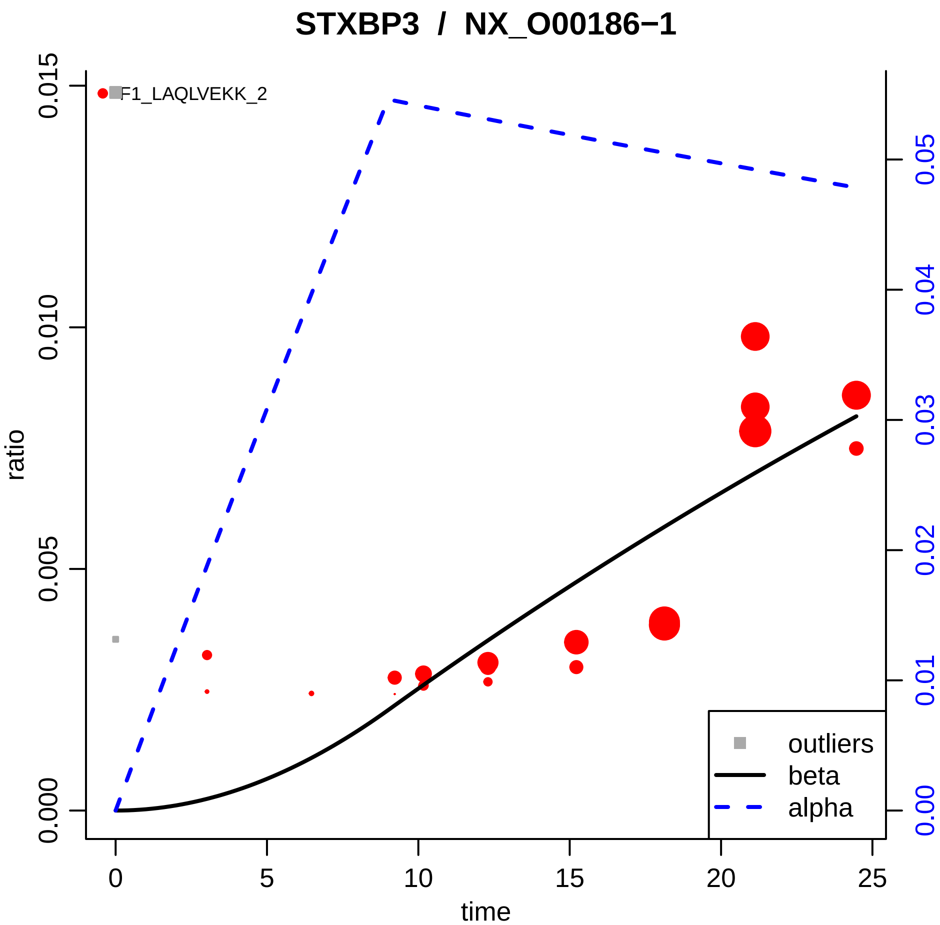
STXBP3 / NX_O00186−1
F1_LAQLVEKK_2
ratio
0.000
0.005
0.010
0.015
time
0
5
10
15
20
25
0.00
0.01
0.02
0.03
0.04
0.05
outliers
beta
alpha

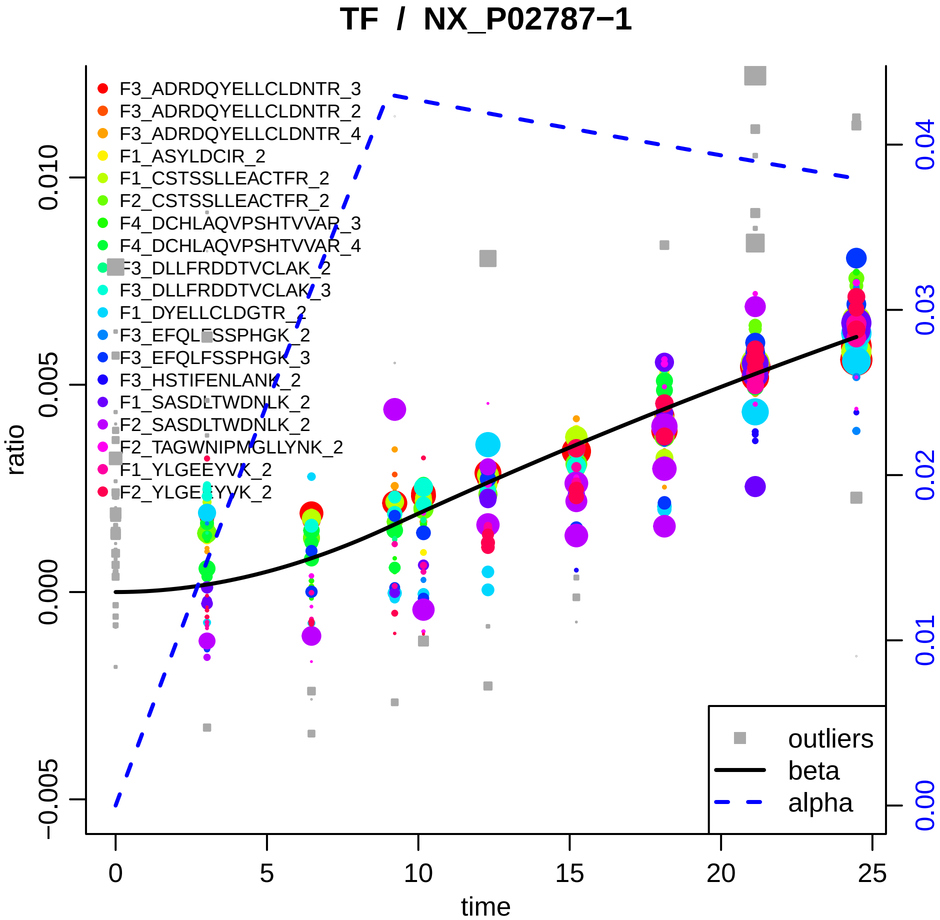

TF / NX_P02787−1
F3_ADRDQYELLCLDNTR_3
F3_ADRDQYELLCLDNTR_2
F3_ADRDQYELLCLDNTR_4
F1_ASYLDCIR_2
F1_CSTSSLLEACTFR_2
F2_CSTSSLLEACTFR_2
F4_DCHLAQVPSHTVVAR_3
F4_DCHLAQVPSHTVVAR_4
F3_DLLFRDDTVCLAK_2
F3_DLLFRDDTVCLAK_3
F1_DYELLCLDGTR_2
F3_EFQLFSSPHGK_2
F3_EFQLFSSPHGK_3
F3_HSTIFENLANK_2
F1_SASDLTWDNLK_2
F2_SASDLTWDNLK_2
F2_TAGWNIPMGLLYNK_2
F1_YLGEEYVK_2
F2_YLGEEYVK_2
outliers
beta
alpha
ratio
time
0.010
0.005
0.000
−0.005
0
5
10
15
20
25
0.04
0.03
0.02
0.01
0.00

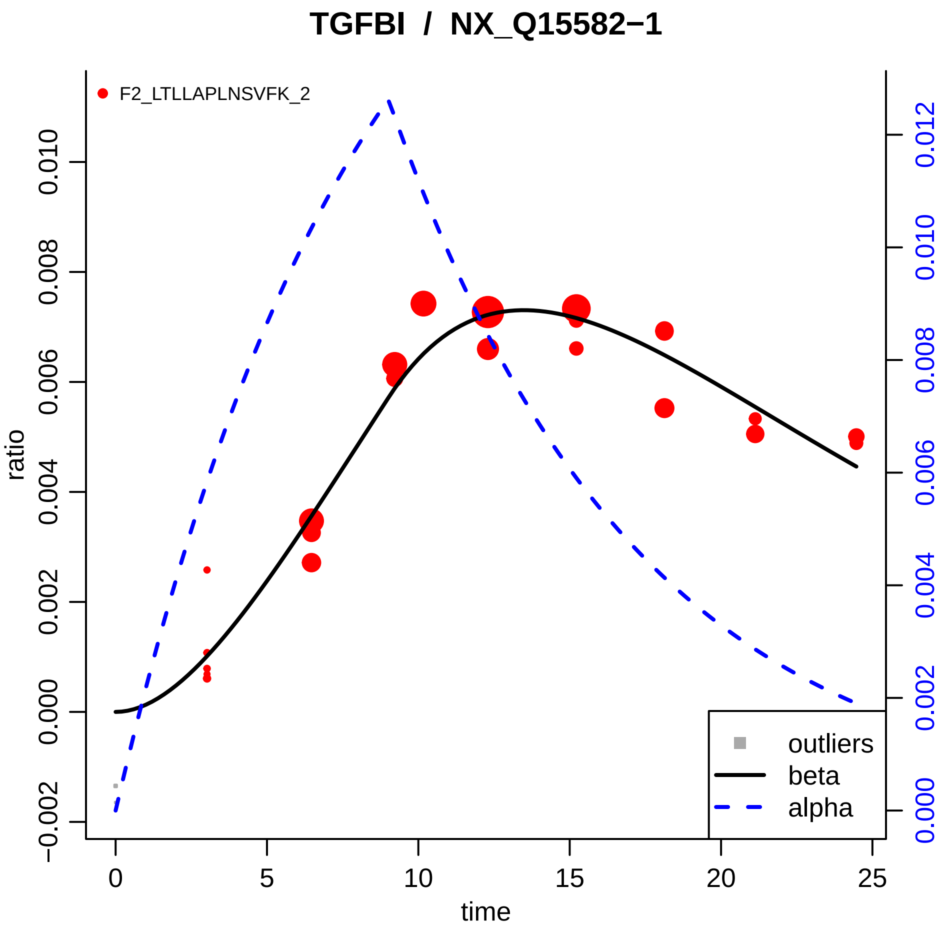
TGFBI / NX_Q15582−1
F2_LTLLAPLNSVFK_2
ratio
time
0.010
0.008
0.006
0.004
0.002
0.000
−0.002
0.012
0
5
10
15
20
25
outliers
beta
alpha

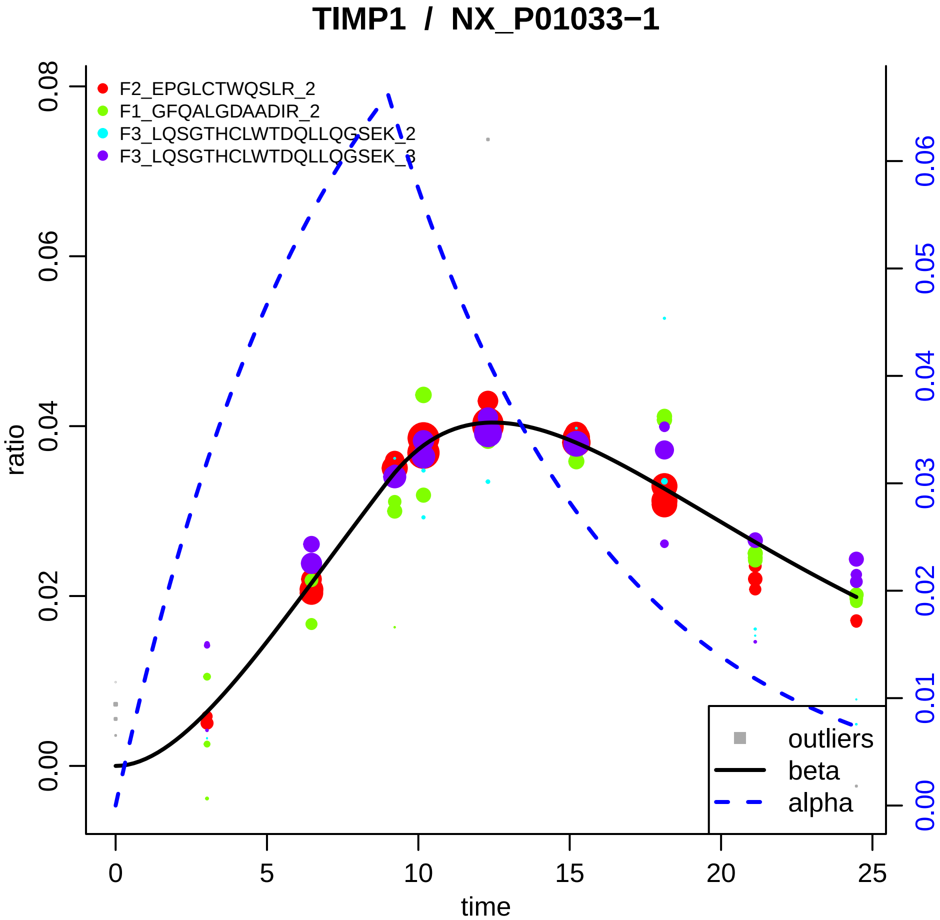
TIMP1 / NX_P01033−1
F2_EPGLCTWQSLR_2
F1_GFQALGDAADIR_2
F3_LQSGTHCLWTDQLLQGSEK_2
F3_LQSGTHCLWTDQLLQGSEK_3
outliers
beta
alpha
ratio
time
0.00
0.02
0.04
0.06
0.08
0
5
10
15
20
25
0.00
0.01
0.02
0.03
0.04
0.05
0.06

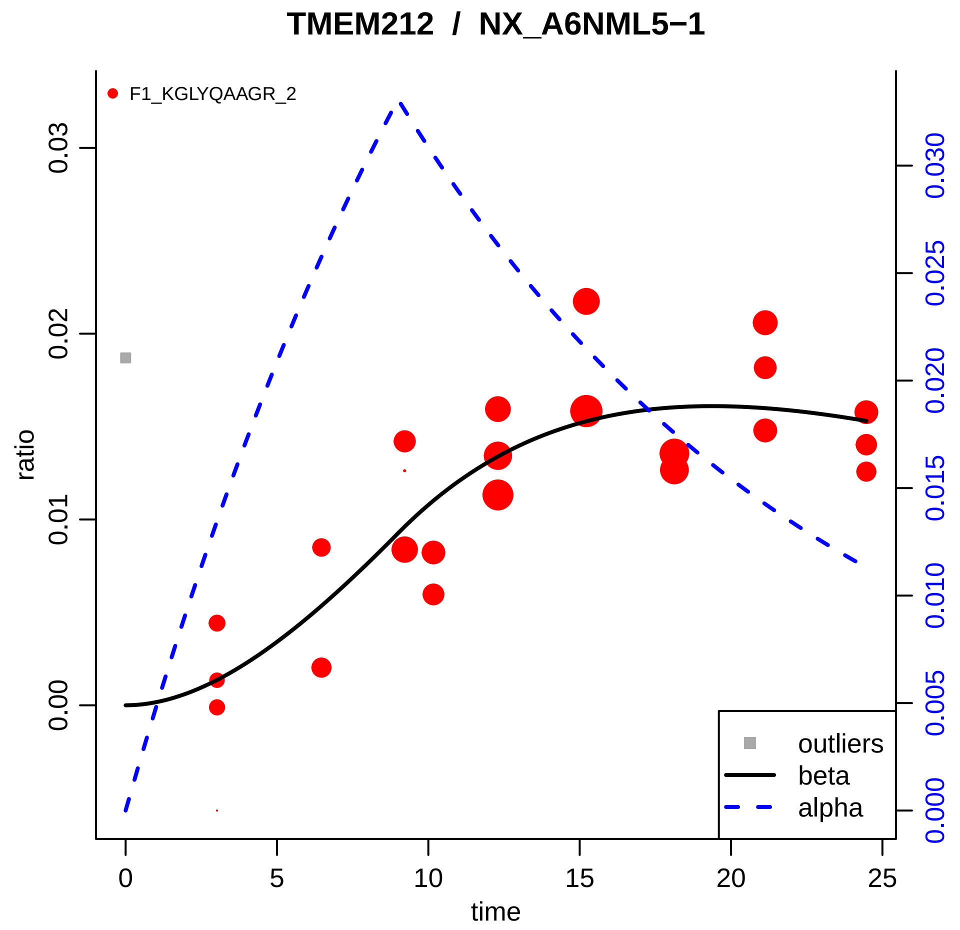

TMEM212 / NX_A6NML5−1
F1_KGLYQAAGR_2
ratio
0.00
0.01
0.02
0.03
time
0
5
10
15
20
25
0.000
0.005
0.010
0.015
0.020
0.025
0.030
outliers
beta
alpha

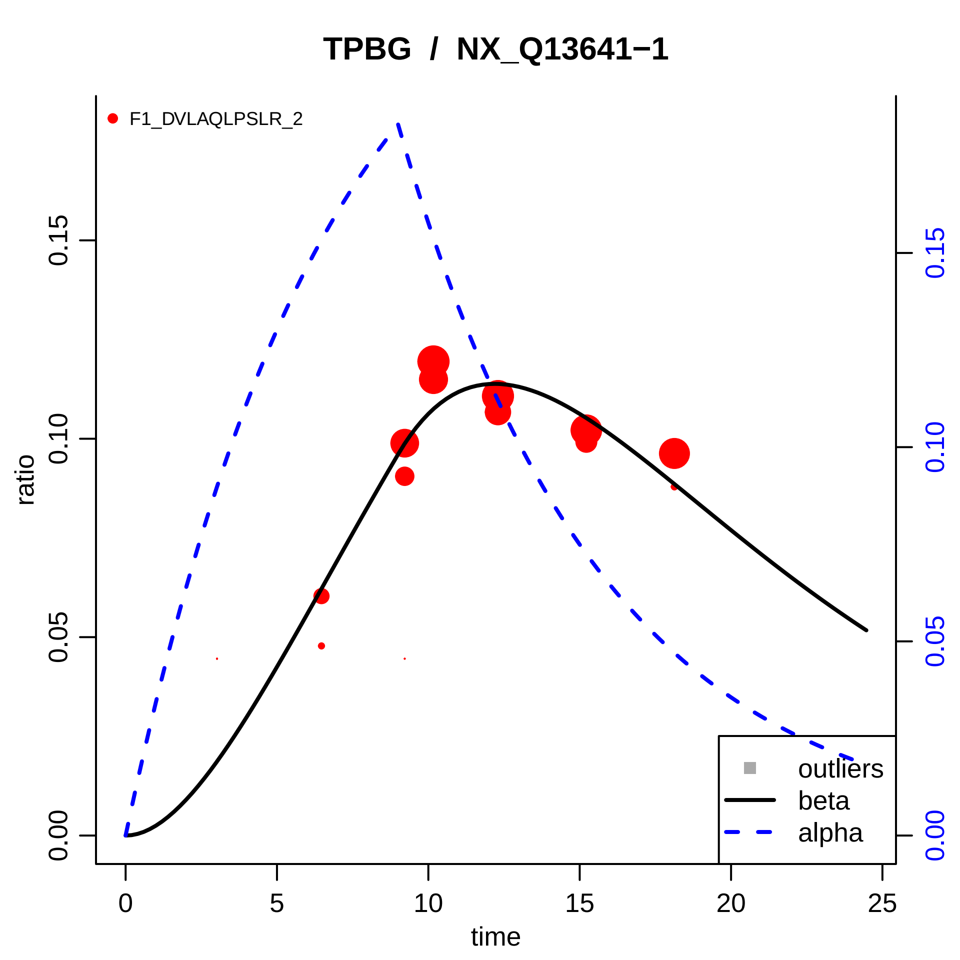

TPBG / NX_Q13641−1
F1_DVLAQLPSLR_2
ratio
time
0.00
0.05
0.10
0.15
0
5
10
15
20
25
outliers
beta
alpha

### TTR / NX_P02766−1

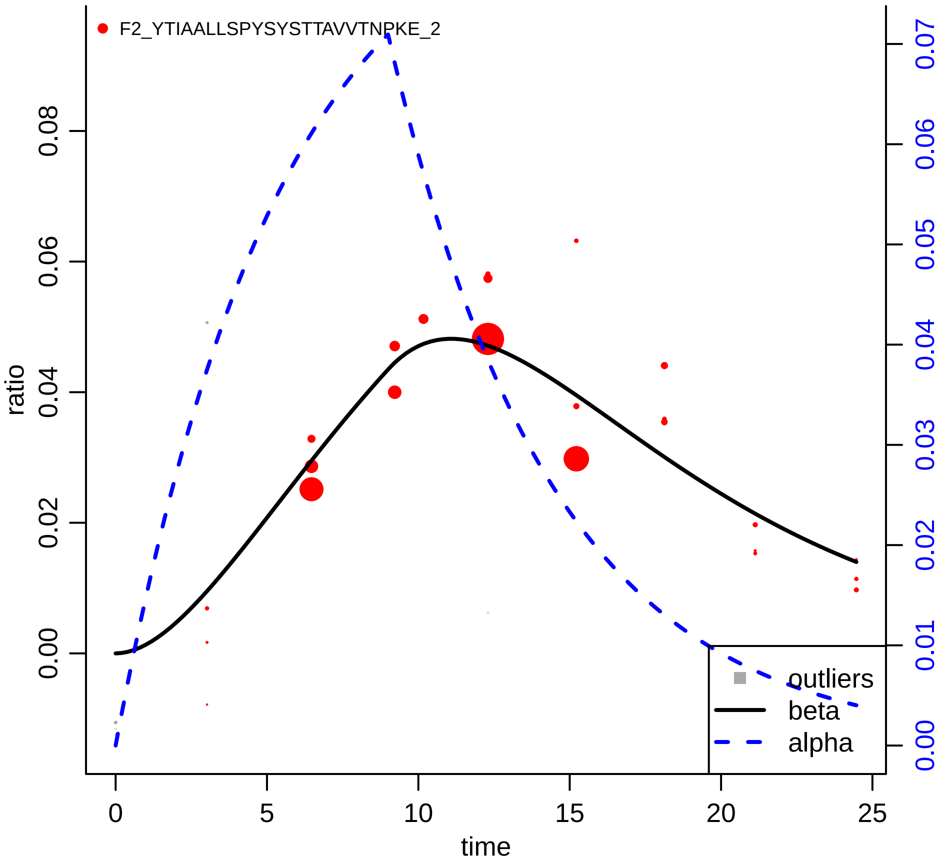

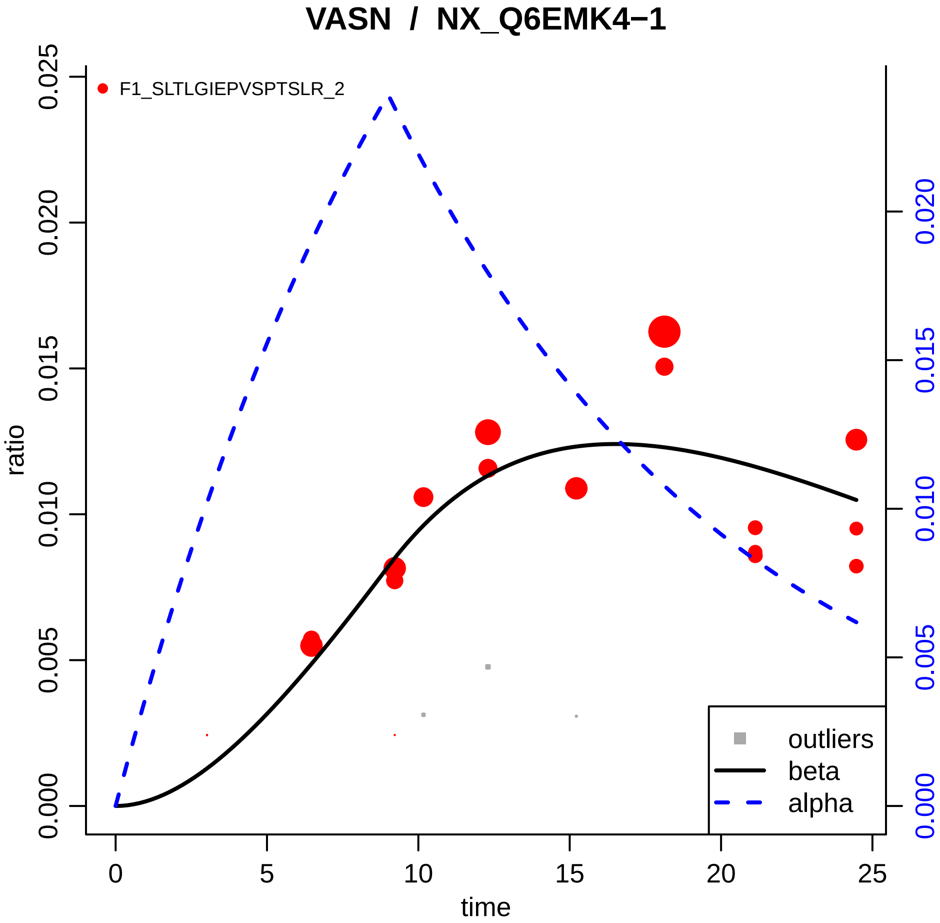

VASN / NX_Q6EMK4−1
F1_SLTLGIEPVSPTSLR_2
ratio
time
0.000
0.005
0.010
0.015
0.020
0.025
0
5
10
15
20
25
outliers
beta
alpha

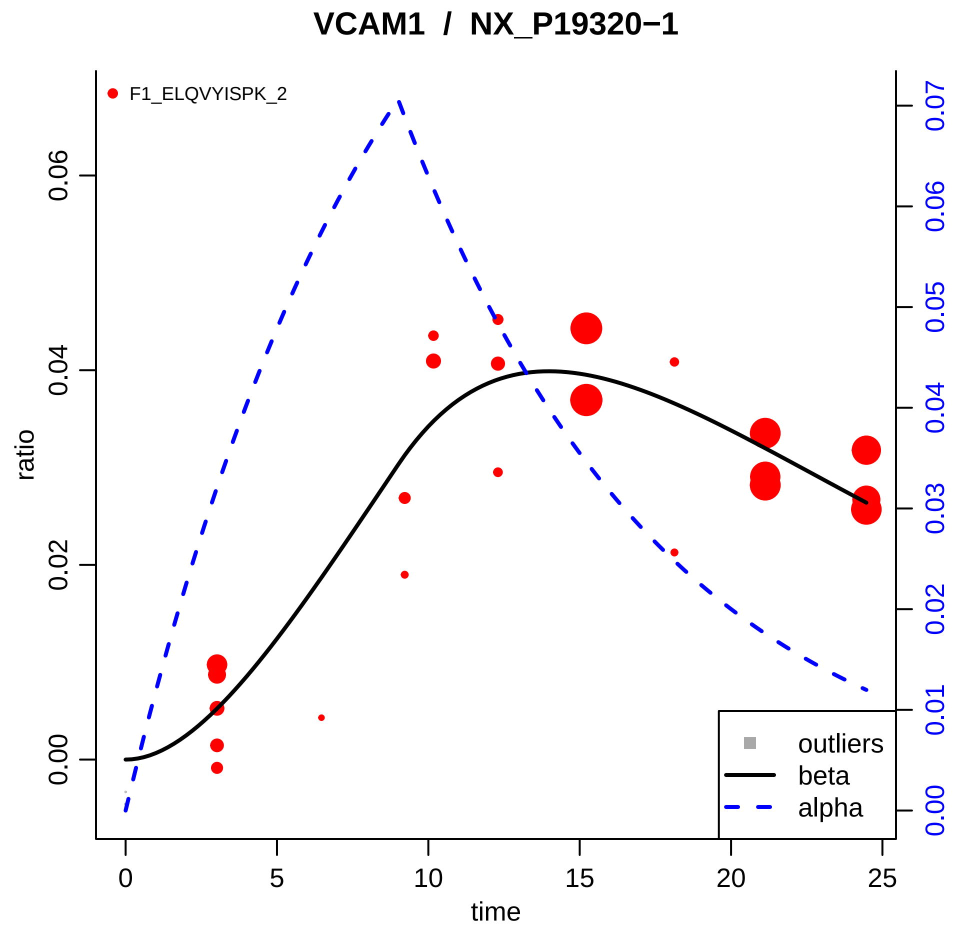
VCAM1 / NX_P19320−1
F1_ELQVYISPK_2
ratio
time
0.00
0.02
0.04
0.06
0
5
10
15
20
25
0.00
0.01
0.02
0.03
0.04
0.05
0.06
0.07
outliers
beta
alpha

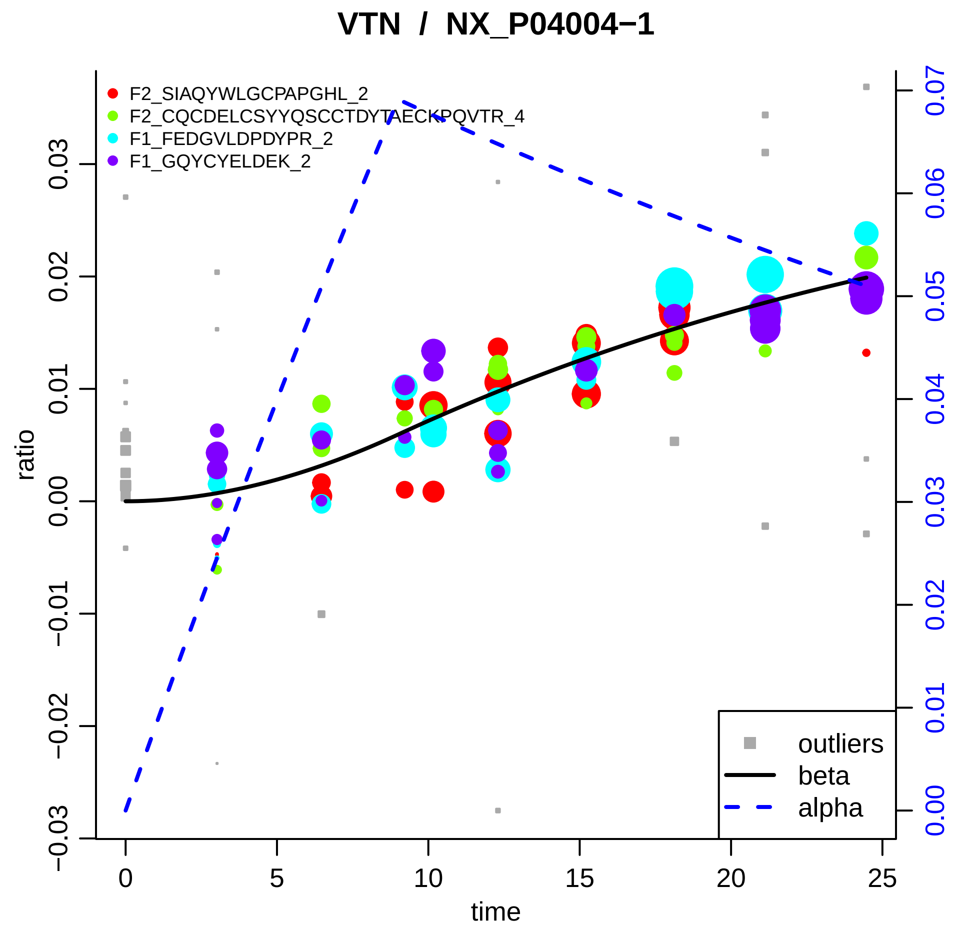

VTN / NX_P04004−1
F2_SIAQYWLGCPAPGHL_2
F2_CQCDELCSYYQSCCTDYTAECKPQVTR_4
F1_FEDGVLDPDYPR_2
F1_GQYCYELDEK_2
outliers
beta
alpha
ratio
time
0
5
10
15
20
25
−0.03
−0.02
−0.01
0.00
0.01
0.02
0.03
0.00
0.01
0.02
0.03
0.04
0.05
0.06
0.07
