## Supplementary material for "*In Vivo* Large Scale Mapping Of Protein Turnover In The Human Cerebrospinal Fluid": TQ protein models

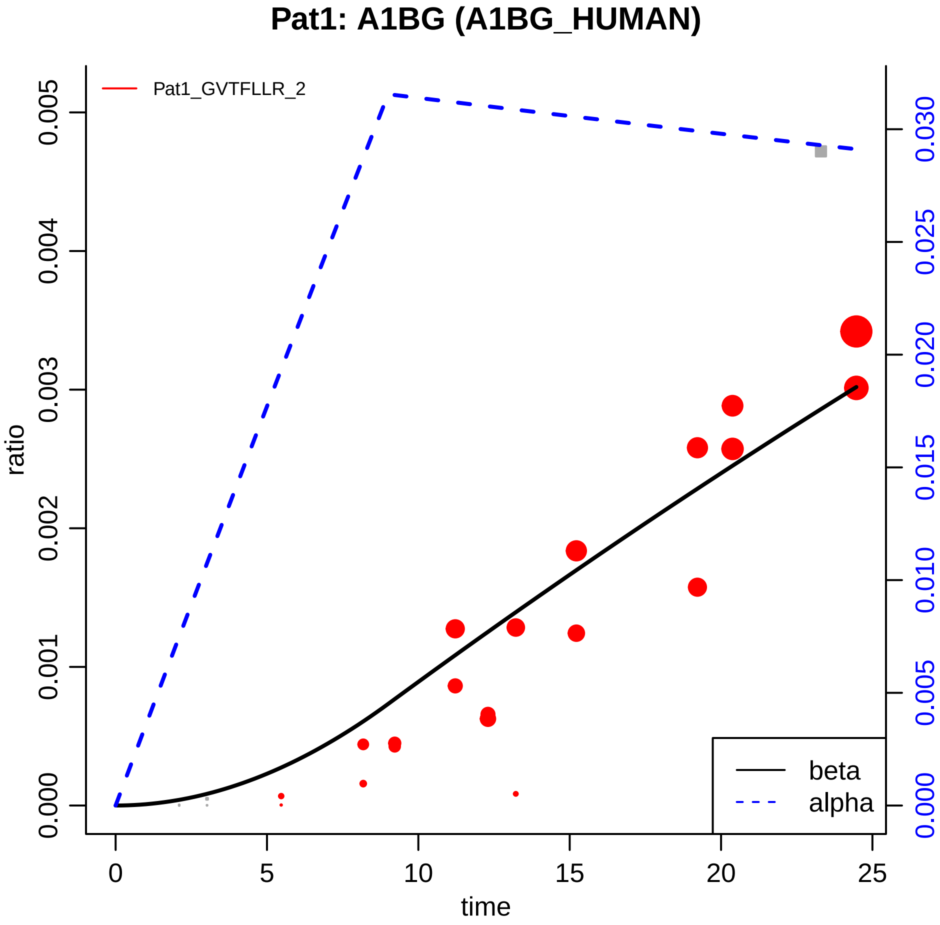

Pat1: A1BG (A1BG_HUMAN)
Pat1_GVTFLLR_2
ratio
0.000
0.001
0.002
0.003
0.004
0.005
0.000
0.005
0.010
0.015
0.020
0.025
0.030
0
5
10
15
20
25
time
beta
alpha

### Pat1: A2M (A2MG_HUMAN)

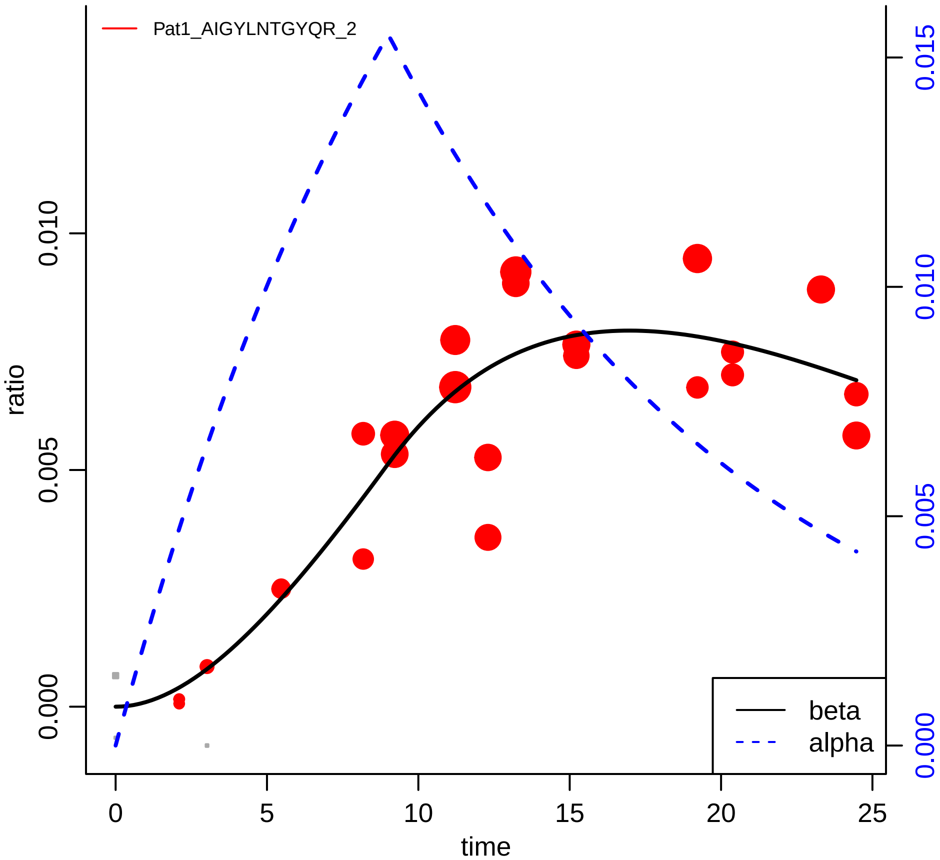

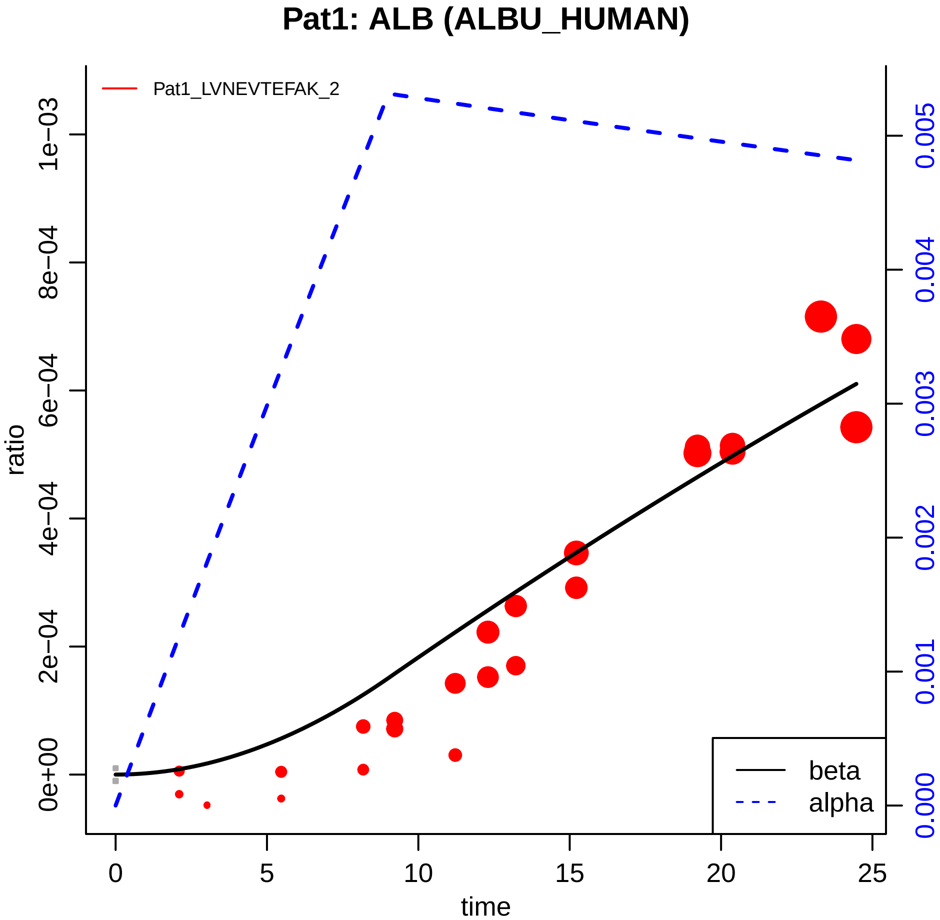

Pat1: ALB (ALBU_HUMAN)
Pat1_LVNEVTEFAK_2
ratio
0e+00
2e−04
4e−04
6e−04
8e−04
1e−03
0.000
0.001
0.002
0.003
0.004
0.005
0
5
10
15
20
25
time
beta
alpha

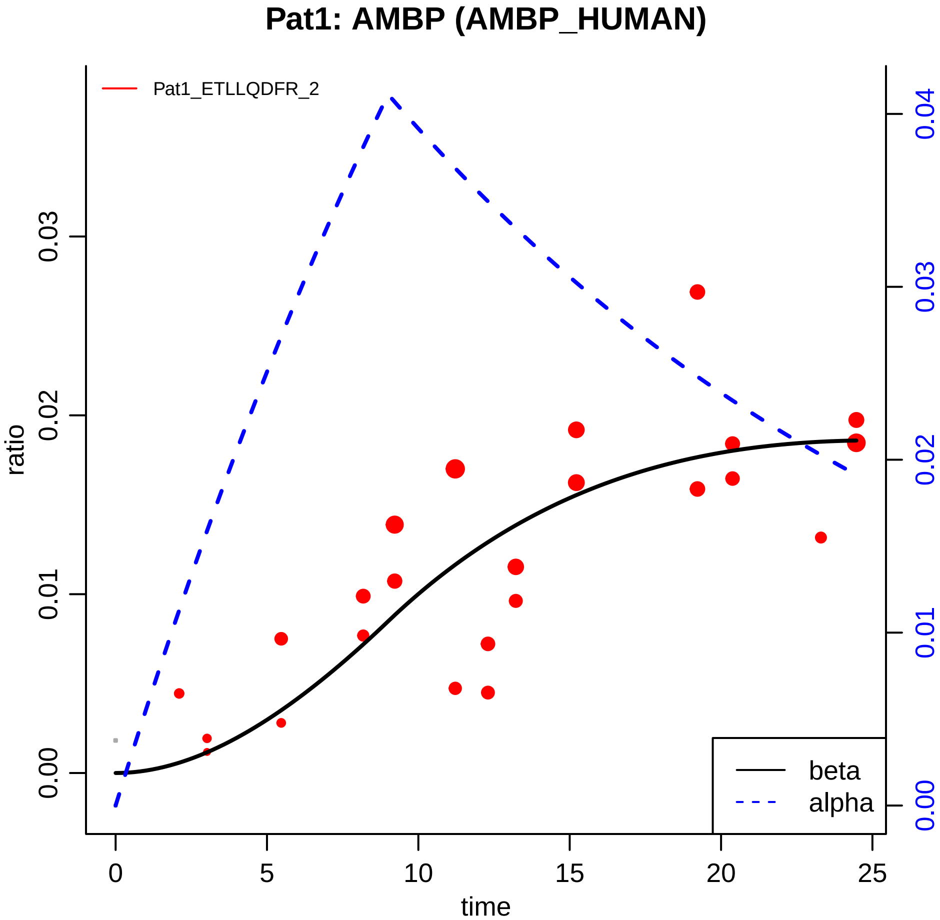

Pat1: AMBP (AMBP_HUMAN)
Pat1_ETLLQDFR_2
ratio
0.00
0.01
0.02
0.03
0.00
0.01
0.02
0.03
0.04
time
0
5
10
15
20
25
beta
alpha

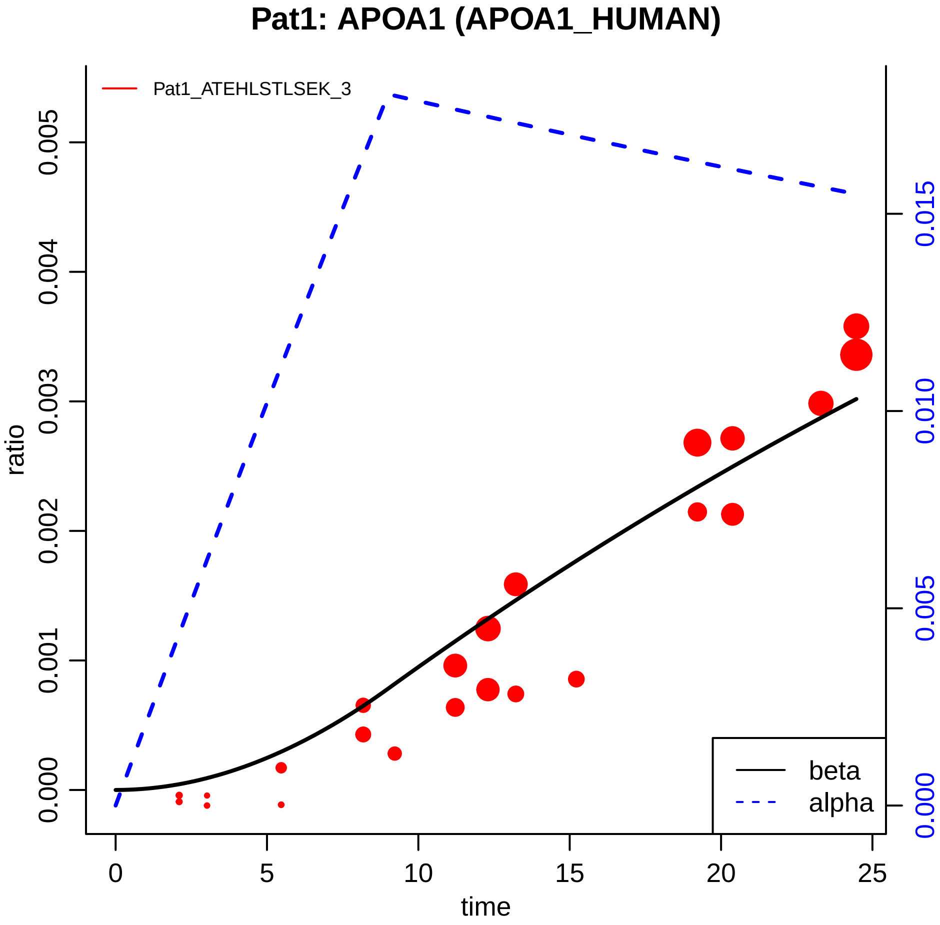

Pat1: APOA1 (APOA1_HUMAN)
Pat1_ATEHLSTLSEK_3
ratio
0.000
0.001
0.002
0.003
0.004
0.005
0.000
0.005
0.010
0.015
0
5
10
15
20
25
time
beta
alpha

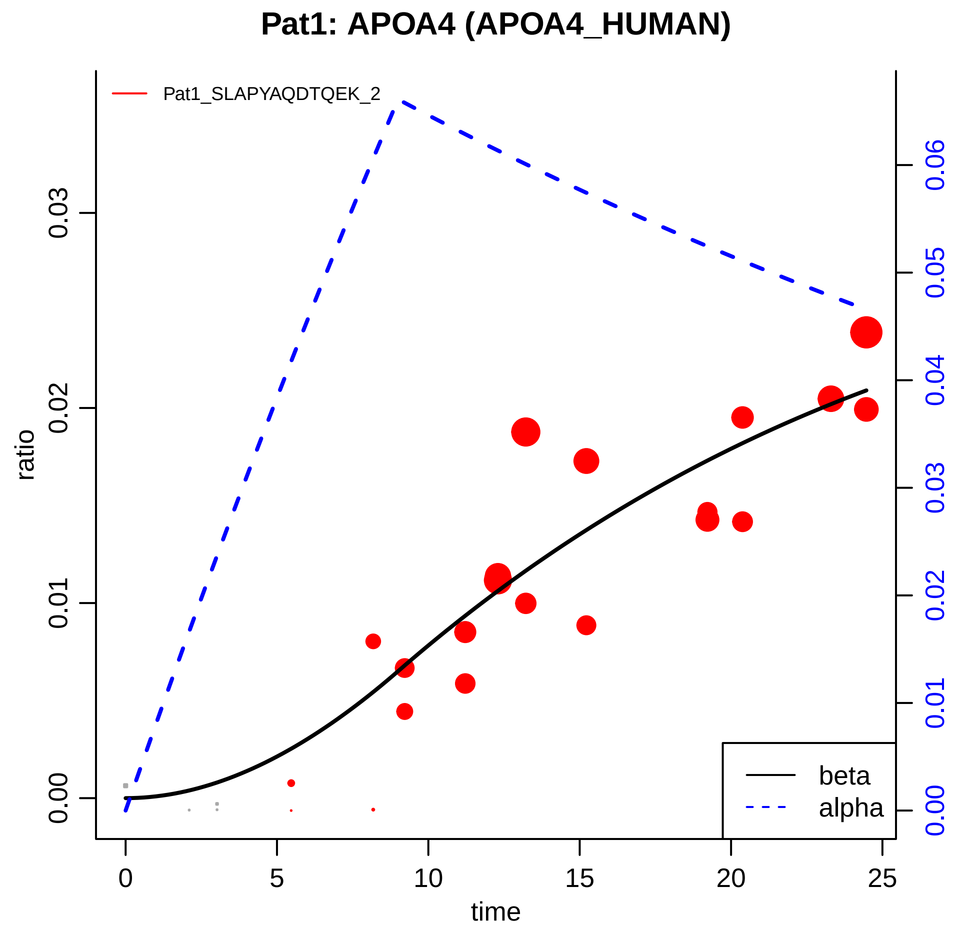

Pat1: APOA4 (APOA4_HUMAN)
Pat1_SLAPYAQDTQEK_2
ratio
0.00
0.01
0.02
0.03
time
0
5
10
15
20
25
0.00
0.01
0.02
0.03
0.04
0.05
0.06
beta
alpha

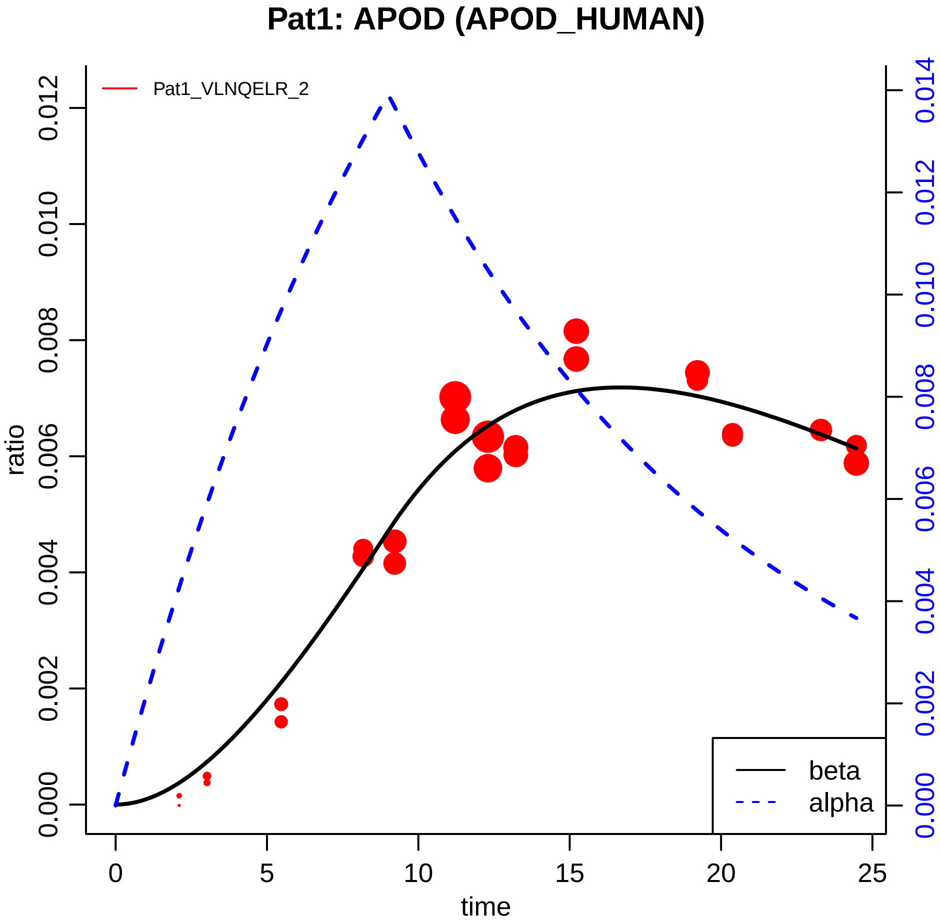

Pat1: APOD (APOD_HUMAN)
Pat1_VLNQELR_2
ratio
time
0.000
0.002
0.004
0.006
0.008
0.010
0.012
0.014
0
5
10
15
20
25
beta
alpha

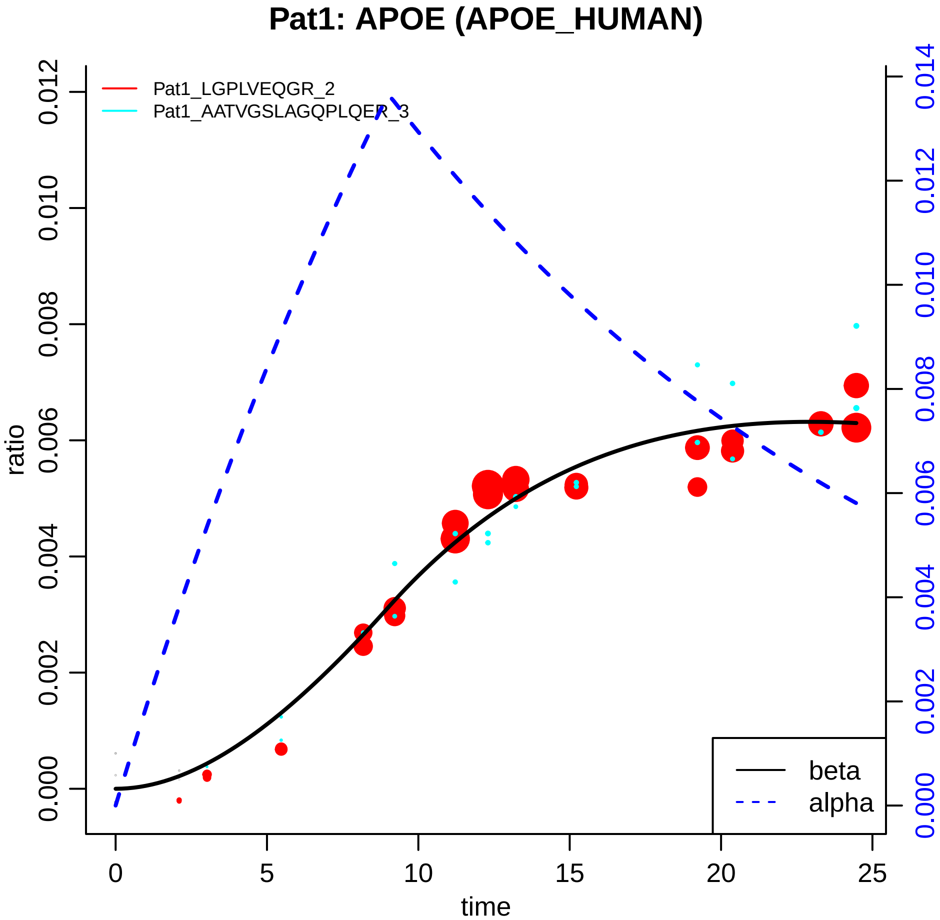

Pat1: APOE (APOE_HUMAN)
Pat1_LGPLVEQGR_2
Pat1_AATVGSLAGQPLQER_3
ratio
time
0.000
0.002
0.004
0.006
0.008
0.010
0.012
0.014
0
5
10
15
20
25
beta
alpha

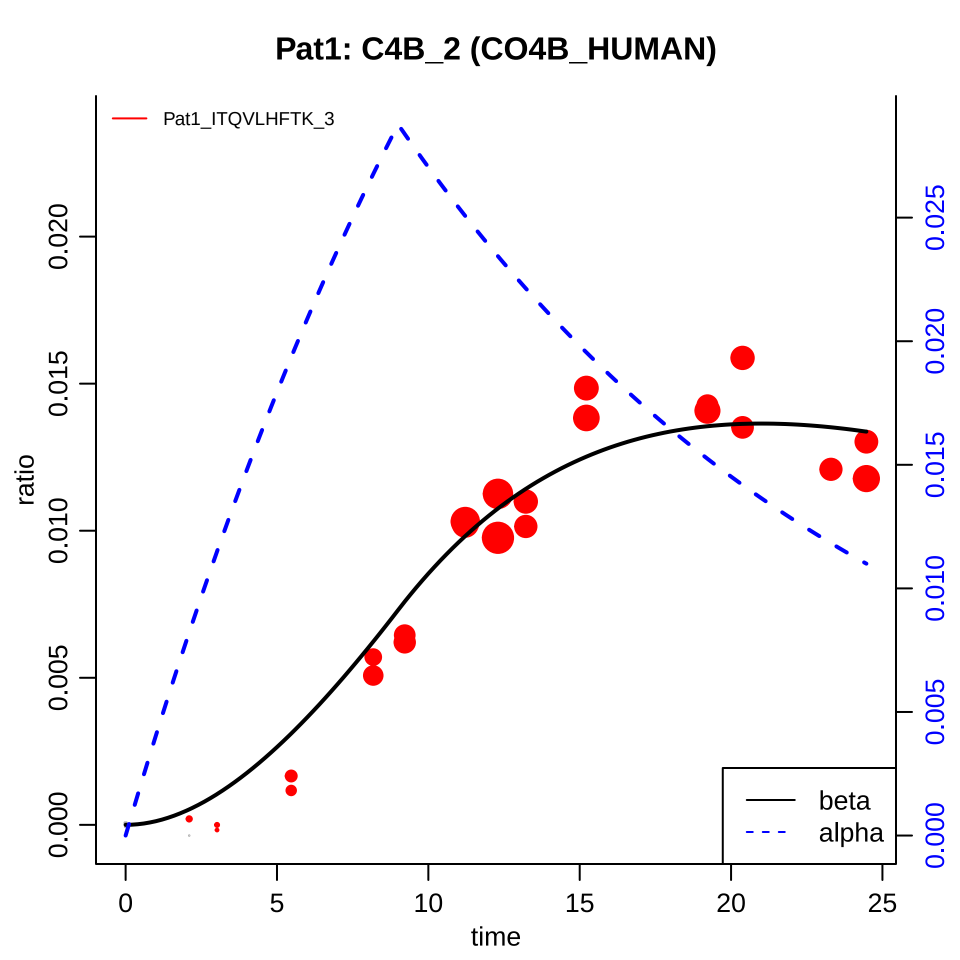

Pat1: C4B_2 (CO4B_HUMAN)
Pat1_ITQVLHFTK_3
ratio
0.000
0.005
0.010
0.015
0.020
0.000
0.005
0.010
0.015
0.020
0.025
0
5
10
15
20
25
time
beta
alpha

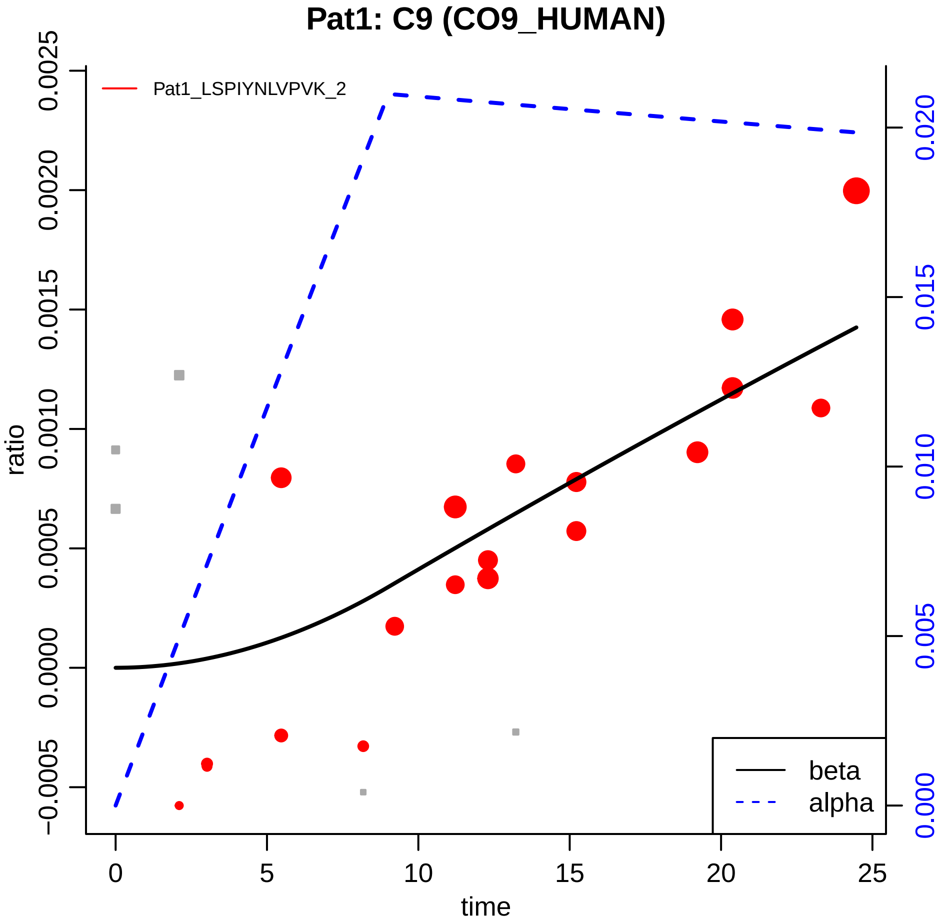

Pat1: C9 (CO9_HUMAN)
Pat1_LSPIYNLVPVK_2
ratio
time
beta
alpha
0
5
10
15
20
25
−0.0005
0.0000
0.0005
0.0010
0.0015
0.0020
0.0025
0.000
0.005
0.010
0.015
0.020

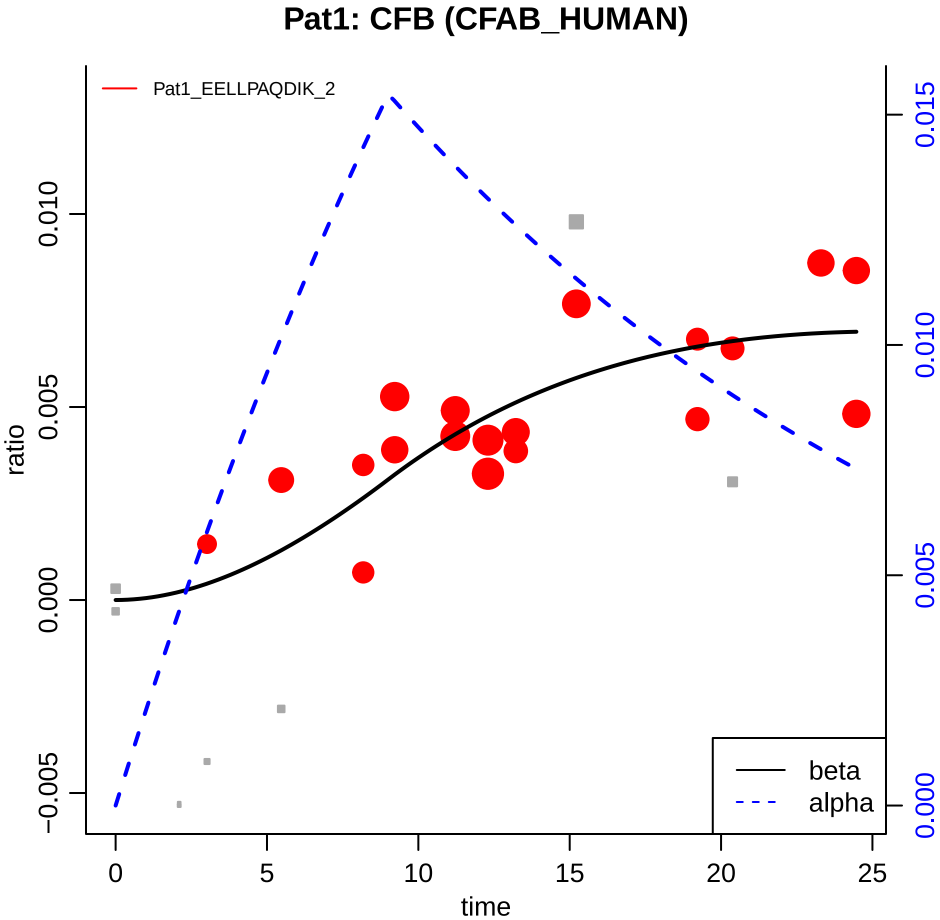

Pat1: CFB (CFAB_HUMAN)
Pat1_EELLPAQDIK_2
ratio
0.010
0.005
0.000
−0.005
0.015
0.010
0.005
0.000
0
5
10
15
20
25
time
beta
alpha

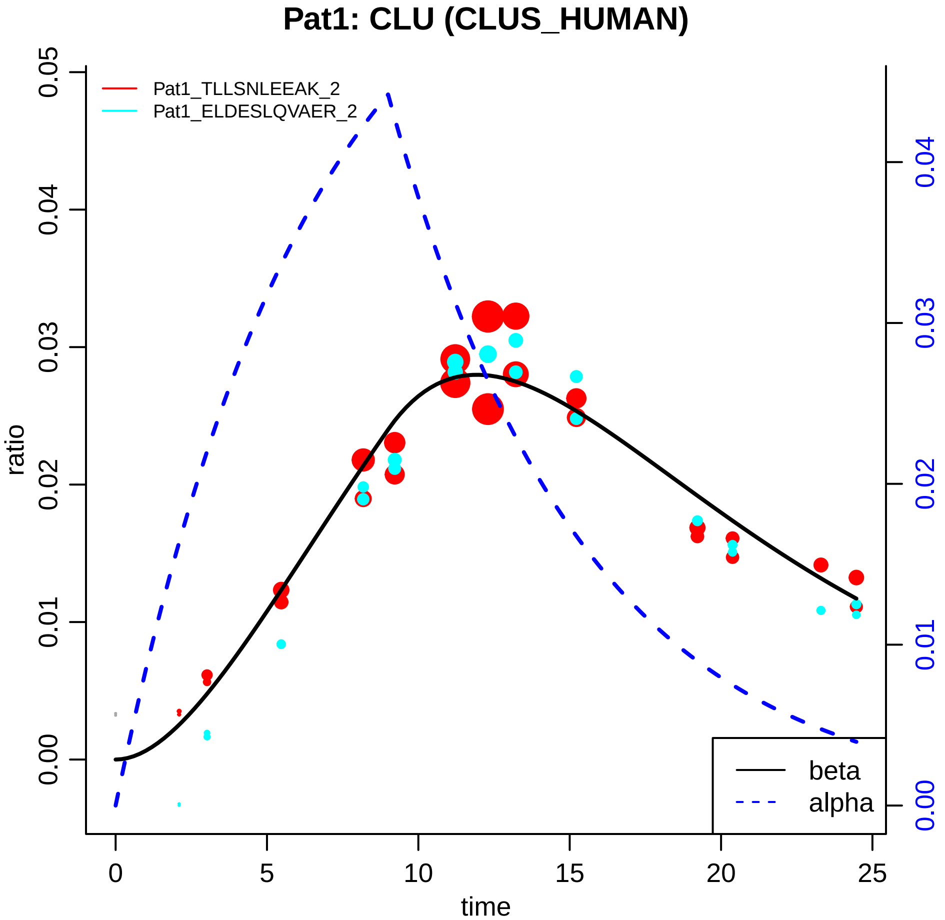

Pat1: CLU (CLUS_HUMAN)
Pat1_TLLSNLEEAK_2
Pat1_ELDESLQVAER_2
ratio
time
0.00
0.01
0.02
0.03
0.04
0.05
0
5
10
15
20
25
beta
alpha

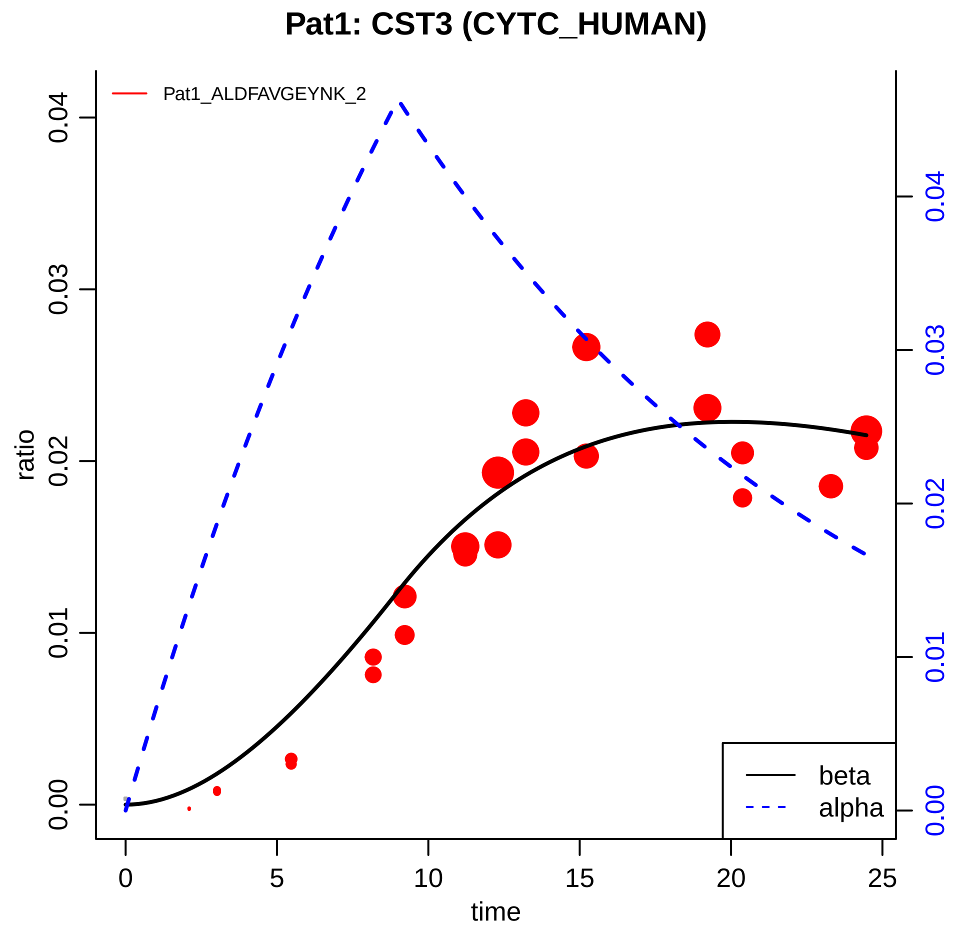

Pat1: CST3 (CYTC_HUMAN)
Pat1_ALDFAVGEYNK_2
ratio
0.00
0.01
0.02
0.03
0.04
time
0
5
10
15
20
25
beta
alpha

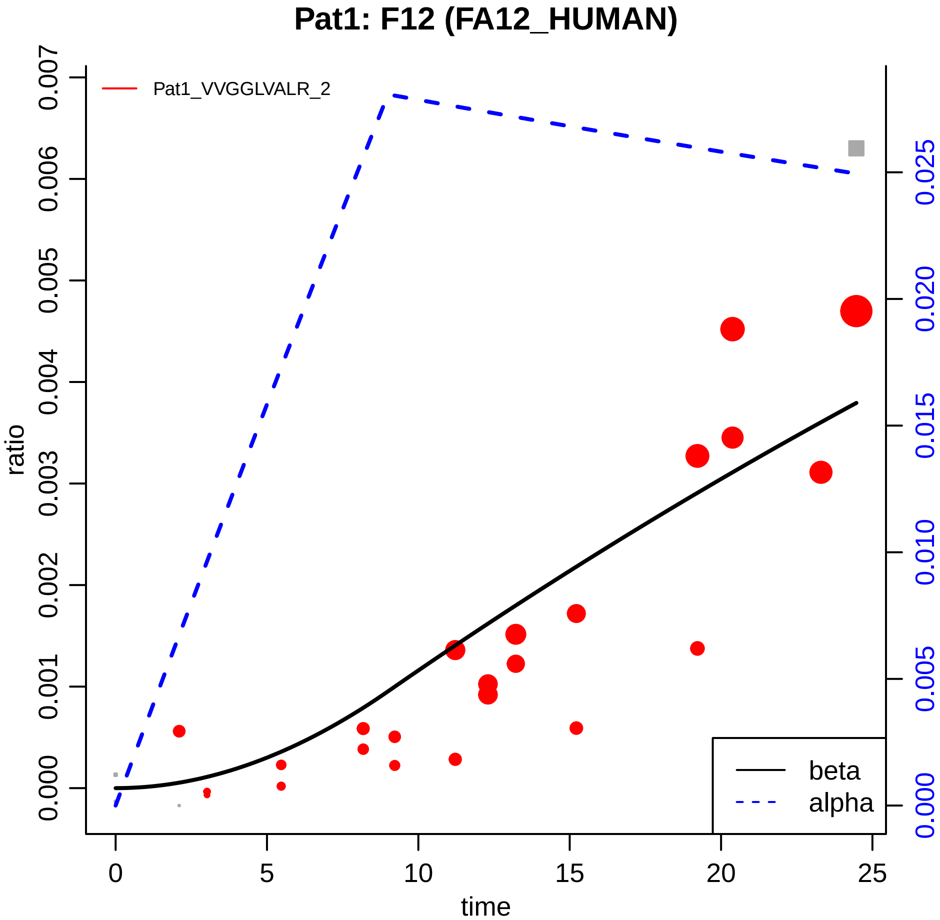
Pat1: F12 (FA12_HUMAN)
Pat1_VVGGLVALR_2
ratio
time
0.000
0.001
0.002
0.003
0.004
0.005
0.006
0.007
0
5
10
15
20
25
0.000
0.005
0.010
0.015
0.020
0.025
beta
alpha

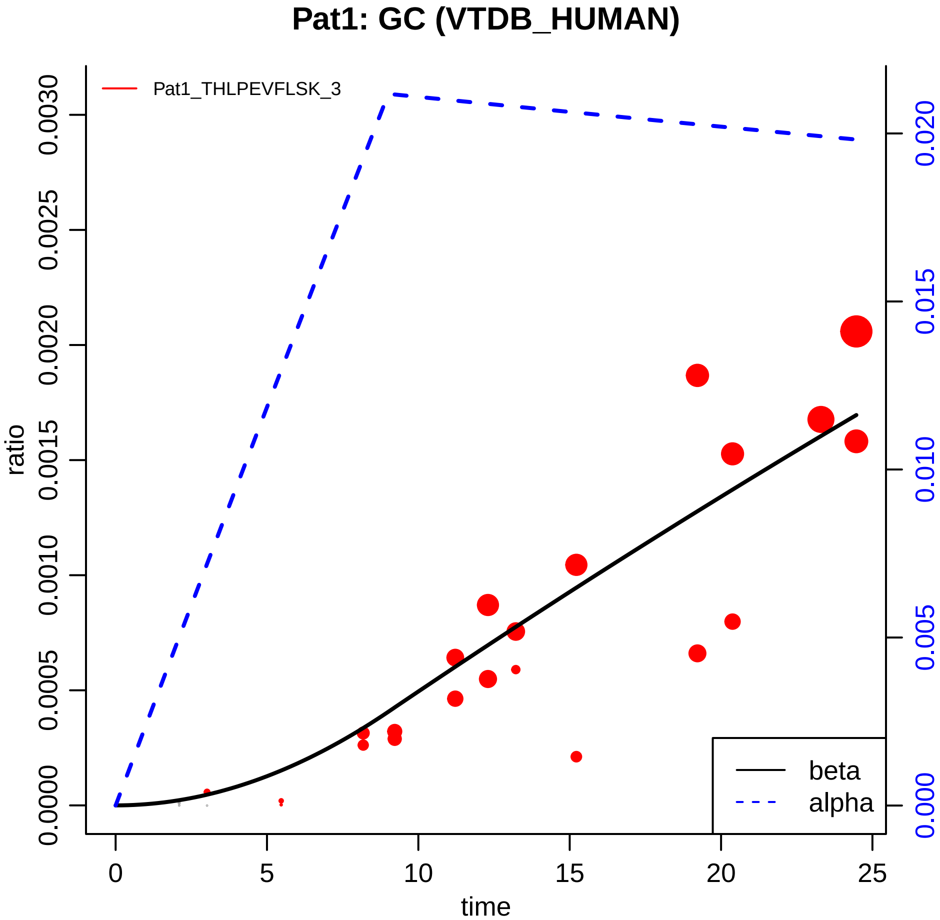
Pat1: GC (VTDB_HUMAN)
Pat1_THLPEVFLSK_3
ratio
0.0000
0.0005
0.0010
0.0015
0.0020
0.0025
0.0030
0.000
0.005
0.010
0.015
0.020
0
5
10
15
20
25
time
beta
alpha

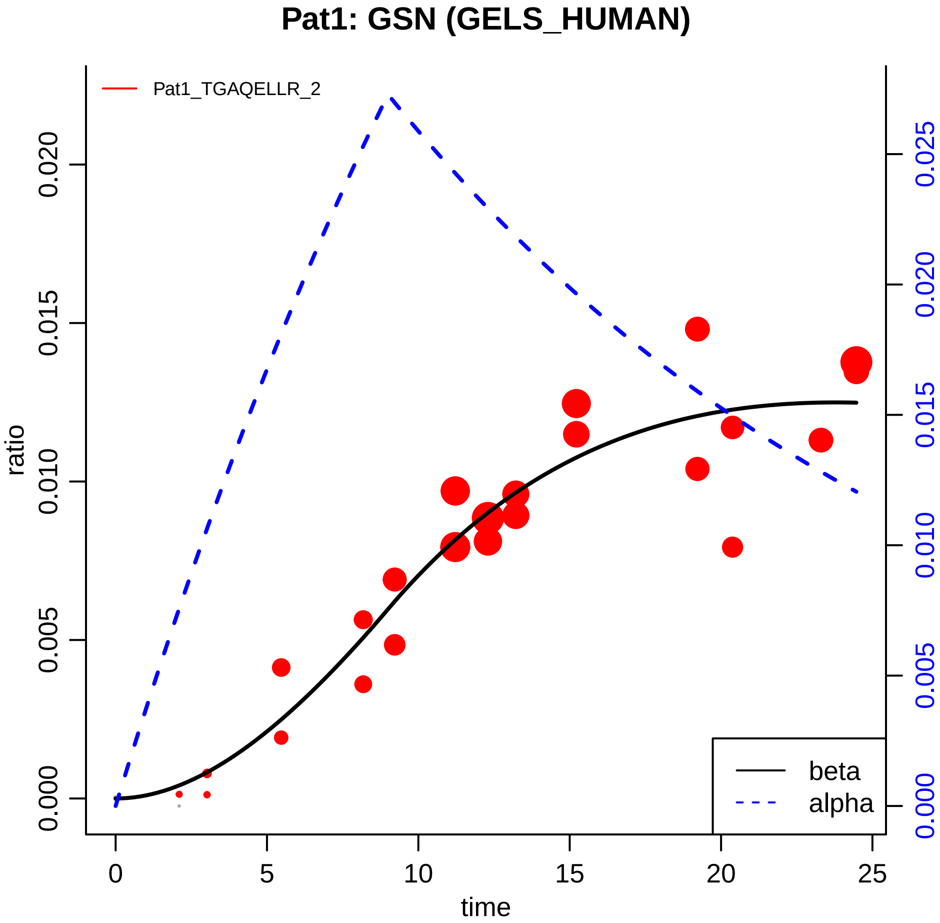

Pat1: GSN (GELS_HUMAN)
Pat1_TGAQELLR_2
ratio
0.000
0.005
0.010
0.015
0.020
0.025
time
0
5
10
15
20
25
beta
alpha

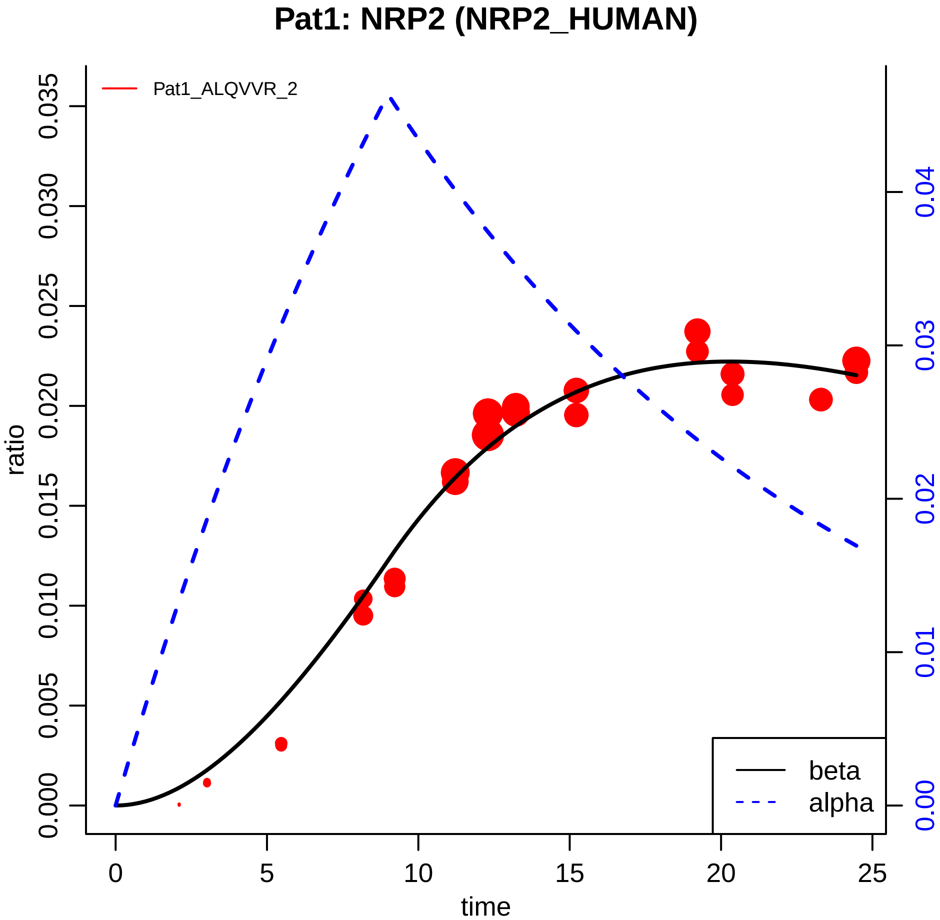

Pat1: NRP2 (NRP2_HUMAN)
Pat1_ALQVVR_2
ratio
0.000
0.005
0.010
0.015
0.020
0.025
0.030
0.035
0.00
0.01
0.02
0.03
0.04
time
0
5
10
15
20
25
beta
alpha

### Pat1: RBP4 (RET4_HUMAN)

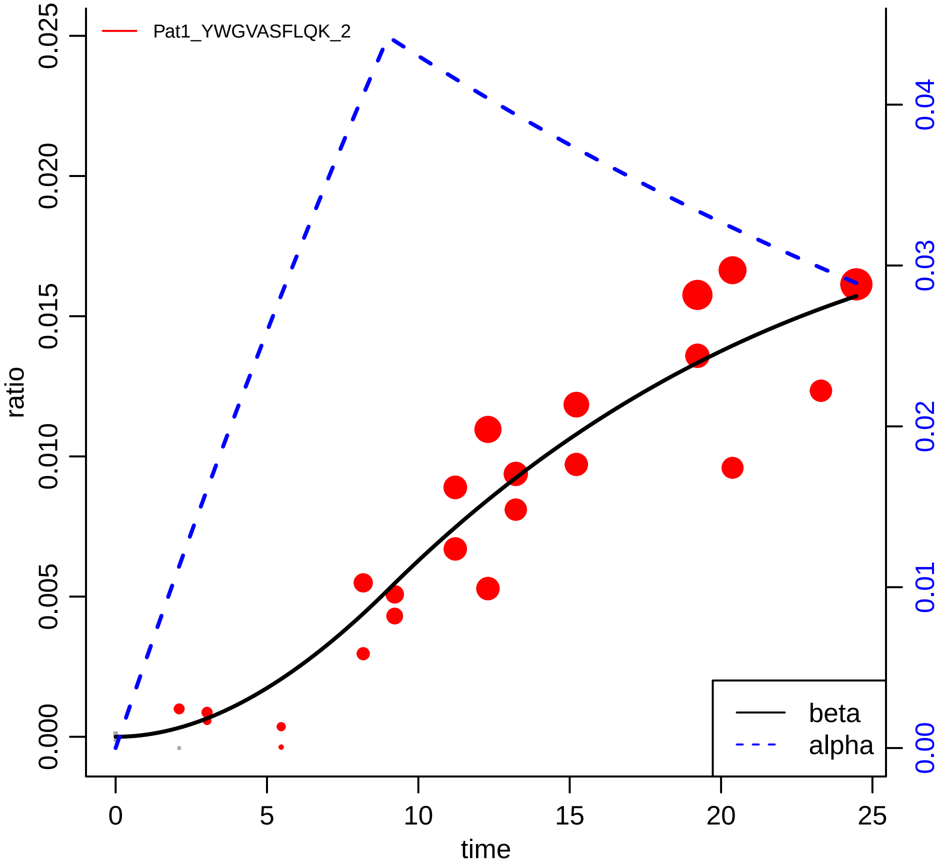

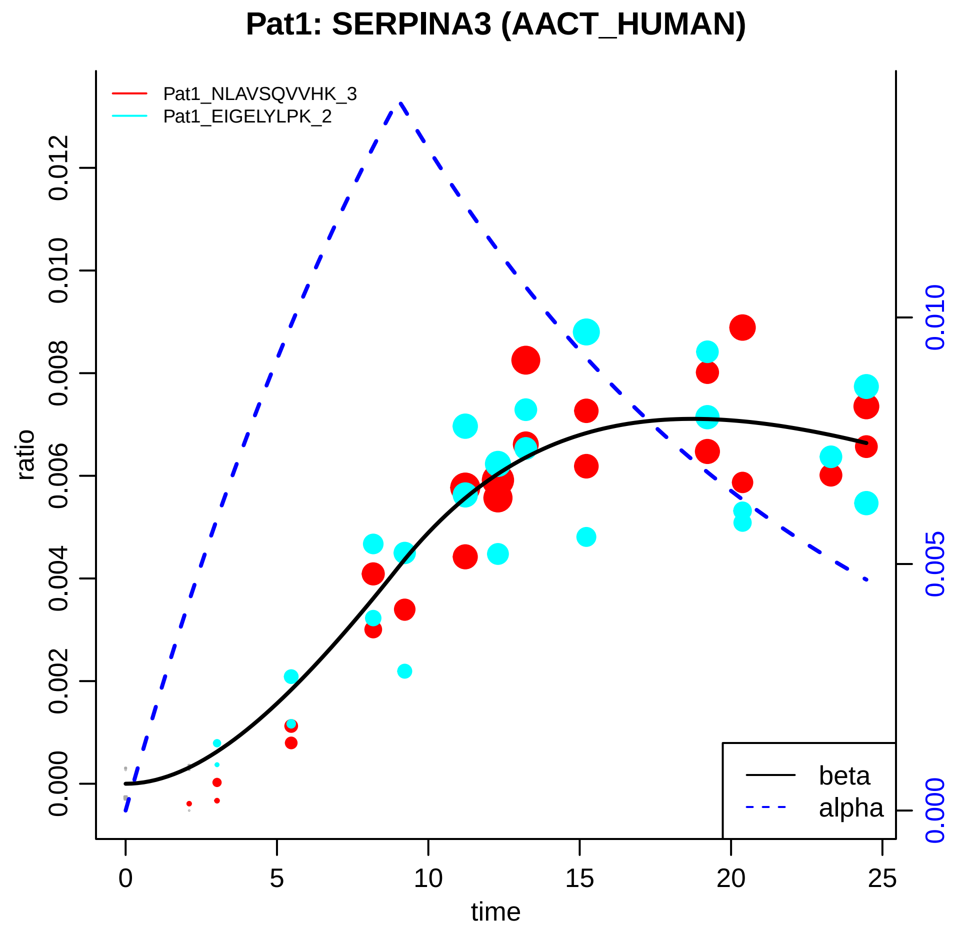

Pat1: SERPINA3 (AACT_HUMAN)
Pat1_NLAVSQVVHK_3
Pat1_EIGELYLPK_2
ratio
0.000
0.002
0.004
0.006
0.008
0.010
0.012
0.000
0.005
0.010
0
5
10
15
20
25
time
beta
alpha

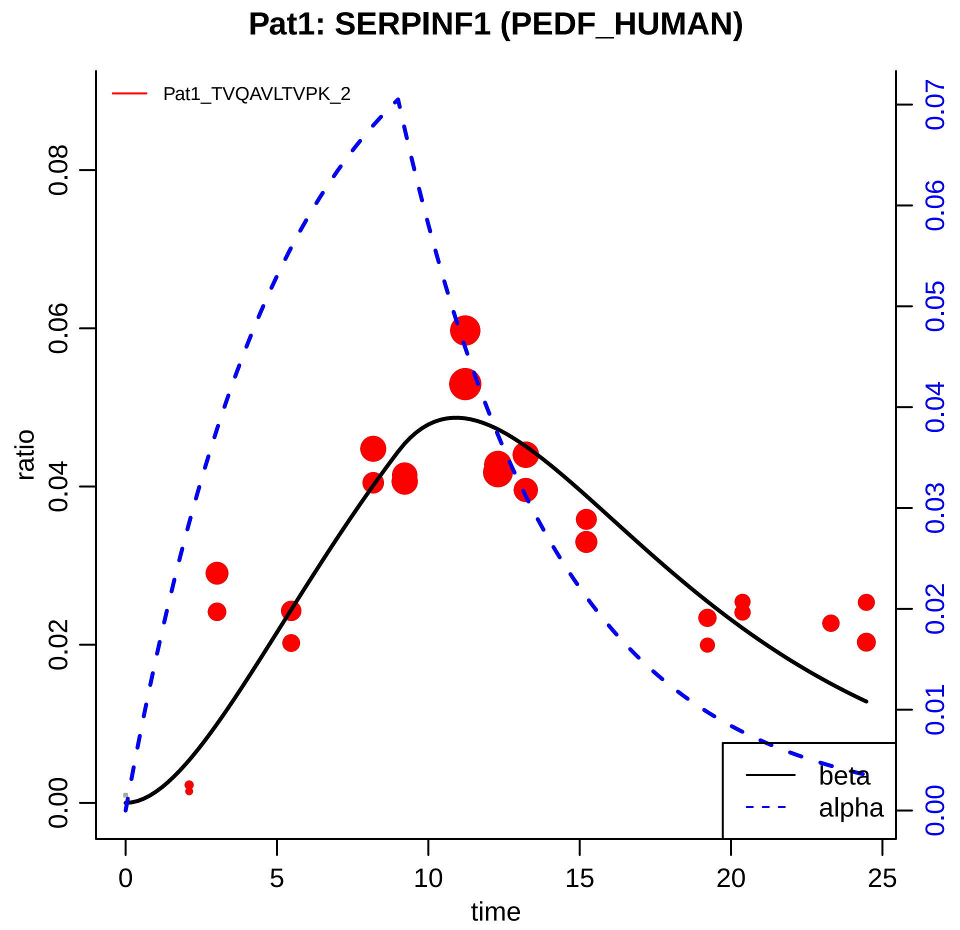

Pat1: SERPINF1 (PEDF_HUMAN)
Pat1_TVQAVLTVPK_2
ratio
time
0.00
0.02
0.04
0.06
0.08
0
5
10
15
20
25
0.00
0.01
0.02
0.03
0.04
0.05
0.06
0.07
beta
alpha

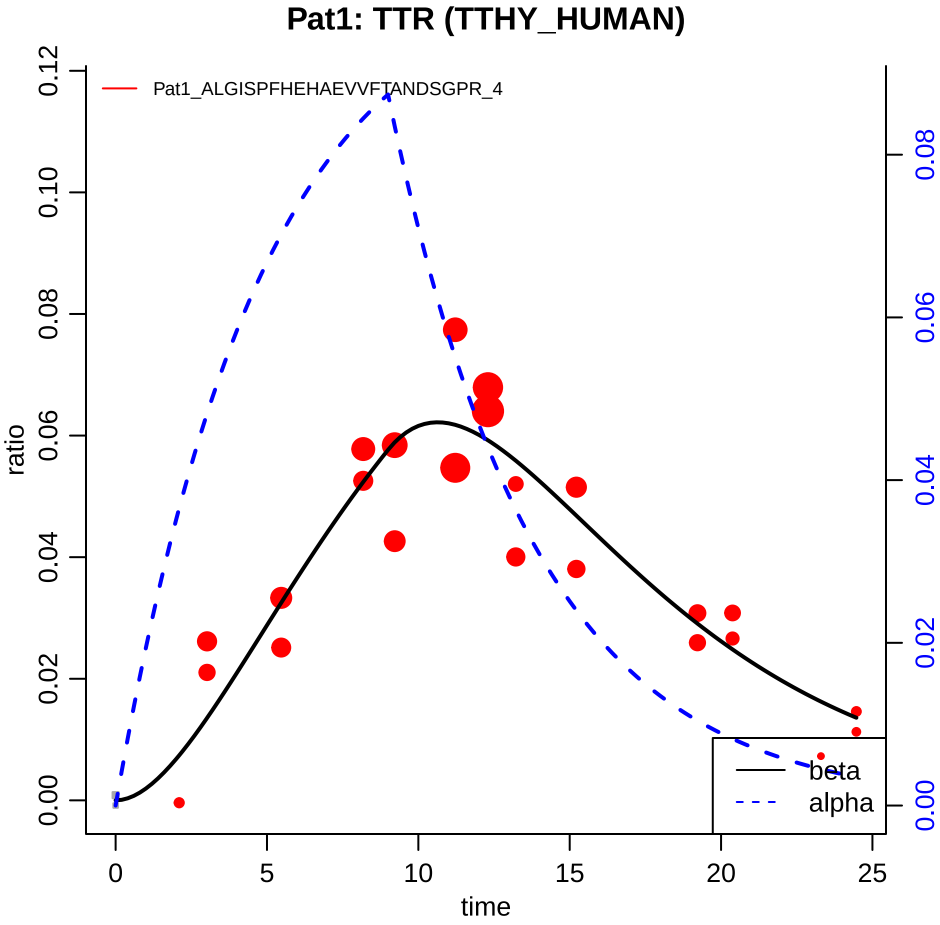

Pat1: TTR (TTHY_HUMAN)
Pat1_ALGISPFHEHAEVVFTANDSGPR_4
ratio
0.00
0.02
0.04
0.06
0.08
0.10
0.12
0.00
0.02
0.04
0.06
0.08
0
5
10
15
20
25
time
beta
alpha

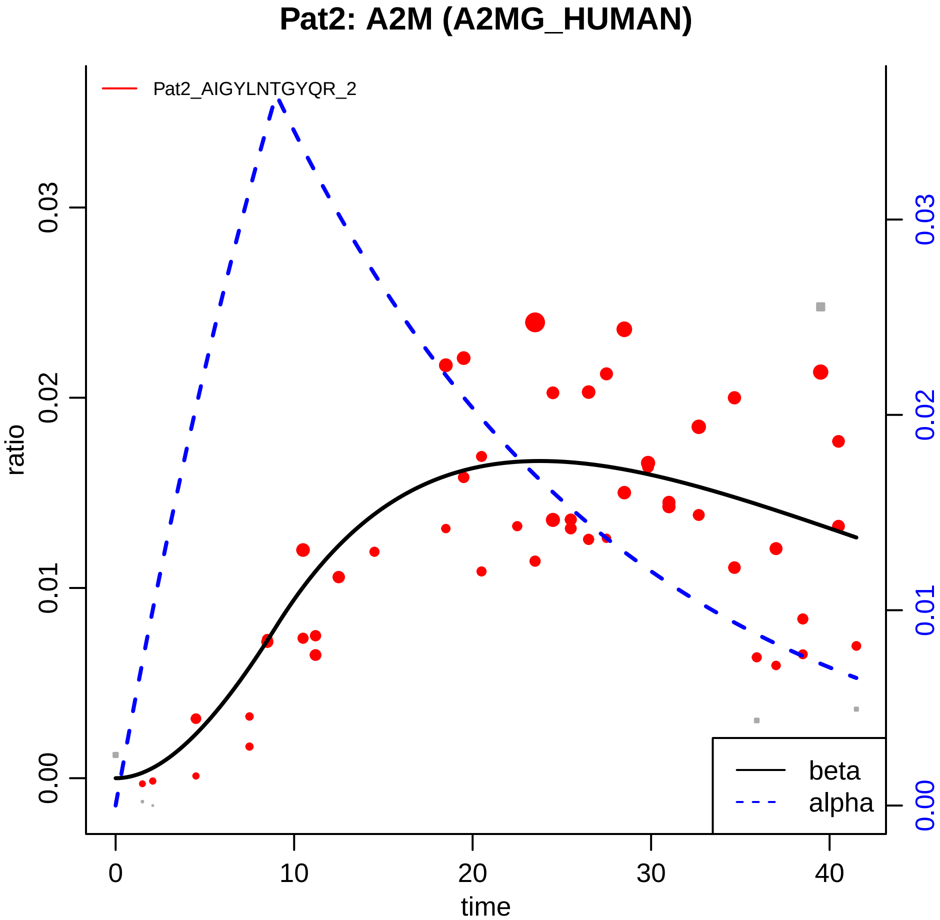
Pat2: A2M (A2MG_HUMAN)
Pat2_AIGYLNTGYQR_2
ratio
0.00
0.01
0.02
0.03
time
0
10
20
30
40
beta
alpha
0.00
0.01
0.02
0.03

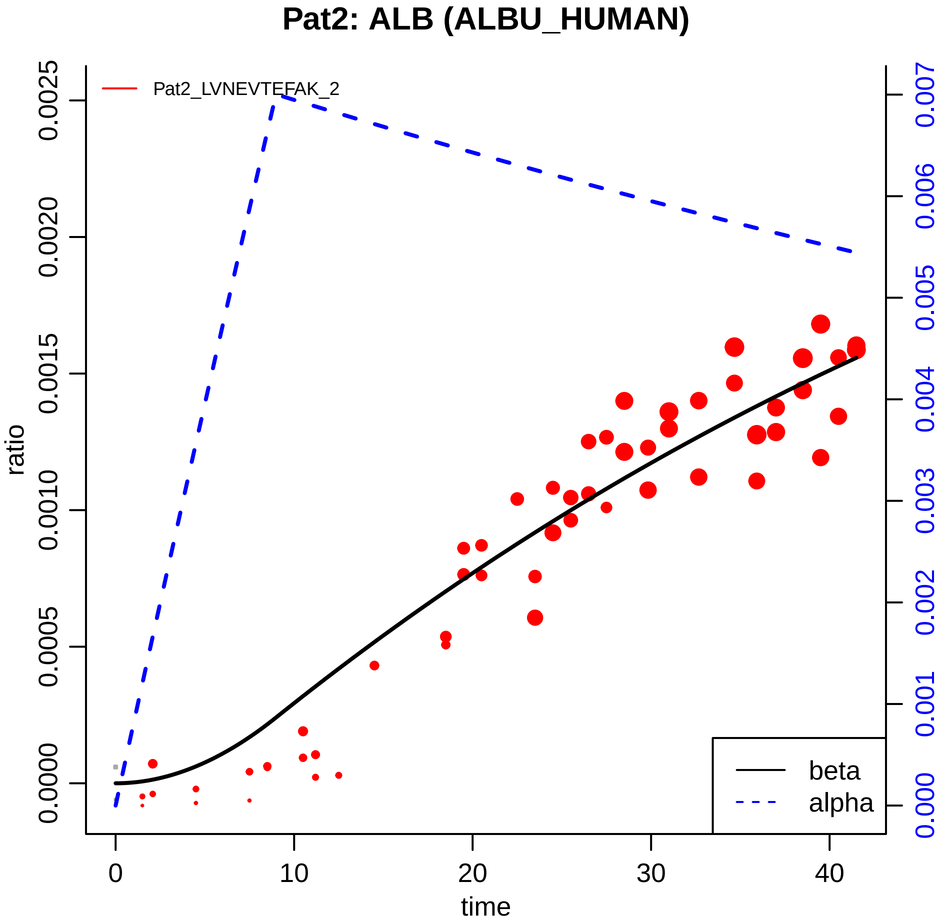

Pat2: ALB (ALBU_HUMAN)
Pat2_LVNEVTEFAK_2
ratio
0.0000
0.0005
0.0010
0.0015
0.0020
0.0025
0.000
0.001
0.002
0.003
0.004
0.005
0.006
0.007
0
10
20
30
40
time
beta
alpha

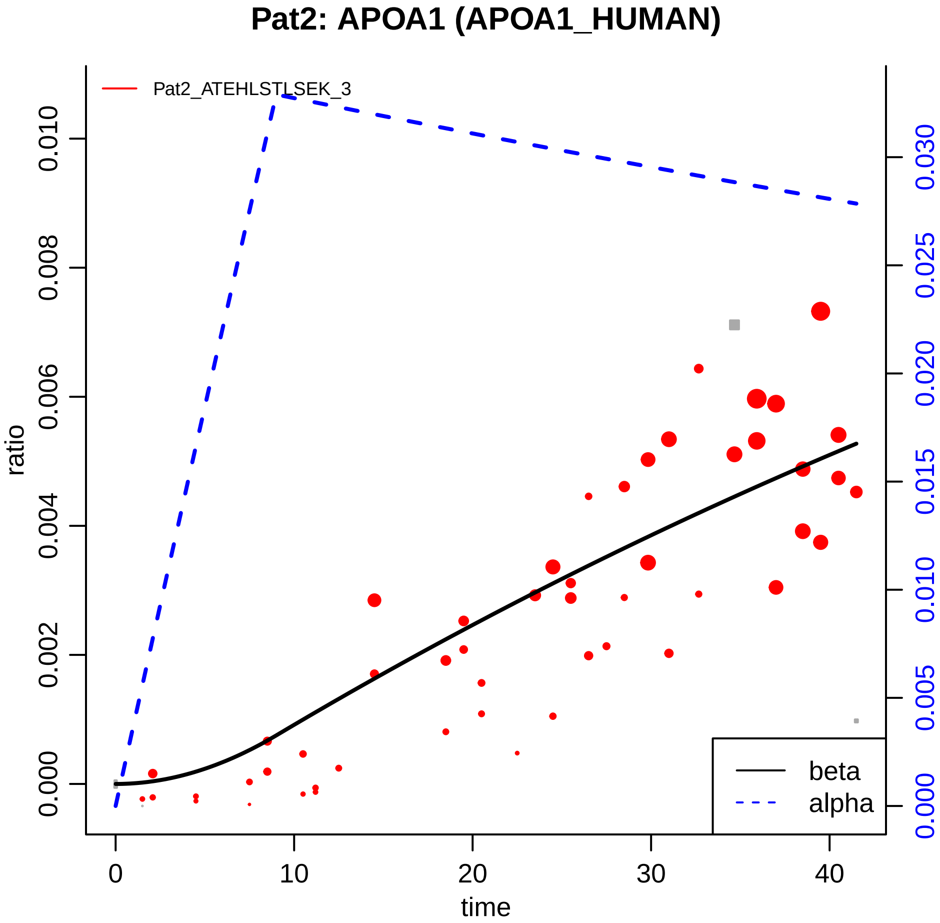

Pat2: APOA1 (APOA1_HUMAN)
Pat2_ATEHLSTLSEK_3
ratio
0.000
0.002
0.004
0.006
0.008
0.010
0.000
0.005
0.010
0.015
0.020
0.025
0.030
0
10
20
30
40
time
beta
alpha

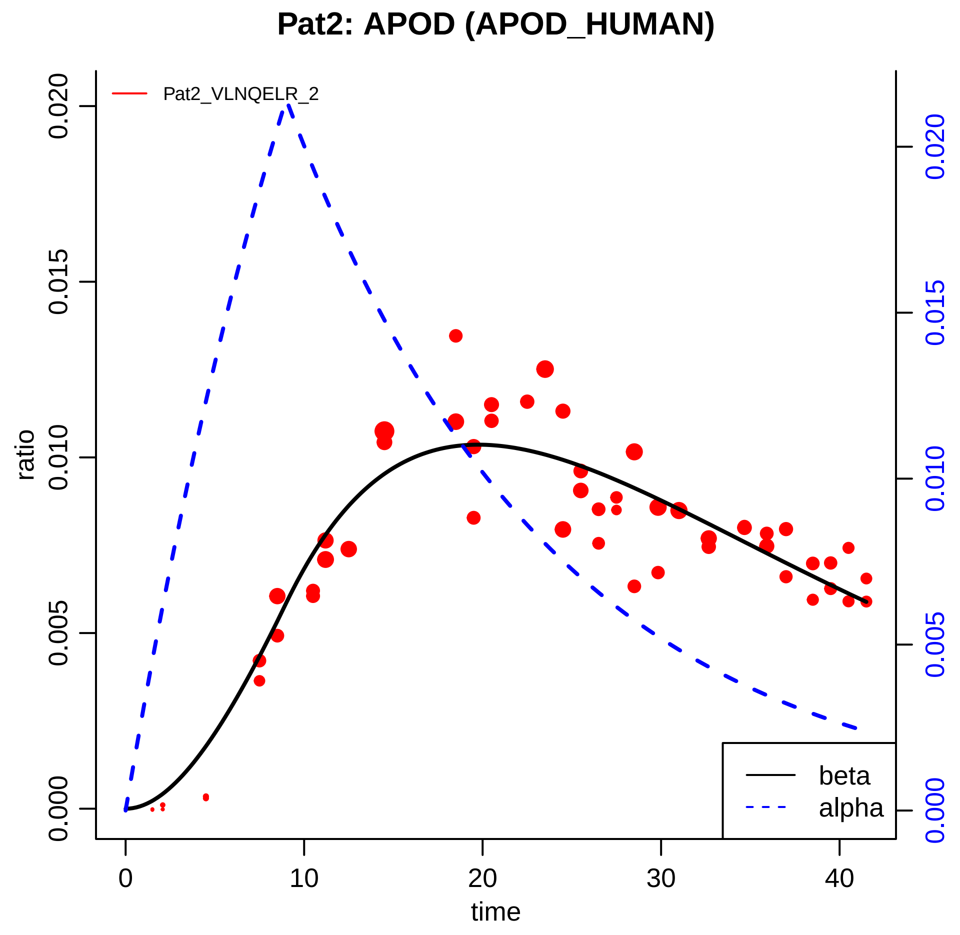

Pat2: APOD (APOD_HUMAN)
Pat2_VLNQELR_2
ratio
time
0.000
0.005
0.010
0.015
0.020
0
10
20
30
40
beta
alpha

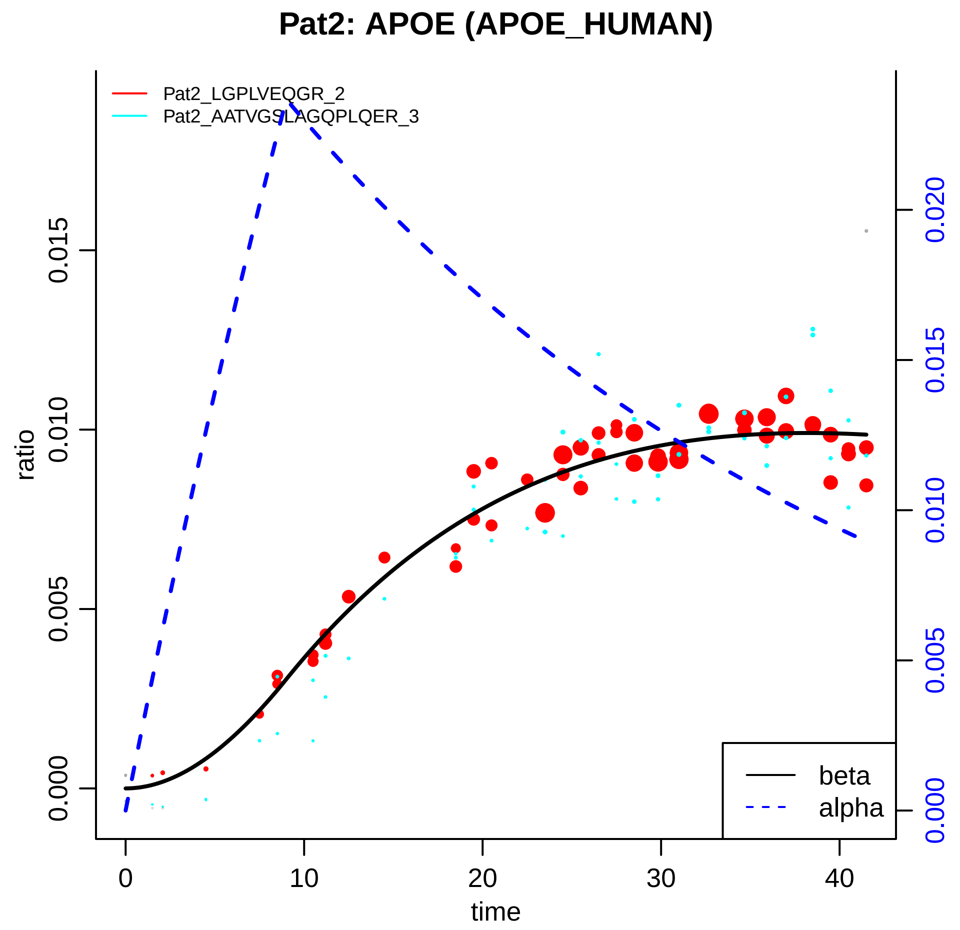

Pat2: APOE (APOE_HUMAN)
Pat2_LGPLVEQGR_2
Pat2_AATVGSLAGQPLQER_3
ratio
0.000
0.005
0.010
0.015
0.020
time
0
10
20
30
40
beta
alpha

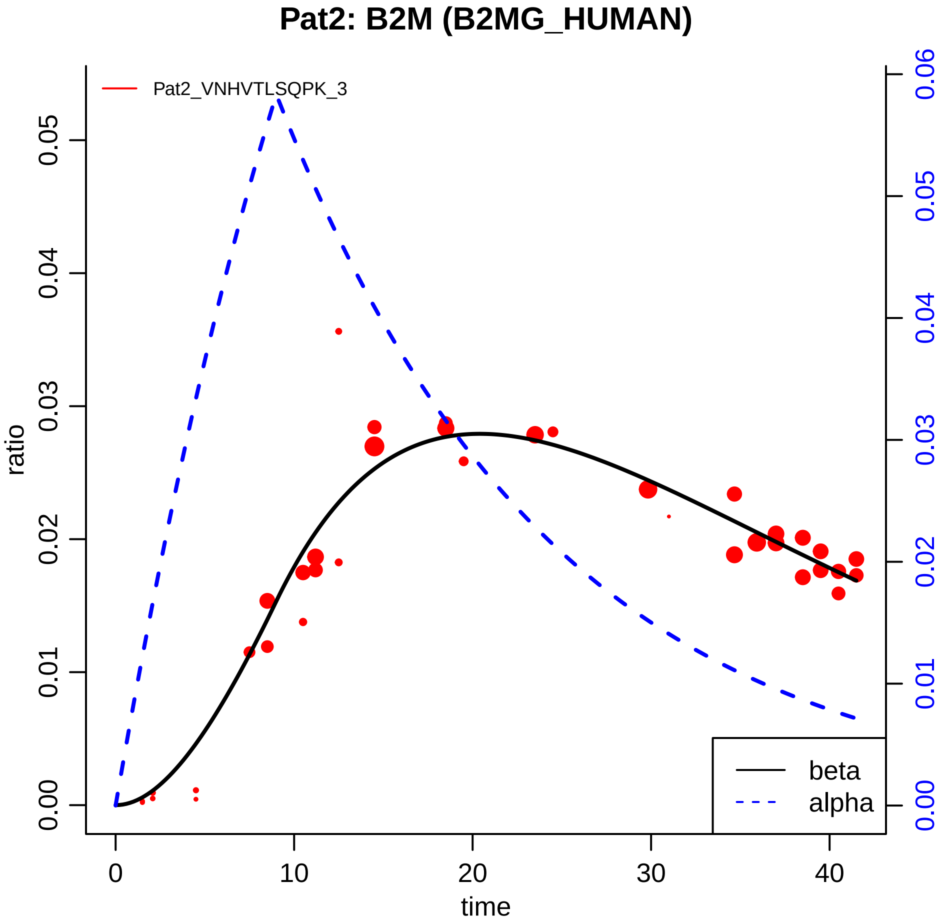

Pat2: B2M (B2MG_HUMAN)
Pat2_VNHVTLSQPK_3
ratio
time
0.00
0.01
0.02
0.03
0.04
0.05
0.06
0
10
20
30
40
beta
alpha

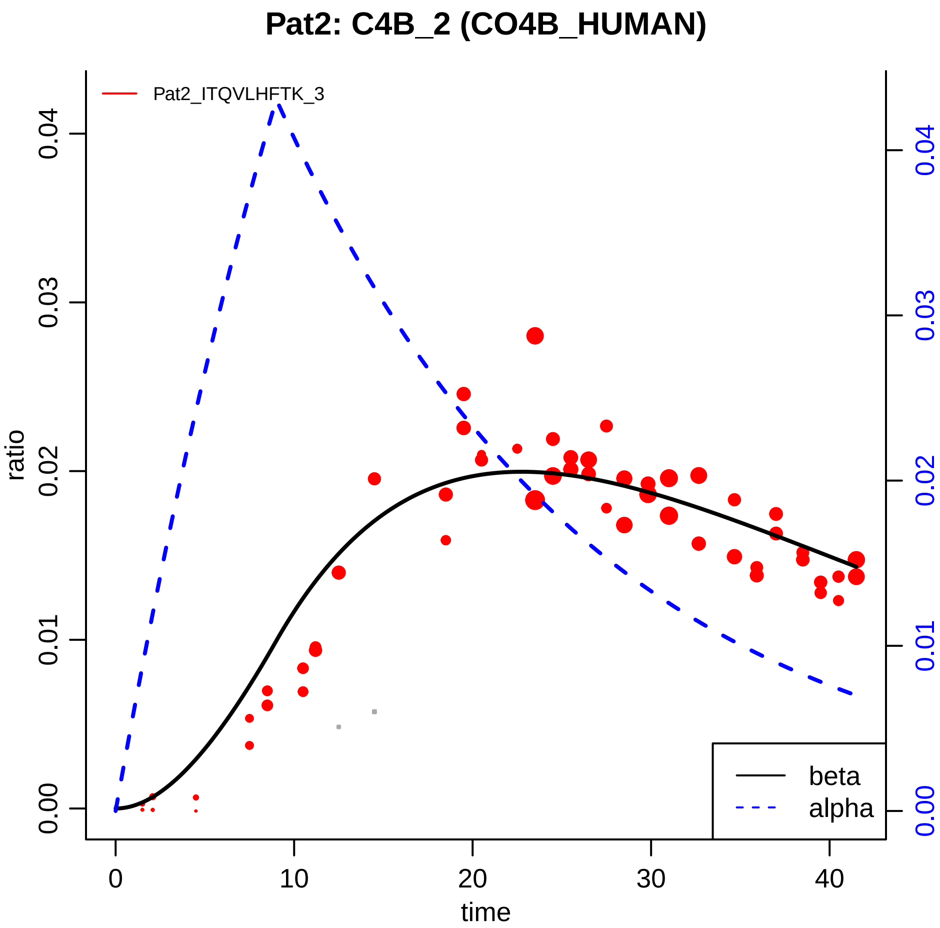

Pat2: C4B_2 (CO4B_HUMAN)
Pat2_ITQVLHFTK_3
ratio
0.00
0.01
0.02
0.03
0.04
time
0
10
20
30
40
beta
alpha

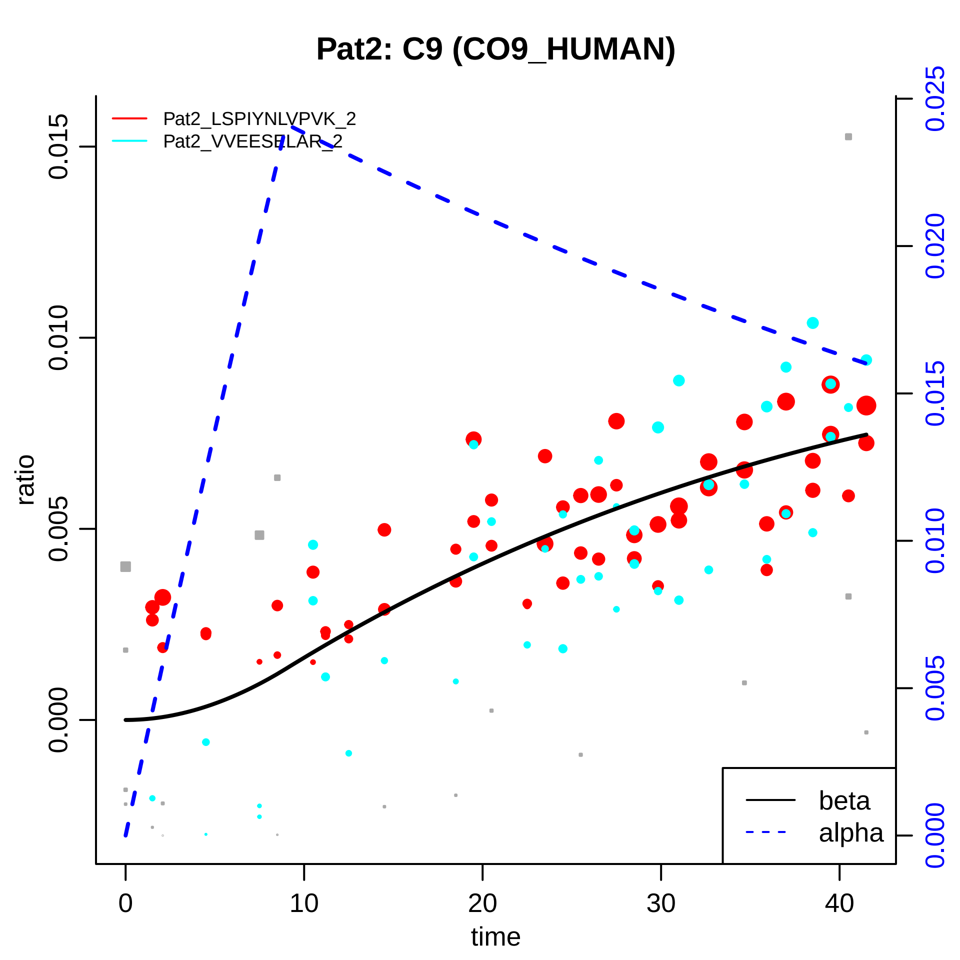

Pat2: C9 (CO9_HUMAN)
Pat2_LSPIYNLVPVK_2
Pat2_VVEESELAR_2
ratio
time
beta
alpha
0.000
0.005
0.010
0.015
0.020
0.025
0
10
20
30
40

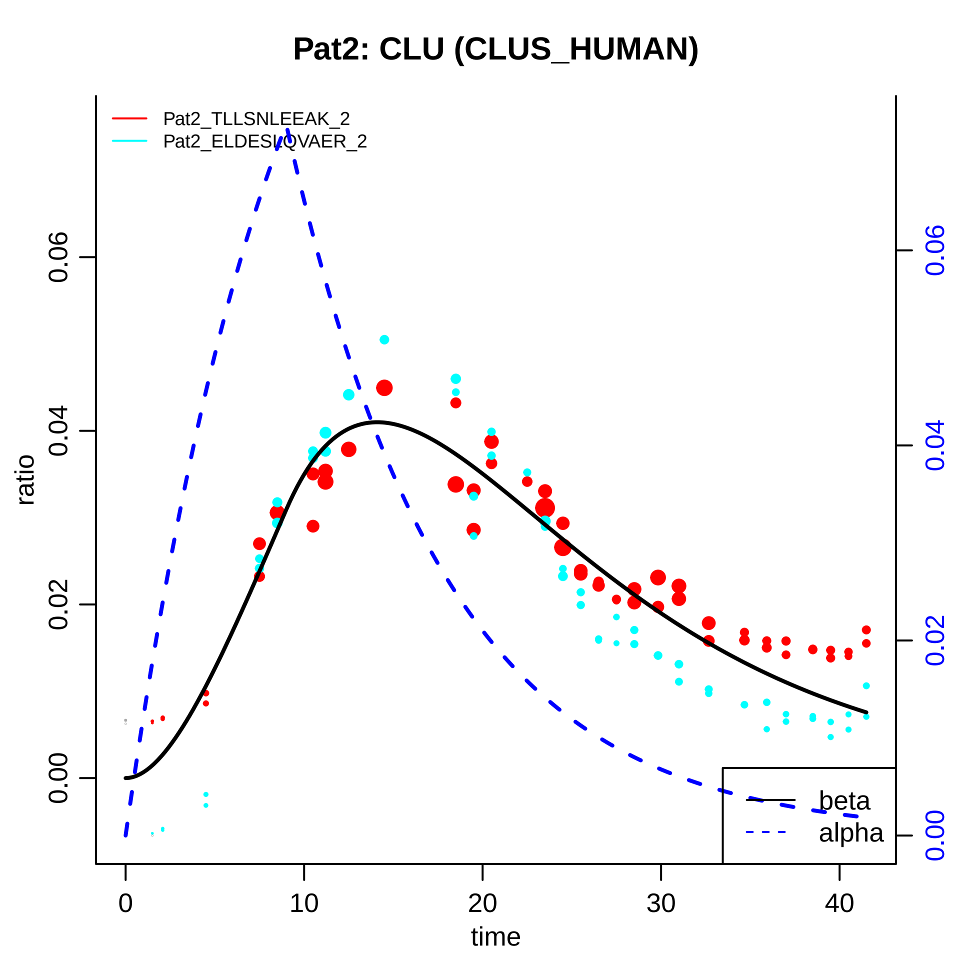

Pat2: CLU (CLUS_HUMAN)
Pat2_TLLSNLEEAK_2
Pat2_ELDESLQVAER_2
ratio
0.00
0.02
0.04
0.06
time
0
10
20
30
40
beta
alpha

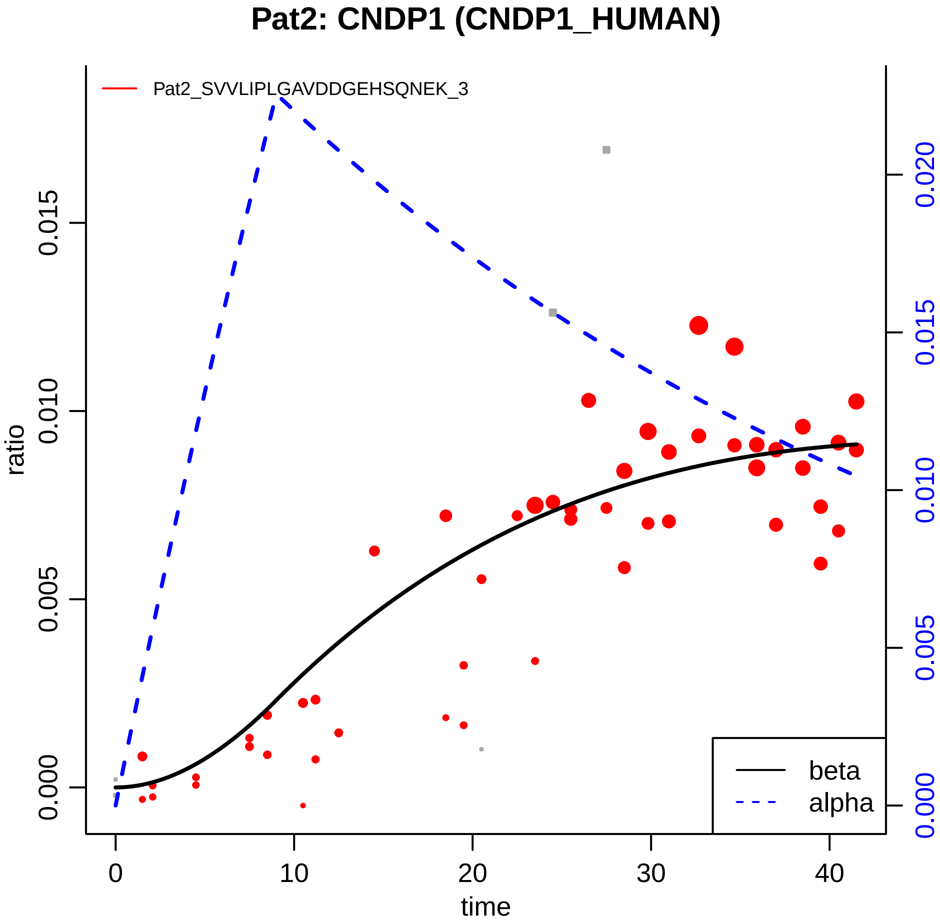
Pat2: CNDP1 (CNDP1_HUMAN)
Pat2_SVVLIPLGAVDDGEHSQNEK_3
ratio
0.000
0.005
0.010
0.015
0.020
time
0
10
20
30
40
beta
alpha

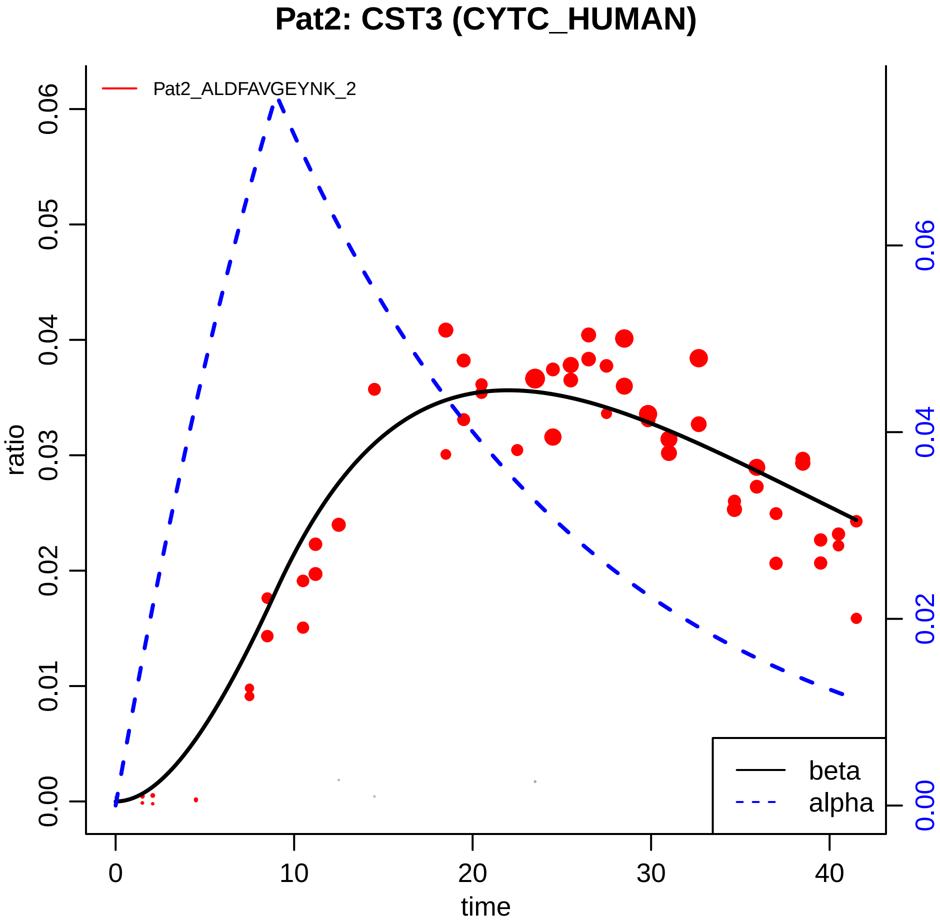

Pat2: CST3 (CYTC_HUMAN)
Pat2_ALDFAVGEYNK_2
ratio
time
beta
alpha
0.00
0.01
0.02
0.03
0.04
0.05
0.06
0
10
20
30
40
0.00
0.02
0.04
0.06

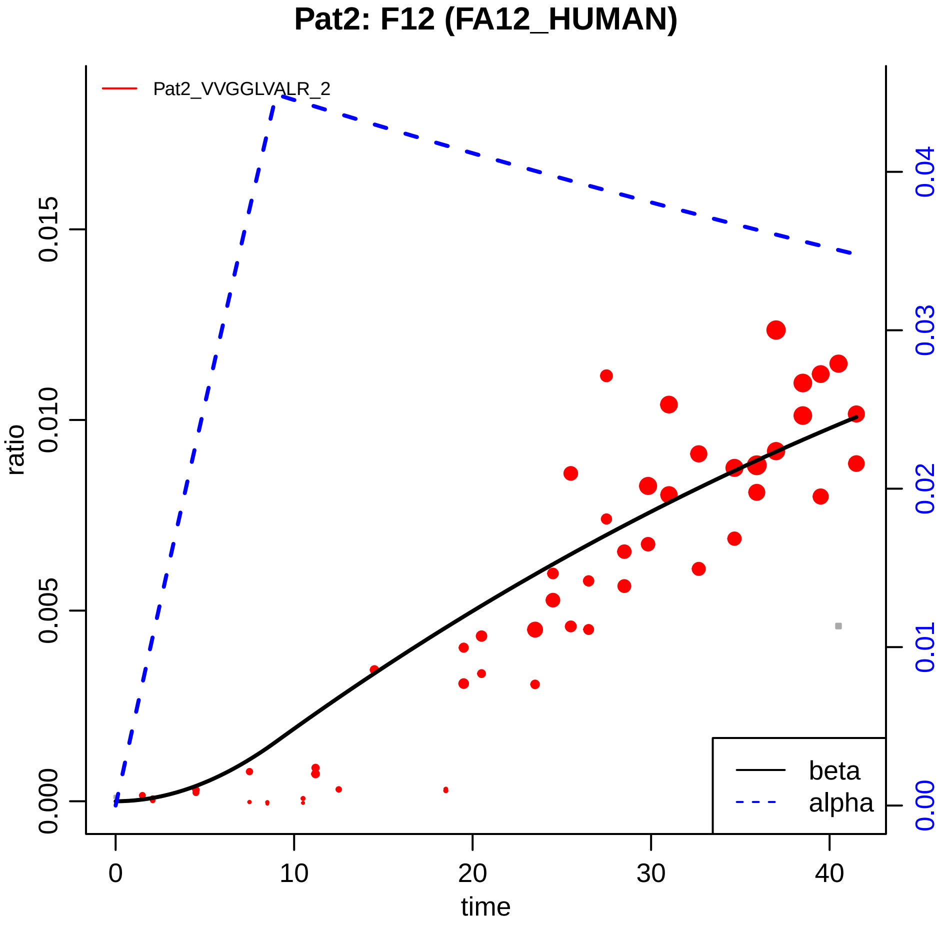
Pat2: F12 (FA12_HUMAN)
Pat2_VVGGLVALR_2
ratio
0.000
0.005
0.010
0.015
0.00
0.01
0.02
0.03
0.04
time
0
10
20
30
40
beta
alpha

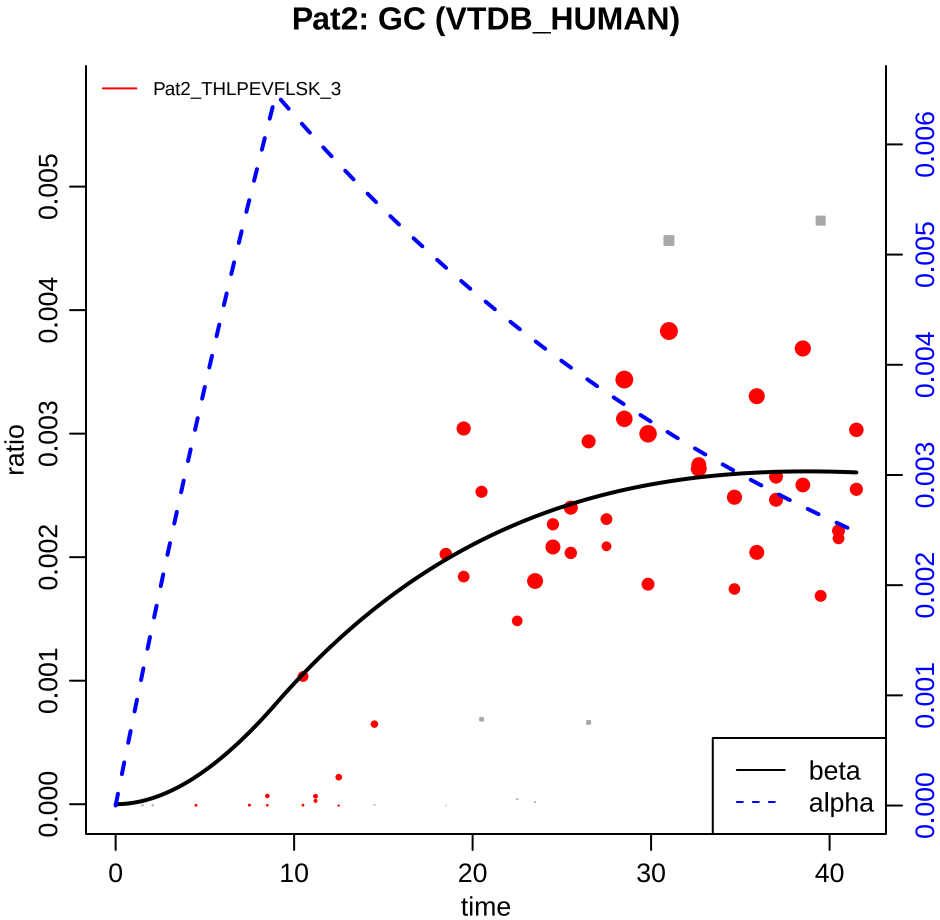

Pat2: GC (VTDB_HUMAN)
Pat2_THLPEVFLSK_3
ratio
time
0.000
0.001
0.002
0.003
0.004
0.005
0.006
0
10
20
30
40
beta
alpha

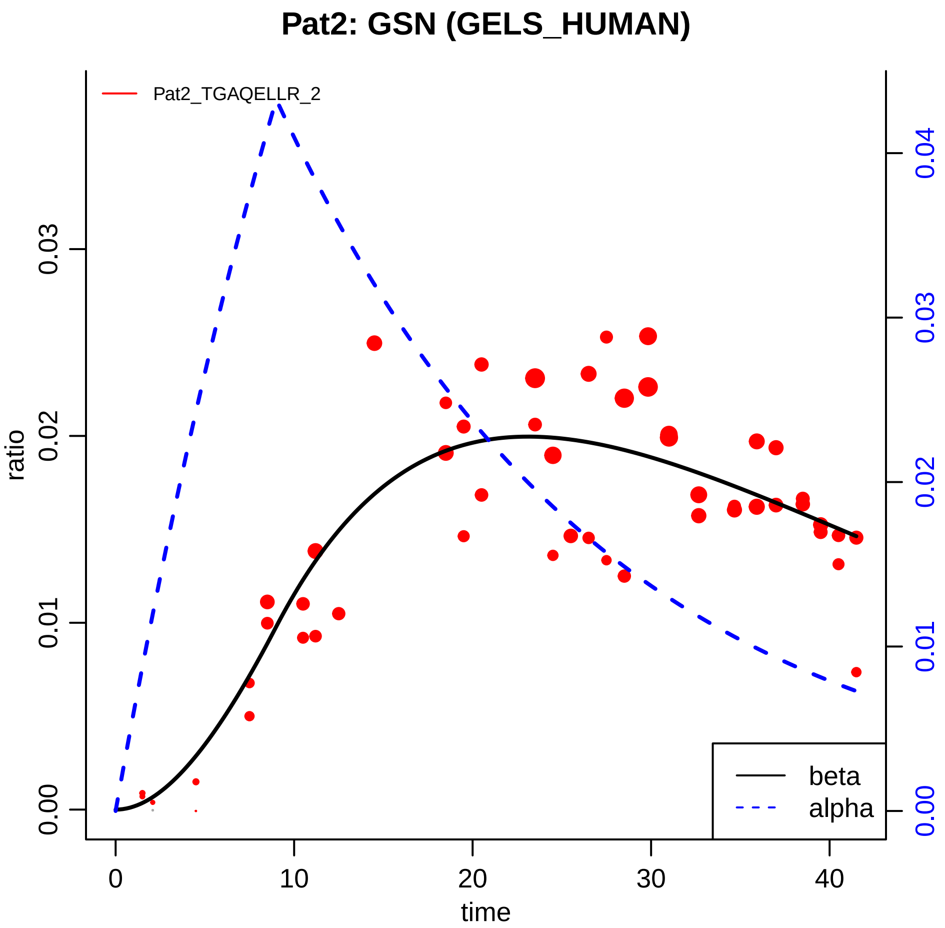

Pat2: GSN (GELS_HUMAN)
Pat2_TGAQELLR_2
ratio
time
0.00
0.01
0.02
0.03
0.04
0
10
20
30
40
beta
alpha

### Pat2: NRP2 (NRP2_HUMAN)

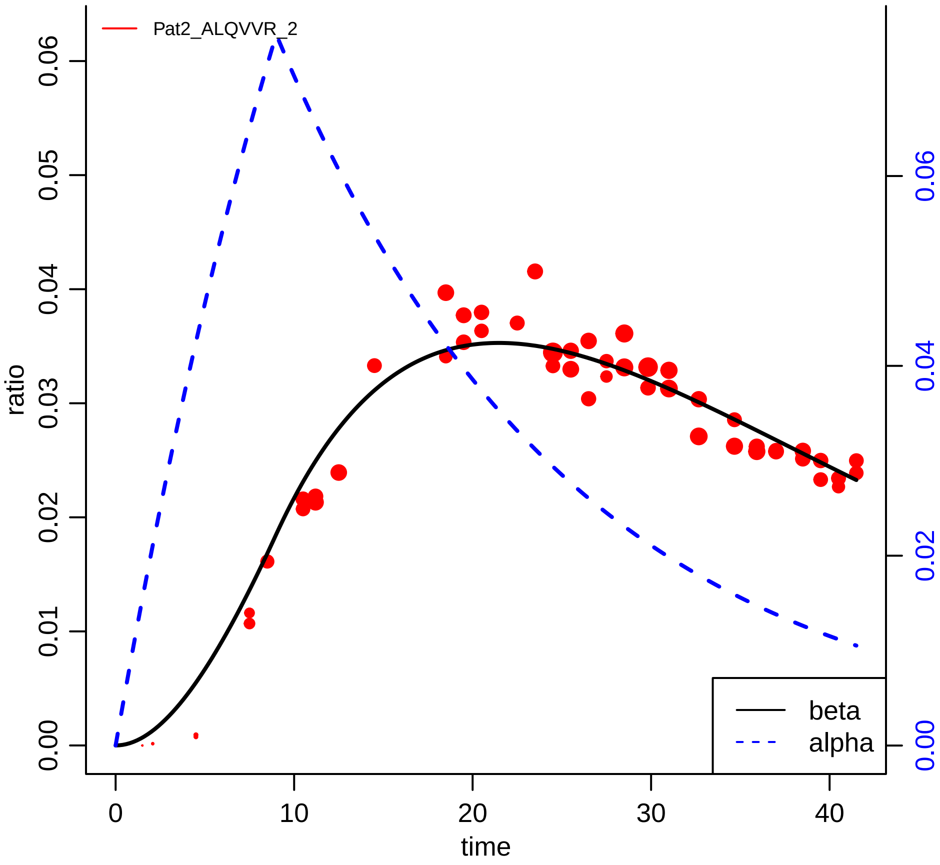

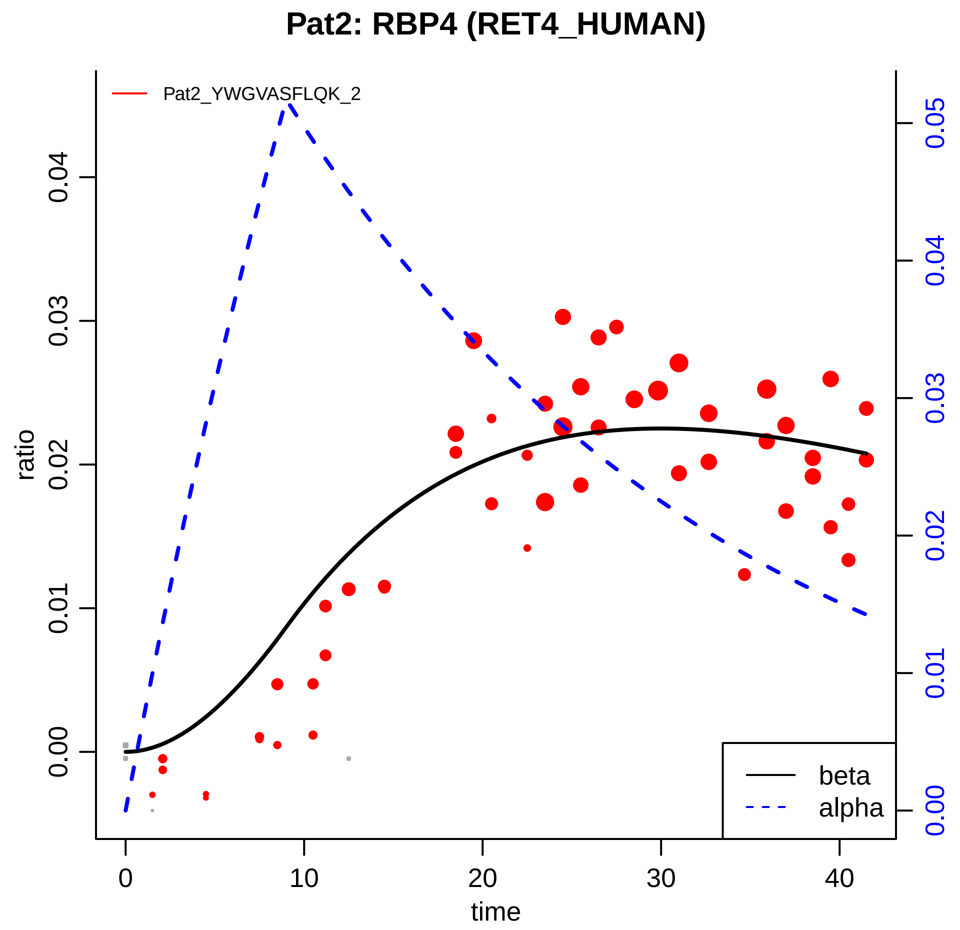

Pat2: RBP4 (RET4_HUMAN)
Pat2_YWGVASFLQK_2
ratio
time
0.00
0.01
0.02
0.03
0.04
0
10
20
30
40
0.00
0.01
0.02
0.03
0.04
0.05
beta
alpha

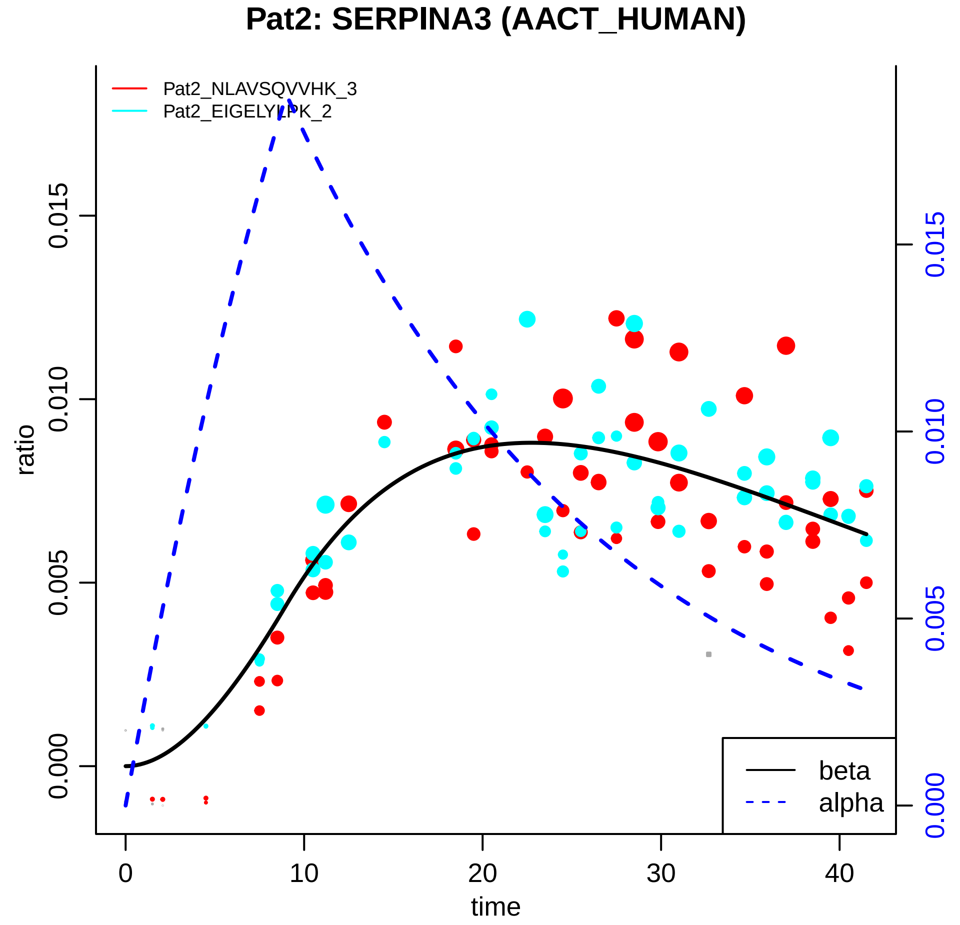

Pat2: SERPINA3 (AACT_HUMAN)
Pat2_NLAVSQVVHK_3
Pat2_EIGELYLPK_2
ratio
0.000
0.005
0.010
0.015
time
0
10
20
30
40
beta
alpha

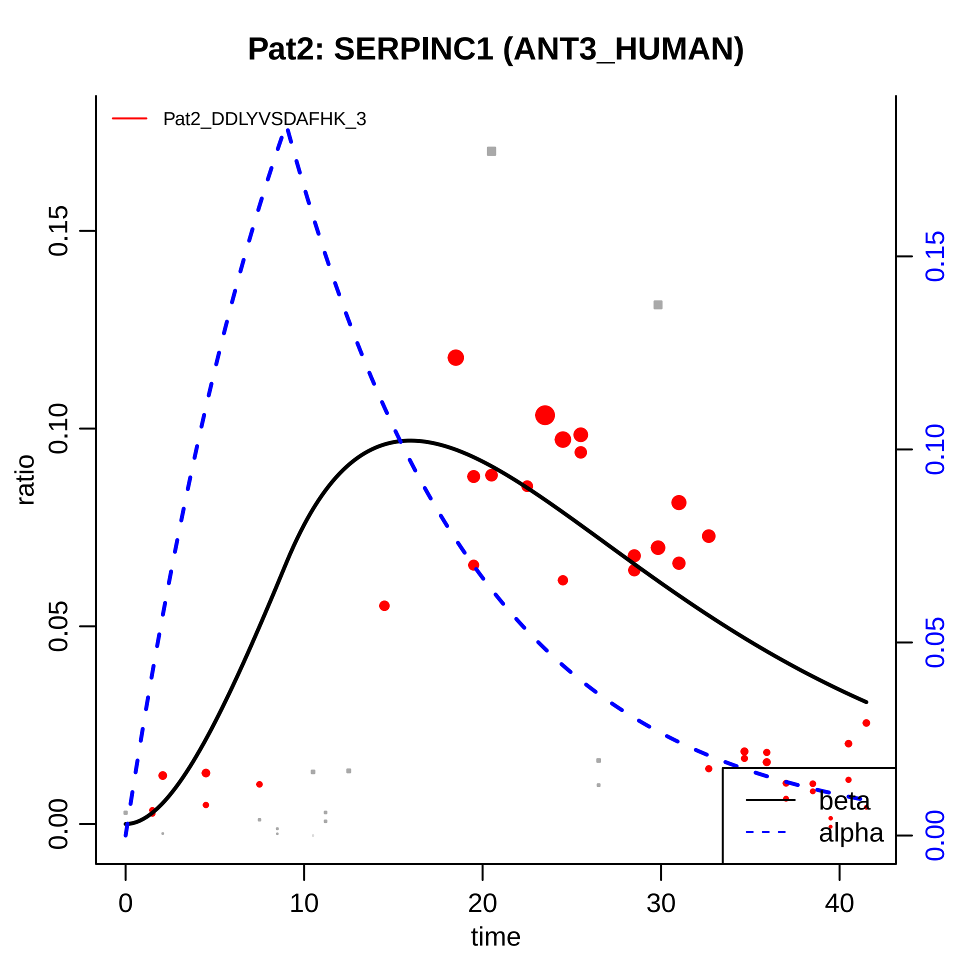

Pat2: SERPINC1 (ANT3_HUMAN)
Pat2_DDLYVSDAFHK_3
ratio
0.00
0.05
0.10
0.15
time
0
10
20
30
40
beta
alpha

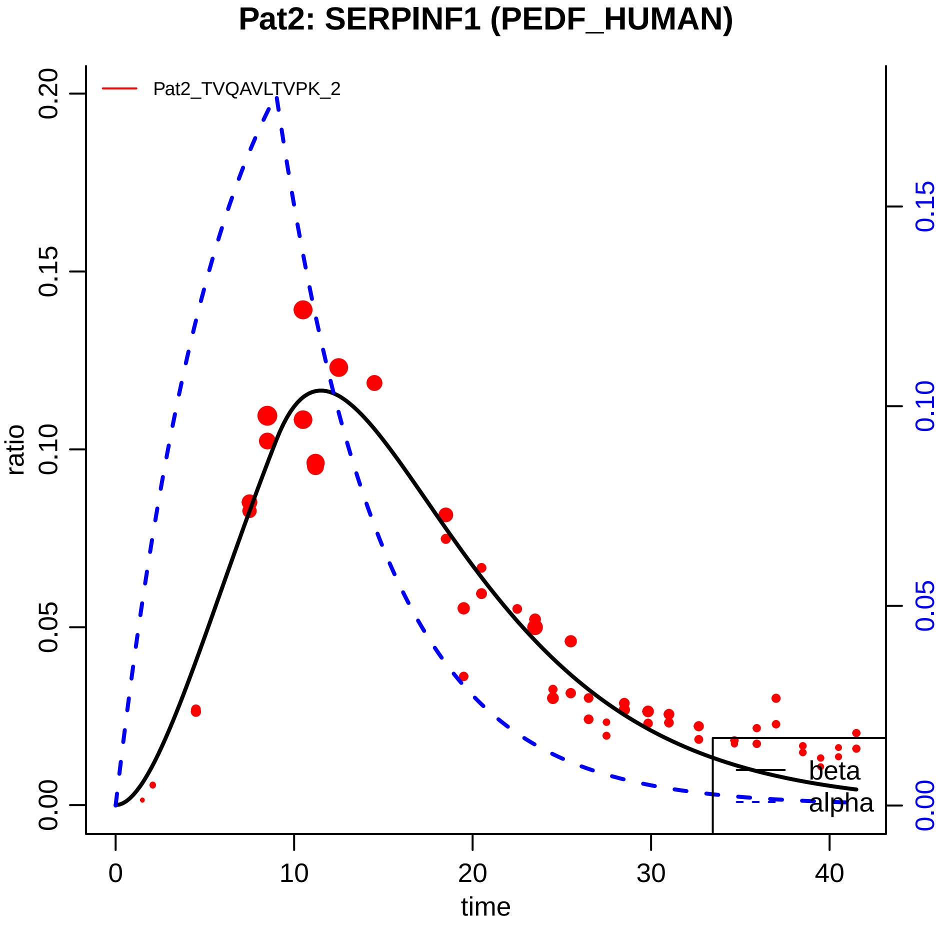

Pat2: SERPINF1 (PEDF_HUMAN)
Pat2_TVQAVLTVPK_2
ratio
0.00
0.05
0.10
0.15
0.20
time
0
10
20
30
40
0.00
0.05
0.10
0.15
beta
alpha

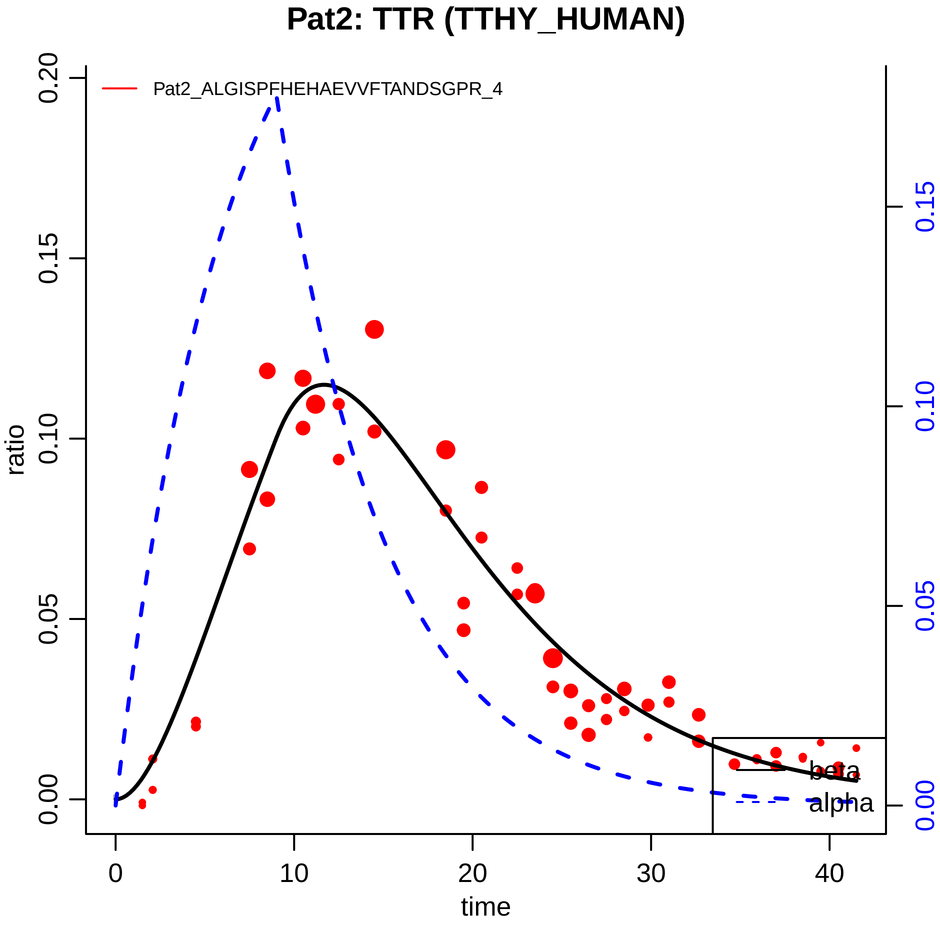

Pat2: TTR (TTHY_HUMAN)
Pat2_ALGISPFHEHAEVVFTANDSGPR_4
ratio
0.00
0.05
0.10
0.15
0.20
time
0
10
20
30
40
beta
alpha

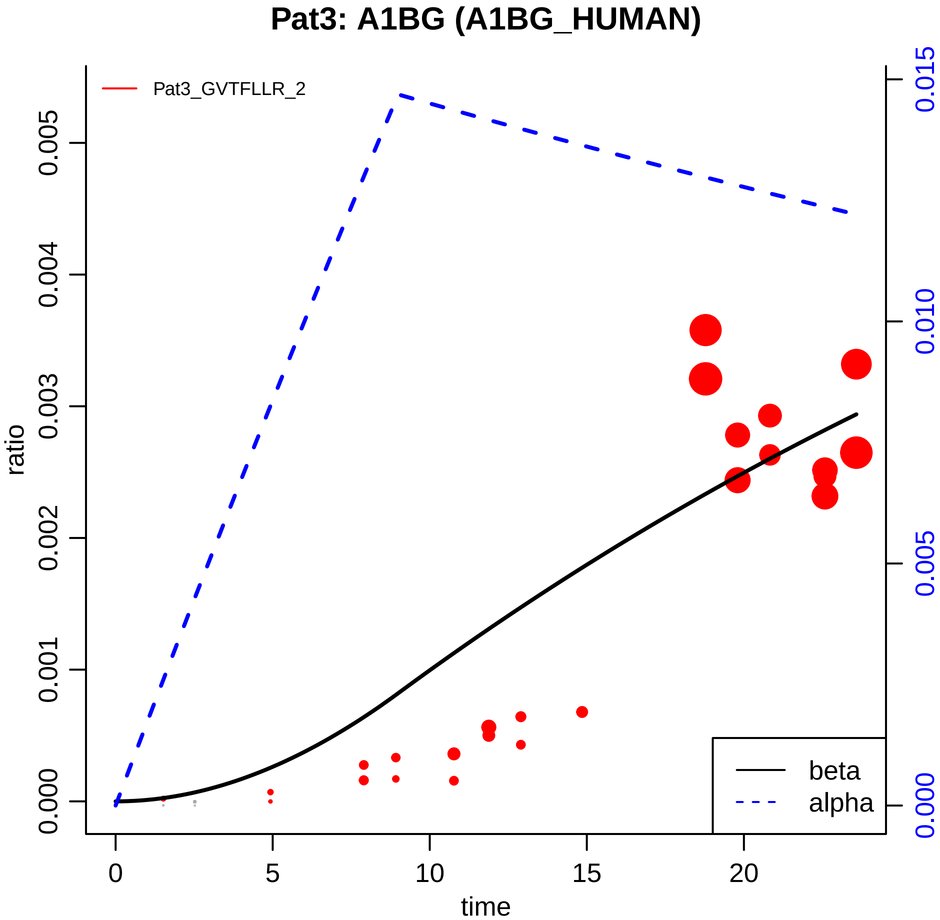

Pat3: A1BG (A1BG_HUMAN)
Pat3_GVTFLLR_2
ratio
0.000
0.001
0.002
0.003
0.004
0.005
0.000
0.005
0.010
0.015
0
5
10
15
20
time
beta
alpha

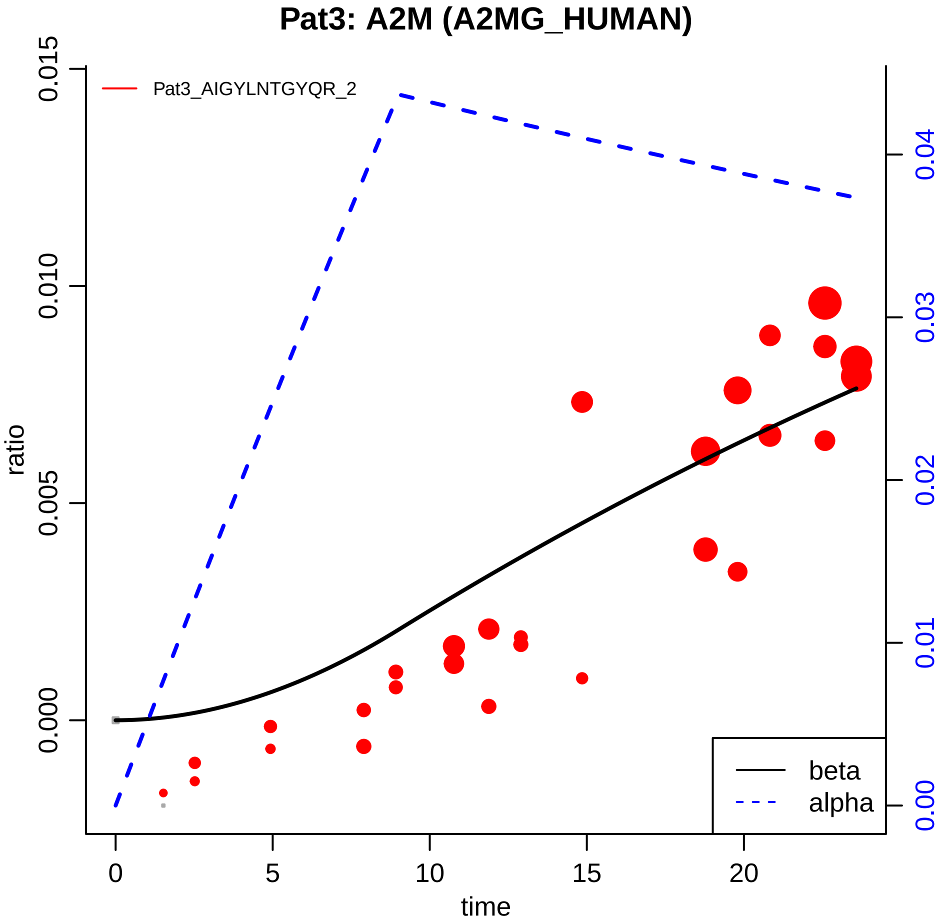

Pat3: A2M (A2MG_HUMAN)
Pat3_AIGYLNTGYQR_2
ratio
0.000
0.005
0.010
0.015
time
0
5
10
15
20
0.00
0.01
0.02
0.03
0.04
beta
alpha

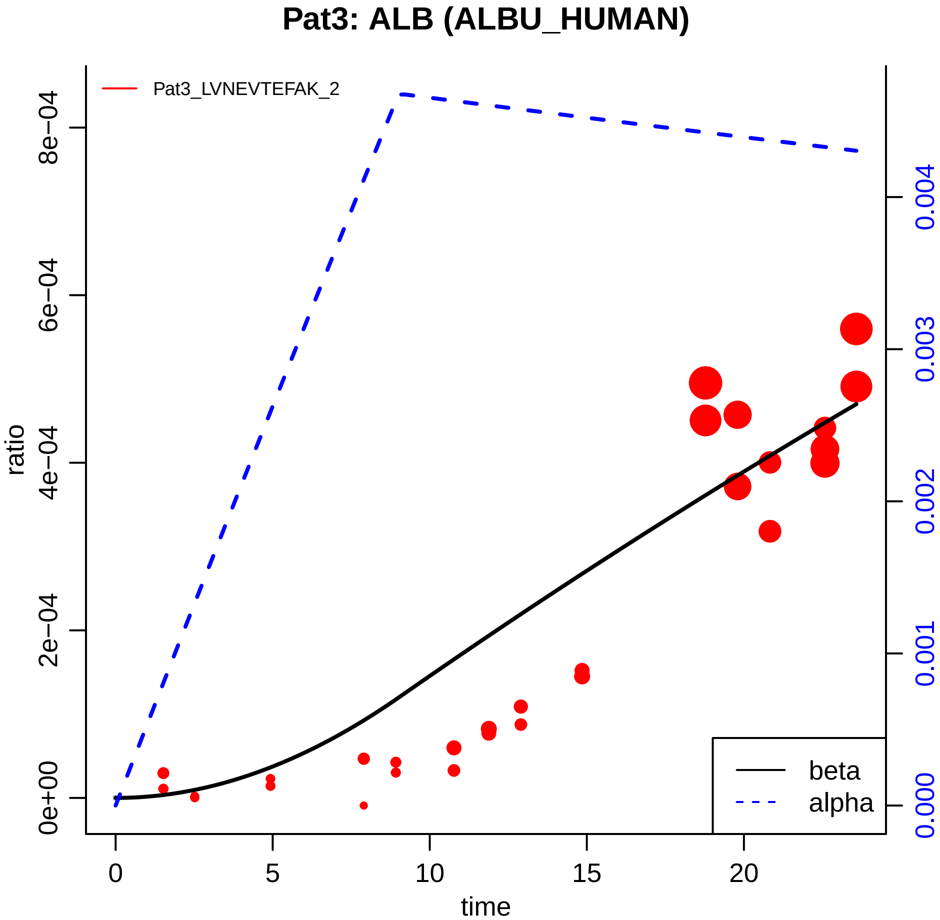

Pat3: ALB (ALBU_HUMAN)
Pat3_LVNEVTEFAK_2
ratio
0e+00
2e−04
4e−04
6e−04
8e−04
0.000
0.001
0.002
0.003
0.004
0
5
10
15
20
time
beta
alpha

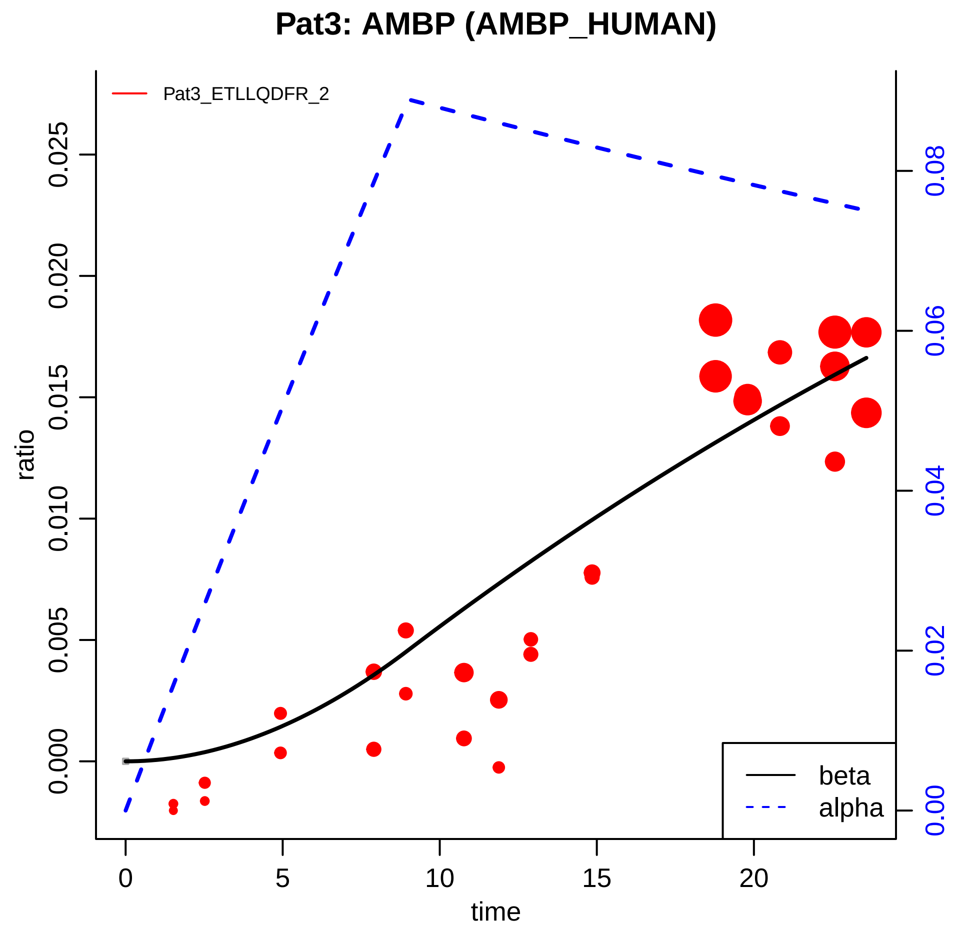

Pat3: AMBP (AMBP_HUMAN)
Pat3_ETLLQDFR_2
ratio
0.000
0.005
0.010
0.015
0.020
0.025
time
0
5
10
15
20
0.00
0.02
0.04
0.06
0.08
beta
alpha

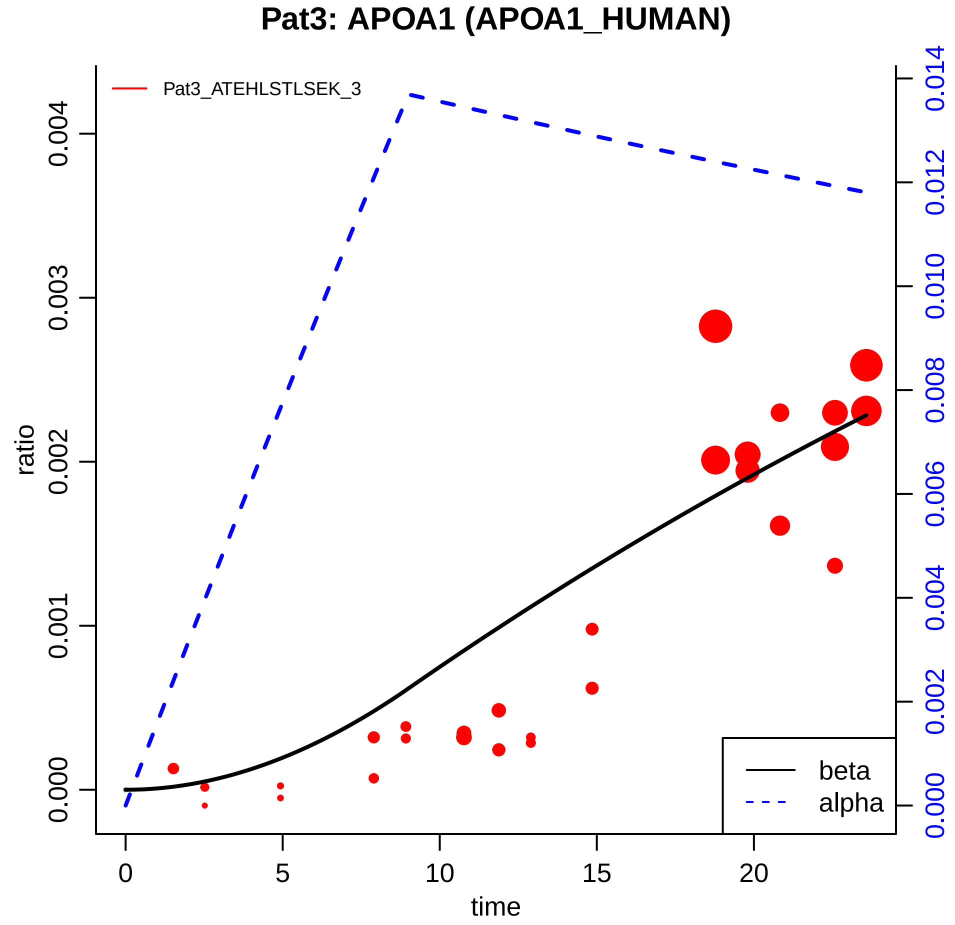

Pat3: APOA1 (APOA1_HUMAN)
Pat3_ATEHLSTLSEK_3
ratio
0.000
0.001
0.002
0.003
0.004
0.000
0.002
0.004
0.006
0.008
0.010
0.012
0.014
0
5
10
15
20
time
beta
alpha

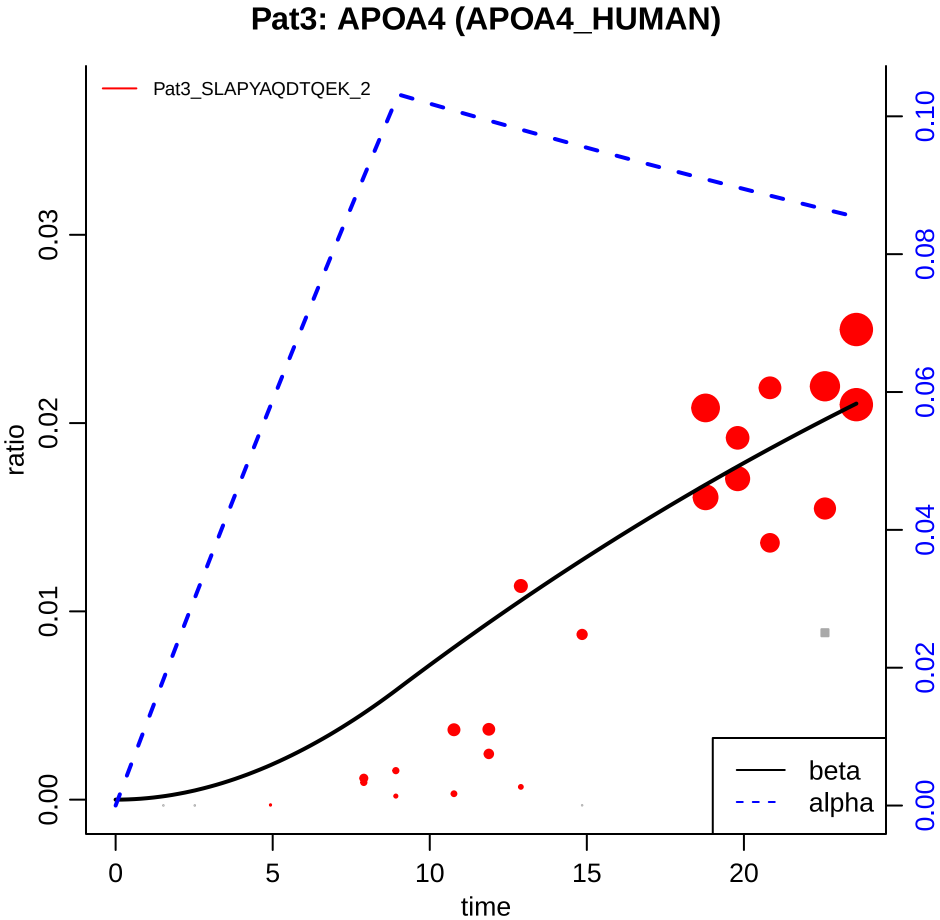

Pat3: APOA4 (APOA4_HUMAN)
Pat3_SLAPYAQDTQEK_2
ratio
0.00
0.01
0.02
0.03
0.00
0.02
0.04
0.06
0.08
0.10
0
5
10
15
20
time
beta
alpha

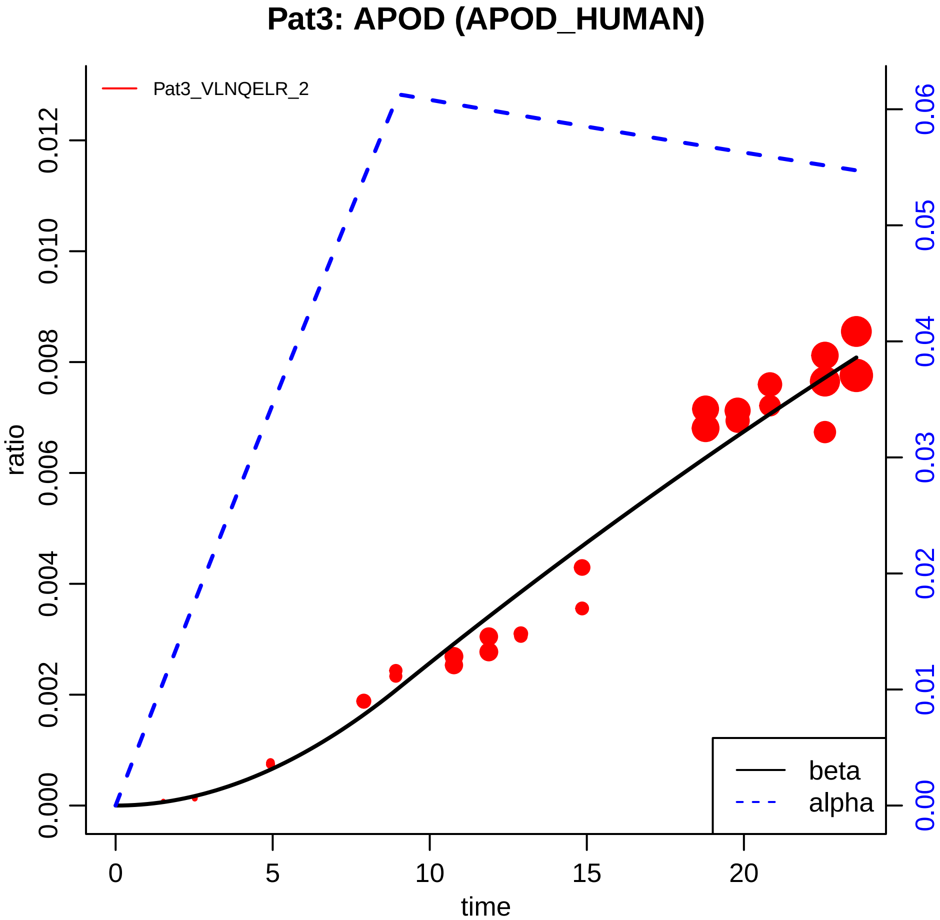

Pat3: APOD (APOD_HUMAN)
Pat3_VLNQELR_2
ratio
0.000
0.002
0.004
0.006
0.008
0.010
0.012
0.00
0.01
0.02
0.03
0.04
0.05
0.06
0
5
10
15
20
time
beta
alpha

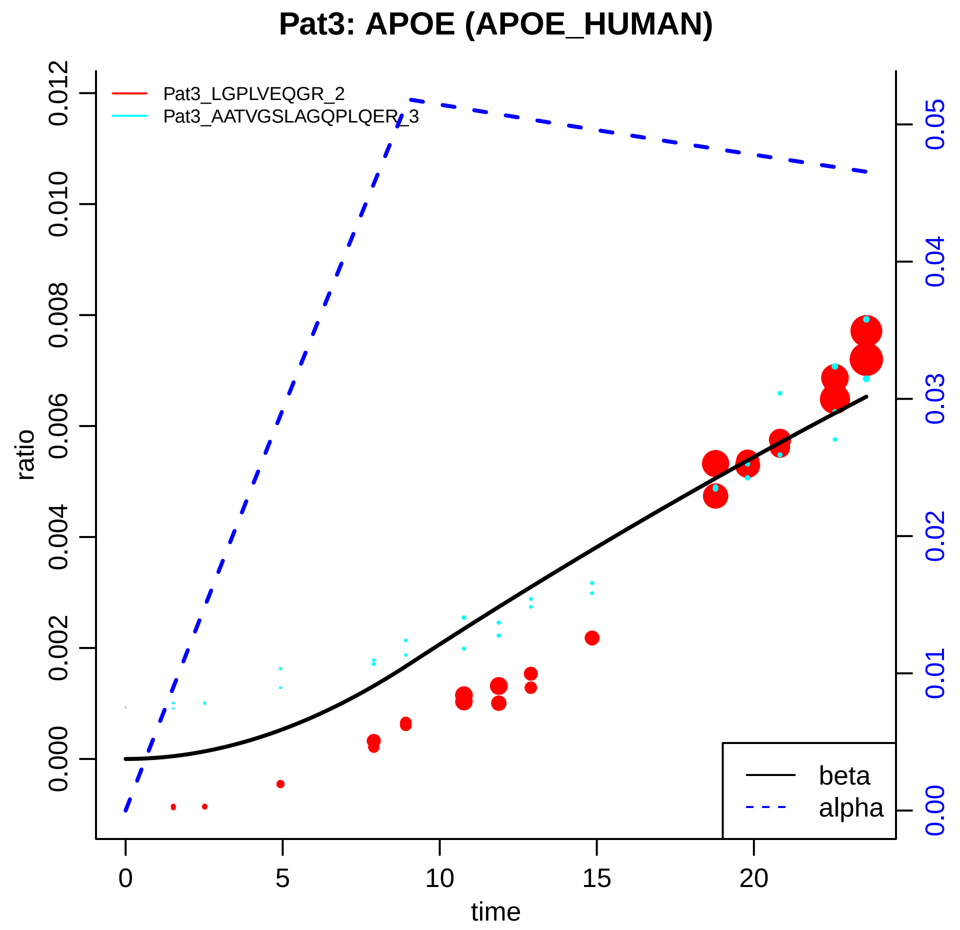

Pat3: APOE (APOE_HUMAN)
Pat3_LGPLVEQGR_2
Pat3_AATVGSLAGQPLQER_3
ratio
time
beta
alpha
0.000
0.002
0.004
0.006
0.008
0.010
0.012
0
5
10
15
20
0.00
0.01
0.02
0.03
0.04
0.05

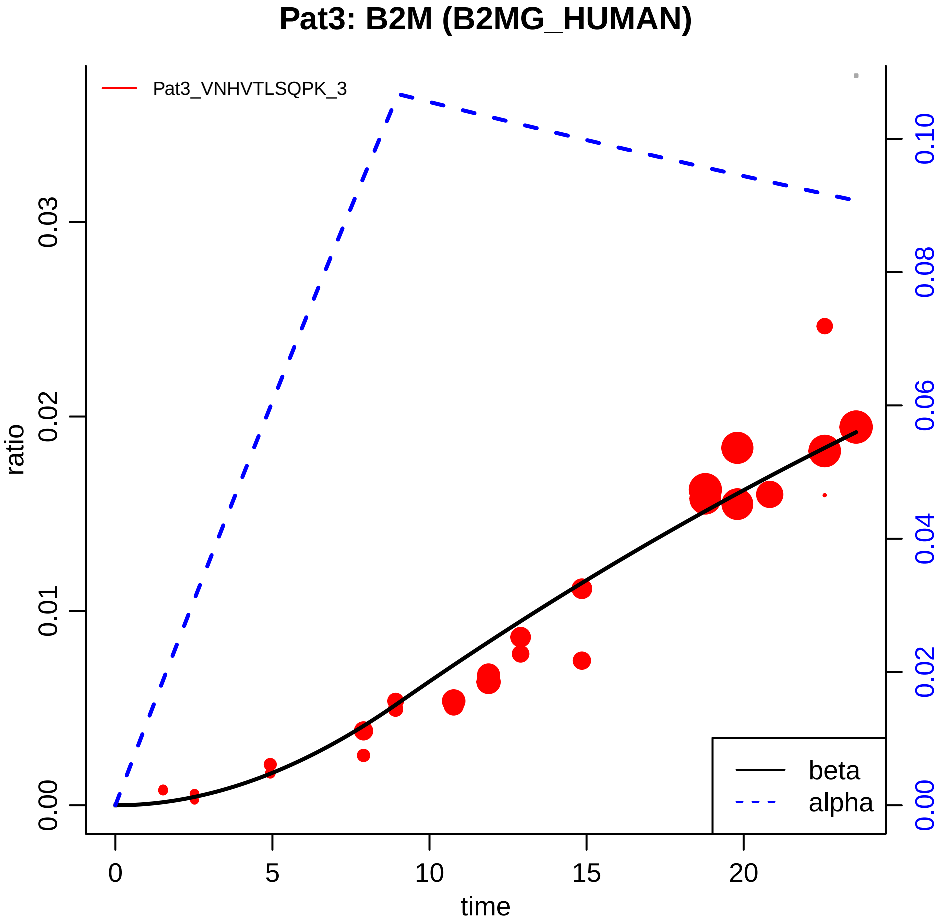

Pat3: B2M (B2MG_HUMAN)
Pat3_VNHVTLSQPK_3
ratio
0.00
0.01
0.02
0.03
time
0
5
10
15
20
0.00
0.02
0.04
0.06
0.08
0.10
beta
alpha

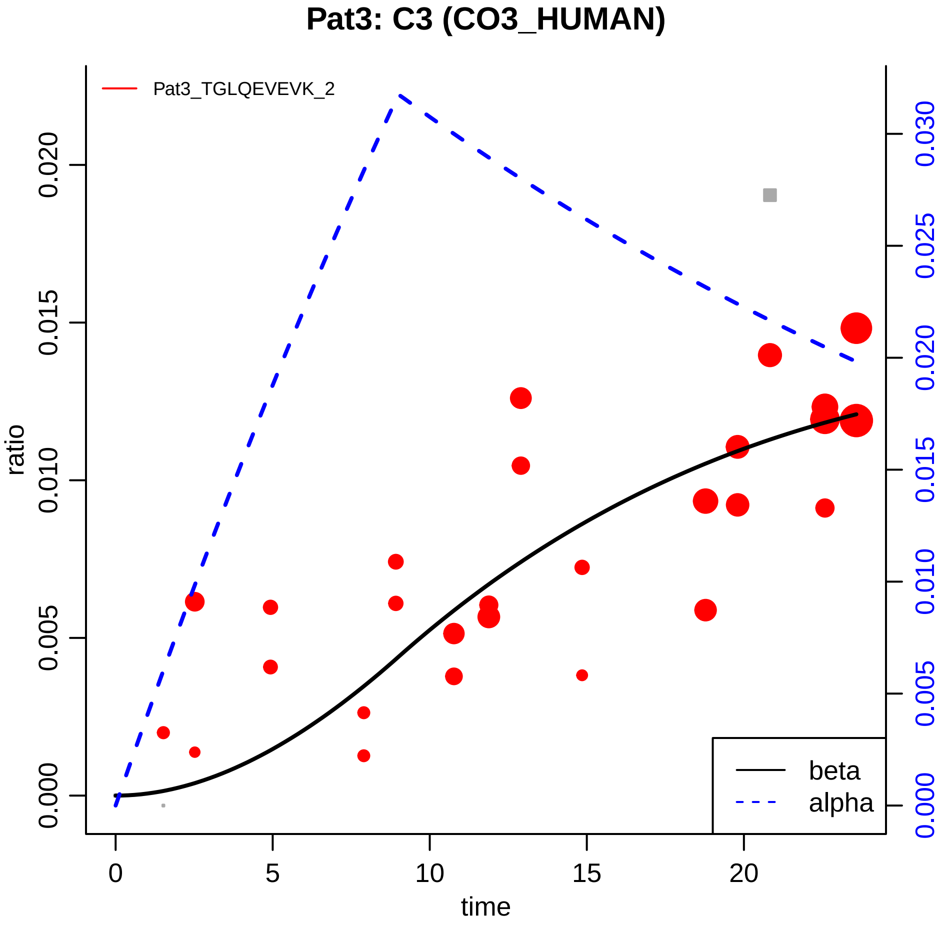

Pat3: C3 (CO3_HUMAN)
Pat3_TGLQEVEVK_2
ratio
time
0.000
0.005
0.010
0.015
0.020
0.025
0.030
0
5
10
15
20
beta
alpha

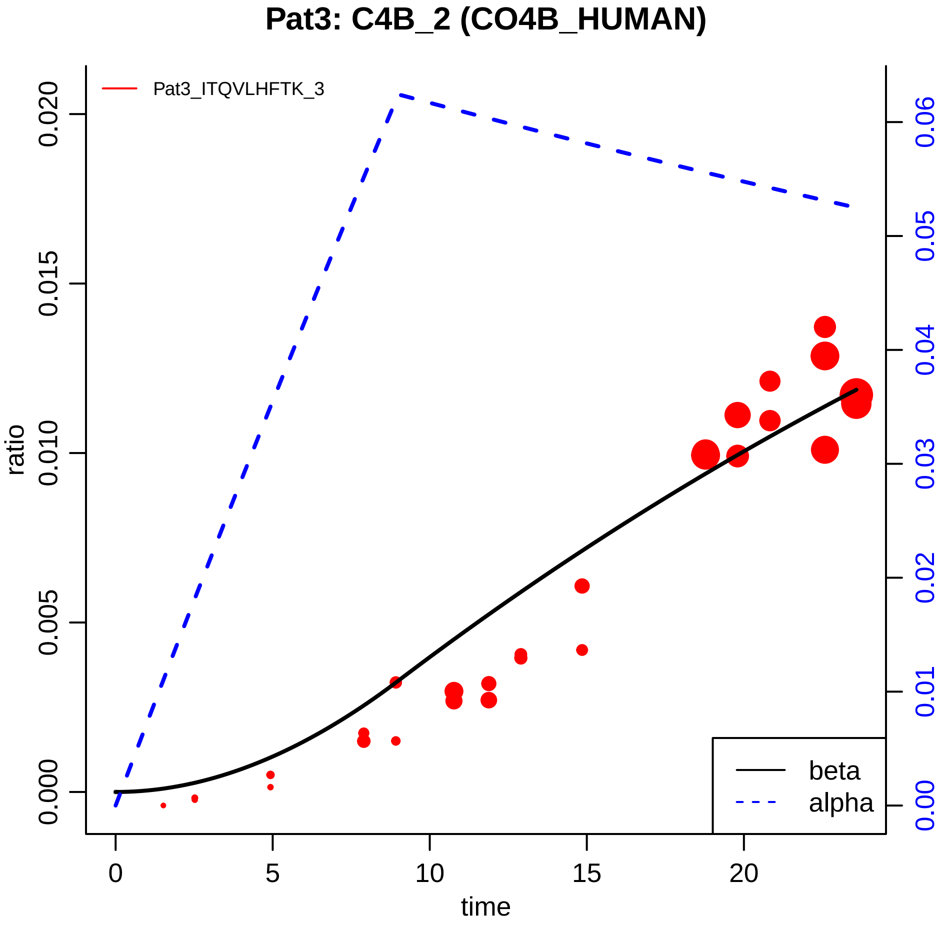

Pat3: C4B_2 (CO4B_HUMAN)
Pat3_ITQVLHFTK_3
ratio
0.000
0.005
0.010
0.015
0.020
0.00
0.01
0.02
0.03
0.04
0.05
0.06
0
5
10
15
20
time
beta
alpha

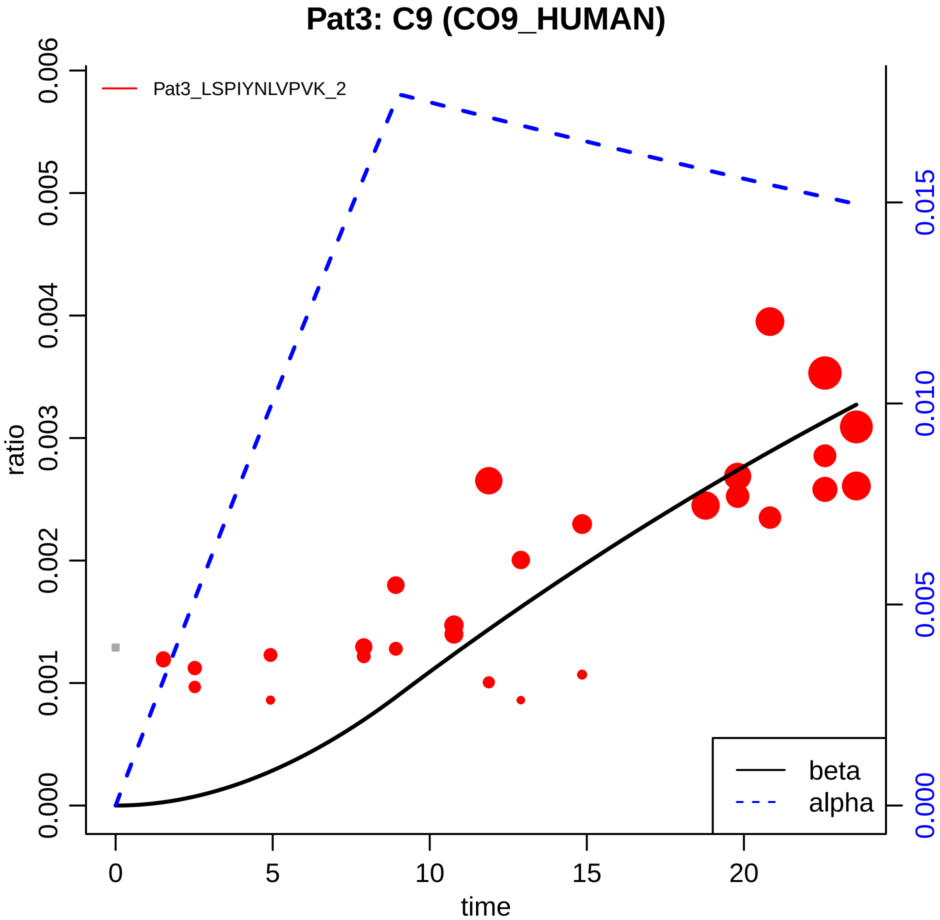

Pat3: C9 (CO9_HUMAN)
Pat3_LSPIYNLVPVK_2
ratio
0.000
0.001
0.002
0.003
0.004
0.005
0.006
0.000
0.005
0.010
0.015
0
5
10
15
20
time
beta
alpha

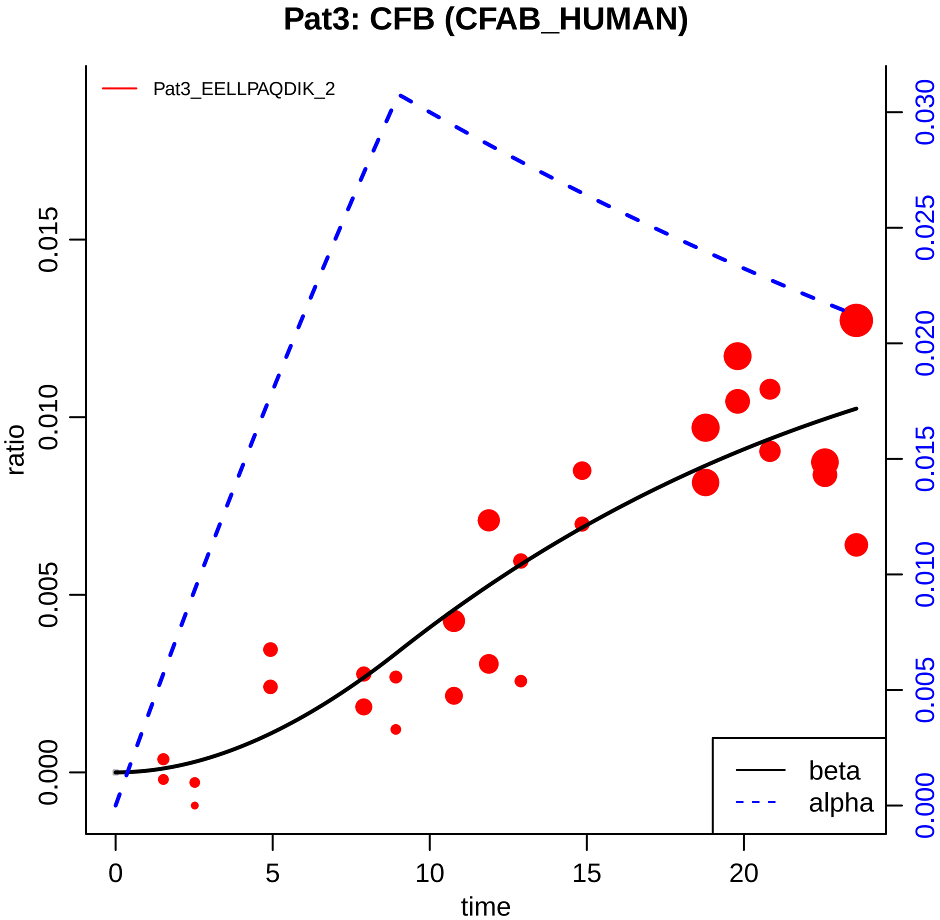

Pat3: CFB (CFAB_HUMAN)
Pat3_EELLPAQDIK_2
ratio
0.000
0.005
0.010
0.015
time
0
5
10
15
20
0.000
0.005
0.010
0.015
0.020
0.025
0.030
beta
alpha

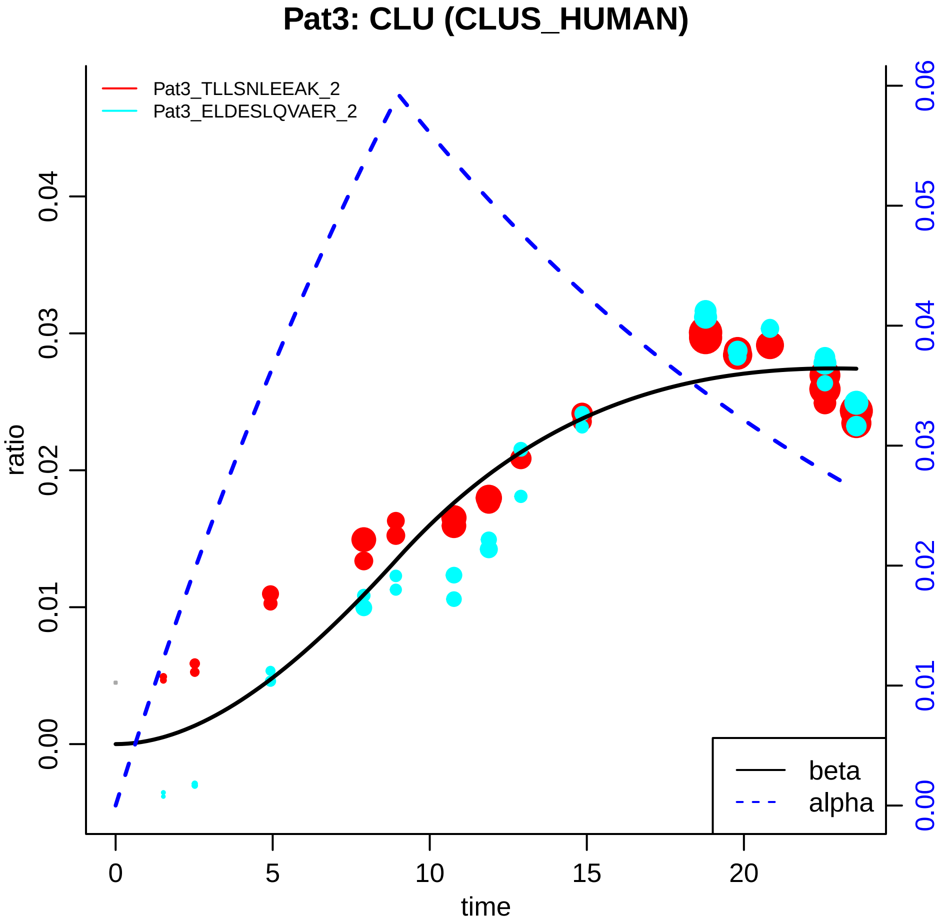
Pat3: CLU (CLUS_HUMAN)
Pat3_TLLSNLEEAK_2
Pat3_ELDESLQVAER_2
ratio
0.00
0.01
0.02
0.03
0.04
time
0
5
10
15
20
0.00
0.01
0.02
0.03
0.04
0.05
0.06
beta
alpha

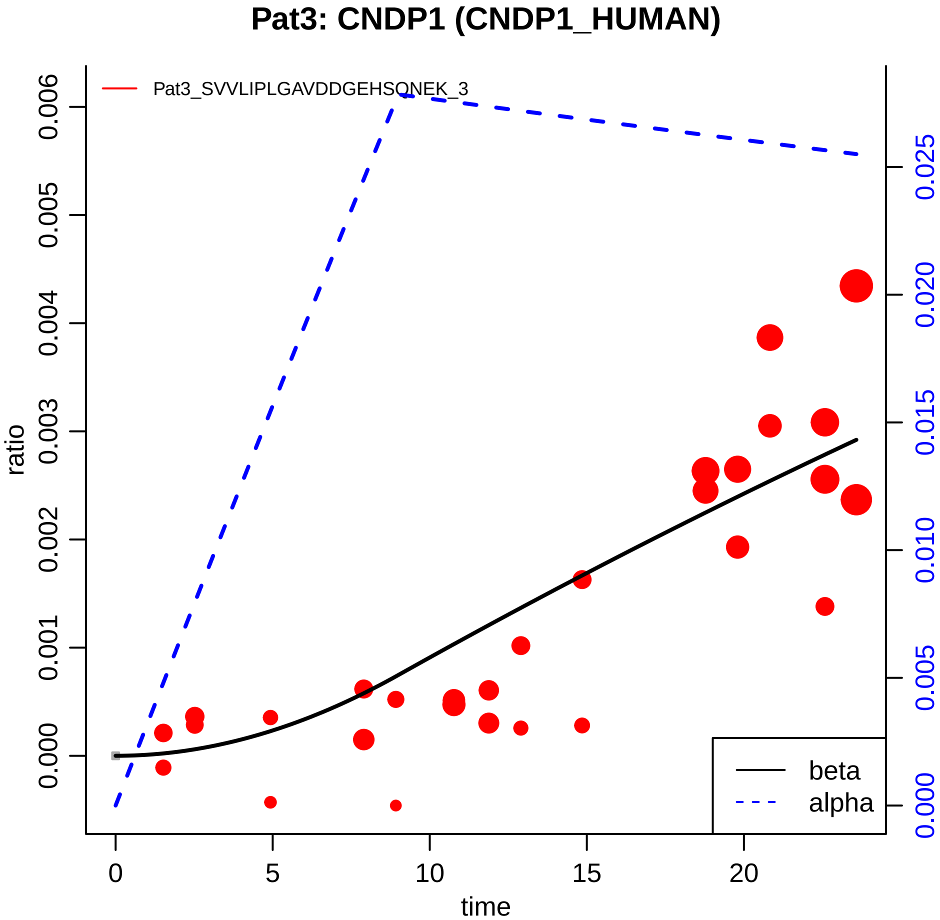

Pat3: CNDP1 (CNDP1_HUMAN)
Pat3_SVVLIPLGAVDDGEHSQNEK_3
ratio
0.000
0.001
0.002
0.003
0.004
0.005
0.006
0.000
0.005
0.010
0.015
0.020
0.025
0
5
10
15
20
time
beta
alpha

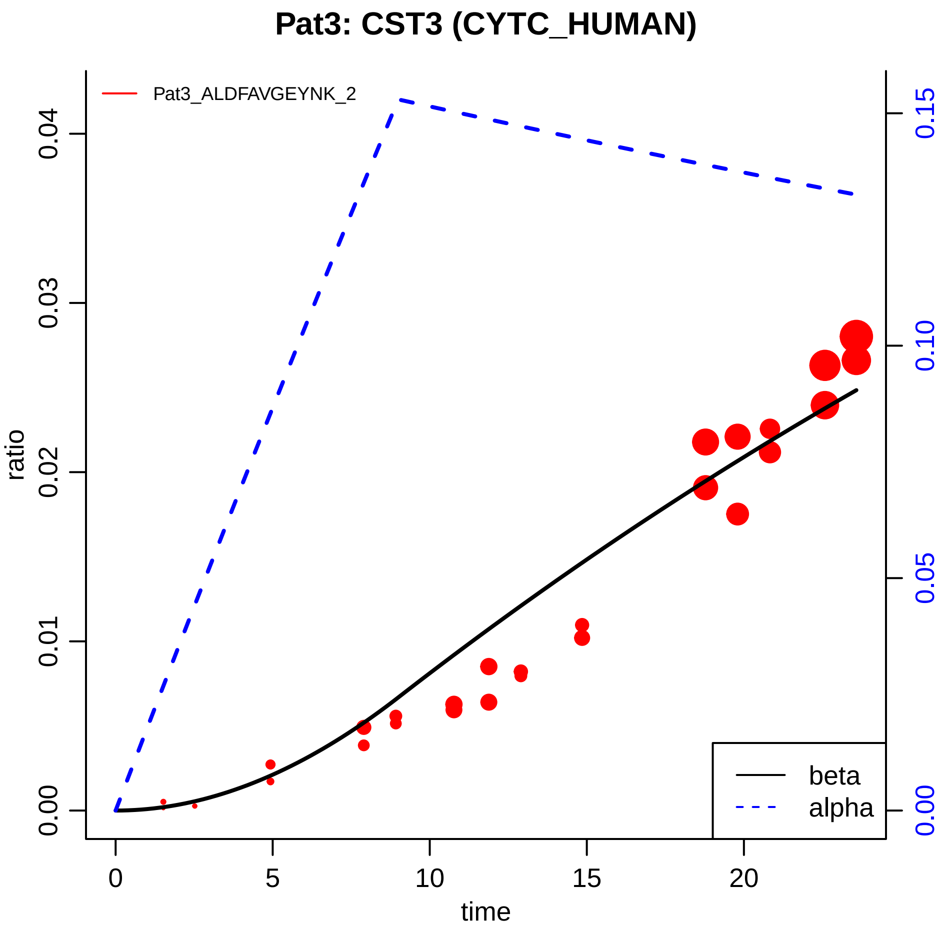

Pat3: CST3 (CYTC_HUMAN)
Pat3_ALDFAVGEYNK_2
ratio
time
0.00
0.01
0.02
0.03
0.04
0
5
10
15
20
0.00
0.05
0.10
0.15
beta
alpha

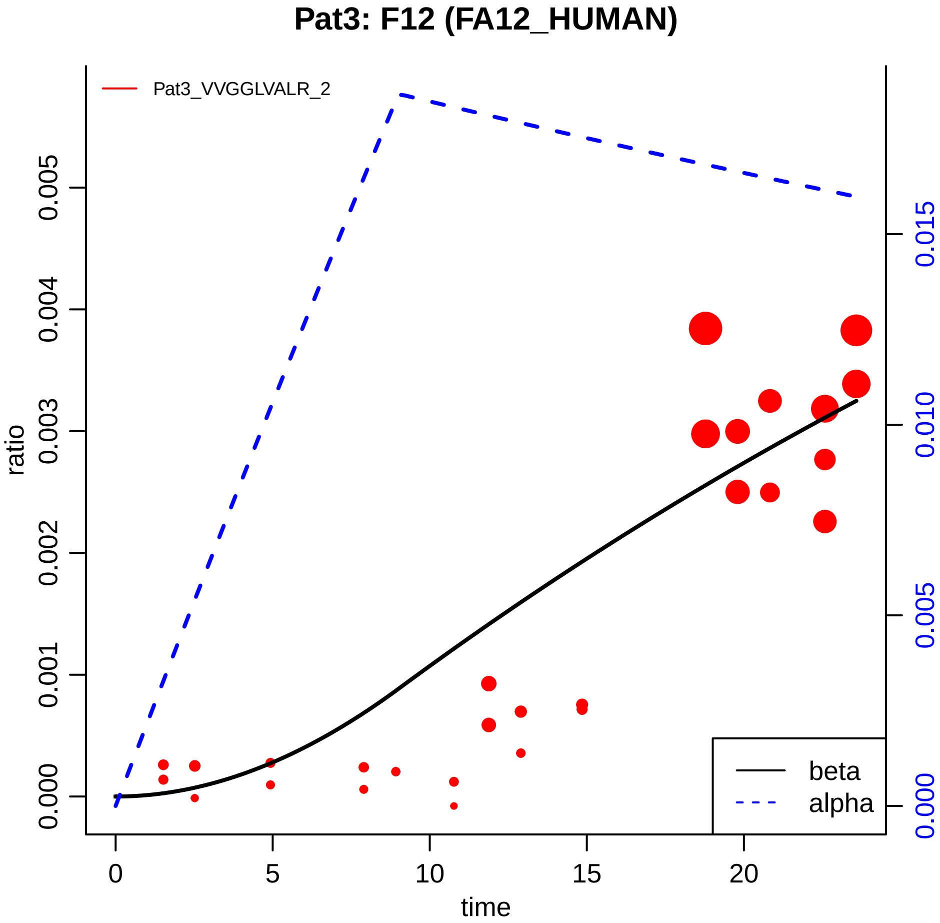
Pat3: F12 (FA12_HUMAN)
Pat3_VVGGLVALR_2
ratio
0.000
0.001
0.002
0.003
0.004
0.005
0.000
0.005
0.010
0.015
0
5
10
15
20
time
beta
alpha

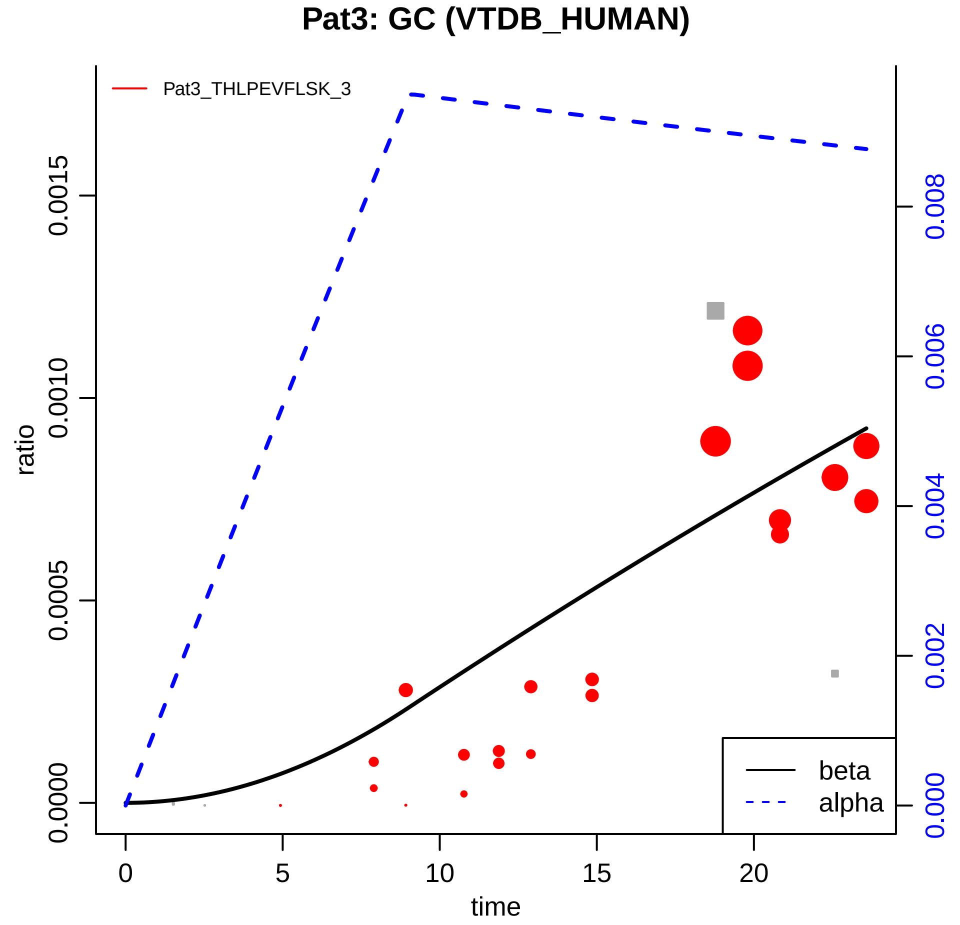

Pat3: GC (VTDB_HUMAN)
Pat3_THLPEVFLSK_3
ratio
0.0000
0.0005
0.0010
0.0015
0.000
0.002
0.004
0.006
0.008
0
5
10
15
20
time
beta
alpha

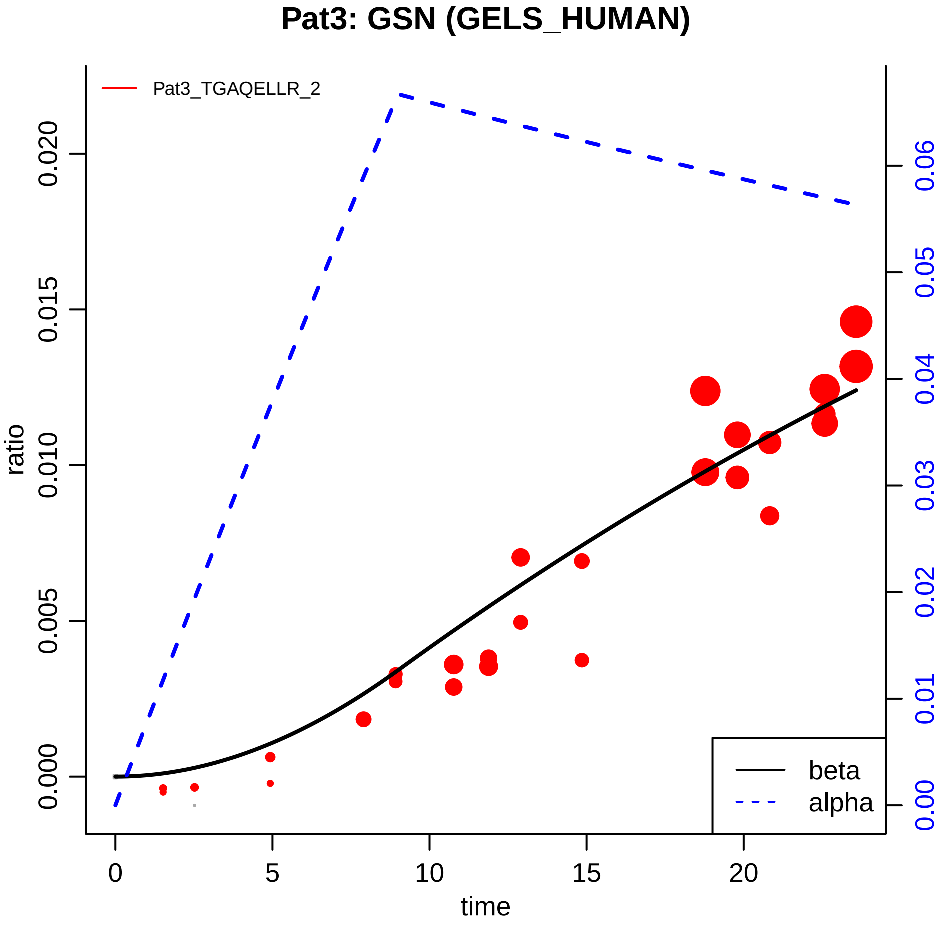

Pat3: GSN (GELS_HUMAN)
Pat3_TGAQELLR_2
ratio
0.000
0.005
0.010
0.015
0.020
time
0
5
10
15
20
0.00
0.01
0.02
0.03
0.04
0.05
0.06
beta
alpha

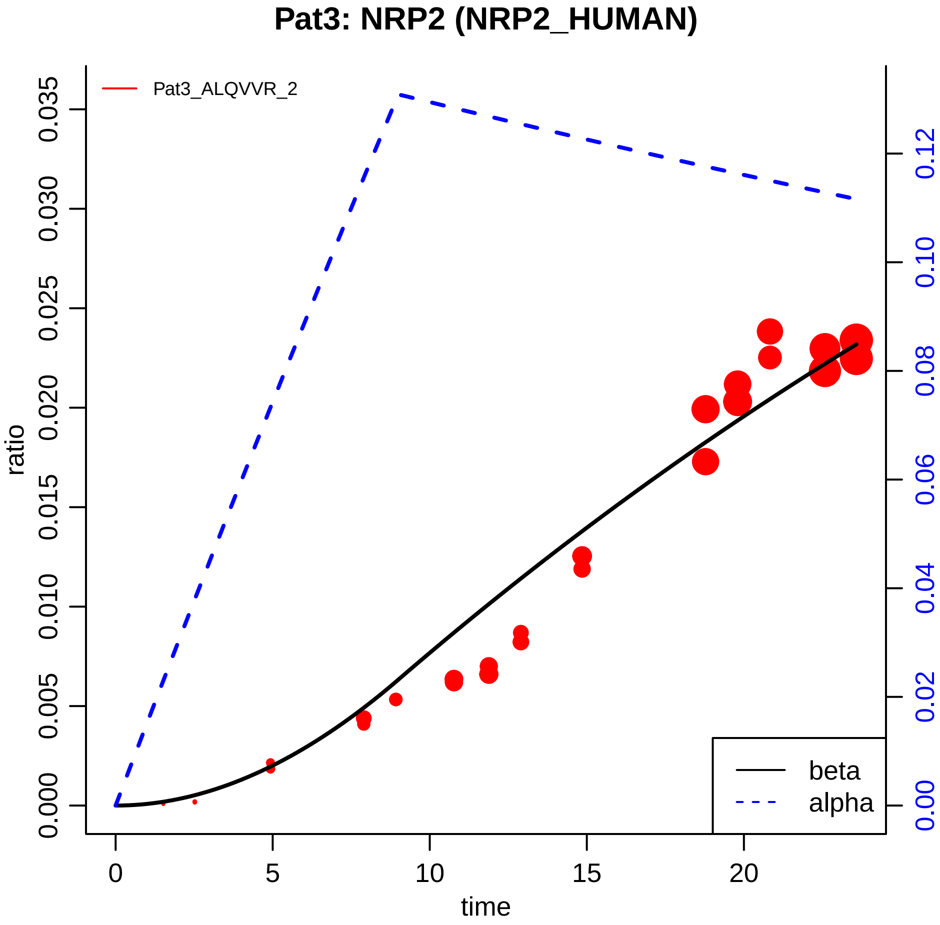

Pat3: NRP2 (NRP2_HUMAN)
Pat3_ALQVVR_2
ratio
time
0.000
0.005
0.010
0.015
0.020
0.025
0.030
0.035
0
5
10
15
20
0.00
0.02
0.04
0.06
0.08
0.10
0.12
beta
alpha

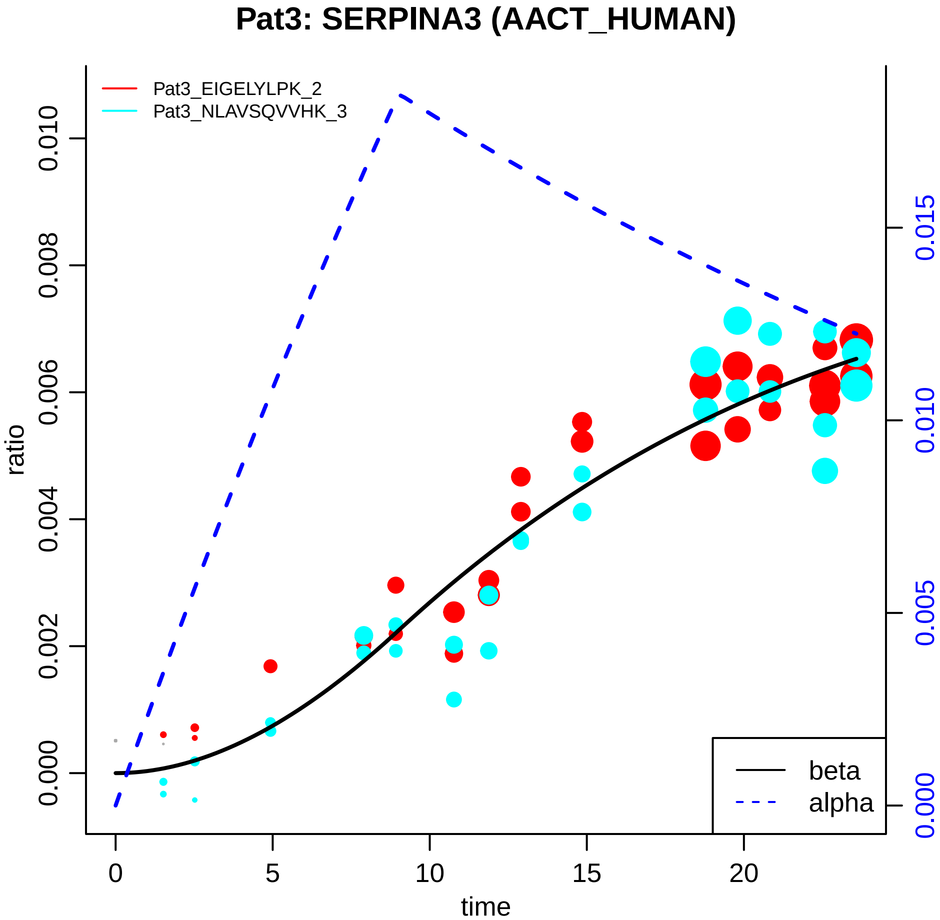

Pat3: SERPINA3 (AACT_HUMAN)
Pat3_EIGELYLPK_2
Pat3_NLAVSQVVHK_3
ratio
0.000
0.002
0.004
0.006
0.008
0.010
time
0
5
10
15
20
0.000
0.005
0.010
0.015
beta
alpha

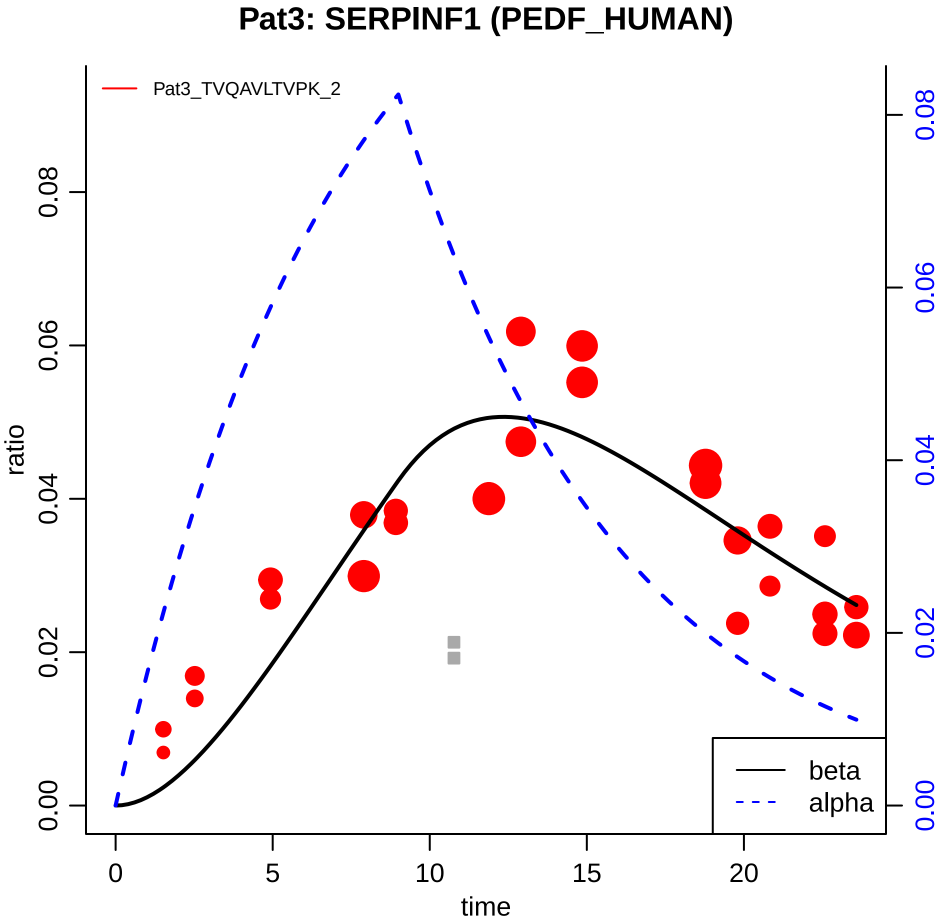

Pat3: SERPINF1 (PEDF_HUMAN)
Pat3_TVQAVLTVPK_2
ratio
0.00
0.02
0.04
0.06
0.08
time
0
5
10
15
20
beta
alpha

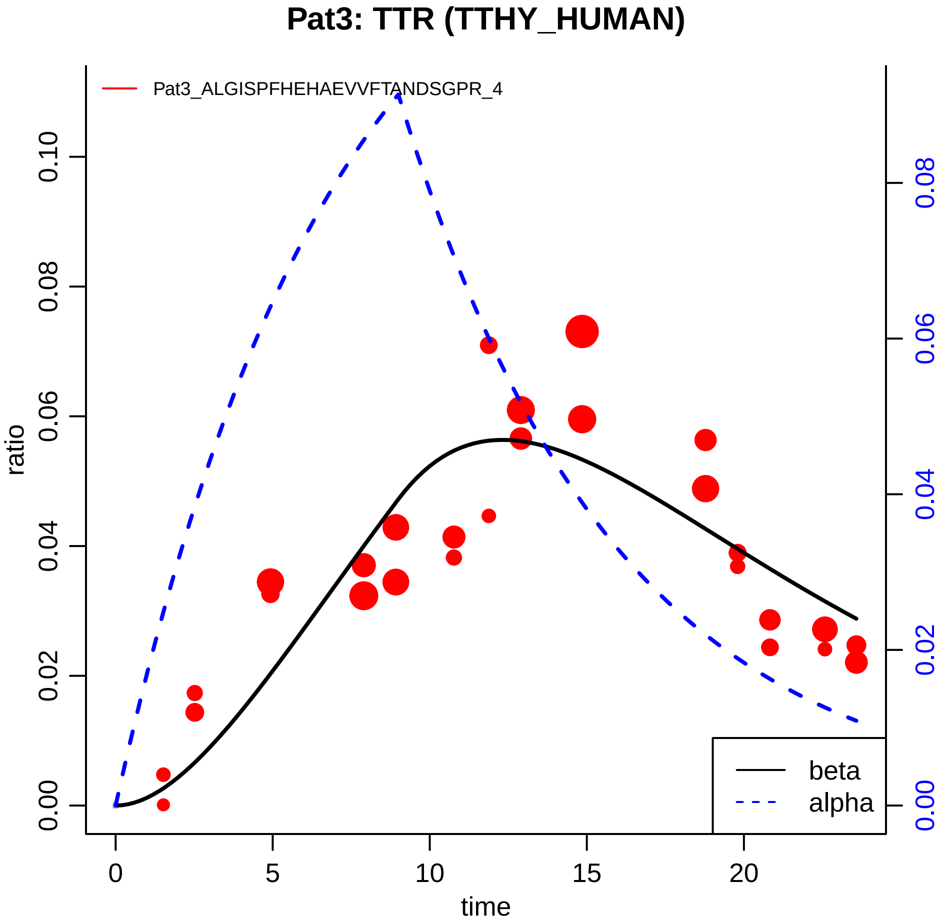

Pat3: TTR (TTHY_HUMAN)
Pat3_ALGISPFHEHAEVVFTANDSGPR_4
ratio
0.00
0.02
0.04
0.06
0.08
0.10
0.00
0.02
0.04
0.06
0.08
0
5
10
15
20
time
beta
alpha

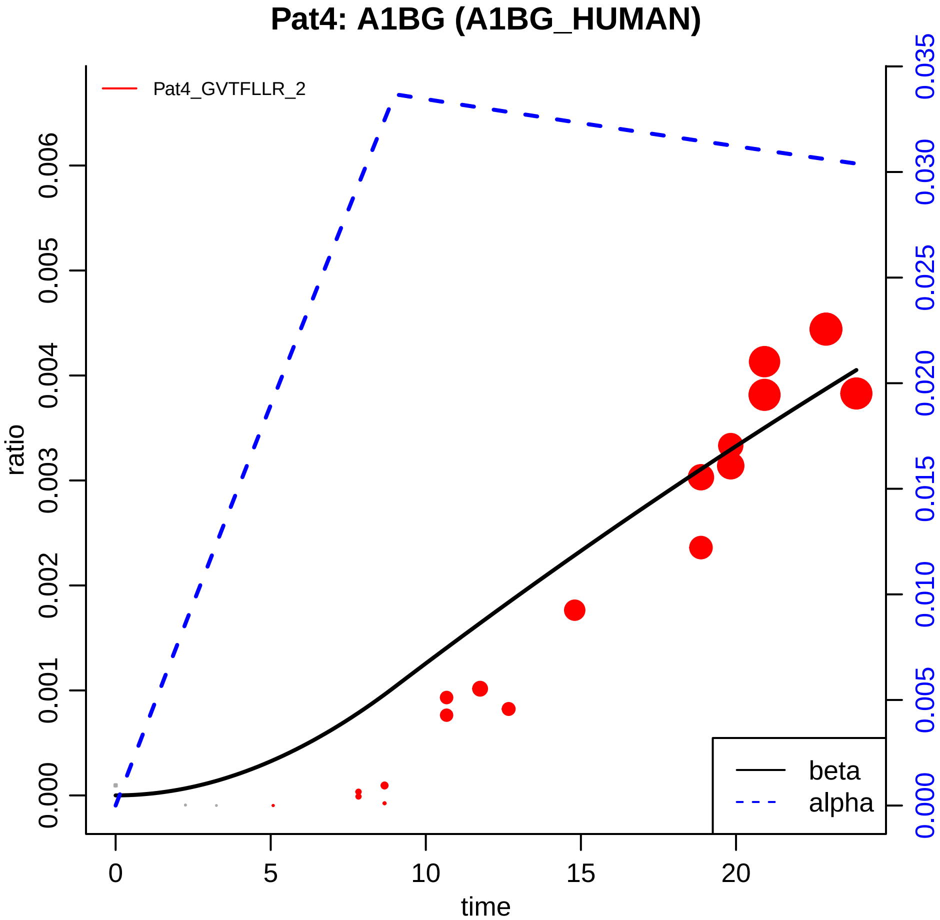

Pat4: A1BG (A1BG_HUMAN)
Pat4_GVTFLLR_2
ratio
0.000
0.001
0.002
0.003
0.004
0.005
0.006
time
0
5
10
15
20
0.000
0.005
0.010
0.015
0.020
0.025
0.030
0.035
beta
alpha

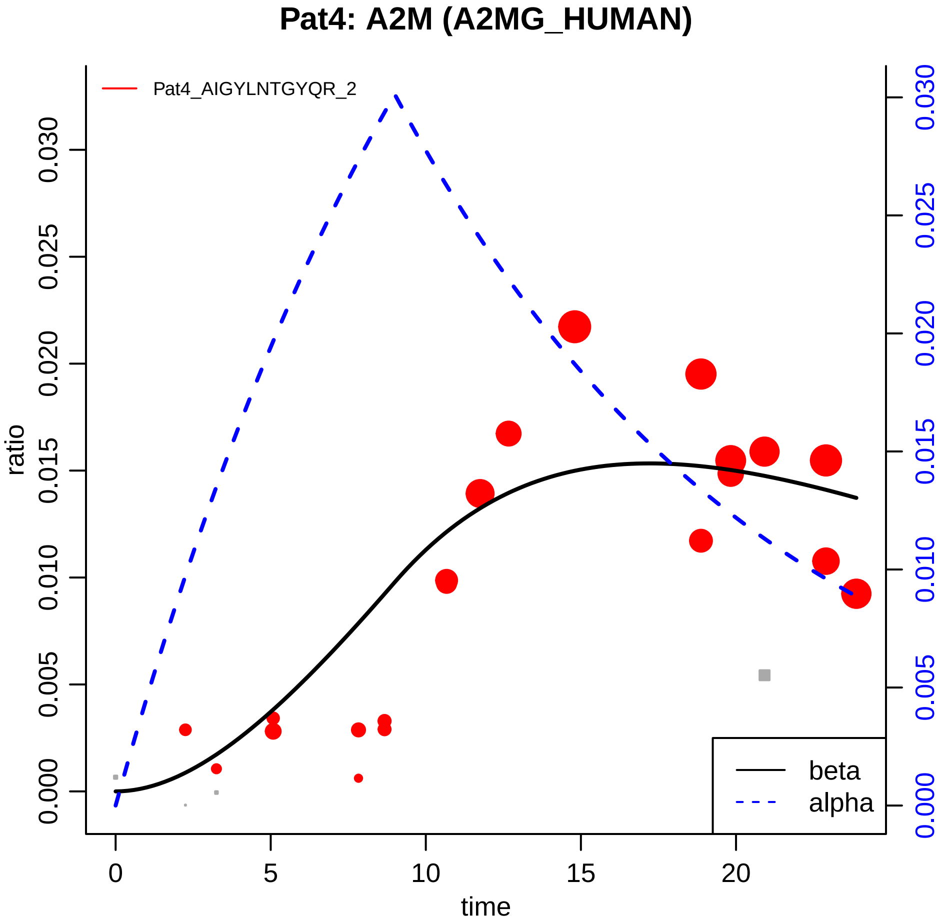

Pat4: A2M (A2MG_HUMAN)
Pat4_AIGYLNTGYQR_2
ratio
0.000
0.005
0.010
0.015
0.020
0.025
0.030
time
0
5
10
15
20
beta
alpha

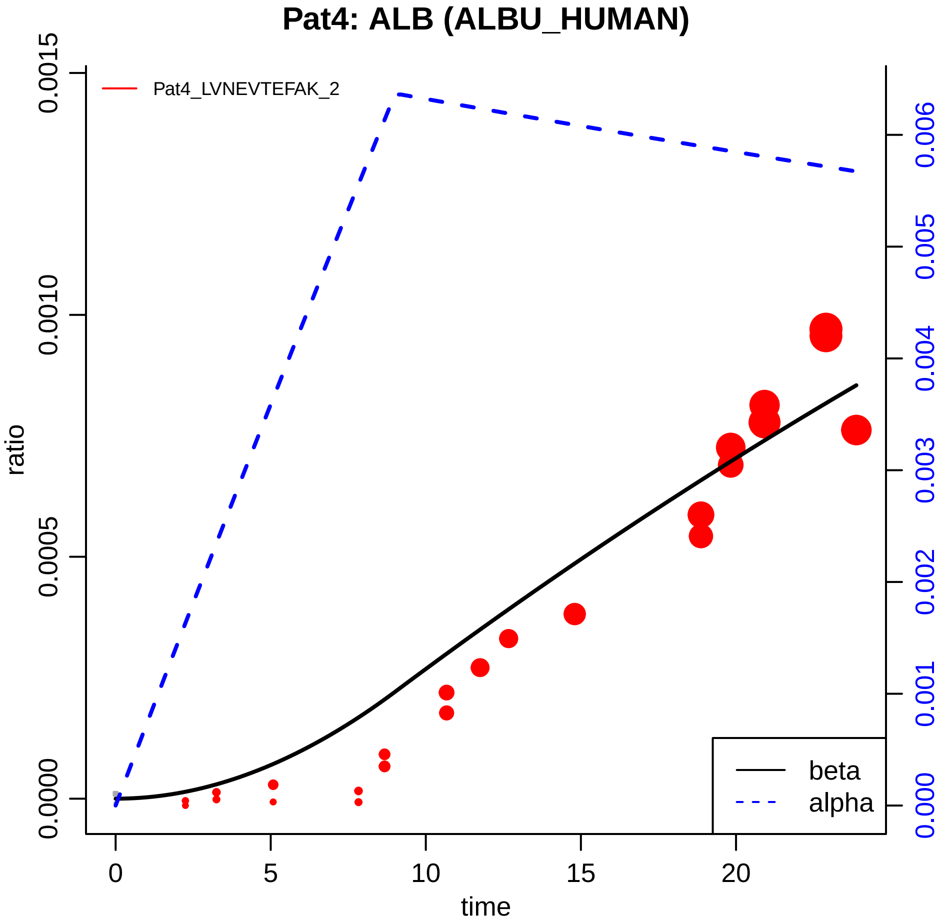

Pat4: ALB (ALBU_HUMAN)
Pat4_LVNEVTEFAK_2
ratio
0.0000
0.0005
0.0010
0.0015
0.000
0.001
0.002
0.003
0.004
0.005
0.006
0
5
10
15
20
time
beta
alpha

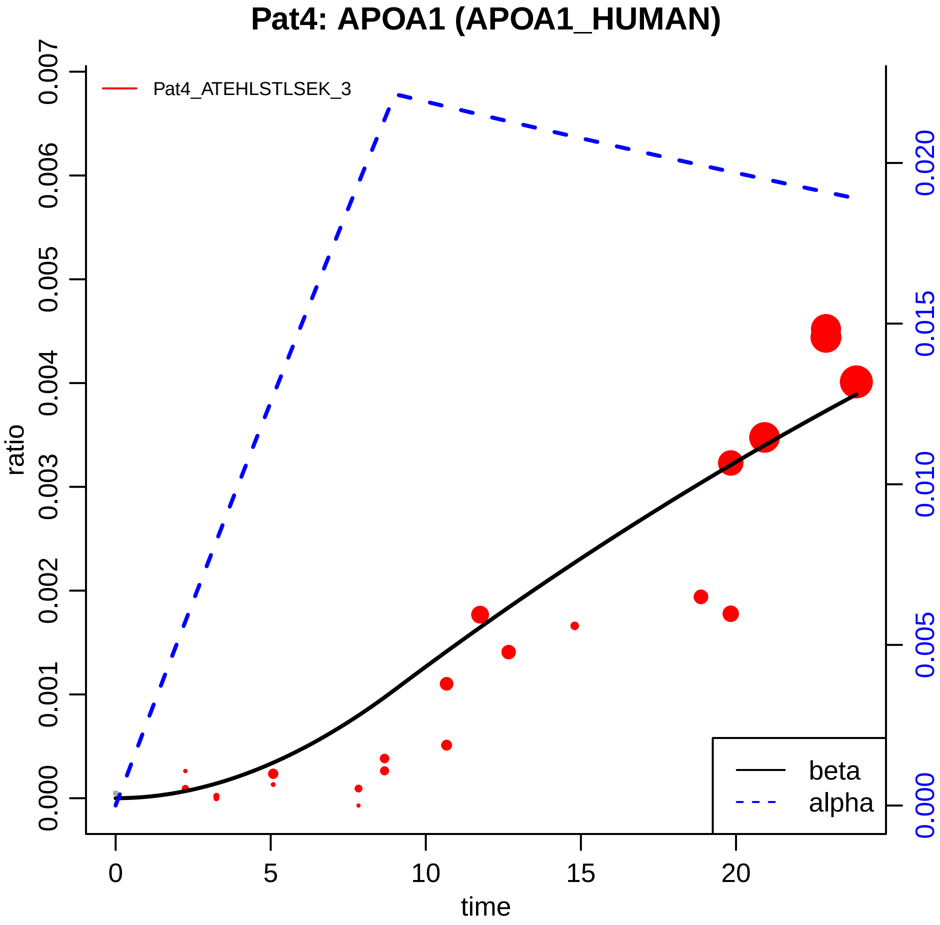

Pat4: APOA1 (APOA1_HUMAN)
Pat4_ATEHLSTLSEK_3
ratio
0.000
0.001
0.002
0.003
0.004
0.005
0.006
0.007
0.000
0.005
0.010
0.015
0.020
0
5
10
15
20
time
beta
alpha

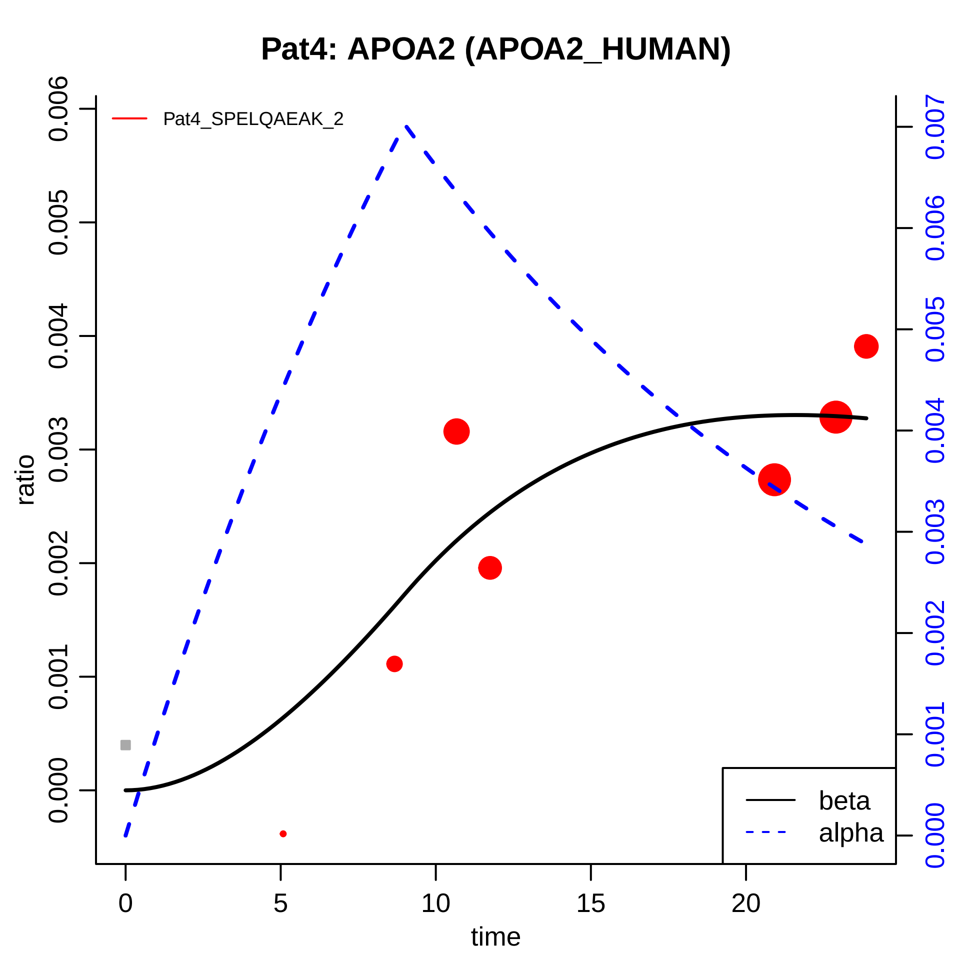

Pat4: APOA2 (APOA2_HUMAN)
Pat4_SPELQAEAK_2
ratio
0.000
0.001
0.002
0.003
0.004
0.005
0.006
0.000
0.001
0.002
0.003
0.004
0.005
0.006
0.007
time
0
5
10
15
20
beta
alpha

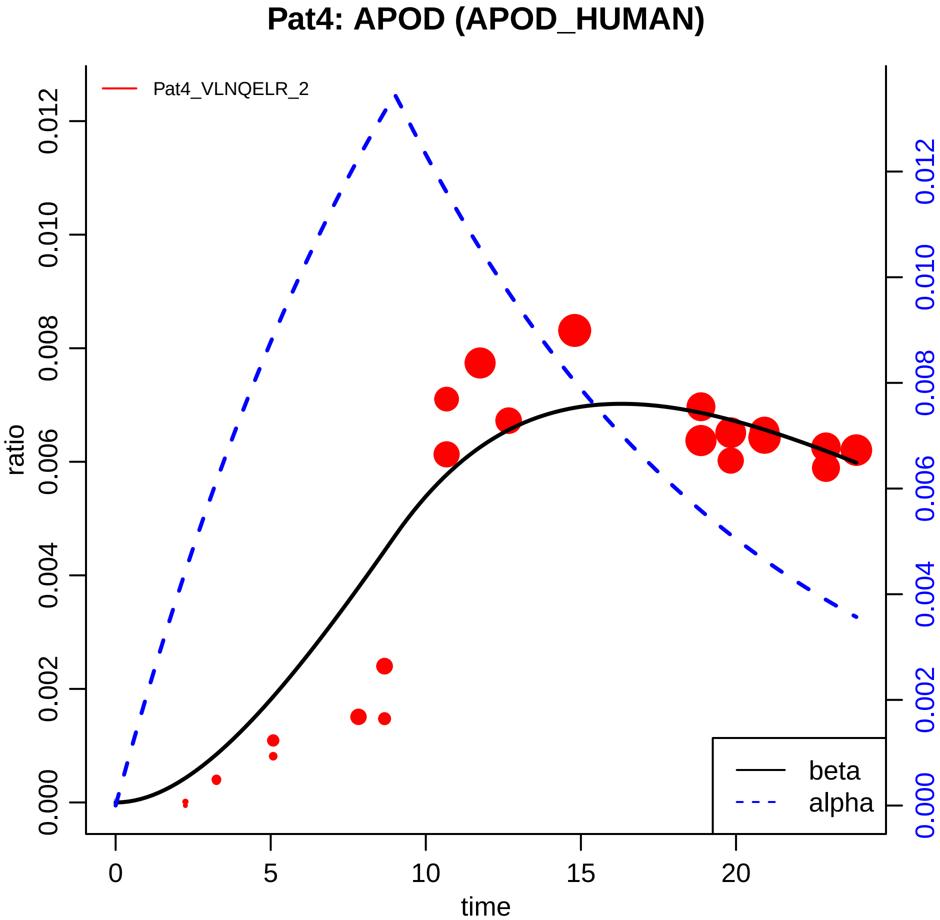

Pat4: APOD (APOD_HUMAN)
Pat4_VLNQELR_2
ratio
time
beta
alpha
0
5
10
15
20
0.000
0.002
0.004
0.006
0.008
0.010
0.012

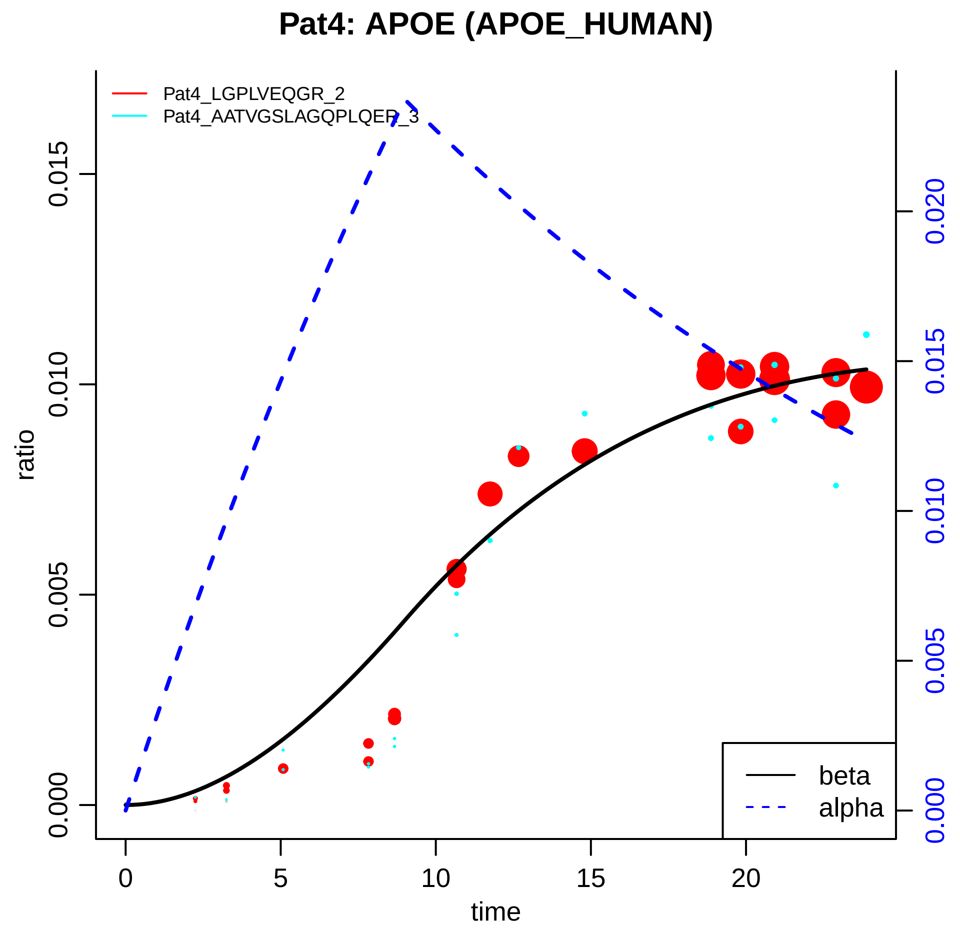

Pat4: APOE (APOE_HUMAN)
Pat4_LGPLVEQGR_2
Pat4_AATVGSLAGQPLQER_3
ratio
time
beta
alpha
0.000
0.005
0.010
0.015
0.020
0
5
10
15
20

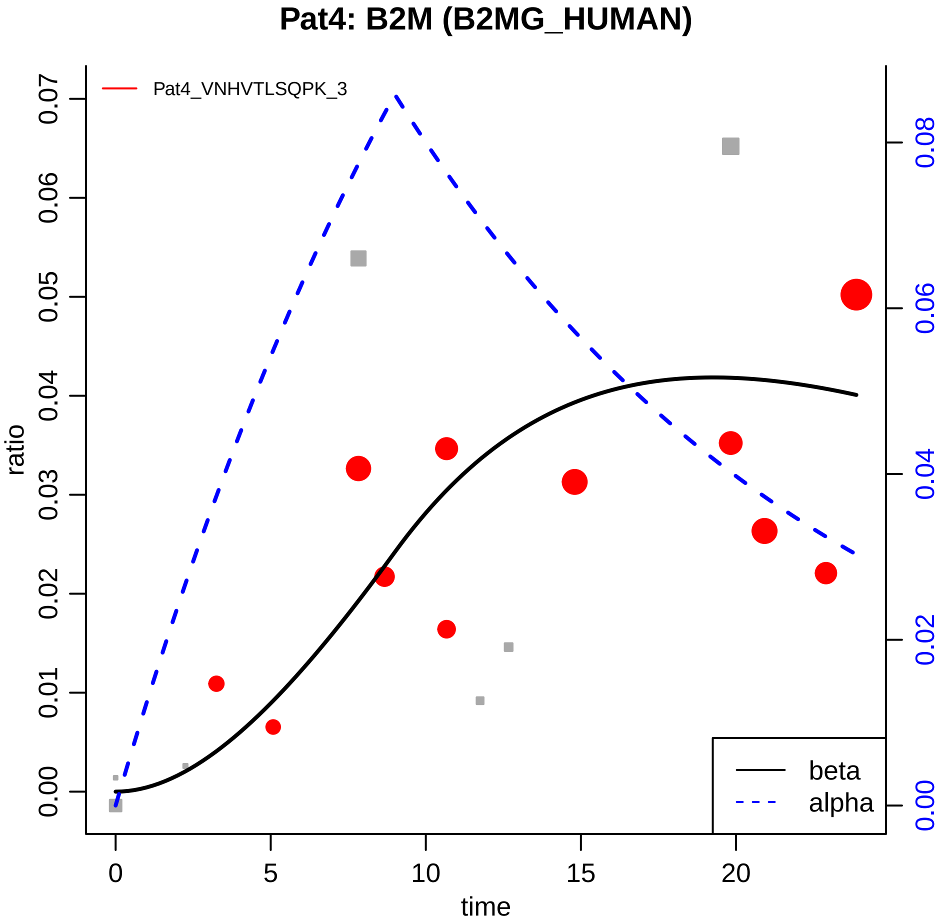

Pat4: B2M (B2MG_HUMAN)
Pat4_VNHVTLSQPK_3
ratio
time
beta
alpha
0.00
0.01
0.02
0.03
0.04
0.05
0.06
0.07
0.00
0.02
0.04
0.06
0.08
0
5
10
15
20

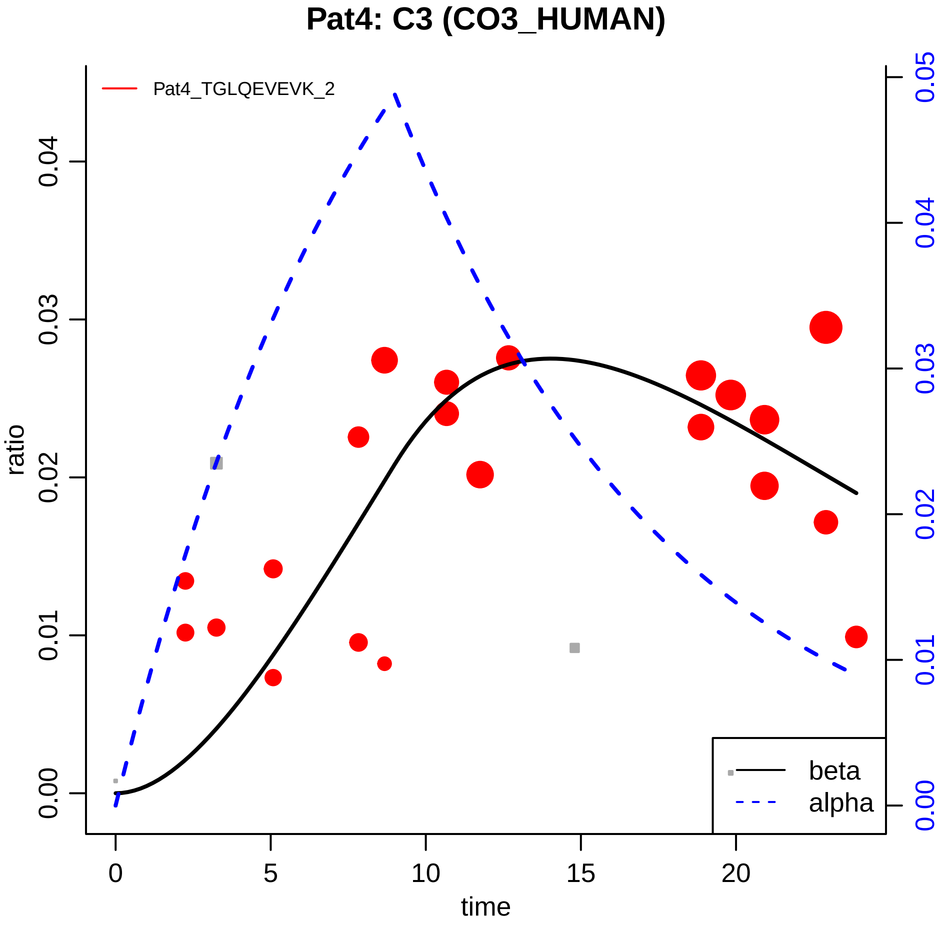

Pat4: C3 (CO3_HUMAN)
Pat4_TGLQEVEVK_2
ratio
time
0.00
0.01
0.02
0.03
0.04
0.05
0
5
10
15
20
beta
alpha

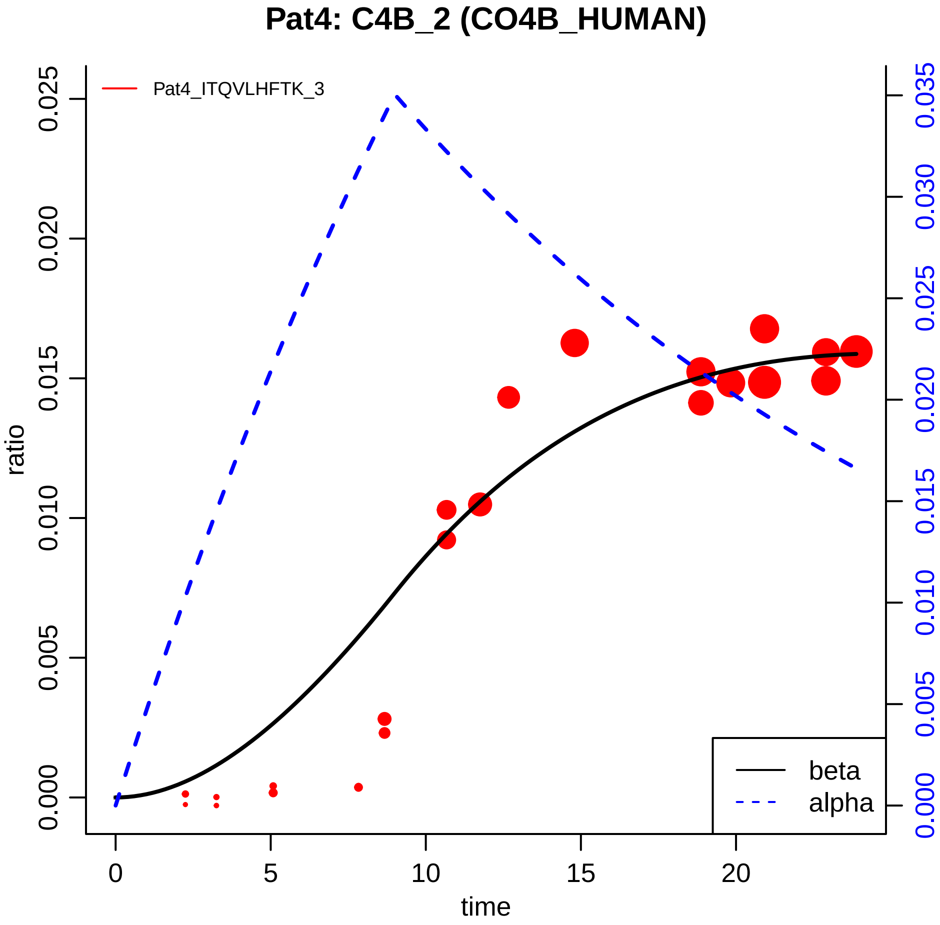

Pat4: C4B_2 (CO4B_HUMAN)
Pat4_ITQVLHFTK_3
ratio
time
0.000
0.005
0.010
0.015
0.020
0.025
0.030
0.035
0
5
10
15
20
beta
alpha

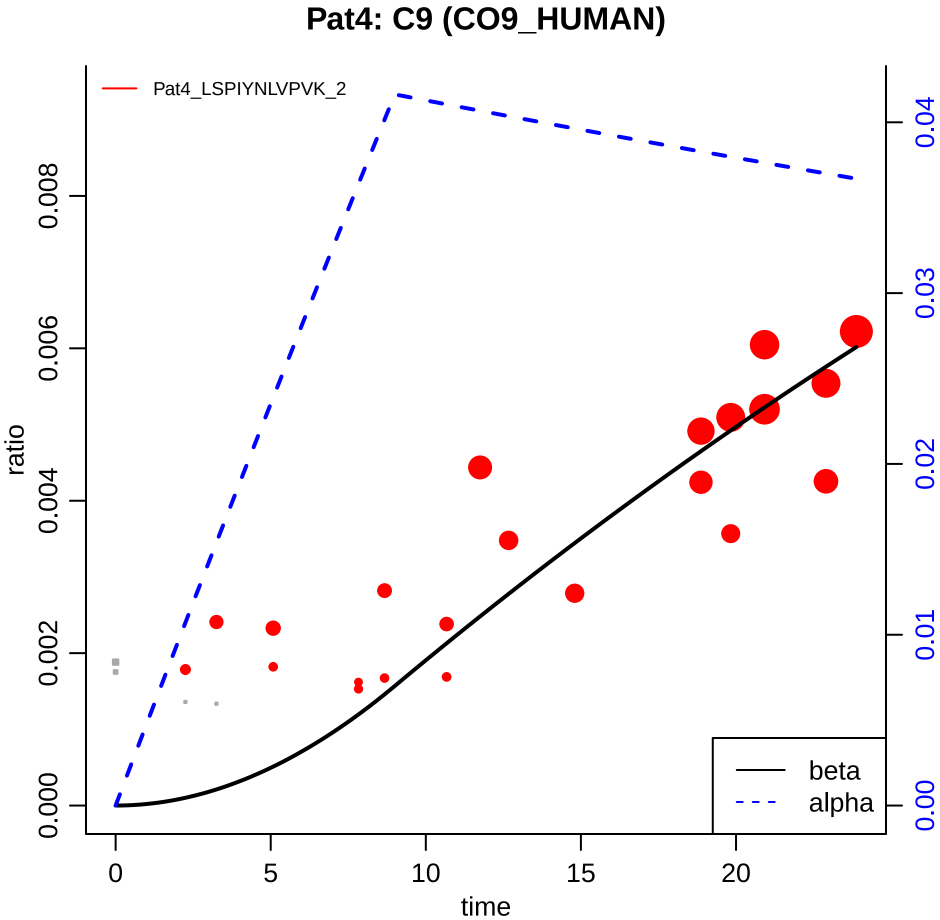

Pat4: C9 (CO9_HUMAN)
Pat4_LSPIYNLVPVK_2
ratio
0.000
0.002
0.004
0.006
0.008
0.00
0.01
0.02
0.03
0.04
0
5
10
15
20
time
beta
alpha

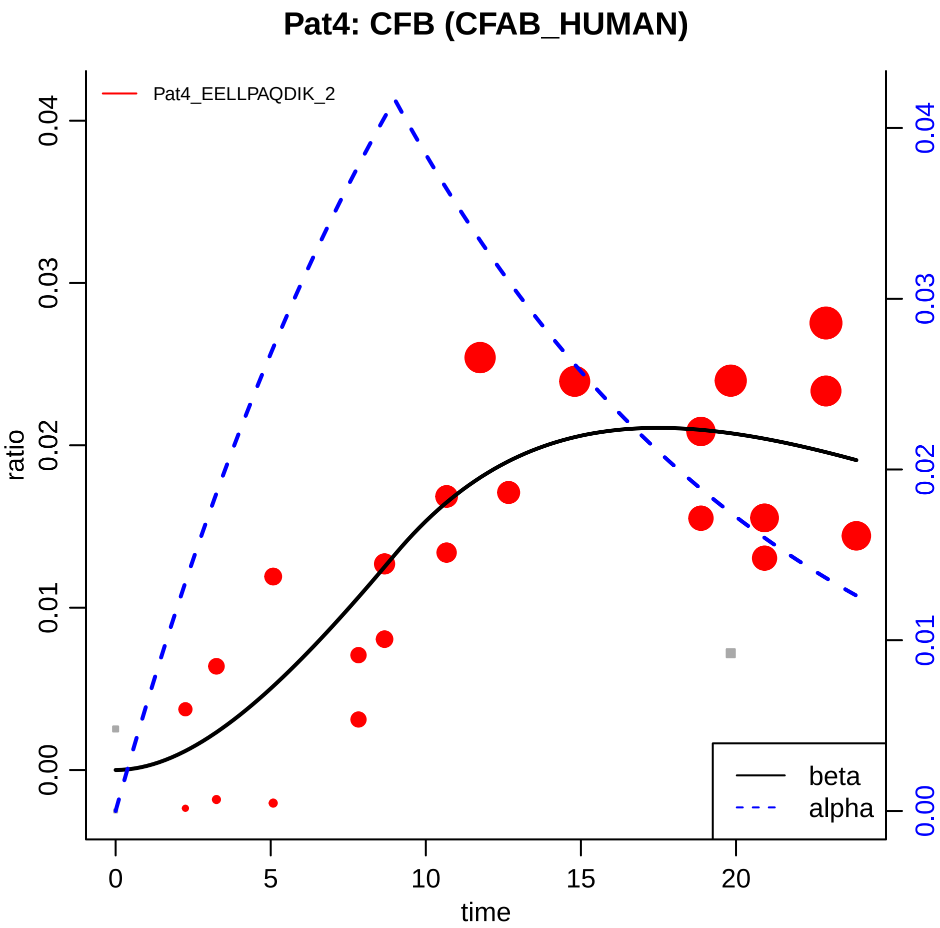

Pat4: CFB (CFAB_HUMAN)
Pat4_EELLPAQDIK_2
ratio
0.00
0.01
0.02
0.03
0.04
time
0
5
10
15
20
beta
alpha

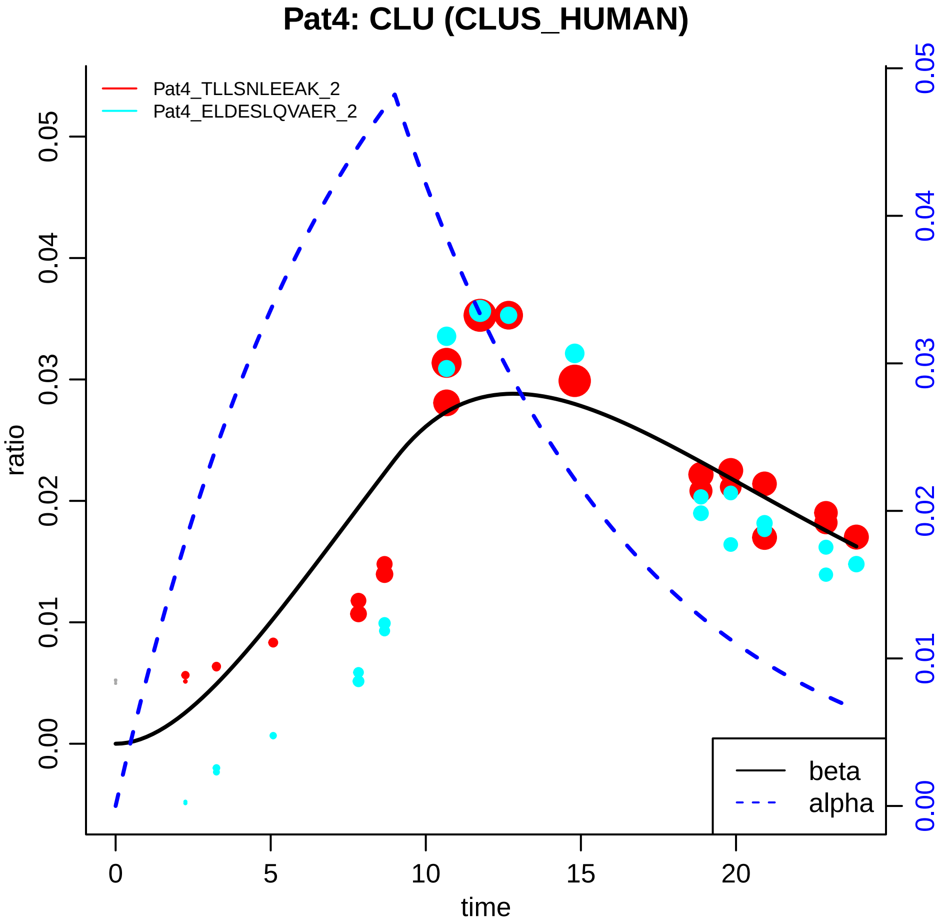
Pat4: CLU (CLUS_HUMAN)
Pat4_TLLSNLEEAK_2
Pat4_ELDESLQVAER_2
ratio
time
beta
alpha
0.00
0.01
0.02
0.03
0.04
0.05
0
5
10
15
20

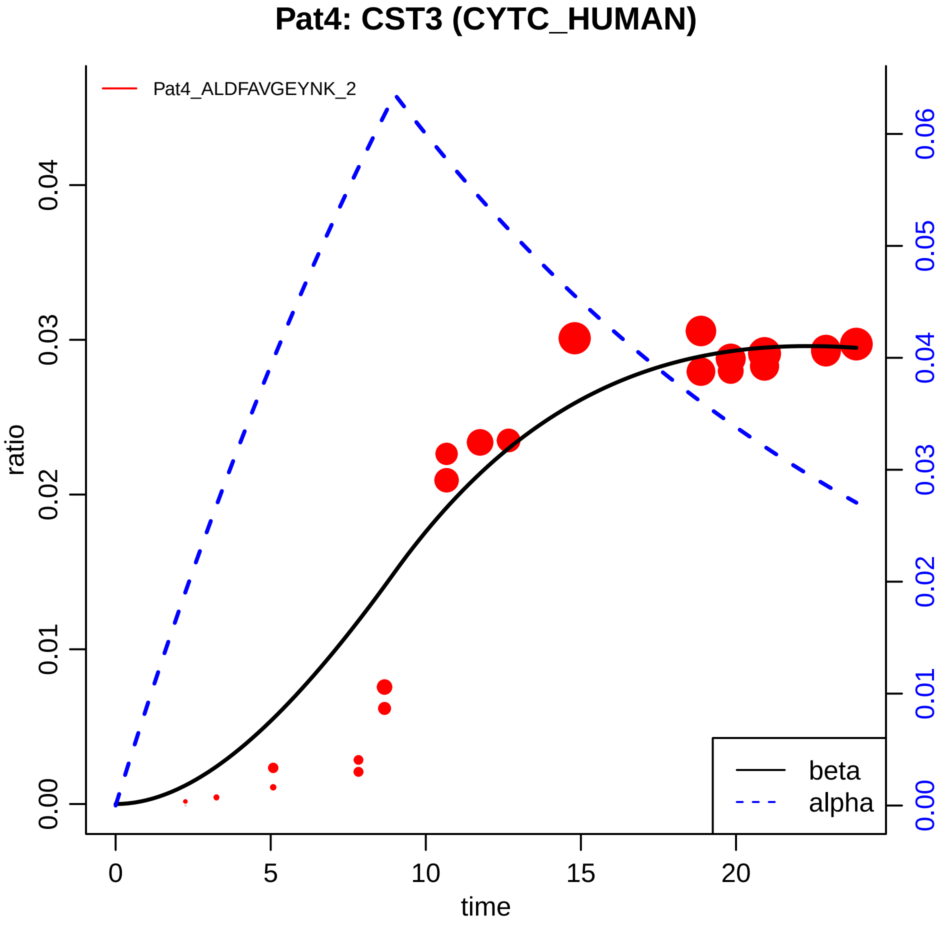

Pat4: CST3 (CYTC_HUMAN)
Pat4_ALDFAVGEYNK_2
ratio
time
beta
alpha
0.00
0.01
0.02
0.03
0.04
0.05
0.06
0
5
10
15
20

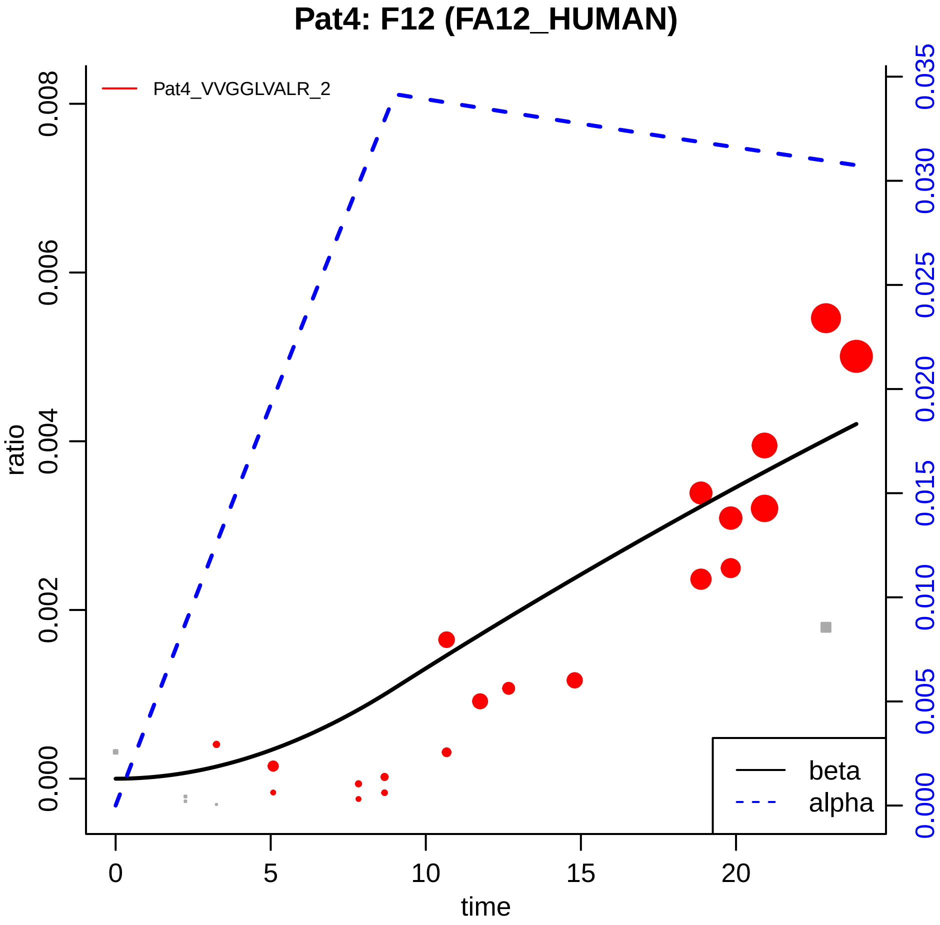

Pat4: F12 (FA12_HUMAN)
Pat4_VVGGLVALR_2
ratio
0.000
0.002
0.004
0.006
0.008
time
0
5
10
15
20
0.000
0.005
0.010
0.015
0.020
0.025
0.030
0.035
beta
alpha

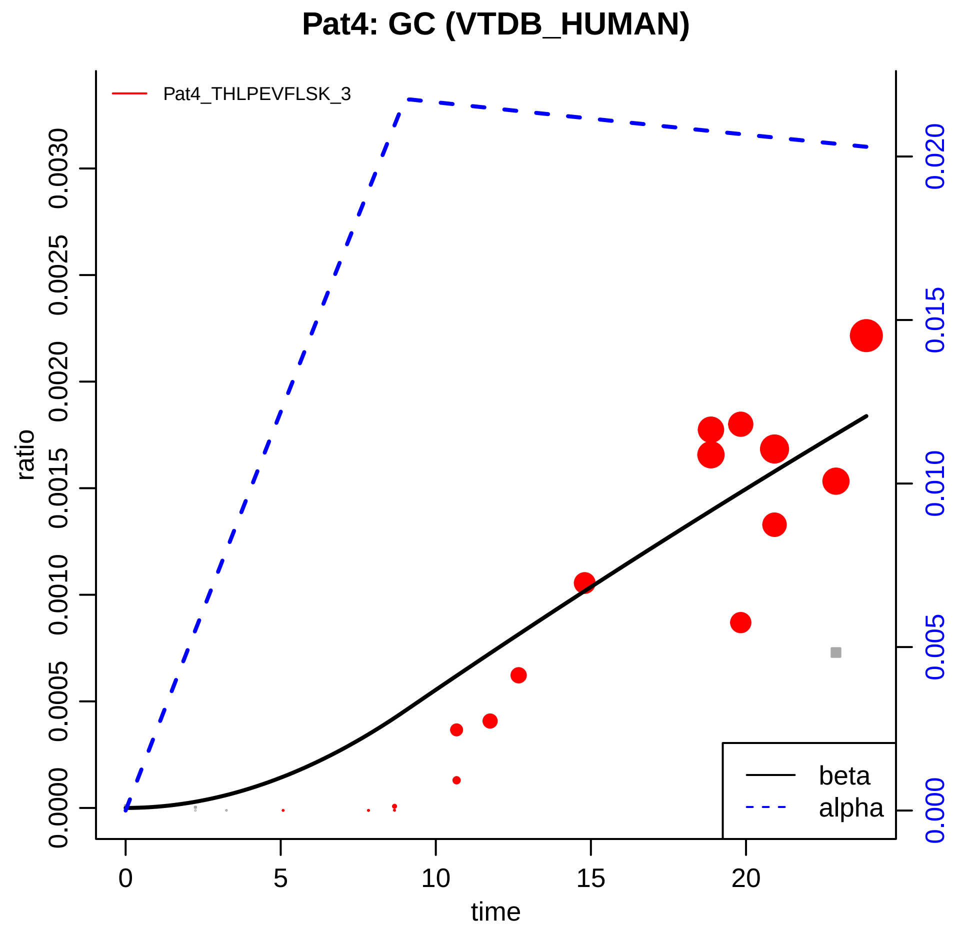
Pat4: GC (VTDB_HUMAN)
Pat4_THLPEVFLSK_3
ratio
0.0000
0.0005
0.0010
0.0015
0.0020
0.0025
0.0030
0.000
0.005
0.010
0.015
0.020
0
5
10
15
20
time
beta
alpha

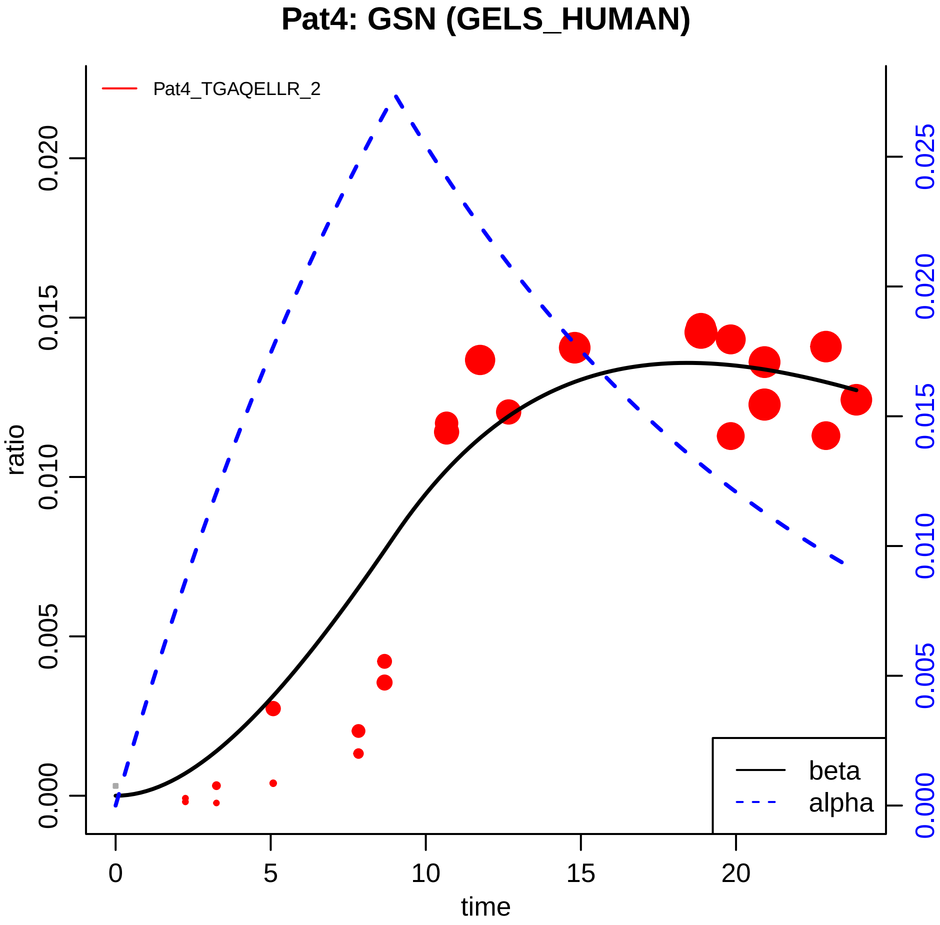
Pat4: GSN (GELS_HUMAN)
Pat4_TGAQELLR_2
ratio
0.000
0.005
0.010
0.015
0.020
0.000
0.005
0.010
0.015
0.020
0.025
0
5
10
15
20
time
beta
alpha

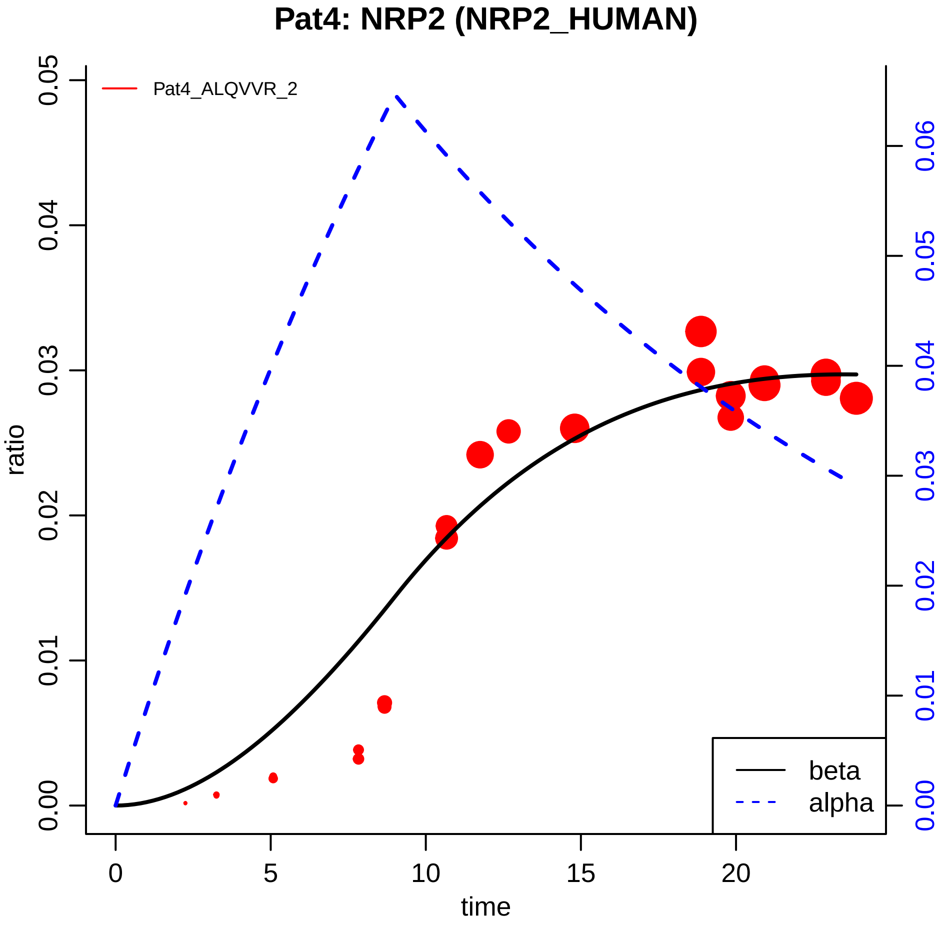

Pat4: NRP2 (NRP2_HUMAN)
Pat4_ALQVVR_2
ratio
0.00
0.01
0.02
0.03
0.04
0.05
0.00
0.01
0.02
0.03
0.04
0.05
0.06
time
0
5
10
15
20
beta
alpha

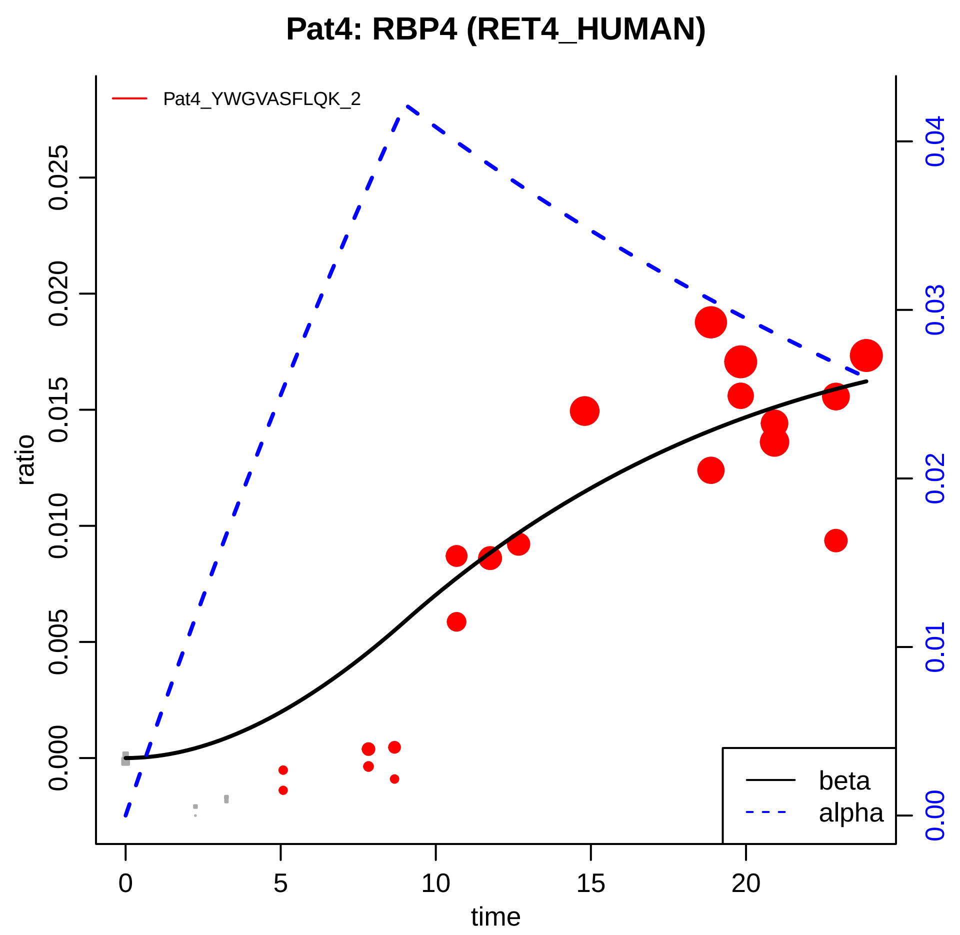

Pat4: RBP4 (RET4_HUMAN)
Pat4_YWGVASFLQK_2
ratio
0.000
0.005
0.010
0.015
0.020
0.025
0.00
0.01
0.02
0.03
0.04
0
5
10
15
20
time
beta
alpha

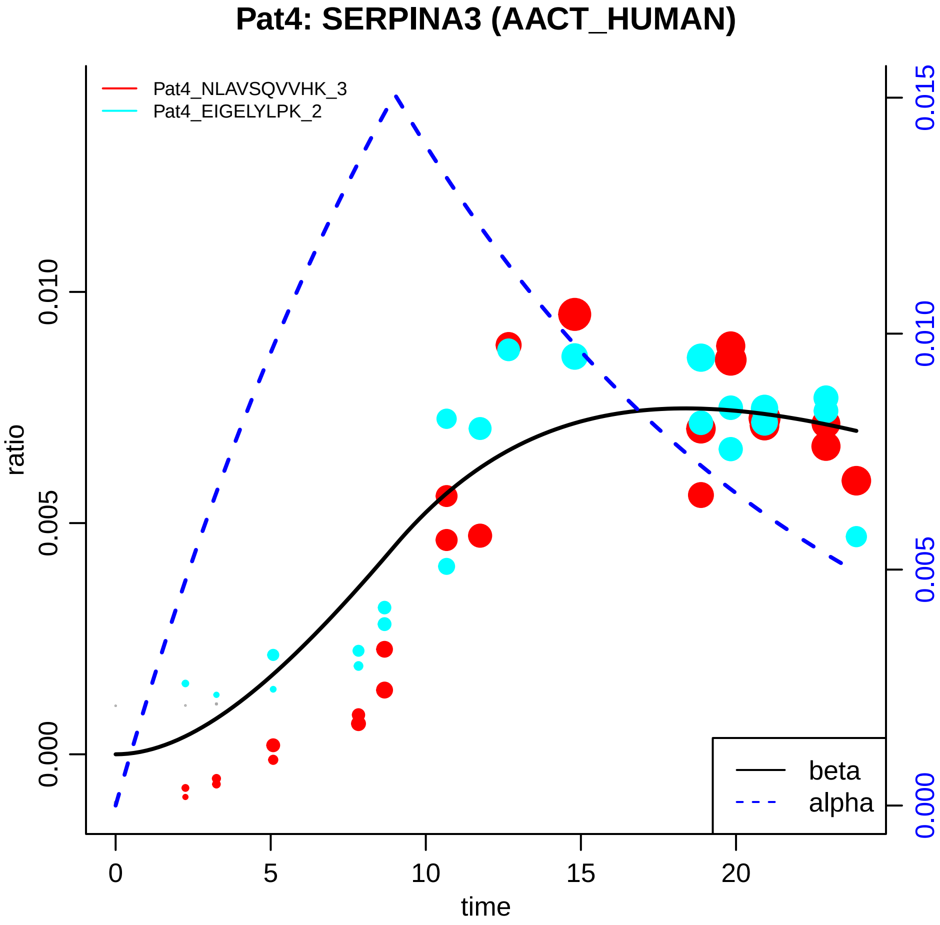

Pat4: SERPINA3 (AACT_HUMAN)
Pat4_NLAVSQVVHK_3
Pat4_EIGELYLPK_2
ratio
0.000
0.005
0.010
0.015
time
0
5
10
15
20
beta
alpha

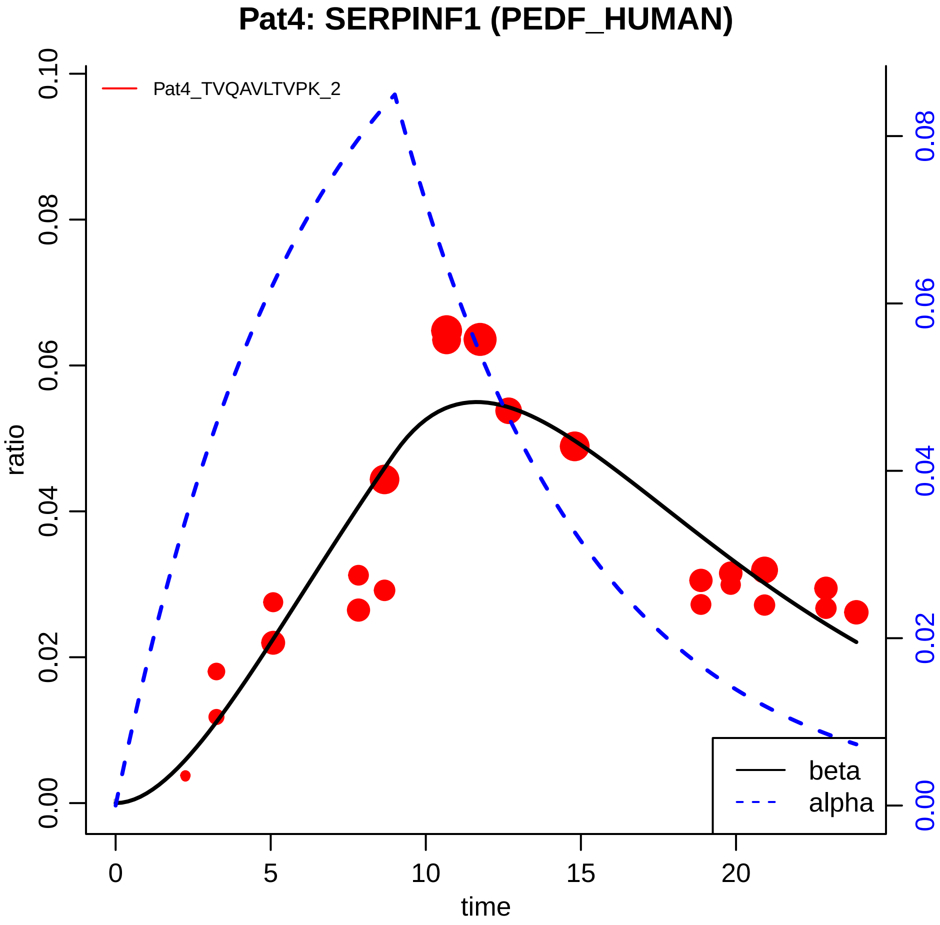
Pat4: SERPINF1 (PEDF_HUMAN)
Pat4_TVQAVLTVPK_2
ratio
time
0.00
0.02
0.04
0.06
0.08
0.10
0
5
10
15
20
beta
alpha

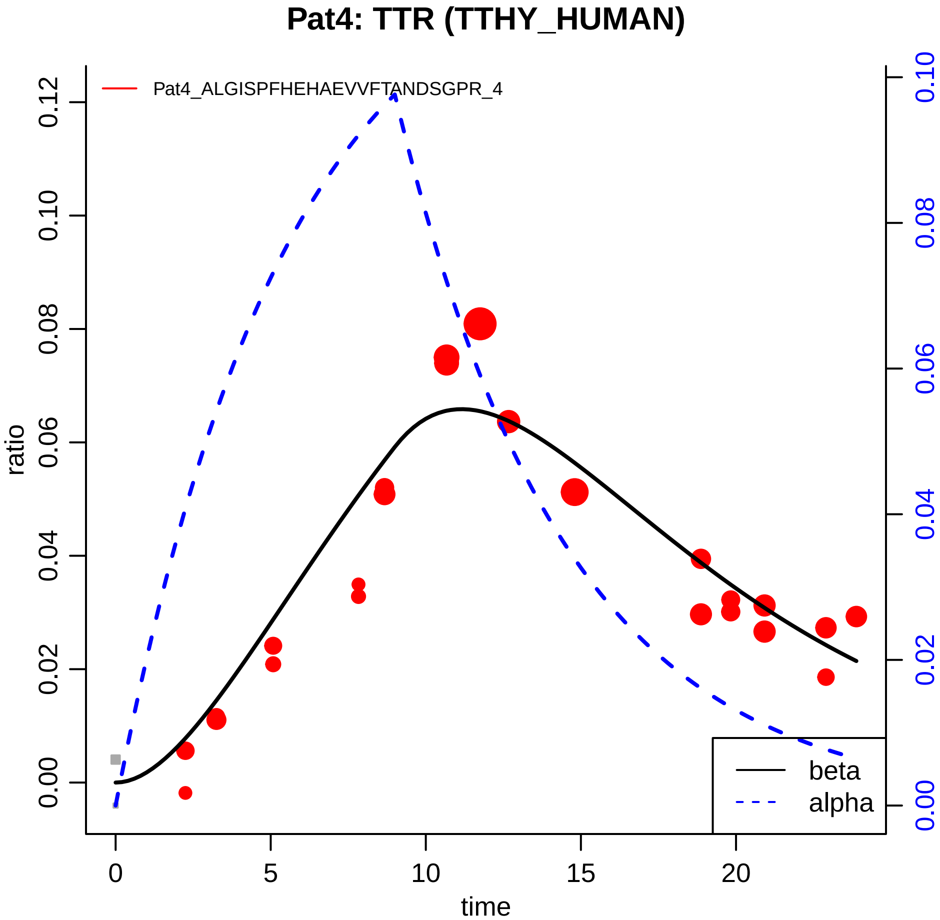

Pat4: TTR (TTHY_HUMAN)
Pat4_ALGISPFHEHAEVVFTANDSGPR_4
ratio
0.00
0.02
0.04
0.06
0.08
0.10
0.12
time
0
5
10
15
20
0.00
0.02
0.04
0.06
0.08
0.10
beta
alpha
