## Supplementary material for "*In Vivo* Large Scale Mapping Of Protein Turnover In The Human Cerebrospinal Fluid": original TQ protein models

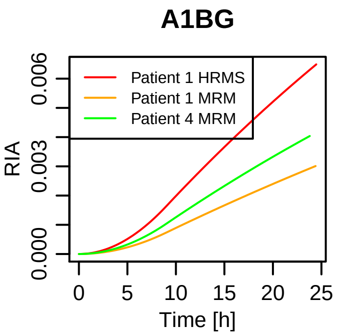

A1BG
Patient 1 HRMS
Patient 1 MRM
Patient 4 MRM
RIA
0.000
0.003
0.006
0
5
10
15
20
25
Time [h]

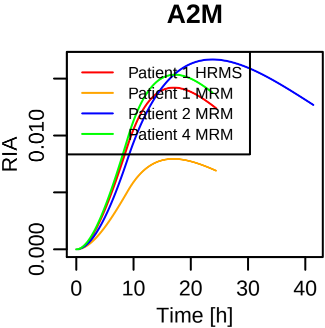

A2M
Patient 1 HRMS
Patient 1 MRM
Patient 2 MRM
Patient 4 MRM
RIA
0.000
0.010
0
10
20
30
40
Time [h]

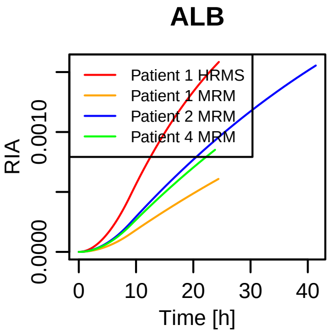
ALB
Patient 1 HRMS
Patient 1 MRM
Patient 2 MRM
Patient 4 MRM
RIA
0.0000
0.0010
Time [h]
0
10
20
30
40

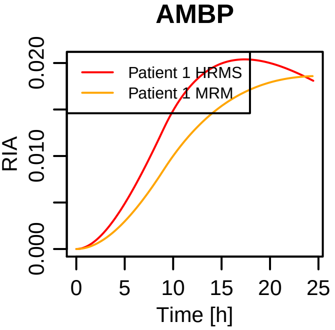

AMBP
Patient 1 HRMS
Patient 1 MRM
RIA
0.000
0.010
0.020
0
5
10
15
20
25
Time [h]

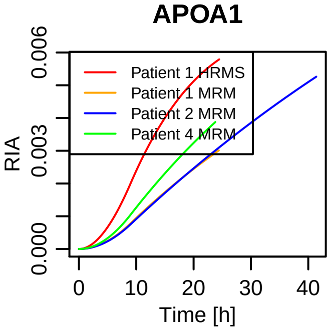
APOA1
Patient 1 HRMS
Patient 1 MRM
Patient 2 MRM
Patient 4 MRM
RIA
0.000
0.003
0.006
0
10
20
30
40
Time [h]

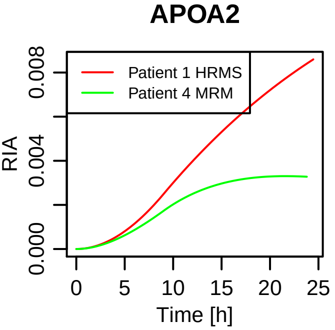

APOA2
Patient 1 HRMS
Patient 4 MRM
RIA
0.000
0.004
0.008
0
5
10
15
20
25
Time [h]

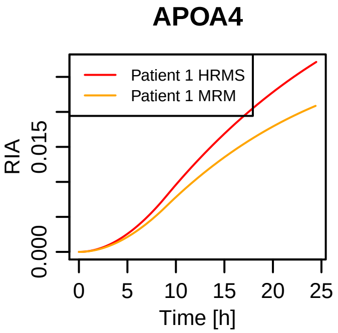

APOA4
Patient 1 HRMS
Patient 1 MRM
RIA
0.000
0.015
0
5
10
15
20
25
Time [h]

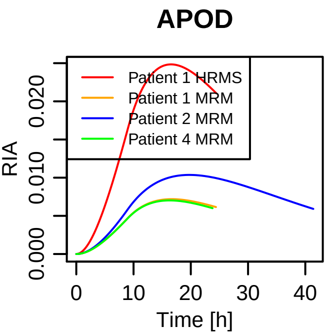

APOD
Patient 1 HRMS
Patient 1 MRM
Patient 2 MRM
Patient 4 MRM
RIA
0.000
0.010
0.020
0
10
20
30
40
Time [h]

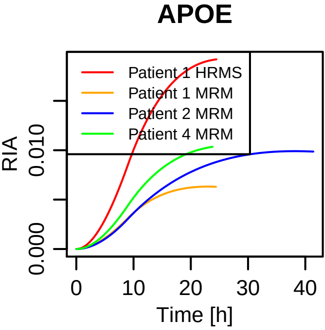

APOE
Patient 1 HRMS
Patient 1 MRM
Patient 2 MRM
Patient 4 MRM
RIA
0.000
0.010
0
10
20
30
40
Time [h]

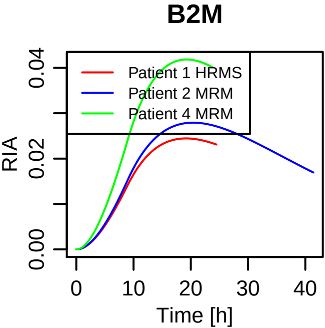
B2M
Patient 1 HRMS
Patient 2 MRM
Patient 4 MRM
RIA
0.00
0.02
0.04
0
10
20
30
40
Time [h]

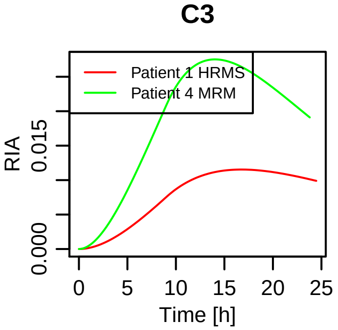

C3
Patient 1 HRMS
Patient 4 MRM
RIA
0.000
0.015
0
5
10
15
20
25
Time [h]

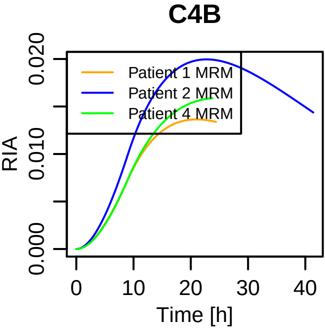

C4B
Patient 1 MRM
Patient 2 MRM
Patient 4 MRM
RIA
0.000
0.010
0.020
0
10
20
30
40
Time [h]

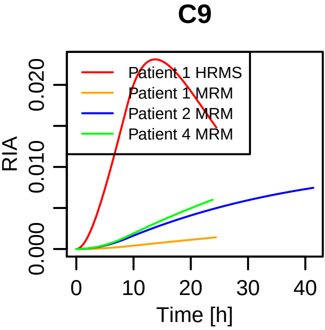

C9
Patient 1 HRMS
Patient 1 MRM
Patient 2 MRM
Patient 4 MRM
RIA
0.000
0.010
0.020
0
10
20
30
40
Time [h]

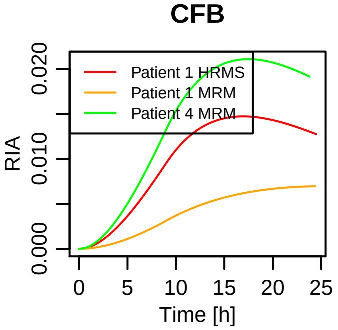
CFB
Patient 1 HRMS
Patient 1 MRM
Patient 4 MRM
RIA
0.000
0.010
0.020
0
5
10
15
20
25
Time [h]

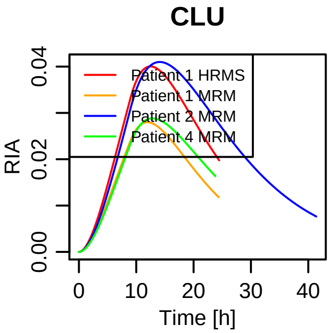
CLU
Patient 1 HRMS
Patient 1 MRM
Patient 2 MRM
Patient 4 MRM
RIA
0.00
0.02
0.04
0
10
20
30
40
Time [h]

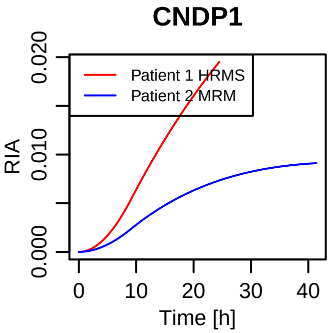
CNDP1
Patient 1 HRMS
Patient 2 MRM
RIA
0.000
0.010
0.020
0
10
20
30
40
Time [h]

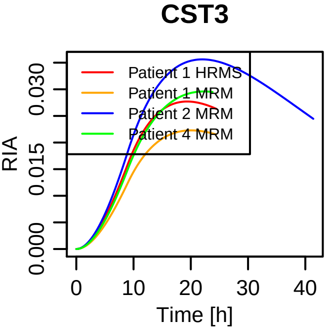

CST3
Patient 1 HRMS
Patient 1 MRM
Patient 2 MRM
Patient 4 MRM
RIA
0.000
0.015
0.030
0
10
20
30
40
Time [h]

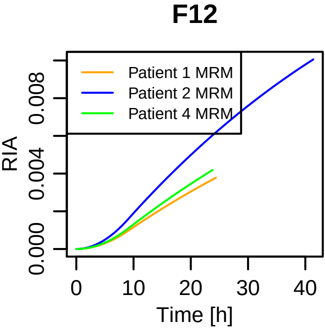

F12
Patient 1 MRM
Patient 2 MRM
Patient 4 MRM
RIA
0.000
0.004
0.008
0
10
20
30
40
Time [h]

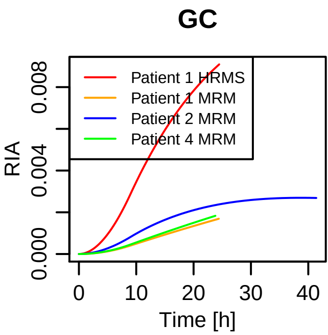

GC
Patient 1 HRMS
Patient 1 MRM
Patient 2 MRM
Patient 4 MRM
RIA
0.000
0.004
0.008
0
10
20
30
40
Time [h]

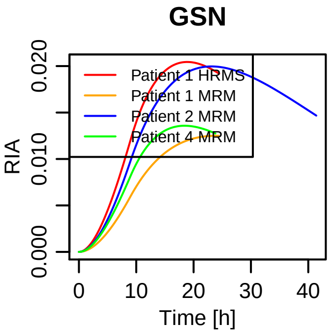
GSN
Patient 1 HRMS
Patient 1 MRM
Patient 2 MRM
Patient 4 MRM
RIA
0.000
0.010
0.020
0
10
20
30
40
Time [h]

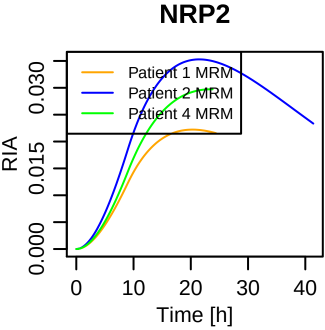

NRP2
Patient 1 MRM
Patient 2 MRM
Patient 4 MRM
RIA
0.000
0.015
0.030
0
10
20
30
40
Time [h]

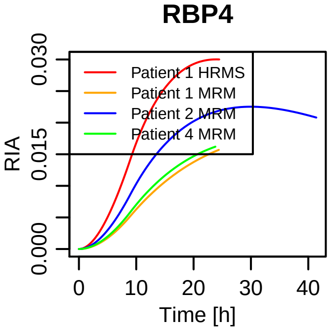
RBP4
Patient 1 HRMS
Patient 1 MRM
Patient 2 MRM
Patient 4 MRM
RIA
0.000
0.015
0.030
0
10
20
30
40
Time [h]

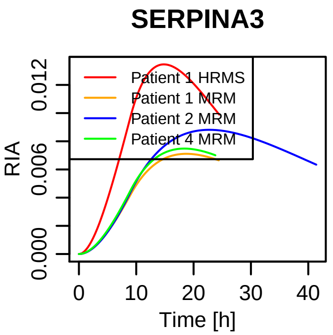

SERPINA3
Patient 1 HRMS
Patient 1 MRM
Patient 2 MRM
Patient 4 MRM
RIA
0.000
0.006
0.012
0
10
20
30
40
Time [h]

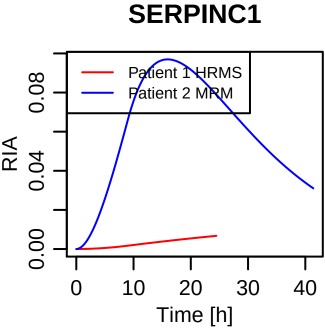

SERPINC1
Patient 1 HRMS
Patient 2 MRM
RIA
0.00
0.04
0.08
0
10
20
30
40
Time [h]

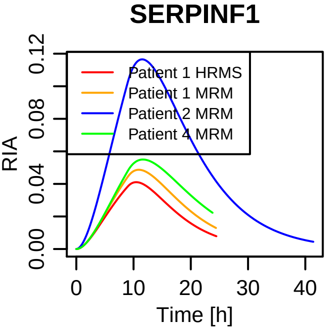
SERPINF1
Patient 1 HRMS
Patient 1 MRM
Patient 2 MRM
Patient 4 MRM
RIA
0.00
0.04
0.08
0.12
0
10
20
30
40
Time [h]

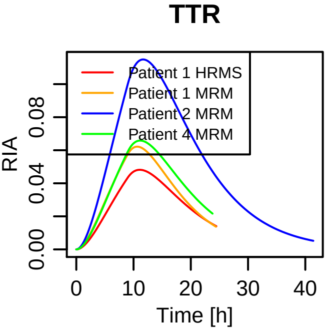
TTR
Patient 1 HRMS
Patient 1 MRM
Patient 2 MRM
Patient 4 MRM
RIA
0.00
0.04
0.08
0
10
20
30
40
Time [h]
