## Supplementary material for "*In Vivo* Large Scale Mapping Of Protein Turnover In The Human Cerebrospinal Fluid": aligned TQ protein models

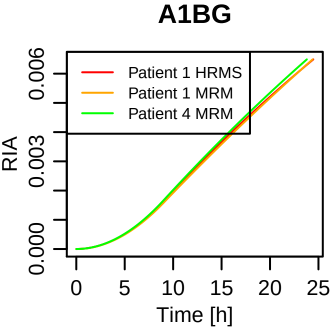

A1BG
Patient 1 HRMS
Patient 1 MRM
Patient 4 MRM
RIA
0.000
0.003
0.006
0
5
10
15
20
25
Time [h]

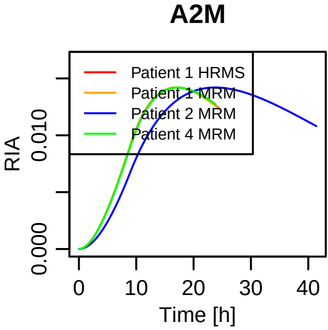

A2M
Patient 1 HRMS
Patient 1 MRM
Patient 2 MRM
Patient 4 MRM
RIA
0.000
0.010
0
10
20
30
40
Time [h]

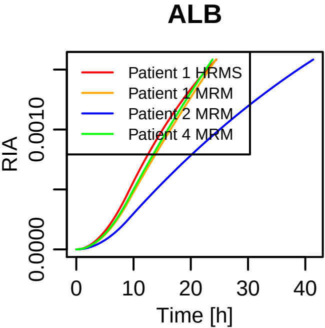

ALB
Patient 1 HRMS
Patient 1 MRM
Patient 2 MRM
Patient 4 MRM
RIA
0.0000
0.0010
0
10
20
30
40
Time [h]

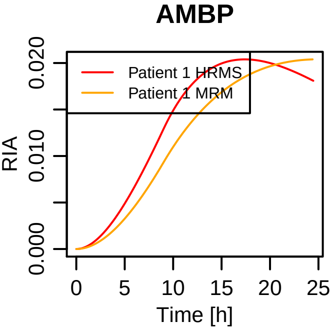
AMBP
Patient 1 HRMS
Patient 1 MRM
RIA
0.000
0.010
0.020
0
5
10
15
20
25
Time [h]

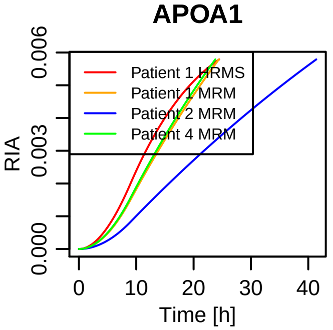
APOA1
RIA
0.000
0.003
0.006
Time [h]
0
10
20
30
40
Patient 1 HRMS
Patient 1 MRM
Patient 2 MRM
Patient 4 MRM

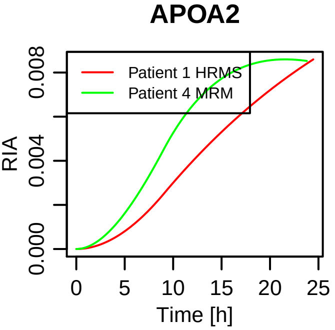

APOA2
Patient 1 HRMS
Patient 4 MRM
RIA
0.000
0.004
0.008
0
5
10
15
20
25
Time [h]

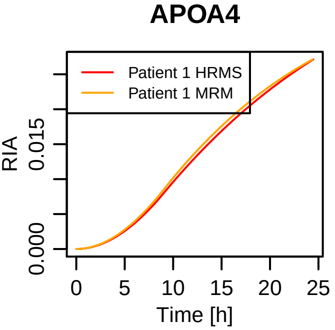

APOA4
Patient 1 HRMS
Patient 1 MRM
RIA
0.000
0.015
0
5
10
15
20
25
Time [h]

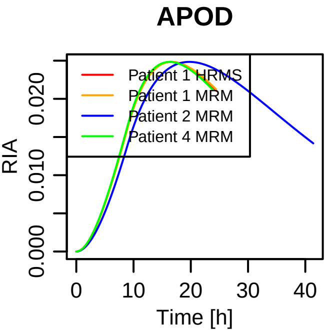

APOD
Patient 1 HRMS
Patient 1 MRM
Patient 2 MRM
Patient 4 MRM
RIA
0.000
0.010
0.020
Time [h]
0
10
20
30
40

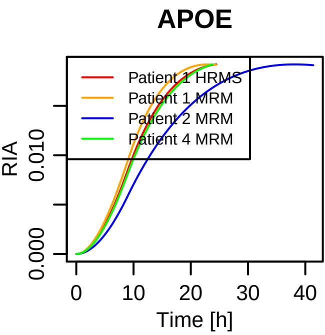
APOE
Patient 1 HRMS
Patient 1 MRM
Patient 2 MRM
Patient 4 MRM
RIA
0.000
0.010
0
10
20
30
40
Time [h]

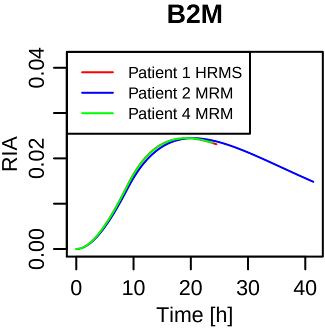

B2M
Patient 1 HRMS
Patient 2 MRM
Patient 4 MRM
RIA
0.00
0.02
0.04
0
10
20
30
40
Time [h]

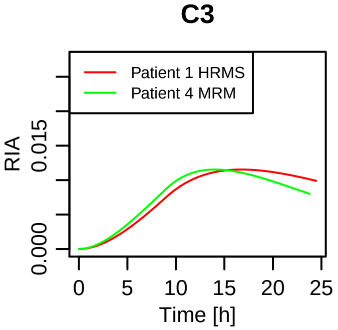
C3
Patient 1 HRMS
Patient 4 MRM
RIA
0.000
0.015
0
5
10
15
20
25
Time [h]

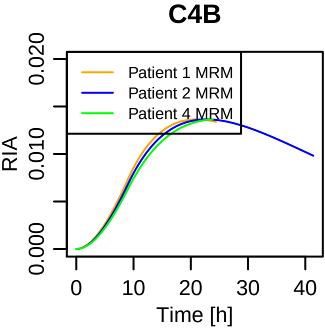
C4B
Patient 1 MRM
Patient 2 MRM
Patient 4 MRM
RIA
0.000
0.010
0.020
0
10
20
30
40
Time [h]

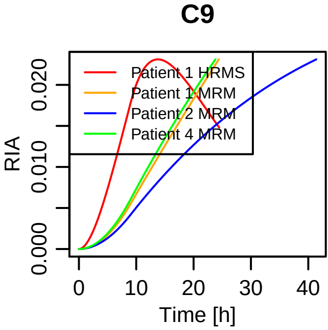

C9
RIA
0.000
0.010
0.020
Time [h]
0
10
20
30
40
Patient 1 HRMS
Patient 1 MRM
Patient 2 MRM
Patient 4 MRM

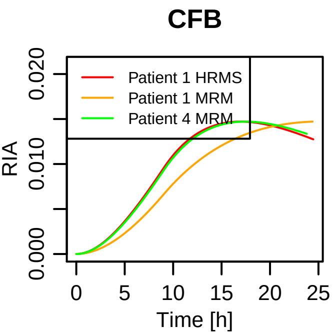
CFB
Patient 1 HRMS
Patient 1 MRM
Patient 4 MRM
RIA
0.000
0.010
0.020
0
5
10
15
20
25
Time [h]

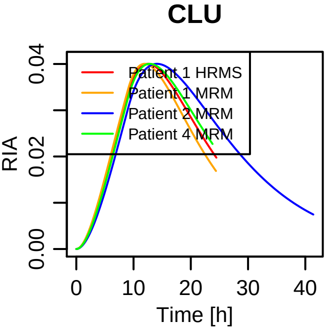
CLU
Patient 1 HRMS
Patient 1 MRM
Patient 2 MRM
Patient 4 MRM
RIA
0.00
0.02
0.04
0
10
20
30
40
Time [h]

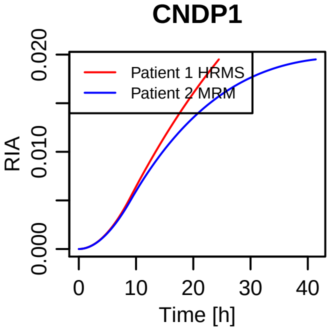
CNDP1
Patient 1 HRMS
Patient 2 MRM
RIA
0.000
0.010
0.020
0
10
20
30
40
Time [h]

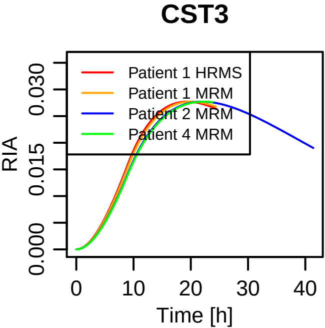
CST3
Patient 1 HRMS
Patient 1 MRM
Patient 2 MRM
Patient 4 MRM
RIA
0.000
0.015
0.030
0
10
20
30
40
Time [h]

# F12

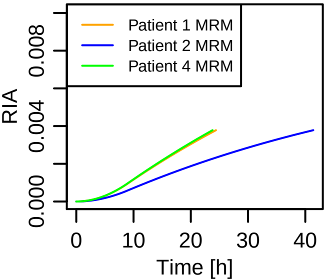
Patient 1 MRM
Patient 2 MRM
Patient 4 MRM
RIA
0.000
0.004
0.008
0
10
20
30
40
Time [h]

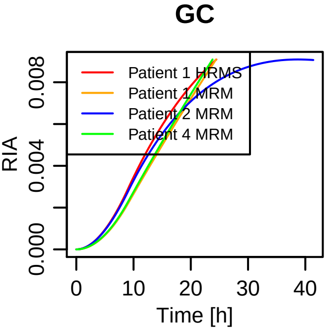

GC
Patient 1 HRMS
Patient 1 MRM
Patient 2 MRM
Patient 4 MRM
RIA
0.000
0.004
0.008
0
10
20
30
40
Time [h]

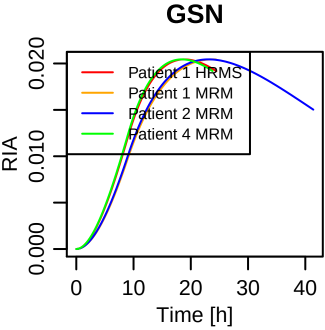
GSN
RIA
0.000
0.010
0.020
Time [h]
0
10
20
30
40
Patient 1 HRMS
Patient 1 MRM
Patient 2 MRM
Patient 4 MRM

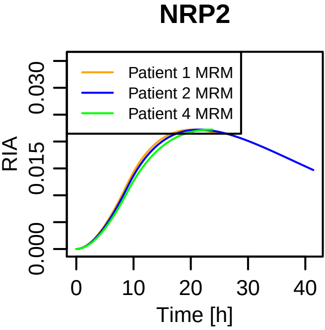

NRP2
Patient 1 MRM
Patient 2 MRM
Patient 4 MRM
RIA
0.000
0.015
0.030
0
10
20
30
40
Time [h]

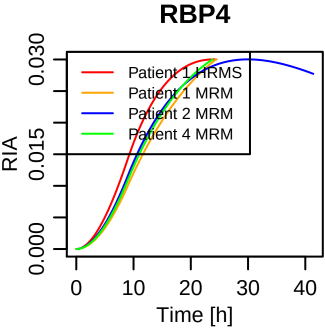

RBP4
Patient 1 HRMS
Patient 1 MRM
Patient 2 MRM
Patient 4 MRM
RIA
0.000
0.015
0.030
Time [h]
0
10
20
30
40

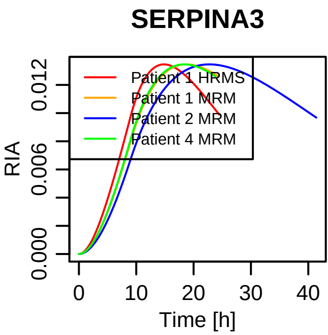

SERPINA3
Patient 1 HRMS
Patient 1 MRM
Patient 2 MRM
Patient 4 MRM
RIA
0.000
0.006
0.012
0
10
20
30
40
Time [h]

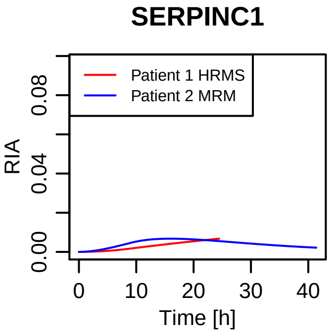

SERPINC1
Patient 1 HRMS
Patient 2 MRM
RIA
0.00
0.04
0.08
0
10
20
30
40
Time [h]

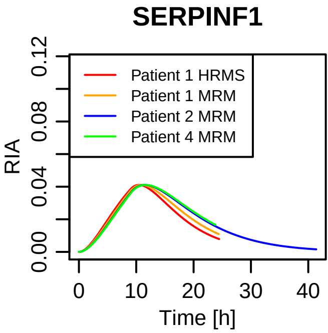
SERPINF1
Patient 1 HRMS
Patient 1 MRM
Patient 2 MRM
Patient 4 MRM
RIA
0.00
0.04
0.08
0.12
0
10
20
30
40
Time [h]

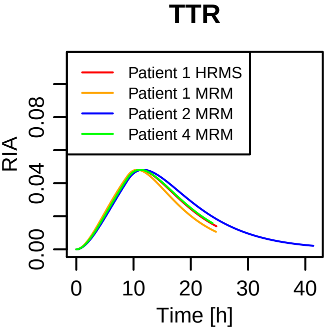

TTR
Patient 1 HRMS
Patient 1 MRM
Patient 2 MRM
Patient 4 MRM
RIA
0.00
0.04
0.08
0
10
20
30
40
Time [h]
